# The Establishment of Cell-Type Specific Gene Regulation in the Sea Urchin Embryo

**DOI:** 10.1101/2025.08.12.669999

**Authors:** Jonas Maurice Brandenburg, Alexandra Trinks, Dominika Voijtasova, Anna Alessandra Monaco, William Chang, Roberto Arsie, Marian Hu, Amro Hamdoun, Chloe Jenniches, Markus Landthaler, Nils Blüthgen, David Aaron Garfield

**Affiliations:** Institute of Biology, Humboldt Universität zu Berlin; Max Delbruck Center for Molecular Medicine, Berlin; Institute of Physiology, Christian-Albrechts Universität zu Kiel; Scrips Institution of Oceanography, University of California San Diego; Charité - Universitätsmedizin Berlin; Genentech, South San Francisco, USA

## Abstract

Cell-fate commitment in metazoan development relies on precise gene regulatory programs. This study presents a comprehensive single-cell atlas of gene expression (scRNA-seq), nascent transcription (scSLAM-seq), and chromatin accessibility (scATAC-seq) in the purple sea urchin, *Strongylocentrotus purpuratus*, from early cleavage to pluteus larva stages. Our findings reveal a dynamic regulatory landscape with extensive usage of distal and intronic regulatory elements, which often exhibit cell-type-specific motifs and accessibility profiles that closely track gene expression. We identify a major wave of zygotic genome activation (ZGA) at the 128-cell stage, coinciding with the loss of developmental plasticity, alongside evidence of restricted, lineage-specific gene activation preceding widespread ZGA. Motif analysis highlights distinct regulatory grammars for these early accessible regions. Regulatory element usage largely clusters by germ layer, indicating shared accessibility among related cell types. We delve into the regulatory intricacies of neurons and skeletogenic cells. Sea urchin neurodevelopment proceeds through three distinct lineages, utilizing transcription factors with conserved roles in mammalian neurogenesis. Surprisingly, skeletogenic cells show significant transcriptional and regulatory diversity across their subpopulations, and we identify novel genes associated with calcification. This research offers unprecedented insights into the dynamic regulatory genome of a non-chordate deuterostome, highlighting both conserved principles of gene regulation and unique features that underscore the sea urchin’s importance as a model for understanding developmental and evolutionary genomics in ecologically critical marine species.

## Introduction

Across metazoan development, cell-fate commitment is orchestrated by two inter-related events: Initial cellular asymmetries (e.g. sperm entry or protein localization) are transmitted into stable axis formation in the embryo and maternally deposited developmental information (in the form of proteins and mRNA) are translated to the zygotically encoded gene expression programs that underlie cell-fate specification and cellular differentiation (Schulz & Harrison, 2019; Vastenhouw *et al*, 2019). But while these events are nearly universal in metazoan development, the specifics vary across taxa (Schulz & Harrison, 2019; Vastenhouw *et al*, 2019; Goldstein & Freeman, 1997; Beddington & Robertson, 1999) with little known about how these first steps in development are encoded in the regulatory genome outside of a few model species.

In this study, we make use of a combination of single-cell RNA-sequencing (scRNA-seq), metabolic labeling (scSLAM-seq), and single-cell ATAC-sequencing (scATAC-seq) to examine changes in the regulatory genome of the purple sea urchin *Strongylocentrotus purpuratus* from the earliest cleavage stages through cell fate specification and the development of the early pluteus larvae. *S. purpuratus* has been a model in developmental biology for more than a century, resulting in a detailed understanding of cell fates and lineage relationships among all early blastomeres from the 8-cell embryo onwards (McClay, 2011) including a map of the “logic” of many of the TF interactions necessary for early cell fate specification (Ransick & Davidson, 2006; Yuh *et al*, 1996; Barsi & Davidson, 2016; Yuh *et al*, 2004).

This map is robust under normal developmental conditions, but can be highly plastic in the face of perturbations, including the removal of entire cell lineages (Hörstadius, 1936). Elements of this plasticity remain at later stages of development, for example the replacement of the skeletogeneic mesoderm (and, likely, embryonic PGCs) from specified, non-skeletogenic mesoderm cells following the removal of the micromere/PMC lineage (Ettensohn *et al*, 2007; McClay & Logan, 1996; Yajima & Wessel, 2011). How this plasticity is regulated, and why it is restricted at later stages of development, is unknown.

Many of the molecular players driving early cell-fate specification in the sea urchin are known, for example, the specification of the animal-vegetal axis via the asymmetric distribution of maternal factors and the nuclear localization of β-catenin (Weitzel *et al*, 2004; Lyons *et al*, 2014), and for some genes, the promoter-proximal regulatory elements at which transcription factors bind in early development have been described in extensive detail (Garfield *et al*, 2012). The extent to which these examples exemplify generalities in CRE architecture for this species, however, is unclear, as is the role for for non-promoter proximal regulatory elements, which constitute an extensive component of the regulatory genome in most metazoan species (Cornejo-Páramo *et al*, 2022; Reddington *et al*, 2020; Oudelaar & Higgs, 2021; Noonan & McCallion, 2010).

In this study, we expand our understanding of gene regulation in the sea urchin through the construction of a detailed, cell-type-specific atlas of gene expression and genome-wide regulatory element usage spanning from early cleavage, through the activation of the zygotic genome, the loss of trans-germ-layer plasticity, through the terminal differentiation of diverse cell types in early pluteus larvae. We identify a major wave of zygotic genome activation (ZGA) between the 60- and 128-cell stage, with evidence of restricted activation, pre-ZGA, for a core set of lineage-specific TFs. We further identify a diverse array of cell-type-specific regulatory elements with clear enrichment of cell-type-specific regulatory motifs, indicating a conserved regulatory grammar with strong similarities between sea urchins and mammalian species. Many of these elements are intronic or intergenic, highlighting an essential role for regulatory interactions outside of the proximal promoter region in the sea urchin. Finally, we focus on two key differentiated cell types – neurons and skeletogenic cells, uncovering the regulatory events underlying subpopulations of these diverse cell lineages.

## Material & Methods

### Sea urchin husbandry and handling

Sea urchins were obtained from Monterey Abalone Company (Monterey, California), housed in 12°C artificial sea water, and fed *ad libitum* with kelp. Gametes were obtained by intracoelomic injection of 0.5ml KCl (0.55M). Eggs were collected for 15 minutes and washed in filtered artificial sea water (FASW) [437mM NaCl, 9mM KCl, 22.9mM MgCl_2_, 25.5mM MgSO_4_, 9.3mM CaCl_2_, 2.1mM NaHCO_3_; pH adjusted to 8.2] twice before use. For each female, a sample of 100 eggs was inspected for signs of fertilization and egg quality under a microscope. Fertilization occurred in vitro by adding sperm to eggs (final dilution of sperm 1:20 000). Eggs were fertilized within 4h of gamete collection. All gametes were kept on ice until fertilization and only cultures with fertilization rates >95% were used for this study. Newly fertilized cultures were washed twice with FASW to remove excess sperm and divided in culture dishes in monolayer within 1h of fertilization. Final cultures were covered with aluminum foil to prevent evaporation and incubated at 15°C for the indicated period.

### Cell dissociation for single-cell RNA sequencing

For scRNA-seq experiments, cells were dissociated using previously described methods (McClay, 2004). Briefly, staged embryos were pelleted at 150g at 4°C for 5 minutes, washed with calcium-free artificial sea water (CFSW) [454mM NaCl, 9mM KCl, 48mM MgSO_4_, 6mM NaHCO_3_; pH adjusted to 8.2], and incubated in hyaline extraction medium (HEM) supplemented with BSA [NaCl 300mM, KCl 10mM, MgSO_4_ 10mM, Glycine 300mM, Tris 10mM, EGTA 2mM; pH adjusted to 8.2; supplemented with BSA 2mg/mL] on ice for 5 minutes. Embryos were dissociated by gentle trituration with a 25ml serological pipet until reaching >95% single cells. Dissociated cells were then washed twice in CFSW and remaining clumps of cells were filtered out using a 40μm cell strainer. Cell viability was assessed using propidium iodine (Sigma Aldrich) staining. Only samples with > 90% viability were processed further. Cells were then diluted to 5x10^6^ cells/ml in CFSW and fixed by adding ice-cold MeOH (to 90%) dropwise while gently vortexing and stored at -20°C until processing. To rehydrate fixed cells prior to library preparation, cells were spun down (1000g, 5min at 4°C) and hydrated by two washes in CFSW with 0.1% BSA and 1x SUPERaseIn RNase inhibitor (both Fisher scientific).

Rehydrated cells were counted using a hematocytometer and diluted in Hanks balances salt solution (HBSS) with 0.04% BSA, followed by library preparation according to manufacturer’s instructions (10x Genomics). Quality of final libraries was assessed using a Tape Station D1000 kit (Agilent) and quantified on a Qubit 3 fluorometer (Fischer scientific). Libraries were sequenced with 2x150 cycles on a NovaSeq 6000 (scSLAM-seq of Hpf3 to Hpf15) or a NextSeq system (R1:28, R2:91, i7:8 cycles; scSlam-seq of Hpf24 and scRNA-seq of Hpf20 to Hpf72; both Illiumina) to 45 000 reads per cell, resulting in a median of 2 889 unique molecular identifiers (UMIs) and 1,301 unique genes detected per cell. Cells from multiple donors and time points were pooled in three batches, with the resulting reads later separated based on single nucleotide polymorphisms (SNPs) present (described below, Table S1).

### scRNA-seq read processing

Raw data was converted to fastq format using cellranger mkfastq (10x Genomics, version 6.0.2). Cutadapt v1.18 (Martin, 2011) was used to trim TSO and polyA sequences and the length of read 2 was limited to 91bp. Fastq files were then converted to bam format and counted using cellranger count. High quality cells were determined based on percentage of mitochondrial UMI (< 30%), number of UMIs (≥ 1,000 for post-cleavage stages, and ≥ 3 000 for cleavage stages and below 50,000 to exclude doublets) and number of genes detected per cell (≥ 500 and < 10,000). Donors within each pooled run were deconvoluted using cellSNP-lite and Vireo v0.5.7 with genotype information (Huang & Huang, 2021; Huang *et al*, 2019a). Given the use of eggs from the same mothers at multiple time points (with different fathers) in our early scRNA-seq dataset, Vireo was only able to assign cells to the probable mother. In cases of ambiguity, cells were assigned to the corresponding timepoint (and thus father of origin) using enrichments of stage specific gene-scores based on our bulk-RNA-seq time series.

These thresholds resulted in a total of 20,928 retained cells, originating from 20 donors and 12 developmental stages across three libraries (Table S1). Each developmental stage was clustered independently following the recommended Seurat v4 workflow (Hao *et al*, 2021). Briefly, the 2000 most variable genes were log-normalized and scaled, while regressing out the percentage of mitochondrial reads. Principal component analysis (PCA) was performed on the scaled data and the top 30 principal components were used for Uniform Manifold Approximation and Projection (UMAP). Clustering of cells was archived by building a kNN-graph based on the top 30 principal components of the data and applying the Louvain algorithm with a resolution of 0.8. If multiple donors were present at a developmental stage, donors were integrated using the CCA-based integration workflow of Seurat (Stuart *et al*, 2019). The resulting Louvain clustering was used to annotate cell types using known marker genes (Table S2). For refined cell type annotation, cells for each developmental stage were sub-clustered at the germ layer level (based on 15 principal components, resolution of 0.8 to 2, depending on the complexity of the cell type). As skeletogenic and neuronal cells already constitute highly distinct cell states, these cell types were subclustered without other cell types of their germ layer. Doublets were removed from the datasets using DoubletFinder v2.0.3 (McGinnis *et al*, 2019) with default settings, followed by the removal of sub-clusters of cells expressing incompatible, germ-layer specific marker genes. Finally, the combined projections across developmental time points were created by merging annotated and filtered datasets from all developmental stages and performing PCA and UMAP using the intersection of highly variable genes assessed between individual experiments. This, in combination with regressing out the fraction of mitochondrial RNA (percent.mt) reduced batch-specific effects. As above, the top 30 principal components were then used for UMAP projections. Lineage and cell-type-specific differentially expressed genes (DEG) were identified using a Wilcoxon Rank Sum test (min.pct = 0.25, logFC threshold = 0.25; adjusted p-value < 0.1; Table S3. To infer neurodevelopmental trajectories in our dataset, we used a diffusion map approach (Haghverdi *et al*, 2015), ordering neuronal cells on a diffusion gradient based on variable features, and further validating the trajectories, using RNA velocities, as described (Bergen *et al*, 2020).

For the analysis of skeletogenic gene regulation, skeletogenic cells were subclustered separately and skeletogenic cell-specific transcription factors (TFs) and effector genes were identified based on differentially expression within skeletogenic cells vs. all other cell types. Skeletogenic cell-specific genes were then assigned to subpopulation specific expression patterns by comparing differentially expression within subsets of skeletogenic cells and vs. all other cells. A calcification module score was then calculated based on the expression of a set of genes known to be involved in calcification (Otop2l, Slc4a10, Cara7, sAC (Chang *et al*, 2021; Hu *et al*, 2020)). Genes with expression significantly correlated with the calcification module score (Pearson correlation coefficient, FDR < 0.1) were identified as probable calcification genes.

### Validation by qPCR

To validate candidate calcification genes, recalcification assay and qPCR were performed. RNA samples from control and recalcifying larvae were extracted by using the Direct-zol RNA Miniprep Kits (Zymo Research) followed by complementary DNA (cDNA) synthesis via SuperScript IV Reverse Transcriptase (Invitrogen). Gene expression analysis by qPCR was performed on a ABI 7500 Real-Time PCR system (Fisher Scientific) and gene expression levels were normalized to the reference gene Ef1a (LOC548620) that was demonstrated to be stable during larval development and recalcification conditions (Hu *et al*, 2020). All amplification primers used for qPCR analysis are listed in Table S4.

### Bulk RNA sequencing and genotyping

For bulk RNA analysis, approximatively 1000 staged embryos were pelleted in a minifuge, resuspended in RLT-buffer (Qiagen RNeasy kit) with β-Mercaptoethanol and dissolved by vortexing. Dissolved samples were flash-frozen in liquid nitrogen and stored at -80°C. The procedure from sampling to freezing took less than 2 minutes. Total RNA was extracted from these samples using the RNeasy mini Kit (Qiagen) according to manufacturer’s instructions.

Two library preparation methods were used for bulk-RNA sequencing in this study. Matching bulk RNA libraries for scHpf24_rna samples (used for genotyping; i.e. Hpf24_1 and Hpf24_2, Hpf10_1 and Hpf8_1) and the bulk-RNA seq time course were prepared using the TruSeq RNA Library Prep Kit v2 (Illumina) according to manufacturer’s instructions. For the preparation of bulk RNA libraries for genotyping all other scRNA-seq samples, we made use of a 3’-tag sequencing approach (BRB-seq) that more directly complemented the 3’-biased libraries resulting from scRNA-seq (Alpern *et al*, 2019).

After sequencing on a HiSeq 4000 machine (Illumina) with 2x150bp reads, data from TruSeq V2 generated libraries were converted to fastq files and demultiplexed based on sample specific i5 barcodes using bcl2fastq v2.20 (Illumina). Adapter sequences were trimmed from the fastq file using skewer v0.2.2 [adaptor_x: NNNNNNNAGATCGGAAGAGCGGTTCAGCAGGAATGCCGAG, adaptor_y=NNNNNNNAGATCGGAAGAGCGTCGTGTAGGGAAAGAGTGT, paired end mode (Jiang *et al*, 2014)], and trimmed reads were aligned to Spur_5.0 genome using STAR v2.6.1 (Dobin *et al*, 2013). Uniquely mapping, high quality reads were filtered using samtools v1.9 [settings -F 2308 -q 20 (Li *et al*, 2009)]. For the bulk-RNA-seq time-course dataset, differential expression analysis was performed using the DEseq2 framework (Love *et al*, 2014). Differential expression analysis was performed relative to Hpf3 (4-cell stage), as delayed polyadenylation ceases at this time point and all genes upregulated after this stage are likely zygotic transcripts (Slater *et al*, 1972).

Due to the inherently different library structure created by BRB-seq, demultiplexing and adapter trimming was performed using the tool suite designed for BRB-seq (Alpern *et al*, 2019). Following conversion of sequencing data to fastq format using bcl2fastq v2.20 (Illumina), adapters and poly-A stretches were trimmed from read1, and reads were aligned to Spur_5.0 genome using STAR v2.6.1 (Dobin *et al*, 2013). The resulting bam file was used for calling of SNPs using the CellSNP-lite software v1.2.2 (Huang & Huang, 2021) and only SNPs also called based on scRNA-seq samples (minor allele fraction (MAF) > 0.1, Depth > 100) were retained. The resulting vcf file was used for assigning genotypes using Vireo (see above).

### Single-cell SLAM-sequencing with metabolic labeling (scSLAM-seq)

ScSLAM-seq was performed following a protocol previously published (Uhlitz *et al*, 2021). Briefly, unfertilized sea urchin eggs were incubated with 250μM 4sU in FASW for 30 minutes prior to fertilization. For best fertilization success, 4sU was washed out shortly before fertilization. Following fertilization and washing of cultures, 4sU was again added to a final concentration of 250μM in FASW (30 minutes after fertilization) and embryos were incubated at 15°C for indicated times. No effect of 4sU on developmental progression and growth was noted at the doses used and incorporation efficiency appeared to be similar between cell types, with only slightly lower 4sU incorporation rated in mesodermal cell types and neurons at Hpf24. Cells were dissociated as described for scRNA-seq and fixed for 30 minutes at -20°C. Single cells were next incubated with 10mM iodoacetamide overnight in 80% MeOH in CFSW at room temperature with gentle rotation. Cells were rehydrated in presence of 100mM DTT to quench the acetylation. Reads from scSLAM-seq was initially processed using cell-ranger (version 6.0.2, 10x Genomics), as described above. Labeled and unlabeled reads were counted using a custom pipeline, as previously described (Uhlitz *et al*, 2021). To deal with the high number of polymorphisms in the sea urchin, sites with >10% T to C transitions relative to the reference genome were excluded. UMIs containing more than one T to C transition were counted as “new” mRNA (i.e. zygotic), while UMIs without T to C transition were assigned as “old” (i.e. maternal). Overall, we obtained an incorporation efficiency of 3.2% across all potential incorporation sites.

### Nuclei isolation and single-cell ATAC-sequencing

Nuclei were isolated from embryos using the BITS-nuclei extraction protocol (Bonn *et al*, 2012) with minor modification for sea urchins: Sea urchin embryos were pelleted by centrifugation for 5 minutes at 150g and 4°C. Next, FASW was removed, and embryos were resuspended in HB buffer [Tris pH 7.4 15mM, Sucrose 340mM, NaCl 15mM, KCl 60mM, EDTA 0.2mM, EGTA 0.2mM, filtered and stored at 4°C]. Embryos were dissociated using a dounce homogenizer (Wheaton Scientific, 15mL, 15 loose, 5 tight pestle strokes) and passed through double-layered Miracloth. Nuclei were washed once in HB buffer and resuspended in PBT [PBS with 0.1% Triton X-100] resulting in a single-nuclei preparation. To remove any remaining clumps, nuclei were passed through a 20μm nitex mesh. Nuclei were stained with propidium iodide and counted on a Neubauer chamber. Isolated nuclei of early development samples (i.e. Hpf3 to Hpf15) and one Hpf72 sample (used to compare quality of NFB-frozen vs. fresh nuclei) were diluted in 90% Nuclear Freezing Buffer [NFB; 50 mM Tris at pH 8.0, 25% glycerol, 5 mM Mg(OAc)2, 0.1 mM EDTA, 5 mM DTT, 1X protease inhibitor cocktail (Roche), 1:2500 SUPERaseIn (Fischer scientific)] (Cusanovich *et al*, 2018) and flash frozen in liquid nitrogen. Prior to use, nuclei were thawed in a dry bath at 37°C, filtered again and diluted to appropriate concentration in Diluted Nuclei Buffer (10x Genomics). Library preparation for scATAC-seq was performed according to manufacturer’s instructions (10x Genomics).

Sequencing was carried out on a NovaSeq 6000 (scATAC Hpf3 to Hpf15 and Hpf20 to Hpf72 samples) or HiSeq 4000 system (scATAC_Hpf24 sample) in 2x50bp mode, yielding on average ∼125 000 reads per nuclei. Sequencing data was processed using cellranger-atac (version 1.2, 10x Genomics), and nuclei were filtered based on total reads (>1 000), percent reads in peaks (> 10%), nucleosomal score (<4) and TSS Enrichment (>2), with the lower than typical percent reads in peaks being selected to keep nuclei from pre-ZGA samples. This resulted in a total of 20,928 nuclei, originating from 17 donors and 12 developmental stages (Table S5). Donor deconvolution was performed as indicated for scRNA-seq. Each cross, and stage was clustered independently as suggested in the Signac v4 workflow (Stuart *et al*, 2021). Briefly, counts were normalized, and linear dimensional reduction was performed using latent semantic indexing (LSI). UMAP and Leiden clustering was carried out as described for scRNA-seq data while excluding the first dimension (which is correlated with sequencing depths) and doublets between cells of the same cross were removed using ArchR doublet scoring (Granja *et al*, 2021). Cell types were assigned through label transfer from scRNA-seq dataset, as described (Stuart *et al*, 2021). In some cases, it was necessary to merge cluster annotations (e.g. multiple neuronal subtypes into “neurons”), resulting in fewer annotated states in our scATAC-seq dataset. Given the low overlap between the transcriptome and accessible genome in the transcriptionally quiescent primordial germ cells (PGCs) of the blastula (Oulhen & Wessel, 2017) the cluster of PGCs was annotated manually.

Combined clustering of scATAC-seq data from all stages was performed using Harmony (Korsunsky *et al*, 2019), based on 20000 most accessible peaks, followed by UMAP projection based on 30 dimensions of the harmony-corrected LSI. Differential peak accessibility was assessed on the merged dataset for lineage specific peaks and on developmental stage clustering for cell-type-specific peaks (Table S6, S7), using logistic regression and the total number of fragments as a latent variable and min.pct of 0.05, as suggested in the Signac framework (Stuart *et al*, 2021).

Regulatory state clustering was performed using scregseg v0.1.1 (McGarvey *et al*, 2022). Cell-type and stage specific counts were merged per peak for all cell types with more than 10 cells, and the resulting aggregated count matrix was used as input for the scregseg segmentation step. Scregseg classifies the genome into regulatory states based on cross-cell-type accessibility profiles, using a HMM-base segmentation model. To reduce batch-effects, only the 20,000 most accessible peaks per experiment were used for this analysis and batch-specific regulatory states were removed. Using these cut-offs, the final round of segmentation was performed on 23,421 peaks.

### Bulk ATAC sequencing

Nuclei were isolated as described for scATAC-seq. Following nuclear preparation, nuclei were flash frozen in 90% NFB and stored in 20μl aliquots (100,000 nuclei each) at -80°C. For bulk ATAC-seq library preparation, nuclei were thawed at 37°C, and ATAC enzyme mix [3μl in-House Tn5 (MDC Protein Production & Characterization Platform), 1x Taps-DMF,16.5μl PBS, 0.5μl Digitonin 1%, 0.5μl Tween 20 10%] (Picelli *et al*, 2014; Corces *et al*, 2017; McGarvey *et al*, 2022) was added directly onto the nuclei. Reaction mixes were mixed by pipetting, incubated for 30 minutes at 37°C with gentle rotation and cleaned up using a MinElute PCR purification kit (Qiagen). Following PCR amplification, libraries were size selected (range 200 -1 000bp) using AmPURE XP beads (Beckman Coulter) and libraries were checked for characteristic nucleosomal banding patterns using a TapeStation D1000 kit (Agilent).

Final libraries were sequenced on a HiSeq 4000 machine (Illumina) in 2x150bp mode, and reads aligned to Spur_5.0 genome using BWA MEM [version=0.7.17(Li, 2013)]. Reads were filtered for unique mapping and duplicates. Peaks were called on the filtered bam file using MACS2 v2.1.1 (Zhang *et al*, 2008) and a reference peak set was created for each developmental time point (i.e. only peaks overlapping between replicates were retained; Table S8). Peaks were annotated using ChIPSeeker v1.36 (Yu *et al*, 2015) using a promoter range of 1kb upstream of the transcription start site (TSS). Throughout this manuscript, the following peak type definitions are used: Promoter (1kb upstream of, and including the TSS), 5’UTR, 3’UTR, intron and exon refer to positions on the appropriate gene feature, downstream (within 5kb of the end of a gene) and distal intergenic (being more than 5kb downstream, or >1kb upstream of a gene). Elements lying outside of these ranges were not assigned to a gene.

To generate references for genotyping scATAC-seq samples, filtered bulkATAC bam files were subject to SNP calling using cellSNP-lite and restricted to SNPs also called based on scATAC-seq data (MAF > 0.1, Depth > 300). The resulting vcf file was used for assigning genotypes in Vireo v0.5.7 (Huang *et al*, 2019b).

### Ancestry voting and candidate transcription factor identification

Ancestry voting was performed as described (Qiu *et al*, 2022). Briefly, cells from each developmental time point were integrated with cells from the corresponding previous time point. Dimensionality reduction was performed using principal component analysis (PCA) and uniform manifold approximation and projection (UMAP), and Euclidean distances between individual cells from both time points (i.e. ancestor and daughter time point) were calculated. For each cell of the daughter time point, the 5 closest neighbors from the ancestor time point were identified, and the fraction of the cell types of predicted ancestry states constitute the edge weight. This step was repeated 500 times with 80% subsampling for bootstrapping. Edges were defined as the median proportions of these weights. Edge weights >0.3 were retained for constructing the cell lineage tree.

Candidate lineage TFs were identified as differentially expressed TFs (Wilcoxon Rank Sum test, min.pct=0.25, logFC threshold=0.25; adjusted p-value < 0.1) between daughter cell types that also showed significantly increased expression compared to the “pseudo-ancestor” state, following previous reports (Qiu *et al*, 2022). Only ancestor-daughter pairs corresponding to known biological cell fate commitments were retained for this analysis.

### Temporal analysis of gene expression and regulatory element opening

To compare mRNA expression and regulatory peak opening, cells profiled by scATAC-seq and scRNA-seq were ordered based on true developmental time, and average gene expression (based on unspliced reads, to capture current transcription only, given strong maternal contributions at early time points) and regulatory element accessibility were calculated for each cell type. The time point where accessibility or expression reached >50% of the gene specific maximum was retained as the time point of onset of gene expression or opening of regulatory element, respectively.

### Motif enrichments

Enriched motifs were identified using the Homer tool suite (Heinz *et al*, 2010). For cell-type-specific motifs, cell-type-specific differentially accessible elements were used as foreground and differentially accessible peaks found in other cell types but not in the target cell type at the respective time of development as background. To identify motifs associated with the activation of the zygotic genome, newly accessible peaks per developmental interval (i.e. before 16-cell, 16- to 32-cell stage (pre-ZGA), 60- to 256 cell stage (ZGA) peaks were compared to all accessible peaks at prior time points as background. Combining developmental stages was necessary to obtain more robust peak sets. In the case of the pre-16-cell peak set, peaks called on tagmented genomic DNA of the same genotypes were used as background. Enriched motifs were compared to Homer database(Heinz *et al*, 2010), and similarities >0.8 with vertebrate or fly TFBS were considered meaningful.

### Categorical enrichment

Categorical enrichments for differentially expressed genes and differentially accessible peaks were carried out via Fisher’s exact tests implemented in suites of custom Ruby and Python scripts. In these tests, peaks or genes showing differential accessibility or differential expression, respectively, at a significance threshold of adjusted p-value of <0.1 were treated as foreground and all other assessed peaks/genes as background. For annotation purposes, peaks were mapped to the nearest gene such that all features could be assigned to an annotated gene for subsequent steps. We carried out enrichment using three sets of features. The first consists of GO-terms downloaded from Echinobase (https://www.echinobase.org/). In the second set, we mapped each sea urchin gene to its most similar human sequence via reciprocal best BLAST mapping to access the GO, GO-SLIM, and PANTHER annotation categories developed for humans. Our third set consisted of custom mapping of developmental genes to specific developmental lineages and was used primarily as an additional check on our cluster annotations (Ashby, 2006). Categories were considered significant at a p-value of 0.05. Because of extensive hierarchical relationships among the categories, no multiple testing correction was performed.

### Genome and gene sets

Throughout this manuscript, the spur_5.0 version of the *S. purpuratus* genome was used (downloaded from https://www.echinobase.org/, (Foley *et al*, 2021), with the following modifications for scRNA-seq. Due to significant losses at the 3’ end of several genes, the last exon of each transcript was extended under the following circumstances: (1) MACS2-called peak within 10kb downstream of the 3’end of the gene on the same strand and upstream of the next gene, based on our scRNA-seq data, and (2) continuous coverage between current annotation and the above called MACS2 peak, based on combined bulk and scRNA-seq data. This affected a total of 9,475 transcripts (out of 43,104) from 6,401 genes (out of 30,906), which were extended on average 519.10bp (maximum 4,725bp) and included genes with known function in early sea urchin development (e.g. Wnt8, Endo16).

Gene sets used in this manuscript are based on gene ontology (GO) terms and the following terms were used to define gene sets: TF (GO:0000981, GO:0006355, GO:0003700 or GO:0000976), translation (GO:0006412), development-related genes (GO:003250) and metabolism related genes (GO:0008152).

### Statistical analysis

For this study, significance levels of p <0.05 were considered significant for GOterm analysis and TSS Enrichment. For differential gene expression, significance was calculated using a Wilcoxon Rank Sum test and for cell-type-specific regulatory elements (scATAC-seq), a logistic regression framework was used (LR test in Seurat). P-values were adjusted using Bonferroni correction. In both cases, a p-adjusted value of 0.1 and a logFC threshold of 0.25 was considered significant.

### Code and Data availability

Data used in this study have been uploaded to GEO [coming soon], the code used for this analysis is available at https://github.com/BrandeJo/scEarlyDev_SPU. BigWig files and peak sets for visualization can be found at Zenodo: **10.5281/zenodo.16815709**

## Results

### Cell fate and lineage commitment in the sea urchin profiled by single-cell sequencing

To understand the gene regulatory events driving cell type diversification in the sea urchin embryo, we profiled gene expression changes and regulatory element usage from early cleavage stages through early larval development (Fig 1A). Using scRNA-seq and scATAC-seq, we identified 47 distinct transcriptomic states based on unsupervised clustering of gene expression data, and 37 distinct regulatory states based on patterns of open chromatin, which we annotated based on previously reported marker genes (Fig. 1 B,C, Table S2, Fig. S1). The recovered cell types include all major developmental lineages known in the sea urchin (PMCs, Oral and Aboral NSM, Apical, Oral, Aboral and Ciliary Band Ectoderm, Neurons and Endoderm), including primordial germ cells (small micromere descendants), and immune cell subsets (filopodial cells, globular cells). At higher levels of resolution, we also obtain clear sub-clusters of skeletogenic and neuronal cells within the mesoderm and ectoderm, respectively (Fig S1).

**Figure 1.**
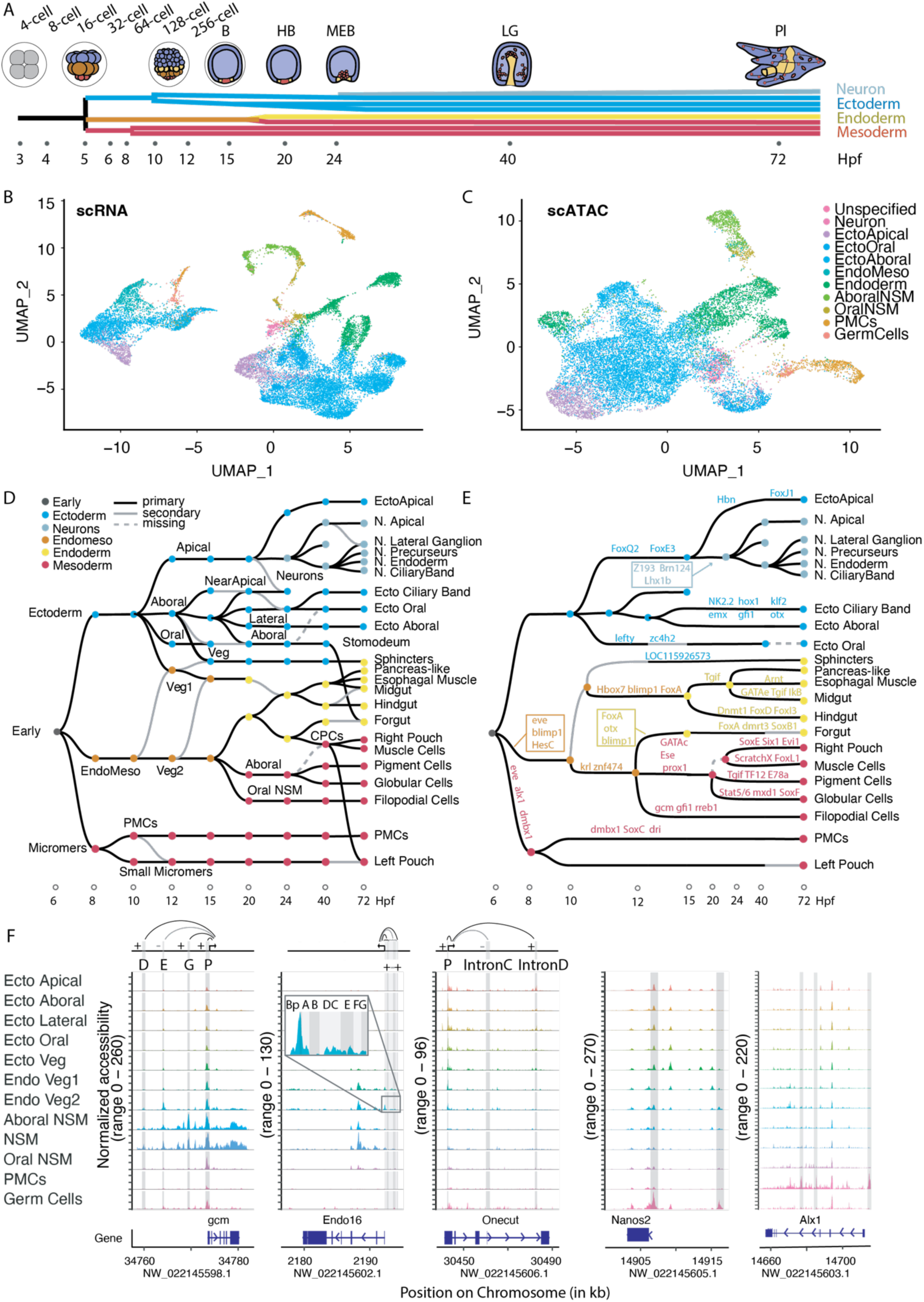
Cell fate and lineage commitment in the sea urchin profiled by single-cell sequencing. (A) Lineage cell fate map in the sea urchin, highlighting germ layer and neuron lineage specification. Sampling timepoints in hours are indicated as dots. (B,C) UMAP representations of 4-cell embryo to pluteus stage sea urchin profiled by scRNA-seq (B) and scATAC-seq (C) colored by major cell lineages in the developing sea urchin embryo. (D) Lineage tree of sea urchin development inferred by “ancestry voting” from scRNA-seq data. Edges are either marked as primary links (black), secondary links (gray line, see Methods) or missing in the case of known lineage edges that are not captured in the analysis (dashed line). Colors of nodes indicate mayor developmental lineages. (E) Lineage tree of major lineages from D (modified for better legibility) with most DE candidate lineage specifying TFs throughout development. Colors of nodes indicate mayor developmental lineages, as in D. (F) Coverage plot for mayor lineages in scATAC-seq, showing aggregated normalized read coverage at lineage level, gene location of one marker gene for each lineage is shown. Regulatory elements of interest (i.e. known CRM for gcm, Endo16 and onecut, and lineage specific regulatory elements for Nanos2 and Alx1) are highlighted in gray. For gcm, Endo16 and onecut, the respective effect on transcription for each known regulatory element is indicated by + (enhancing transcription) or – (repressive). For better legibility, the promoter region of Endo16 has be enlarged.

To gain insights into genes and TFs driving cell type diversification in the sea urchin, we reconstructed developmental lineage relationships using ‘ancestry voting’ and used these relationships to identify pattern of TF expression concurrent with changes in cell state. The resulting lineage tree closely matches the lineage relationships known from classic lineage tracing experiments, with micromere descendants (small micromeres and prospective PMCs) being distinct lineages starting at the 64-cell stage (Fig. 1D) and neuronal cell types emerging from apical ectoderm cells around mesenchyme blastula (MEB, Hpf24) stage. Distinct oral and aboral NSM lineages are evident from hatched blastula (HB, Hpf20) stage onwards (Fig. 1D, Fig. S2A).

In a few cases, incomplete lineage resolution or converging transcriptomic states potentially hinder inference of the correct lineage tree using our approach. For example, the foregut and the stomodeal ectoderm together form the esophagus in pluteus larvae, which have highly similar gene expression profiles. Here, the stomodeal ectoderm is predicted to be the origin of 41% of foregut cells (Fig. S2A). A similar case occurs in the blastula endoderm, where Veg1 and Veg2 endoderm are both predicted to mainly stem from Veg2 Endoderm (52% and 100% respectively, Fig. S2A), likely reflecting the fact that Veg1 Endoderm acquires a transcriptomic state similar to the Veg2 Endoderm in the previous stage. Finally, the aboral ectoderm appears to be a significant contributor to many ectodermal lineages, potentially reflecting that ectodermal lineages (with the exception of the apical ectoderm) are only fully resolved at late blastula stages (Li *et al*, 2014) (Fig. 1D, Fig. S2A).

Many of the TFs we identified (e.g. *Alx1* in PMCs and *FoxA* in the endoderm, Table S8) have been previously identified, highlighting the accuracy of the method. However, our data further suggest developmental roles for previously uncharacterized TFs, most prominently *Znf875/Z193* (LOC581084) in neurogenesis, *Zc4h2* (LOC575413) in the oral ectoderm, and *Rreb1* (LOC578691) in globular cell specification (Fig. S2B). In support of this inference, multiple *cis*-regulatory elements associated with these TFs are also specifically accessible in the respective lineages (Fig. S2C). Most candidate lineage defining TFs are used in only a single territory in the measured timespan, but, interestingly, TFs that are reused in multiple territories are not necessarily redeployed in related territories, with at least 8 candidate TFs shared between ectoderm and NSM and 4 candidate TFs shared between ectoderm and PMCs (Fig S2D-F). While TF expression may be shared, regulatory site usage, particularly in introns, is typically distinct between territories. For example, the TF *Rreb* (LOC578691) is expressed in PMCs until blastula stage, and subsequently restricted to the aboral NSM and, eventually, the globular cells (Fig S2B). However, distinct regulatory elements flanking this gene are accessible in each of these tissues (Fig. S2C), highlighting the diversity of regulatory states that can be encoded by TF::DNA interactions with a limited set of TFs.

To assess the quality of our scATAC-seq data and annotations, we compared our data to validated *cis*-regulatory elements associated with lineage specific gene expression (Ransick & Davidson, 2006; Yuh *et al*, 1996; Barsi & Davidson, 2016; Ransick & Davidson, 2012) and PMC-specific regulatory elements identified in a recent genome-wide screen (Shashikant *et al*, 2018). Convincingly, we recover cell type-specific regulatory element patterns corresponding to published reporter assays (Fig. 1F). In the case of *gcm*, for example, four regulatory elements have been previously defined (Ransick & Davidson, 2006, 2012) (D,E,G and P), out of which D and G are cell-type-specific distal elements (NSM and Aboral NSM), while the promoter, as for many developmental genes, shows reduced cell type-specificity (Reddington *et al*, 2020). Functionally, the distal E-element is important for the repression of *gcm* expression in endodermal cells. Consequently, this element is detected most prominently in the Veg2 endoderm in our dataset. Similar patterns are observed for *Endo16* and *Onecut*, where we readily detect accessible elements known to regulate gene expression. Notably, the promoters are often accessibly even in tissues in which the gene is not expressed. While the published promoter-centric, upstream regulatory elements appear to be sufficient recapitulate endogenous gene expression patterns in laboratory studies, we observe many more specifically accessible regions in proximity to and within the gene body of these genes, suggesting much more extensive gene regulation than captured in most reporter assays or strictly necessary for driving gene expression. Newly identified cell-type-specific regulatory elements include those associated with PMCs (e.g. *Alx1*) and primordial germ cells (*Nanos2*, Fig 1F), as well as all other major lineages in sea urchin development (Tables S6, S7). Taken together, our dataset gives insights into the regulatory complexity of the developing sea urchin, both on the transcriptomic and *cis*-regulatory level.

### Zygotic genome activation and chromatin maturation occur simultaneously with the loss of developmental plasticity in the sea urchin

Zygotic transcription in sea urchin embryos has been detected as early as 8-cell stage (Wilt, 1970) and has been linked to the earliest cell-fate specification events at 4th cleavage, when micromeres specifically express Pmar1 and Alx1 (Ettensohn *et al*, 2003; Oliveri *et al*, 2003). It remains unclear, however, if these earliest transcriptional events are part of widespread zygotic genome activation or represent individually controlled, pre-ZGA gene expression events.

A challenge in evaluating ZGA in the sea urchin is that an estimated 76% of the zygotically expressed transcripts in the sea urchin are also maternally loaded (Flytzanis *et al*, 1982). To overcome this issue, we made use of scRNA-seq combined with metabolic labeling (scSLAM-seq (Uhlitz *et al*, 2021; Holler *et al*, 2021)), allowing us to distinguish zygotic from maternal transcripts on the basis of T>C transitions induced by 4sU incorporation into nascent mRNA (Fig S3A-G). While the low cell numbers sampled at early time points combined with high levels of SNPs in the sea urchin do not allow for confident estimates at the level of individual transcripts, the resulting genome and category-wide estimates are sufficiently robust to allow for general insights. For example, while most lineages show a uniform incorporation of 4sU, PGCs show noticeably reduced levels of 4sU incorporation, consistent with previously reported reduced transcriptional activity (Oulhen & Wessel, 2017) (Fig S3H,I).

Within our data, distinct clusters in the scRNA and scATAC data can first be observed at the 128-cell stage (Hpf10, Fig. S1), when a cluster of micromeres becomes distinct from the remainder of cells. Concurrently, per-cell 4sU incorporation increased markedly from the 128-cell to 512-cell stage, as does the fraction of unspliced reads and the number of upregulated transcripts observed in our bulkRNA dataset (Fig. 2A, Fig. S4A-C). Collectively, these data suggests a primary activation of the zygotic genome at the 128 cell stage concomitant with the loss of cell-fate plasticity observed in classical micromere transplantation experiments (Hörstadius, 1936).

**Figure 2.**
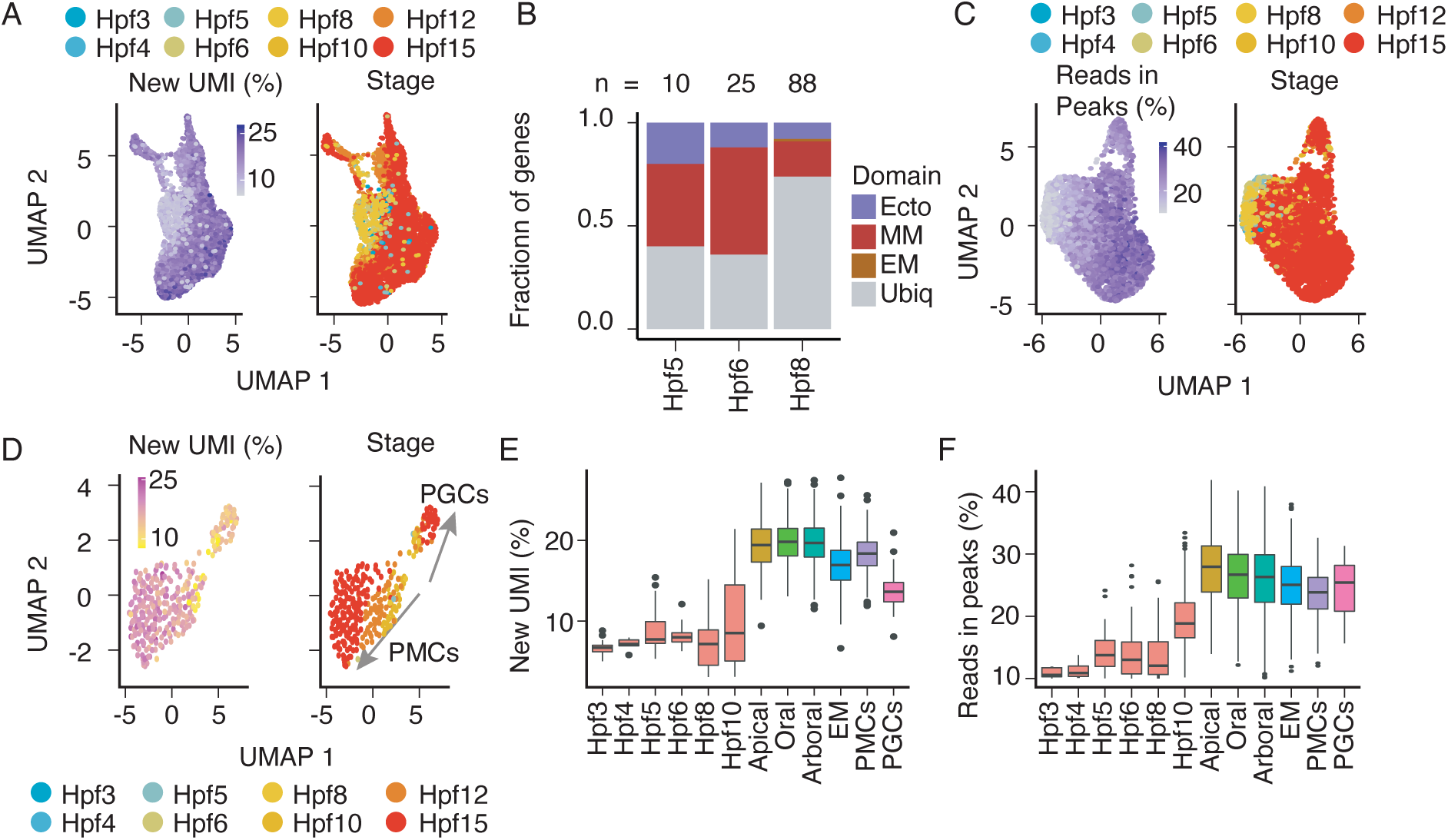
Zygotic genome activation in the sea urchin occurs simultaneously with cell type diversification and chromatin maturation. (A) UMAP plot of early sea urchin development obtained from scRNA-seq (Hpf3 to Hpf15) colored by fraction of new (zygotic) mRNA molecules (left) or developmental stage (right). (B) Stacked bar chart showing the fraction of genes upregulated pre-ZGA in each embryonic territory. Total numbers of (detected) newly upregulated genes per stage are indicated on top. (C) UMAP plot of early sea urchin development obtained from scATAC-seq (Hpf3 to Hpf15) colored by fraction of reads in peaks (left) or developmental stage (right). (D) UMAP plot of sub-clustering of the micromere lineage from B and colored by fraction of new (zygotic) mRNA molecules or developmental stage (right). Arrows indicate the developmental progression of each lineage. Note the reduced fraction of new UMIs in the primordial germ cells (PGCs). (E,F) Box and whiskers plots of fraction of new mRNA molecules per cell lineage (scRNA-seq, E), or fraction of reads in peaks (scATAC-seq, F) in *S. purpuratus* development, following ZGA and until blastula stage (Hpf12-Hpf20), colored by cell lineage. Earlier timepoints (Hpf3 – Hpf10) are indicated at whole embryo level for reference. EM = Endomesoderm, MM=Micromeres, sMM=small micromeres, Ecto= Ectoderm, Ubiq=Ubiquitous, Apical and Aboral refer to the respective ectoderm.

In line with this observation, changes in the developmental direction in PCA occur at the whole embryo level in my bulk RNA-seq developmental time course.

Similar changes in the developmental direction, and thus changes to the transcriptome, occur also at the 16-cell stage, when the earliest upregulated genes are detected in the bulk RNA-seq time course. Further, the 512-cell stage (Hpf15), an early blastula stage marked by the patterning of ectodermal and endomesodermal territories and the onset of gastrulation (MEB stage, Hpf24) represent stages where large shifts to the zygotic transcriptome occur (Fig. S4D).

While most of the nascent transcription appears to be associated with the major wave of ZGA, a set of 89 genes shows clear evidence of upregulation as early as 16-cell stage (Hpf5, Fig S4E) based on increased expression relative to the 4-cell embryo (potentially missing genes with smooth transitions from maternal to zygotic expression). Interestingly, a large fraction of the upregulated genes are marked by their cell-type-specificity even before Hpf10 (60% of newly expressed genes at 16-cells (Hpf5), 63% at 32-cell (Hpf6) and <25% at 64-cell (Hpf8) and later stages, Fig. 2B), suggestive of local regulatory effects that precede the major wave of ZGA in a cell-type specific manner.

In other metazoan animals, including the sea urchin *Paracentrorus lividus*, the chromatin becomes locally more compacted, and a more mature regulatory landscape appears at the time of ZGA (Pálfy *et al*, 2020; Vastenhouw *et al*, 2019; Marlétaz *et al*, 2023a). Similarly, major organization of the regulatory genome also starts to emerge at the 128-cell stage (Hpf10, Fig. 2C, Fig. S4F. The fraction of reads in peaks, as well as transcription start site enrichment (TSSE) increases from the 128-cell (Hpf10) stage onwards, reaching a plateau of per cell TSSE scores by 512-cell (Hpf15) stage, suggesting a maturation of the regulatory landscape at this developmental interval (Fig. S5D) with few peaks appearing before ZGA (Table S9 discussed below).

It is noteworthy that at the time of the major wave of ZGA (Hpf10, 128-cell stage), the potential of cells of the animal half to acquire mesodermal fates is lost in classic experiments (Hörstadius, 1936). Thus, the major activation of the zygotic genome co-occurs with the establishment of cell-type-specific transcriptional states, the establishment of a defined cell-type specific regulatory landscape and the loss of developmental plasticity in the sea urchin.

Of note, while PGCs show reduced 4sU incorporation consistent with transcriptional repression (Fig 2E), the fraction of reads in peaks and the transcription start site enrichment - markers of mature and accessible chromatin - is similar between small micromeres (contributing to the PGCs) and related lineages (e.g. PMCs, Fig 2F).Furthermore, the levels of 4sU incorporation are higher than for pre-ZGA stages, suggesting an activation of the zygotic genome within these cells, followed by widespread, active transcriptional repression (Fig. 2E,F, S3H,I, S4G) that leaves intact the mature regulatory architecture of transcriptionally active cells.

### A distinct set of accessible regions is present in the sea urchin before the emergence of a defined regulatory landscape

The earliest gene expression events in our data (pre ZGA) are enriched for GOterms associated with development, including TF binding (e.g. regulatory region nucleic acid binding) and signaling (e.g. signaling receptor activity, cytokine activity, Fig 3A), and genes in the vicinity of early accessible regulatory elements in bulk ATAC-seq data are likewise enriched for GOterms related to TF binding with additional enrichments for nucleotidase/ATPase and hydrolase activity (Fig 3B). Establishing direct links between specific open chromatin peaks and gene expression is difficult, but these shared enrichments suggested a coordinated biological function to these early opening peaks related to priming the regulatory genome.

**Figure 3.**
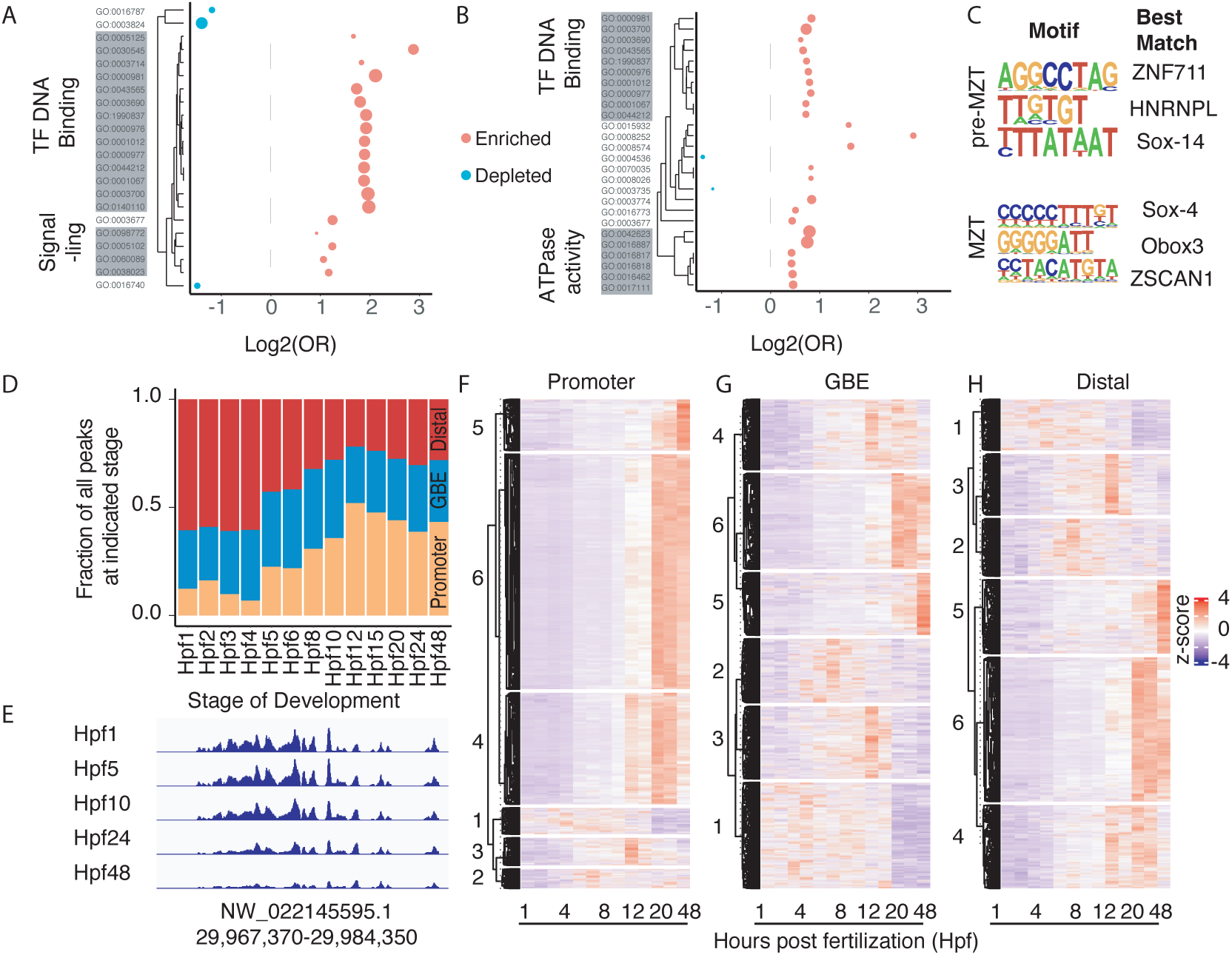
Early expressed genes are dominated by lineage-specific transcription factors. (A,B) Dot plot of GOterms enriched in genes upregulated prior to ZGA (A) and genes associated with regulatory elements present before ZGA (B). Size of dots is inversely proportional to the p-value of the enrichment, values on the x axis represent the log2 odds ratio. GOterms are clustered by overlaps in the gene sets. (C) Motif plot of top 3 DNA motifs significantly enriched in accessible regions and with similarity >0.8 (Homer score) to mammalian or insect TFB-motif, pre-ZGA (16 to 32 cell stage) and around ZGA (64-to 256 cells). Best match refers to most similar human/insect TFB-motif. (D) Stacked bar chart of the position of accessible DNA regions with respect to closest gene for each stage in sea urchin development. (E) Genome Browser tracks of bulk ATAC-seq at representative stages of sea urchin development at a genomic region that is well defined at Hpf1. (F-H) Clustered heatmap of accessibility at Promoters (F), Gene body elements (G) or distal elements (H) by developmental stage, based on bulk ATAC-seq dataset.

To better understand these early regulatory peaks, we searched for motifs enriched in pre- vs. post-ZGA regulatory elements. To avoid biases associated with having low representation of cells at the earliest stages in our single-cell data, we conducted these analyses using a bulk time course. Contrasting pre-and post-ZGA peaks, we identified a clear set of motifs that are present in newly established, pre-ZGA peaks (appearing at 16-32 cell stage), with motifs bearing similarity to human ZNF711 and Sox14 being the most dominant (Fig 3C, Table S10). A different Sox4-like motif is enriched at peaks opening at ZGA/MZT (60-to 256-cell stages), along with a ZSCAN1-like and a MZT1/Obox3-like motif. While the corresponding TFs and functions in the sea urchin are unknown, the enriched motifs indicate a distinct grammar for these early peaks and suggest a role of Sox-like TFs prior to and at ZGA in the sea urchin and highlight the specificity of early accessible chromatin.

Regulatory elements accessible in early development, particularly those that remain open in subsequent stages, have been shown in other species to be enriched for structural elements and the binding of insulator factors (e.g. TAD boundaries) (Reddington *et al*, 2020). In our bulk ATAC-seq dataset, around 4,500 regions are accessible before Hpf10, a large portion of which (61.4%) are distal to any gene active in early development. As development proceeds, promoter peaks are established (increasing from ∼10% to >50% of all peaks) at which point fewer than 25% of accessible regions are distal (Fig 3D).

Interestingly, among the earliest accessible peaks (Hpf1, zygote), several peaks are grouped together in regions with apparently mature chromatin landscape well before ZGA (Fig 3E). In total, we identified 11 groups of early peaks, encompassing 52 genes (enriched for ribosomal RNA, tRNAs and early histone genes), and accounting for 21% of accessible regions at Hpf1. Many of these early peak regions remain accessible throughout development, suggesting a structural role, while others close as development proceeds (Fig 3E; Table S9).

### Regulatory elements open with distinct temporal dynamics in developing sea urchins

To characterize the temporal dynamics of accessibility more globally, we clustered all accessible regions (peaks) identified in our bulk ATAC dataset (Fig 3F-H). We identified 6 groups of temporal dynamics, based on hierarchical clustering, including the above mentioned transient early peaks. Other sets of dynamic elements are early transitory peaks (accessible by Hpf5 through early blastula stages), peaks that are accessible mainly around blastula stage, peaks accessible mainly around at the gastrula stage, and, most common, peaks that are accessible at all late (post-gastrulation) stages. These sets of peaks differ notably in the relative proportion of distal and promoter-proximal peaks. While most promoter proximal peaks tend to gain accessibility around Hpf10 to Hpf12 (early blastula stages) and remain accessible throughout development, distal elements tend to become fully accessible on later in development to be more stage specific (Fig 3F-H, S5A-D). This trend parallels the establishment of new peaks by stage, which appears to proceed in two waves. The first one (around ZGA) is dominated by promoter proximal peaks and the second, potentially associated with an increasing diversity of cell types, shows an increase in the proportion of distal peaks. These findings are similar to patterns observed in other metazoans (Reddington *et al*, 2020) and highlight the importance of examining intergenic peaks for understanding cell type specification during development.

### The cell-type-specific regulatory landscape is defined by germ layer identity and gene body regulatory elements in the sea urchin

Correct spatiotemporal expression of genes during development is coordinated by complex, and often overlapping, activities of diverse regulatory sites (Spitz & Furlong, 2012). While there is clear differential CRE usage across tissues from late cleavage stages onwards, there are also instances, mainly among related cell types, of shared regulatory site usage. To characterize shared usage regulatory information across the genome and among cell types, we used a HMM-based approach (scregseg, (McGarvey *et al*, 2022)) to group accessible regions from cell-type level pseudo-bulk data from cell types with > 10 cells into distinct regulatory states. Using this approach, we find that cell types primarily cluster by germ layer at later stages of development (Fig. 4A): Mesoderm, Endoderm and Ectoderm each cluster together, suggesting extensively shared regulatory site usage by diverse cell lineages of a common germ-layer of origin. Within the mesodermal clade, we find a subgroup for skeletogeneic cells (PMCs) and one for the non-skeletogenic mesoderm, highlighting their distinct fates early on, while within the late mesodermal clade, two subgroups are present, one for late skeletogenic cells (PMCs) and filopodial cells, and one for the aboral NSM derived cell types, revealing a surprising overlap in regulatory element usage between filopodial and skeletogenic cells. Interestingly, early PMCs and late blastula GermCells (small micromere descendants, mesodermal origin) cluster together in a distinct group, highlighting the shared origin of these cell types.

**Figure 4.**
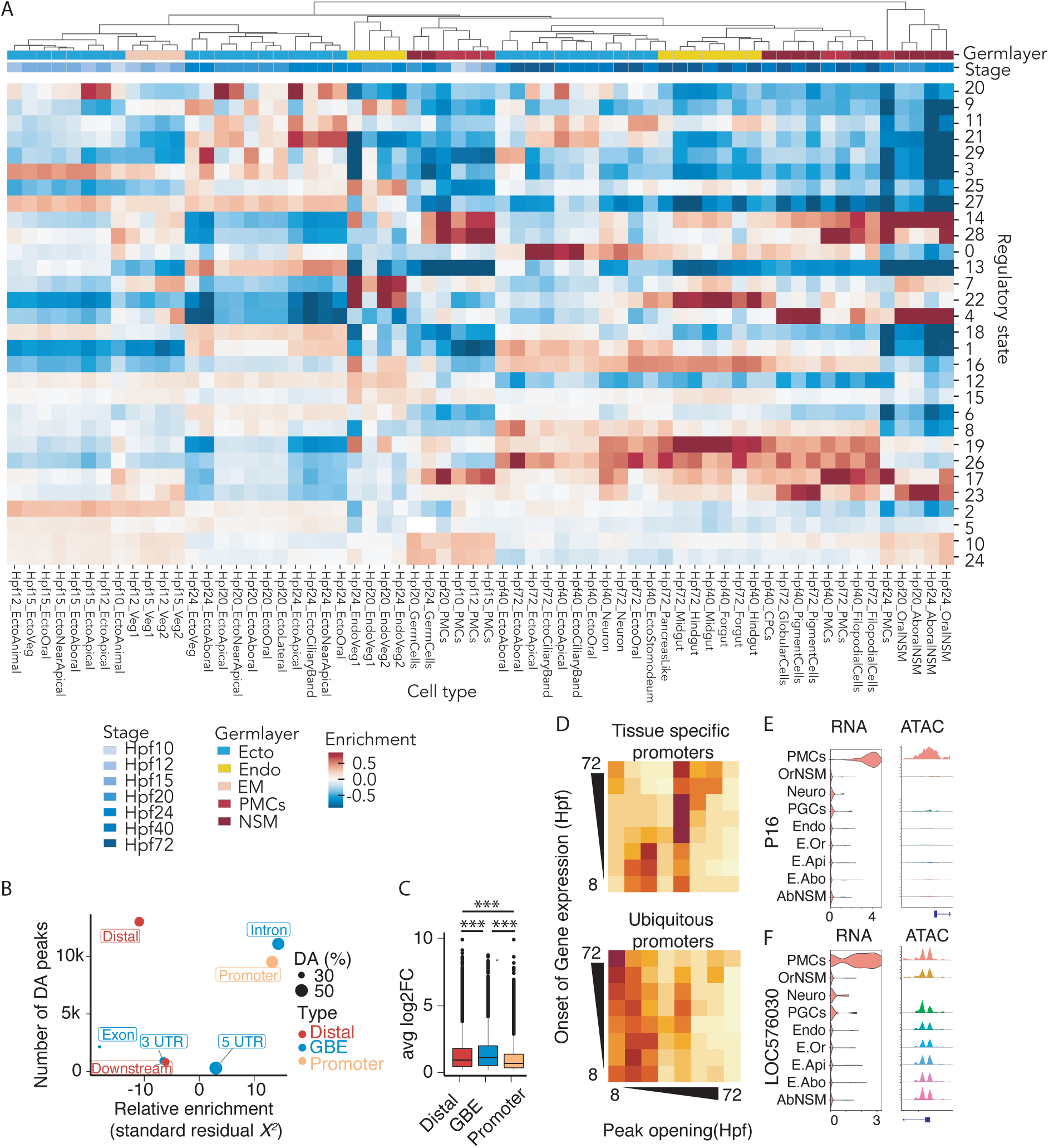
Sea urchin cell-type-specific regulatory landscape is clustering by germ layer and defined by gene body regulatory elements. (A) Heatmap of hierarchical clustering of cell types throughout sea urchin development, columns represent individual cell types and rows indicate regulatory states identifies by scregseg segmentation approach. (B) Bubble plot showing the number of differentially accessible regulatory elements (y-axis) and the over/underrepresentation (standardized chi-squared residual x-axis) for each type of regulatory elements. Size of points is proportional to the fraction of regulatory elements of each class that are cell-type-specific at any point of development. (C) Box and whiskers plot of average log2fold change of differentially accessible elements across cell types. (D) Heatmap of temporal difference between promoter opening and onset of gene expression for lineage specific genes in sea urchin development, plotted for cell-type-specific promoters and ubiquitously accessible promoters. (E,F) Violin plot of expression and coverage plot of promoter accessibility of P16 (E) and LOC576030 (F) by cell type at Hpf24 (MEB). Note that no neurons were found at this time of development in the ATAC dataset. *** indicate significance levels of p < 0.001, ns=non-significant ,Tukey-Kramer test.

Within the ectodermal clade, several subgroups were identified, namely neurogenic ectoderm (late ciliary band ectoderm, apical ectoderm, and oral ectoderm) and blastula ectoderm.

While clustering of ectodermal cell types appears to be stage specific, especially at earlier developmental stages, specific regulatory states can be assigned to the apical ectoderm (State 20) and the aboral ectoderm (State 29, Fig 4A).

Pre-blastula cell types formed a distinct group, where early PMCs and endomesoderm were distinct from early ectodermal accessibility profiles, suggesting that the vegetal and animal half of the embryo share distinct regulatory landscapes from an early stage. Interestingly, a predominantly endodermal state (State 7) is already enriched in the “veg1” and “veg2” endomesoderm, both of which eventually contribute to the endoderm. In addition, a predominantly mesodermal regulatory state (State 4) is enriched in “veg2” but not “veg1” endomesoderm of early stages (Hpf12, Hpf15), reflecting the exclusive “veg2”-origin of this tissue (Fig 4A). Taken together, these results indicate widespread shared accessibility of *cis*-regulatory elements between cell types, especially those of the same germ layer.

Underlying our inferred regulatory states are distinct regulatory sequence grammars that can be identified in the form of motif enrichments. Motifs enriched in each regulatory state are mostly associated with well-known lineage TFs (Table S11), e.g. PMC states dominated by Alx1, Ets1 and Jun/Fos), mirroring motif enrichment seen via differential accessibility tests in cell types and stages (Tables S12, S13). Among the regulatory states associated with skeletogenic fates, only state 17 is enriched in late skeletogenic cells, while state 28 shows pan-PMC accessibility. Interestingly, states 17 and 28 were also enriched in late oral NSM (Hpf24 to Hpf72), highlighting the similarity of the regulatory landscape in these cell types during a period in which there appears to be some plasticity between NSM and skeletogenic lineages. Taken together, the data suggests that regulatory element accessibility is strongly associated with germ layer identity throughout sea urchin development, with noteworthy similarities in Ets1-dominated filopodial and skeletogenic cell regulatory landscapes.

### Promoter accessibility is often decoupled from gene expression

Most annotated regulatory elements in the sea urchin, including those characterized as sufficient for recapitulating endogenous expression in transgenic assays, are located proximal to the TSS. However, in vertebrates and other metazoans, distal and intronic elements have been found to be of importance (Davidson, 2006). We therefore wondered which types of peaks were most associated with the cell-type-specific gene expression in the sea urchin. To address this question, we investigated correlations between peak accessibility and gene expression, the number of and types of peaks with differential accessibility per time point, and the average log fold change of differentially accessible peaks as measures of distinctness between cell-types.

In terms of average log fold change among cell types, distal regulatory are elements are the most distinct as well as the most numerous cell-type specific element (Tukey-Kramer test, p < 0.001 for all comparisons, Fig. 4B) with ∼11 000 cell-type-specific gene body elements, ∼12 500 cell-type-specific distal elements, and 9 500 cell-type-specific promoters identified in our dataset. In terms of logFC between cell types, promoters show the least variation among cell types (Fig 4C), though differentially accessible promoters, 5’UTR, and intronic elements are overrepresented among all DA peaks. Conversely distal, 3’UTR, downstream and exonic elements are underrepresented (Fig 4B Chi-squared test, p-value < 2.2e-16)

Overall, gene expression correlated well with promoter accessibility, especially from Hpf10 onwards (r∼0.5) (Fig 4D, Fig S6 A-B,F). Following the blastula stage, gene body elements show a slightly higher correlation with gene expression than promoters, while distal elements (>1kb from TSS) only exhibit weak correlation with gene expression (r∼0.2). However, while many known cell-type specific regulators do show cell-type specific promoter accessibility (e.g. P16, Fig. 4E), a large fraction does not (e.g. >60% at Hpf24), showing instead ubiquitous accessibility despite cell type specific expression (e.g. LOC576030, Fig 4F).

To gain further insight into the timing of gene expression changes relative to changes in promoter accessibility, we examined correlations between the accumulation of unspliced reads and the opening of promoter-proximal peaks for cell-type specific genes. We observe a bimodal distribution of relationships, with about half of the promoters opening around Hpf10 (128-cell stage) and gene expression starting at later stages of development and the other half opening shortly before the onset of gene expression (Fig. 4D). Interestingly, the former group of promoters corresponds largely to ubiquitously accessible promoters, while the latter group is enriched for cell-type specific promoters (Fig S6A). Other regulatory elements (distal and gene body) show a higher correspondence between promoter opening and onset of gene expression, as seen for enhancers in other species (Fig. S6B,C).

As observed in other metazoans, genes with more complex expression patterns tend to have more regulatory elements (more complex local regulatory landscapes, Fig S6D) (Floc’hlay *et al*, 2021). We further note that cell-type-specific (or differentially accessible) regulatory elements are also more frequent in these groups. Other groups of genes with a high number of cell-type-specific regulatory elements are developmental genes and TFs (Fig S6E). Thus, complex expression patterns and pleiotropic function correlates with more complex cell-type-specific regulatory landscapes in the sea urchin.

### Neurodevelopment within sea urchin ectoderm proceeds in three distinct lineages

Neurogenesis in the sea urchin begins during the mesenchyme blastula stage (Hpf24) (Garner *et al*, 2016) with the first neurons formed through the differentiation of apical ectodermal cells into serotonergic apical neurons followed by neurodevelopment originating in the oral ectoderm, ciliary band, and endoderm (McClay *et al*, 2018; Wei *et al*, 2016). To better understand the mechanism of this differentiation, we pooled all neuronal cells (∼260) identified at all time points and performed additional sub-clustering, resulting in nine distinct neuronal transcriptional states in the sea urchin pluteus, corresponding to neuronal precursors (1 cluster), apical neurons (3 clusters), ciliary band neurons (2 clusters), dopaminergic neurons (lateral ganglion, 2 clusters) and endodermal neurons (1 cluster, Fig. 5A,B). However, the presence of small groups of cells with highly specific marker gene expression within these clusters suggests an even greater diversity of neurons in the relatively simple sea urchin larva.

**Figure 5.**
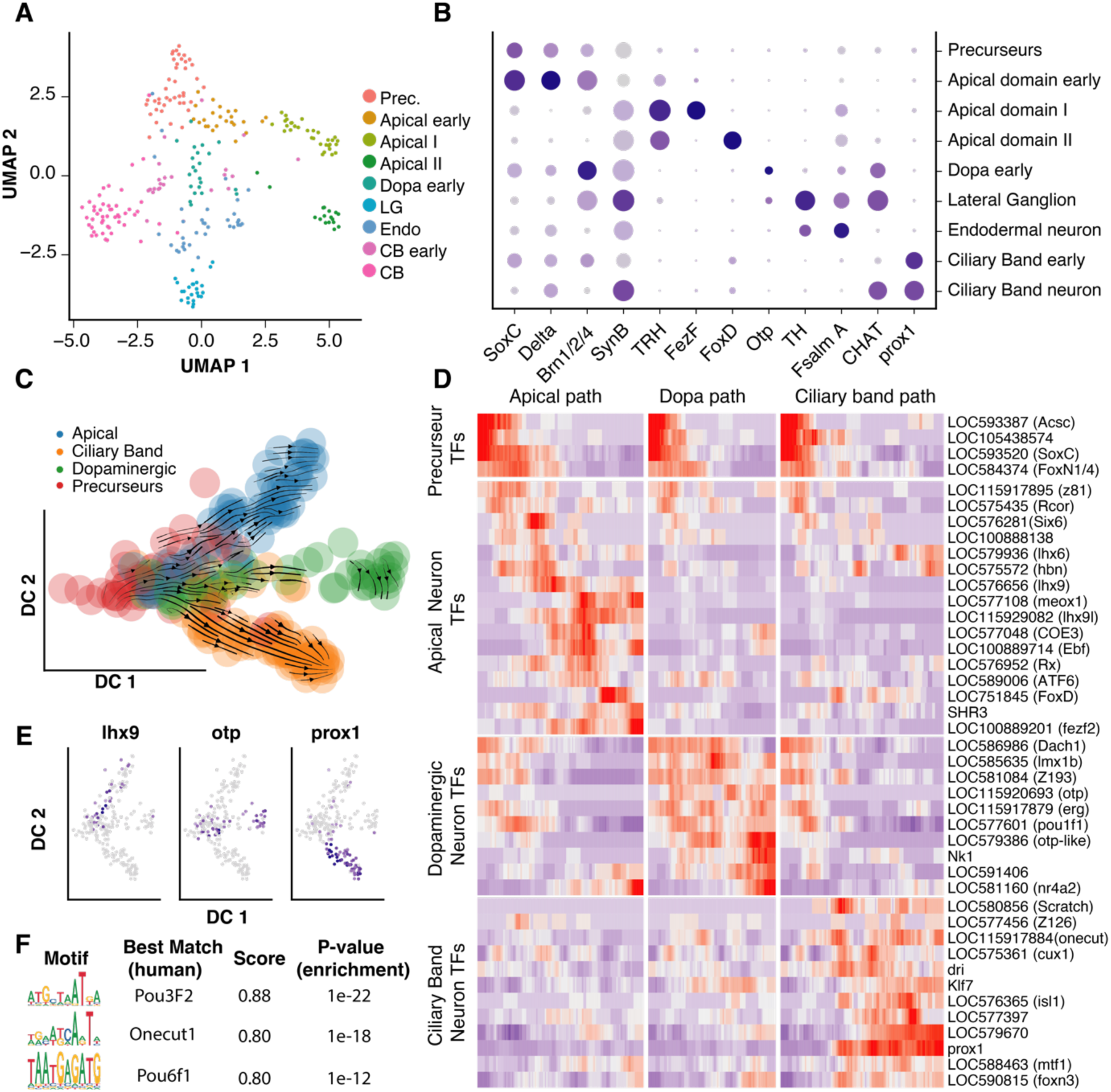
Neurodevelopment in the ectoderm of developing sea urchins proceeds in three lineages. (A) UMAP clustering and annotation of neuronal cells from scRNA-seq dataset, colored by neuronal type. (B) Dot plot of neuronal marker genes used for annotation of neuronal subsets. (C) Diffusion map of neurodevelopment in sea urchin development, colored by neuronal lineage (note that endodermal neurons were removed here). Arrows indicate RNA velocities, colored by neuronal lineage. (D) Smoothed heatmap of expression levels of neuron-specific TFs in each branch of neurodifferentiation in the sea urchin. (E) Diffusion map of neurodevelopment in sea urchin colored by expression level of three lineage specific TFs (*lhx9* – apical neurons, *otp* – dopaminergic neurons, *prox1* – Ciliary band neurons) (F) Enriched motifs in neuron specific peaks at pluteus stage (Hpf72) with high similarity to human/insect TFB-motifs (homer score > 0.8) and their closest match in humans. For a full list of enriched motifs, see supplementary table 12. Prec.= precursors, LG = Lateral ganglion, CB = Ciliary band.

Using diffusion maps (Haghverdi *et al*, 2015), we were able to link the nine clusters into three distinct developmental trajectories corresponding to serotonergic (apical), dopaminergic and ciliary band trajectories, with distinct transcriptional regulation (Fig. 5C-E). Neuronal precursors were defined by expression of SoxC and FoxN1/4 TFs while later neuronal populations were demarcated by the expression of TFs such as prox1 for ciliary band neurons, fezf2 and lhx9 for serotonergic neurons, and otp and lmx1 for dopaminergic neurons (Fig. 5 B,D,E, Table S11). Among the TFs that were upregulated in either of these developmental trajectories (Fig. 5D), 32 could be linked to orthologs in mammals or insects, 26 of which have a known function in neuro- or brain development. Further, six of the remaining orthologs have been found to be involved in the formation of related structures (e.g. retina, somite, face development; Table S14).

In contrast to scRNA-seq data of neurons, we only identified three sub-clusters of neurons in the scATAC-seq dataset, a difference we suspect resulted from the limited resolution of the late-stage scATAC-seq data. Out of these clusters, two clearly correspond to neuronal identities based on the overlap in marker genes scores with our scRNA-seq dataset: Ciliary Band Neurons and remaining ectodermal neurons (Fig. S7). To further characterized these clusters, we looked for TF motifs enriched in neuron specific peaks. Three enriched motifs could be confidently assigned to vertebrate TF binding motifs (homer score > 0.8), namely those matching mammalian/fly Oct6, Onecut1, and Pou6f1 (Fig. 5F, Table S12), supporting a highly conserved regulatory code with mammalian neurons.

### Diversity of gene regulation in skeletogeneic cells of the sea urchin pluteus

Skeletogenic cells in the sea urchin have previously been reported to be transcriptionally (Sun & Ettensohn, 2014; Guss & Ettensohn, 1997; Illies *et al*, 2002) and functionally diverse (Hu *et al*, 2020; Chang *et al*, 2021). We therefore sought to profile these cells and the TFs driving differentiation during skeletal formation as well as to more fully characterize the diversity of skeletogenic states. After sub-clustering annotated skeletogenetic cells, we identified three distinct sets of skeletogenic cells that we could annotate on the basis of known gene expression patterns including MSP130 and P16 are in line with previously reported expression patterns (Sun & Ettensohn, 2014) (Fig. 6A, B, Fig. S8A,B).

**Figure 6.**
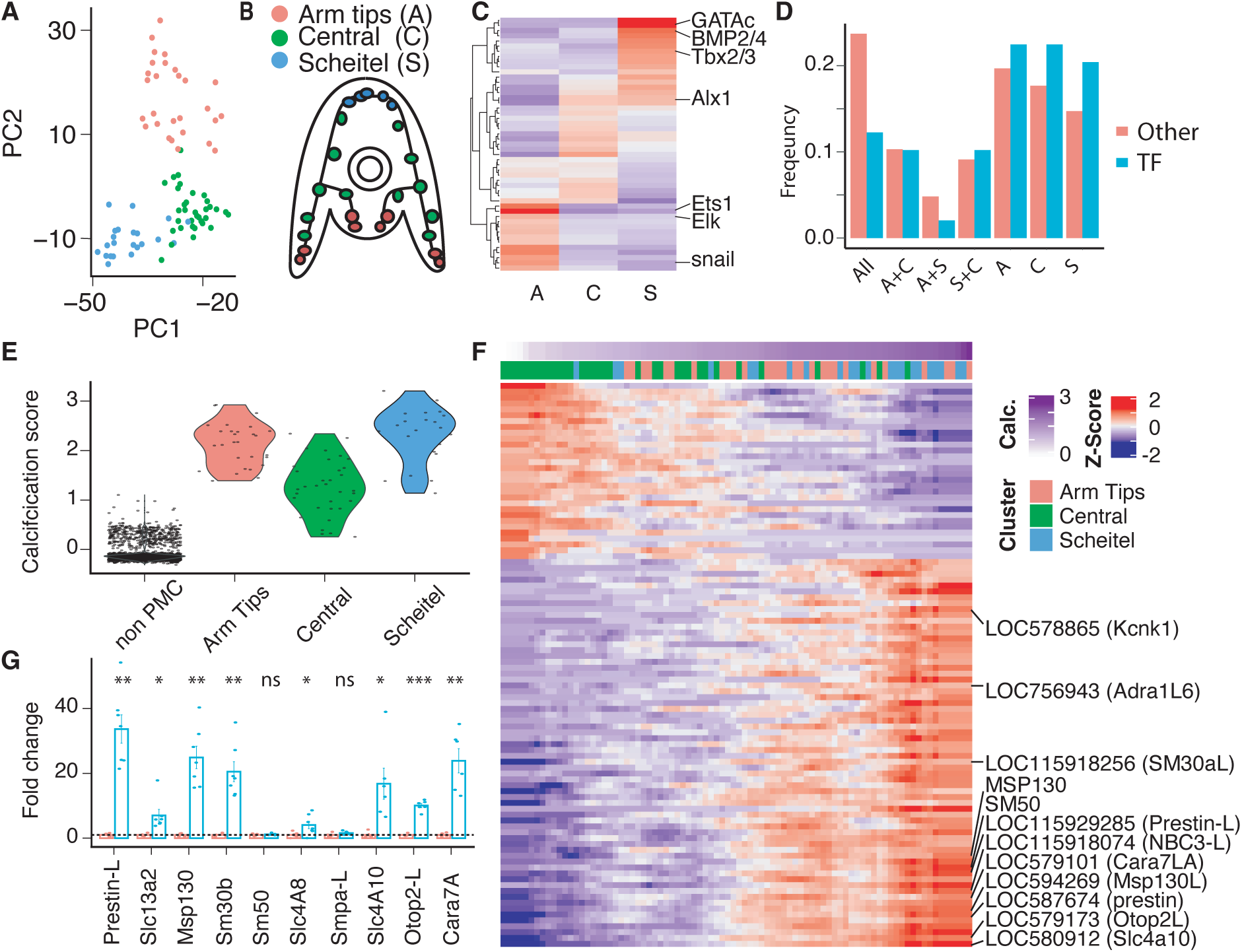
Diversity of gene regulation in skeletogenic cells. PCA plot of skeletogenic cells at Hpf72 (A) and cartoon indicating the localization of each subset of cells within the sea urchin embryo (B). (C)Heatmap of TFs expressed in different subpopulations of skeletogenic cells at Hpf72. (D) Stacked bar chart showing the expression domain of skeletogenic cell specific TFs and non-TF genes at pluteus stage (Hpf72). (E) Violin plot of calcification score calculated based on known calcification genes in sea urchins grouped by subtype of PMC. (F) Heatmap of genes significantly correlated with the calcification score, columns colored by subset of skeletogeneic cells at Hpf72. (G) Bar charts of relative expression of a subset of genes identified in this study in a recalcification assay, measured by qPCR (names based on human homology). A = Arm tips, C=Central rods, S= Scheitel rods. ns, *, **, *** indicate significance levels of p>0.05, p< 0.05, p< 0.01, p< 0.001 respectively, based on FDR-adjusted t-test.

Despite their extensive commonalities in function at the pluteus stage, the major clusters of skeletogenic cells identified in our study show a surprising diversity of subtype-specific regulatory states (12% shared by all subtypes, vs. > 60% cell-type-specific TFs at pluteus stage, Fig. 6C, D). Only six TFs were differentially expressed in all skeletogenic subsets when compared to non-skeletogenic cells, including *Alx1*, *Ets1*, and *Mitf*, which are involved in skeletogenic cell specification. A similar, albeit less pronounced pattern is observed for non-TF genes, and at gastrula stage (Fig. 6D, Fig. S8 C,D). This low transcriptional overlap of TFs and putative effector genes between skeletogenic subtypes highlights distinct functional gene regulation in subpopulations of skeletogenic cells (Valencia & Peter, 2024).

An important tasks of skeletogenic cells in the sea urchin is the deposition of calcite to form the larval skeleton (Mcintyre *et al*, 2014). In order to identify genes potentially involved in active calcification in the sea urchin, we calculated a ‘calcification score’ based on genes previously reported to be directly involved in the process of calcification (Otop2l, Slc4a10, CARA7, and sAC) (Hu *et al*, 2020; Chang *et al*, 2021) and identified genes significantly correlated with this score. In line with known patterns of calcification in the pluteus larva, the ‘calcification score’ is highest in Scheitel and Arm rod cells (Fig. 6E) (Guss & Ettensohn, 1997). We observed ∼100 genes to correlate with the calcification score (Fig. 6G), most of which are specifically expressed in skeletogenic cells (Fig. S8E). Some of the significantly correlated genes have previously been associated with calcification, (e.g. Msp130, SM50, SM30 (Guss & Ettensohn, 1997)), but most are uncharacterized.

To validate this set of genes, we measured relative expression of 10 genes that were positively correlated with the calcification score in a recalcification assay. Genes that are involved in calcification process were demonstrated to be upregulated in this assay (Hu *et al*, 2020). Out of these genes, 8 were significantly upregulated in this assay under actively recalcifying conditions, including prestin-like (LOC115929285), Slc13a2 (LOC588117) and NBC3-like (LOC115918074, with human homolog Slc4a8), which were previously not linked to calcification. Taken together, these results illustrate the functional heterogeneity at the level of transcriptional regulation of skeletogenic cells in developing sea urchins.

## 4. Discussion

### Patterns of accessibility over development

In this study, we used scRNA-seq, scATAC-seq, and scSLAM-seq to examine the regulatory landscape driving cell fate specification in the purple sea urchin from early cleavage to pluteus larval stages. We mapped a dynamic regulatory landscape that includes distal and intronic regulatory elements that are enriched for cell-type specific motifs and with accessibility profiles that often track changes in gene expression more closely than does promoter accessibility. This suggests that regulatory information is more complex and distributed in the sea urchin than suggested by promoter-based reporter assays. However, using co-accessibility, we did not find evidence for the highly distal regulatory interactions often seen in mammals that are mediated by chromatin loops and the actions of CTCF and the mediator complex, despite the sea urchin having a relatively large genome (as compared to, *e.g. Drosophila*). This suggests to us a regulatory genome that, while more complex than seen in cnidarians, does not feature the full suite of regulatory innovations seen in chordates. This view is supported by the real, but relatively modest, synteny blocks observed across sea urchin evolution (Marlétaz *et al*, 2023).

Temporally, we observed an increasingly complex regulatory landscapes that initiates with a major wave of zygotic genome activation (ZGA) at the 128-cell stage. The onset of accessibility at this time point appears well coordinated across cell types and coincides with an experimentally observed loss of developmental plasticity, suggesting that prior to this point, changes in inductive signals can induce cells to take on alternative fates. Not all is quiescent before this point, however: We found evidence for a minor wave of ZGA beginning at the 16-cell stage involving the activation of a number of genes with cell type specific function. Given the enrichment of genes in this set associated with the skeletogenic network (e.g. Alx), it is tempting to propose an early activation of the micromeres, possibly as a result of a changed cytoplasmic to nuclear ratio following asymmetric division at the 8-cell stage. However, we also observe the upregulation of genes with more ubiquitous later expression profiles (e.g. SpAN and HE) complicating an easy interpretation. More high-resolution datasets focusing on these early time points will be needed to resolve this hypothesis.

Several of the motifs enriched in pre-ZGA open chromatin correspond to TFs with known function in early sea urchin development (i.e. SoxB1 and SoxB2) and to TFs of importance in ZGA of other animals. For example, Sox19b is required for ZGA in zebrafish, while Zscan4 has been identified a specific TF in 2-cell mouse embryos (onset of ZGA) (Pálfy *et al*, 2020; Vastenhouw *et al*, 2019; Falco *et al*, 2007). These suggest potentially shared mechanisms of ZGA across related species, a hypothesis that can be tested by further experiments in sea urchins and other species.

Regulatory element usage between cell types of the same germ layer exhibits a large degree of overlap, even at later developmental states, reminiscent of pattens seen in *Drosophila* (##). This generality, however, may mask some interesting sea-urchin specific features. For example, the fraction of shared differentially accessible (DA) peaks between PMCs and the oral NSM is high until late blastula stages (>30%, not shown), while at the latest observed timepoints during skeletogenesis (pluteus stage) only a small overlap exists (<5%). Whether this overlap in DA elements is permissive for, or even supports, transfating of NSM into skeletogenic cells upon PMC removal is speculative at this point, but it is consistent with the observation that ets1-like motifs are dominant in both PMC- and oral NSM-specific regulatory elements, highlighting the potential role of Ets1 in NSM transfating into PMCs (Sharma & Ettensohn, 2011).

### Sea urchin neural development

Several neuronal subtypes have been described in the sea urchin, with distinct spatial origin (i.e. apical ectoderm, oral ectoderm, ciliary band and endoderm) (Garner *et al*, 2016; Wei *et al*, 2016; McClay *et al*, 2018). Our results support these distinct origins and identify a distinct pattern of TFs usage for ciliary band neurons, lateral ganglion neurons, and the apical organ differentiation. We identified a total of 7 neuronal transcriptomic states, including transitory ones. This diversity among a quite modest number of sampled neuronal cells (just 178 at the pluteus stage) suggests a much more extensive neuronal specification than has usually been reported for this species (Wood *et al*, 2018), a finding supported by other scRNA-seq projects that report similarly high levels of transcriptomic diversity from modestly sampled neuronal populations (Paganos *et al*, 2021).

Coverage in our scATAC data for neurons was modest, and lacked sufficient resolution to clearly distinguish neuronal subtypes. Nonetheless, we were able to identify clear patterns of TF motif usage that was dominated by TFs also involved in neurogenesis in human and flies (Table S11). This indicates a conserved neuronal regulatory program at least within the ectodermal neurons and highlights the utility of the sea urchin as a model for understanding neuronal evolution. Whether these features are also shared with endodermal neurons warrants further investigation.

#### Skeletogenic cell diversity in sea urchin

The larval skeleton of the sea urchin is deposited by specialized skeletogenic cells with diverse function and gene expression (Chang et al, 2021; Hu et al, 2020; Sun & Ettensohn, 2014; Guss & Ettensohn, 1997). We investigated the transcriptional regulation of these subsets and identified transcription factors driving their specification and, presumably function. To our surprise, few TFs were shared between the identified subpopulations, with the lineage defining TFs Alx1, Ets1, and Mitf being notable exceptions, indicating unexpected diversity among skeletogenic cells.

In addition to TFs, we identified ∼100 genes associated with calcification in the sea urchin. While many of these genes are shared between Arm and Scheitel rods at the pluteus stage, the sites of most active calcification (Guss & Ettensohn, 1997), both populations cluster distinctly and exhibit distinct transcriptional profiles at the pluteus stage. Some of these distinct transcriptional regulation might stem from spatial information; however, it is also noteworthy that these populations are involved in different modes and rates of calcification (Guss & Ettensohn, 1997). Further work will be needed to elucidate the specific role of the identified genes for calcification in developing sea urchins, a functionality that is impaired by acidifying oceans (Dupont et al, 2010).

### Final remarks

With the advent of more advanced cell culture and organoids, interest in model systems has waned in some quarters. This study is a reminder that while simplified with respect to genome duplication and with a radically re-arranged adult body plans, there is much we can recognize from mammalian gene regulation in early sea urchin development. While we saw no evidence for very long-distance regulation in this study, we did observe a complex regulatory landscape with dynamic distal and regulatory elements containing motif combinations that often show striking similarity to what is observed in mammals. But unlike mammalian development, early development in the sea urchin is striking accessible offering opportunities to use methods like metabolic labeling that would be essentially impossible in mammals and difficult even in systems like the zebrafish. As genome technologies continue to improve, and as costs go down, we think that systems like sea urchins will be increasingly relevant for understanding the evolution of our own genome as well as as crucial model systems for the billions and billions of ecologically and environmentally critical animals species whose genomes and life histories look more like sea urchin than they do mammals, nematodes, or fruit flies.

## Supporting information

Supplemental Tabel 1

---

**Figure S1.**
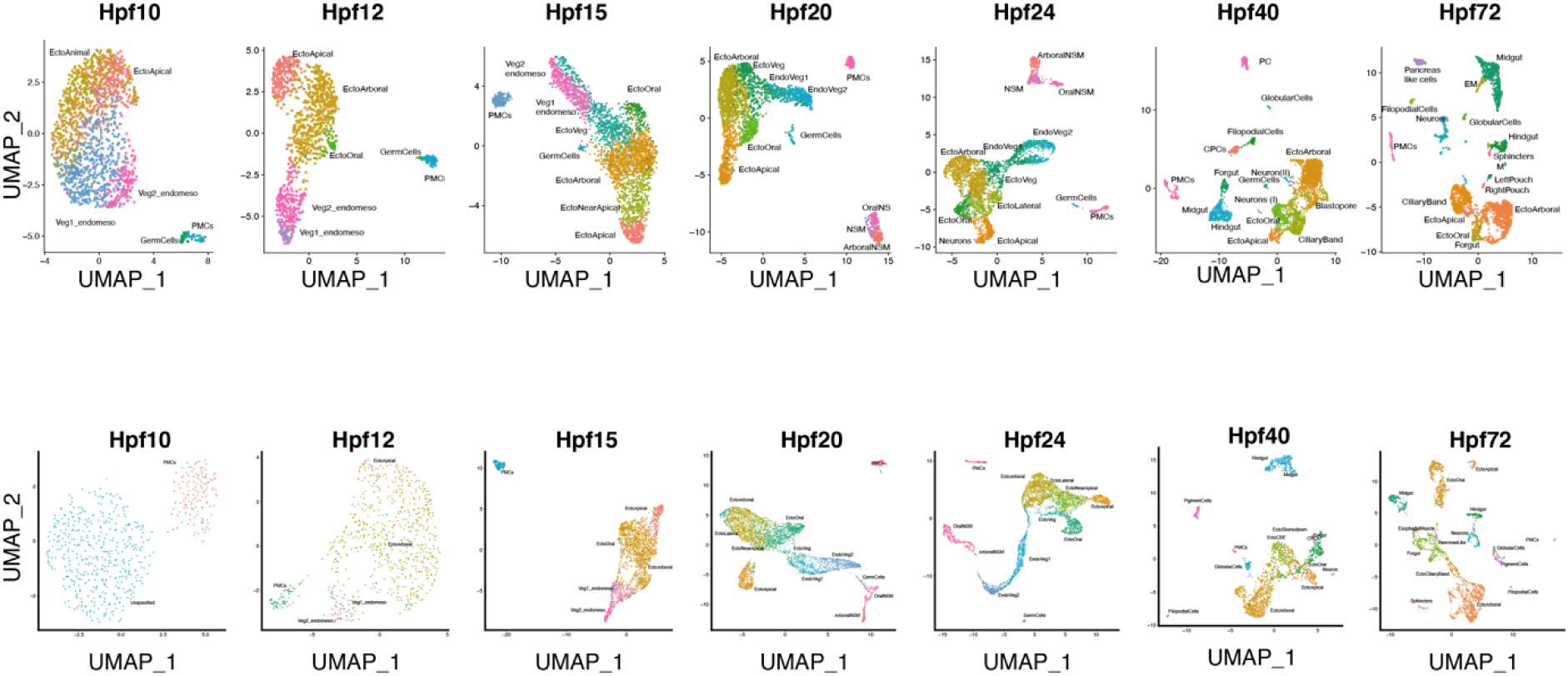
ScRNA-seq and scATAC-seq cell type assignment by developmental timepoint. Top panels: UMAP projections of scRNA-seq cells, mapped by developmental time point and colored by assigned cell type. Bottom panels: UMAP projections of scATAC-seq nuclei, mapped by developmental time point and colored by predicted cell type. To better resolve cell type complexity, nuclei were first coembedded with scRNA-seq data of appropriate stage, followed by dimensional reduction. There is no evidence of distinct clusters prior to 10hpf.

**Figure S2.**
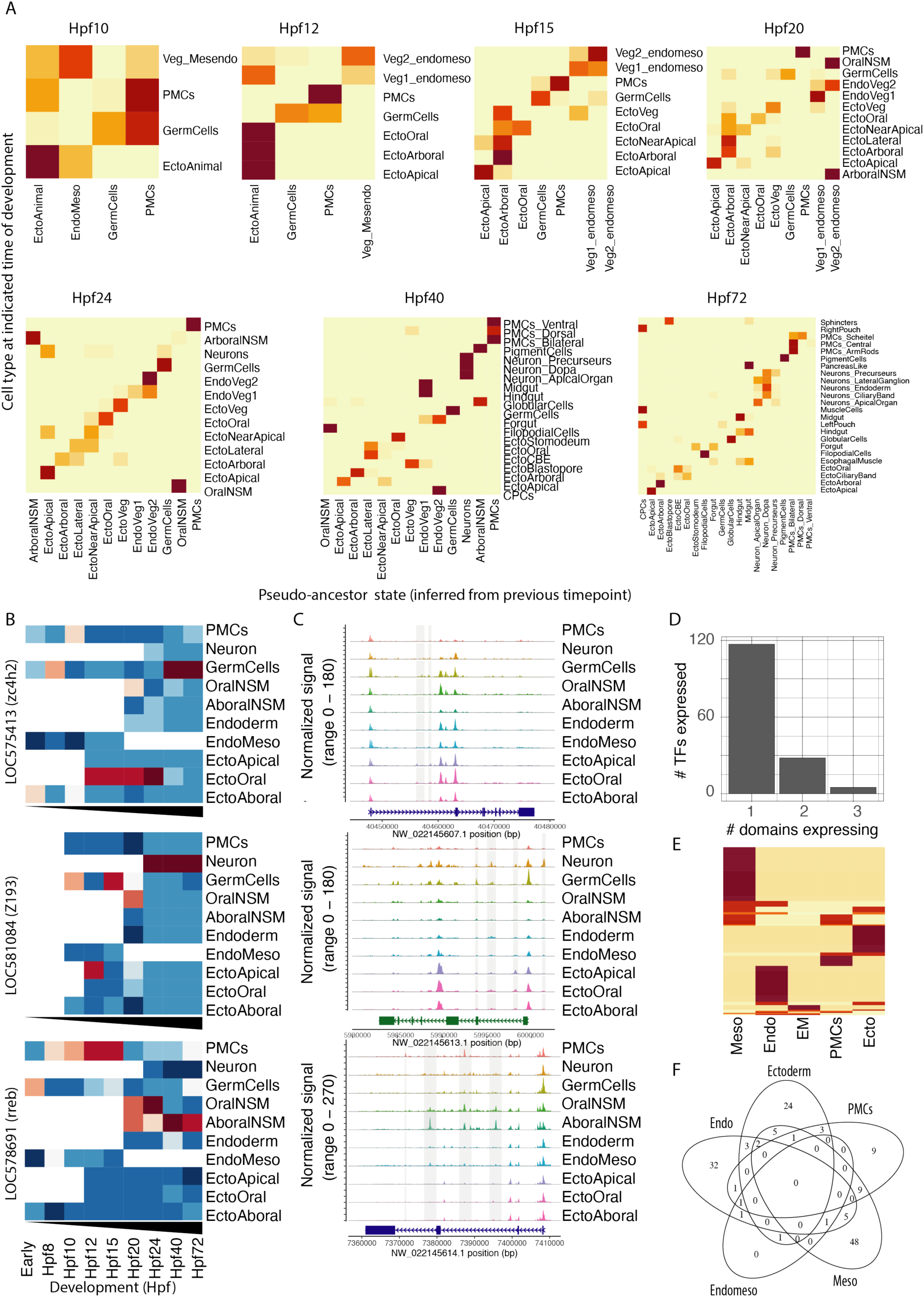
Ancestry voting identifies cellular trajectories across developmental timepoints in the sea urchin. (A) Heatmaps of edge weight for each cell type and each time point following Hpf8, based on ancestry voting. More likely ancestor states are marked in dark red, potentially ancestor states are labeled horizontally and descendant states vertically. (B) Expression of candidate lineage TFs Z193 (Neurons), zc4h2 (Oral ectoderm), and rreb1 (globular cells, Aboral NSM) in different cell lineages at different developmental times of the sea urchin. Darker red indicates stronger expression. (C) Coverage plot of the regulatory landscape around each of the candidate lineage TFs (Z193, zc4h2, rreb1), showing the genomic region covering the gene and extending 2kb on either side. Highlighted in gray are putative CREs with accessibility profile that is coherent with the expression pattern of the respective gene. (D) Frequency histogram showing the number of developmental lineages each candidate TFs is expressed. (E) Heatmap of candidate TF expression in different domains of the sea urchin across development. Expression has been binarized (expressed or not expressed) and scaled across domains, causing in more restricted expressed candidate TFs being shown in darker red. (F) Venn diagram of overlap between candidate lineage specifying TFs in different domains of the developing sea urchin embryo.

**Figure S3.**
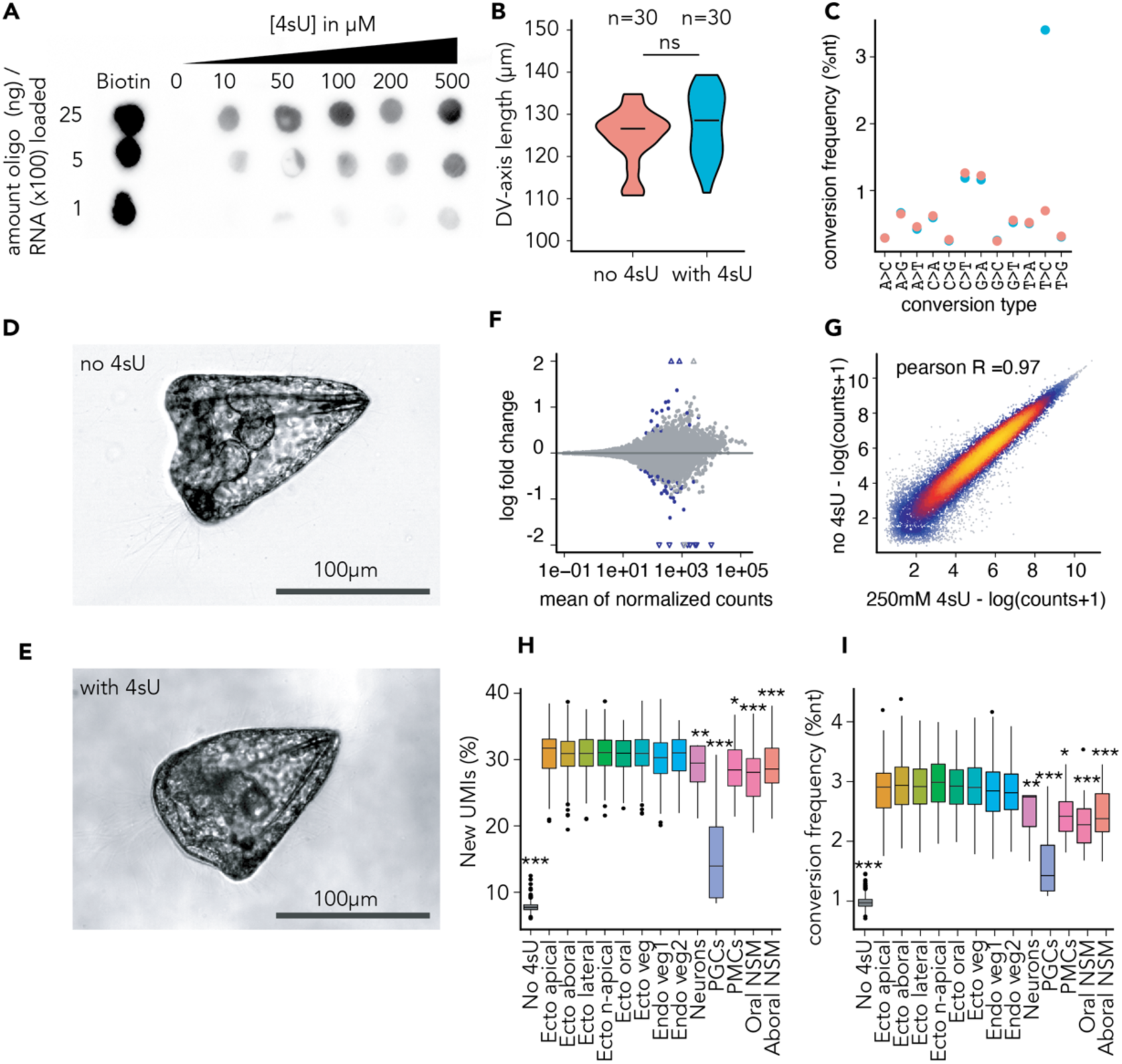
Method validation scSLAM-seq in the sea urchin. (A) 4sU incorporation in sea urchin embryos visualized by biotin labeling and dot blot. Positive control (biotinylated RNA) is shown on the left. (B) Violin plot of main axis lengths of Hpf72 sea urchin embryos incubated with or without 4sU. Significance assessed by student t-test. Measured embryos per condition are indicated on top. (C) Dot plot showing the frequency of different conversions in presence or absence of 4sU incorporation in scSLAM-seq data from the sea urchin at Hpf24. (D, E) Representative images of an early pluteus stage control embryo (C) and a pluteus stage embryo exposed to 4sU (D). Embryos are oriented with the dorsal side to the right. Scale bar in the lower right corner represents 100μm. Note that the orientation of the animal is slightly shifted. (F) MA plot showing differential gene expression between an unlabeled control and a 500μM labeled (6h) set of embryos, analysed by bulkRNA-seq. (G) Heat scatter plot of log transformed counts of Hpf24 samples labeled with 4sU and not labeled at 24hpf, analyzed by scSLAM-seq. Pearson correlation coefficient is indicated in the upper left corner. (H) Box-and-whiskers plot of the fraction of transcripts assigned as “new” by cell type in Hpf24 sea urchin embryos labeled with 4sU. (I) Box-and-whiskers plot of T-to-C conversion rate by cell type in Hpf24 sea urchin embryos labeled with 4sU. ns=non-significant. PMCs = primary mesenchyme cells. PGCs = primordial germ cells.

**Figure S4.**
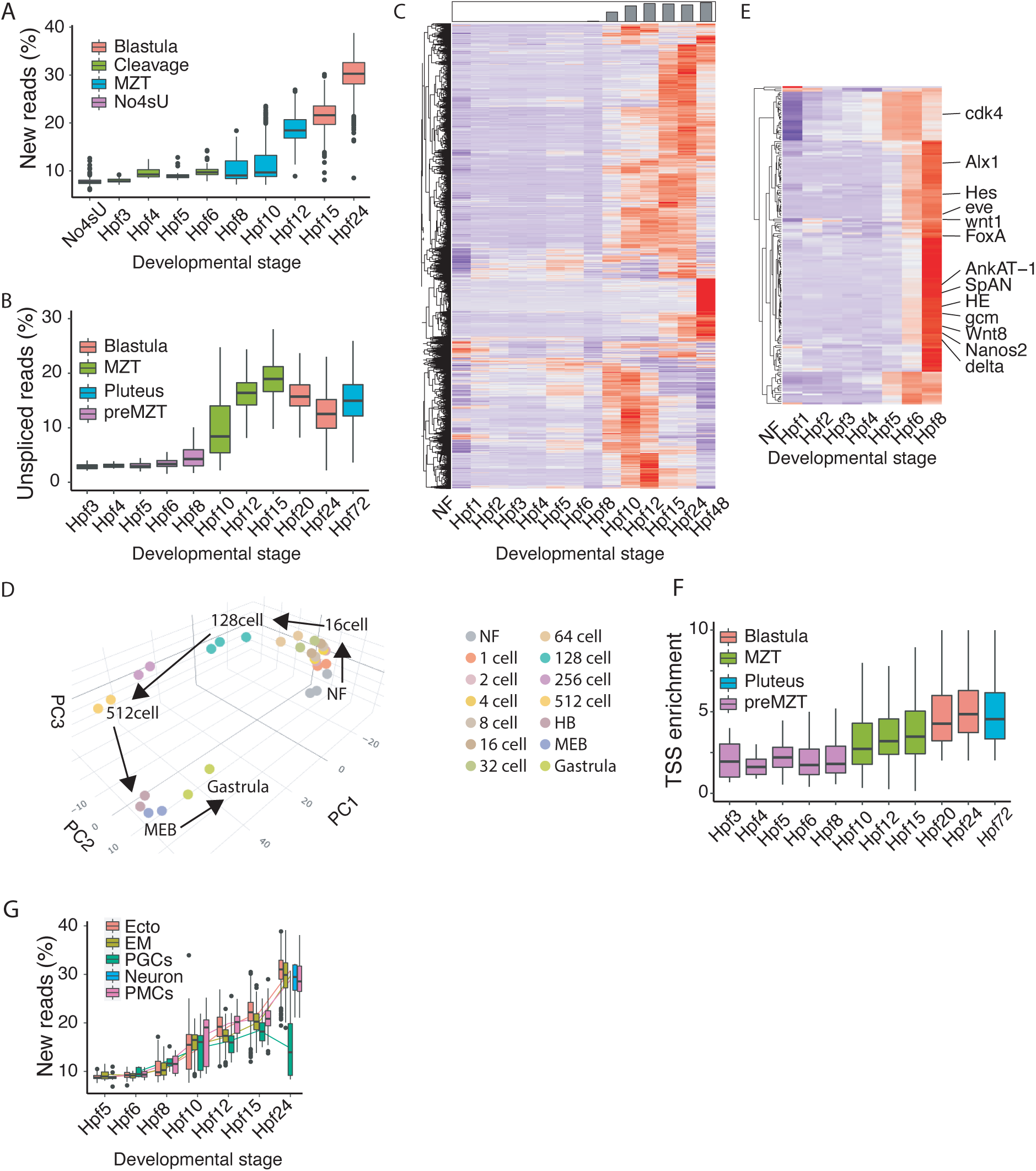
Zygotic genome activation in the sea urchin *S. purpuratus*. (A) 3D Principal component plot of cleavage resolution bulk RNA-seq samples. Developmental stages are indicated by colors. Arrows indicate “breaks” in directionality of temporal progression. (B) Heatmap of all newly upregulated genes in sea urchin development by developmental time. Expression levels are scaled and centered. Top panel is a bar chart representing the fraction of newly upregulated genes at the indicated developmental time point (out of all genes upregulated during development up to the late gastrula stage). (C) Heatmap of early upregulated genes (prior to ZGA at Hpf10) by developmental time point. For legibility, only the names of a few well-known early genes are indicated on the right. (D) Box-and-whiskers plot showing the fraction of unspliced UMIs per cell for each developmental stage based on scRNA-seq. (E) Box-and-whiskers plot showing the fraction of new UMIs (determined by 4sU incorporation) per cell for each developmental stage based on scSLAM-seq. (F) Box plot showing the TSS enrichment from scATAC-seq data at each time point. Note the systematic increase after 10hpf/128 cells. (G) Box-and-whiskers plot showing the fraction of new UMIs (determined by 4sU incorporation) per cell for each developmental stage based on scSLAM-seq, and split by cell-type. Lines connect average fraction of new UMIs by cell type and stage. Ecto=Ectoderm, EM=Endomesoderm, PGCs=germ cells, PMCs=primary mesenchyme cells.

**Figure S5.**
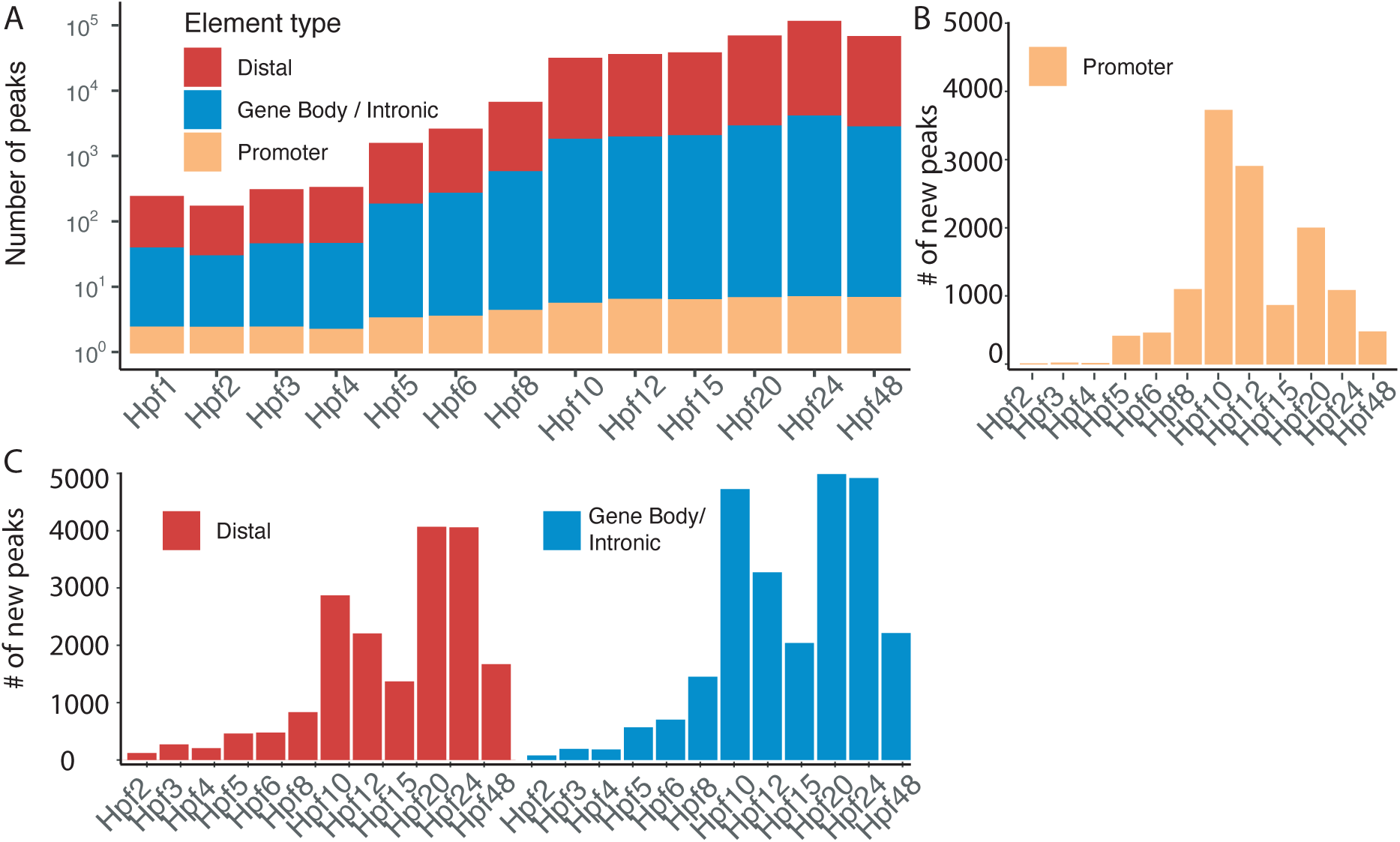
Opening of regulatory landscape in sea urchin development. (A) Frequency histogram of the number of regulatory elements detected at the bulk level for each developmental time point in the bulkATAC dataset, colored by peak annotation respective to closest gene. Note that the scale of the y axis is log-transformed. (B,C) Frequency histogram of newly accessible promoter (B), or gene body and distal (C) regulatory elements per developmental stage.

**Figure S6.**
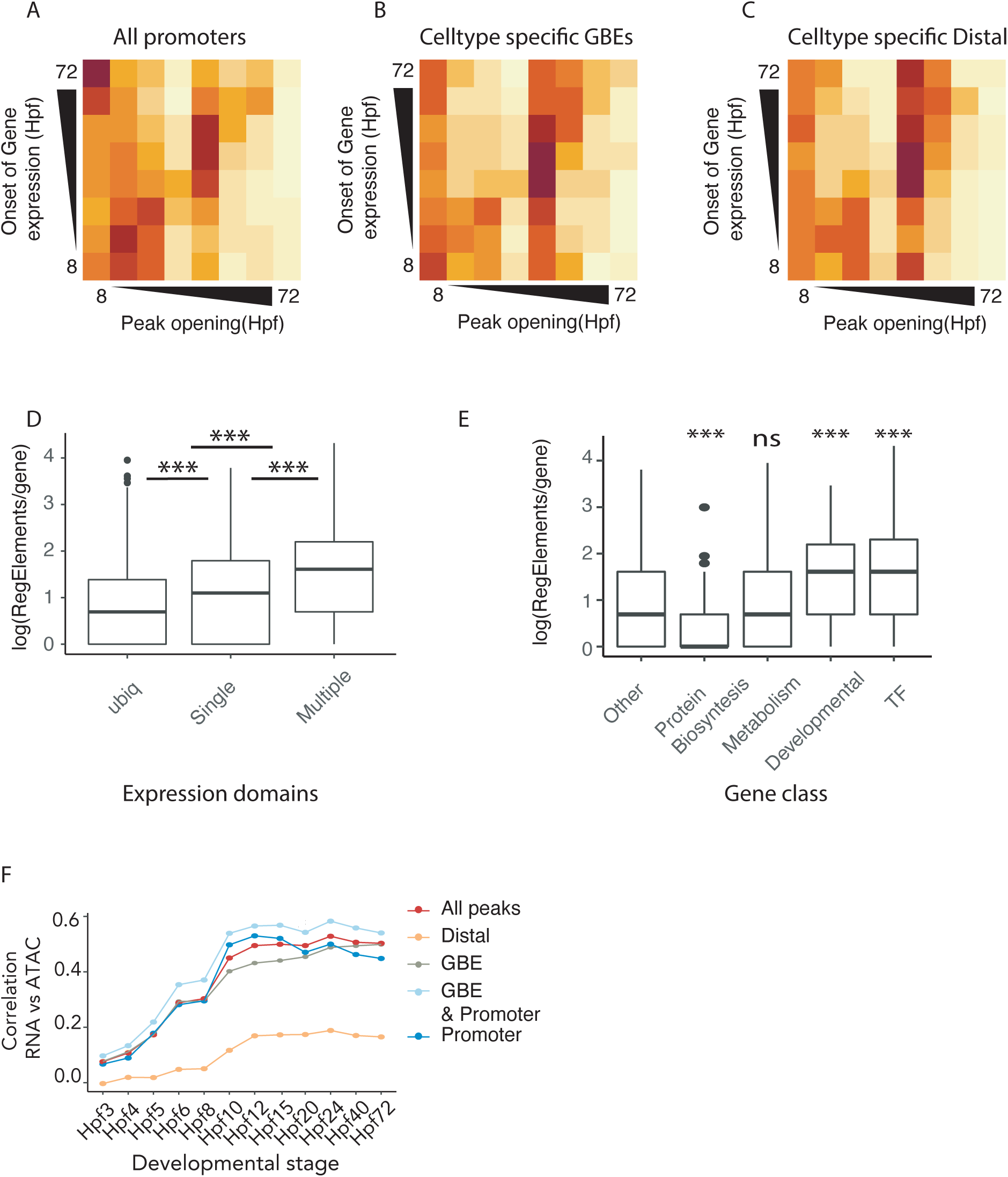
(A-C) Heatmap of temporal difference between promoter opening and onset of gene expression (by unspliced UMI count) for lineage specific genes in sea urchin development, plotted for all promoters(A), all cell-type-specific GBEs (B) and all cell-type-specific distal elements (C). (D,E) Box and whiskers plot of log-transformed number of accessible regulatory regions associated with expression domain usage (D) or different gene classes (E). (F) The correlation between RNA expression level and open chromatin for different categories of regulatory features. Peaks were assigned to gene based on shortest distance.

**Figure S7.**
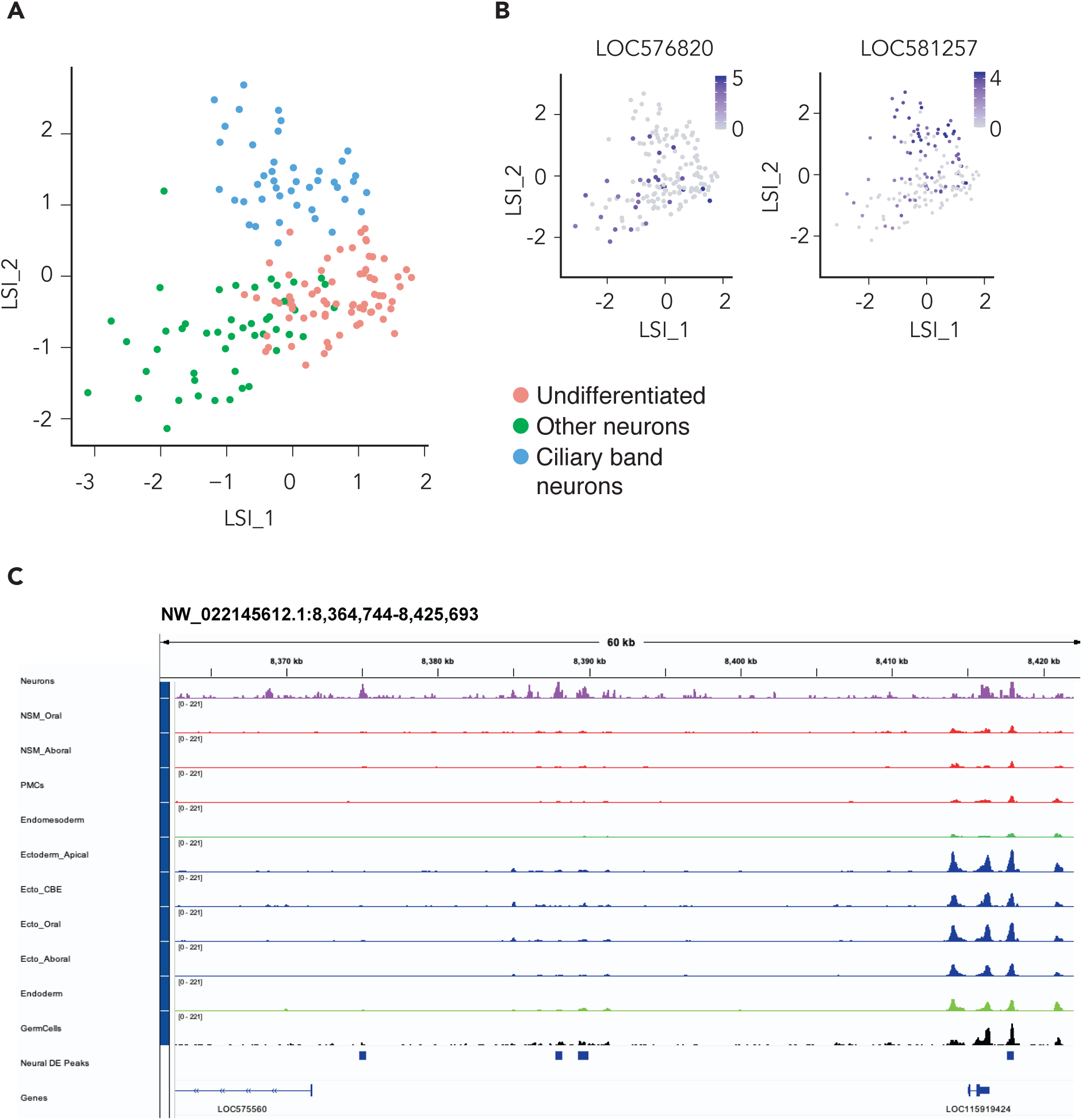
Regulatory landscape of neurodevelopment in sea urchins. (A) scATAC-seq LSI sub-clustering of neurons colored by neuronal subset. (B) LSI clustering from (A) colored by the gene activity score of two marker genes for ciliary band neurons and other neurons. (C) Representative open chromatin tracks around two Elav paralogues highlighting neuron-specific accessibility.

**Figure S8.**
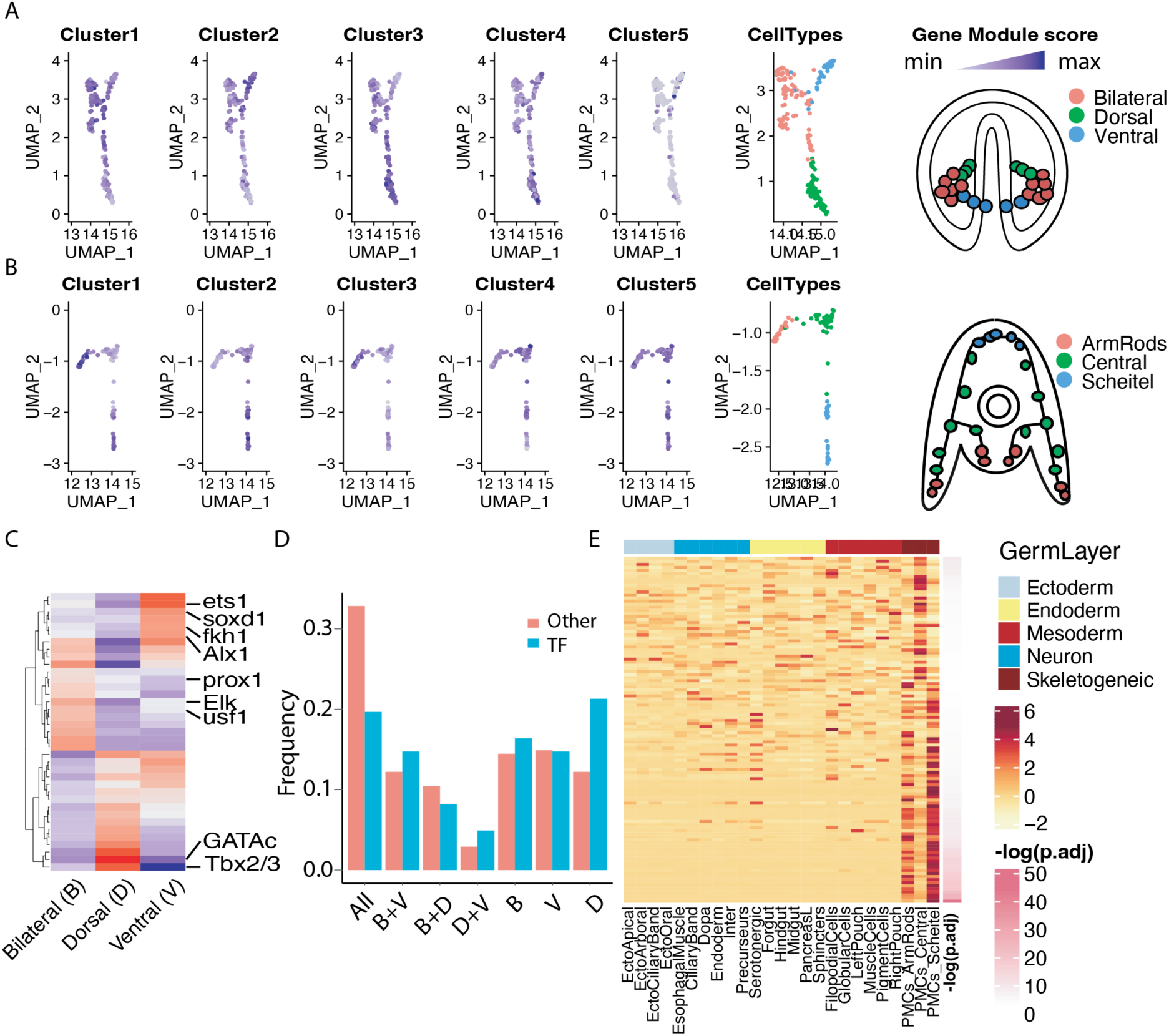
Diversity of skeletogeneic cells of the sea urchin development. (A) UMAP clustering of skeletogenic cells at Hpf40, colored by expression of gene clusters from Sun & Ettensohn (2014) or by skeletogeneic cell sub-type and a cartoon indicating the localization of each subset of cells within the sea urchin embryo. (B) UMAP clustering of skeletogenic cells at Hpf40, colored by expression of gene clusters from Sun & Ettensohn (2014) or by skeletogeneic cell sub-type and a cartoon indicating the localization of each subset of cells within the sea urchin embryo. (C) Heatmap of skeletogenic-cell specific TFs expressed in different subpopulations of skeletogenic cells at Hpf40. (D) Stacked bar chart showing the expression domain of skeletogenic cell specific TFs and non-TF genes at the gastrula stage (Hpf40). (E) Heatmap of genes significantly correlated with calcification score, ordered by correlation with calcification score. Columns colored by germ layer, rows colored by negative log adjusted p-value. Note the skeletogenic subset on the far right.

## References

Alpern D, Gardeux V, Russeil J, Mangeat B, Meireles-Filho ACA, Breysse R, Hacker D & Deplancke B (2019) BRB-seq: Ultra-affordable high-throughput transcriptomics enabled by bulk RNA barcoding and sequencing. Genome Biology 20: 1–15

Ashby MH (2006) The Sea Urchin Regulome in Development.

Barsi JC & Davidson EH (2016) Cis-Regulatory control of the initial neurogenic pattern of onecut gene expression in the sea urchin embryo. Developmental Biology 409: 310– 318

Beddington RSP & Robertson EJ (1999) Axis development and early asymmetry in mammals. Cell 96: 195–209

Bergen V, Lange M, Peidli S, Wolf FA & Theis FJ (2020) Generalizing RNA velocity to transient cell states through dynamical modeling. Nature Biotechnology 38: 1408– 1414

Bonn S, Zinzen RP, Perez-Gonzalez A, Riddell A, Gavin AC & Furlong EEM (2012) Cell type-specific chromatin immunoprecipitation from multicellular complex samples using bits-chip. Nature Protocols 7: 978–994

Chang WW, Matt AS, Schewe M, Musinszki M, Grüssel S, Brandenburg J, Garfield D, Bleich M, Baukrowitz T & Hu MY (2021) An otopetrin family proton channel promotes cellular acid efflux critical for biomineralization in a marine calcifier. Proceedings of the National Academy of Sciences of the United States of America 118: 1–8

Corces MR, Trevino AE, Hamilton EG, Greenside PG, Sinnott-Armstrong NA, Vesuna S, Satpathy AT, Rubin AJ, Montine KS, Wu B, et al. (2017) An improved ATAC-seq protocol reduces background and enables interrogation of frozen tissues. Nature Methods 14: 959–962

Cornejo-Páramo P, Roper K, Degnan SM, Degnan BM & Wong ES (2022) Distal regulation, silencers, and a shared combinatorial syntax are hallmarks of animal embryogenesis. Genome Res 32: 474–487

Cusanovich DA, Reddington JP, Garfield DA, Daza RM, Aghamirzaie D, Marco-Ferreres R, Pliner HA, Christiansen L, Qiu X, Steemers FJ, et al. (2018) The cis-regulatory dynamics of embryonic development at single-cell resolution. Nature 555: 538–542

Davidson EH (2006) cis-Regulatory Modules , and the Structure / Function Basis of Regulatory Logic. In The regulatory genome: gene regulatory networks in development and evolution. pp 31–86.

Dobin A, Davis CA, Schlesinger F, Drenkow J, Zaleski C, Jha S, Batut P, Chaisson M & Gingeras TR (2013) STAR: ultrafast universal RNA-seq aligner. Bioinformatics 29: 15–21

Ettensohn CA, Illies MR, Oliveri P & De Jong DL (2003) Alx1, a member of the Cart1/Alx3/Alx4 subfamily of paired-class homeodomain proteins, is an essential component of the gene network controlling skeletogenic fate specification in the sea urchin embryo. Development 130: 2917–2928

Ettensohn CA, Kitazawa C, Cheers MS, Leonard JD & Sharma T (2007) Gene regulatory networks and developmental plasticity in the early sea urchin embryo: Alternative deployment of the skeletogenic gene regulatory network. Development 134: 3077– 3087

Floc’hlay S, Wong ES, Zhao B, Viales RR, Thomas-Chollier M, Thieffry D, Garfield DA & Furlong EEM (2021) Cis-acting variation is common across regulatory layers but is often buffered during embryonic development. Genome Research 31: 211–224

Flytzanis CN, Brandhorst BP, Britten RJ & Davidson EH (1982) Developmental patterns of cytoplasmic transcript prevalence in sea urchin embryos. Dev Biol 91: 27–35

Foley S, Ku C, Arshinoff B, Lotay V, Karimi K, Vize PD & Hinman V (2021) Integration of 1:1 orthology maps and updated datasets into Echinobase. Database 2021: 1–7

Garfield D, Haygood R, Nielsen WJ & Wray GA (2012) Population genetics of cis-regulatory sequences that operate during embryonic development in the sea urchin Strongylocentrotus purpuratus. Evol Dev 14: 152–167

Garner S, Zysk I, Byrne G, Kramer M, Moller D, Taylor V & Burke RD (2016) Neurogenesis in sea urchin embryos and the diversity of deuterostome neurogenic mechanisms. Development (Cambridge) 143: 286–297

Goldstein B & Freeman G (1997) Axis specification in animal development. BioEssays 19: 105–116

Granja JM, Corces MR, Pierce SE, Bagdatli ST, Choudhry H, Chang HY & Greenleaf WJ (2021) ArchR is a scalable software package for integrative single-cell chromatin accessibility analysis. Nature Genetics 53: 935

Guss KA & Ettensohn CA (1997) Skeletal morphogenesis in the sea urchin embryo: Regulation of primary mesenchyme gene expression and skeletal rod growth by ectoderm-derived cues. Development 124: 1899–1908

Haghverdi L, Buettner F & Theis FJ (2015) Diffusion maps for high-dimensional single-cell analysis of differentiation data. Bioinformatics 31: 2989–2998

Hao Y, Hao S, Andersen-Nissen E, Mauck WM, Zheng S, Butler A, Lee MJ, Wilk AJ, Darby C, Zager M, et al. (2021) Integrated analysis of multimodal single-cell data. Cell 184: 3573–3587.e29

Heinz S, Benner C, Spann N, Bertolino E, Lin YC, Laslo P, Cheng JX, Murre C, Singh H & Glass CK (2010) Simple Combinations of Lineage-Determining Transcription Factors Prime cis-Regulatory Elements Required for Macrophage and B Cell Identities. Molecular Cell 38: 576–589

Holler K, Neuschulz A, Drewe-Boß P, Mintcheva J, Spanjaard B, Arsiè R, Ohler U, Landthaler M & Junker JP (2021) Spatio-temporal mRNA tracking in the early zebrafish embryo. Nature Communications 12

Hörstadius S (1936) Über die zeitliche Determination im Keim von Paracentrotus Lividus LK. Wilhelm Roux’ Archiv für Entwicklungsmechanik der Organismen 135: 1–39

Hu MY, Petersen I, Chang WW, Blurton C & Stumpp M (2020) Cellular bicarbonate accumulation and vesicular proton transport promote calcification in the sea urchin larva. Proceedings of the royal society B 287: 20201506

Huang X & Huang Y (2021) Cellsnp-lite: an efficient tool for genotyping single cells. Bioinformatics: 4–7

Huang Y, McCarthy DJ & Stegle O (2019a) Vireo: Bayesian demultiplexing of pooled single-cell RNA-seq data without genotype reference. Genome Biology 20: 1–12

Huang Y, McCarthy DJ & Stegle O (2019b) Vireo: Bayesian demultiplexing of pooled single-cell RNA-seq data without genotype reference. Genome Biology 20: 273

Illies MR, Peeler MT, Dechtiaruk AM & Ettensohn CA (2002) Identification and developmental expression of new biomineralization proteins in the sea urchin Strongylocentrotus purpuratus. Development Genes and Evolution 212: 419–431

Jiang H, Lei R, Ding S-W & Zhu S (2014) Skewer: a fast and accurate adapter trimmer for next-generation sequencing paired-end reads. BMC Bioinformatics 15: 182

Korsunsky I, Millard N, Fan J, Slowikowski K, Zhang F, Wei K, Baglaenko Y, Brenner M, Loh P ru & Raychaudhuri S (2019) Fast, sensitive and accurate integration of single-cell data with Harmony. Nature Methods 16: 1289–1296

Li E, Cui M, Peter IS & Davidson EH (2014) Encoding regulatory state boundaries in the pregastrular oral ectoderm of the sea urchin embryo. Proceedings of the National Academy of Sciences of the United States of America 111

Li H (2013) Aligning sequence reads, clone sequences and assembly contigs with BWA-MEM. 00: 1–3

Li H, Handsaker B, Wysoker A, Fennell T, Ruan J, Homer N, Marth G, Abecasis G, Durbin R, & 1000 Genome Project Data Processing Subgroup (2009) The Sequence Alignment/Map format and SAMtools. Bioinformatics 25: 2078–2079

Love MI, Huber W & Anders S (2014) Moderated estimation of fold change and dispersion for RNA-seq data with DESeq2. Genome Biology 15: 550

Lyons DC, Martik ML, Saunders LR & McClay DR (2014) Specification to biomineralization: following a single cell type as it constructs a skeleton. Integrative and comparative biology 54: 723–733

Marlétaz F, Couloux A, Poulain J, Labadie K, Da Silva C, Mangenot S, Noel B, Poustka AJ, Dru P, Pegueroles C, et al. (2023a) Analysis of the P. lividus sea urchin genome highlights contrasting trends of genomic and regulatory evolution in deuterostomes. Cell Genomics 3: 100295

Marlétaz F, Couloux A, Poulain J, Labadie K, Da Silva C, Mangenot S, Noel B, Poustka AJ, Dru P, Pegueroles C, et al. (2023b) Analysis of the P. lividus sea urchin genome highlights contrasting trends of genomic and regulatory evolution in deuterostomes. Cell Genomics 3: 100295

Martin M (2011) Cutadapt removes adapter sequences from high-throughput sequencing reads. EMBnet 17: 10–12

McClay DR (2004) Methods for embryo dissociation and analysis of cell adhesion. In Methods in Cell Biology pp 311–329.

McClay DR (2011) Evolutionary crossroads in developmental biology: Sea urchins. Development 138: 2639–2648

McClay DR & Logan CY (1996) Regulative capacity of the archenteron during gastrulation in the sea urchin. Development 122: 607–616

McClay DR, Miranda E & Feinberg SL (2018) Neurogenesis in the sea urchin embryo is initiated uniquely in three domains. Development (Cambridge) 145

McGarvey AC, Kopp W, Vučićević D, Mattonet K, Kempfer R, Hirsekorn A, Bilić I, Gil M, Trinks A, Merks AM, et al. (2022) Single-cell-resolved dynamics of chromatin architecture delineate cell and regulatory states in zebrafish embryos. Cell Genomics 2: 100083

McGinnis CS, Murrow LM & Gartner ZJ (2019) DoubletFinder: Doublet Detection in Single-Cell RNA Sequencing Data Using Artificial Nearest Neighbors. Cell Systems 8: 329–337.e4

Mcintyre DC, Lyons DC, Martik M & Mcclay DR (2014) Branching out: Origins of the sea urchin larval skeleton in development and evolution. Genesis 52: 173–185 doi:10.1002/dvg.22756 [PREPRINT]

Noonan JP & McCallion AS (2010) Genomics of Long-Range Regulatory Elements. Annu Rev Genom Hum Genet 11: 1–23

Oliveri P, Davidson EH & McClay DR (2003) Activation of pmar1 controls specification of micromeres in the sea urchin embryo. Developmental Biology 258: 32–43

Oudelaar AM & Higgs DR (2021) The relationship between genome structure and function. Nat Rev Genet 22: 154–168

Oulhen N & Wessel G (2017) A quiet space during rush hour: Quiescence in primordial germ cells. Stem Cell Research 25: 296–299

Pálfy M, Schulze G, Valen E & Vastenhouw NL (2020) Chromatin accessibility established by Pou5f3, Sox19b and Nanog primes genes for activity during zebrafish genome activation. PLoS Genetics 16: 1–25

Picelli S, Björklund ÅK, Reinius B, Sagasser S, Winberg G & Sandberg R (2014) Tn5 transposase and tagmentation procedures for massively scaled sequencing projects. Genome Research 24: 2033–2040

Qiu C, Cao J, Martin BK, Li T, Welsh IC, Srivatsan S, Huang X, Calderon D, Noble WS, Disteche CM, et al. (2022) Systematic reconstruction of cellular trajectories across mouse embryogenesis. Nature Genetics 54: 328–341

Ransick A & Davidson EH (2006) cis-regulatory processing of Notch signaling input to the sea urchin glial cells missing gene during mesoderm specification. Developmental Biology 297: 587–602

Ransick A & Davidson EH (2012) Cis-regulatory logic driving glial cells missing: Self-sustaining circuitry in later embryogenesis. Developmental Biology 364: 259–267

Reddington JP, Garfield DA, Sigalova OM, Karabacak Calviello A, Marco-Ferreres R, Girardot C, Viales RR, Degner JF, Ohler U & Furlong EEM (2020) Lineage-Resolved Enhancer and Promoter Usage during a Time Course of Embryogenesis. Developmental Cell 55: 648–664.e9

Schulz KN & Harrison MM (2019) Mechanisms regulating zygotic genome activation. Nature Reviews Genetics 20: 221–234

Shashikant T, Khor JM & Ettensohn CA (2018) Global analysis of primary mesenchyme cell cis-regulatory modules by chromatin accessibility profiling. BMC Genomics 19: 1–18

Slater DW, Slater I & Gillespie D (1972) Post-fertilization Synthesis of Polyadenylic Acid in Sea Urchin Embryos. Nature 240: 333–337

Spitz F & Furlong EEM (2012) Transcription factors: From enhancer binding to developmental control. Nature Reviews Genetics 13: 613–626

Stuart T, Butler A, Hoffman P, Hafemeister C, Papalexi E, Mauck WM, Hao Y, Stoeckius M, Smibert P & Satija R (2019) Comprehensive Integration of Single-Cell Data. Cell 177: 1888–1902.e21

Stuart T, Srivastava A, Madad S, Lareau CA & Satija R (2021) Single-cell chromatin state analysis with Signac. Nature Methods 18: 1333–1341

Sun Z & Ettensohn CA (2014) Signal-dependent regulation of the sea urchin skeletogenic gene regulatory network. Gene Expression Patterns 16: 93–103

Uhlitz F, Bischoff P, Peidli S, Sieber A, Trinks A, Lüthen M, Obermayer B, Blanc E, Ruchiy Y, Sell T, et al. (2021) Mitogen-activated protein kinase activity drives cell trajectories in colorectal cancer. EMBO Molecular Medicine 13

Valencia JE & Peter IS (2024) Combinatorial regulatory states define cell fate diversity during embryogenesis. Nat Commun 15: 6841

Vastenhouw NL, Cao WX & Lipshitz HD (2019) The maternal-to-zygotic transition revisited. Development 146

Wei Z, Angerer LM & Angerer RC (2016) Neurogenic gene regulatory pathways in the sea urchin embryo. Development (Cambridge) 143: 298–305

Weitzel HE, Illies MR, Byrum CA, Xu R, Wikramanayake AH & Ettensohn CA (2004) Differential stability of β-catenin along the animal-vegetal axis of the sea urchin embryo mediated by dishevelled. Development 131: 2947–2956

Wilt FH (1970) The synthesis of ribonucleic acid in sea urchin embryos. Developmental Biology 23: 444–455

Yajima M & Wessel GM (2011) Small micromeres contribute to the germline in the sea urchin. Development 138: 237–243

Yu G, Wang LG & He QY (2015) ChIP seeker: An R/Bioconductor package for ChIP peak annotation, comparison and visualization. Bioinformatics 31: 2382–2383

Yuh CH, Dorman ER, Howard ML & Davidson EH (2004) An otx cis-regulatory module: A key node in the sea urchin endomesoderm gene regulatory network. Developmental Biology 269: 536–551

Yuh CH, Moore JG & Davidson EH (1996) Quantitative functional interrelations within the cis-regulatory system of the S. purpuratus Endo16 gene. Development 122: 4045– 4056

Zhang Y, Liu T, Meyer CA, Eeckhoute J, Johnson DS, Bernstein BE, Nussbaum C, Myers RM, Brown M, Li W, et al. (2008) Model-based analysis of ChIP-Seq (MACS). Genome Biology 9

