## Supplemental Tables for "The Establishment of Cell-Type Specific Gene Regulation in the Sea Urchin Embryo"

**Supplementary Table 1**. Number of cells and quality metrics for each cross in scRNA-seq.

| Cross | Library | Total cells | UMI per cell | Genes per cell | Mitochondrial UMIs |
|---|---|---|---|---|---|
| Hpf3_F4 | scRNA_EarlyDev_v3_1 | 30 | 1319.5 | 813 | 19.5 |
| Hpf4_F1 | | 7 | 1276 | 790 | 17.4 |
| Hpf5_F3 | | 21 | 1442 | 812 | 20.3 |
| Hpf5_F5 | | 24 | 1276 | 755.5 | 20.3 |
| Hpf6_F8 | | 64 | 1693.5 | 1012 | 16.2 |
| Hpf6_F9 | | 66 | 1736.5 | 1010.5 | 18.2 |
| Hpf8_F6 | | 66 | 5016.5 | 2115.5 | 19.3 |
| Hpf8_F7 | | 89 | 1395 | 832 | 20.2 |
| Hpf10_F5 | | 350 | 4750.5 | 2109 | 15.4 |
| Hpf10_F7 | | 189 | 5261 | 2343 | 15.6 |
| Hpf12_F1 | | 593 | 3695 | 1645 | 17.4 |
| Hpf12_F3 | | 606 | 3985 | 1673.5 | 20.2 |
| Hpf15_F4 | | 3043 | 3183 | 1447 | 18.8 |
| Hpf8_1 | scRNA_4sU_v3_1 | 131 | 1968 | 1102 | 16.9 |
| Hpf10_1 | | 307 | 2848 | 1435 | 17.6 |
| Hpf24_1 | | 2248 | 3010.5 | 1212 | 22.2 |
| Hpf24_2 | | 2723 | 3630 | 1412 | 17.4 |
| Hpf20 | scRNA_Late_v3_1 | 2825 | 3026 | 1383 | 18.0 |
| Hpf40 | | 4785 | 2270 | 1162 | 14.4 |
| Hpf72 | | 2761 | 1999 | 995 | 17.1 |
| | **Total** | **20928** | **2889** | **1301** | **17.5** |

**Supplementary Table 2**. Marker genes used for cell type assignment in scRNA-seq dataset.

| Gene Names | Loc number | Cell type |
|---|---|---|
| Brn124<br>FoxP<br>Isl1<br>FoxA | LOC577601<br>LOC577677<br>LOC576365<br>LOC578584 | Forgut |
| Six3<br>myosin regulatory light chain 12A<br>myosin light chain 3, skeletal muscle isoform<br>myosin light chain kinase, smooth muscle | LOC576281<br>LOC578403<br>LOC578241<br>LOC576786 | Muscle |
| blimp1/krox<br>GATAe<br>Endo16<br>macrophage mannose receptor 1 | LOC751833<br>LOC408052<br>LOC373279<br>LOC757247 | Midgut |
| Ptf1a<br>Cpa2L<br>Pnlp | LOC115926162<br>LOC115918269<br>LOC594532 | Pancreas-like cells |
| cdx<br>FoxD<br>xlox<br>Hox11/13<br>Bra | LOC584191<br>LOC751845<br>LOC373527<br>LOC373462<br>LOC576774 | Hindgut |
| SM30A<br>Alx1<br>SM50 | LOC115917938<br>LOC373521<br>LOC373464 | PMCs |
| P16 negative | LOC373523 | PMCs_Central |

| | | |
|---|---|---|
| FGF | LOC582058 | PMCs_ArmRods |
| BMP2/4<br>FGF negative | LOC373196<br>LOC582058 | PMCs_Scheitel |
| SM30A low | LOC115917938 | PMCs_Ventral |
| FGF+ | LOC582058 | PMCs_bilateral |
| BMP2/4<br>GATAc<br>Tbx2/3 | LOC373196<br>LOC373316<br>LOC 575938 | PMCs_Dorsal |
| 14-3-3<br>GST tetha1<br>FoxG*<br>CHRD<br>Gsc | LOC575018<br>LOC115918827<br>LOC583688<br>LOC580173<br>LOC373244 | Oral Ectoderm |
| CyIIIa<br>SPEC1<br>EGF2 | LOC373298<br>LOC373443<br>LOC373304 | Aboral Ectoderm |
| hnf6<br>FoxG*<br>Bicc1 | LOC378468<br>LOC583688<br>LOC100888728 | Ciliary Band Ectoderm |
| AnkAT-1<br>NK2.1<br>Frizz5/8<br>Six3<br>FoxQ2 | LOC590011<br>LOC 373506<br>LOC582578<br>LOC576281<br>LOC100891362 | Apical Ectoderm |
| Bra<br>FoxA | LOC576774<br>LOC578584 | Stomodeal Ectoderm |
| FGF | LOC582058 | Lateral Ectoderm |
| Bra<br>FoxA | LOC576774<br>LOC578584 | Blastopore |
| Eve | LOC373529 | Veg1 Ectoderm |

| | | |
|---|---|---|
| CyIIIa<br>EGF2<br>Endo16_negative | LOC373298<br>LOC373304<br>LOC115926321 | |
| Pks1<br>Srcr142 | LOC588806<br>LOC115929142 | Pigment Cells |
| MacpfA2 | LOC763692 | Globular Cells |
| c3<br>COLP1alpha | LOC373284<br>LOC373267 | Filopodial Cells |
| Vasa | LOC576055 | Left Pouch, CPCs |
| SoxE | LOC581729 | Right Pouch, CPCs |
| SynB<br>CHAT<br>TH<br>TPH | LOC115917880<br>LOC574696<br>LOC581094<br>LOC578903 | Neuron |
| SoxC<br>Delta | LOC593520<br>LOC115921237 | Neuron_precurseurs |
| Tph<br>Np18<br>Sp-AN | LOC578903<br>LOC752246<br>LOC763123 | Neuron_ApicalOrgan |
| FSalmFa<br>NGFFFa | LOC763080<br>LOC752998 | Neron_Dopaminergic |
| prox1<br>ScratchX | LOC576148<br>LOC580246 | Neuron_CiliaryBand |
| Nanos2<br>vasa | LOC751842<br>LOC576055 | Primordial germ cells<br>(PGCs) |
| Gcm<br>Six1 | LOC378470<br>LOC576117 | AboralNSM |
| GATAc<br>Ese<br>prox1 | LOC373316<br>LOC582205<br>LOC576148 | OralNSM |

| | | |
|---|---|---|
| Endo16 positive<br>FoxA | LOC115926321<br>LOC578584 | Veg2_Endoderm |
| FoxA<br>Endo16 negative | LOC578584<br>LOC115926321 | Veg1_Endoderm |
| Eve<br>Wnt8 | LOC373529<br>LOC397068 | Veg1_EndoMeso |
| FoxA<br>Gcm<br>GATAe | LOC578584<br>LOC378470<br>LOC408052 | Veg2_EndoMeso |

*FoxG expression is restricted to ciliary band at pluteus stage, while at earlier stages, FoxG appears to be a marker of oral ectoderm.

**Supplementary Table 3**. Cell-type-specific marker genes obtained by scRNA-seq of individual time points.

**Separate excel file.**

**Supplementary Table 4**. Primers used for qPCR validation of calcification genes.

| | Gene | Primer sequence | product size (bp) |
|---|---|---|---|
| **Primer sequence** | PrestinL (LOC115929285) | F-GACAAGACCCCGGAAATCGT | 189 |
| | | R-GGAGATGGCGACCACGATGA | |
| | Slc13a2 (LOC588117) | F-TTGGGGGCCCAGTTTTCG | 157 |
| | | R-GTCCGATGCTCTTGCTCAAT | |
| | Msp130L (LOC594269) | F-CCTAGTCCTGGCAATGCTCG | 186 |
| | | R-CTCGACGAACTCTCCACCAC | |
| | Sm30b (LOC115918134) | F-ACAGCTGCTACCTGTTCGAC | 197 |
| | | R-CATTCTGACAGCCCACTCGT | |
| | Sm50 (LOC105437370) | F-TGCTCTGGCTTCAGTTTCGT | 168 |
| | | R-GCGGCGAATCCGTTAGGATA | |
| | NBC3L/SLC4A8 (LOC115918074 ) | F-GAGGACTATATCGCGGCCTT | 190 |
| | | R-ATCCGATCCACACTCGGAAC | |
| | Sepal (LOC115918256) | F-GTGTTGGTGTGCGTATTGGC | 183 |
| | | R-ACCACTGTCGAACAGGTAGC | |
| | NBC/SLC4A10 (LOC580912) | F-GTTCTTGTTCTCTTGCGCCCTC | 139 |
| | | R-AGCCAGGAAAGCCATGAAGAC | |
| | Otop2L (LOC579173) | F-GTTGGCAGACGGATCACAGC | 200 |
| | | R-TAGAAGCACCTGCATGACGG | |
| | Cara7 (LOC579101) | F-GAACGGTAACGGATGGGGAG | 148 |
| | | R-GGGTCCGTTCATGCCAAAAG | |

**Supplementary Table 5**. Number of cells and quality metrics for each cross in scATAC-seq.

| Cross | Library | Final cells | Fragments per cell | FRiP | TSSE score |
|---|---|---|---|---|---|
| Hpf3 | scEarlyDev (with NFB) | | 2138.5 | 14.85 | 1.94 |
| Hpf4 | | 10 | 2422 | 14.48 | 1.61 |
| Hpf5_1 | | 29 | 2586 | 19.86 | 2.10 |
| Hpf5_2 | | 10 | 3767 | 20.46 | 2.53 |
| Hpf6_1 | | 56 | 4728.5 | 18.91 | 2.36 |
| Hpf6_2 | | 40 | 2466.5 | 15.45 | 1.66 |
| Hpf8_1 | | 103 | 2393 | 15.23 | 1.80 |
| Hpf10_1 | | 384 | 3064 | 24.06 | 2.82 |
| Hpf12_1 | | 291 | 8228 | 28.43 | 3.49 |
| Hpf12_2 | | 555 | 3745 | 28.96 | 3.04 |
| Hpf15_1 | | 535 | 6434 | 27.79 | 3.66 |
| Hpf15_2 | | 3085 | 3932.5 | 31.28 | 3.59 |
| Hpf24 | scHpf24 | 161 | 14043 | 40.97 | 5.14 |
| Hpf20 | scLateDev (Hpf72_2 frozen in NFB) | 850 | 4851 | 29.89 | 4.64 |
| Hpf40 | | 2472 | 4240 | 26.44 | 4.19 |
| Hpf72_1 | | 1824 | 4542.5 | 29.97 | 4.89 |
| Hpf72_2 | | 2732 | 4931 | 31.9 | 5.11 |
| Total | --- | 19412 | 6429.5 | 33.14 | 4.69 |

**Supplementary Table 6**. Lineage specific differentially accessible peaks obtained by scATAC-seq.

**Separate excel file.**

**Supplementary Table 7**. Cell-type-specific differentially accessible peaks obtained by scATAC-seq of individual time points.

**Separate excel file.**

**Supplementary Table 8**. List of lineage-specific candidate transcription factors.

**Separate excel file.**

**Supplementary Table 9.** Coordinates of regions with groups of accessible elements and defined regulatory landscapes in the sea urchin at the 1-cell stage (Hpf1).

| Chromosome | Start position | End position |
|---|---|---|
| NW_022145595.1 | 29897988 | 30069416 |
| NW_022145596.1 | 5692 | 55742 |
| NW_022145597.1 | 1829761 | 1857386 |
| NW_022145597.1 | 5119622 | 5260036 |
| NW_022145597.1 | 34049012 | 34147416 |
| NW_022145601.1 | 28999832 | 29449762 |
| NW_022145603.1 | 19840077 | 19919760 |
| NW_022145605.1 | 599955 | 699894 |
| NW_022145606.1 | 18179815 | 18339928 |
| NW_022145607.1 | 42896847 | 44998651 |
| NW_022145611.1 | 3790727 | 3820506 |

**Supplementary Table 10**. Motifs derived from early opening peak sets

**Provided as PDF files**

**Supplementary Table 11**. Top motifs associated with each regulatory state

Unless states otherwise "best match" refers to human TFs. Only motifs top3 with strong similarity to human or fly TF binding motifs are reported. States that are missing were not enriched for any motif with high similarity to human or fly motifs.

| State | Motif | P-value | % of Targets | % of Background | Best match | Score |
|---|---|---|---|---|---|---|
| 0 | 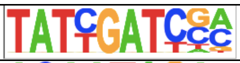 | 1e-112 | 82.05 | 18.77 | Hnf6 | 0.95 |
| 0 | 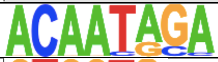 | 1e-38 | 70.33 | 31.39 | SOX15 | 0.86 |
| 0 | 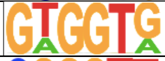 | 1e-29 | 89.38 | 58.16 | Fly::run | 0.87 |
| 1 | 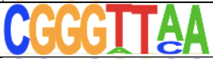 | 1e-30 | 81.71 | 50.73 | Fly::kr | 0.88 |
| 1 | 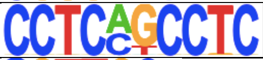 | 1e-27 | 79.88 | 50.66 | ZNF460 | 0.83 |
| 2 | 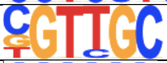 | 1e-79 | 86.98% | 49.62% | Rfx7 | 0.89 |
| 2 | 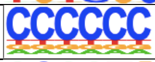 | 1e-44 | 96.53% | 75.21% | VEZF1 | 0.96 |
| 3 | 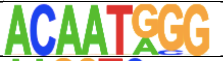 | 1e-38 | 44.08% | 11.16% | SOX13 | 0.90 |
| 3 | 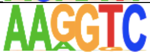 | 1e-34 | 92.24% | 56.68% | NR2F2 | 0.89 |
| 3 | 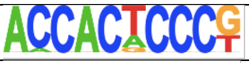 | 1e-30 | 19.59% | 2.18% | KLF2 | 0.85 |
| 4 | 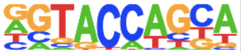 | 1e-133 | 72.53% | 12.85% | GCM1 | 0.77 |
| 4 | 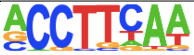 | 1e-44 | 82.41% | 44.25 | Fly::PUM | 0.86 |
| 4 | 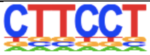 | 1e-37 | 70.68% | 35.41 | ETV4 | 0.98 |
| 5 | 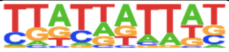 | 1e-64 | 80.22% | 44.81% | Fly::AP | 0.80 |
| 5 | 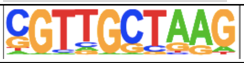 | 1e-58 | 84.21% | 51.38% | RFX4 | 0.91 |
| 5 | 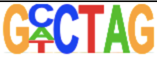 | 1e-56 | 74.59% | 41.00% | Smad4 | 0.82 |
| 6 | 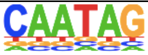 | 1e-61 | 74.58% | 34.45 | SOX9 | 0.81 |
| 7 | 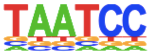 | 1e-52 | 77.19% | 32.91 | PITX1 | 0.99 |
| 7 | 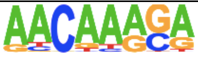 | 1e-47 | 82.11% | 39.90 | Sox10 | 0.88 |
| 7 | 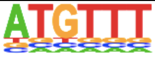 | 1e-37 | 92.28% | 58.21% | FoxA1 | 0.84 |
| 8 | 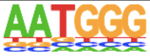 | 1e-50 | 86.43% | 51.72% | FOXL2 | 0.80 |
| 9 | 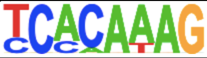 | 1e-29 | 82.96% | 35.30% | Sox10 | 0.81 |
| 10 | 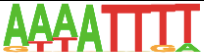 | 1e-199 | 87.65% | 48.38% | Fly::cad | 0.84 |
| 11 | 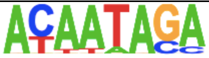 | 1e-44 | 83.23% | 30.17% | Sox15 | 0.82 |
| 11 | 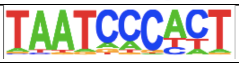 | 1e-33 | 82.04% | 36.08% | Pitx2 | 0.83 |

| | | | | | | |
|---|---|---|---|---|---|---|
| 11 | TGATTGGC | 1e-32 | 77.84% | 32.56% | Pknox1 | 0.85 |
| 12 | TAATCC | 1e-36 | 63.47% | 31.55% | Pitx1 | 0.99 |
| 12 | AGACAAAGCC | 1e-34 | 86.40% | 56.89% | Sox10 | 0.82 |
| 13 | CTATTC | 1e-38 | 87.27% | 49.88% | Fly:: br(var.2) | 0.82 |
| 13 | GAAAGT | 1e-34 | 76.73% | 39.87% | PTBP1(RRM) | 0.88 |
| 13 | TAATCC | 1e-30 | 66.91% | 32.67% | PITX1 | 0.97 |
| 14 | CAGGAAGT | 1e-55 | 80.47% | 32.44% | ELF3(ETS) | 0.95 |
| 15 | CGTCGC | 1e-52 | 79.42% | 40.89% | RSF1(RRM) | 0.82 |
| 15 | GTGGTG | 1e-52 | 90.77% | 54.89% | Fly::run | 0.93 |
| 16 | CTATTT | 1e-48 | 83.60% | 47.23% | MEF2C | 0.84 |
| 16 | TGATTACG | 1e-46 | 71.69% | 35.27% | FOXD1 | 0.89 |
| 16 | ACTGTC | 1e-44 | 64.02% | 28.90% | ENOX1(RRM) | 0.85 |
| 17 | GGAAGT | 1e-111 | 96.07% | 52.97% | Elf4(ETS) | 0.90 |
| 17 | ATGACTCA | 1e-90 | 72.71% | 30.24% | FOS::JUND | 0.96 |
| 17 | CAGCTGTC | 1e-77 | 71.78% | 32.30% | MYF5 | 0.95 |
| 18 | CAATAG | 1e-55 | 75.99% | 33.54% | SOX9 | 0.81 |
| 19 | GCAAACAG | 1e-95 | 91.39% | 48.56% | FOXP3 | 0.85 |
| 19 | AGATAA | 1e-71 | 82.39% | 43.79% | GATA2 | 0.95 |
| 19 | TGAGTCAG | 1e-64 | 82.19% | 45.59% | BATF::JUN | 0.8 |
| 20 | GATTGACAGG | 1e-24 | 36.43% | 5.80% | PBX3 | 0.88 |
| 20 | ACACTTGA | 1e-21 | 58.14% | 19.50% | NKX2-8 | 0.82 |
| 20 | GCTAATGA | 1e-19 | 68.99% | 29.31% | HoxB4 | 0.86 |
| 21 | TATTGATT | 1e-36 | 76.17% | 31.45% | ONECUT1 | 0.93 |
| 21 | ATCGGATTAA | 1e-31 | 86.53% | 46.07% | Ptx1 | 0.82 |
| 22 | TGTTTGCGAT | 1e-65 | 78.60% | 30.40% | Fly::fkh | 0.83 |
| 22 | TGTAAACATT | 1e-39 | 84.62% | 47.90% | FoxA1 | 0.914 |
| 23 | TATGCTGGTA | 1e-134 | 78.95% | 28.86% | GCM1 | 0.81 |
| 25 | GGGGATTA | 1e-36 | 78.93% | 38.77% | Pitx2 | 0.93 |
| 27 | CCCATTGT | 1e-45 | 70.52% | 27.06% | SOX15 | 0.85 |
| 27 | CTTAATCC | 1e-34 | 71.71% | 33.33% | PITX3 | 0.96 |

| | | | | | | |
|---|---|---|---|---|---|---|
| 28 | 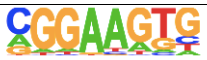 | 1e-57 | 76.54% | 27.72% | ETV5(ETS) | 0.96 |
| 28 | 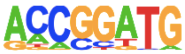 | 1e-43 | 84.23% | 42.08% | Ets2 | 0.82 |

**Supplementary Table 12**. Table of cell-type-specific enriched motifs

**Provided as PDF**

**Supplementary Table 13**. Table of motifs enriched per developmental stage

**Provided as PDF**

**Supplementary Table 14**. List of transcription factors involved in neurodevelopment in *S. purpuratus* and evidence for their role in neurodevelopment in other animals.

| **Gene name** | **LOC Number** | **homolog** | **Reference** | **Function neurodevelopment** | **Related function** |
|---|---|---|---|---|---|
| Acsl1 | LOC593387 | Acsl1 | https://www.nature.com/articles/nn1247 | T | NA |
| SoxC | LOC593520 | Sox4 | https://www.jneurosci.org/content/jneuro/32/9/3067.full.pdf | T | NA |
| FoxN1/4 | LOC584374 | FoxN4 | https://journals.biologists.com/dev/article/134/19/3427/64495/A-regulatory-network-involving-Foxn4-Mash1-and | T | NA |
| ZEB2 | LOC115917895 | ZEB2 | https://www.sciencedirect.com/science/article/abs/pii/S0006899318304992 | T | NA |
| Rcor | LOC575435 | | | NA | NA |
| Six3/6 | LOC576281 | Six3 | https://academic.oup.com/cercor/article/18/3/553/286638 | T | NA |
| lhx6 | LOC579936 | LHX6 | https://www.jneurosci.org/content/27/12/3078.short | T | NA |
| hbn | LOC575572 | hbn:fly | https://www.sciencedirect.com/science/article/pii/S2667290121000048 | T | NA |
| lhx9 | LOC576656 | lhx9 | https://www.ncbi.nlm.nih.gov/pmc/articles/PMC3236734/ | T | NA |
| meox1 | LOC577108 | MEOX1 | https://pubmed.ncbi.nlm.nih.gov/12925591/ | F | Somite |
| lhx9l | LOC115929082 | Lhx2 | https://www.ncbi.nlm.nih.gov/pmc/articles/PMC3236734/ | T | NA |
| COE3 | LOC577048 | | | NA | NA |

| | | | | | |
|---|---|---|---|---|---|
| Ebf | LOC100889714 | EBF2 | https://pubmed.ncbi.nlm.nih.gov/22042145/ | T | NA |
| Rx | LOC576952 | Rx1:zebrafish | https://core.ac.uk/download/pdf/82169985.pdf | F | Retina |
| ATF6 | LOC589006 | Atf6 | https://jbiomedsci.biomedcentral.com/articles/10.1186/s12929-018-0453-1 | T | NA |
| FoxD | LOC751845 | FoxD3 | https://journals.biologists.com/dev/article/135/9/1615/64940/Requirement-for-Foxd3-in-the-maintenance-of-neural | F | Neural crest |
| SHR3 | LOC574072 | nr6a1:zebrafish | https://febs.onlinelibrary.wiley.com/doi/epdf/10.1111/j.1432-1033.1997.t01-1-00826.x | T | NA |
| fezf2 | LOC100889201 | FezF2 | https://onlinelibrary.wiley.com/doi/10.1002/bies.201400039 | T | NA |
| Dach1 | LOC586986 | Dach1 | https://www.ncbi.nlm.nih.gov/pmc/articles/PMC6458905/ | T | NA |
| lmx1b | LOC585635 | Lmx1b | https://www.jneurosci.org/content/31/35/12413.long | T | NA |
| otp | LOC115920693 | OTP | https://www.sciencedirect.com/science/article/pii/S0012160600999020 | T | NA |
| erg | LOC115917879 | ERG | https://www.sciencedirect.com/science/article/pii/S0925477399002725?via%3Dihub | F | Neural crest |
| pou1f1 | LOC577601 | POU1F1 | https://www.tandfonline.com/doi/pdf/10.1080/07853890600994963 | F | Pituitary |
| otp | LOC579386 | OTP | https://www.sciencedirect.com/science/article/pii/S0012160600999020 | T | NA |
| Nk1 | NK1 | | | NA | NA |
| nr4a2 | LOC580856 | NR4A2 | https://linkinghub.elsevier.com/retrieve/pii/S1044-7431(08)00163-2 | T | NA |

| | | | | | |
|---|---|---|---|---|---|
| Scratch | LOC580856 | Scratch1 | https://anatomypubs.onlinelibrary.wiley.com/doi/10.1002/dvdy.20869 | T | NA |
| onecut | LOC115917884 | ONECUT2 | https://journals.biologists.com/dev/article/146/12/dev173807/19473/Single-cell-transcriptomics-reveals-spatial-and | T | NA |
| cux1 | LOC575361 | CUX1 | https://www.ncbi.nlm.nih.gov/pmc/articles/PMC2894581/ | T | NA |
| dri | dri | arid3a | https://academic.oup.com/gigascience/article/7/11/giy117/5099469 | T | NA |
| Klf7 | Klf7 | Klf6 | https://journals.plos.org/plosbiology/article/file?id=10.1371/journal.pbio.1002467&type=printable | T | NA |
| isl1 | LOC576365 | Isl1 | https://www.nature.com/articles/nn.2209 | T | NA |
| prox1 | LOC576148 | Prox1 | https://www.frontiersin.org/articles/10.3389/fncel.2014.00454/full | T | NA |
| mtf1 | LOC588463 | MTF1 | | F | NA |
| foxn3 | LOC590815 | foxn2 | https://reader.elsevier.com/reader/sd/pii/S0925477302002204?token=726BAF999305C6D2E8B2BCE0E2D218F7AFDBBF18EC041E8EE2C3F9FE97DEC52B4EDF285190C4FA249358CA8ABA665CC4&originRegion=eu-west-1&originCreation=20211210204632 | F | craniofacial |
| | | | | 26/32 (total 35) | 6/6 (total 9) |

**Supplementary table 15**. Reference peak sets per stage.

**Separate excel file.**
