## Supplemental Table 10 for "The Establishment of Cell-Type Specific Gene Regulation in the Sea Urchin Embryo": BlastulaPeaks_homerResults.html

/fast/users/jbrande\_m/work/scEarlyDev/bulkATAC//earlyMotifs//Blastula/ - Homer de novo Motif Results


### Homer *de novo* Motif Results (/fast/users/jbrande\_m/work/scEarlyDev/bulkATAC//earlyMotifs//Blastula/)

Known Motif Enrichment Results  
Gene Ontology Enrichment Results  
If Homer is having trouble matching a motif to a known motif, try copy/pasting the matrix file into
STAMP  
More information on motif finding results: HOMER
| Description of Results
| Tips
  
Total target sequences = 22593  
Total background sequences = 18368  
\* - possible false positive  

|  |  |  |  |  |  |  |  |  |
| --- | --- | --- | --- | --- | --- | --- | --- | --- |
| Rank | Motif | P-value | log P-pvalue | % of Targets | % of Background | STD(Bg STD) | Best Match/Details | Motif File |
| 1 | G T A C A C T G A C G T A G T C A G T C A C G T C G T A C G T A A C G T A G C T | 1e-44 | -1.027e+02 | 0.18% | 0.01% | 145.9bp (115.3bp) | ZmHOX2a(1)(HD-HOX)/Zea mays/AthaMap(0.769) More Information \| Similar Motifs Found | motif file (matrix) |
| 2 | A C G T A G T C A G T C A C T G A G T C A C G T A G T C A C G T A G T C A C G T | 1e-27 | -6.354e+01 | 0.18% | 0.02% | 144.5bp (139.8bp) | Trl/dmmpmm(Pollard)/fly(0.775) More Information \| Similar Motifs Found | motif file (matrix) |
| 3 | C G T A A C G T A G T C C G T A A G T C C T A G A G T C A G T C C G T A G A C T | 1e-22 | -5.222e+01 | 0.54% | 0.19% | 150.2bp (196.2bp) | Srebp1a(bHLH)/HepG2-Srebp1a-ChIP-Seq(GSE31477)/Homer(0.922) More Information \| Similar Motifs Found | motif file (matrix) |
| 4 | A G T C C G T A A C G T A C G T A G T C C T G A A C T G C G T A A G T C A G T C | 1e-21 | -4.961e+01 | 0.13% | 0.01% | 135.0bp (274.2bp) | TARDBP(RRM)/Homo\_sapiens-RNCMPT00076-PBM/HughesRNA(0.694) More Information \| Similar Motifs Found | motif file (matrix) |
| 5 | T A G C T G A C G A T C G C A T G A C T C G T A C G T A C G T A C T A G A T C G | 1e-21 | -4.880e+01 | 7.95% | 6.34% | 155.3bp (188.0bp) | RPH1/Literature(Harbison)/Yeast(0.760) More Information \| Similar Motifs Found | motif file (matrix) |
| 6 | A C T G A G T C G T C A A G T C C G T A A C G T A C T G C G T A A C T G A C G T | 1e-16 | -3.706e+01 | 0.18% | 0.03% | 168.0bp (131.5bp) | MXI1/MA1108.2/Jaspar(0.729) More Information \| Similar Motifs Found | motif file (matrix) |
| 7 \* | C T A G T C A G A T C G C T A G A G T C C A G T G C A T C G A T A G C T G C T A | 1e-11 | -2.631e+01 | 12.15% | 10.72% | 153.1bp (187.4bp) | dl/MA0022.1/Jaspar(0.618) More Information \| Similar Motifs Found | motif file (matrix) |
| 8 \* | T C G A T C A G C G T A T A C G C A T G T A G C G C A T G A C T A C G T C G T A | 1e-8 | -1.878e+01 | 8.02% | 7.03% | 160.5bp (196.8bp) | TRP2(MYBrelated)/colamp-TRP2-DAP-Seq(GSE60143)/Homer(0.775) More Information \| Similar Motifs Found | motif file (matrix) |
| 9 \* | G A C T C T A G A C G T G C A T C A G T T A C G A T G C G C A T G A C T C G T A | 1e-8 | -1.850e+01 | 1.64% | 1.21% | 144.5bp (202.1bp) | fkh/dmmpmm(Noyes)/fly(0.900) More Information \| Similar Motifs Found | motif file (matrix) |
| 10 \* | A G C T T A C G C G A T C G T A G T A C G T C A A C G T C T A G A G C T C G T A | 1e-6 | -1.496e+01 | 4.24% | 3.60% | 161.6bp (222.6bp) | PH0084.1\_Irx3\_2/Jaspar(0.886) More Information \| Similar Motifs Found | motif file (matrix) |
| 11 \* | G C A T A T C G T G A C A G T C T A G C A T G C G T A C G T A C G C A T T G A C | 1e-6 | -1.488e+01 | 16.15% | 14.96% | 158.2bp (206.0bp) | VEZF1/MA1578.1/Jaspar(0.761) More Information \| Similar Motifs Found | motif file (matrix) |
| 12 \* | T C A G C A G T G T A C A G T C G C T A C G A T C A G T T G C A | 1e-3 | -8.327e+00 | 8.30% | 7.68% | 156.3bp (195.8bp) | Isl1/MA1608.1/Jaspar(0.848) More Information \| Similar Motifs Found | motif file (matrix) |
| 13 \* | T C G A T C G A A T C G T A C G A T G C C A T G A G T C C G A T G T C A G C A T | 1e-2 | -5.301e+00 | 2.65% | 2.39% | 161.6bp (176.7bp) | STP4/MA0397.1/Jaspar(0.731) More Information \| Similar Motifs Found | motif file (matrix) |
| 14 \* | A T C G C G T A C A G T C G A T C G T A C G T A G C T A A T G C T G C A A T C G | 1e-2 | -4.740e+00 | 1.18% | 1.02% | 152.9bp (194.1bp) | Arid3a/MA0151.1/Jaspar(0.756) More Information \| Similar Motifs Found | motif file (matrix) |
| 15 \* | C T G A C G T A A C G T A G T C A C G T A G T C A C G T A G T C A C G T A G T C | 1e-1 | -4.072e+00 | 0.34% | 0.26% | 195.6bp (218.1bp) | SeqBias: GA-repeat(0.809) More Information \| Similar Motifs Found | motif file (matrix) |
| 16 \* | A C T G C G T A C G T A C G T A C G T A C G T A C G T A C G T A C G T A A C T G | 1e-1 | -3.494e+00 | 0.74% | 0.64% | 147.6bp (207.6bp) | REM19(REM)/colamp-REM19-DAP-Seq(GSE60143)/Homer(0.853) More Information \| Similar Motifs Found | motif file (matrix) |
| 17 \* | T C G A A G C T A C T G G A T C A T C G A C G T A G T C G C T A | 1e0 | -1.810e+00 | 2.72% | 2.62% | 156.1bp (196.2bp) | FHL1/MA0295.1/Jaspar(0.814) More Information \| Similar Motifs Found | motif file (matrix) |
| 18 \* | C G T A A C G T A C G T C G T A A G T C A G T C A G T C T G C A | 1e0 | -1.632e+00 | 3.50% | 3.40% | 168.7bp (201.9bp) | REB1/MA0363.1/Jaspar(0.848) More Information \| Similar Motifs Found | motif file (matrix) |
| 19 \* | A T C G A T C G C A T G C A G T C G A T C G A T C G A T C T G A | 1e0 | -7.805e-01 | 3.73% | 3.72% | 157.8bp (192.7bp) | TRP2(MYBrelated)/colamp-TRP2-DAP-Seq(GSE60143)/Homer(0.729) More Information \| Similar Motifs Found | motif file (matrix) |
| 20 \* | G T A C C G T A A C G T C T A G A G C T A G C T | 1e0 | 0.000e+00 | 24.05% | 27.76% | 163.3bp (189.8bp) | Tv\_0259(RRM)/Trichomonas\_vaginalis-RNCMPT00259-PBM/HughesRNA(0.841) More Information \| Similar Motifs Found | motif file (matrix) |
