## Supplemental Table 10 for "The Establishment of Cell-Type Specific Gene Regulation in the Sea Urchin Embryo": Gastrula_homerResults.html

/fast/users/jbrande\_m/work/scEarlyDev/bulkATAC//earlyMotifs//Gastrula/ - Homer de novo Motif Results


### Homer *de novo* Motif Results (/fast/users/jbrande\_m/work/scEarlyDev/bulkATAC//earlyMotifs//Gastrula/)

Known Motif Enrichment Results  
Gene Ontology Enrichment Results  
If Homer is having trouble matching a motif to a known motif, try copy/pasting the matrix file into
STAMP  
More information on motif finding results: HOMER
| Description of Results
| Tips
  
Total target sequences = 3667  
Total background sequences = 22561  
\* - possible false positive  

|  |  |  |  |  |  |  |  |  |
| --- | --- | --- | --- | --- | --- | --- | --- | --- |
| Rank | Motif | P-value | log P-pvalue | % of Targets | % of Background | STD(Bg STD) | Best Match/Details | Motif File |
| 1 | C A T G A G T C T G C A C G T A G C T A A G T C T C G A A G T C G C A T C A T G | 1e-23 | -5.480e+01 | 46.50% | 38.24% | 161.2bp (180.9bp) | PHA-4(Forkhead)/cElegans-Embryos-PHA4-ChIP-Seq(modEncode)/Homer(0.907) More Information \| Similar Motifs Found | motif file (matrix) |
| 2 | C G T A T G C A C T G A C G T A G C T A T A G C T G A C T G A C C T G A C T A G | 1e-17 | -3.949e+01 | 1.88% | 0.55% | 173.9bp (185.4bp) | NCU02182/MA1434.1/Jaspar(0.749) More Information \| Similar Motifs Found | motif file (matrix) |
| 3 | C G A T T G A C G T A C G T C A C G T A C A G T A G T C C T A G C G T A A G C T | 1e-14 | -3.402e+01 | 12.76% | 8.84% | 159.8bp (170.0bp) | Cux2(Homeobox)/Liver-Cux2-ChIP-Seq(GSE35985)/Homer(0.734) More Information \| Similar Motifs Found | motif file (matrix) |
| 4 | A C T G G C A T A G C T T C G A C G T A A G C T A G T C C G T A A C G T A G C T | 1e-14 | -3.323e+01 | 5.37% | 2.95% | 148.3bp (187.7bp) | HNF1b(Homeobox)/PDAC-HNF1B-ChIP-Seq(GSE64557)/Homer(0.839) More Information \| Similar Motifs Found | motif file (matrix) |
| 5 | A C G T A G C T A C G T A G C T T C G A A T C G T C G A C G T A C T G A C T A G | 1e-13 | -3.169e+01 | 4.80% | 2.58% | 153.0bp (179.2bp) | ORB2(RRM)/Drosophila\_melanogaster-RNCMPT00126-PBM/HughesRNA(0.696) More Information \| Similar Motifs Found | motif file (matrix) |
| 6 | A C T G A C G T A C T G C G T A A G T C C G T A A C G T G T C A A C T G A T G C | 1e-13 | -3.032e+01 | 0.57% | 0.06% | 185.5bp (168.7bp) | Hth/dmmpmm(Noyes\_hd)/fly(0.717) More Information \| Similar Motifs Found | motif file (matrix) |
| 7 | T G C A A G C T C T G A C A G T T C G A A T G C A C T G A G C T G C T A C G A T | 1e-12 | -2.940e+01 | 10.36% | 7.08% | 155.7bp (187.9bp) | RBMS1(RRM)/Homo\_sapiens-RNCMPT00152-PBM/HughesRNA(0.711) More Information \| Similar Motifs Found | motif file (matrix) |
| 8 \* | A C T G C T A G A G T C A C G T A C T G G T C A A C T G C G T A A C G T A C G T | 1e-11 | -2.747e+01 | 1.04% | 0.26% | 171.5bp (196.6bp) | Gfi1b/MA0483.1/Jaspar(0.801) More Information \| Similar Motifs Found | motif file (matrix) |
| 9 \* | C G T A G T C A G T C A A C T G G T C A A C T G A C G T A G C T | 1e-11 | -2.575e+01 | 11.75% | 8.47% | 163.7bp (172.7bp) | ZNF652/MA1657.1/Jaspar(0.842) More Information \| Similar Motifs Found | motif file (matrix) |
| 10 \* | A G T C A C G T A C T G A C T G A C G T A G T C C G T A C G T A C G T A A C G T | 1e-10 | -2.445e+01 | 0.44% | 0.05% | 106.9bp (217.1bp) | Bcl11a(Zf)/HSPC-BCL11A-ChIP-Seq(GSE104676)/Homer(0.793) More Information \| Similar Motifs Found | motif file (matrix) |
| 11 \* | A G T C A C T G A C G T A G T C A C G T A G C T C G T A C G T A A G T C A C G T | 1e-10 | -2.375e+01 | 0.57% | 0.09% | 150.4bp (155.8bp) | RSF1(RRM)/Drosophila\_melanogaster-RNCMPT00061-PBM/HughesRNA(0.674) More Information \| Similar Motifs Found | motif file (matrix) |
| 12 \* | C G T A A C T G C A T G C T A G G T C A T C G A C G T A T C A G | 1e-9 | -2.215e+01 | 27.76% | 23.32% | 165.8bp (180.7bp) | PCBP3(KH)/Mus\_musculus-RNCMPT00215-PBM/HughesRNA(0.932) More Information \| Similar Motifs Found | motif file (matrix) |
| 13 \* | T A C G G C T A G C T A G T C A A G T C A G T C C G T A G T A C | 1e-9 | -2.183e+01 | 10.09% | 7.29% | 177.2bp (178.6bp) | RUNX1(Runt)/Jurkat-RUNX1-ChIP-Seq(GSE29180)/Homer(0.868) More Information \| Similar Motifs Found | motif file (matrix) |
| 14 \* | A C G T A C G T A C T G A C T G C G T A A G T C A C G T C G T A A G T C A C G T | 1e-8 | -1.989e+01 | 0.22% | 0.01% | 51.3bp (40.9bp) | MSI(RRM)/Drosophila\_melanogaster-RNCMPT00040-PBM/HughesRNA(0.681) More Information \| Similar Motifs Found | motif file (matrix) |
| 15 \* | C A G T A T C G G T C A G A T C C T A G A C G T G T A C C G T A G T A C A T C G | 1e-8 | -1.906e+01 | 6.95% | 4.80% | 134.8bp (147.6bp) | FEA4(bZIP)/Corn-FEA4-ChIP-Seq(GSE61954)/Homer(0.949) More Information \| Similar Motifs Found | motif file (matrix) |
| 16 \* | C T G A A T C G A C T G C T G A G T C A G A C T C G T A A G T C | 1e-8 | -1.880e+01 | 8.40% | 6.04% | 159.8bp (167.9bp) | Ik-1(0.816) More Information \| Similar Motifs Found | motif file (matrix) |
| 17 \* | C G T A A C T G A G T C A C G T A C T G A C T G C G T A A G T C A C G T A C T G | 1e-7 | -1.679e+01 | 0.22% | 0.02% | 122.1bp (130.8bp) | SAMD4A(SAM)/Homo\_sapiens-RNCMPT00063-PBM/HughesRNA(0.761) More Information \| Similar Motifs Found | motif file (matrix) |
| 18 \* | C G T A G T A C G T A C C G A T A C T G A C G T T G C A G A T C | 1e-6 | -1.469e+01 | 21.57% | 18.35% | 165.8bp (191.3bp) | SNAI2/MA0745.2/Jaspar(0.774) More Information \| Similar Motifs Found | motif file (matrix) |
| 19 \* | A G T C A C T G A C T G A C G T C G T A A C T G A C G T C G A T | 1e-5 | -1.358e+01 | 1.53% | 0.76% | 137.0bp (158.9bp) | Rbm42(RRM)/Xenopus\_tropicalis-RNCMPT00282-PBM/HughesRNA(0.816) More Information \| Similar Motifs Found | motif file (matrix) |
| 20 \* | A C T G A C T G A C T G A C T G G T C A A C T G A G T C G T C A C T A G A C T G | 1e-4 | -1.075e+01 | 0.35% | 0.08% | 110.8bp (161.5bp) | ZNF263/MA0528.2/Jaspar(0.777) More Information \| Similar Motifs Found | motif file (matrix) |
| 21 \* | A G T C A C G T A C G T C G T A A C T G C G T A A C T G A C T G A C T G C G T A | 1e-1 | -3.554e+00 | 0.11% | 0.03% | 122.3bp (128.9bp) | XBP1/Literature(Harbison)/Yeast(0.665) More Information \| Similar Motifs Found | motif file (matrix) |
