## Supplemental Table 10 for "The Establishment of Cell-Type Specific Gene Regulation in the Sea Urchin Embryo": Pre16cells_homerResults.html

/fast/users/jbrande\_m/work/scEarlyDev/bulkATAC//earlyMotifs//Pre16/ - Homer de novo Motif Results


### Homer *de novo* Motif Results (/fast/users/jbrande\_m/work/scEarlyDev/bulkATAC//earlyMotifs//Pre16/)

Known Motif Enrichment Results  
Gene Ontology Enrichment Results  
If Homer is having trouble matching a motif to a known motif, try copy/pasting the matrix file into
STAMP  
More information on motif finding results: HOMER
| Description of Results
| Tips
  
Total target sequences = 1147  
Total background sequences = 1163  
\* - possible false positive  

|  |  |  |  |  |  |  |  |  |
| --- | --- | --- | --- | --- | --- | --- | --- | --- |
| Rank | Motif | P-value | log P-pvalue | % of Targets | % of Background | STD(Bg STD) | Best Match/Details | Motif File |
| 1 | C G A T A C G T A C T G A G T C A C T G A G T C G T C A C G T A A C G T G T C A | 1e-15 | -3.608e+01 | 2.26% | 0.34% | 133.2bp (189.9bp) | CEBPD/MA0836.2/Jaspar(0.850) More Information \| Similar Motifs Found | motif file (matrix) |
| 2 | A G T C A T C G G A C T C T G A A G T C T A C G A T G C T C A G G A C T C T A G | 1e-15 | -3.554e+01 | 1.91% | 0.19% | 121.1bp (160.7bp) | TCFL5/MA0632.2/Jaspar(0.748) More Information \| Similar Motifs Found | motif file (matrix) |
| 3 | C T A G C A T G A C T G G T C A A G T C C T G A A C T G A G T C A G T C C T A G | 1e-15 | -3.475e+01 | 1.48% | 0.14% | 148.6bp (42.4bp) | ZNF341(Zf)/EBV-ZNF341-ChIP-Seq(GSE113194)/Homer(0.788) More Information \| Similar Motifs Found | motif file (matrix) |
| 4 \* | C T A G A C G T G A T C C T G A A G T C C T A G A G C T A T G C C G T A G T A C | 1e-11 | -2.635e+01 | 1.22% | 0.09% | 145.8bp (0.0bp) | FEA4(bZIP)/Corn-FEA4-ChIP-Seq(GSE61954)/Homer(0.868) More Information \| Similar Motifs Found | motif file (matrix) |
| 5 \* | G A C T C G A T A C T G A G T C G T C A C G T A C G T A A C G T C T A G A C G T | 1e-11 | -2.552e+01 | 1.83% | 0.34% | 123.2bp (141.5bp) | slbo/dmmpmm(Pollard)/fly(0.720) More Information \| Similar Motifs Found | motif file (matrix) |
| 6 \* | A C G T A C T G A T C G C G A T A C T G A C T G A G C T A C T G A T C G G C A T | 1e-9 | -2.297e+01 | 2.18% | 0.49% | 100.4bp (118.0bp) | Run/dmmpmm(Papatsenko)/fly(0.738) More Information \| Similar Motifs Found | motif file (matrix) |
| 7 \* | C G T A A C G T A C T G C G A T A C G T C G T A A C T G A C G T A G C T C A G T | 1e-7 | -1.663e+01 | 1.83% | 0.46% | 133.8bp (151.2bp) | Foxd3/MA0041.1/Jaspar(0.809) More Information \| Similar Motifs Found | motif file (matrix) |
| 8 \* | C T G A C G T A G T A C A G T C C G T A C G T A A C T G A G T C A T G C C G T A | 1e-6 | -1.546e+01 | 1.57% | 0.40% | 151.0bp (420.7bp) | HNRNPK(KH)/Homo\_sapiens-RNCMPT00026-PBM/HughesRNA(0.710) More Information \| Similar Motifs Found | motif file (matrix) |
| 9 \* | A T C G A G C T A T C G A C T G A C T G C G A T A C T G C G A T T C A G A G C T | 1e-6 | -1.516e+01 | 1.74% | 0.50% | 99.7bp (456.8bp) | KLF10(Zf)/HEK293-KLF10.GFP-ChIP-Seq(GSE58341)/Homer(0.823) More Information \| Similar Motifs Found | motif file (matrix) |
| 10 \* | C G T A G T A C A T G C T G A C T C A G T G C A C A T G G C T A C G A T G C T A | 1e-6 | -1.424e+01 | 5.05% | 2.55% | 134.9bp (320.7bp) | PB0126.1\_Gata5\_2/Jaspar(0.715) More Information \| Similar Motifs Found | motif file (matrix) |
| 11 \* | A G T C A G T C C T G A C T A G A G T C A G T C C G T A T C A G A T G C A G T C | 1e-5 | -1.373e+01 | 1.65% | 0.48% | 284.8bp (389.2bp) | ARF34/MA1693.1/Jaspar(0.722) More Information \| Similar Motifs Found | motif file (matrix) |
| 12 \* | A C T G A G T C A C G T A G T C A C T G A T C G A C G T G T A C C T A G A T G C | 1e-4 | -1.026e+01 | 1.74% | 0.63% | 84.3bp (173.9bp) | ZNF519(Zf)/HEK293-ZNF519.GFP-ChIP-Seq(GSE58341)/Homer(0.800) More Information \| Similar Motifs Found | motif file (matrix) |
| 13 \* | A C T G C G T A C G T A C G T A C G T A A C G T A C T G A C T G A T G C C T A G | 1e-3 | -8.452e+00 | 0.78% | 0.24% | 112.6bp (368.9bp) | ROX8(RRM)/Drosophila\_melanogaster-RNCMPT00148-PBM/HughesRNA(0.801) More Information \| Similar Motifs Found | motif file (matrix) |
| 14 \* | A C G T A T C G C T G A A C G T A T C G G T C A G A C T A T C G T G C A A G C T | 1e-3 | -8.352e+00 | 2.26% | 1.08% | 216.6bp (269.8bp) | PH0017.1\_Cux1\_2/Jaspar(0.802) More Information \| Similar Motifs Found | motif file (matrix) |
| 15 \* | A T C G C G A T A G T C A C G T A G T C A C G T A T G C C T A G G T A C C G T A | 1e-3 | -8.350e+00 | 1.13% | 0.36% | 142.8bp (1014.2bp) | SRSF10(RRM)/Homo\_sapiens-RNCMPT00090-PBM/HughesRNA(0.752) More Information \| Similar Motifs Found | motif file (matrix) |
| 16 \* | G T A C A G T C A C G T C G T A A C G T T A C G G T C A G T A C T A G C A G T C | 1e-3 | -8.264e+00 | 0.96% | 0.33% | 127.2bp (348.4bp) | THRB/MA1574.1/Jaspar(0.696) More Information \| Similar Motifs Found | motif file (matrix) |
| 17 \* | A C T G C T A G A C G T C A G T C G T A A C T G A C T G A C T G A C G T A G C T | 1e-2 | -6.022e+00 | 3.05% | 1.82% | 190.7bp (221.2bp) | AT1G72740/MA1353.1/Jaspar(0.746) More Information \| Similar Motifs Found | motif file (matrix) |
| 18 \* | G T A C A C G T C G T A G T A C A C G T G T C A A G T C A C G T C T G A A G T C | 1e-2 | -4.905e+00 | 0.87% | 0.39% | 85.6bp (225.5bp) | An\_0287(RRM)/Aspergillus\_nidulans-RNCMPT00287-PBM/HughesRNA(0.718) More Information \| Similar Motifs Found | motif file (matrix) |
| 19 \* | A C T G A C G T C T A G A C T G A G T C C T A G C G T A A C G T | 1e-1 | -4.253e+00 | 2.00% | 1.23% | 132.1bp (242.6bp) | MET31/MA0333.1/Jaspar(0.805) More Information \| Similar Motifs Found | motif file (matrix) |
| 20 \* | A C T G A T G C T A C G A C T G A T C G G A T C C G T A A C G T A T G C T G A C | 1e-1 | -3.457e+00 | 0.61% | 0.33% | 97.9bp (205.4bp) | TEA1/MA0405.1/Jaspar(0.752) More Information \| Similar Motifs Found | motif file (matrix) |
| 21 \* | A C T G A C G T A C G T C T A G A C G T A C T G A C G T A C G T | 1e-1 | -2.479e+00 | 3.57% | 2.88% | 206.8bp (357.6bp) | ETR-1(RRM)/Caenorhabditis\_elegans-RNCMPT00183-PBM/HughesRNA(0.817) More Information \| Similar Motifs Found | motif file (matrix) |
| 22 \* | A T G C T A G C A T C G A T C G C A G T G C A T A T G C A G T C | 1e0 | -1.882e+00 | 1.48% | 1.14% | 172.1bp (621.2bp) | YDR026C(MacIsaac)/Yeast(0.865) More Information \| Similar Motifs Found | motif file (matrix) |
| 23 \* | A C T G A C G T A G T C A C T G C G T A C G T A A G T C A C G T | 1e0 | -1.403e+00 | 0.70% | 0.57% | 100.1bp (280.4bp) | OPI1/Literature(Harbison)/Yeast(0.702) More Information \| Similar Motifs Found | motif file (matrix) |
| 24 \* | T C G A T G A C A T G C A G T C C G T A G T A C C T A G T A G C | 1e0 | -1.195e+00 | 1.22% | 1.04% | 77.4bp (167.2bp) | Egr1(Zf)/K562-Egr1-ChIP-Seq(GSE32465)/Homer(0.746) More Information \| Similar Motifs Found | motif file (matrix) |
| 25 \* | C T A G G T C A A G T C A C T G A G T C A C T G | 1e0 | -2.223e-01 | 8.36% | 9.09% | 104.8bp (254.3bp) | SWI6(MacIsaac)/Yeast(0.956) More Information \| Similar Motifs Found | motif file (matrix) |
| 26 \* | A C T G A G T C C G A T G T A C C G T A A G T C C G T A A G T C | 1e0 | -1.973e-01 | 2.35% | 2.83% | 90.9bp (171.2bp) | PB0130.1\_Gm397\_2/Jaspar(0.835) More Information \| Similar Motifs Found | motif file (matrix) |
| 27 \* | A C G T A C G T C G T A A G T C A C T G A G T C | 1e0 | -3.377e-02 | 4.96% | 6.21% | 442.5bp (229.9bp) | YAP3/MA0416.1/Jaspar(0.904) More Information \| Similar Motifs Found | motif file (matrix) |
| 28 \* | A C T G A G T C G T C A A C G T A C T G G T A C | 1e0 | -1.490e-04 | 6.09% | 9.01% | 80.5bp (284.2bp) | EIF-2ALPHA(S1)/Drosophila\_melanogaster-RNCMPT00273-PBM/HughesRNA(0.933) More Information \| Similar Motifs Found | motif file (matrix) |
| 29 \* | C A G T A G C T A G T C G T C A C G A T A G T C | 1e0 | 0.000e+00 | 17.75% | 29.39% | 216.4bp (260.6bp) | RBM47(RRM)/Gallus\_gallus-RNCMPT00279-PBM/HughesRNA(0.938) More Information \| Similar Motifs Found | motif file (matrix) |
