## Supplemental Table 10 for "The Establishment of Cell-Type Specific Gene Regulation in the Sea Urchin Embryo": PreZGAPeaks_homerResults.html

/fast/users/jbrande\_m/work/scEarlyDev/bulkATAC//earlyMotifs//Early/ - Homer de novo Motif Results


### Homer *de novo* Motif Results (/fast/users/jbrande\_m/work/scEarlyDev/bulkATAC//earlyMotifs//Early/)

Known Motif Enrichment Results  
Gene Ontology Enrichment Results  
If Homer is having trouble matching a motif to a known motif, try copy/pasting the matrix file into
STAMP  
More information on motif finding results: HOMER
| Description of Results
| Tips
  
Total target sequences = 2521  
Total background sequences = 1113  
\* - possible false positive  

|  |  |  |  |  |  |  |  |  |
| --- | --- | --- | --- | --- | --- | --- | --- | --- |
| Rank | Motif | P-value | log P-pvalue | % of Targets | % of Background | STD(Bg STD) | Best Match/Details | Motif File |
| 1 | G C T A C T A G C T A G T A G C A T G C C A G T C G T A A T C G | 1e-108 | -2.492e+02 | 17.18% | 5.09% | 174.2bp (320.9bp) | ZNF711(Zf)/SHSY5Y-ZNF711-ChIP-Seq(GSE20673)/Homer(0.924) More Information \| Similar Motifs Found | motif file (matrix) |
| 2 | A C G T C G A T T C A G A G C T A C T G A C G T | 1e-96 | -2.223e+02 | 63.98% | 43.25% | 178.9bp (216.9bp) | HNRNPL(RRM)/Homo\_sapiens-RNCMPT00091-PBM/HughesRNA(0.914) More Information \| Similar Motifs Found | motif file (matrix) |
| 3 | A C T G C T G A A C G T A G T C A C G T C G T A | 1e-93 | -2.162e+02 | 55.61% | 35.45% | 173.6bp (223.3bp) | GAT3/MA0301.1/Jaspar(0.883) More Information \| Similar Motifs Found | motif file (matrix) |
| 4 | C G T A G T C A C G T A C G T A A T C G C G T A | 1e-92 | -2.124e+02 | 65.21% | 44.98% | 174.2bp (205.0bp) | pan/dmmpmm(Down)/fly(0.885) More Information \| Similar Motifs Found | motif file (matrix) |
| 5 | G T A C C G A T C T G A C A T G C T A G T A G C C A G T C G T A A T C G C T G A | 1e-91 | -2.116e+02 | 3.17% | 0.17% | 206.9bp (81.2bp) | HRB27C(RRM)/Drosophila\_melanogaster-RNCMPT00093-PBM/HughesRNA(0.674) More Information \| Similar Motifs Found | motif file (matrix) |
| 6 | A C T G A C G T C T G A C G T A C G T A A C G T | 1e-87 | -2.007e+02 | 71.88% | 52.57% | 175.6bp (190.4bp) | GTL1(Trihelix)/colamp-GTL1-DAP-Seq(GSE60143)/Homer(0.883) More Information \| Similar Motifs Found | motif file (matrix) |
| 7 | A C G T A C G T A C T G A C T G A G T C C G T A A C G T C G T A A G T C A G T C | 1e-82 | -1.898e+02 | 84.49% | 67.64% | 170.3bp (214.1bp) | NFIA/MA0670.1/Jaspar(0.763) More Information \| Similar Motifs Found | motif file (matrix) |
| 8 | G T A C A C G T C G T A A C T G A G T C A C G T C G T A A C T G | 1e-73 | -1.687e+02 | 4.20% | 0.43% | 175.6bp (89.8bp) | MSI1(RRM)/Homo\_sapiens-RNCMPT00176-PBM/HughesRNA(0.766) More Information \| Similar Motifs Found | motif file (matrix) |
| 9 | C G T A A G T C C T G A C G T A G T C A A G T C A G C T A T C G | 1e-68 | -1.576e+02 | 61.76% | 44.35% | 166.5bp (178.9bp) | PB0149.1\_Myb\_2/Jaspar(0.781) More Information \| Similar Motifs Found | motif file (matrix) |
| 10 | A C G T A C T G A G T C A C T G C T G A A C T G | 1e-67 | -1.562e+02 | 21.06% | 9.50% | 181.0bp (247.5bp) | CG7903(RRM)/Drosophila\_melanogaster-RNCMPT00144-PBM/HughesRNA(0.842) More Information \| Similar Motifs Found | motif file (matrix) |
| 11 | C G T A C G T A A C G T A C G T A G T C A C G T A C T G C G T A | 1e-66 | -1.526e+02 | 3.09% | 0.24% | 162.4bp (44.2bp) | MAFG/MA0659.2/Jaspar(0.696) More Information \| Similar Motifs Found | motif file (matrix) |
| 12 | A G C T A C G T A C G T C G T A A C G T C G T A C G T A A C G T | 1e-64 | -1.476e+02 | 16.26% | 6.53% | 162.9bp (267.4bp) | PB0079.1\_Sry\_1/Jaspar(0.829) More Information \| Similar Motifs Found | motif file (matrix) |
| 13 | A T C G A C T G A G T C A T C G C G T A A C G T | 1e-64 | -1.475e+02 | 61.72% | 44.94% | 172.8bp (235.9bp) | pho/MA1460.1/Jaspar(0.732) More Information \| Similar Motifs Found | motif file (matrix) |
| 14 | T G A C G C A T C A T G A C T G G C A T C G T A C G A T A G T C C G T A C A T G | 1e-61 | -1.424e+02 | 79.06% | 63.80% | 178.1bp (229.8bp) | so/MA0246.1/Jaspar(0.767) More Information \| Similar Motifs Found | motif file (matrix) |
| 15 | A G T C C T G A A G T C A C G T C G T A G T C A C G A T A G T C | 1e-61 | -1.416e+02 | 10.59% | 3.26% | 173.7bp (441.1bp) | PFF0320c(RRM)/Plasmodium\_falciparum-RNCMPT00235-PBM/HughesRNA(0.864) More Information \| Similar Motifs Found | motif file (matrix) |
| 16 | A C T G A G T C A C T G A C G T C G T A A G T C | 1e-60 | -1.400e+02 | 80.40% | 65.46% | 173.3bp (219.7bp) | PB0094.1\_Zfp128\_1/Jaspar(0.841) More Information \| Similar Motifs Found | motif file (matrix) |
| 17 | C G T A A C T G A C G T A G T C A G T C A C T G | 1e-60 | -1.394e+02 | 80.76% | 65.90% | 171.7bp (187.2bp) | YLR278C/MA0430.1/Jaspar(0.783) More Information \| Similar Motifs Found | motif file (matrix) |
| 18 | A G T C C G T A A C T G A T C G A C T G A C G T | 1e-60 | -1.382e+02 | 32.09% | 18.46% | 168.9bp (183.4bp) | YDR026C(MacIsaac)/Yeast(0.786) More Information \| Similar Motifs Found | motif file (matrix) |
| 19 | A C T G C G T A A C T G A T C G C G T A A C T G C T A G T G C A | 1e-58 | -1.340e+02 | 9.16% | 2.69% | 194.3bp (167.2bp) | TF3A(C2H2)/col-TF3A-DAP-Seq(GSE60143)/Homer(0.900) More Information \| Similar Motifs Found | motif file (matrix) |
| 20 | C G T A A C T G A C T G A C T G C G T A A T C G A T G C A C G T | 1e-57 | -1.317e+02 | 4.52% | 0.64% | 206.7bp (561.6bp) | HNRNPA2B1(RRM)/Homo\_sapiens-RNCMPT00024-PBM/HughesRNA(0.759) More Information \| Similar Motifs Found | motif file (matrix) |
| 21 | C G T A A G T C A G T C A C T G A C T G A C G T | 1e-53 | -1.224e+02 | 66.72% | 51.51% | 173.8bp (201.8bp) | TFCP2/MA0145.3/Jaspar(0.823) More Information \| Similar Motifs Found | motif file (matrix) |
| 22 | A C G T C G T A A G T C A C G T A C G T C G T A | 1e-50 | -1.155e+02 | 22.41% | 11.79% | 169.6bp (258.0bp) | YAP6(MacIsaac)/Yeast(0.848) More Information \| Similar Motifs Found | motif file (matrix) |
| 23 | G C A T C A T G G C T A G C T A G A T C C G A T A G C T A G T C C G T A G A T C | 1e-42 | -9.895e+01 | 5.39% | 1.28% | 158.6bp (50.0bp) | NR1I3/MA1534.1/Jaspar(0.769) More Information \| Similar Motifs Found | motif file (matrix) |
| 24 | A T C G A G C T A T G C C G A T A T G C G A C T A G C T C A G T T G A C A T C G | 1e-42 | -9.684e+01 | 91.59% | 81.99% | 178.4bp (189.0bp) | BPC1(BBRBPC)/colamp-BPC1-DAP-Seq(GSE60143)/Homer(0.749) More Information \| Similar Motifs Found | motif file (matrix) |
| 25 | A C T G C G T A T C G A A C G T C T G A C G T A C G T A C G T A C G A T G T A C | 1e-40 | -9.419e+01 | 2.94% | 0.44% | 149.1bp (181.0bp) | AHL20/MA0933.1/Jaspar(0.777) More Information \| Similar Motifs Found | motif file (matrix) |
| 26 | G T A C C G T A C A T G G T A C C T G A C T A G C G T A T G C A A G C T C G T A | 1e-38 | -8.834e+01 | 3.37% | 0.59% | 163.6bp (9.1bp) | CUP2/MA0287.1/Jaspar(0.772) More Information \| Similar Motifs Found | motif file (matrix) |
| 27 | C G T A A C T G A G T C A C G T A G T C A G T C A C T G A G T C A G T C A C G T | 1e-37 | -8.744e+01 | 78.46% | 66.75% | 178.1bp (209.7bp) | CHA4/MA0283.1/Jaspar(0.769) More Information \| Similar Motifs Found | motif file (matrix) |
| 28 | A C G T A G T C A C G T G T A C G T C A A C T G A T G C A C G T | 1e-36 | -8.361e+01 | 3.97% | 0.88% | 159.5bp (656.6bp) | PDR8/MA0354.1/Jaspar(0.705) More Information \| Similar Motifs Found | motif file (matrix) |
| 29 | A T G C C G A T T C A G T G A C C G T A G T C A C G A T A T C G G C A T G T C A | 1e-36 | -8.320e+01 | 83.42% | 72.80% | 180.0bp (203.3bp) | TEAD1/MA0090.3/Jaspar(0.758) More Information \| Similar Motifs Found | motif file (matrix) |
| 30 | C G T A A G T C A C G T A C G T A G T C A G T C | 1e-35 | -8.081e+01 | 16.86% | 9.02% | 184.5bp (151.5bp) | ETV5/MA0765.2/Jaspar(0.938) More Information \| Similar Motifs Found | motif file (matrix) |
| 31 | A C T G A G T C A G T C C G T A A G T C A C T G | 1e-34 | -7.939e+01 | 81.48% | 70.76% | 174.4bp (194.0bp) | GBF2/MA1672.1/Jaspar(0.856) More Information \| Similar Motifs Found | motif file (matrix) |
| 32 | A G T C C G T A A C G T A C T G A C G T A C T G A G T C C G T A | 1e-27 | -6.374e+01 | 2.06% | 0.31% | 189.4bp (17.2bp) | MXI1/MA1108.2/Jaspar(0.864) More Information \| Similar Motifs Found | motif file (matrix) |
| 33 | C A T G A C G T C A G T T G A C T C G A G A C T G T A C C T A G C G T A A C T G | 1e-21 | -4.957e+01 | 92.74% | 86.72% | 172.7bp (153.3bp) | XBP1(MacIsaac)/Yeast(0.734) More Information \| Similar Motifs Found | motif file (matrix) |
| 34 | A C T G A C G T A G T C A C T G C G T A A G T C | 1e-19 | -4.405e+01 | 7.30% | 3.52% | 183.2bp (138.4bp) | CBF4(AP2EREBP)/colamp-CBF4-DAP-Seq(GSE60143)/Homer(0.813) More Information \| Similar Motifs Found | motif file (matrix) |
| 35 | A G T C A C G T A C G T A C G T A G T C C G T A A C G T C G T A | 1e-12 | -2.961e+01 | 2.10% | 0.67% | 162.6bp (18.6bp) | TATA-box/Drosophila-Promoters/Homer(0.761) More Information \| Similar Motifs Found | motif file (matrix) |
| 36 \* | A G T C A G T C C G T A A C G T A G T C A G T C C G T A A G T C | 1e-8 | -1.939e+01 | 2.62% | 1.24% | 139.0bp (36.0bp) | ZNF354C/MA0130.1/Jaspar(0.830) More Information \| Similar Motifs Found | motif file (matrix) |
| 37 \* | A G T C A G T C A G T C A G T C A G T C A G T C C G T A C G T A A G T C A G T C | 1e-8 | -1.903e+01 | 98.33% | 96.36% | 172.0bp (204.4bp) | ZNF740/MA0753.2/Jaspar(0.748) More Information \| Similar Motifs Found | motif file (matrix) |
