## Supplemental Table 10 for "The Establishment of Cell-Type Specific Gene Regulation in the Sea Urchin Embryo": ZGAPeaks_homerResults.html

/fast/users/jbrande\_m/work/scEarlyDev/bulkATAC//earlyMotifs//MZT/ - Homer de novo Motif Results


### Homer *de novo* Motif Results (/fast/users/jbrande\_m/work/scEarlyDev/bulkATAC//earlyMotifs//MZT/)

Known Motif Enrichment Results  
Gene Ontology Enrichment Results  
If Homer is having trouble matching a motif to a known motif, try copy/pasting the matrix file into
STAMP  
More information on motif finding results: HOMER
| Description of Results
| Tips
  
Total target sequences = 18254  
Total background sequences = 2521  
\* - possible false positive  

|  |  |  |  |  |  |  |  |  |
| --- | --- | --- | --- | --- | --- | --- | --- | --- |
| Rank | Motif | P-value | log P-pvalue | % of Targets | % of Background | STD(Bg STD) | Best Match/Details | Motif File |
| 1 | G A C T C G T A A T C G C G T A C G T A A C G T C G T A A C T G | 1e-191 | -4.399e+02 | 37.66% | 27.61% | 185.7bp (199.4bp) | br-Z2/dmmpmm(Bigfoot)/fly(0.824) More Information \| Similar Motifs Found | motif file (matrix) |
| 2 | G A T C G A T C G A T C G A T C G A T C G C A T A G C T A G C T A C T G G C A T | 1e-151 | -3.496e+02 | 38.67% | 29.56% | 192.3bp (191.9bp) | Sox3(HMG)/NPC-Sox3-ChIP-Seq(GSE33059)/Homer(0.801) More Information \| Similar Motifs Found | motif file (matrix) |
| 3 | A C G T C G A T A C T G A C T G C A G T A C T G G T A C A C G T A C G T A C T G | 1e-143 | -3.304e+02 | 3.69% | 1.19% | 192.4bp (165.4bp) | MZF1/MA0056.2/Jaspar(0.711) More Information \| Similar Motifs Found | motif file (matrix) |
| 4 | T C A G C T A G T C A G A C T G C T A G G T C A A C G T A C G T | 1e-129 | -2.972e+02 | 11.41% | 6.57% | 191.4bp (179.7bp) | MZF1/MA0056.2/Jaspar(0.842) More Information \| Similar Motifs Found | motif file (matrix) |
| 5 | G C T A A C G T A C T G C A T G C A T G A C G T A T G C C T A G C T A G C A G T | 1e-117 | -2.717e+02 | 3.34% | 1.14% | 189.7bp (165.0bp) | PB0167.1\_Sox13\_2/Jaspar(0.775) More Information \| Similar Motifs Found | motif file (matrix) |
| 6 | A T G C A G C T A G T C A G T C C G T A A T C G A C G T A C T G C T A G A C G T | 1e-116 | -2.672e+02 | 0.93% | 0.10% | 186.6bp (63.5bp) | tin/MA0247.2/Jaspar(0.744) More Information \| Similar Motifs Found | motif file (matrix) |
| 7 | C G T A C G T A A G T C A C G T A G T C A G T C A C G T C G T A | 1e-113 | -2.621e+02 | 0.72% | 0.03% | 197.3bp (0.0bp) | SUT2/MA0400.1/Jaspar(0.767) More Information \| Similar Motifs Found | motif file (matrix) |
| 8 | C G A T A C T G C G T A C G T A A C G T A T C G C G T A C G T A A C G T C G T A | 1e-107 | -2.476e+02 | 0.70% | 0.08% | 193.8bp (10.5bp) | PB0028.1\_Hbp1\_1/Jaspar(0.875) More Information \| Similar Motifs Found | motif file (matrix) |
| 9 | A C G T C A G T A C G T A C T G A C G T A G T C A T C G C G T A A G T C C G T A | 1e-106 | -2.457e+02 | 1.04% | 0.12% | 215.0bp (215.6bp) | ARF18/MA1689.1/Jaspar(0.745) More Information \| Similar Motifs Found | motif file (matrix) |
| 10 | G C A T C T G A G T C A C G A T A C T G A C T G G T C A A C G T A C G T G C T A | 1e-98 | -2.272e+02 | 2.76% | 0.91% | 211.1bp (166.4bp) | PAX7/MA0680.1/Jaspar(0.891) More Information \| Similar Motifs Found | motif file (matrix) |
| 11 | A C G T A T C G A C T G A C G T A C G T A C G T A G T C A G T C | 1e-96 | -2.221e+02 | 2.59% | 0.84% | 179.3bp (173.0bp) | che-1/MA0260.1/Jaspar(0.802) More Information \| Similar Motifs Found | motif file (matrix) |
| 12 | C G A T A C G T C G T A C A T G G A T C C T A G C T A G C G T A C G A T G A T C | 1e-92 | -2.137e+02 | 0.63% | 0.06% | 190.4bp (0.0bp) | GATA20/MA1324.1/Jaspar(0.689) More Information \| Similar Motifs Found | motif file (matrix) |
| 13 | C T G A C A T G C G A T T C G A C T A G A C T G | 1e-89 | -2.064e+02 | 47.24% | 39.90% | 189.6bp (184.3bp) | ZKSCAN1/MA1585.1/Jaspar(0.816) More Information \| Similar Motifs Found | motif file (matrix) |
| 14 | G A T C G T A C G C A T C G T A G T A C G C T A G C A T C A T G G C A T G C T A | 1e-81 | -1.874e+02 | 5.01% | 2.50% | 198.8bp (147.0bp) | PH0082.1\_Irx2/Jaspar(0.906) More Information \| Similar Motifs Found | motif file (matrix) |
| 15 | C G T A C T G A A C T G A G T C A C G T C G A T A C G T A G T C C G T A C G T A | 1e-78 | -1.810e+02 | 0.87% | 0.16% | 184.3bp (248.6bp) | PH0170.1\_Tgif2/Jaspar(0.799) More Information \| Similar Motifs Found | motif file (matrix) |
| 16 | A C G T A C G T A C G T C G T A C G T A C G T A C G T A A C G T A C T G A C G T | 1e-76 | -1.764e+02 | 1.18% | 0.27% | 167.1bp (230.3bp) | Lm\_0212(RRM)/Leishmania\_major-RNCMPT00212-PBM/HughesRNA(0.812) More Information \| Similar Motifs Found | motif file (matrix) |
| 17 | A G T C C T G A C T A G A C G T C T G A A C T G A G T C C G T A C G T A A C G T | 1e-75 | -1.734e+02 | 1.35% | 0.34% | 204.0bp (120.6bp) | odd/dmmpmm(Noyes)/fly(0.814) More Information \| Similar Motifs Found | motif file (matrix) |
| 18 | C G T A A C G T A C T G C T A G C G A T C G T A G T A C A G C T | 1e-74 | -1.704e+02 | 4.49% | 2.24% | 192.6bp (241.6bp) | HSF1/MA0319.1/Jaspar(0.712) More Information \| Similar Motifs Found | motif file (matrix) |
| 19 | T G A C C G T A C G A T A G T C C T A G C G A T G A C T C G T A A C T G C T A G | 1e-71 | -1.646e+02 | 5.71% | 3.15% | 192.4bp (218.1bp) | SOX14/MA1562.1/Jaspar(0.771) More Information \| Similar Motifs Found | motif file (matrix) |
| 20 | A C G T A T C G G C A T T G A C G A T C G T A C A G T C C G T A G A T C C T A G | 1e-69 | -1.602e+02 | 0.68% | 0.11% | 202.5bp (175.1bp) | YPR022C/MA0436.1/Jaspar(0.748) More Information \| Similar Motifs Found | motif file (matrix) |
| 21 | A C T G A G C T A T C G A G C T C T A G A G C T A C T G A C G T | 1e-68 | -1.584e+02 | 18.14% | 13.52% | 177.7bp (170.6bp) | BRUNOL5(RRM)/Homo\_sapiens-RNCMPT00166-PBM/HughesRNA(0.891) More Information \| Similar Motifs Found | motif file (matrix) |
| 22 | C G A T A C T G A G T C A G C T A G T C G T A C A C G T T A G C A G T C C T A G | 1e-68 | -1.581e+02 | 0.67% | 0.10% | 172.8bp (38.7bp) | SRSF1(RRM)/Homo\_sapiens-RNCMPT00163-PBM/HughesRNA(0.761) More Information \| Similar Motifs Found | motif file (matrix) |
| 23 | A C T G A C T G A C T G A G T C C G T A A C T G A C G T A G T C | 1e-63 | -1.469e+02 | 1.72% | 0.58% | 170.7bp (268.5bp) | THAP1/MA0597.1/Jaspar(0.786) More Information \| Similar Motifs Found | motif file (matrix) |
| 24 | C G T A C A T G C A T G C T A G A C T G G T C A C G A T A C T G C T G A A T G C | 1e-62 | -1.438e+02 | 0.97% | 0.20% | 190.5bp (446.3bp) | WRKY48/MA1088.1/Jaspar(0.722) More Information \| Similar Motifs Found | motif file (matrix) |
| 25 | A G C T A G T C C G A T A C G T A C G T A G T C A G C T A C G T A G T C A C G T | 1e-51 | -1.190e+02 | 1.79% | 0.71% | 182.9bp (244.1bp) | Tb\_0220(RRM)/Trypanosoma\_brucei-RNCMPT00220-PBM/HughesRNA(0.760) More Information \| Similar Motifs Found | motif file (matrix) |
| 26 | A G T C A C T G C G T A A C T G C G T A A G T C C G T A C G T A | 1e-49 | -1.137e+02 | 92.82% | 89.66% | 189.9bp (189.4bp) | ARF16/MA1688.1/Jaspar(0.784) More Information \| Similar Motifs Found | motif file (matrix) |
| 27 | A C G T G T A C A C G T C G T A C G T A C G T A | 1e-48 | -1.115e+02 | 45.81% | 40.46% | 190.8bp (195.1bp) | POL012.1\_TATA-Box/Jaspar(0.817) More Information \| Similar Motifs Found | motif file (matrix) |
| 28 | C G T A C G T A C G T A A C T G A G T C A C T G A C T G C G T A | 1e-41 | -9.583e+01 | 2.78% | 1.43% | 179.0bp (177.7bp) | RDR1/MA0360.1/Jaspar(0.766) More Information \| Similar Motifs Found | motif file (matrix) |
| 29 | A C T G A C G T A G T C A C T G A G T C A C T G A C T G C G T A | 1e-38 | -8.952e+01 | 78.41% | 74.26% | 187.5bp (181.2bp) | nit-4/MA1435.1/Jaspar(0.724) More Information \| Similar Motifs Found | motif file (matrix) |
| 30 | C G T A A C T G A C T G A G C T A G T C A G T C | 1e-35 | -8.286e+01 | 18.34% | 14.93% | 195.8bp (175.8bp) | TCP16(TCP)/colamp-TCP16-DAP-Seq(GSE60143)/Homer(0.746) More Information \| Similar Motifs Found | motif file (matrix) |
| 31 | A G C T A G T C C G T A A C T G C G T A A C T G | 1e-31 | -7.258e+01 | 49.94% | 45.60% | 189.0bp (197.3bp) | ZNF768(Zf)/Rajj-ZNF768-ChIP-Seq(GSE111879)/Homer(0.797) More Information \| Similar Motifs Found | motif file (matrix) |
| 32 | A C G T A G T C A C G T A G T C C G T A A C G T A C T G A G T C | 1e-29 | -6.832e+01 | 1.17% | 0.49% | 198.2bp (196.2bp) | ASD-1(RRM)/Caenorhabditis\_elegans-RNCMPT00180-PBM/HughesRNA(0.806) More Information \| Similar Motifs Found | motif file (matrix) |
| 33 | G A T C G A C T G C T A G A T C G T A C G T A C G A T C A G T C C A T G T C G A | 1e-29 | -6.742e+01 | 0.42% | 0.10% | 218.5bp (118.8bp) | ZBTB7B/MA0694.1/Jaspar(0.755) More Information \| Similar Motifs Found | motif file (matrix) |
| 34 | A C G T A C T G A G T C A G T C A G C T A C G T | 1e-25 | -5.937e+01 | 54.24% | 50.32% | 189.9bp (184.7bp) | MOT3/MA0340.1/Jaspar(0.860) More Information \| Similar Motifs Found | motif file (matrix) |
| 35 | A C T G C G T A A G T C A G T C C G T A A C T G A G C T A G T C | 1e-23 | -5.380e+01 | 2.30% | 1.38% | 193.2bp (213.5bp) | ACE2/ACE2\_YPD/2-SWI5(Harbison)/Yeast(0.713) More Information \| Similar Motifs Found | motif file (matrix) |
| 36 | A G T C A G T C A C T G A C G T A C G T C G T A | 1e-16 | -3.753e+01 | 8.78% | 7.16% | 191.4bp (185.5bp) | OVOL2/MA1545.1/Jaspar(0.955) More Information \| Similar Motifs Found | motif file (matrix) |
| 37 | A C G T A G T C A G T C A C G T A C T G A G T C A C T G A C G T | 1e-16 | -3.697e+01 | 0.66% | 0.29% | 203.2bp (94.0bp) | SOK2/SOK2\_BUT14/4-SUT1(Harbison)/Yeast(0.745) More Information \| Similar Motifs Found | motif file (matrix) |
| 38 \* | A G T C A C T G C G T A A G T C A C G T A C G T A C G T A C T G | 1e-9 | -2.226e+01 | 0.99% | 0.63% | 165.5bp (142.4bp) | HNF4G/MA0484.2/Jaspar(0.793) More Information \| Similar Motifs Found | motif file (matrix) |
| 39 \* | A C G T A C T G A C G T C G T A A C G T A C T G A C T G A G T C | 1e-7 | -1.658e+01 | 1.17% | 0.80% | 226.5bp (181.0bp) | Abd-B/dmmpmm(Bergman)/fly(0.760) More Information \| Similar Motifs Found | motif file (matrix) |
