## Supplemental Tabel 1 for "The Establishment of Cell-Type Specific Gene Regulation in the Sea Urchin Embryo": AboralNSM_homerResults.html

/data/gpfs-1/users/jbrande\_m/work/scEarlyDev/scATACseq\_analysis/peaksPerLineage/Motifs/AboralNSM/ - Homer de novo Motif Results


### Homer *de novo* Motif Results (/data/gpfs-1/users/jbrande\_m/work/scEarlyDev/scATACseq\_analysis/peaksPerLineage/Motifs/AboralNSM/)

Known Motif Enrichment Results  
Gene Ontology Enrichment Results  
If Homer is having trouble matching a motif to a known motif, try copy/pasting the matrix file into
STAMP  
More information on motif finding results: HOMER
| Description of Results
| Tips
  
Total target sequences = 4595  
Total background sequences = 5927  
\* - possible false positive  

|  |  |  |  |  |  |  |  |  |
| --- | --- | --- | --- | --- | --- | --- | --- | --- |
| Rank | Motif | P-value | log P-pvalue | % of Targets | % of Background | STD(Bg STD) | Best Match/Details | Motif File |
| 1 | C T A G T A C G G A C T T C G A A T G C A G T C G T C A A C T G A G T C G C T A | 1e-811 | -1.868e+03 | 46.96% | 11.05% | 1161.8bp (1525.0bp) | GCM1/MA0646.1/Jaspar(0.839) More Information \| Similar Motifs Found | motif file (matrix) |
| 2 | C G A T G C A T T C G A G C A T A C T G G A T C C A T G C T A G C A T G G A T C | 1e-66 | -1.529e+02 | 29.31% | 18.76% | 1399.5bp (1483.9bp) | gcm2/MA0917.1/Jaspar(0.811) More Information \| Similar Motifs Found | motif file (matrix) |
| 3 | A C T G A C T G A T C G A T C G A G T C A G T C A G T C T C A G C G A T G T C A | 1e-36 | -8.312e+01 | 0.65% | 0.03% | 1908.2bp (1395.1bp) | PLAGL2/MA1548.1/Jaspar(0.789) More Information \| Similar Motifs Found | motif file (matrix) |
| 4 | A C T G T A C G A C G T C G T A T C G A A C G T C G T A C G T A | 1e-21 | -4.863e+01 | 34.21% | 27.77% | 1393.5bp (1413.8bp) | MEX-5(Znf)/Caenorhabditis\_elegans-RNCMPT00039-PBM/HughesRNA(0.812) More Information \| Similar Motifs Found | motif file (matrix) |
| 5 | C G T A C G T A A G T C A G T C A G T C T G C A A C T G A C G T A C G T C G T A | 1e-18 | -4.149e+01 | 0.63% | 0.07% | 1087.9bp (1991.8bp) | grh/MA1457.1/Jaspar(0.720) More Information \| Similar Motifs Found | motif file (matrix) |
| 6 | A C G T C T A G A G T C G C T A A C G T A C T G C G A T C T G A T A G C G T C A | 1e-17 | -3.938e+01 | 37.37% | 31.44% | 1318.3bp (1735.5bp) | FUS3/MA0565.2/Jaspar(0.819) More Information \| Similar Motifs Found | motif file (matrix) |
| 7 | G T C A T A C G A C T G G A C T G T C A G T C A A T C G A T C G C T A G A G C T | 1e-15 | -3.486e+01 | 0.57% | 0.08% | 1135.6bp (1425.7bp) | SFP1/SFP1\_SM/50-RAP1(Harbison)/Yeast(0.745) More Information \| Similar Motifs Found | motif file (matrix) |
| 8 | C G T A A C T G A C G T C G T A C G T A A C G T A C T G A G T C C G T A A C T G | 1e-12 | -2.921e+01 | 0.70% | 0.15% | 951.3bp (880.7bp) | TEAD1(TEAD)/HepG2-TEAD1-ChIP-Seq(Encode)/Homer(0.672) More Information \| Similar Motifs Found | motif file (matrix) |
| 9 \* | A C T G A C G T C G T A A C G T A G T C C T G A C G T A A G T C A C G T A G T C | 1e-11 | -2.601e+01 | 0.52% | 0.09% | 838.9bp (2610.3bp) | so/MA0246.1/Jaspar(0.757) More Information \| Similar Motifs Found | motif file (matrix) |
| 10 \* | A G T C C G T A C G T A A C T G C G T A A G T C A G T C A G T C A C G T A G T C | 1e-10 | -2.459e+01 | 0.46% | 0.07% | 672.8bp (306.5bp) | LRF(Zf)/Erythroblasts-ZBTB7A-ChIP-Seq(GSE74977)/Homer(0.759) More Information \| Similar Motifs Found | motif file (matrix) |
| 11 \* | C G T A A C G T C G T A A C G T A C T G C T G A A C G T C G T A | 1e-10 | -2.420e+01 | 31.40% | 27.04% | 1389.8bp (1508.4bp) | CRC(C2C2YABBY)/col-CRC-DAP-Seq(GSE60143)/Homer(0.777) More Information \| Similar Motifs Found | motif file (matrix) |
| 12 \* | A C T G A G C T A C T G A G T C A G T C A G T C A C G T A T C G | 1e-10 | -2.394e+01 | 5.44% | 3.53% | 1395.2bp (1498.1bp) | PHD1(MacIsaac)/Yeast(0.778) More Information \| Similar Motifs Found | motif file (matrix) |
| 13 \* | C G T A C G T A A C T G C T G A A T C G A C G T C G T A A G T C A G C T A C T G | 1e-10 | -2.332e+01 | 2.66% | 1.40% | 1451.8bp (1473.2bp) | OSR1/MA1542.1/Jaspar(0.649) More Information \| Similar Motifs Found | motif file (matrix) |
| 14 \* | A G T C A C T G A G T C C G A T A G T C A C T G A T C G A C G T | 1e-9 | -2.084e+01 | 6.81% | 4.80% | 1294.5bp (1454.9bp) | PB0140.1\_Irf6\_2/Jaspar(0.689) More Information \| Similar Motifs Found | motif file (matrix) |
| 15 \* | A G T C A C G T A G T C C G T A A C G T A C G T A G T C A G T C A C G T A C G T | 1e-8 | -1.991e+01 | 0.67% | 0.20% | 1236.8bp (388.2bp) | TEC1/TEC1\_YPD/[](Harbison)/Yeast(0.782) More Information \| Similar Motifs Found | motif file (matrix) |
| 16 \* | A G T C A G T C A G T C A G T C A T G C A G T C A G T C A C G T | 1e-8 | -1.874e+01 | 20.24% | 17.02% | 1345.1bp (1478.8bp) | VEZF1/MA1578.1/Jaspar(0.894) More Information \| Similar Motifs Found | motif file (matrix) |
| 17 \* | A C G T C T G A C T G A A C G T A C T G A C G T A C T G A G T C | 1e-7 | -1.798e+01 | 11.32% | 8.90% | 1325.3bp (1600.3bp) | MEC-8(RRM)/Caenorhabditis\_elegans-RNCMPT00181-PBM/HughesRNA(0.806) More Information \| Similar Motifs Found | motif file (matrix) |
| 18 \* | A C G T C G T A A G T C A C T G A C T G A G T C G T C A A C G T | 1e-7 | -1.773e+01 | 2.00% | 1.07% | 879.3bp (1200.9bp) | SKO1(MacIsaac)/Yeast(0.781) More Information \| Similar Motifs Found | motif file (matrix) |
| 19 \* | A C T G C T G A C T A G C G T A A C G T A C G T A G T C A C T G | 1e-7 | -1.757e+01 | 7.62% | 5.67% | 1332.9bp (1573.5bp) | ARR14/MA0947.1/Jaspar(0.841) More Information \| Similar Motifs Found | motif file (matrix) |
| 20 \* | C G T A C G T A A C G T A C G T A C G T A C G T | 1e-7 | -1.747e+01 | 66.05% | 62.18% | 1407.8bp (1490.9bp) | Ng\_0261(RRM)/Naegleria\_gruberi-RNCMPT00261-PBM/HughesRNA(0.893) More Information \| Similar Motifs Found | motif file (matrix) |
| 21 \* | C A T G C G T A C A T G T C G A C A T G C T G A C T A G A G T C C T A G G C T A | 1e-7 | -1.624e+01 | 9.27% | 7.19% | 1319.1bp (1276.4bp) | SeqBias: GA-repeat(0.873) More Information \| Similar Motifs Found | motif file (matrix) |
| 22 \* | A C G T G T A C A G T C A C T G A C G T G T A C A C T G A C T G | 1e-6 | -1.437e+01 | 2.72% | 1.72% | 1333.9bp (1356.0bp) | AT3G60490(AP2EREBP)/colamp-AT3G60490-DAP-Seq(GSE60143)/Homer(0.731) More Information \| Similar Motifs Found | motif file (matrix) |
| 23 \* | A C T G A C G T C G T A A C G T C G T A C G T A C G T A A G T C A C T G A C G T | 1e-3 | -8.998e+00 | 0.41% | 0.15% | 1376.4bp (1930.1bp) | KHDRBS3(KH)/Homo\_sapiens-RNCMPT00034-PBM/HughesRNA(0.726) More Information \| Similar Motifs Found | motif file (matrix) |
| 24 \* | A C T G A C T G C G T A A C G T A C T G A C T G C G T A A C T G A C T G C G T A | 1e-1 | -2.354e+00 | 0.26% | 0.18% | 998.3bp (721.8bp) | PF10\_0068(RRM)/Plasmodium\_falciparum-RNCMPT00199-PBM/HughesRNA(0.760) More Information \| Similar Motifs Found | motif file (matrix) |
