## Supplemental Tabel 1 for "The Establishment of Cell-Type Specific Gene Regulation in the Sea Urchin Embryo": EctoAboral_homerResults.html

/data/gpfs-1/users/jbrande\_m/work/scEarlyDev/scATACseq\_analysis/peaksPerLineage/Motifs/EctoAboral/ - Homer de novo Motif Results


### Homer *de novo* Motif Results (/data/gpfs-1/users/jbrande\_m/work/scEarlyDev/scATACseq\_analysis/peaksPerLineage/Motifs/EctoAboral/)

Known Motif Enrichment Results  
Gene Ontology Enrichment Results  
If Homer is having trouble matching a motif to a known motif, try copy/pasting the matrix file into
STAMP  
More information on motif finding results: HOMER
| Description of Results
| Tips
  
Total target sequences = 159  
Total background sequences = 9167  
\* - possible false positive  

|  |  |  |  |  |  |  |  |  |
| --- | --- | --- | --- | --- | --- | --- | --- | --- |
| Rank | Motif | P-value | log P-pvalue | % of Targets | % of Background | STD(Bg STD) | Best Match/Details | Motif File |
| 1 | A C G T A G T C A G T C C G T A A C T G A C G T A C T G C G T A A C G T A G T C | 1e-50 | -1.164e+02 | 18.87% | 0.18% | 1134.8bp (1697.4bp) | PB0195.1\_Zbtb3\_2/Jaspar(0.696) More Information \| Similar Motifs Found | motif file (matrix) |
| 2 | C G T A A C G T A G T C A C G T A C T G C G T A A C T G A C T G A C T G A G T C | 1e-50 | -1.154e+02 | 18.24% | 0.16% | 1373.3bp (5398.7bp) | THAP1/MA0597.1/Jaspar(0.647) More Information \| Similar Motifs Found | motif file (matrix) |
| 3 | A C T G A C G T A C G T C G T A A C T G C G T A C G T A A C T G A G T C A C G T | 1e-46 | -1.067e+02 | 15.72% | 0.11% | 1556.9bp (2181.7bp) | SFL1(MacIsaac)/Yeast(0.671) More Information \| Similar Motifs Found | motif file (matrix) |
| 4 | A C G T A C T G A G T C C T G A A G T C A C G T A C T G A G T C A G T C A G T C | 1e-45 | -1.050e+02 | 17.61% | 0.19% | 1611.4bp (1574.3bp) | THAP1/MA0597.1/Jaspar(0.694) More Information \| Similar Motifs Found | motif file (matrix) |
| 5 | A C T G C G T A C G T A G T C A A G T C A C T G A G T C C G T A A G T C A C G T | 1e-41 | -9.570e+01 | 15.72% | 0.16% | 1159.0bp (1138.4bp) | CST6(MacIsaac)/Yeast(0.647) More Information \| Similar Motifs Found | motif file (matrix) |
| 6 | A C T G C G T A A G T C A C G T A C G T A C G T A G T C C G T A C G T A A C T G | 1e-40 | -9.217e+01 | 17.61% | 0.30% | 1296.2bp (1204.2bp) | vnd/dmmpmm(Noyes\_hd)/fly(0.712) More Information \| Similar Motifs Found | motif file (matrix) |
| 7 | C G T A A C T G C T A G A G T C C G T A A C T G A G T C C G T A C T G A C T A G | 1e-38 | -8.906e+01 | 20.13% | 0.57% | 1644.2bp (5039.1bp) | ARF34/MA1693.1/Jaspar(0.774) More Information \| Similar Motifs Found | motif file (matrix) |
| 8 | C T G A C G T A C G T A A C T G C G T A A C T G A T G C C G T A A G T C C G T A | 1e-35 | -8.250e+01 | 19.50% | 0.62% | 1553.5bp (3805.6bp) | NCU08034(RRM)/Neurospora\_crassa-RNCMPT00209-PBM/HughesRNA(0.725) More Information \| Similar Motifs Found | motif file (matrix) |
| 9 | A G T C A C G T A C G T A G C T A G T C A C G T A C G T A G T C A C T G A G T C | 1e-35 | -8.095e+01 | 16.35% | 0.33% | 1693.0bp (1242.5bp) | Tb\_0220(RRM)/Trypanosoma\_brucei-RNCMPT00220-PBM/HughesRNA(0.794) More Information \| Similar Motifs Found | motif file (matrix) |
| 10 | C G T A A C G T A C G T C G T A A C T G C G T A A C T G A C G T A C G T A C G T | 1e-35 | -8.069e+01 | 18.24% | 0.51% | 1889.2bp (1805.8bp) | AT1G72740/MA1353.1/Jaspar(0.785) More Information \| Similar Motifs Found | motif file (matrix) |
| 11 | A G T C A G T C C G T A A C G T A G T C A G T C A G T C A C G T A C G T A C T G | 1e-34 | -7.911e+01 | 15.09% | 0.26% | 806.3bp (10164.9bp) | HNRNPK(KH)/Homo\_sapiens-RNCMPT00026-PBM/HughesRNA(0.721) More Information \| Similar Motifs Found | motif file (matrix) |
| 12 | C G T A A C G T A C T G C G T A A G C T A C T G A G T C A G T C A G T C C G T A | 1e-34 | -7.873e+01 | 17.61% | 0.48% | 811.2bp (1786.3bp) | AGL42/MA1201.1/Jaspar(0.752) More Information \| Similar Motifs Found | motif file (matrix) |
| 13 | C G T A A C G T A C G T A G T C A G T C C G T A A C G T A G T C A C G T A G T C | 1e-32 | -7.554e+01 | 16.98% | 0.48% | 1746.3bp (7275.0bp) | Pp\_0237(RRM)/Physcomitrella\_patens-RNCMPT00237-PBM/HughesRNA(0.726) More Information \| Similar Motifs Found | motif file (matrix) |
| 14 | C A G T A G T C C G A T C T G A T C G A C T G A T C G A A G C T A C T G C T A G | 1e-29 | -6.691e+01 | 25.79% | 2.45% | 1505.3bp (6760.3bp) | Lm\_0212(RRM)/Leishmania\_major-RNCMPT00212-PBM/HughesRNA(0.745) More Information \| Similar Motifs Found | motif file (matrix) |
| 15 | C G T A C G T A A C G T C G T A C G T A A G T C A G T C A C G T A G T C A C G T | 1e-27 | -6.220e+01 | 13.84% | 0.37% | 1679.3bp (7057.5bp) | NR1H4/MA1110.1/Jaspar(0.697) More Information \| Similar Motifs Found | motif file (matrix) |
| 16 | A G T C A G T C A G T C A C G T A C G T A C G T A G T C C G T A A C G T A C G T | 1e-26 | -6.167e+01 | 16.98% | 0.79% | 1823.9bp (5440.7bp) | Hoxa10(Homeobox)/ChickenMSG-Hoxa10.Flag-ChIP-Seq(GSE86088)/Homer(0.743) More Information \| Similar Motifs Found | motif file (matrix) |
| 17 | A G T C A C G T A C T G A C T G A C T G A C T G A G T C A C G T A G T C A G T C | 1e-26 | -6.015e+01 | 8.81% | 0.06% | 401.6bp (1796.2bp) | ZNF519(Zf)/HEK293-ZNF519.GFP-ChIP-Seq(GSE58341)/Homer(0.743) More Information \| Similar Motifs Found | motif file (matrix) |
| 18 | A C G T A C G T A G T C A C G T A G T C C G A T A G T C A G C T A C G T C G T A | 1e-25 | -5.843e+01 | 20.75% | 1.68% | 1568.9bp (8210.8bp) | Trl/MA0205.2/Jaspar(0.824) More Information \| Similar Motifs Found | motif file (matrix) |
| 19 | A C T G C G T A A G T C A G T C C G T A C T A G C G T A C G T A | 1e-25 | -5.798e+01 | 50.31% | 14.78% | 1496.0bp (15326.1bp) | ACE2/ACE2\_YPD/2-SWI5(Harbison)/Yeast(0.722) More Information \| Similar Motifs Found | motif file (matrix) |
| 20 | A C T G C G T A A C G T A C G T A G T C A C G T A C G T A C T G A G T C A C G T | 1e-25 | -5.758e+01 | 10.69% | 0.17% | 1749.9bp (864.7bp) | NAC078/MA1677.1/Jaspar(0.702) More Information \| Similar Motifs Found | motif file (matrix) |
| 21 | C G T A C G T A A C T G C G T A A G T C A C G T A G T C A G T C A G T C A C T G | 1e-24 | -5.664e+01 | 9.43% | 0.10% | 2259.2bp (15499.3bp) | PB0203.1\_Zfp691\_2/Jaspar(0.704) More Information \| Similar Motifs Found | motif file (matrix) |
| 22 | A G T C C G T A A G C T A C G T A T C G C G T A C T G A A C G T A C T G A C G T | 1e-23 | -5.491e+01 | 86.16% | 47.33% | 1611.2bp (12636.7bp) | HDG7(HB)/col-HDG7-DAP-Seq(GSE60143)/Homer(0.772) More Information \| Similar Motifs Found | motif file (matrix) |
| 23 | C G T A A G T C A C G T A G T C A C G T A G T C A C G T A G T C A G T C C G T A | 1e-22 | -5.279e+01 | 11.95% | 0.34% | 1682.6bp (12303.2bp) | Trl(Zf)/S2-GAGAfactor-ChIP-Seq(GSE40646)/Homer(0.809) More Information \| Similar Motifs Found | motif file (matrix) |
| 24 | C T G A A C G T A C G T A C G T T C A G C G T A G T C A A C G T | 1e-22 | -5.118e+01 | 61.01% | 24.14% | 1715.0bp (14080.6bp) | AT2G20110(CPP)/colamp-AT2G20110-DAP-Seq(GSE60143)/Homer(0.869) More Information \| Similar Motifs Found | motif file (matrix) |
| 25 | C G T A A C T G C G T A C G T A C G T A A C G T A G C T A C T G | 1e-21 | -4.997e+01 | 44.03% | 12.71% | 1973.5bp (14861.6bp) | PABPC5(RRM)/Homo\_sapiens-RNCMPT00171-PBM/HughesRNA(0.830) More Information \| Similar Motifs Found | motif file (matrix) |
| 26 | A T G C A G T C A C T G A C T G G T A C A T G C | 1e-21 | -4.889e+01 | 86.16% | 49.83% | 1792.3bp (12496.4bp) | RDS1/MA0361.1/Jaspar(0.850) More Information \| Similar Motifs Found | motif file (matrix) |
| 27 | A C T G A C T G C G T A A G T C C G T A A G T C C G T A C G T A | 1e-16 | -3.871e+01 | 84.91% | 52.93% | 1644.6bp (11658.8bp) | HNRNPL(RRM)/Homo\_sapiens-RNCMPT00091-PBM/HughesRNA(0.813) More Information \| Similar Motifs Found | motif file (matrix) |
| 28 | A C T G C G T A A G T C A C T G C G T A A G T C A C G T C G T A | 1e-16 | -3.803e+01 | 13.84% | 1.16% | 1352.4bp (11722.0bp) | RBM45(RRM)/Homo\_sapiens-RNCMPT00241-PBM/HughesRNA(0.789) More Information \| Similar Motifs Found | motif file (matrix) |
| 29 | C G T A C G T A A C T G A C T G A G T C A C G T A G T C A C G T | 1e-14 | -3.288e+01 | 76.10% | 45.67% | 1833.9bp (14146.3bp) | MYB88(MYB)/col-MYB88-DAP-Seq(GSE60143)/Homer(0.677) More Information \| Similar Motifs Found | motif file (matrix) |
