## Supplemental Tabel 1 for "The Establishment of Cell-Type Specific Gene Regulation in the Sea Urchin Embryo": EctoApical_homerResults.html

/data/gpfs-1/users/jbrande\_m/work/scEarlyDev/scATACseq\_analysis/peaksPerLineage/Motifs/EctoApical/ - Homer de novo Motif Results


### Homer *de novo* Motif Results (/data/gpfs-1/users/jbrande\_m/work/scEarlyDev/scATACseq\_analysis/peaksPerLineage/Motifs/EctoApical/)

Known Motif Enrichment Results  
Gene Ontology Enrichment Results  
If Homer is having trouble matching a motif to a known motif, try copy/pasting the matrix file into
STAMP  
More information on motif finding results: HOMER
| Description of Results
| Tips
  
Total target sequences = 368  
Total background sequences = 8507  
\* - possible false positive  

|  |  |  |  |  |  |  |  |  |
| --- | --- | --- | --- | --- | --- | --- | --- | --- |
| Rank | Motif | P-value | log P-pvalue | % of Targets | % of Background | STD(Bg STD) | Best Match/Details | Motif File |
| 1 | A T G C T A G C G A C T T C A G G A C T A T G C C G T A G C T A C G A T G A T C | 1e-32 | -7.494e+01 | 42.39% | 16.01% | 1083.3bp (10421.9bp) | Pbx3(Homeobox)/GM12878-PBX3-ChIP-Seq(GSE32465)/Homer(0.910) More Information \| Similar Motifs Found | motif file (matrix) |
| 2 | C T A G C G T A A G T C C G T A C G T A A C G T C T G A A T C G | 1e-30 | -7.042e+01 | 50.82% | 22.93% | 1116.8bp (10445.5bp) | Sox7(HMG)/ESC-Sox7-ChIP-Seq(GSE133899)/Homer(0.884) More Information \| Similar Motifs Found | motif file (matrix) |
| 3 | C T A G T A G C T G A C A C G T G T A C C G T A A C G T G A C T C G T A C T A G | 1e-21 | -4.895e+01 | 26.36% | 9.10% | 1201.6bp (14434.9bp) | zen/MA0256.1/Jaspar(0.910) More Information \| Similar Motifs Found | motif file (matrix) |
| 4 | T A G C G T C A A G T C G A C T A G C T C A G T A G T C G T A C G C T A A G C T | 1e-18 | -4.153e+01 | 58.42% | 35.79% | 1429.0bp (11002.3bp) | NFAT(RHD)/Jurkat-NFATC1-ChIP-Seq(Jolma\_et\_al.)/Homer(0.731) More Information \| Similar Motifs Found | motif file (matrix) |
| 5 | C G A T C T A G A C T G A C T G A G T C A C G T A C G T A G T C | 1e-17 | -4.111e+01 | 41.85% | 21.48% | 1222.7bp (9616.2bp) | che-1/MA0260.1/Jaspar(0.871) More Information \| Similar Motifs Found | motif file (matrix) |
| 6 | C G A T C A T G C G T A A C G T A C T G A G T C A C G T A T G C A C G T G T A C | 1e-16 | -3.787e+01 | 68.75% | 47.06% | 1316.0bp (10155.6bp) | At3g24120(G2like)/col-At3g24120-DAP-Seq(GSE60143)/Homer(0.697) More Information \| Similar Motifs Found | motif file (matrix) |
| 7 | C G A T A T G C A C T G T A G C G C A T C T A G C T G A C A G T C T A G T A C G | 1e-15 | -3.668e+01 | 51.90% | 31.15% | 1273.7bp (10285.6bp) | RBP1-LIKE(RRM)/Drosophila\_melanogaster-RNCMPT00127-PBM/HughesRNA(0.756) More Information \| Similar Motifs Found | motif file (matrix) |
| 8 | T C A G C T A G G T C A A C G T A C G T T C G A T C A G A G T C | 1e-15 | -3.566e+01 | 71.74% | 50.92% | 1279.5bp (10375.0bp) | CRX(Homeobox)/Retina-Crx-ChIP-Seq(GSE20012)/Homer(0.968) More Information \| Similar Motifs Found | motif file (matrix) |
| 9 | C G T A G A C T G T C A C G T A A G T C A G T C A C T G C G T A | 1e-15 | -3.559e+01 | 55.43% | 34.69% | 1354.6bp (10855.9bp) | MYB33/MA1391.2/Jaspar(0.764) More Information \| Similar Motifs Found | motif file (matrix) |
| 10 | A G T C G T A C C T G A G T A C A G C T C G A T C T G A A T C G | 1e-15 | -3.540e+01 | 49.46% | 29.33% | 1257.2bp (11386.8bp) | Nkx3-1/MA0124.2/Jaspar(0.856) More Information \| Similar Motifs Found | motif file (matrix) |
| 11 | C G T A C G T A A G C T A C T G A C T G C G T A A C T G A C T G C G T A A G T C | 1e-14 | -3.299e+01 | 73.37% | 53.56% | 1423.8bp (11141.8bp) | SRSF1(RRM)/Homo\_sapiens-RNCMPT00106-PBM/HughesRNA(0.781) More Information \| Similar Motifs Found | motif file (matrix) |
| 12 | C T G A T A C G T A G C T G C A G A T C G T A C A G T C T G A C G A C T G A T C | 1e-13 | -3.112e+01 | 50.00% | 31.06% | 1259.8bp (11132.3bp) | AFT2/AFT2\_H2O2Lo/10-RCS1[~AFT2](Harbison)/Yeast(0.753) More Information \| Similar Motifs Found | motif file (matrix) |
| 13 \* | T G A C C A T G G A C T A G T C T C A G A C G T A T G C T A C G | 1e-11 | -2.747e+01 | 60.60% | 42.27% | 1407.0bp (9472.3bp) | SRSF7(RRM,Znf)/Homo\_sapiens-RNCMPT00073-PBM/HughesRNA(0.911) More Information \| Similar Motifs Found | motif file (matrix) |
| 14 \* | A C G T A C G T C G T A C G T A A C T G C G T A | 1e-11 | -2.617e+01 | 63.32% | 45.46% | 1282.1bp (10922.0bp) | ZCRB1(RRM)/Homo\_sapiens-RNCMPT00087-PBM/HughesRNA(0.785) More Information \| Similar Motifs Found | motif file (matrix) |
| 15 \* | A G C T A C G T G T C A C T G A A C G T A G T C A C G T A G T C | 1e-11 | -2.555e+01 | 40.49% | 24.41% | 1653.6bp (10806.6bp) | Ptx1/dmmpmm(Noyes\_hd)/fly(0.766) More Information \| Similar Motifs Found | motif file (matrix) |
| 16 \* | T C G A G T C A C A T G A T C G C T G A A G T C A T G C C G A T A C T G C A T G | 1e-10 | -2.463e+01 | 10.05% | 2.74% | 1304.7bp (12277.1bp) | ttk/dmmpmm(Pollard)/fly(0.748) More Information \| Similar Motifs Found | motif file (matrix) |
| 17 \* | A G T C G T A C A G T C G T A C A G C T A C T G A G T C G A T C T C A G A C G T | 1e-10 | -2.358e+01 | 8.15% | 1.92% | 1149.4bp (8221.5bp) | Unknown-ESC-element(?)/mES-Nanog-ChIP-Seq(GSE11724)/Homer(0.787) More Information \| Similar Motifs Found | motif file (matrix) |
| 18 \* | C G T A A C G T C G T A C G T A A C T G A C T G C G T A A C G T A C T G C G T A | 1e-9 | -2.286e+01 | 82.61% | 67.84% | 1308.9bp (11334.2bp) | G3BP2(RRM)/Homo\_sapiens-RNCMPT00021-PBM/HughesRNA(0.784) More Information \| Similar Motifs Found | motif file (matrix) |
| 19 \* | A C T G C A T G A T G C A C T G A C T G A C G T A G T C C A G T C G T A A C G T | 1e-9 | -2.280e+01 | 4.89% | 0.68% | 642.9bp (7696.6bp) | Run/dmmpmm(Papatsenko)/fly(0.666) More Information \| Similar Motifs Found | motif file (matrix) |
| 20 \* | A G C T C G T A G T A C A G T C C T G A A T G C A G C T G C T A G T C A A G C T | 1e-9 | -2.192e+01 | 61.14% | 44.93% | 1253.0bp (12394.5bp) | schlank/MA0193.1/Jaspar(0.711) More Information \| Similar Motifs Found | motif file (matrix) |
| 21 \* | C G T A A T C G C G T A G C A T A C T G A G T C A G T C C G A T A T C G A C T G | 1e-6 | -1.560e+01 | 3.53% | 0.55% | 1503.7bp (3536.7bp) | ZFX(Zf)/mES-Zfx-ChIP-Seq(GSE11431)/Homer(0.654) More Information \| Similar Motifs Found | motif file (matrix) |
| 22 \* | C G A T A C T G A C G T G T C A A C T G G T C A C G T A C G T A C G T A A C G T | 1e-6 | -1.461e+01 | 11.14% | 4.73% | 1629.1bp (9704.6bp) | RLR1/RLR1\_YPD/[](Harbison)/Yeast(0.806) More Information \| Similar Motifs Found | motif file (matrix) |
| 23 \* | A C G T G T C A A C T G A C G T C G T A A C G T A C G T A G T C G T C A A G T C | 1e-6 | -1.387e+01 | 76.09% | 64.41% | 1399.2bp (11152.4bp) | vvl/dmmpmm(Pollard)/fly(0.714) More Information \| Similar Motifs Found | motif file (matrix) |
