## Supplemental Tabel 1 for "The Establishment of Cell-Type Specific Gene Regulation in the Sea Urchin Embryo": EctoCiliaryBand_homerResults.html

/data/gpfs-1/users/jbrande\_m/work/scEarlyDev/scATACseq\_analysis/peaksPerLineage/Motifs/EctoCiliaryBand/ - Homer de novo Motif Results


### Homer *de novo* Motif Results (/data/gpfs-1/users/jbrande\_m/work/scEarlyDev/scATACseq\_analysis/peaksPerLineage/Motifs/EctoCiliaryBand/)

Known Motif Enrichment Results  
Gene Ontology Enrichment Results  
If Homer is having trouble matching a motif to a known motif, try copy/pasting the matrix file into
STAMP  
More information on motif finding results: HOMER
| Description of Results
| Tips
  
Total target sequences = 46  
Total background sequences = 8008  
\* - possible false positive  

|  |  |  |  |  |  |  |  |  |
| --- | --- | --- | --- | --- | --- | --- | --- | --- |
| Rank | Motif | P-value | log P-pvalue | % of Targets | % of Background | STD(Bg STD) | Best Match/Details | Motif File |
| 1 | G T A C C T G A G T C A C G A T A T G C C T G A C G T A A G C T T C G A C G A T | 1e-20 | -4.682e+01 | 69.57% | 10.72% | 518.8bp (12253.0bp) | ONECUT1/MA0679.2/Jaspar(0.896) More Information \| Similar Motifs Found | motif file (matrix) |
| 2 \* | G C T A C G T A C G A T C G A T C T A G C G T A C A G T G A T C C G T A C T G A | 1e-6 | -1.475e+01 | 67.39% | 30.92% | 1033.1bp (12540.6bp) | CUX1/MA0754.1/Jaspar(0.802) More Information \| Similar Motifs Found | motif file (matrix) |
| 3 \* | A C G T A G T C C A T G G C T A A C G T A C G T C T G A A C G T A C G T T C A G | 1e-6 | -1.437e+01 | 32.61% | 7.37% | 596.4bp (6521.3bp) | Tv\_0226(RRM)/Trichomonas\_vaginalis-RNCMPT00226-PBM/HughesRNA(0.873) More Information \| Similar Motifs Found | motif file (matrix) |
| 4 \* | A C T G A C T G A G T C A C G T A C T G A C G T A G T C A G T C A G T C A C G T | 1e-5 | -1.253e+01 | 6.52% | 0.07% | 75.1bp (823.1bp) | ZNF341(Zf)/EBV-ZNF341-ChIP-Seq(GSE113194)/Homer(0.830) More Information \| Similar Motifs Found | motif file (matrix) |
| 5 \* | C G T A A C T G A C T G C G T A A C G T A C T G A C T G A C T G | 1e-5 | -1.205e+01 | 21.74% | 3.72% | 757.6bp (14586.4bp) | G3BP2(RRM)/Homo\_sapiens-RNCMPT00021-PBM/HughesRNA(0.802) More Information \| Similar Motifs Found | motif file (matrix) |
| 6 \* | T G A C A C T G C G T A C T G A A C G T G C T A A C T G C G T A | 1e-5 | -1.165e+01 | 47.83% | 18.98% | 1493.7bp (11796.5bp) | RBM28(RRM)/Homo\_sapiens-RNCMPT00049-PBM/HughesRNA(0.772) More Information \| Similar Motifs Found | motif file (matrix) |
| 7 \* | G T C A A G T C A C G T G T A C C G A T A G T C G C T A A G T C A C G T A G T C | 1e-5 | -1.159e+01 | 30.43% | 8.04% | 1532.8bp (11073.4bp) | GAGA-repeat/Arabidopsis-Promoters/Homer(0.824) More Information \| Similar Motifs Found | motif file (matrix) |
| 8 \* | A C G T A C G T A G T C A C T G C G T A A C G T | 1e-4 | -9.279e+00 | 63.04% | 34.92% | 1782.5bp (10549.6bp) | OPI1/Literature(Harbison)/Yeast(0.811) More Information \| Similar Motifs Found | motif file (matrix) |
| 9 \* | A C G T A G T C A C G T A C G T A T G C C A G T A C G T A T C G A C T G A C G T | 1e-3 | -7.703e+00 | 8.70% | 0.78% | 200.6bp (11757.6bp) | PABPN1(RRM)/Homo\_sapiens-RNCMPT00157-PBM/HughesRNA(0.692) More Information \| Similar Motifs Found | motif file (matrix) |
