## Supplemental Tabel 1 for "The Establishment of Cell-Type Specific Gene Regulation in the Sea Urchin Embryo": EctoOral_homerResults.html

/data/gpfs-1/users/jbrande\_m/work/scEarlyDev/scATACseq\_analysis/peaksPerLineage/Motifs/EctoOral/ - Homer de novo Motif Results


### Homer *de novo* Motif Results (/data/gpfs-1/users/jbrande\_m/work/scEarlyDev/scATACseq\_analysis/peaksPerLineage/Motifs/EctoOral/)

Known Motif Enrichment Results  
Gene Ontology Enrichment Results  
If Homer is having trouble matching a motif to a known motif, try copy/pasting the matrix file into
STAMP  
More information on motif finding results: HOMER
| Description of Results
| Tips
  
Total target sequences = 247  
Total background sequences = 6908  
\* - possible false positive  

|  |  |  |  |  |  |  |  |  |
| --- | --- | --- | --- | --- | --- | --- | --- | --- |
| Rank | Motif | P-value | log P-pvalue | % of Targets | % of Background | STD(Bg STD) | Best Match/Details | Motif File |
| 1 | C A G T C A G T T G A C C A G T C A G T T G C A G C T A C G T A C G A T T A C G | 1e-31 | -7.189e+01 | 65.99% | 29.69% | 1447.6bp (14340.7bp) | BH1/dmmpmm(Noyes\_hd)/fly(0.754) More Information \| Similar Motifs Found | motif file (matrix) |
| 2 | A C T G C T G A G C A T A G C T C G T A A C G T G A T C C A T G T A C G T C A G | 1e-30 | -7.027e+01 | 87.04% | 52.29% | 1478.2bp (13574.0bp) | DPRX/MA1480.1/Jaspar(0.803) More Information \| Similar Motifs Found | motif file (matrix) |
| 3 | A G T C A G T C C G A T A C G T A C G T A G T C C G T A C G T A | 1e-29 | -6.891e+01 | 64.78% | 29.32% | 1417.4bp (11814.0bp) | At2g41835(C2H2)/col-At2g41835-DAP-Seq(GSE60143)/Homer(0.766) More Information \| Similar Motifs Found | motif file (matrix) |
| 4 | A T C G A T C G A G C T A C G T A C G T A G T C A C G T A G C T | 1e-26 | -5.987e+01 | 70.04% | 36.48% | 1531.9bp (14547.6bp) | che-1/MA0260.1/Jaspar(0.837) More Information \| Similar Motifs Found | motif file (matrix) |
| 5 | C G T A A C T G A C T G C G T A A C G T A C G T | 1e-23 | -5.491e+01 | 76.11% | 44.12% | 1509.3bp (13537.5bp) | G3BP2(RRM)/Homo\_sapiens-RNCMPT00021-PBM/HughesRNA(0.844) More Information \| Similar Motifs Found | motif file (matrix) |
| 6 | A C G T A C T G A C T G A T C G T G C A A C G T C T G A C G T A | 1e-23 | -5.490e+01 | 77.33% | 45.44% | 1472.5bp (14249.6bp) | At1g49010(MYBrelated)/col-At1g49010-DAP-Seq(GSE60143)/Homer(0.790) More Information \| Similar Motifs Found | motif file (matrix) |
| 7 | A C G T G A T C G A C T A G C T G T A C G A C T G A T C G A T C T G A C T G A C | 1e-23 | -5.352e+01 | 83.00% | 52.39% | 1681.1bp (12613.5bp) | LIN28A(CSD,Znf)/Homo\_sapiens-RNCMPT00036-PBM/HughesRNA(0.730) More Information \| Similar Motifs Found | motif file (matrix) |
| 8 | C G T A C G T A A C T G A C T G A C G T A C T G A C T G C G T A | 1e-22 | -5.196e+01 | 83.81% | 53.90% | 1533.7bp (13869.3bp) | PF10\_0068(RRM)/Plasmodium\_falciparum-RNCMPT00199-PBM/HughesRNA(0.778) More Information \| Similar Motifs Found | motif file (matrix) |
| 9 | C G T A C G T A A C T G G T C A C T A G A C G T A C T G C G T A | 1e-21 | -5.047e+01 | 49.80% | 21.53% | 1384.7bp (15936.1bp) | AT3G46070/MA1381.1/Jaspar(0.834) More Information \| Similar Motifs Found | motif file (matrix) |
| 10 | C G T A C G T A C G T A A C T G A T C G G A T C C G A T G C A T G A T C G C A T | 1e-21 | -5.025e+01 | 55.87% | 26.43% | 1576.3bp (11666.8bp) | Kr/dmmpmm(SeSiMCMC)/fly(0.717) More Information \| Similar Motifs Found | motif file (matrix) |
| 11 | C T G A G T A C C G A T C G A T A G C T C T G A C T G A C T A G A T G C G C T A | 1e-19 | -4.450e+01 | 39.68% | 15.51% | 1529.6bp (14045.8bp) | FoxL2(Forkhead)/Ovary-FoxL2-ChIP-Seq(GSE60858)/Homer(0.725) More Information \| Similar Motifs Found | motif file (matrix) |
| 12 | A G T C A G T C A C G T A G T C A G T C A C G T C G T A A C G T A C G T A G T C | 1e-18 | -4.361e+01 | 83.00% | 55.82% | 1585.5bp (17069.2bp) | TF3A(C2H2)/col-TF3A-DAP-Seq(GSE60143)/Homer(0.692) More Information \| Similar Motifs Found | motif file (matrix) |
| 13 | C G A T A C T G C G T A A T C G A C T G T G C A A C T G G T C A | 1e-18 | -4.322e+01 | 57.89% | 30.16% | 1464.7bp (11722.9bp) | SRSF2(RRM)/Homo\_sapiens-RNCMPT00072-PBM/HughesRNA(0.832) More Information \| Similar Motifs Found | motif file (matrix) |
| 14 | A C G T A G T C C G T A A C G T C G T A A G T C A C G T C G T A C G T A C G T A | 1e-16 | -3.856e+01 | 91.50% | 69.08% | 1597.4bp (13502.7bp) | PFF0320c(RRM)/Plasmodium\_falciparum-RNCMPT00235-PBM/HughesRNA(0.699) More Information \| Similar Motifs Found | motif file (matrix) |
| 15 | A C T G A C G T A G C T A G T C A C G T C G T A G T C A A C T G A C T G A C G T | 1e-15 | -3.478e+01 | 80.16% | 55.84% | 1475.7bp (13160.0bp) | PB0194.1\_Zbtb12\_2/Jaspar(0.816) More Information \| Similar Motifs Found | motif file (matrix) |
| 16 | G T A C A C G T A C G T A C G T C G T A C T G A A C T G A C G T A C T G A C G T | 1e-14 | -3.392e+01 | 87.45% | 65.26% | 1480.7bp (14640.7bp) | bap/MA0211.1/Jaspar(0.827) More Information \| Similar Motifs Found | motif file (matrix) |
| 17 | A C T G C G T A A C G T A C G T C G T A A C T G A C G T C G A T A C T G G T A C | 1e-13 | -3.202e+01 | 91.09% | 71.10% | 1562.1bp (14830.1bp) | PFF0320c(RRM)/Plasmodium\_falciparum-RNCMPT00235-PBM/HughesRNA(0.777) More Information \| Similar Motifs Found | motif file (matrix) |
| 18 | A C T G G T A C A C T G A C T G A C G T A C T G A C T G A C T G | 1e-13 | -3.170e+01 | 77.33% | 53.85% | 1622.7bp (12853.2bp) | AT3G57600(AP2EREBP)/col-AT3G57600-DAP-Seq(GSE60143)/Homer(0.859) More Information \| Similar Motifs Found | motif file (matrix) |
| 19 | A C T G C G T A A C G T C G T A A C T G C G T A A C T G A G T C C G T A A G T C | 1e-13 | -3.036e+01 | 84.62% | 63.10% | 1499.2bp (13498.1bp) | ZNF415(Zf)/HEK293-ZNF415.GFP-ChIP-Seq(GSE58341)/Homer(0.728) More Information \| Similar Motifs Found | motif file (matrix) |
| 20 | A G T C A G T C A C G T A C G T A G T C A G T C A G T C A C G T A C G T A G T C | 1e-12 | -2.879e+01 | 82.59% | 61.30% | 1737.7bp (16446.7bp) | PCBP2(KH)/Homo\_sapiens-RNCMPT00044-PBM/HughesRNA(0.764) More Information \| Similar Motifs Found | motif file (matrix) |
| 21 \* | A G T C A G T C A C G T A G T C A C G T A C G T A G C T A C G T A G T C A C G T | 1e-10 | -2.530e+01 | 92.31% | 75.75% | 1529.6bp (13722.1bp) | PTBP1(RRM)/Homo\_sapiens-RNCMPT00268-PBM/HughesRNA(0.773) More Information \| Similar Motifs Found | motif file (matrix) |
| 22 \* | A C G T A G T C A G T C A C G T A T C G A C G T C G T A A C G T A T C G A C G T | 1e-7 | -1.792e+01 | 7.69% | 1.56% | 925.5bp (5337.2bp) | HNRPLL(RRM)/Homo\_sapiens-RNCMPT00178-PBM/HughesRNA(0.720) More Information \| Similar Motifs Found | motif file (matrix) |
