## Supplemental Tabel 1 for "The Establishment of Cell-Type Specific Gene Regulation in the Sea Urchin Embryo": Endoderm_homerResults.html

/data/gpfs-1/users/jbrande\_m/work/scEarlyDev/scATACseq\_analysis/peaksPerLineage/Motifs/Endoderm/ - Homer de novo Motif Results


### Homer *de novo* Motif Results (/data/gpfs-1/users/jbrande\_m/work/scEarlyDev/scATACseq\_analysis/peaksPerLineage/Motifs/Endoderm/)

Known Motif Enrichment Results  
Gene Ontology Enrichment Results  
If Homer is having trouble matching a motif to a known motif, try copy/pasting the matrix file into
STAMP  
More information on motif finding results: HOMER
| Description of Results
| Tips
  
Total target sequences = 1504  
Total background sequences = 4513  
\* - possible false positive  

|  |  |  |  |  |  |  |  |  |
| --- | --- | --- | --- | --- | --- | --- | --- | --- |
| Rank | Motif | P-value | log P-pvalue | % of Targets | % of Background | STD(Bg STD) | Best Match/Details | Motif File |
| 1 | G C A T T C A G C T G A A C T G G A T C G C T A C G T A C G T A A G T C G T C A | 1e-160 | -3.699e+02 | 80.78% | 46.92% | 1379.5bp (9331.4bp) | fkh/dmmpmm(Noyes)/fly(0.933) More Information \| Similar Motifs Found | motif file (matrix) |
| 2 | A G T C C T G A G T C A A C G T C G T A A C G T A C G T A C T G C G T A A G T C | 1e-45 | -1.037e+02 | 80.98% | 64.33% | 1487.3bp (11465.9bp) | YOX1/Literature(Harbison)/Yeast(0.806) More Information \| Similar Motifs Found | motif file (matrix) |
| 3 | A C T G C G T A A C G T C G T A C G T A A C T G | 1e-41 | -9.522e+01 | 52.66% | 35.50% | 1401.6bp (11026.6bp) | GZF3/Literature(Harbison)/Yeast(1.000) More Information \| Similar Motifs Found | motif file (matrix) |
| 4 | C T A G A C T G A C G T A G T C A G T C C G T A | 1e-41 | -9.488e+01 | 71.61% | 54.59% | 1456.3bp (10186.3bp) | HSF1/MA0319.1/Jaspar(0.799) More Information \| Similar Motifs Found | motif file (matrix) |
| 5 | C G T A C G T A C G T A A T G C A C G T C G T A C G T A C T G A C G T A C G T A | 1e-39 | -9.181e+01 | 77.53% | 61.45% | 1460.0bp (13855.1bp) | HuR(?)/HEK293-HuR-CLIP-Seq(GSE87887)/Homer(0.774) More Information \| Similar Motifs Found | motif file (matrix) |
| 6 | C G T A A C T G C G T A A C G T C G A T C T G A A G T C C T A G | 1e-39 | -9.127e+01 | 67.29% | 50.36% | 1422.5bp (10170.4bp) | GATA3(Zf)/iTreg-Gata3-ChIP-Seq(GSE20898)/Homer(0.837) More Information \| Similar Motifs Found | motif file (matrix) |
| 7 | A C G T A G T C C G A T A C T G A C G T C A G T A G T C A C G T | 1e-37 | -8.624e+01 | 79.65% | 64.43% | 1481.9bp (11496.4bp) | pros/dmmpmm(Bergman)/fly(0.751) More Information \| Similar Motifs Found | motif file (matrix) |
| 8 | C G T A G C A T C G A T C G A T A C G T C G T A A G T C C G T A C T G A C G T A | 1e-36 | -8.310e+01 | 71.61% | 55.76% | 1432.7bp (11457.0bp) | CG11617/dmmpmm(Noyes\_hd)/fly(0.762) More Information \| Similar Motifs Found | motif file (matrix) |
| 9 | A G T C C T G A C G T A A G T C C G T A A C G T C G T A A C T G | 1e-35 | -8.272e+01 | 89.69% | 77.16% | 1503.4bp (11878.6bp) | RAV1(RAV)/colamp-RAV1-DAP-Seq(GSE60143)/Homer(0.769) More Information \| Similar Motifs Found | motif file (matrix) |
| 10 | T G A C T C A G G T C A C G T A A T C G G T C A T C G A A C T G C A G T T C A G | 1e-35 | -8.088e+01 | 78.86% | 64.05% | 1473.9bp (10915.0bp) | TRA2(RRM)/Drosophila\_melanogaster-RNCMPT00078-PBM/HughesRNA(0.799) More Information \| Similar Motifs Found | motif file (matrix) |
| 11 | C G A T A C T G A C G T A G T C A G T C A C G T C G T A A C T G | 1e-34 | -8.056e+01 | 76.66% | 61.59% | 1517.9bp (12276.0bp) | SRSF1(RRM)/Homo\_sapiens-RNCMPT00110-PBM/HughesRNA(0.792) More Information \| Similar Motifs Found | motif file (matrix) |
| 12 | A C T G A G T C A G T C A C G T A G C T C G T A A C T G C G T A | 1e-34 | -7.902e+01 | 79.06% | 64.47% | 1442.0bp (9406.8bp) | TFAP2A/MA0003.4/Jaspar(0.683) More Information \| Similar Motifs Found | motif file (matrix) |
| 13 | A G T C C G A T A G T C A G T C A C G T A G T C G T A C A C G T | 1e-31 | -7.330e+01 | 65.16% | 50.00% | 1424.9bp (10088.3bp) | SRSF1(RRM)/Homo\_sapiens-RNCMPT00163-PBM/HughesRNA(0.872) More Information \| Similar Motifs Found | motif file (matrix) |
| 14 | A C T G T A C G A T C G A C T G A C G T A C G T C G T A C A G T | 1e-28 | -6.586e+01 | 75.47% | 61.85% | 1463.5bp (11451.1bp) | Bgb::run/MA0242.1/Jaspar(0.788) More Information \| Similar Motifs Found | motif file (matrix) |
| 15 | C G T A A C G T A G T C A C G T A C G T C G T A C G T A A C T G | 1e-27 | -6.379e+01 | 63.16% | 49.04% | 1492.1bp (12271.9bp) | bap/dmmpmm(Noyes\_hd)/fly(0.734) More Information \| Similar Motifs Found | motif file (matrix) |
| 16 | A G T C A G T C A G T C A G T C A G T C A G T C A G T C A G T C | 1e-27 | -6.362e+01 | 54.72% | 40.64% | 1519.8bp (16130.1bp) | Maz(Zf)/HepG2-Maz-ChIP-Seq(GSE31477)/Homer(0.974) More Information \| Similar Motifs Found | motif file (matrix) |
| 17 | A C G T C G T A C G T A C G T A C G T A C G T A A C G T A C T G A C G T A G T C | 1e-27 | -6.295e+01 | 86.77% | 75.38% | 1481.6bp (11843.7bp) | Lm\_0212(RRM)/Leishmania\_major-RNCMPT00212-PBM/HughesRNA(0.792) More Information \| Similar Motifs Found | motif file (matrix) |
| 18 | A T G C C G T A C G T A C G T A A C G T A C G T A C G T C T A G A G T C A G T C | 1e-26 | -6.146e+01 | 76.99% | 64.08% | 1498.7bp (11271.0bp) | CG17838(RRM)/Drosophila\_melanogaster-RNCMPT00131-PBM/HughesRNA(0.774) More Information \| Similar Motifs Found | motif file (matrix) |
| 19 | A G T C A C G T A C G T A C T G C G T A C G T A A C G T C G T A A C T G C G T A | 1e-26 | -6.042e+01 | 79.59% | 67.13% | 1505.8bp (11823.5bp) | vvl/dmmpmm(Bigfoot)/fly(0.706) More Information \| Similar Motifs Found | motif file (matrix) |
| 20 | C G T A A G T C G T A C A C T G C G A T A C T G C G T A T A G C C G A T G T A C | 1e-26 | -5.993e+01 | 73.60% | 60.52% | 1477.7bp (10705.9bp) | prd/dmmpmm(Noyes)/fly(0.756) More Information \| Similar Motifs Found | motif file (matrix) |
| 21 | C G T A C G T A A G T C A C G T A C G T A G C T A T C G A C T G C G T A C T G A | 1e-25 | -5.853e+01 | 78.79% | 66.45% | 1454.3bp (11962.1bp) | MET31/Literature(Harbison)/Yeast(0.652) More Information \| Similar Motifs Found | motif file (matrix) |
| 22 | C G T A C G T A A C T G A C T G A G C T A C T G A C G T A C T G C G T A C G T A | 1e-24 | -5.654e+01 | 77.06% | 64.74% | 1504.4bp (11441.8bp) | TBX21/MA0690.1/Jaspar(0.968) More Information \| Similar Motifs Found | motif file (matrix) |
| 23 | C G A T A G C T C G T A A C G T C G T A T C G A A C G T G T A C A G T C A C T G | 1e-23 | -5.479e+01 | 59.04% | 45.94% | 1492.2bp (12695.0bp) | PB0174.1\_Sox30\_2/Jaspar(0.839) More Information \| Similar Motifs Found | motif file (matrix) |
| 24 | C G T A A G T C A C G T A C G T C G T A A C T G A C T G A C T G | 1e-23 | -5.327e+01 | 98.54% | 93.01% | 1515.2bp (11927.9bp) | HNRNPA1L2(RRM)/Homo\_sapiens-RNCMPT00023-PBM/HughesRNA(0.812) More Information \| Similar Motifs Found | motif file (matrix) |
| 25 | A C T G A C T G A C T G A C T G A G T C A C G T G T C A A C T G | 1e-20 | -4.717e+01 | 94.88% | 87.73% | 1490.7bp (11277.7bp) | ZNF682/MA1599.1/Jaspar(0.738) More Information \| Similar Motifs Found | motif file (matrix) |
| 26 | A G T C A G T C A G T C A G T C A C G T A C T G A T C G C T A G | 1e-18 | -4.371e+01 | 55.05% | 43.44% | 1594.1bp (10699.8bp) | EBF1(EBF)/Near-E2A-ChIP-Seq(GSE21512)/Homer(0.882) More Information \| Similar Motifs Found | motif file (matrix) |
| 27 | A C T G A G T C A C T G A G T C C G T A A G T C A G T C A G C T | 1e-18 | -4.301e+01 | 92.15% | 84.43% | 1509.5bp (11114.6bp) | PB0089.1\_Tcfe2a\_1/Jaspar(0.758) More Information \| Similar Motifs Found | motif file (matrix) |
