## Supplemental Tabel 1 for "The Establishment of Cell-Type Specific Gene Regulation in the Sea Urchin Embryo": Endomeso_homerResults.html

/data/gpfs-1/users/jbrande\_m/work/scEarlyDev/scATACseq\_analysis/peaksPerLineage/Motifs/Endomeso/ - Homer de novo Motif Results


### Homer *de novo* Motif Results (/data/gpfs-1/users/jbrande\_m/work/scEarlyDev/scATACseq\_analysis/peaksPerLineage/Motifs/Endomeso/)

Known Motif Enrichment Results  
Gene Ontology Enrichment Results  
If Homer is having trouble matching a motif to a known motif, try copy/pasting the matrix file into
STAMP  
More information on motif finding results: HOMER
| Description of Results
| Tips
  
Total target sequences = 710  
Total background sequences = 5276  
\* - possible false positive  

|  |  |  |  |  |  |  |  |  |
| --- | --- | --- | --- | --- | --- | --- | --- | --- |
| Rank | Motif | P-value | log P-pvalue | % of Targets | % of Background | STD(Bg STD) | Best Match/Details | Motif File |
| 1 | G A C T C G T A A C G T A G T C A G T C A C T G A C T G A C T G | 1e-28 | -6.618e+01 | 15.63% | 4.54% | 965.9bp (5950.5bp) | SPDEF/MA0686.1/Jaspar(0.848) More Information \| Similar Motifs Found | motif file (matrix) |
| 2 | T C G A C G T A A G T C C G T A C G T A C G T A C T G A A C T G | 1e-28 | -6.499e+01 | 67.89% | 47.16% | 1054.0bp (5147.8bp) | Sox3(HMG)/NPC-Sox3-ChIP-Seq(GSE33059)/Homer(0.932) More Information \| Similar Motifs Found | motif file (matrix) |
| 3 | C G T A T G A C C T A G C G T A G C T A A G T C C T G A A C T G A C T G C G T A | 1e-22 | -5.080e+01 | 9.44% | 2.21% | 1139.5bp (6259.6bp) | IKZF1/MA1508.1/Jaspar(0.791) More Information \| Similar Motifs Found | motif file (matrix) |
| 4 | C A T G T A G C T C A G C T G A G C A T A G T C C A T G G T A C G T A C C T A G | 1e-18 | -4.266e+01 | 51.55% | 35.15% | 1098.4bp (5251.2bp) | wc-1/MA1437.1/Jaspar(0.726) More Information \| Similar Motifs Found | motif file (matrix) |
| 5 | A C T G A C T G C G T A A C G T A C G T C G T A | 1e-18 | -4.238e+01 | 53.66% | 37.22% | 1126.3bp (5525.4bp) | bcd/MA0212.1/Jaspar(0.998) More Information \| Similar Motifs Found | motif file (matrix) |
| 6 | A C G T C G T A A G T C A C T G C G T A C G T A A G T C A C T G | 1e-16 | -3.753e+01 | 83.24% | 69.61% | 1250.1bp (5623.2bp) | ZNF638(RRM)/Homo\_sapiens-RNCMPT00164-PBM/HughesRNA(0.795) More Information \| Similar Motifs Found | motif file (matrix) |
| 7 | A C G T C G T A T C A G A C T G G A T C G A T C G A T C T C G A C T G A G C A T | 1e-15 | -3.651e+01 | 60.14% | 44.77% | 1212.2bp (5265.9bp) | PCF/Arabidopsis-Promoters/Homer(0.765) More Information \| Similar Motifs Found | motif file (matrix) |
| 8 | G A T C T A G C C T G A T G C A G T C A T C A G T A C G C T A G C G A T A G T C | 1e-15 | -3.492e+01 | 78.31% | 64.43% | 1241.2bp (5558.1bp) | Kr/dmmpmm(Noyes)/fly(0.738) More Information \| Similar Motifs Found | motif file (matrix) |
| 9 | A T C G A T C G G T C A T A G C T G C A T G C A G T A C A C T G | 1e-15 | -3.476e+01 | 13.94% | 5.76% | 1377.3bp (4603.6bp) | RBP1(RRM)/Drosophila\_melanogaster-RNCMPT00058-PBM/HughesRNA(0.792) More Information \| Similar Motifs Found | motif file (matrix) |
| 10 | C T A G A G T C A G T C C T G A A C T G C G T A C G T A A G T C | 1e-13 | -3.120e+01 | 28.03% | 16.68% | 1209.0bp (5126.8bp) | Hand1::Tcf3/MA0092.1/Jaspar(0.755) More Information \| Similar Motifs Found | motif file (matrix) |
| 11 | C G T A A C G T A T C G A C G T C T G A C G A T C G T A C G T A | 1e-13 | -3.028e+01 | 26.90% | 15.93% | 1267.4bp (6869.1bp) | Cf2-II/dmmpmm(Pollard)/fly(0.810) More Information \| Similar Motifs Found | motif file (matrix) |
| 12 | A C T G C G T A A C G T A G T C A C G T C G T A A C G T A C G T | 1e-13 | -3.013e+01 | 67.46% | 53.79% | 1090.2bp (6710.3bp) | GATA10/MA1013.1/Jaspar(0.779) More Information \| Similar Motifs Found | motif file (matrix) |
| 13 | A G T C A C G T A C T G A G T C A C G T C G T A A C G T A C G T C G T A A G C T | 1e-12 | -2.906e+01 | 4.51% | 0.91% | 817.5bp (5515.5bp) | QKR58E-1(KH)/Drosophila\_melanogaster-RNCMPT00142-PBM/HughesRNA(0.679) More Information \| Similar Motifs Found | motif file (matrix) |
| 14 \* | C G T A A T C G A C G T A C G T A C G T A C G T A C T G C G T A A C G T A G T C | 1e-11 | -2.734e+01 | 2.96% | 0.39% | 894.7bp (786.0bp) | pan/dmmpmm(Pollard)/fly(0.786) More Information \| Similar Motifs Found | motif file (matrix) |
| 15 \* | A G T C A C T G A C T G A G T C A C T G A C T G G A C T C T A G | 1e-11 | -2.698e+01 | 25.77% | 15.58% | 1127.2bp (4764.9bp) | LEP(AP2EREBP)/col-LEP-DAP-Seq(GSE60143)/Homer(0.899) More Information \| Similar Motifs Found | motif file (matrix) |
| 16 \* | A C T G A C G T C G T A C G T A C T G A C T G A G T C A A G T C A C T G A G C T | 1e-10 | -2.528e+01 | 5.49% | 1.50% | 973.8bp (8389.0bp) | gt/dmmpmm(Bigfoot)/fly(0.760) More Information \| Similar Motifs Found | motif file (matrix) |
| 17 \* | C G T A A G T C A G T C A G T C C G T A A C T G A C G T A C G T C G T A G T C A | 1e-10 | -2.463e+01 | 2.25% | 0.24% | 244.0bp (485.4bp) | PB0048.1\_Nkx3-1\_1/Jaspar(0.721) More Information \| Similar Motifs Found | motif file (matrix) |
| 18 \* | A G T C A G T C A G T C A C G T A G T C C G T A A C G T C G T A A C G T A C G T | 1e-10 | -2.445e+01 | 3.10% | 0.49% | 1040.5bp (1499.1bp) | PUF68(RRM)/Drosophila\_melanogaster-RNCMPT00141-PBM/HughesRNA(0.800) More Information \| Similar Motifs Found | motif file (matrix) |
| 19 \* | A C G T A C T G A C T G A G T C A C G T A C G T A C G T A G T C A C G T C T A G | 1e-10 | -2.359e+01 | 2.39% | 0.29% | 634.1bp (927.7bp) | Tb\_0220(RRM)/Trypanosoma\_brucei-RNCMPT00220-PBM/HughesRNA(0.763) More Information \| Similar Motifs Found | motif file (matrix) |
| 20 \* | C G T A A C T G A C G T A C G T A G T C A G T C A C G T A G T C | 1e-10 | -2.324e+01 | 8.17% | 3.14% | 1033.2bp (6258.7bp) | PB0058.1\_Sfpi1\_1/Jaspar(0.808) More Information \| Similar Motifs Found | motif file (matrix) |
| 21 \* | A C G T A C G T A C T G A C T G C G T A A C G T C G T A A C T G A C G T A G T C | 1e-9 | -2.148e+01 | 69.86% | 58.69% | 1248.1bp (5859.3bp) | HNRNPAB(RRM)/Tetraodon\_nigroviridis-RNCMPT00245-PBM/HughesRNA(0.713) More Information \| Similar Motifs Found | motif file (matrix) |
| 22 \* | C G T A A C G T C G T A A C T G A G T C A G T C A G T C A C G T A G T C A C G T | 1e-9 | -2.137e+01 | 98.17% | 93.16% | 1220.5bp (5515.6bp) | Trl(Zf)/S2-GAGAfactor-ChIP-Seq(GSE40646)/Homer(0.642) More Information \| Similar Motifs Found | motif file (matrix) |
| 23 \* | A C G T A C T G A G T C A G T C A G T C A C G T A C T G A C T G A G T C A G T C | 1e-9 | -2.095e+01 | 72.82% | 62.00% | 1158.3bp (5060.3bp) | SKN7/SKN7\_H2O2Lo/[](Harbison)/Yeast(0.738) More Information \| Similar Motifs Found | motif file (matrix) |
| 24 \* | A C T G C G T A A T C G A C T G A C T G A C G T G T C A A T C G | 1e-8 | -2.070e+01 | 7.04% | 2.66% | 1024.3bp (4539.3bp) | RBM5(Znf)/Homo\_sapiens-RNCMPT00055-PBM/HughesRNA(0.712) More Information \| Similar Motifs Found | motif file (matrix) |
| 25 \* | A T G C A C T G A G T C A C T G G C T A C G T A | 1e-8 | -2.068e+01 | 52.11% | 40.90% | 1259.0bp (5160.7bp) | SWI4/MA0401.1/Jaspar(0.916) More Information \| Similar Motifs Found | motif file (matrix) |
| 26 \* | C T G A C T G A A G T C C G T A A C G T A G T C A G T C A G T C A C G T C G A T | 1e-8 | -2.063e+01 | 3.10% | 0.61% | 839.1bp (1337.0bp) | YBX1(CSD)/Homo\_sapiens-RNCMPT00083-PBM/HughesRNA(0.736) More Information \| Similar Motifs Found | motif file (matrix) |
| 27 \* | A G T C A C G T C G T A A C G T C G T A A C T G C G T A C G T A C G T A A C T G | 1e-8 | -1.981e+01 | 4.65% | 1.39% | 998.8bp (14173.2bp) | TATA-box/Drosophila-Promoters/Homer(0.721) More Information \| Similar Motifs Found | motif file (matrix) |
| 28 \* | A T G C T A G C T C G A T G A C A T C G C T A G T A C G T A G C | 1e-7 | -1.760e+01 | 33.24% | 24.08% | 1105.5bp (4456.7bp) | Pho2(bHLH)/Yeast-Pho2-ChIP-Seq(GSE29506)/Homer(0.750) More Information \| Similar Motifs Found | motif file (matrix) |
| 29 \* | A G T C C G T A A C T G C G T A A G T C A G T C A G T C C G T A A G T C C G T A | 1e-6 | -1.440e+01 | 97.04% | 92.72% | 1253.5bp (5195.7bp) | SeqBias: CA-repeat(0.734) More Information \| Similar Motifs Found | motif file (matrix) |
| 30 \* | A C T G C T A G A G T C C G T A C G T A A C T G A G T C C G T A C G A T A C G T | 1e-5 | -1.244e+01 | 2.96% | 0.92% | 643.9bp (5708.6bp) | MBNL1(Znf)/Homo\_sapiens-RNCMPT00038-PBM/HughesRNA(0.763) More Information \| Similar Motifs Found | motif file (matrix) |
| 31 \* | A C T G A C T G A G T C A C G T C G T A A C T G A C G T A C G T | 1e-5 | -1.178e+01 | 83.94% | 77.34% | 1242.9bp (5424.6bp) | PH0040.1\_Hmbox1/Jaspar(0.690) More Information \| Similar Motifs Found | motif file (matrix) |
| 32 \* | A G T C A C G T A C T G A G T C C G T A A C T G C G T A A C T G | 1e-4 | -1.002e+01 | 45.35% | 38.09% | 1290.2bp (6168.9bp) | PB0091.1\_Zbtb3\_1/Jaspar(0.725) More Information \| Similar Motifs Found | motif file (matrix) |
| 33 \* | A C G T A G T C A C G T A C T G A C G T A G T C G T C A A C T G A C G T A G T C | 1e-1 | -4.515e+00 | 2.96% | 1.70% | 893.1bp (956.8bp) | GRF9(GRF)/colamp-GRF9-DAP-Seq(GSE60143)/Homer(0.781) More Information \| Similar Motifs Found | motif file (matrix) |
