## Supplemental Tabel 1 for "The Establishment of Cell-Type Specific Gene Regulation in the Sea Urchin Embryo": GermCells_homerResults.html

/data/gpfs-1/users/jbrande\_m/work/scEarlyDev/scATACseq\_analysis/peaksPerLineage/Motifs/GermCells/ - Homer de novo Motif Results


### Homer *de novo* Motif Results (/data/gpfs-1/users/jbrande\_m/work/scEarlyDev/scATACseq\_analysis/peaksPerLineage/Motifs/GermCells/)

Known Motif Enrichment Results  
Gene Ontology Enrichment Results  
If Homer is having trouble matching a motif to a known motif, try copy/pasting the matrix file into
STAMP  
More information on motif finding results: HOMER
| Description of Results
| Tips
  
Total target sequences = 182  
Total background sequences = 4990  
\* - possible false positive  

|  |  |  |  |  |  |  |  |  |
| --- | --- | --- | --- | --- | --- | --- | --- | --- |
| Rank | Motif | P-value | log P-pvalue | % of Targets | % of Background | STD(Bg STD) | Best Match/Details | Motif File |
| 1 | C G T A A C G T C G T A C G T A C G A T C G T A C G T A A C G T | 1e-16 | -3.832e+01 | 65.38% | 34.52% | 1678.5bp (10133.3bp) | QKR58E-1(KH)/Drosophila\_melanogaster-RNCMPT00142-PBM/HughesRNA(0.870) More Information \| Similar Motifs Found | motif file (matrix) |
| 2 | C T A G T G A C A C G T C G T A A C T G A G C T A G T C A C G T | 1e-16 | -3.757e+01 | 65.38% | 34.82% | 1771.1bp (8663.5bp) | Smad3(MAD)/NPC-Smad3-ChIP-Seq(GSE36673)/Homer(0.775) More Information \| Similar Motifs Found | motif file (matrix) |
| 3 | T A G C A G C T A C T G G T A C C G T A A G C T C T G A G T A C C G T A A T G C | 1e-15 | -3.517e+01 | 45.05% | 18.79% | 1455.1bp (7882.0bp) | ABI3/MA0564.1/Jaspar(0.733) More Information \| Similar Motifs Found | motif file (matrix) |
| 4 | A G T C A C G T G T A C G T A C C T G A A G T C A G T C A G T C C G T A A G T C | 1e-13 | -3.130e+01 | 9.89% | 0.84% | 1370.4bp (7751.0bp) | Sp5(Zf)/mES-Sp5.Flag-ChIP-Seq(GSE72989)/Homer(0.853) More Information \| Similar Motifs Found | motif file (matrix) |
| 5 | A T G C A C G T A T G C A G T C G A T C A C G T C T A G A C T G A C G T G A T C | 1e-13 | -3.045e+01 | 10.44% | 1.02% | 836.7bp (11672.4bp) | Tag/dmmpmm(Papatsenko)/fly(0.723) More Information \| Similar Motifs Found | motif file (matrix) |
| 6 | C G A T C A T G A G T C C T G A G C T A T C A G A G T C A C G T | 1e-12 | -2.915e+01 | 76.92% | 50.51% | 1658.3bp (8282.9bp) | MBNL1(Znf)/Homo\_sapiens-RNCMPT00038-PBM/HughesRNA(0.805) More Information \| Similar Motifs Found | motif file (matrix) |
| 7 | A C G T A C T G A C G T A C T G C G A T C T A G A C G T C G T A | 1e-12 | -2.895e+01 | 60.99% | 34.46% | 1481.9bp (7827.7bp) | SM(RRM)/Drosophila\_melanogaster-RNCMPT00069-PBM/HughesRNA(0.899) More Information \| Similar Motifs Found | motif file (matrix) |
| 8 \* | A C T G C G T A C T A G T A G C A C G T A C T G C G T A A C T G A T G C G C T A | 1e-11 | -2.758e+01 | 17.58% | 3.90% | 1299.6bp (4985.6bp) | ZKSCAN5/MA1652.1/Jaspar(0.789) More Information \| Similar Motifs Found | motif file (matrix) |
| 9 \* | A C G T C T G A G T C A A C G T A G T C A G T C A G T C A G T C | 1e-11 | -2.626e+01 | 64.29% | 38.90% | 1554.0bp (7641.8bp) | PITX2/MA1547.1/Jaspar(0.905) More Information \| Similar Motifs Found | motif file (matrix) |
| 10 \* | A C G T A T G C C T G A C G T A C T A G G T A C C G T A C G A T A G T C A C T G | 1e-11 | -2.548e+01 | 73.63% | 48.85% | 1681.7bp (8909.6bp) | BEAF-32B/dmmpmm(Pollard)/fly(0.672) More Information \| Similar Motifs Found | motif file (matrix) |
| 11 \* | A T G C C G A T T G C A A T C G C G T A A G T C G A T C A C G T | 1e-11 | -2.539e+01 | 74.73% | 50.08% | 1658.1bp (10681.4bp) | GATA9/MA1018.1/Jaspar(0.820) More Information \| Similar Motifs Found | motif file (matrix) |
| 12 \* | A C T G G A T C C A T G A T C G G A T C T G A C T G A C T C A G | 1e-10 | -2.414e+01 | 39.01% | 18.13% | 1489.8bp (9470.5bp) | ERF069/MA0997.1/Jaspar(0.744) More Information \| Similar Motifs Found | motif file (matrix) |
| 13 \* | G T C A A G T C C G A T A C T G C G T A A G T C A G T C G T C A | 1e-10 | -2.313e+01 | 35.71% | 16.07% | 1433.0bp (8476.8bp) | ERF018/MA1048.1/Jaspar(0.776) More Information \| Similar Motifs Found | motif file (matrix) |
| 14 \* | A G T C A C T G A C T G A C G T A G T C C G T A A G T C A G T C C G T A G T C A | 1e-10 | -2.310e+01 | 4.40% | 0.14% | 1255.9bp (9179.7bp) | ASHR1(ND)/col-ASHR1-DAP-Seq(GSE60143)/Homer(0.790) More Information \| Similar Motifs Found | motif file (matrix) |
| 15 \* | A G T C C G T A A G T C A C T G A G T C A C T G A C G T C T G A A G T C A C T G | 1e-9 | -2.078e+01 | 70.33% | 48.04% | 1369.4bp (7538.1bp) | SPL7/MA1060.1/Jaspar(0.740) More Information \| Similar Motifs Found | motif file (matrix) |
| 16 \* | A C G T A G T C A C G T A G T C A C T G C G T A A C T G C G T A A C T G C G T A | 1e-8 | -1.867e+01 | 60.99% | 39.90% | 1367.7bp (8767.5bp) | GFX(?)/Promoter/Homer(0.847) More Information \| Similar Motifs Found | motif file (matrix) |
| 17 \* | A C G T C G T A A C T G A T C G A C G T A C T G A C T G A C G T A G T C A C G T | 1e-7 | -1.627e+01 | 73.08% | 53.91% | 1618.8bp (8075.1bp) | Gli2(Zf)/GM2-Gli2-ChIP-Chip(GSE112702)/Homer(0.791) More Information \| Similar Motifs Found | motif file (matrix) |
| 18 \* | C G A T A T G C C G A T A C T G C G A T A C T G C G T A C G T A C G T A A C T G | 1e-6 | -1.485e+01 | 20.88% | 8.72% | 894.0bp (5066.2bp) | PRDM1/MA0508.3/Jaspar(0.783) More Information \| Similar Motifs Found | motif file (matrix) |
| 19 \* | G C A T A G T C A G C T A C G T A G C T A G C T A G C T T C G A T C A G G C A T | 1e-6 | -1.391e+01 | 8.79% | 1.98% | 1554.8bp (3893.2bp) | SA0001.1\_at\_AC\_acceptor/Jaspar(0.676) More Information \| Similar Motifs Found | motif file (matrix) |
| 20 \* | A C G T G T C A A C G T A G T C A C G T A G T C C G T A A T C G A C T G A C T G | 1e-5 | -1.363e+01 | 80.22% | 63.89% | 1689.4bp (8687.1bp) | REI1/MA0364.1/Jaspar(0.685) More Information \| Similar Motifs Found | motif file (matrix) |
| 21 \* | A G T C A G T C A C T G C G T A A C T G A C G T | 1e-5 | -1.254e+01 | 98.35% | 89.58% | 1514.6bp (8453.3bp) | CMTA3/MA0970.1/Jaspar(0.791) More Information \| Similar Motifs Found | motif file (matrix) |
| 22 \* | A C G T C G A T G T A C C T G A A C T G A G T C A G T C C G T A A C G T C G T A | 1e-5 | -1.153e+01 | 3.30% | 0.27% | 429.9bp (3697.2bp) | pho/dmmpmm(Pollard)/fly(0.713) More Information \| Similar Motifs Found | motif file (matrix) |
| 23 \* | A C G T A C G T A C G T A C G T A C G T A C G T A C G T A C G T A C G T A C G T | 1e-4 | -1.139e+01 | 57.69% | 41.80% | 1520.6bp (10984.1bp) | SeqBias: polyA-repeat(1.000) More Information \| Similar Motifs Found | motif file (matrix) |
| 24 \* | A C G T A C G T A C T G C G T A A C T G A C G T C G T A A C T G A C G T A C G T | 1e-4 | -1.069e+01 | 71.43% | 56.46% | 1834.5bp (8432.2bp) | MSI(RRM)/Drosophila\_melanogaster-RNCMPT00040-PBM/HughesRNA(0.763) More Information \| Similar Motifs Found | motif file (matrix) |
