## Supplemental Tabel 1 for "The Establishment of Cell-Type Specific Gene Regulation in the Sea Urchin Embryo": Neuron_homerResults.html

/data/gpfs-1/users/jbrande\_m/work/scEarlyDev/scATACseq\_analysis/peaksPerLineage/Motifs/Neuron/ - Homer de novo Motif Results


### Homer *de novo* Motif Results (/data/gpfs-1/users/jbrande\_m/work/scEarlyDev/scATACseq\_analysis/peaksPerLineage/Motifs/Neuron/)

Known Motif Enrichment Results  
Gene Ontology Enrichment Results  
If Homer is having trouble matching a motif to a known motif, try copy/pasting the matrix file into
STAMP  
More information on motif finding results: HOMER
| Description of Results
| Tips
  
Total target sequences = 589  
Total background sequences = 4709  
\* - possible false positive  

|  |  |  |  |  |  |  |  |  |
| --- | --- | --- | --- | --- | --- | --- | --- | --- |
| Rank | Motif | P-value | log P-pvalue | % of Targets | % of Background | STD(Bg STD) | Best Match/Details | Motif File |
| 1 | G A C T A T C G A G T C C G T A A C G T A G T C C G A T A T C G | 1e-32 | -7.549e+01 | 55.18% | 31.16% | 1461.2bp (4610.4bp) | Tal1(0.879) More Information \| Similar Motifs Found | motif file (matrix) |
| 2 | C G T A A C G T A C T G A T G C C T G A A C G T A T C G A G T C | 1e-32 | -7.460e+01 | 48.39% | 25.41% | 1526.1bp (4110.6bp) | A2BP1(RRM)/Drosophila\_melanogaster-RNCMPT00123-PBM/HughesRNA(0.907) More Information \| Similar Motifs Found | motif file (matrix) |
| 3 | T C G A C G T A A C G T A G T C G T C A C G T A A C G T C T G A | 1e-32 | -7.399e+01 | 51.78% | 28.39% | 1358.4bp (5108.4bp) | Cux2(Homeobox)/Liver-Cux2-ChIP-Seq(GSE35985)/Homer(0.947) More Information \| Similar Motifs Found | motif file (matrix) |
| 4 | C A T G G C A T A C T G C G A T C A T G C G A T C T A G C G A T C T A G G C A T | 1e-31 | -7.159e+01 | 41.09% | 19.94% | 903.1bp (3228.0bp) | SeqBias: CA-repeat(0.893) More Information \| Similar Motifs Found | motif file (matrix) |
| 5 | C T A G A G C T G C T A C G A T C G T A G C T A A G C T A T C G A T G C G T C A | 1e-24 | -5.595e+01 | 28.69% | 12.67% | 1530.0bp (4691.7bp) | OCT:OCT-short(POU,Homeobox)/NPC-OCT6-ChIP-Seq(GSE43916)/Homer(0.733) More Information \| Similar Motifs Found | motif file (matrix) |
| 6 | T C G A C T A G G A T C T A C G C T G A A C T G G T A C C T A G | 1e-23 | -5.310e+01 | 67.06% | 46.54% | 1200.1bp (3290.0bp) | Clamp/MA1700.1/Jaspar(0.754) More Information \| Similar Motifs Found | motif file (matrix) |
| 7 | T G A C C G A T T C G A G C A T T A C G G T A C C G T A A C T G G A T C C G T A | 1e-21 | -4.900e+01 | 40.24% | 22.53% | 1427.7bp (5040.8bp) | vvl/dmmpmm(Pollard)/fly(0.703) More Information \| Similar Motifs Found | motif file (matrix) |
| 8 | C A T G G A T C G C A T C T G A C G T A A G C T G C A T T A C G A T G C C T G A | 1e-20 | -4.748e+01 | 29.88% | 14.60% | 1560.8bp (5011.2bp) | Isl1(Homeobox)/Neuron-Isl1-ChIP-Seq(GSE31456)/Homer(0.783) More Information \| Similar Motifs Found | motif file (matrix) |
| 9 | T C G A C T A G T C G A A G C T C T A G A T C G C G T A T C G A C G A T G T C A | 1e-19 | -4.427e+01 | 14.26% | 4.59% | 1717.4bp (5302.1bp) | NFATC3/MA0625.1/Jaspar(0.777) More Information \| Similar Motifs Found | motif file (matrix) |
| 10 | T A G C C A G T C T G A C T G A A G C T A C T G T G C A T A C G | 1e-18 | -4.358e+01 | 71.14% | 52.90% | 1485.4bp (4864.5bp) | POU6F1(var.2)/MA1549.1/Jaspar(0.894) More Information \| Similar Motifs Found | motif file (matrix) |
| 11 | C G A T G A T C A G C T A G T C A G C T C T G A G C A T T A G C A C G T A G T C | 1e-18 | -4.305e+01 | 26.15% | 12.47% | 1163.9bp (4433.3bp) | SeqBias: GA-repeat(0.834) More Information \| Similar Motifs Found | motif file (matrix) |
| 12 | A G C T G A C T A G T C G A C T G A T C G A C T A G T C G T A C G C A T G T A C | 1e-17 | -4.080e+01 | 44.14% | 27.35% | 1409.6bp (5631.1bp) | SRSF2(RRM)/Homo\_sapiens-RNCMPT00072-PBM/HughesRNA(0.802) More Information \| Similar Motifs Found | motif file (matrix) |
| 13 | A T G C A C T G A G T C C G A T A C T G G C A T A C T G C G A T | 1e-16 | -3.780e+01 | 25.64% | 12.82% | 1218.3bp (5665.8bp) | POL009.1\_DCE\_S\_II/Jaspar(0.786) More Information \| Similar Motifs Found | motif file (matrix) |
| 14 | G A C T C G T A A C G T T A C G G A T C T G C A T A G C T C A G C A T G C G A T | 1e-16 | -3.718e+01 | 6.45% | 1.18% | 1170.7bp (8813.7bp) | SNRPA(RRM)/Homo\_sapiens-RNCMPT00071-PBM/HughesRNA(0.649) More Information \| Similar Motifs Found | motif file (matrix) |
| 15 | A G T C C G T A A G T C A C G T A G T C A C G T | 1e-13 | -3.078e+01 | 51.95% | 36.77% | 1408.9bp (4843.8bp) | z/dmmpmm(Down)/fly(0.837) More Information \| Similar Motifs Found | motif file (matrix) |
| 16 | C G T A A C G T A C T G A T G C A G T C T A C G G T C A A C G T T A G C T A G C | 1e-13 | -3.072e+01 | 3.06% | 0.26% | 1304.8bp (758.7bp) | PF10\_0214(RRM)/Plasmodium\_falciparum-RNCMPT00240-PBM/HughesRNA(0.693) More Information \| Similar Motifs Found | motif file (matrix) |
| 17 \* | A T G C A G T C C T G A A T G C T A G C A T C G A C T G A G C T | 1e-7 | -1.817e+01 | 8.15% | 3.28% | 1614.6bp (3244.3bp) | MET4(MacIsaac)/Yeast(0.805) More Information \| Similar Motifs Found | motif file (matrix) |
| 18 \* | A C G T A G T C A C T G A G C T C T A G A C G T C G T A A G T C | 1e-6 | -1.417e+01 | 7.81% | 3.55% | 747.0bp (5313.5bp) | Lm\_0223(RRM)/Leishmania\_major-RNCMPT00223-PBM/HughesRNA(0.826) More Information \| Similar Motifs Found | motif file (matrix) |
| 19 \* | C G T A A G T C A T G C C T A G C T G A A G T C A C T G A G T C A G T C C T G A | 1e-6 | -1.405e+01 | 1.70% | 0.22% | 929.5bp (3833.1bp) | Os05g0497200/MA1034.1/Jaspar(0.736) More Information \| Similar Motifs Found | motif file (matrix) |
