## Supplemental Tabel 1 for "The Establishment of Cell-Type Specific Gene Regulation in the Sea Urchin Embryo": OralNSM_homerResults.html

/data/gpfs-1/users/jbrande\_m/work/scEarlyDev/scATACseq\_analysis/peaksPerLineage/Motifs/OralNSM/ - Homer de novo Motif Results


### Homer *de novo* Motif Results (/data/gpfs-1/users/jbrande\_m/work/scEarlyDev/scATACseq\_analysis/peaksPerLineage/Motifs/OralNSM/)

Known Motif Enrichment Results  
Gene Ontology Enrichment Results  
If Homer is having trouble matching a motif to a known motif, try copy/pasting the matrix file into
STAMP  
More information on motif finding results: HOMER
| Description of Results
| Tips
  
Total target sequences = 3061  
Total background sequences = 2279  
\* - possible false positive  

|  |  |  |  |  |  |  |  |  |
| --- | --- | --- | --- | --- | --- | --- | --- | --- |
| Rank | Motif | P-value | log P-pvalue | % of Targets | % of Background | STD(Bg STD) | Best Match/Details | Motif File |
| 1 | T A G C T G C A A C T G A C T G C G T A C G T A T C A G A G C T | 1e-263 | -6.074e+02 | 45.02% | 17.78% | 1299.4bp (1629.8bp) | Eip74EF/dmmpmm(Bigfoot)/fly(0.968) More Information \| Similar Motifs Found | motif file (matrix) |
| 2 | A G T C T A G C A T C G A C T G C T G A C G T A C T G A C A T G | 1e-25 | -5.762e+01 | 11.34% | 6.31% | 1179.2bp (1269.7bp) | LBD18/MA1673.1/Jaspar(0.819) More Information \| Similar Motifs Found | motif file (matrix) |
| 3 | C G A T A G T C C T G A A C G T C G T A A G T C A G T C G T C A A C G T A C T G | 1e-23 | -5.410e+01 | 2.71% | 0.74% | 1123.1bp (1569.7bp) | GAT3(MacIsaac)/Yeast(0.772) More Information \| Similar Motifs Found | motif file (matrix) |
| 4 | A C G T A C G T A C G T A C T G C G T A A G T C C G T A C G T A A C T G A C G T | 1e-19 | -4.525e+01 | 1.11% | 0.14% | 1411.5bp (131.4bp) | exd/dmmpmm(Noyes\_hd)/fly(0.825) More Information \| Similar Motifs Found | motif file (matrix) |
| 5 | A G T C A G C T A T C G A C G T A C G T A G T C C G T A A G T C C G T A A C G T | 1e-19 | -4.478e+01 | 2.03% | 0.50% | 1432.2bp (2667.9bp) | INO4/INO4\_YPD/4-INO4,37-INO2(Harbison)/Yeast(0.799) More Information \| Similar Motifs Found | motif file (matrix) |
| 6 | A G T C A C T G A C G T C G A T A C G T A G T C A G T C A C G T | 1e-17 | -4.065e+01 | 6.70% | 3.50% | 1326.0bp (1725.5bp) | UME1/UME1\_YPD/[](Harbison)/Yeast(0.749) More Information \| Similar Motifs Found | motif file (matrix) |
| 7 | A C T G A C T G A C T G A C T G A G T C A T C G A T C G C G T A A C G T A G T C | 1e-13 | -3.127e+01 | 1.01% | 0.20% | 959.8bp (1128.1bp) | btd/dmmpmm(Noyes)/fly(0.822) More Information \| Similar Motifs Found | motif file (matrix) |
| 8 | C G T A A G T C A C G T C G T A A G T C A C G T A C T G A G T C | 1e-13 | -3.042e+01 | 4.12% | 1.98% | 1531.3bp (796.3bp) | MSI(RRM)/Drosophila\_melanogaster-RNCMPT00040-PBM/HughesRNA(0.801) More Information \| Similar Motifs Found | motif file (matrix) |
| 9 | A C T G A G C T C G T A G T A C G T C A A G T C A G T C G T C A A C G T C T A G | 1e-12 | -2.871e+01 | 2.12% | 0.76% | 1263.9bp (1801.5bp) | NR3C1/MA0113.3/Jaspar(0.706) More Information \| Similar Motifs Found | motif file (matrix) |
| 10 | C G T A A C G T A C T G A C G T A C G T A G T C A C G T A C G T A C G T A G T C | 1e-12 | -2.776e+01 | 0.95% | 0.19% | 1987.0bp (1783.3bp) | MGA1/MA0336.1/Jaspar(0.755) More Information \| Similar Motifs Found | motif file (matrix) |
| 11 \* | A G T C C G T A C G A T C T A G A C G T C G A T A G T C G T C A | 1e-10 | -2.462e+01 | 28.62% | 23.44% | 1431.9bp (1724.6bp) | ESRRA/MA0592.3/Jaspar(0.867) More Information \| Similar Motifs Found | motif file (matrix) |
| 12 \* | A C G T A G T C C G T A A G T C A C G T A C T G A C G T A G T C C G T A C G T A | 1e-9 | -2.075e+01 | 1.18% | 0.39% | 1361.4bp (1417.2bp) | PB0195.1\_Zbtb3\_2/Jaspar(0.781) More Information \| Similar Motifs Found | motif file (matrix) |
| 13 \* | C G T A C G T A A C G T A G T C A C G T A C G T A G T C C G T A A C G T C G T A | 1e-8 | -2.028e+01 | 1.08% | 0.34% | 1015.4bp (1538.0bp) | Brn2(POU,Homeobox)/NPC-Brn2-ChIP-Seq(GSE35496)/Homer(0.714) More Information \| Similar Motifs Found | motif file (matrix) |
| 14 \* | A C G T A C G T C G T A A C G T C G T A A C T G | 1e-8 | -1.962e+01 | 48.15% | 42.95% | 1427.1bp (1495.8bp) | TBP(- other)/several species/AthaMap(0.794) More Information \| Similar Motifs Found | motif file (matrix) |
| 15 \* | A C T G A C G T A C T G C G T A A G T C A G T C A C T G C G T A | 1e-8 | -1.954e+01 | 2.94% | 1.50% | 1551.4bp (1483.4bp) | RTG3/Literature(Harbison)/Yeast(0.866) More Information \| Similar Motifs Found | motif file (matrix) |
| 16 \* | A C G T C G T A A C T G A C T G A C G T A G T C A C T G A C T G | 1e-7 | -1.779e+01 | 1.18% | 0.41% | 1277.2bp (1254.0bp) | CBF2(AP2EREBP)/colamp-CBF2-DAP-Seq(GSE60143)/Homer(0.785) More Information \| Similar Motifs Found | motif file (matrix) |
| 17 \* | A G T C A C T G A G T C A C G T C G T A A C T G A C G T A C G T | 1e-7 | -1.719e+01 | 1.60% | 0.68% | 1354.5bp (2043.8bp) | STP3/MA0396.1/Jaspar(0.786) More Information \| Similar Motifs Found | motif file (matrix) |
| 18 \* | C G T A C G T A A C G T C G T A C G T A A C G T A G T C A C G T A C G T A C T G | 1e-7 | -1.715e+01 | 1.08% | 0.36% | 1333.0bp (982.8bp) | MATR3(RRM)/Homo\_sapiens-RNCMPT00037-PBM/HughesRNA(0.791) More Information \| Similar Motifs Found | motif file (matrix) |
| 19 \* | A C G T C G T A A C G T A G T C A G T C A G T C A C G T A C T G | 1e-5 | -1.313e+01 | 4.08% | 2.66% | 1064.3bp (1660.6bp) | Pcbp2(KH)/Danio\_rerio-RNCMPT00246-PBM/HughesRNA(0.767) More Information \| Similar Motifs Found | motif file (matrix) |
| 20 \* | A C T G A C G T A G T C A G T C C G T A A G T C C G T A A G T C C G T A A G T C | 1e-5 | -1.249e+01 | 0.62% | 0.18% | 1529.5bp (163.8bp) | Rbm38(RRM)/Danio\_rerio-RNCMPT00283-PBM/HughesRNA(0.803) More Information \| Similar Motifs Found | motif file (matrix) |
| 21 \* | C G T A C G T A C G T A A C G T A G T C C G T A C G T A A G T C A C G T A C T G | 1e-4 | -1.141e+01 | 1.05% | 0.46% | 782.0bp (2066.6bp) | onecut/MA0235.1/Jaspar(0.789) More Information \| Similar Motifs Found | motif file (matrix) |
| 22 \* | C G T A A C G T A C G T A T C G A C T G A C T G C G T A A C T G A C T G A C T G | 1e-4 | -1.098e+01 | 0.82% | 0.33% | 1134.7bp (1691.0bp) | HNRNPH2(RRM)/Homo\_sapiens-RNCMPT00160-PBM/HughesRNA(0.798) More Information \| Similar Motifs Found | motif file (matrix) |
| 23 \* | A C T G C G T A A G T C A G T C A C T G A C T G A C G T A G T C | 1e-4 | -1.061e+01 | 0.65% | 0.23% | 1568.5bp (1765.4bp) | PH0140.1\_Pknox1/Jaspar(0.664) More Information \| Similar Motifs Found | motif file (matrix) |
| 24 \* | A C T G C G T A A C T G C G T A A C G T A G T C A C G T A G T C | 1e-3 | -9.039e+00 | 1.27% | 0.70% | 1495.1bp (1575.2bp) | SRD1/MA0389.1/Jaspar(0.839) More Information \| Similar Motifs Found | motif file (matrix) |
| 25 \* | C G T A A C G T A C T G A C G T C G T A A G T C A C T G C G A T | 1e-3 | -7.981e+00 | 10.81% | 9.02% | 1445.0bp (1445.3bp) | SPL5(SBP)/colamp-SPL5-DAP-Seq(GSE60143)/Homer(0.825) More Information \| Similar Motifs Found | motif file (matrix) |
| 26 \* | A G T C A G T C A G T C C G T A A G T C A G T C A G T C A G T C A G T C A G T C | 1e-2 | -5.865e+00 | 32.60% | 30.28% | 1394.7bp (1557.8bp) | SeqBias: polyC-repeat(0.867) More Information \| Similar Motifs Found | motif file (matrix) |
| 27 \* | A G T C C G T A A C G T C G T A C G T A C G T A A G T C C G T A C G T A A G T C | 1e-2 | -5.163e+00 | 0.78% | 0.47% | 1348.9bp (1222.5bp) | Foxj2/MA0614.1/Jaspar(0.828) More Information \| Similar Motifs Found | motif file (matrix) |
