## Supplemental Tabel 1 for "The Establishment of Cell-Type Specific Gene Regulation in the Sea Urchin Embryo": PMCs_homerResults.html

/data/gpfs-1/users/jbrande\_m/work/scEarlyDev/scATACseq\_analysis/peaksPerLineage/Motifs/PMCs/ - Homer de novo Motif Results


### Homer *de novo* Motif Results (/data/gpfs-1/users/jbrande\_m/work/scEarlyDev/scATACseq\_analysis/peaksPerLineage/Motifs/PMCs/)

Known Motif Enrichment Results  
Gene Ontology Enrichment Results  
If Homer is having trouble matching a motif to a known motif, try copy/pasting the matrix file into
STAMP  
More information on motif finding results: HOMER
| Description of Results
| Tips
  
Total target sequences = 4050  
Total background sequences = 8370  
\* - possible false positive  

|  |  |  |  |  |  |  |  |  |
| --- | --- | --- | --- | --- | --- | --- | --- | --- |
| Rank | Motif | P-value | log P-pvalue | % of Targets | % of Background | STD(Bg STD) | Best Match/Details | Motif File |
| 1 | T A C G C T G A T A G C T C G A A C T G A C T G C G T A C G T A T C A G G A C T | 1e-103 | -2.377e+02 | 57.41% | 40.52% | 1113.1bp (5354.8bp) | ERG(ETS)/VCaP-ERG-ChIP-Seq(GSE14097)/Homer(0.958) More Information \| Similar Motifs Found | motif file (matrix) |
| 2 | T A G C T C G A G A C T C A T G C G T A T A G C G C A T T G A C T C G A A G C T | 1e-70 | -1.630e+02 | 40.35% | 27.34% | 1125.9bp (5112.8bp) | Fra1(bZIP)/BT549-Fra1-ChIP-Seq(GSE46166)/Homer(0.982) More Information \| Similar Motifs Found | motif file (matrix) |
| 3 | A G C T T G C A C G T A G A C T G A C T C T G A T C A G A T G C T C A G T G C A | 1e-36 | -8.337e+01 | 43.78% | 34.18% | 1114.4bp (5564.2bp) | DLX2(Homeobox)/BasalGanglia-Dlx2-ChIP-seq(GSE124936)/Homer(0.970) More Information \| Similar Motifs Found | motif file (matrix) |
| 4 | C T G A A C G T A C G T G T C A C T G A A G T C C T G A A G T C | 1e-17 | -3.983e+01 | 27.93% | 22.16% | 1094.6bp (4490.1bp) | Eomes(T-box)/H9-Eomes-ChIP-Seq(GSE26097)/Homer(0.856) More Information \| Similar Motifs Found | motif file (matrix) |
| 5 \* | T A G C A C T G C T A G C G T A C G A T A C T G A G T C G T A C G A T C A T G C | 1e-11 | -2.565e+01 | 1.53% | 0.57% | 986.0bp (2109.3bp) | RELB/MA1117.1/Jaspar(0.714) More Information \| Similar Motifs Found | motif file (matrix) |
| 6 \* | C G T A A C T G A C T G A T G C A G T C A T G C C G T A A C G T A C T G A C T G | 1e-9 | -2.280e+01 | 0.44% | 0.07% | 734.5bp (300.3bp) | PCF/Arabidopsis-Promoters/Homer(0.707) More Information \| Similar Motifs Found | motif file (matrix) |
| 7 \* | G T A C A C G T A G T C A C G T A C T G A C G T A G T C A G T C A C G T C G T A | 1e-9 | -2.077e+01 | 0.79% | 0.22% | 727.4bp (14718.8bp) | CNOT4(RRM)/Homo\_sapiens-RNCMPT00156-PBM/HughesRNA(0.778) More Information \| Similar Motifs Found | motif file (matrix) |
| 8 \* | C T A G A T C G T C G A C G A T C G T A A C G T A G T C A T G C C G A T C A G T | 1e-8 | -2.072e+01 | 5.21% | 3.38% | 1280.8bp (5236.9bp) | SD0003.1\_at\_AC\_acceptor/Jaspar(0.859) More Information \| Similar Motifs Found | motif file (matrix) |
| 9 \* | A C G T C G T A C G T A A G T C A G T C A C G T A C T G C G T A A G T C C G T A | 1e-8 | -1.998e+01 | 0.44% | 0.08% | 795.0bp (1232.9bp) | SIX1/MA1118.1/Jaspar(0.836) More Information \| Similar Motifs Found | motif file (matrix) |
| 10 \* | A C G T A G T C A C T G A C G T A G T C A C G T A C T G A C G T C G T A A C T G | 1e-7 | -1.686e+01 | 0.30% | 0.04% | 455.9bp (879.7bp) | Smad4(MAD)/ESC-SMAD4-ChIP-Seq(GSE29422)/Homer(0.663) More Information \| Similar Motifs Found | motif file (matrix) |
| 11 \* | C A G T C T A G C A T G T A C G A C T G G T C A T G C A C T A G C A T G A T C G | 1e-5 | -1.340e+01 | 21.43% | 18.52% | 1129.6bp (8959.6bp) | ZNF467(Zf)/HEK293-ZNF467.GFP-ChIP-Seq(GSE58341)/Homer(0.889) More Information \| Similar Motifs Found | motif file (matrix) |
| 12 \* | C G T A A C T G C G T A A C G T C G T A A G T C A C T G C G T A C G T A A C G T | 1e-3 | -8.449e+00 | 0.30% | 0.09% | 818.3bp (5068.3bp) | ARR18/MA0948.1/Jaspar(0.810) More Information \| Similar Motifs Found | motif file (matrix) |
| 13 \* | A C G T A G T C A G T C A G T C C G T A C T G A C T A G C T G A A C G T C G T A | 1e-2 | -4.833e+00 | 3.19% | 2.57% | 1128.5bp (9283.6bp) | Ebf2/MA1604.1/Jaspar(0.692) More Information \| Similar Motifs Found | motif file (matrix) |
| 14 \* | A C G T G T A C A G T C C G T A A C G T C T A G A C G T C T G A | 1e-1 | -2.641e+00 | 7.51% | 6.91% | 1289.5bp (6004.6bp) | MXI1/MA1108.2/Jaspar(0.778) More Information \| Similar Motifs Found | motif file (matrix) |
| 15 \* | C G T A A C T G A C T G A C G T A C T G A C G T | 1e0 | -2.209e+00 | 38.64% | 37.70% | 1163.3bp (5632.6bp) | TBX5/MA0807.1/Jaspar(0.917) More Information \| Similar Motifs Found | motif file (matrix) |
| 16 \* | A C T G C G T A C G T A A C T G A G T C C G T A | 1e0 | -4.045e-02 | 36.42% | 37.75% | 1186.3bp (5689.2bp) | POL008.1\_DCE\_S\_I/Jaspar(0.856) More Information \| Similar Motifs Found | motif file (matrix) |
| 17 \* | A G T C A G T C A G T C A G T C A T G C A C G T | 1e0 | -2.418e-02 | 46.67% | 48.21% | 1168.1bp (5594.6bp) | msn-1/MA1433.1/Jaspar(0.918) More Information \| Similar Motifs Found | motif file (matrix) |
| 18 \* | A G C T A G T C A C G T A C G T A C G T A G T C | 1e0 | -0.000e+00 | 68.86% | 72.38% | 1213.0bp (5538.8bp) | br-Z4/dmmpmm(Bergman)/fly(0.853) More Information \| Similar Motifs Found | motif file (matrix) |
