## Supplemental Tabel 1 for "The Establishment of Cell-Type Specific Gene Regulation in the Sea Urchin Embryo": Unspecified_homerResults.html

/data/gpfs-1/users/jbrande\_m/work/scEarlyDev/scATACseq\_analysis/peaksPerLineage/Motifs/Unspecified/ - Homer de novo Motif Results


### Homer *de novo* Motif Results (/data/gpfs-1/users/jbrande\_m/work/scEarlyDev/scATACseq\_analysis/peaksPerLineage/Motifs/Unspecified/)

Known Motif Enrichment Results  
Gene Ontology Enrichment Results  
If Homer is having trouble matching a motif to a known motif, try copy/pasting the matrix file into
STAMP  
More information on motif finding results: HOMER
| Description of Results
| Tips
  
Total target sequences = 147  
Total background sequences = 10222  
\* - possible false positive  

|  |  |  |  |  |  |  |  |  |
| --- | --- | --- | --- | --- | --- | --- | --- | --- |
| Rank | Motif | P-value | log P-pvalue | % of Targets | % of Background | STD(Bg STD) | Best Match/Details | Motif File |
| 1 \* | A G T C A G T C C G T A A C T G C G T A A G T C C G T A A C T G A C G T A G T C | 1e-8 | -1.934e+01 | 4.08% | 0.09% | 448.7bp (2079.4bp) | sma-4/MA0925.1/Jaspar(0.832) More Information \| Similar Motifs Found | motif file (matrix) |
| 2 \* | A G T C A G T C A G T C A C T G A C T G C G T A | 1e-7 | -1.842e+01 | 36.73% | 17.11% | 2315.6bp (2487.1bp) | CAT8/MA0280.1/Jaspar(0.891) More Information \| Similar Motifs Found | motif file (matrix) |
| 3 \* | C G T A A C G T A C T G A C T G C G T A A C G T A C T G A C T G C G T A A C G T | 1e-7 | -1.811e+01 | 8.84% | 1.11% | 916.3bp (1010.3bp) | SRSF9(RRM)/Homo\_sapiens-RNCMPT00067-PBM/HughesRNA(0.704) More Information \| Similar Motifs Found | motif file (matrix) |
| 4 \* | A C T G A G T C A C T G A C T G A C T G C G T A A T G C A C G T A G T C A G T C | 1e-6 | -1.601e+01 | 2.72% | 0.03% | 2319.1bp (963.8bp) | UGA3/MA0410.1/Jaspar(0.735) More Information \| Similar Motifs Found | motif file (matrix) |
| 5 \* | A C T G A G T C A G T C A G T C A T G C A G T C A C T G A C G T | 1e-5 | -1.277e+01 | 12.24% | 3.39% | 531.4bp (3187.7bp) | PB0010.1\_Egr1\_1/Jaspar(0.754) More Information \| Similar Motifs Found | motif file (matrix) |
| 6 \* | G T C A A C G T A C G T A G T C A C T G C G T A A C T G A G T C | 1e-5 | -1.271e+01 | 14.29% | 4.51% | 736.2bp (3872.0bp) | XBP1/Literature(Harbison)/Yeast(0.784) More Information \| Similar Motifs Found | motif file (matrix) |
| 7 \* | A C T G A C T G A G T C A G T C A C T G A C G T A C T G A C G T A G T C A C G T | 1e-5 | -1.262e+01 | 2.04% | 0.03% | 2537.0bp (535.3bp) | ARG80/MA0271.1/Jaspar(0.655) More Information \| Similar Motifs Found | motif file (matrix) |
| 8 \* | C T A G C G T A T C G A T C A G C T G A T A C G C A G T A C T G A C G T A G C T | 1e-5 | -1.224e+01 | 31.29% | 16.28% | 1515.7bp (2994.5bp) | AT2G40260(G2like)/colamp-AT2G40260-DAP-Seq(GSE60143)/Homer(0.647) More Information \| Similar Motifs Found | motif file (matrix) |
| 9 \* | A C G T C T G A G T A C G T A C T C G A A T G C G T C A C A T G T C A G T G C A | 1e-5 | -1.213e+01 | 29.25% | 14.78% | 676.7bp (4339.3bp) | MET28/MA0332.1/Jaspar(0.773) More Information \| Similar Motifs Found | motif file (matrix) |
| 10 \* | G T A C T G A C T C G A C G T A A G T C A T C G T C G A T A C G | 1e-4 | -1.016e+01 | 21.77% | 10.36% | 2091.8bp (2517.8bp) | HAP2(MacIsaac)/Yeast(0.831) More Information \| Similar Motifs Found | motif file (matrix) |
| 11 \* | C T G A G T A C A G T C C G T A C G T A A G T C A G T C C G T A C G T A A G T C | 1e-4 | -9.753e+00 | 4.76% | 0.65% | 466.8bp (2015.0bp) | MYB96(MYB)/colamp-MYB96-DAP-Seq(GSE60143)/Homer(0.787) More Information \| Similar Motifs Found | motif file (matrix) |
| 12 \* | A T C G G T C A C T G A A C G T C T G A C T G A A C T G C T G A T C G A A C T G | 1e-4 | -9.555e+00 | 14.97% | 6.01% | 1010.0bp (10299.7bp) | PEND/MA0127.1/Jaspar(0.863) More Information \| Similar Motifs Found | motif file (matrix) |
| 13 \* | C G T A A G T C A G T C C T A G A G C T A C T G A C T G A G C T G T C A A G C T | 1e-4 | -9.217e+00 | 68.03% | 52.55% | 1563.3bp (3500.5bp) | Tv\_0259(RRM)/Trichomonas\_vaginalis-RNCMPT00259-PBM/HughesRNA(0.639) More Information \| Similar Motifs Found | motif file (matrix) |
| 14 \* | A C T G A C G T G T A C A G T C A G T C A G T C C G A T A C T G A G T C A G T C | 1e-3 | -8.606e+00 | 2.72% | 0.19% | 2502.0bp (881.8bp) | CTCFL/MA1102.2/Jaspar(0.710) More Information \| Similar Motifs Found | motif file (matrix) |
| 15 \* | A C T G A C G T A C T G A C G T C T A G A G C T A G T C A G T C | 1e-3 | -8.555e+00 | 13.61% | 5.55% | 841.2bp (2800.1bp) | SM(RRM)/Drosophila\_melanogaster-RNCMPT00069-PBM/HughesRNA(0.808) More Information \| Similar Motifs Found | motif file (matrix) |
| 16 \* | A C T G A C T G A G T C A C T G C G T A C G T A C T A G T A C G A C T G A G T C | 1e-3 | -8.076e+00 | 3.40% | 0.40% | 702.7bp (7048.9bp) | ZNF467(Zf)/HEK293-ZNF467.GFP-ChIP-Seq(GSE58341)/Homer(0.726) More Information \| Similar Motifs Found | motif file (matrix) |
| 17 \* | C G T A A G T C C T A G A G T C G T A C C T A G A G T C A G T C C G A T G T C A | 1e-3 | -7.969e+00 | 3.40% | 0.40% | 265.8bp (2389.7bp) | CRF4(AP2EREBP)/colamp-CRF4-DAP-Seq(GSE60143)/Homer(0.765) More Information \| Similar Motifs Found | motif file (matrix) |
| 18 \* | C G T A A C G T A C T G C G T A C G T A A C G T C G T A A C T G | 1e-3 | -7.345e+00 | 12.93% | 5.64% | 1686.1bp (3507.0bp) | Brn2(POU,Homeobox)/NPC-Brn2-ChIP-Seq(GSE35496)/Homer(0.793) More Information \| Similar Motifs Found | motif file (matrix) |
| 19 \* | A G C T C G A T A C G T A G T C A G T C A G T C C G A T A G C T G A C T G A C T | 1e-3 | -7.063e+00 | 31.97% | 20.65% | 1670.3bp (4816.0bp) | NDD1(MacIsaac)/Yeast(0.871) More Information \| Similar Motifs Found | motif file (matrix) |
| 20 \* | A C G T C G T A A G T C A G T C A T G C A G T C A C T G C G T A C G T A A C T G | 1e-2 | -6.105e+00 | 1.36% | 0.05% | 460.6bp (659.5bp) | ELF2/MA1483.1/Jaspar(0.720) More Information \| Similar Motifs Found | motif file (matrix) |
| 21 \* | C T A G A G C T C T A G A T C G A C T G C G A T T A C G A C G T | 1e-2 | -5.606e+00 | 20.41% | 12.33% | 1015.3bp (2545.0bp) | Aef1/dmmpmm(Pollard)/fly(0.770) More Information \| Similar Motifs Found | motif file (matrix) |
| 22 \* | A G T C A G T C C G T A A G T C A C G T A G T C A C T G A C T G A G T C C G T A | 1e-2 | -5.598e+00 | 94.56% | 87.52% | 1347.6bp (3284.8bp) | PB0114.1\_Egr1\_2/Jaspar(0.696) More Information \| Similar Motifs Found | motif file (matrix) |
| 23 \* | A C T G C G T A A G T C A C T G A C T G A G T C C G T A A C G T | 1e-2 | -5.391e+00 | 4.76% | 1.39% | 231.3bp (2978.7bp) | TGA10(bZIP)/colamp-TGA10-DAP-Seq(GSE60143)/Homer(0.797) More Information \| Similar Motifs Found | motif file (matrix) |
| 24 \* | A C G T A G T C A C T G A G T C A G T C A C T G C G T A A G T C C G T A A G T C | 1e-2 | -5.140e+00 | 57.14% | 46.43% | 1298.0bp (2922.9bp) | ERF039/MA0995.2/Jaspar(0.846) More Information \| Similar Motifs Found | motif file (matrix) |
| 25 \* | A G T C A G T C C G T A A C G T A G T C A C G T A G T C A C G T A C T G A G T C | 1e-2 | -4.763e+00 | 1.36% | 0.10% | 1188.8bp (3989.1bp) | ARF27/MA1691.1/Jaspar(0.728) More Information \| Similar Motifs Found | motif file (matrix) |
| 26 \* | G T A C C G T A C A G T G A C T G T A C T A G C G T C A A G C T G T A C A G T C | 1e-2 | -4.759e+00 | 30.61% | 21.88% | 1616.9bp (3903.6bp) | TEAD2/MA1121.1/Jaspar(0.749) More Information \| Similar Motifs Found | motif file (matrix) |
| 27 \* | G A C T C G T A G A C T A G T C C T G A A G T C A G T C A G T C | 1e-2 | -4.703e+00 | 68.03% | 58.20% | 975.0bp (3166.1bp) | AFT2/AFT2\_H2O2Lo/10-RCS1[~AFT2](Harbison)/Yeast(0.883) More Information \| Similar Motifs Found | motif file (matrix) |
| 28 \* | C G T A C G T A C G T A A C T G C G T A C G T A | 1e-1 | -2.913e+00 | 61.22% | 54.32% | 1441.1bp (3976.3bp) | AZF1/MA0277.1/Jaspar(0.843) More Information \| Similar Motifs Found | motif file (matrix) |
| 29 \* | A G T C C G T A A G T C A G T C T G C A A G T C A C G T A G T C A C G T A G T C | 1e0 | -2.203e+00 | 10.88% | 7.81% | 1381.6bp (2086.1bp) | Trl(Zf)/S2-GAGAfactor-ChIP-Seq(GSE40646)/Homer(0.708) More Information \| Similar Motifs Found | motif file (matrix) |
| 30 \* | A C G T A G T C A C G T A C T G A G T C C G T A A C T G C G T A | 1e0 | -2.107e+00 | 3.40% | 1.78% | 1239.9bp (2812.3bp) | SOK2/MA0385.1/Jaspar(0.710) More Information \| Similar Motifs Found | motif file (matrix) |
| 31 \* | C G T A A G T C C G T A A G T C A G T C A C T G A C T G A G C T | 1e0 | -2.049e+00 | 5.44% | 3.40% | 1491.8bp (2239.4bp) | Ptf1a(var.3)/MA1620.1/Jaspar(0.786) More Information \| Similar Motifs Found | motif file (matrix) |
