## Supplemental Table 1 for "The Establishment of Cell-Type Specific Gene Regulation in the Sea Urchin Embryo": Hpf10_EctoAnimal_homerResults.html

homer\_motifs\_wbackground/128cell\_EctoAnimal/ - Homer de novo Motif Results


### Homer *de novo* Motif Results (homer\_motifs\_wbackground/128cell\_EctoAnimal/)

Known Motif Enrichment Results  
Gene Ontology Enrichment Results  
If Homer is having trouble matching a motif to a known motif, try copy/pasting the matrix file into
STAMP  
More information on motif finding results: HOMER
| Description of Results
| Tips
  
Total target sequences = 73  
Total background sequences = 586  
\* - possible false positive  

|  |  |  |  |  |  |  |  |  |
| --- | --- | --- | --- | --- | --- | --- | --- | --- |
| Rank | Motif | P-value | log P-pvalue | % of Targets | % of Background | STD(Bg STD) | Best Match/Details | Motif File |
| 1 | T C G A C G A T T A C G A C T G G T A C G A C T G A T C G T A C G T C A C G T A | 1e-12 | -2.854e+01 | 23.29% | 2.32% | 403.8bp (351.8bp) | MYB88(MYB)/col-MYB88-DAP-Seq(GSE60143)/Homer(0.650) More Information \| Similar Motifs Found | motif file (matrix) |
| 2 | A C T G A C G T A C T G C G T A A G C T A T C G A T G C G T A C A G T C A C G T | 1e-12 | -2.778e+01 | 10.96% | 0.19% | 341.4bp (0.0bp) | RARB(var.3)/MA1552.1/Jaspar(0.700) More Information \| Similar Motifs Found | motif file (matrix) |
| 3 \* | A G T C G T A C C G A T C G T A A C G T A C G T C G T A C G A T G T A C C T A G | 1e-11 | -2.636e+01 | 15.07% | 0.76% | 296.4bp (312.9bp) | DAL80/Literature(Harbison)/Yeast(0.743) More Information \| Similar Motifs Found | motif file (matrix) |
| 4 \* | G T C A A G T C C G T A A T C G C G T A G T C A A G T C A C G T G T A C C G T A | 1e-11 | -2.603e+01 | 12.33% | 0.48% | 356.7bp (170.6bp) | ZNF274/MA1592.1/Jaspar(0.722) More Information \| Similar Motifs Found | motif file (matrix) |
| 5 \* | A C G T A G T C A T G C T G A C A C G T A G T C G T A C T A C G A C G T A C G T | 1e-11 | -2.579e+01 | 19.18% | 1.68% | 427.0bp (430.8bp) | B52(RRM)/Drosophila\_melanogaster-RNCMPT00134-PBM/HughesRNA(0.705) More Information \| Similar Motifs Found | motif file (matrix) |
| 6 \* | A C T G A C T G A G T C C G T A A G C T A G T C C T A G C G A T A G T C A G T C | 1e-10 | -2.351e+01 | 9.59% | 0.10% | 463.4bp (0.0bp) | AGL42/MA1201.1/Jaspar(0.665) More Information \| Similar Motifs Found | motif file (matrix) |
| 7 \* | C G T A C G T A G T C A A C T G C G T A C T A G C G T A A G T C C G A T C G T A | 1e-9 | -2.233e+01 | 10.96% | 0.37% | 415.0bp (399.3bp) | SGR5(C2H2)/colamp-SGR5-DAP-Seq(GSE60143)/Homer(0.670) More Information \| Similar Motifs Found | motif file (matrix) |
| 8 \* | C G T A A C G T C G T A A G T C C G T A A C T G A G T C A C G T A G T C A G T C | 1e-9 | -2.099e+01 | 13.70% | 0.91% | 350.3bp (572.3bp) | POL013.1\_MED-1/Jaspar(0.644) More Information \| Similar Motifs Found | motif file (matrix) |
| 9 \* | A G T C C T G A C G T A A C T G A G T C A C T G C G T A C T A G C G T A A C T G | 1e-8 | -1.940e+01 | 8.22% | 0.30% | 116.2bp (0.0bp) | Clamp/MA1700.1/Jaspar(0.767) More Information \| Similar Motifs Found | motif file (matrix) |
| 10 \* | A C G T C G T A C G T A G A T C A C G T A C T G A G T C A C G T G T A C A G C T | 1e-8 | -1.926e+01 | 13.70% | 1.15% | 267.6bp (460.2bp) | MYB77(MYB)/col-MYB77-DAP-Seq(GSE60143)/Homer(0.709) More Information \| Similar Motifs Found | motif file (matrix) |
| 11 \* | A C G T A C T G A G T C A C G T C G T A C G T A A T G C A G T C | 1e-8 | -1.921e+01 | 19.18% | 2.71% | 332.4bp (432.9bp) | PB0029.1\_Hic1\_1/Jaspar(0.761) More Information \| Similar Motifs Found | motif file (matrix) |
| 12 \* | G T C A A T G C C A T G G T C A A C T G A C G T C A G T C A T G A C G T G A T C | 1e-8 | -1.919e+01 | 10.96% | 0.63% | 486.2bp (108.7bp) | MGP(C2H2)/colamp-MGP-DAP-Seq(GSE60143)/Homer(0.652) More Information \| Similar Motifs Found | motif file (matrix) |
| 13 \* | A C G T A C G T A G T C C G T A A T G C A C G T A G T C A C T G A G T C C G T A | 1e-8 | -1.919e+01 | 10.96% | 0.60% | 269.8bp (54.8bp) | AT3G46070/MA1381.1/Jaspar(0.778) More Information \| Similar Motifs Found | motif file (matrix) |
| 14 \* | A C G T A G T C A G T C A C G T A C G T A C G T A T C G C G A T A G T C C A G T | 1e-7 | -1.824e+01 | 16.44% | 1.92% | 491.3bp (486.8bp) | PB0166.1\_Sox12\_2/Jaspar(0.862) More Information \| Similar Motifs Found | motif file (matrix) |
| 15 \* | A G T C C G T A C G T A A C G T C G T A C G T A A C G T A C T G | 1e-7 | -1.803e+01 | 26.03% | 5.68% | 581.9bp (579.1bp) | Tv\_0226(RRM)/Trichomonas\_vaginalis-RNCMPT00226-PBM/HughesRNA(0.887) More Information \| Similar Motifs Found | motif file (matrix) |
| 16 \* | A G T C A C G T A C T G G C A T G T C A A G C T A C T G A C T G | 1e-7 | -1.756e+01 | 26.03% | 5.95% | 381.5bp (430.1bp) | RAP1/MA0359.1/Jaspar(0.781) More Information \| Similar Motifs Found | motif file (matrix) |
| 17 \* | A C T G C G T A A C G T A C G T C G T A C G T A T A G C A C G T A G T C A G T C | 1e-7 | -1.698e+01 | 10.96% | 0.72% | 321.6bp (402.6bp) | PH0124.1\_Obox5\_1/Jaspar(0.753) More Information \| Similar Motifs Found | motif file (matrix) |
| 18 \* | C G A T G T A C G C T A C T G A G T A C G C T A C A T G C T A G C G T A G A T C | 1e-7 | -1.687e+01 | 15.07% | 1.83% | 309.3bp (486.7bp) | Foxo1/MA0480.1/Jaspar(0.754) More Information \| Similar Motifs Found | motif file (matrix) |
| 19 \* | G T A C A T C G A C T G A G T C C G T A A G T C C G T A C T G A A C G T A C T G | 1e-6 | -1.602e+01 | 9.59% | 0.62% | 349.6bp (503.0bp) | MATA1(MacIsaac)/Yeast(0.820) More Information \| Similar Motifs Found | motif file (matrix) |
| 20 \* | A C G T C G T A A C T G A C T G A C G T C G T A A G T C C G T A | 1e-6 | -1.550e+01 | 13.70% | 1.64% | 550.5bp (297.6bp) | MYB121(MYB)/col-MYB121-DAP-Seq(GSE60143)/Homer(0.797) More Information \| Similar Motifs Found | motif file (matrix) |
| 21 \* | G C T A A C T G T G C A C G A T G T A C A G T C T A G C A C T G C T A G G T A C | 1e-6 | -1.533e+01 | 8.22% | 0.41% | 401.4bp (378.1bp) | GAT4/MA0302.1/Jaspar(0.667) More Information \| Similar Motifs Found | motif file (matrix) |
| 22 \* | C G A T A C G T A G T C A C G T A C T G A C T G A C G T A G T C | 1e-5 | -1.301e+01 | 16.44% | 3.17% | 382.2bp (477.2bp) | ACE2/ACE2\_YPD/2-SWI5(Harbison)/Yeast(0.737) More Information \| Similar Motifs Found | motif file (matrix) |
| 23 \* | A C G T A C T G A C G T A C T G C G T A A C G T A C G T A C G T A C G T A C T G | 1e-5 | -1.234e+01 | 12.33% | 1.80% | 259.1bp (259.8bp) | tll/dmmpmm(Papatsenko)/fly(0.738) More Information \| Similar Motifs Found | motif file (matrix) |
| 24 \* | A G T C A G T C A C G T A G T C C G T A A C T G C T G A A C G T | 1e-4 | -1.027e+01 | 12.33% | 2.29% | 430.7bp (721.8bp) | MET4(MacIsaac)/Yeast(0.736) More Information \| Similar Motifs Found | motif file (matrix) |
| 25 \* | C G T A A C T G A G T C A G T C A G T C A C G T C G T A A G T C | 1e-3 | -9.181e+00 | 12.33% | 2.72% | 495.7bp (403.0bp) | HRB98DE(RRM)/Drosophila\_melanogaster-RNCMPT00096-PBM/HughesRNA(0.822) More Information \| Similar Motifs Found | motif file (matrix) |
| 26 \* | A C G T C T G A C T A G A C G T A C T G A C G T C T G A A C G T C G T A A C T G | 1e-3 | -9.133e+00 | 8.22% | 1.07% | 205.7bp (383.8bp) | PUM(PUF)/Drosophila\_melanogaster-RNCMPT00102-PBM/HughesRNA(0.703) More Information \| Similar Motifs Found | motif file (matrix) |
| 27 \* | A C T G A G T C T A C G A C T G A G T C A G T C A C T G A G T C A G T C A C T G | 1e-3 | -9.015e+00 | 5.48% | 0.42% | 371.1bp (659.3bp) | ERF094/MA1049.1/Jaspar(0.821) More Information \| Similar Motifs Found | motif file (matrix) |
| 28 \* | C G T A A G T C A C G T A C T G A C T G C G T A A G T C A G T C | 1e-3 | -7.784e+00 | 6.85% | 0.95% | 359.2bp (189.9bp) | NRG1(MacIsaac)/Yeast(0.699) More Information \| Similar Motifs Found | motif file (matrix) |
| 29 \* | G C T A A C G T C G T A C G T A A C G T A G T C A G T C A C T G A C G T G T C A | 1e-3 | -6.968e+00 | 6.85% | 1.05% | 223.1bp (240.9bp) | dve/MA0915.1/Jaspar(0.863) More Information \| Similar Motifs Found | motif file (matrix) |
| 30 \* | A C T G A C T G A G T C A C G T A C T G A C T G A G T C A C G T | 1e-1 | -4.073e+00 | 6.85% | 2.14% | 317.2bp (536.5bp) | Vts1p(SAM)/Saccharomyces\_cerevisiae-RNCMPT00111-PBM/HughesRNA(0.795) More Information \| Similar Motifs Found | motif file (matrix) |
| 31 \* | A C T G A C G T A C T G A C G T A C T G A C G T A T C G A C G T A T C G A C G T | 1e-1 | -4.041e+00 | 12.33% | 5.46% | 603.4bp (276.8bp) | SeqBias: CA-repeat(0.891) More Information \| Similar Motifs Found | motif file (matrix) |
