## Supplemental Table 1 for "The Establishment of Cell-Type Specific Gene Regulation in the Sea Urchin Embryo": Hpf10_EctoApical_homerResults.html

homer\_motifs\_wbackground/128cell\_EctoApical/ - Homer de novo Motif Results


### Homer *de novo* Motif Results (homer\_motifs\_wbackground/128cell\_EctoApical/)

Known Motif Enrichment Results  
Gene Ontology Enrichment Results  
If Homer is having trouble matching a motif to a known motif, try copy/pasting the matrix file into
STAMP  
More information on motif finding results: HOMER
| Description of Results
| Tips
  
Total target sequences = 167  
Total background sequences = 492  
\* - possible false positive  

|  |  |  |  |  |  |  |  |  |
| --- | --- | --- | --- | --- | --- | --- | --- | --- |
| Rank | Motif | P-value | log P-pvalue | % of Targets | % of Background | STD(Bg STD) | Best Match/Details | Motif File |
| 1 | A C T G A C T G C T A G C G T A A C G T A C T G G A T C C G T A A C G T A C G T | 1e-16 | -3.737e+01 | 7.78% | 0.24% | 377.0bp (0.0bp) | RCS1/RCS1\_H2O2Hi/35-RCS1(Harbison)/Yeast(0.741) More Information \| Similar Motifs Found | motif file (matrix) |
| 2 | A C T G A T C G A C G T T G A C A C G T A T C G A C T G C G A T A C G T A C G T | 1e-16 | -3.737e+01 | 7.78% | 0.35% | 211.1bp (171.4bp) | MET31(MacIsaac)/Yeast(0.686) More Information \| Similar Motifs Found | motif file (matrix) |
| 3 | T C A G C G A T A G T C C G T A C G A T A C T G A C T G A C G T A C G T A C G T | 1e-13 | -3.199e+01 | 9.58% | 0.77% | 374.2bp (253.5bp) | Tv\_0259(RRM)/Trichomonas\_vaginalis-RNCMPT00259-PBM/HughesRNA(0.771) More Information \| Similar Motifs Found | motif file (matrix) |
| 4 | A G T C C G T A A C T G C A T G G A T C C G A T A G T C A G T C A T G C A T C G | 1e-13 | -3.175e+01 | 8.38% | 0.54% | 350.5bp (559.7bp) | ZNF460/MA1596.1/Jaspar(0.705) More Information \| Similar Motifs Found | motif file (matrix) |
| 5 | A C T G G T C A A C T G T G A C A C T G C G T A C T G A C G T A C G T A C G T A | 1e-13 | -3.051e+01 | 16.77% | 3.03% | 403.9bp (568.0bp) | MAC1(MacIsaac)/Yeast(0.834) More Information \| Similar Motifs Found | motif file (matrix) |
| 6 \* | A C T G A C G T C T G A A C G T A C G T C T A G G T A C A G T C C T A G C G T A | 1e-11 | -2.647e+01 | 5.99% | 0.35% | 325.2bp (138.9bp) | slbo/dmmpmm(Bergman)/fly(0.707) More Information \| Similar Motifs Found | motif file (matrix) |
| 7 \* | A C T G A C T G A G T C G T A C A C G T C A G T A T C G C G A T A C T G G T C A | 1e-11 | -2.600e+01 | 13.77% | 2.28% | 320.6bp (150.0bp) | ZNF711(Zf)/SHSY5Y-ZNF711-ChIP-Seq(GSE20673)/Homer(0.716) More Information \| Similar Motifs Found | motif file (matrix) |
| 8 \* | C G A T G T A C A T C G G T C A A C G T C T G A A C T G G T A C A G C T C G T A | 1e-10 | -2.367e+01 | 7.78% | 0.69% | 379.3bp (294.7bp) | pnr/MA0536.1/Jaspar(0.761) More Information \| Similar Motifs Found | motif file (matrix) |
| 9 \* | C T G A G T A C C T G A A T C G C T A G A C T G A G T C T G A C A G T C G C T A | 1e-10 | -2.315e+01 | 17.96% | 4.57% | 329.0bp (505.7bp) | OsI\_08196/MA1050.1/Jaspar(0.774) More Information \| Similar Motifs Found | motif file (matrix) |
| 10 \* | G T C A T G C A C T A G G T C A T G A C C T A G T A G C T C A G A C G T G A T C | 1e-9 | -2.268e+01 | 6.59% | 0.57% | 268.4bp (670.0bp) | MBP1(MacIsaac)/Yeast(0.847) More Information \| Similar Motifs Found | motif file (matrix) |
| 11 \* | G T A C T A C G A T C G G C A T C G T A G A C T C T G A T C A G C G T A T C A G | 1e-9 | -2.194e+01 | 11.98% | 2.13% | 375.7bp (408.3bp) | RBMS1(RRM)/Homo\_sapiens-RNCMPT00152-PBM/HughesRNA(0.651) More Information \| Similar Motifs Found | motif file (matrix) |
| 12 \* | A T C G C A G T C A T G C A T G C G A T C G T A A T G C G A T C G C T A C G A T | 1e-9 | -2.105e+01 | 7.19% | 0.72% | 259.1bp (198.8bp) | PTF1(TCP)/colamp-PTF1-DAP-Seq(GSE60143)/Homer(0.659) More Information \| Similar Motifs Found | motif file (matrix) |
| 13 \* | G T A C T C G A A C T G A C G T C G T A T A G C G T A C A G T C A G T C A C G T | 1e-9 | -2.105e+01 | 7.19% | 0.73% | 405.5bp (657.7bp) | RGM1/MA0366.1/Jaspar(0.714) More Information \| Similar Motifs Found | motif file (matrix) |
| 14 \* | A G T C C G A T A G T C A C T G G A C T A T G C C G T A C G T A C G T A A C T G | 1e-9 | -2.105e+01 | 7.19% | 0.80% | 283.1bp (144.9bp) | TCF7/MA0769.2/Jaspar(0.768) More Information \| Similar Motifs Found | motif file (matrix) |
| 15 \* | G T A C T A C G A C T G C G T A A C T G C G T A A C T G C A T G A C G T A C G T | 1e-9 | -2.105e+01 | 7.19% | 0.62% | 208.7bp (455.8bp) | PDR8/MA0354.1/Jaspar(0.706) More Information \| Similar Motifs Found | motif file (matrix) |
| 16 \* | A C G T C G T A A C T G C G T A A C G T A G C T C G T A A G T C | 1e-9 | -2.084e+01 | 19.76% | 6.09% | 479.8bp (520.2bp) | GATA1/MA0035.4/Jaspar(0.743) More Information \| Similar Motifs Found | motif file (matrix) |
| 17 \* | A T G C A G T C C T G A C G T A A C T G C A T G C G A T C G T A C T G A G T C A | 1e-8 | -2.022e+01 | 7.78% | 0.96% | 280.4bp (355.2bp) | NF1:FOXA1(CTF,Forkhead)/LNCAP-FOXA1-ChIP-Seq(GSE27824)/Homer(0.737) More Information \| Similar Motifs Found | motif file (matrix) |
| 18 \* | C G T A A C G T C G T A A T G C C G T A A C T G C G T A G T A C C G T A A C T G | 1e-7 | -1.708e+01 | 5.39% | 0.51% | 215.4bp (616.0bp) | ARF34/MA1693.1/Jaspar(0.774) More Information \| Similar Motifs Found | motif file (matrix) |
| 19 \* | C A G T T A G C A G C T A C T G G T C A G A C T T G A C T C A G G C A T T G C A | 1e-7 | -1.708e+01 | 5.39% | 0.59% | 242.1bp (131.1bp) | DAL82/DAL82\_SM/3-DAL82(Harbison)/Yeast(0.785) More Information \| Similar Motifs Found | motif file (matrix) |
| 20 \* | C G T A A G T C C T A G A G T C A G T C A C G T A C T G A C T G | 1e-7 | -1.637e+01 | 11.98% | 2.89% | 453.2bp (454.8bp) | RAV1(2)(AP2/EREBP)/Arabidopsis thaliana/AthaMap(0.815) More Information \| Similar Motifs Found | motif file (matrix) |
| 21 \* | A C T G A G T C A G T C A C T G A C T G A C G T C G T A C G T A A C T G A C T G | 1e-6 | -1.607e+01 | 5.99% | 0.64% | 316.8bp (215.0bp) | PCBP1(KH)/Homo\_sapiens-RNCMPT00186-PBM/HughesRNA(0.744) More Information \| Similar Motifs Found | motif file (matrix) |
| 22 \* | T C A G C G T A G A C T T C G A A T G C A G T C G C A T A G T C C G T A A C T G | 1e-6 | -1.550e+01 | 7.19% | 1.10% | 320.8bp (201.4bp) | RUNX3/MA0684.2/Jaspar(0.713) More Information \| Similar Motifs Found | motif file (matrix) |
| 23 \* | A C T G T A C G C A T G T G C A G T C A A T G C T A G C C T A G | 1e-6 | -1.550e+01 | 7.19% | 1.09% | 376.2bp (413.3bp) | OPI1/MA0349.1/Jaspar(0.860) More Information \| Similar Motifs Found | motif file (matrix) |
| 24 \* | A G C T C T G A A T C G T C A G A T C G G C A T C G A T A C T G G A T C C T A G | 1e-6 | -1.445e+01 | 4.79% | 0.49% | 297.1bp (339.3bp) | MUB(KH)/Drosophila\_melanogaster-RNCMPT00137-PBM/HughesRNA(0.760) More Information \| Similar Motifs Found | motif file (matrix) |
| 25 \* | A G C T A C G T A C T G A G T C A G T C A C T G A C T G C G T A | 1e-5 | -1.372e+01 | 5.39% | 0.81% | 361.3bp (445.4bp) | LBD18/MA1673.1/Jaspar(0.823) More Information \| Similar Motifs Found | motif file (matrix) |
| 26 \* | A G T C A C T G A C G T C G T A A C T G A G T C C G T A A C T G | 1e-5 | -1.288e+01 | 10.18% | 2.72% | 383.1bp (301.4bp) | POL013.1\_MED-1/Jaspar(0.715) More Information \| Similar Motifs Found | motif file (matrix) |
| 27 \* | A G T C A G T C A C G T C T G A A G T C A C T G A G T C G T C A | 1e-5 | -1.237e+01 | 7.78% | 1.78% | 467.2bp (440.6bp) | SRP54(RRM)/Drosophila\_melanogaster-RNCMPT00272-PBM/HughesRNA(0.750) More Information \| Similar Motifs Found | motif file (matrix) |
| 28 \* | A C T G A C T G C G T A A C G T A C G T A C G T A C T G A C G T | 1e-5 | -1.237e+01 | 7.78% | 1.65% | 353.0bp (464.9bp) | sd/dmmpmm(Bergman)/fly(0.738) More Information \| Similar Motifs Found | motif file (matrix) |
| 29 \* | A T C G C G A T C G T A G T C A A T C G A G T C C G T A C G T A | 1e-4 | -1.150e+01 | 48.50% | 32.47% | 354.6bp (447.8bp) | FOXN3/MA1489.1/Jaspar(0.793) More Information \| Similar Motifs Found | motif file (matrix) |
| 30 \* | A C G T C G T A C G T A C G T A A G T C C G T A A C T G A C T G | 1e-4 | -1.071e+01 | 10.78% | 3.55% | 366.8bp (352.2bp) | Foxo1/MA0480.1/Jaspar(0.877) More Information \| Similar Motifs Found | motif file (matrix) |
| 31 \* | A C G T A G T C C G T A A C G T A G T C A C T G A G T C A C G T | 1e-4 | -1.035e+01 | 6.59% | 1.60% | 389.7bp (353.2bp) | CHA4(MacIsaac)/Yeast(0.825) More Information \| Similar Motifs Found | motif file (matrix) |
| 32 \* | C G T A A C G T C G T A A C G T G T C A A C G T C G T A A G C T G T C A A G T C | 1e-4 | -9.473e+00 | 4.79% | 1.01% | 201.8bp (377.4bp) | SeqBias: TA-repeat(0.878) More Information \| Similar Motifs Found | motif file (matrix) |
| 33 \* | C G T A A C G T A C T G A G T C A C T G A G T C C G T A A C G T | 1e-4 | -9.473e+00 | 4.79% | 0.92% | 292.4bp (341.6bp) | LEC2/MA0581.1/Jaspar(0.745) More Information \| Similar Motifs Found | motif file (matrix) |
| 34 \* | A C T G C G T A A C T G A C G T C G T A A C G T A C T G A G T C | 1e-4 | -9.376e+00 | 4.19% | 0.76% | 422.3bp (409.5bp) | CG11360(KH)/Drosophila\_melanogaster-RNCMPT00129-PBM/HughesRNA(0.809) More Information \| Similar Motifs Found | motif file (matrix) |
| 35 \* | A T G C C T G A A C G T A C G T C T G A C G T A C T A G A G T C | 1e-3 | -8.664e+00 | 18.56% | 9.36% | 439.8bp (379.1bp) | PHO2(MacIsaac)/Yeast(0.801) More Information \| Similar Motifs Found | motif file (matrix) |
| 36 \* | A T C G A C G T A G T C C G A T A G T C A C T G A C T G A C G T | 1e-3 | -7.329e+00 | 8.38% | 3.16% | 372.4bp (588.2bp) | PB0140.1\_Irf6\_2/Jaspar(0.824) More Information \| Similar Motifs Found | motif file (matrix) |
| 37 \* | A G T C C G T A A C T G C G T A A C T G A C G T | 1e-2 | -5.238e+00 | 38.32% | 28.93% | 356.0bp (460.2bp) | CG11360(KH)/Drosophila\_melanogaster-RNCMPT00129-PBM/HughesRNA(0.769) More Information \| Similar Motifs Found | motif file (matrix) |
