## Supplemental Table 1 for "The Establishment of Cell-Type Specific Gene Regulation in the Sea Urchin Embryo": Hpf10_PMCs_homerResults.html

homer\_motifs\_wbackground/128cell\_PMCs/ - Homer de novo Motif Results


### Homer *de novo* Motif Results (homer\_motifs\_wbackground/128cell\_PMCs/)

Known Motif Enrichment Results  
Gene Ontology Enrichment Results  
If Homer is having trouble matching a motif to a known motif, try copy/pasting the matrix file into
STAMP  
More information on motif finding results: HOMER
| Description of Results
| Tips
  
Total target sequences = 392  
Total background sequences = 277  
\* - possible false positive  

|  |  |  |  |  |  |  |  |  |
| --- | --- | --- | --- | --- | --- | --- | --- | --- |
| Rank | Motif | P-value | log P-pvalue | % of Targets | % of Background | STD(Bg STD) | Best Match/Details | Motif File |
| 1 | A C T G C T A G C A G T C G A T C G T A C T G A C G A T A C G T C G T A C T G A | 1e-41 | -9.618e+01 | 12.24% | 0.94% | 411.3bp (22.9bp) | LMX1A/MA0702.2/Jaspar(0.875) More Information \| Similar Motifs Found | motif file (matrix) |
| 2 | C G T A G T A C G A C T C A T G C T A G G C A T C G A T G T C A A G C T T G C A | 1e-31 | -7.301e+01 | 8.16% | 0.66% | 377.3bp (286.1bp) | grh/dmmpmm(Bigfoot)/fly(0.749) More Information \| Similar Motifs Found | motif file (matrix) |
| 3 | C A T G A C T G G C A T C G T A C G A T C T G A A G T C G A C T A G C T G C T A | 1e-27 | -6.352e+01 | 7.40% | 0.55% | 515.9bp (132.1bp) | PB0198.1\_Zfp128\_2/Jaspar(0.737) More Information \| Similar Motifs Found | motif file (matrix) |
| 4 | C T A G C T A G C T G A A G T C C T A G C A T G C G T A T C G A C T A G A G C T | 1e-27 | -6.352e+01 | 7.40% | 0.65% | 409.6bp (488.1bp) | ETV5/MA0765.2/Jaspar(0.760) More Information \| Similar Motifs Found | motif file (matrix) |
| 5 | C A T G C T A G G T A C G T C A A C T G G A C T C A T G A C G T C A T G G A C T | 1e-27 | -6.243e+01 | 9.18% | 1.07% | 425.7bp (78.2bp) | RBM24(RRM)/Homo\_sapiens-RNCMPT00184-PBM/HughesRNA(0.738) More Information \| Similar Motifs Found | motif file (matrix) |
| 6 | A C T G A G T C G T C A G T A C A C T G A G T C A C T G A T G C | 1e-26 | -6.101e+01 | 15.82% | 2.95% | 347.2bp (508.9bp) | TSAR2/MA1412.1/Jaspar(0.831) More Information \| Similar Motifs Found | motif file (matrix) |
| 7 | A T C G C G A T C G A T T A C G C G T A G T A C G A T C C A G T C G A T A G T C | 1e-24 | -5.737e+01 | 6.89% | 0.38% | 530.1bp (0.0bp) | WRKY28/MA1311.2/Jaspar(0.846) More Information \| Similar Motifs Found | motif file (matrix) |
| 8 | C G A T C T A G T A C G G T C A A T C G A G T C C G A T G C T A T C A G G C A T | 1e-23 | -5.463e+01 | 8.42% | 0.89% | 295.5bp (364.6bp) | br-Z3/dmmpmm(Pollard)/fly(0.649) More Information \| Similar Motifs Found | motif file (matrix) |
| 9 | G C T A C A T G A T C G C A G T A C T G C T G A G T A C C A G T A G C T C T A G | 1e-22 | -5.185e+01 | 11.73% | 2.13% | 387.0bp (572.7bp) | WRKY22(WRKY)/colamp-WRKY22-DAP-Seq(GSE60143)/Homer(0.720) More Information \| Similar Motifs Found | motif file (matrix) |
| 10 | A C T G A C G T C G T A C G T A A G T C C G T A A G T C A G T C A G T C A C G T | 1e-22 | -5.138e+01 | 6.38% | 0.72% | 402.7bp (118.9bp) | AFT2/MA0270.1/Jaspar(0.710) More Information \| Similar Motifs Found | motif file (matrix) |
| 11 | A G C T C T G A T A G C T A C G G A T C T A G C A C G T T C G A A T G C C T A G | 1e-20 | -4.712e+01 | 7.65% | 0.90% | 359.4bp (126.9bp) | EGR2/MA0472.2/Jaspar(0.669) More Information \| Similar Motifs Found | motif file (matrix) |
| 12 | A C T G A C G T A C G T A C T G C T A G G A T C A C G T A T C G T A G C A C G T | 1e-19 | -4.469e+01 | 7.40% | 1.07% | 373.3bp (531.7bp) | AT1G24250(Orphan)/col-AT1G24250-DAP-Seq(GSE60143)/Homer(0.705) More Information \| Similar Motifs Found | motif file (matrix) |
| 13 | C G T A A G T C G T C A A C T G A C T G C G T A C G T A C T G A | 1e-17 | -3.995e+01 | 21.43% | 7.78% | 409.0bp (430.0bp) | EWS:FLI1-fusion(ETS)/SK\_N\_MC-EWS:FLI1-ChIP-Seq(SRA014231)/Homer(0.948) More Information \| Similar Motifs Found | motif file (matrix) |
| 14 | G A C T A G T C C T A G G A T C G C A T C A G T A G T C C A T G G T C A G C T A | 1e-17 | -3.993e+01 | 6.89% | 0.87% | 557.8bp (176.4bp) | YER051W(MacIsaac)/Yeast(0.784) More Information \| Similar Motifs Found | motif file (matrix) |
| 15 | A C T G C T G A C G T A C T A G T A G C C T G A T G A C C T A G C T G A C T A G | 1e-16 | -3.768e+01 | 10.71% | 2.41% | 576.3bp (569.2bp) | POL008.1\_DCE\_S\_I/Jaspar(0.674) More Information \| Similar Motifs Found | motif file (matrix) |
| 16 | A G T C C G T A A C T G A C G T A G T C C G T A A C T G A G C T | 1e-16 | -3.716e+01 | 5.10% | 0.70% | 247.4bp (434.4bp) | ASH1/Literature(Harbison)/Yeast(0.846) More Information \| Similar Motifs Found | motif file (matrix) |
| 17 | A G T C C T A G T C A G G C T A C G A T A G C T T G C A C A T G C G T A C G T A | 1e-15 | -3.623e+01 | 7.65% | 1.30% | 271.5bp (294.6bp) | dve/MA0915.1/Jaspar(0.820) More Information \| Similar Motifs Found | motif file (matrix) |
| 18 | A C T G A G T C A G T C A C G T A C T G A C T G A G T C A G T C | 1e-15 | -3.623e+01 | 7.65% | 1.42% | 432.6bp (143.1bp) | SKN7/SKN7\_H2O2Lo/[](Harbison)/Yeast(0.762) More Information \| Similar Motifs Found | motif file (matrix) |
| 19 | G A T C G C A T G C A T C G T A C A T G G C T A G A T C G C A T C G T A C T A G | 1e-15 | -3.616e+01 | 8.67% | 1.59% | 439.5bp (414.7bp) | GATA1/MA0035.4/Jaspar(0.704) More Information \| Similar Motifs Found | motif file (matrix) |
| 20 | A T G C G C T A C A T G G A T C C G A T C A G T C G T A A G C T G A T C A C T G | 1e-15 | -3.533e+01 | 6.38% | 0.92% | 281.4bp (168.6bp) | DAL80/MA0289.1/Jaspar(0.777) More Information \| Similar Motifs Found | motif file (matrix) |
| 21 | A C T G A C G T C G T A C G T A A C G T G T C A C G T A A C T G | 1e-12 | -2.820e+01 | 8.42% | 2.09% | 351.9bp (318.0bp) | gt/dmmpmm(Bigfoot)/fly(0.819) More Information \| Similar Motifs Found | motif file (matrix) |
| 22 \* | C G A T A G C T G A C T G A T C C T A G C G T A G A T C G A T C C G A T G A T C | 1e-11 | -2.647e+01 | 6.38% | 1.31% | 329.7bp (651.3bp) | ZNF135/MA1587.1/Jaspar(0.715) More Information \| Similar Motifs Found | motif file (matrix) |
| 23 \* | A G T C C G T A A C T G A C T G A C T G G T C A A C T G A C T G | 1e-11 | -2.543e+01 | 7.14% | 1.46% | 428.0bp (504.5bp) | HNRNPH2(RRM)/Homo\_sapiens-RNCMPT00160-PBM/HughesRNA(0.816) More Information \| Similar Motifs Found | motif file (matrix) |
| 24 \* | C G T A A C T G A C T G C G T A A C T G A C T G G T C A A C G T | 1e-10 | -2.377e+01 | 6.89% | 1.74% | 330.2bp (447.6bp) | SF2(RRM)/Drosophila\_melanogaster-RNCMPT00066-PBM/HughesRNA(0.867) More Information \| Similar Motifs Found | motif file (matrix) |
| 25 \* | A T G C C G T A C G T A A G C T A C G T A C G T C G T A A C T G | 1e-9 | -2.259e+01 | 12.24% | 4.67% | 390.5bp (235.8bp) | CG17838(RRM)/Drosophila\_melanogaster-RNCMPT00131-PBM/HughesRNA(0.767) More Information \| Similar Motifs Found | motif file (matrix) |
| 26 \* | C G T A A G T C C G T A C G T A A G T C G C T A A G T C A G T C | 1e-9 | -2.257e+01 | 4.85% | 0.84% | 392.8bp (330.7bp) | UNC-75(RRM)/Caenorhabditis\_elegans-RNCMPT00081-PBM/HughesRNA(0.798) More Information \| Similar Motifs Found | motif file (matrix) |
| 27 \* | C G T A A C T G C G T A A C G T A C G T A C G T A G T C A C T G | 1e-9 | -2.110e+01 | 5.61% | 1.26% | 423.0bp (310.5bp) | PH0037.1\_Hdx/Jaspar(0.715) More Information \| Similar Motifs Found | motif file (matrix) |
| 28 \* | A C G T C G T A A C G T C G T A A C T G A C G T C G T A C G T A | 1e-9 | -2.110e+01 | 5.61% | 1.29% | 576.0bp (228.2bp) | br-Z2/dmmpmm(SeSiMCMC)/fly(0.782) More Information \| Similar Motifs Found | motif file (matrix) |
| 29 \* | A C G T A C T G A C T G A G T C A G T C C G T A A C G T A C G T | 1e-9 | -2.076e+01 | 11.22% | 4.16% | 353.6bp (422.9bp) | pho/dmmpmm(Bergman)/fly(0.822) More Information \| Similar Motifs Found | motif file (matrix) |
| 30 \* | A C G T C G T A T G C A G T A C G A T C A C T G C T G A G C A T T C G A C T G A | 1e-8 | -2.046e+01 | 7.14% | 1.93% | 381.4bp (405.2bp) | CUX2/MA0755.1/Jaspar(0.780) More Information \| Similar Motifs Found | motif file (matrix) |
| 31 \* | A C T G A C T G C G T A C G T A C G T A A G T C A G T C C G T A A G T C A G T C | 1e-8 | -1.901e+01 | 6.12% | 1.63% | 443.7bp (307.9bp) | MF0003.1\_REL\_class/Jaspar(0.732) More Information \| Similar Motifs Found | motif file (matrix) |
| 32 \* | A C T G A C T G A C T G A C G T A T C G A G T C C G T A A C G T | 1e-7 | -1.793e+01 | 7.40% | 2.27% | 336.7bp (525.6bp) | RCS1(MacIsaac)/Yeast(0.896) More Information \| Similar Motifs Found | motif file (matrix) |
| 33 \* | A G T C C G T A A C T G A C G T A G T C A C G T A G T C C G T A | 1e-7 | -1.775e+01 | 5.10% | 1.20% | 468.5bp (414.2bp) | YML081W(MacIsaac)/Yeast(0.788) More Information \| Similar Motifs Found | motif file (matrix) |
| 34 \* | A C G T A C G T A C G T A C T G C G T A A C G T | 1e-2 | -6.536e+00 | 67.35% | 60.09% | 442.0bp (431.7bp) | pan/dmmpmm(Bigfoot)/fly(0.979) More Information \| Similar Motifs Found | motif file (matrix) |
| 35 \* | A C G T A C T G A G T C G T A C C T A G C G T A | 1e-2 | -6.391e+00 | 22.45% | 16.69% | 423.4bp (456.5bp) | NFIA/MA0670.1/Jaspar(0.839) More Information \| Similar Motifs Found | motif file (matrix) |
| 36 \* | A C G T C G T A A C T G A C T G A G T C A G T C | 1e0 | -2.219e+00 | 37.76% | 34.67% | 423.1bp (474.9bp) | ZNF711(Zf)/SHSY5Y-ZNF711-ChIP-Seq(GSE20673)/Homer(0.927) More Information \| Similar Motifs Found | motif file (matrix) |
