## Supplemental Table 1 for "The Establishment of Cell-Type Specific Gene Regulation in the Sea Urchin Embryo": Hpf10_Veg1_homerResults.html

homer\_motifs\_wbackground/128cell\_Veg1/ - Homer de novo Motif Results


### Homer *de novo* Motif Results (homer\_motifs\_wbackground/128cell\_Veg1/)

Known Motif Enrichment Results  
Gene Ontology Enrichment Results  
If Homer is having trouble matching a motif to a known motif, try copy/pasting the matrix file into
STAMP  
More information on motif finding results: HOMER
| Description of Results
| Tips
  
Total target sequences = 38  
Total background sequences = 628  
\* - possible false positive  

|  |  |  |  |  |  |  |  |  |
| --- | --- | --- | --- | --- | --- | --- | --- | --- |
| Rank | Motif | P-value | log P-pvalue | % of Targets | % of Background | STD(Bg STD) | Best Match/Details | Motif File |
| 1 \* | C G A T C G T A C G T A C G T A C G T A A C G T A C T G A C T G C A G T C G T A | 1e-10 | -2.400e+01 | 28.95% | 1.83% | 240.7bp (494.6bp) | ROX8(RRM)/Drosophila\_melanogaster-RNCMPT00148-PBM/HughesRNA(0.836) More Information \| Similar Motifs Found | motif file (matrix) |
| 2 \* | A C G T C T A G A C T G A G T C A C T G A C G T A C T G G T A C | 1e-8 | -1.945e+01 | 28.95% | 2.73% | 297.9bp (309.9bp) | NAC058/MA0938.1/Jaspar(0.824) More Information \| Similar Motifs Found | motif file (matrix) |
| 3 \* | A C G T A C G T A C G T A C T G A C T G C G T A A C G T A C T G A G T C C G T A | 1e-7 | -1.720e+01 | 23.68% | 1.94% | 237.2bp (315.6bp) | AtLEC2(ABI3/VP1)/Arabidopsis thaliana/AthaMap(0.741) More Information \| Similar Motifs Found | motif file (matrix) |
| 4 \* | C G T A C G T A A C T G C G T A C G T A A C T G A C G T C G T A A C T G C T A G | 1e-6 | -1.478e+01 | 23.68% | 2.57% | 183.6bp (393.8bp) | MSI(RRM)/Drosophila\_melanogaster-RNCMPT00100-PBM/HughesRNA(0.755) More Information \| Similar Motifs Found | motif file (matrix) |
| 5 \* | A C G T G T C A A C G T C G T A A G T C A C T G A C T G A G T C C G T A A G T C | 1e-5 | -1.333e+01 | 15.79% | 1.08% | 251.2bp (198.6bp) | STP2/MA0395.1/Jaspar(0.682) More Information \| Similar Motifs Found | motif file (matrix) |
| 6 \* | A C G T A C T G A C G T A G T C A G T C A C T G | 1e-4 | -9.752e+00 | 60.53% | 29.25% | 486.1bp (441.4bp) | FMR1(KH)/Drosophila\_melanogaster-RNCMPT00015-PBM/HughesRNA(0.876) More Information \| Similar Motifs Found | motif file (matrix) |
| 7 \* | C T A G C G T A A C T G A C T G A G T C A C G T A C T G G T A C C T A G A C T G | 1e-3 | -8.287e+00 | 7.89% | 0.45% | 591.7bp (141.6bp) | CRZ1(MacIsaac)/Yeast(0.816) More Information \| Similar Motifs Found | motif file (matrix) |
| 8 \* | A C T G T C A G A C G T C T G A G T C A C T A G A C T G A C T G A G C T C G T A | 1e-3 | -7.273e+00 | 21.05% | 5.30% | 273.8bp (320.5bp) | SFP1/SFP1\_SM/50-RAP1(Harbison)/Yeast(0.668) More Information \| Similar Motifs Found | motif file (matrix) |
| 9 \* | A C T G C G T A A G T C C G T A C G T A A G T C A C G T C G T A A T G C A G T C | 1e-2 | -6.365e+00 | 5.26% | 0.30% | 414.2bp (79.4bp) | MYB30(MYB)/colamp-MYB30-DAP-Seq(GSE60143)/Homer(0.766) More Information \| Similar Motifs Found | motif file (matrix) |
| 10 \* | A C G T C G T A C G T A A C T G A G T C A G T C | 1e-1 | -4.544e+00 | 39.47% | 21.88% | 560.8bp (440.7bp) | bcd/dmmpmm(Bigfoot)/fly(0.854) More Information \| Similar Motifs Found | motif file (matrix) |
| 11 \* | A T C G A C G T A C G T C G T A A C T G A C T G A C T G A C T G | 1e0 | -9.873e-01 | 36.84% | 33.28% | 676.3bp (474.5bp) | RPH1/MA0372.1/Jaspar(0.928) More Information \| Similar Motifs Found | motif file (matrix) |
