## Supplemental Table 1 for "The Establishment of Cell-Type Specific Gene Regulation in the Sea Urchin Embryo": Hpf10_Veg2_homerResults.html

homer\_motifs\_wbackground/128cell\_Veg2/ - Homer de novo Motif Results


### Homer *de novo* Motif Results (homer\_motifs\_wbackground/128cell\_Veg2/)

Known Motif Enrichment Results  
Gene Ontology Enrichment Results  
If Homer is having trouble matching a motif to a known motif, try copy/pasting the matrix file into
STAMP  
More information on motif finding results: HOMER
| Description of Results
| Tips
  
Total target sequences = 308  
Total background sequences = 651  
\* - possible false positive  

|  |  |  |  |  |  |  |  |  |
| --- | --- | --- | --- | --- | --- | --- | --- | --- |
| Rank | Motif | P-value | log P-pvalue | % of Targets | % of Background | STD(Bg STD) | Best Match/Details | Motif File |
| 1 \* | A C T G A C G T C G T A A C T G A C G T A C G T G T C A C G T A A T C G A C G T | 1e-11 | -2.711e+01 | 5.84% | 0.65% | 188.2bp (323.5bp) | HMBOX1/MA0895.1/Jaspar(0.787) More Information \| Similar Motifs Found | motif file (matrix) |
| 2 \* | C T G A T A G C C T A G G T A C C T G A A C G T A C T G C T A G G A C T C G T A | 1e-11 | -2.711e+01 | 5.84% | 0.74% | 307.2bp (154.1bp) | PL0008.1\_hlh-29/Jaspar(0.832) More Information \| Similar Motifs Found | motif file (matrix) |
| 3 \* | G A T C A C T G A C T G C T A G T A C G C G T A G T C A G A T C C T A G A G T C | 1e-11 | -2.711e+01 | 4.55% | 0.38% | 272.8bp (174.6bp) | RELB/MA1117.1/Jaspar(0.760) More Information \| Similar Motifs Found | motif file (matrix) |
| 4 \* | A C T G T A G C A C G T A T G C C T G A C A T G G C A T T G C A A C T G A T G C | 1e-11 | -2.711e+01 | 4.55% | 0.43% | 284.4bp (183.7bp) | OSR1/MA1542.1/Jaspar(0.725) More Information \| Similar Motifs Found | motif file (matrix) |
| 5 \* | C G T A A G C T A C T G G T A C A C G T A G C T C G T A C G T A A G T C G T C A | 1e-10 | -2.480e+01 | 5.52% | 0.75% | 285.1bp (697.7bp) | YAP5(MacIsaac)/Yeast(0.717) More Information \| Similar Motifs Found | motif file (matrix) |
| 6 \* | A C G T T A G C C A T G A C T G A C T G C G T A A G T C A C G T A G T C A G T C | 1e-9 | -2.255e+01 | 5.19% | 0.65% | 526.3bp (542.9bp) | PB0203.1\_Zfp691\_2/Jaspar(0.705) More Information \| Similar Motifs Found | motif file (matrix) |
| 7 \* | A G T C A C T G C T A G A C G T A C T G A C T G A C T G C G A T A T C G C G T A | 1e-8 | -1.913e+01 | 3.57% | 0.45% | 199.5bp (299.5bp) | MYB55/MA1041.1/Jaspar(0.767) More Information \| Similar Motifs Found | motif file (matrix) |
| 8 \* | G C T A C G A T C G A T C A T G G T A C G T A C G A T C C A T G A C T G C G A T | 1e-8 | -1.913e+01 | 3.57% | 0.33% | 211.5bp (130.2bp) | REB1(MacIsaac)/Yeast(0.714) More Information \| Similar Motifs Found | motif file (matrix) |
| 9 \* | C T A G C T A G A T C G C T G A G T A C C G T A A C T G A C G T A G T C A G T C | 1e-8 | -1.913e+01 | 3.57% | 0.45% | 226.6bp (270.0bp) | RELA/MA0107.1/Jaspar(0.742) More Information \| Similar Motifs Found | motif file (matrix) |
| 10 \* | A C T G C G T A A C G T A C T G C A T G C G T A A C G T A C G T A C G T A C T G | 1e-8 | -1.913e+01 | 3.57% | 0.32% | 392.5bp (458.3bp) | Pp\_0237(RRM)/Physcomitrella\_patens-RNCMPT00237-PBM/HughesRNA(0.726) More Information \| Similar Motifs Found | motif file (matrix) |
| 11 \* | A G T C A C G T A C T G A G T C A C G T A G T C C G T A A G T C | 1e-8 | -1.913e+01 | 3.57% | 0.31% | 254.6bp (543.5bp) | MafA(bZIP)/Islet-MafA-ChIP-Seq(GSE30298)/Homer(0.690) More Information \| Similar Motifs Found | motif file (matrix) |
| 12 \* | A G T C G T A C A C G T A G T C G T A C C G A T G A C T A G C T C G A T T A C G | 1e-7 | -1.779e+01 | 6.49% | 1.46% | 387.0bp (483.8bp) | TF3A(C2H2)/col-TF3A-DAP-Seq(GSE60143)/Homer(0.732) More Information \| Similar Motifs Found | motif file (matrix) |
| 13 \* | A C T G A T G C C G T A A C T G A C G T A G T C A C T G A C G T | 1e-7 | -1.709e+01 | 2.60% | 0.28% | 199.2bp (372.2bp) | Adf1/dmmpmm(Pollard)/fly(0.808) More Information \| Similar Motifs Found | motif file (matrix) |
| 14 \* | C G T A A G T C C T G A C A T G A C T G A C G T A C G T A G T C G A T C C G T A | 1e-6 | -1.608e+01 | 6.82% | 1.83% | 326.4bp (510.3bp) | GRHL2/MA1105.2/Jaspar(0.796) More Information \| Similar Motifs Found | motif file (matrix) |
| 15 \* | C G A T A C G T T A C G A G T C C G A T A G C T A G T C A C G T A T G C C A G T | 1e-6 | -1.592e+01 | 11.04% | 4.06% | 395.3bp (443.2bp) | RNP4F(RRM)/Drosophila\_melanogaster-RNCMPT00060-PBM/HughesRNA(0.798) More Information \| Similar Motifs Found | motif file (matrix) |
| 16 \* | A G C T C G T A A T G C G C A T A G T C G C A T A G T C A G T C C G T A A C T G | 1e-6 | -1.554e+01 | 4.55% | 0.90% | 422.8bp (275.7bp) | CG11360(KH)/Drosophila\_melanogaster-RNCMPT00129-PBM/HughesRNA(0.682) More Information \| Similar Motifs Found | motif file (matrix) |
| 17 \* | C A T G A T G C G T C A G C T A C A T G C A T G T A G C A C T G C G A T C G T A | 1e-6 | -1.508e+01 | 3.57% | 0.57% | 274.1bp (292.7bp) | Kr/dmmpmm(Pollard)/fly(0.664) More Information \| Similar Motifs Found | motif file (matrix) |
| 18 \* | G A T C C G T A A G C T T G C A A C T G C G T A C G A T A C T G G T A C G A C T | 1e-5 | -1.167e+01 | 4.55% | 1.23% | 362.8bp (463.5bp) | At3g24120(G2like)/col-At3g24120-DAP-Seq(GSE60143)/Homer(0.655) More Information \| Similar Motifs Found | motif file (matrix) |
| 19 \* | C G T A A C G T T A G C C G T A C A T G C G A T A C T G C G T A G C T A G T C A | 1e-4 | -1.112e+01 | 10.71% | 4.77% | 289.6bp (595.8bp) | At5g04390(C2H2)/col200-At5g04390-DAP-Seq(GSE60143)/Homer(0.743) More Information \| Similar Motifs Found | motif file (matrix) |
| 20 \* | C G A T A T G C G T A C C G T A A G C T G T A C A G T C C T G A A G C T A T G C | 1e-4 | -1.100e+01 | 2.92% | 0.53% | 171.8bp (467.2bp) | HOXA1(Homeobox)/mES-Hoxa1-ChIP-Seq(SRP084292)/Homer(0.806) More Information \| Similar Motifs Found | motif file (matrix) |
| 21 \* | C G T A A C G T A C G T C G T A A C T G A G T C A G T C A G T C | 1e-4 | -1.053e+01 | 3.25% | 0.72% | 186.2bp (459.3bp) | br-Z4/dmmpmm(Pollard)/fly(0.758) More Information \| Similar Motifs Found | motif file (matrix) |
| 22 \* | A G T C A G T C A G T C A G T C A C G T C G T A A C T G C G T A | 1e-4 | -1.019e+01 | 3.90% | 1.07% | 271.2bp (106.6bp) | GIS1/MA0306.1/Jaspar(0.788) More Information \| Similar Motifs Found | motif file (matrix) |
| 23 \* | C G T A A C G T A G T C A C T G A G T C A C T G A C T G A G T C | 1e-3 | -9.116e+00 | 2.60% | 0.47% | 167.5bp (185.6bp) | YY2/MA0748.2/Jaspar(0.776) More Information \| Similar Motifs Found | motif file (matrix) |
| 24 \* | C A T G A C G T C T G A G T C A A C T G G T A C A C T G A G T C | 1e-3 | -9.006e+00 | 6.17% | 2.43% | 409.7bp (472.1bp) | STP3/MA0396.1/Jaspar(0.694) More Information \| Similar Motifs Found | motif file (matrix) |
| 25 \* | C G T A A C T G A C G T A C G T A G C T A C T G A G C T A G T C G T C A A C G T | 1e-3 | -8.988e+00 | 4.55% | 1.45% | 284.2bp (429.9bp) | FXR1(KH)/Homo\_sapiens-RNCMPT00161-PBM/HughesRNA(0.793) More Information \| Similar Motifs Found | motif file (matrix) |
| 26 \* | C G T A A G T C A C T G C A T G C T A G C G A T A C G T A T C G A C G T A C G T | 1e-3 | -8.948e+00 | 1.62% | 0.10% | 302.6bp (0.0bp) | YBX2(CSD)/Homo\_sapiens-RNCMPT00084-PBM/HughesRNA(0.761) More Information \| Similar Motifs Found | motif file (matrix) |
| 27 \* | G C T A C T A G A G T C C T G A T A C G T A G C G C T A A C T G A G T C C G T A | 1e-3 | -8.822e+00 | 2.92% | 0.63% | 279.5bp (240.9bp) | SNAI3/MA1559.1/Jaspar(0.749) More Information \| Similar Motifs Found | motif file (matrix) |
| 28 \* | A C G T C G T A C G T A A G T C A G T C A G T C C G T A A C G T | 1e-3 | -8.701e+00 | 3.57% | 0.93% | 413.8bp (470.9bp) | NCU02182/MA1434.1/Jaspar(0.821) More Information \| Similar Motifs Found | motif file (matrix) |
| 29 \* | C T G A A G T C C G A T G T C A A C T G G A C T A C G T G A C T | 1e-3 | -7.338e+00 | 23.70% | 16.57% | 294.5bp (459.7bp) | br-Z3/dmmpmm(Pollard)/fly(0.869) More Information \| Similar Motifs Found | motif file (matrix) |
| 30 \* | A C G T A C T G A G C T A G T C A C G T A C T G C G T A G A C T A C G T T C A G | 1e-3 | -7.248e+00 | 7.47% | 3.67% | 239.0bp (499.5bp) | Dux/MA0611.1/Jaspar(0.754) More Information \| Similar Motifs Found | motif file (matrix) |
| 31 \* | A G T C A T C G G A C T G T A C A G T C A C T G G A C T T A G C A G T C A T C G | 1e-3 | -7.218e+00 | 2.60% | 0.64% | 175.5bp (257.8bp) | YLL054C/MA0429.1/Jaspar(0.707) More Information \| Similar Motifs Found | motif file (matrix) |
| 32 \* | A C T G C G T A A C G T C G T A C G T A A G T C A C T G G T A C | 1e-2 | -6.710e+00 | 5.19% | 2.25% | 200.1bp (534.4bp) | GATA5/MA0766.2/Jaspar(0.832) More Information \| Similar Motifs Found | motif file (matrix) |
| 33 \* | C G T A A C G T A G T C A C G T A C T G A G T C A T C G A C T G | 1e-2 | -5.706e+00 | 1.95% | 0.48% | 190.4bp (550.1bp) | nit-4/MA1435.1/Jaspar(0.761) More Information \| Similar Motifs Found | motif file (matrix) |
| 34 \* | T G A C C T A G G T A C C T A G T G A C C T G A G T A C C T G A G T A C T C G A | 1e-1 | -4.461e+00 | 3.25% | 1.48% | 162.7bp (478.9bp) | SeqBias: CA-repeat(0.866) More Information \| Similar Motifs Found | motif file (matrix) |
| 35 \* | A C G T C G T A A C G T C G T A A C G T A C T G A C T G A G T C | 1e-1 | -3.257e+00 | 3.25% | 1.74% | 373.3bp (767.3bp) | MOT2/MA0379.1/Jaspar(0.790) More Information \| Similar Motifs Found | motif file (matrix) |
