## Supplemental Table 1 for "The Establishment of Cell-Type Specific Gene Regulation in the Sea Urchin Embryo": Hpf12_EctoApical_homerResults.html

homer\_motifs\_wbackground/512cell\_EctoApical/ - Homer de novo Motif Results


### Homer *de novo* Motif Results (homer\_motifs\_wbackground/512cell\_EctoApical/)

Known Motif Enrichment Results  
Gene Ontology Enrichment Results  
If Homer is having trouble matching a motif to a known motif, try copy/pasting the matrix file into
STAMP  
More information on motif finding results: HOMER
| Description of Results
| Tips
  
Total target sequences = 521  
Total background sequences = 3051  
\* - possible false positive  

|  |  |  |  |  |  |  |  |  |
| --- | --- | --- | --- | --- | --- | --- | --- | --- |
| Rank | Motif | P-value | log P-pvalue | % of Targets | % of Background | STD(Bg STD) | Best Match/Details | Motif File |
| 1 | T A G C T G A C G C A T C T A G A G C T A T G C G T C A T G C A A C G T G T A C | 1e-21 | -5.046e+01 | 24.57% | 9.78% | 392.9bp (485.9bp) | Pbx3(Homeobox)/GM12878-PBX3-ChIP-Seq(GSE32465)/Homer(0.916) More Information \| Similar Motifs Found | motif file (matrix) |
| 2 | T C G A G C A T G T A C C T A G C G T A C A G T T C A G T C A G C G T A A T C G | 1e-20 | -4.743e+01 | 33.59% | 16.60% | 422.6bp (500.7bp) | CUX1(Homeobox)/K562-CUX1-ChIP-Seq(GSE92882)/Homer(0.884) More Information \| Similar Motifs Found | motif file (matrix) |
| 3 | A G C T A C G T A G T C C G T A G A T C A C G T C G T A G T A C A C T G C G T A | 1e-19 | -4.386e+01 | 6.91% | 0.96% | 374.9bp (403.0bp) | PB0106.1\_Arid5a\_2/Jaspar(0.714) More Information \| Similar Motifs Found | motif file (matrix) |
| 4 | A T C G T C A G A G T C A T G C A G C T C G T A A C G T A C G T C T A G A C G T | 1e-17 | -4.076e+01 | 50.10% | 31.69% | 341.4bp (462.2bp) | SOX15/MA1152.1/Jaspar(0.908) More Information \| Similar Motifs Found | motif file (matrix) |
| 5 | A C T G A T G C T C G A C A G T A C G T G A C T G T C A C A G T A C G T A T G C | 1e-13 | -3.168e+01 | 10.75% | 3.29% | 367.7bp (366.5bp) | EDT1/MA0990.1/Jaspar(0.781) More Information \| Similar Motifs Found | motif file (matrix) |
| 6 | G T A C A G T C C G A T T C A G A C T G C G T A A T G C T G A C C G T A A C G T | 1e-13 | -3.091e+01 | 5.18% | 0.80% | 292.6bp (436.6bp) | NRG1/MA0347.1/Jaspar(0.664) More Information \| Similar Motifs Found | motif file (matrix) |
| 7 | A C T G T A C G G A T C A C G T T G A C C G T A A C G T A G C T | 1e-12 | -2.895e+01 | 47.79% | 32.46% | 426.2bp (479.4bp) | POU6F1(var.2)/MA1549.1/Jaspar(0.799) More Information \| Similar Motifs Found | motif file (matrix) |
| 8 \* | C G A T T G A C C G T A T C G A G C A T G T A C G T C A C G A T T C A G G T C A | 1e-11 | -2.589e+01 | 10.36% | 3.59% | 377.9bp (472.3bp) | PB0176.1\_Sox5\_2/Jaspar(0.641) More Information \| Similar Motifs Found | motif file (matrix) |
| 9 \* | C G T A G T C A A C T G C G T A A C T G A C G T A C G T C G A T A C T G A C G T | 1e-10 | -2.305e+01 | 2.88% | 0.30% | 467.4bp (247.4bp) | MYB.PH3(2)(MYB)/Petunia hybrida/AthaMap(0.736) More Information \| Similar Motifs Found | motif file (matrix) |
| 10 \* | C T A G A C G T C G A T G T A C G A C T A C G T A C G T C A G T A C T G C T A G | 1e-9 | -2.270e+01 | 6.14% | 1.58% | 285.6bp (336.4bp) | Sf3b4(RRM)/Danio\_rerio-RNCMPT00224-PBM/HughesRNA(0.765) More Information \| Similar Motifs Found | motif file (matrix) |
| 11 \* | A T G C A C G T C G A T G C A T T C A G T C G A T C G A C G T A C T A G A C T G | 1e-9 | -2.166e+01 | 26.68% | 16.01% | 362.9bp (442.5bp) | TATA-Box(TBP)/Promoter/Homer(0.768) More Information \| Similar Motifs Found | motif file (matrix) |
| 12 \* | A C T G A G T C G T A C A G C T C T A G G A T C C T A G A G T C A C G T A G T C | 1e-8 | -2.066e+01 | 4.03% | 0.76% | 541.1bp (329.0bp) | NRF1/MA0506.1/Jaspar(0.743) More Information \| Similar Motifs Found | motif file (matrix) |
| 13 \* | A C G T A C G T T A G C A G T C C G T A A C T G G A T C C T A G A C T G A C G T | 1e-8 | -1.946e+01 | 3.65% | 0.69% | 361.9bp (254.0bp) | ZBTB12/MA1649.1/Jaspar(0.682) More Information \| Similar Motifs Found | motif file (matrix) |
| 14 \* | G T C A A C G T A G C T C G T A C G T A C G T A A C T G A G T C C G T A A G T C | 1e-8 | -1.920e+01 | 2.88% | 0.42% | 395.0bp (228.8bp) | DOF2(C2C2(Zn) Dof)/Zea mays/AthaMap(0.752) More Information \| Similar Motifs Found | motif file (matrix) |
| 15 \* | C G T A A G C T A C T G A C T G C T G A C G T A C T A G A C T G | 1e-8 | -1.905e+01 | 20.35% | 11.56% | 471.0bp (487.0bp) | HSF1/MA0319.1/Jaspar(0.803) More Information \| Similar Motifs Found | motif file (matrix) |
| 16 \* | T C G A A C T G C G T A T A G C A G T C A G T C G T A C A G C T A G C T A G T C | 1e-8 | -1.900e+01 | 5.57% | 1.56% | 377.6bp (639.4bp) | MSN4/Literature(Harbison)/Yeast(0.737) More Information \| Similar Motifs Found | motif file (matrix) |
| 17 \* | G T C A C A T G C A T G C T A G G T C A C G T A A C T G G C T A A G T C A G T C | 1e-7 | -1.696e+01 | 4.41% | 1.13% | 279.0bp (525.7bp) | GCR1(MacIsaac)/Yeast(0.764) More Information \| Similar Motifs Found | motif file (matrix) |
| 18 \* | A G T C C G A T A G T C T A C G A G T C C T A G A G C T G T C A A T G C A C T G | 1e-6 | -1.533e+01 | 2.69% | 0.47% | 293.0bp (390.9bp) | SPL7/MA1060.1/Jaspar(0.777) More Information \| Similar Motifs Found | motif file (matrix) |
| 19 \* | A C G T A C G T A C G T A C T G A C G T A G T C A C G T C G T A A C G T A G T C | 1e-6 | -1.505e+01 | 1.73% | 0.18% | 266.1bp (449.9bp) | br/MA0010.1/Jaspar(0.748) More Information \| Similar Motifs Found | motif file (matrix) |
| 20 \* | A G T C A G T C A G T C A G T C A C G T A G T C A C T G A C G T | 1e-5 | -1.241e+01 | 3.45% | 0.96% | 392.4bp (513.6bp) | MSN4/MA0342.1/Jaspar(0.785) More Information \| Similar Motifs Found | motif file (matrix) |
| 21 \* | C G T A C G T A A C G T A C T G A G T C A C G T A G T C A C G T | 1e-4 | -1.039e+01 | 6.72% | 3.15% | 506.8bp (519.5bp) | CST6(MacIsaac)/Yeast(0.688) More Information \| Similar Motifs Found | motif file (matrix) |
| 22 \* | A C T G A C G T A C G T C G T A A G T C C G T A A C G T C G T A A G T C C G T A | 1e-4 | -1.027e+01 | 1.54% | 0.23% | 204.1bp (326.9bp) | YAP6(MacIsaac)/Yeast(0.768) More Information \| Similar Motifs Found | motif file (matrix) |
