## Supplemental Table 1 for "The Establishment of Cell-Type Specific Gene Regulation in the Sea Urchin Embryo": Hpf12_EctoArboral_homerResults.html

homer\_motifs\_wbackground/512cell\_EctoArboral/ - Homer de novo Motif Results


### Homer *de novo* Motif Results (homer\_motifs\_wbackground/512cell\_EctoArboral/)

Known Motif Enrichment Results  
Gene Ontology Enrichment Results  
If Homer is having trouble matching a motif to a known motif, try copy/pasting the matrix file into
STAMP  
More information on motif finding results: HOMER
| Description of Results
| Tips
  
Total target sequences = 595  
Total background sequences = 2947  
\* - possible false positive  

|  |  |  |  |  |  |  |  |  |
| --- | --- | --- | --- | --- | --- | --- | --- | --- |
| Rank | Motif | P-value | log P-pvalue | % of Targets | % of Background | STD(Bg STD) | Best Match/Details | Motif File |
| 1 | A T G C A G T C A C G T C G T A A C G T A C G T A T C G C A G T A T C G C G T A | 1e-37 | -8.742e+01 | 22.18% | 6.05% | 493.7bp (438.6bp) | SOX15/MA1152.1/Jaspar(0.764) More Information \| Similar Motifs Found | motif file (matrix) |
| 2 | C T A G C T G A G C T A T C A G A T G C G T A C G T A C C G A T | 1e-29 | -6.904e+01 | 62.86% | 39.48% | 462.9bp (432.8bp) | GCR1(MacIsaac)/Yeast(0.832) More Information \| Similar Motifs Found | motif file (matrix) |
| 3 | C G A T T C A G C G T A G C T A G C A T C T G A T C A G T C A G C T A G T C G A | 1e-28 | -6.615e+01 | 44.71% | 23.65% | 424.8bp (404.6bp) | HNRNPA2B1(RRM)/Homo\_sapiens-RNCMPT00024-PBM/HughesRNA(0.741) More Information \| Similar Motifs Found | motif file (matrix) |
| 4 | A C T G C G T A C G T A C G T A A C T G A C G T | 1e-28 | -6.544e+01 | 65.21% | 42.44% | 471.0bp (460.7bp) | PTBP1(RRM)/Homo\_sapiens-RNCMPT00269-PBM/HughesRNA(0.881) More Information \| Similar Motifs Found | motif file (matrix) |
| 5 | A C G T A C T G A C T G C T G A C G T A C G T A A T G C A G T C | 1e-21 | -4.852e+01 | 41.85% | 24.05% | 487.0bp (468.5bp) | NFATC4/MA1525.1/Jaspar(0.836) More Information \| Similar Motifs Found | motif file (matrix) |
| 6 | G T A C G A C T G T A C C T A G A G C T A G T C G T A C A G T C C G T A A G C T | 1e-19 | -4.406e+01 | 8.07% | 1.59% | 461.1bp (460.8bp) | PB0098.1\_Zfp410\_1/Jaspar(0.750) More Information \| Similar Motifs Found | motif file (matrix) |
| 7 | A C G T A G T C A G T C C G T A A C T G A G C T C T A G C G T A C G A T A G T C | 1e-19 | -4.376e+01 | 6.89% | 1.13% | 523.5bp (468.8bp) | PB0195.1\_Zbtb3\_2/Jaspar(0.677) More Information \| Similar Motifs Found | motif file (matrix) |
| 8 | C T G A A C G T A G T C A G C T A T C G C G T A A C T G A C T G A C T G A T G C | 1e-18 | -4.156e+01 | 5.04% | 0.58% | 546.0bp (356.8bp) | Tal1(0.652) More Information \| Similar Motifs Found | motif file (matrix) |
| 9 | C G T A C G A T A C G T A G T C C G T A C G T A A C T G C G T A C G T A A T G C | 1e-17 | -3.914e+01 | 4.37% | 0.46% | 443.5bp (382.0bp) | Hsf/MA1458.1/Jaspar(0.765) More Information \| Similar Motifs Found | motif file (matrix) |
| 10 | G T A C A T C G C G T A A C T G G T C A C G T A C A T G C G T A | 1e-15 | -3.682e+01 | 57.48% | 40.66% | 470.2bp (427.6bp) | PABPN1(RRM)/Homo\_sapiens-RNCMPT00157-PBM/HughesRNA(0.834) More Information \| Similar Motifs Found | motif file (matrix) |
| 11 | A C T G G T A C G A C T A C T G A G T C A G C T A C G T A G T C A C T G C G T A | 1e-14 | -3.430e+01 | 4.37% | 0.55% | 355.5bp (415.1bp) | XBP1/Literature(Harbison)/Yeast(0.744) More Information \| Similar Motifs Found | motif file (matrix) |
| 12 | A T C G C G T A A C T G A T G C C G T A A G T C G T A C C T G A | 1e-14 | -3.420e+01 | 27.39% | 14.76% | 494.7bp (464.4bp) | PB0196.1\_Zbtb7b\_2/Jaspar(0.747) More Information \| Similar Motifs Found | motif file (matrix) |
| 13 | C G T A G A T C G T C A A T G C A G T C C G A T C G T A G T C A C G T A T A C G | 1e-14 | -3.232e+01 | 57.31% | 41.61% | 469.8bp (453.5bp) | NAC055/MA0937.1/Jaspar(0.707) More Information \| Similar Motifs Found | motif file (matrix) |
| 14 | A C G T C T A G A C T G C G T A A G T C A G T C G T C A C G T A | 1e-13 | -3.194e+01 | 18.49% | 8.54% | 540.9bp (443.9bp) | HAP2/MA0313.1/Jaspar(0.745) More Information \| Similar Motifs Found | motif file (matrix) |
| 15 | C A T G A C T G T G C A C G A T A C T G A C T G G T C A A C G T A G C T C G T A | 1e-13 | -3.086e+01 | 42.35% | 27.97% | 495.8bp (437.8bp) | dve/MA0915.1/Jaspar(0.795) More Information \| Similar Motifs Found | motif file (matrix) |
| 16 | A C T G A G T C A C G T A C G T A C G T A C G T | 1e-13 | -3.064e+01 | 58.49% | 43.22% | 478.3bp (463.5bp) | Dof2/MA0020.1/Jaspar(0.893) More Information \| Similar Motifs Found | motif file (matrix) |
| 17 | A C G T A T C G A C G T C T G A A G C T C G T A C T A G A G C T | 1e-13 | -3.060e+01 | 49.92% | 34.97% | 467.5bp (420.7bp) | PUM(PUF)/Drosophila\_melanogaster-RNCMPT00102-PBM/HughesRNA(0.759) More Information \| Similar Motifs Found | motif file (matrix) |
| 18 | C G T A C T A G G C A T T C A G A T G C A C T G T C A G C A G T C G A T A G T C | 1e-13 | -3.033e+01 | 12.61% | 4.84% | 388.2bp (391.2bp) | RUNX2/MA0511.2/Jaspar(0.738) More Information \| Similar Motifs Found | motif file (matrix) |
| 19 | A C G T A C G T A G C T C A G T C G T A A C G T A G T C A C T G A G T C A C G T | 1e-12 | -2.925e+01 | 4.37% | 0.68% | 371.3bp (288.5bp) | PH0064.1\_Hoxb9/Jaspar(0.798) More Information \| Similar Motifs Found | motif file (matrix) |
| 20 | A C G T A C G T A C G T C G T A A C T G C G T A C G T A C G T A | 1e-12 | -2.847e+01 | 87.73% | 75.99% | 473.2bp (467.9bp) | ARR1/Literature(Harbison)/Yeast(0.757) More Information \| Similar Motifs Found | motif file (matrix) |
| 21 \* | A G T C C G T A G T A C G T C A A G T C A C G T A T C G A C T G A T C G C G T A | 1e-11 | -2.718e+01 | 33.45% | 21.06% | 495.1bp (451.8bp) | Rbm24(RRM)/Tetraodon\_nigroviridis-RNCMPT00285-PBM/HughesRNA(0.698) More Information \| Similar Motifs Found | motif file (matrix) |
| 22 \* | T A C G A T C G A G T C A C T G A T C G G T C A T A G C A G T C A G T C A T G C | 1e-11 | -2.666e+01 | 2.02% | 0.11% | 452.9bp (210.3bp) | HINFP(Zf)/K562-HINFP.eGFP-ChIP-Seq(Encode)/Homer(0.799) More Information \| Similar Motifs Found | motif file (matrix) |
| 23 \* | A C T G A C G T A C T G A C G T C G T A A G T C C G T A C G T A A C T G C G T A | 1e-11 | -2.619e+01 | 2.18% | 0.14% | 611.1bp (113.5bp) | PUM(PUF)/Drosophila\_melanogaster-RNCMPT00101-PBM/HughesRNA(0.715) More Information \| Similar Motifs Found | motif file (matrix) |
| 24 \* | C G T A A C T G A C T G C G T A A C G T C G A T C G T A C G T A A C G T A G T C | 1e-11 | -2.559e+01 | 70.42% | 56.92% | 455.2bp (458.7bp) | PH0138.1\_Pitx2/Jaspar(0.829) More Information \| Similar Motifs Found | motif file (matrix) |
| 25 \* | C G T A A T C G C G T A A G C T A C G T A C T G A C T G A C T G | 1e-11 | -2.554e+01 | 58.82% | 44.97% | 488.5bp (452.3bp) | HAP2/Literature(Harbison)/Yeast(0.798) More Information \| Similar Motifs Found | motif file (matrix) |
| 26 \* | A C G T C G T A A G T C A C G T C T A G A C T G C T A G A G T C C G T A A G T C | 1e-10 | -2.376e+01 | 2.35% | 0.21% | 392.2bp (195.9bp) | ZNF416(Zf)/HEK293-ZNF416.GFP-ChIP-Seq(GSE58341)/Homer(0.766) More Information \| Similar Motifs Found | motif file (matrix) |
| 27 \* | A G T C C G T A A C G T A G T C A G T C A C T G A C G T C G T A | 1e-10 | -2.354e+01 | 4.37% | 0.90% | 355.6bp (464.0bp) | SPL3/MA0577.2/Jaspar(0.778) More Information \| Similar Motifs Found | motif file (matrix) |
| 28 \* | A G T C A G T C A C G T C T G A A C T G C G T A C G T A A G T C C G T A A G T C | 1e-8 | -1.993e+01 | 1.18% | 0.05% | 401.1bp (331.3bp) | AR-halfsite(NR)/LNCaP-AR-ChIP-Seq(GSE27824)/Homer(0.781) More Information \| Similar Motifs Found | motif file (matrix) |
