## Supplemental Table 1 for "The Establishment of Cell-Type Specific Gene Regulation in the Sea Urchin Embryo": Hpf12_EctoNearApical_homerResults.html

homer\_motifs\_wbackground/512cell\_EctoNearApical/ - Homer de novo Motif Results


### Homer *de novo* Motif Results (homer\_motifs\_wbackground/512cell\_EctoNearApical/)

Known Motif Enrichment Results  
Gene Ontology Enrichment Results  
If Homer is having trouble matching a motif to a known motif, try copy/pasting the matrix file into
STAMP  
More information on motif finding results: HOMER
| Description of Results
| Tips
  
Total target sequences = 268  
Total background sequences = 3342  
\* - possible false positive  

|  |  |  |  |  |  |  |  |  |
| --- | --- | --- | --- | --- | --- | --- | --- | --- |
| Rank | Motif | P-value | log P-pvalue | % of Targets | % of Background | STD(Bg STD) | Best Match/Details | Motif File |
| 1 \* | A C T G C G T A A G T C C A T G A T G C A G C T A G T C A G T C A C G T T A C G | 1e-9 | -2.161e+01 | 4.48% | 0.38% | 276.9bp (488.4bp) | MYB88(MYB)/col-MYB88-DAP-Seq(GSE60143)/Homer(0.822) More Information \| Similar Motifs Found | motif file (matrix) |
| 2 \* | A C G T C G T A C G T A A T G C A G T C A C G T A C T G A C G T A C G T A C G T | 1e-7 | -1.841e+01 | 2.99% | 0.16% | 274.2bp (250.0bp) | GRHL2/MA1105.2/Jaspar(0.882) More Information \| Similar Motifs Found | motif file (matrix) |
| 3 \* | A T C G C G T A A T C G A C G T A C G T C G T A A C G T A C G T A C T G A G T C | 1e-7 | -1.679e+01 | 4.10% | 0.47% | 274.0bp (332.0bp) | Tb\_0218(RRM)/Trypanosoma\_brucei-RNCMPT00218-PBM/HughesRNA(0.806) More Information \| Similar Motifs Found | motif file (matrix) |
| 4 \* | C G T A A G C T A G C T C G T A C G T A A C G T G T A C G T C A A C G T C A T G | 1e-7 | -1.669e+01 | 8.21% | 2.07% | 311.1bp (407.6bp) | LIN-39(Homeobox)/cElegans.L3-LIN39-ChIP-Seq(modEncode)/Homer(0.852) More Information \| Similar Motifs Found | motif file (matrix) |
| 5 \* | C G T A C G T A C T G A A C G T A C G T A G T C A C G T C G T A C G T A A G T C | 1e-6 | -1.582e+01 | 3.73% | 0.41% | 154.7bp (334.6bp) | TCX3/MA1682.1/Jaspar(0.687) More Information \| Similar Motifs Found | motif file (matrix) |
| 6 \* | C G T A A G T C C G A T A G T C A C G T C G T A C T A G A C T G A G T C A C G T | 1e-6 | -1.515e+01 | 3.73% | 0.43% | 445.5bp (330.0bp) | WIP5(C2H2)/colamp-WIP5-DAP-Seq(GSE60143)/Homer(0.664) More Information \| Similar Motifs Found | motif file (matrix) |
| 7 \* | C A T G A C T G C T A G C G T A A C T G A G T C T C A G T C G A A C G T A G T C | 1e-6 | -1.461e+01 | 10.07% | 3.36% | 267.2bp (480.5bp) | shn-ZFP1/dmmpmm(Bergman)/fly(0.710) More Information \| Similar Motifs Found | motif file (matrix) |
| 8 \* | A C G T A C T G A G T C C G T A C G T A A G T C C G T A A C G T C G T A A G T C | 1e-5 | -1.376e+01 | 2.24% | 0.13% | 299.7bp (391.5bp) | CEBP:AP1(bZIP)/ThioMac-CEBPb-ChIP-Seq(GSE21512)/Homer(0.771) More Information \| Similar Motifs Found | motif file (matrix) |
| 9 \* | A T G C A G C T T G A C A G T C T G C A C G A T C G T A C G T A T G C A C T A G | 1e-5 | -1.338e+01 | 1.49% | 0.05% | 125.7bp (720.9bp) | cad/dmmpmm(Papatsenko)/fly(0.785) More Information \| Similar Motifs Found | motif file (matrix) |
| 10 \* | A T C G C A G T C T G A A G T C T A G C C G T A C G A T A G C T G A C T T G C A | 1e-5 | -1.295e+01 | 16.79% | 8.07% | 376.8bp (426.3bp) | BARHL2/MA0635.1/Jaspar(0.746) More Information \| Similar Motifs Found | motif file (matrix) |
| 11 \* | A C G T A C G T A C G T C G A T A C G T G A C T G A C T A C T G A G T C C T A G | 1e-5 | -1.277e+01 | 6.34% | 1.66% | 274.7bp (425.3bp) | hb/dmmpmm(Papatsenko)/fly(0.860) More Information \| Similar Motifs Found | motif file (matrix) |
| 12 \* | C A G T A T C G G A C T A C G T C T G A A G T C A C T G T G C A T A C G A C G T | 1e-5 | -1.257e+01 | 5.60% | 1.32% | 236.5bp (422.9bp) | HOXC13/MA0907.1/Jaspar(0.757) More Information \| Similar Motifs Found | motif file (matrix) |
| 13 \* | A G T C A G T C C G T A A C G T A G T C C G T A C G T A A C G T A C T G A G T C | 1e-5 | -1.217e+01 | 1.87% | 0.09% | 321.4bp (298.4bp) | PH0134.1\_Pbx1/Jaspar(0.723) More Information \| Similar Motifs Found | motif file (matrix) |
| 14 \* | A C G T T C A G A T G C T G A C A G T C A G C T A G T C A C G T A T G C A G C T | 1e-4 | -1.129e+01 | 7.46% | 2.46% | 394.1bp (515.6bp) | Trl(Zf)/S2-GAGAfactor-ChIP-Seq(GSE40646)/Homer(0.766) More Information \| Similar Motifs Found | motif file (matrix) |
| 15 \* | A C T G C G A T A C G T A C T G A C G T C G T A C G T A A C T G | 1e-4 | -1.107e+01 | 10.82% | 4.55% | 288.5bp (434.0bp) | PUM(PUF)/Drosophila\_melanogaster-RNCMPT00103-PBM/HughesRNA(0.728) More Information \| Similar Motifs Found | motif file (matrix) |
| 16 \* | T G A C T A G C T A G C A C G T C T G A C T G A T C G A A T G C T C G A A T G C | 1e-4 | -1.067e+01 | 1.49% | 0.08% | 114.3bp (135.5bp) | AT4G12670(MYBrelated)/col-AT4G12670-DAP-Seq(GSE60143)/Homer(0.754) More Information \| Similar Motifs Found | motif file (matrix) |
| 17 \* | G C A T C T G A G C T A A T C G T A C G A C T G G T A C G T A C G T A C C A G T | 1e-3 | -8.502e+00 | 22.39% | 14.21% | 396.5bp (462.9bp) | NRG1/NRG1\_H2O2Hi/[](Harbison)/Yeast(0.730) More Information \| Similar Motifs Found | motif file (matrix) |
| 18 \* | A G T C G T A C A C G T G T A C C G T A C G T A C G T A A G T C A G T C A G T C | 1e-3 | -7.737e+00 | 4.10% | 1.22% | 353.5bp (435.5bp) | RUNX3/MA0684.2/Jaspar(0.649) More Information \| Similar Motifs Found | motif file (matrix) |
| 19 \* | A C G T A C T G A C T G A C T G A C G T A G T C A G T C A G T C C T G A A C G T | 1e-3 | -7.695e+00 | 2.24% | 0.38% | 202.9bp (454.6bp) | At2g45680(TCP)/colamp-At2g45680-DAP-Seq(GSE60143)/Homer(0.816) More Information \| Similar Motifs Found | motif file (matrix) |
| 20 \* | A G C T G T C A C T A G A G T C C T G A C T G A T A C G A T C G A T C G A C T G | 1e-2 | -6.270e+00 | 1.12% | 0.10% | 134.2bp (117.2bp) | MSN4(MacIsaac)/Yeast(0.746) More Information \| Similar Motifs Found | motif file (matrix) |
| 21 \* | C G T A A C T G A C T G C G T A A G T C C G T A A C T G C G T A | 1e-2 | -5.045e+00 | 11.94% | 7.52% | 360.1bp (467.3bp) | SRSF1(RRM)/Homo\_sapiens-RNCMPT00110-PBM/HughesRNA(0.836) More Information \| Similar Motifs Found | motif file (matrix) |
| 22 \* | C G T A A G T C C G T A C G T A A G T C A C G T A G T C A G T C C G T A A T G C | 1e-2 | -4.854e+00 | 1.12% | 0.16% | 37.5bp (314.6bp) | Pp\_0237(RRM)/Physcomitrella\_patens-RNCMPT00237-PBM/HughesRNA(0.701) More Information \| Similar Motifs Found | motif file (matrix) |
| 23 \* | T C G A A G T C G T A C G T C A A G T C A T G C C T G A A G T C A T G C T C G A | 1e-2 | -4.800e+00 | 5.22% | 2.50% | 206.4bp (431.5bp) | ZBTB7C/MA0695.1/Jaspar(0.767) More Information \| Similar Motifs Found | motif file (matrix) |
| 24 \* | A C G T A G T C A G T C G T C A A G T C C G T A A G T C C G T A | 1e-1 | -4.279e+00 | 12.31% | 8.26% | 291.9bp (416.0bp) | BRUNOL6(RRM)/Homo\_sapiens-RNCMPT00187-PBM/HughesRNA(0.841) More Information \| Similar Motifs Found | motif file (matrix) |
| 25 \* | T G A C A T C G A G T C C T G A G T A C A G C T A G T C C G T A A G T C A G T C | 1e-1 | -4.112e+00 | 24.25% | 18.85% | 331.2bp (458.1bp) | MYB51(MYB)/col-MYB51-DAP-Seq(GSE60143)/Homer(0.758) More Information \| Similar Motifs Found | motif file (matrix) |
| 26 \* | A C G T A T C G C G T A A C T G A G T C A C G T A C G T A G T C A G T C A C T G | 1e-1 | -3.714e+00 | 0.75% | 0.10% | 246.1bp (329.0bp) | GCR2(MacIsaac)/Yeast(0.737) More Information \| Similar Motifs Found | motif file (matrix) |
