## Supplemental Table 1 for "The Establishment of Cell-Type Specific Gene Regulation in the Sea Urchin Embryo": Hpf12_EctoOral_homerResults.html

homer\_motifs\_wbackground/512cell\_EctoOral/ - Homer de novo Motif Results


### Homer *de novo* Motif Results (homer\_motifs\_wbackground/512cell\_EctoOral/)

Known Motif Enrichment Results  
Gene Ontology Enrichment Results  
If Homer is having trouble matching a motif to a known motif, try copy/pasting the matrix file into
STAMP  
More information on motif finding results: HOMER
| Description of Results
| Tips
  
Total target sequences = 476  
Total background sequences = 3115  
\* - possible false positive  

|  |  |  |  |  |  |  |  |  |
| --- | --- | --- | --- | --- | --- | --- | --- | --- |
| Rank | Motif | P-value | log P-pvalue | % of Targets | % of Background | STD(Bg STD) | Best Match/Details | Motif File |
| 1 \* | C G T A A C T G A C T G C G T A C G T A A C G T A C T G C G T A A C G T A T C G | 1e-9 | -2.188e+01 | 1.47% | 0.05% | 520.1bp (66.7bp) | TEC1/TEC1\_YPD/[](Harbison)/Yeast(0.787) More Information \| Similar Motifs Found | motif file (matrix) |
| 2 \* | A C T G A C G T A G T C A C G T A C T G A C G T A G C T A G T C C G T A A C T G | 1e-8 | -1.868e+01 | 4.41% | 0.96% | 238.1bp (417.9bp) | Ot\_0263(RRM)/Ostreococcus\_tauri-RNCMPT00263-PBM/HughesRNA(0.705) More Information \| Similar Motifs Found | motif file (matrix) |
| 3 \* | G T C A A T C G G T C A A C T G T G C A T A G C C G A T G C A T T G A C C A G T | 1e-6 | -1.517e+01 | 1.68% | 0.13% | 337.5bp (302.3bp) | REF2(RRM)/Drosophila\_melanogaster-RNCMPT00059-PBM/HughesRNA(0.680) More Information \| Similar Motifs Found | motif file (matrix) |
| 4 \* | G T C A A C G T A C T G A C T G G T C A A C T G A G T C A C G T A C T G A G T C | 1e-6 | -1.450e+01 | 1.89% | 0.20% | 271.2bp (595.9bp) | Pp\_0237(RRM)/Physcomitrella\_patens-RNCMPT00237-PBM/HughesRNA(0.674) More Information \| Similar Motifs Found | motif file (matrix) |
| 5 \* | C G A T A G C T A C T G C G A T A T C G G T C A C T A G G T C A T G A C A C T G | 1e-5 | -1.351e+01 | 6.09% | 2.21% | 195.2bp (444.8bp) | Six4/MA0204.1/Jaspar(0.709) More Information \| Similar Motifs Found | motif file (matrix) |
| 6 \* | T C G A G A T C G T A C G T A C G A T C T C A G C A T G G C T A G T A C G A C T | 1e-5 | -1.233e+01 | 5.04% | 1.73% | 414.2bp (543.8bp) | GSM1/MA0308.1/Jaspar(0.664) More Information \| Similar Motifs Found | motif file (matrix) |
| 7 \* | A C T G A G T C A C T G A C G T A C G T C G T A C G A T A G T C | 1e-4 | -1.144e+01 | 5.04% | 1.84% | 358.8bp (455.3bp) | GATA5/MA0766.2/Jaspar(0.795) More Information \| Similar Motifs Found | motif file (matrix) |
| 8 \* | A T C G A T C G G C A T A C G T T A G C T G A C C T G A T G A C | 1e-4 | -1.114e+01 | 6.72% | 2.92% | 225.1bp (487.1bp) | TCP20(TCP)/col-TCP20-DAP-Seq(GSE60143)/Homer(0.786) More Information \| Similar Motifs Found | motif file (matrix) |
| 9 \* | A C T G C G T A A G T C A G T C A C T G C T A G A C G T A G T C C G T A A C G T | 1e-4 | -1.084e+01 | 0.84% | 0.02% | 311.0bp (1.3bp) | PH0140.1\_Pknox1/Jaspar(0.694) More Information \| Similar Motifs Found | motif file (matrix) |
| 10 \* | A C G T T G C A A T G C C A T G A G T C A G T C A T C G C G T A G A T C G C T A | 1e-4 | -1.080e+01 | 2.10% | 0.40% | 203.4bp (475.1bp) | ERF021/MA1233.2/Jaspar(0.796) More Information \| Similar Motifs Found | motif file (matrix) |
| 11 \* | A C T G A C G T G T C A A G C T C T A G A G T C C G A T C T A G A C G T C G A T | 1e-4 | -1.011e+01 | 4.83% | 1.86% | 197.3bp (482.4bp) | ZNF317/MA1593.1/Jaspar(0.783) More Information \| Similar Motifs Found | motif file (matrix) |
| 12 \* | C G T A C A G T G C T A C T G A C T G A G C T A T G A C G C A T G C A T A C T G | 1e-4 | -9.412e+00 | 5.88% | 2.63% | 295.7bp (438.5bp) | abd-B/dmmpmm(Noyes)/fly(0.747) More Information \| Similar Motifs Found | motif file (matrix) |
| 13 \* | A C T G A G T C A C G T A C G T A G T C C G T A A G T C G T C A A G T C A C G T | 1e-3 | -9.105e+00 | 1.05% | 0.13% | 102.3bp (292.6bp) | RBM24(RRM)/Homo\_sapiens-RNCMPT00184-PBM/HughesRNA(0.673) More Information \| Similar Motifs Found | motif file (matrix) |
| 14 \* | C G T A A C T G A C T G A C G T A C G T A T C G A C T G A C T G A G T C C T G A | 1e-3 | -8.190e+00 | 0.84% | 0.07% | 202.2bp (108.0bp) | ZNF449/MA1656.1/Jaspar(0.773) More Information \| Similar Motifs Found | motif file (matrix) |
| 15 \* | A C T G A G T C C T G A A C G T C G T A A C G T A C G T A G T C C G T A A C T G | 1e-3 | -8.190e+00 | 0.84% | 0.08% | 153.0bp (156.6bp) | Pit1(Homeobox)/GCrat-Pit1-ChIP-Seq(GSE58009)/Homer(0.779) More Information \| Similar Motifs Found | motif file (matrix) |
| 16 \* | A C T G C G T A A C T G C T G A A C T G G T A C A C T G C G T A A C T G G A C T | 1e-3 | -7.837e+00 | 4.83% | 2.18% | 256.1bp (448.9bp) | Trl(Zf)/S2-GAGAfactor-ChIP-Seq(GSE40646)/Homer(0.795) More Information \| Similar Motifs Found | motif file (matrix) |
| 17 \* | A G T C A C G T A C T G A G T C C G T A A C T G A C G T C G T A A C G T A G T C | 1e-2 | -6.688e+00 | 0.84% | 0.11% | 116.7bp (216.6bp) | TUT1(RRM,Znf)/Homo\_sapiens-RNCMPT00075-PBM/HughesRNA(0.728) More Information \| Similar Motifs Found | motif file (matrix) |
| 18 \* | C G T A A C T G A G T C A C G T A C T G A C T G A G T C A C T G | 1e-2 | -5.981e+00 | 2.73% | 1.12% | 195.4bp (522.8bp) | Vts1p(SAM)/Saccharomyces\_cerevisiae-RNCMPT00111-PBM/HughesRNA(0.793) More Information \| Similar Motifs Found | motif file (matrix) |
| 19 \* | A G T C A G T C C G T A A G T C C G T A A G T C A C G T A C T G A G T C A C G T | 1e-2 | -5.593e+00 | 0.63% | 0.08% | 564.5bp (136.7bp) | CG7804(RRM)/Drosophila\_melanogaster-RNCMPT00146-PBM/HughesRNA(0.753) More Information \| Similar Motifs Found | motif file (matrix) |
| 20 \* | T G A C A T C G A C G T C G A T A G C T G C T A A G T C T A G C G T C A G C T A | 1e-2 | -5.229e+00 | 2.52% | 1.06% | 213.6bp (409.7bp) | NFAT5/MA0606.1/Jaspar(0.757) More Information \| Similar Motifs Found | motif file (matrix) |
| 21 \* | A G T C A G T C A C G T A C T G C G T A A C T G C G T A A C G T C G T A C G T A | 1e-1 | -4.562e+00 | 0.42% | 0.05% | 99.7bp (0.0bp) | TRPS1(Zf)/MCF7-TRPS1-ChIP-Seq(GSE107013)/Homer(0.738) More Information \| Similar Motifs Found | motif file (matrix) |
| 22 \* | C G T A A C T G G T C A A C T G A G T C C T G A A G T C A T C G A G T C A G C T | 1e-1 | -4.562e+00 | 0.42% | 0.05% | 184.1bp (320.7bp) | AFT2/AFT2\_H2O2Lo/10-RCS1[~AFT2](Harbison)/Yeast(0.677) More Information \| Similar Motifs Found | motif file (matrix) |
| 23 \* | A C T G A C T G A G T C A C G T C G T A A C T G C G T A A C G T | 1e-1 | -3.776e+00 | 2.94% | 1.63% | 244.6bp (438.6bp) | ABF2/MA0266.1/Jaspar(0.700) More Information \| Similar Motifs Found | motif file (matrix) |
| 24 \* | C G T A A C T G C G T A C G A T C T G A A C T G A C G T A C G T | 1e-1 | -3.213e+00 | 15.34% | 12.54% | 412.0bp (464.9bp) | CCA1/MA0972.1/Jaspar(0.821) More Information \| Similar Motifs Found | motif file (matrix) |
| 25 \* | A G T C A G T C C G T A A C G T A G T C C G T A A G T C A C G T A C T G C G T A | 1e-1 | -3.175e+00 | 0.63% | 0.19% | 481.0bp (287.3bp) | Unknown2/Drosophila-Promoters/Homer(0.720) More Information \| Similar Motifs Found | motif file (matrix) |
| 26 \* | A G T C A C T G A G T C A C T G G T A C A G T C | 1e0 | -2.226e+00 | 14.29% | 12.31% | 286.9bp (432.6bp) | MYB124/MA1426.1/Jaspar(0.908) More Information \| Similar Motifs Found | motif file (matrix) |
| 27 \* | A G C T A C G T C G T A A G T C A G T C C G A T A C G T A C T G | 1e0 | -2.111e+00 | 11.55% | 9.86% | 277.3bp (451.9bp) | Tag/dmmpmm(Papatsenko)/fly(0.804) More Information \| Similar Motifs Found | motif file (matrix) |
| 28 \* | C T G A C T G A T C G A G T C A C G T A G C T A C G T A C G T A G T C A G T C A | 1e0 | -2.107e+00 | 12.82% | 11.05% | 493.1bp (539.7bp) | SeqBias: polyA-repeat(0.957) More Information \| Similar Motifs Found | motif file (matrix) |
| 29 \* | T G A C G T A C C T G A T A G C T C G A G A T C T C A G A G C T | 1e0 | -2.019e+00 | 12.39% | 10.71% | 307.7bp (463.1bp) | BZR1/MA0550.2/Jaspar(0.819) More Information \| Similar Motifs Found | motif file (matrix) |
| 30 \* | A C T G A C T G A T G C A G C T A C T G A C G T | 1e0 | -6.564e-01 | 25.21% | 25.22% | 377.7bp (473.9bp) | AFT2/MA0270.1/Jaspar(0.850) More Information \| Similar Motifs Found | motif file (matrix) |
| 31 \* | A G T C C G T A A C G T C G T A A C G T C G T A | 1e0 | -1.027e-01 | 38.45% | 41.26% | 396.5bp (466.6bp) | MOT2/MA0379.1/Jaspar(0.912) More Information \| Similar Motifs Found | motif file (matrix) |
