## Supplemental Table 1 for "The Establishment of Cell-Type Specific Gene Regulation in the Sea Urchin Embryo": Hpf12_EctoVeg_homerResults.html

homer\_motifs\_wbackground/512cell\_EctoVeg/ - Homer de novo Motif Results


### Homer *de novo* Motif Results (homer\_motifs\_wbackground/512cell\_EctoVeg/)

Known Motif Enrichment Results  
Gene Ontology Enrichment Results  
If Homer is having trouble matching a motif to a known motif, try copy/pasting the matrix file into
STAMP  
More information on motif finding results: HOMER
| Description of Results
| Tips
  
Total target sequences = 197  
Total background sequences = 3391  
\* - possible false positive  

|  |  |  |  |  |  |  |  |  |
| --- | --- | --- | --- | --- | --- | --- | --- | --- |
| Rank | Motif | P-value | log P-pvalue | % of Targets | % of Background | STD(Bg STD) | Best Match/Details | Motif File |
| 1 \* | C G A T A C G T A C T G C G T A C G T A G T A C A G T C G T A C A C T G A C T G | 1e-11 | -2.567e+01 | 6.60% | 0.46% | 333.0bp (310.7bp) | YDR026C(MacIsaac)/Yeast(0.758) More Information \| Similar Motifs Found | motif file (matrix) |
| 2 \* | A G C T A C T G G T C A A G T C A C G T T G A C C A T G T A G C A C G T A C T G | 1e-8 | -2.028e+01 | 9.64% | 1.71% | 257.5bp (507.8bp) | BAS1/BAS1\_SM/2-BAS1(Harbison)/Yeast(0.739) More Information \| Similar Motifs Found | motif file (matrix) |
| 3 \* | A G T C A G T C A G T C A C G T G A T C C G T A C G T A A C T G A G T C A C G T | 1e-8 | -2.019e+01 | 5.58% | 0.45% | 376.8bp (337.1bp) | ZAP1/ZAP1\_YPD/[](Harbison)/Yeast(0.750) More Information \| Similar Motifs Found | motif file (matrix) |
| 4 \* | A T C G G T A C C T A G A C G T A C G T C G T A C G T A A C T G A G T C A G T C | 1e-7 | -1.779e+01 | 5.58% | 0.57% | 388.8bp (505.7bp) | AT3G10030(Trihelix)/colamp-AT3G10030-DAP-Seq(GSE60143)/Homer(0.740) More Information \| Similar Motifs Found | motif file (matrix) |
| 5 \* | C G T A A C T G A G T C A C G T A G T C C G A T A C T G A C T G C G T A A G T C | 1e-7 | -1.778e+01 | 8.12% | 1.37% | 415.1bp (434.2bp) | ZBTB26/MA1579.1/Jaspar(0.698) More Information \| Similar Motifs Found | motif file (matrix) |
| 6 \* | A C G T A C T G C A T G A G T C G A T C C G T A G T A C A G C T A C G T A G T C | 1e-7 | -1.759e+01 | 5.08% | 0.45% | 437.0bp (380.1bp) | bZIP911(1)(bZIP)/Antirrhinum majus/AthaMap(0.693) More Information \| Similar Motifs Found | motif file (matrix) |
| 7 \* | A C G T A C T G A C G T A C T G A C G T A C G T A C T G A C G T A C T G A G T C | 1e-7 | -1.639e+01 | 3.55% | 0.19% | 480.0bp (172.2bp) | UNC-75(RRM)/Caenorhabditis\_elegans-RNCMPT00081-PBM/HughesRNA(0.835) More Information \| Similar Motifs Found | motif file (matrix) |
| 8 \* | A G T C A C T G C G T A G T C A A C G T A C G T A T C G C T G A A G T C C T G A | 1e-6 | -1.531e+01 | 10.15% | 2.59% | 318.2bp (503.7bp) | Vis/dmmpmm(Noyes\_hd)/fly(0.692) More Information \| Similar Motifs Found | motif file (matrix) |
| 9 \* | C T G A C A G T A C G T A C G T A T C G G A C T A T G C A C T G C G T A G T A C | 1e-6 | -1.516e+01 | 12.69% | 3.91% | 346.9bp (502.7bp) | ARF29/MA1692.1/Jaspar(0.705) More Information \| Similar Motifs Found | motif file (matrix) |
| 10 \* | A G T C C G T A C G T A C G T A C G T A C G T A A G T C A G T C C G T A A C T G | 1e-6 | -1.469e+01 | 2.03% | 0.02% | 180.0bp (0.0bp) | AT1G76870(Trihelix)/col-AT1G76870-DAP-Seq(GSE60143)/Homer(0.783) More Information \| Similar Motifs Found | motif file (matrix) |
| 11 \* | T A G C T A G C G T C A A C T G G C A T T G C A G T A C G A T C C T G A G T A C | 1e-6 | -1.431e+01 | 32.99% | 18.33% | 382.9bp (462.0bp) | odd/dmmpmm(Noyes)/fly(0.688) More Information \| Similar Motifs Found | motif file (matrix) |
| 12 \* | A G T C G T C A C G T A G C T A C G T A G A T C T A C G A C G T C G A T A G C T | 1e-5 | -1.376e+01 | 14.72% | 5.41% | 483.4bp (406.5bp) | SFP1/SacCer-Promoters/Homer(0.687) More Information \| Similar Motifs Found | motif file (matrix) |
| 13 \* | A C G T C G T A A C G T C G T A C G T A C G T A C G T A A C T G A C G T C G T A | 1e-5 | -1.370e+01 | 3.55% | 0.28% | 265.3bp (390.2bp) | TATA-box/Drosophila-Promoters/Homer(0.797) More Information \| Similar Motifs Found | motif file (matrix) |
| 14 \* | T G C A T G C A A G C T T G C A C T G A G T C A G T C A G T C A T C A G A T C G | 1e-4 | -1.117e+01 | 41.62% | 27.53% | 372.1bp (453.2bp) | dof42(C2C2dof)/col-dof42-DAP-Seq(GSE60143)/Homer(0.804) More Information \| Similar Motifs Found | motif file (matrix) |
| 15 \* | A G T C C T G A A G T C A T C G A C G T A C T G A G T C C G T A A T G C A C T G | 1e-4 | -1.039e+01 | 2.03% | 0.09% | 190.8bp (461.7bp) | SOK2/MA0385.1/Jaspar(0.770) More Information \| Similar Motifs Found | motif file (matrix) |
| 16 \* | A C T G A T C G C A T G T G A C T C A G A C T G G T A C G A T C | 1e-4 | -9.555e+00 | 36.55% | 24.19% | 440.9bp (462.5bp) | ERF13/MA1004.1/Jaspar(0.807) More Information \| Similar Motifs Found | motif file (matrix) |
| 17 \* | A C T G C G T A A G T C A C G T A C T G C G T A A C T G A C G T A C G T C T G A | 1e-3 | -8.431e+00 | 2.03% | 0.17% | 319.1bp (330.9bp) | Initiator/Drosophila-Promoters/Homer(0.732) More Information \| Similar Motifs Found | motif file (matrix) |
| 18 \* | A C G T C G T A A C T G C T A G A C T G A C G T A C T G A G T C A C G T C G T A | 1e-2 | -6.394e+00 | 1.52% | 0.12% | 279.3bp (375.6bp) | AFT2/AFT2\_H2O2Lo/10-RCS1[~AFT2](Harbison)/Yeast(0.765) More Information \| Similar Motifs Found | motif file (matrix) |
| 19 \* | A C G T C G T A C G T A C G T A A G T C C G T A A C T G A C G T C G T A A G T C | 1e-2 | -6.258e+00 | 2.03% | 0.26% | 66.2bp (416.7bp) | Foxq1/MA0040.1/Jaspar(0.766) More Information \| Similar Motifs Found | motif file (matrix) |
| 20 \* | A G T C C G A T A C T G A C T G A G T C A C T G | 1e-2 | -5.442e+00 | 22.34% | 15.08% | 388.0bp (481.1bp) | brk/dmmpmm(Papatsenko)/fly(0.847) More Information \| Similar Motifs Found | motif file (matrix) |
| 21 \* | A C G T C G T A A C T G A G T C C G T A C G T A A C T G C G T A | 1e-1 | -3.391e+00 | 5.08% | 2.62% | 318.2bp (397.3bp) | MATR3(RRM)/Homo\_sapiens-RNCMPT00037-PBM/HughesRNA(0.699) More Information \| Similar Motifs Found | motif file (matrix) |
| 22 \* | A G T C C G T A C G T A A C G T A G T C C G T A A C T G A C T G A C G T A C T G | 1e-1 | -3.385e+00 | 1.02% | 0.15% | 252.3bp (369.7bp) | SREBF1(var.2)/MA0829.2/Jaspar(0.766) More Information \| Similar Motifs Found | motif file (matrix) |
| 23 \* | A C G T C G T A A G T C A C G T A G T C A G T C C G T A A C T G A C T G A G T C | 1e-1 | -2.888e+00 | 0.51% | 0.06% | 292.0bp (43.3bp) | WIP5(C2H2)/colamp-WIP5-DAP-Seq(GSE60143)/Homer(0.751) More Information \| Similar Motifs Found | motif file (matrix) |
| 24 \* | A G C T A T C G A G C T A C T G C G A T A C T G C G A T A C T G C G A T A T C G | 1e0 | -1.530e+00 | 8.63% | 7.01% | 156.4bp (442.8bp) | SeqBias: CA-repeat(0.991) More Information \| Similar Motifs Found | motif file (matrix) |
| 25 \* | C G T A A C G T A G T C A C G T C G T A A C G T A C G T A G T C A G T C A G T C | 1e0 | -1.389e+00 | 0.51% | 0.16% | 256.7bp (235.0bp) | kni/dmmpmm(Papatsenko)/fly(0.690) More Information \| Similar Motifs Found | motif file (matrix) |
