## Supplemental Table 1 for "The Establishment of Cell-Type Specific Gene Regulation in the Sea Urchin Embryo": Hpf12_PMCs_homerResults.html

homer\_motifs\_wbackground/512cell\_PMCs/ - Homer de novo Motif Results


### Homer *de novo* Motif Results (homer\_motifs\_wbackground/512cell\_PMCs/)

Known Motif Enrichment Results  
Gene Ontology Enrichment Results  
If Homer is having trouble matching a motif to a known motif, try copy/pasting the matrix file into
STAMP  
More information on motif finding results: HOMER
| Description of Results
| Tips
  
Total target sequences = 1355  
Total background sequences = 2271  
\* - possible false positive  

|  |  |  |  |  |  |  |  |  |
| --- | --- | --- | --- | --- | --- | --- | --- | --- |
| Rank | Motif | P-value | log P-pvalue | % of Targets | % of Background | STD(Bg STD) | Best Match/Details | Motif File |
| 1 | G T A C G T C A A G T C C G A T G C A T A T G C A G T C A C G T T A C G G A C T | 1e-157 | -3.624e+02 | 37.56% | 10.16% | 387.5bp (424.4bp) | ETS1(ETS)/Jurkat-ETS1-ChIP-Seq(GSE17954)/Homer(0.975) More Information \| Similar Motifs Found | motif file (matrix) |
| 2 | C G A T C A T G C T G A A G C T G C A T G T A C C T G A G C A T A C T G T G C A | 1e-35 | -8.087e+01 | 19.48% | 8.60% | 390.7bp (404.1bp) | Fos(bZIP)/TSC-Fos-ChIP-Seq(GSE110950)/Homer(0.916) More Information \| Similar Motifs Found | motif file (matrix) |
| 3 | G A C T G T C A C G T A G C A T G A C T C G T A C T A G T G A C G T C A G T A C | 1e-31 | -7.356e+01 | 33.21% | 19.54% | 387.5bp (452.7bp) | Ro/dmmpmm(Noyes\_hd)/fly(0.904) More Information \| Similar Motifs Found | motif file (matrix) |
| 4 | C G T A A C G T A C G T C G T A G T C A A C G T A C G T C G T A C G T A A C G T | 1e-26 | -6.207e+01 | 2.29% | 0.15% | 254.6bp (345.7bp) | PH0136.1\_Phox2b/Jaspar(0.896) More Information \| Similar Motifs Found | motif file (matrix) |
| 5 | A G T C C T A G A G T C A C T G C G T A A C T G A G C T A G T C | 1e-22 | -5.158e+01 | 8.34% | 2.88% | 445.1bp (445.3bp) | PL0018.1\_hlh-25/Jaspar(0.733) More Information \| Similar Motifs Found | motif file (matrix) |
| 6 | A C G T A C T G A C T G C G T A C G A T C G T A A C G T A G T C | 1e-19 | -4.571e+01 | 16.31% | 8.52% | 443.4bp (406.4bp) | SD0003.1\_at\_AC\_acceptor/Jaspar(0.751) More Information \| Similar Motifs Found | motif file (matrix) |
| 7 | G A T C G C T A A C T G A C G T G T C A G A T C C G T A C G A T G T A C G T A C | 1e-19 | -4.512e+01 | 10.18% | 4.30% | 419.9bp (454.5bp) | SPL9/MA1322.1/Jaspar(0.736) More Information \| Similar Motifs Found | motif file (matrix) |
| 8 | C G T A A C G T A C G T G T A C A G T C C T G A A C G T A G T C G T C A C G A T | 1e-17 | -3.957e+01 | 1.18% | 0.05% | 376.3bp (0.0bp) | ATF4/MA0833.2/Jaspar(0.878) More Information \| Similar Motifs Found | motif file (matrix) |
| 9 | C G T A C G T A A C T G A G T C A C G T A C G T A C T G G T A C A C G T A C G T | 1e-16 | -3.751e+01 | 1.62% | 0.16% | 277.5bp (423.4bp) | Nr2e3/MA0164.1/Jaspar(0.790) More Information \| Similar Motifs Found | motif file (matrix) |
| 10 | A C T G C T G A A G T C C T A G A C G T A G T C C G T A A C G T | 1e-14 | -3.387e+01 | 18.82% | 11.48% | 433.7bp (461.0bp) | TGA10(bZIP)/colamp-TGA10-DAP-Seq(GSE60143)/Homer(0.980) More Information \| Similar Motifs Found | motif file (matrix) |
| 11 | G C T A A C T G A C T G A G T C G T A C A C G T C G T A A C T G A C T G A G T C | 1e-13 | -3.171e+01 | 1.92% | 0.28% | 433.1bp (348.2bp) | Zfx/MA0146.2/Jaspar(0.846) More Information \| Similar Motifs Found | motif file (matrix) |
| 12 | C G T A C G A T A G T C C G T A G T C A A G T C A C G T A C G T A C G T A G T C | 1e-12 | -2.987e+01 | 0.96% | 0.07% | 277.6bp (83.5bp) | Six1(Homeobox)/Myoblast-Six1-ChIP-Chip(GSE20150)/Homer(0.727) More Information \| Similar Motifs Found | motif file (matrix) |
| 13 | A T C G G C A T G C T A A G T C C T G A T C A G A C T G A C G T | 1e-12 | -2.866e+01 | 29.52% | 21.21% | 421.2bp (469.8bp) | sna/dmmpmm(Papatsenko)/fly(0.759) More Information \| Similar Motifs Found | motif file (matrix) |
| 14 | A G T C A T G C C A T G G A C T A G T C T A C G C G A T A C T G A T C G C G T A | 1e-12 | -2.827e+01 | 1.92% | 0.32% | 438.6bp (508.5bp) | RSF1(RRM)/Drosophila\_melanogaster-RNCMPT00061-PBM/HughesRNA(0.751) More Information \| Similar Motifs Found | motif file (matrix) |
| 15 \* | C T G A A G C T A T C G C G A T A C T G G T C A G T A C A C G T C G T A C T G A | 1e-11 | -2.678e+01 | 2.80% | 0.74% | 334.7bp (349.7bp) | MITF(bHLH)/MastCells-MITF-ChIP-Seq(GSE48085)/Homer(0.725) More Information \| Similar Motifs Found | motif file (matrix) |
| 16 \* | A G T C A C T G A C G T A C G T A G T C A C T G A C T G A C G T | 1e-11 | -2.641e+01 | 1.85% | 0.32% | 508.9bp (310.6bp) | Lm\_0254(RRM)/Leishmania\_major-RNCMPT00254-PBM/HughesRNA(0.760) More Information \| Similar Motifs Found | motif file (matrix) |
| 17 \* | G T A C G C T A A G T C C G T A A C T G A G T C A C G T A C T G | 1e-10 | -2.439e+01 | 14.69% | 9.14% | 401.3bp (387.1bp) | Ptf1a(var.2)/MA1619.1/Jaspar(0.937) More Information \| Similar Motifs Found | motif file (matrix) |
| 18 \* | A C G T A C T G A C G T A C T G C G T A A C G T A G C T A C T G A C G T A C T G | 1e-10 | -2.428e+01 | 3.03% | 0.91% | 459.8bp (459.2bp) | SeqBias: CA-repeat(0.790) More Information \| Similar Motifs Found | motif file (matrix) |
| 19 \* | A C T G A C G T A C G T A G T C C G T A A C T G C T A G C G T A C G T A C G A T | 1e-10 | -2.307e+01 | 1.33% | 0.21% | 393.7bp (244.4bp) | TEAD(TEA)/Fibroblast-PU.1-ChIP-Seq(Unpublished)/Homer(0.703) More Information \| Similar Motifs Found | motif file (matrix) |
| 20 \* | G T A C G T A C A G T C A C G T A G T C A G T C A G T C A C G T A C G T A G T C | 1e-9 | -2.210e+01 | 1.55% | 0.29% | 303.5bp (476.3bp) | HNRNPH2(RRM)/Homo\_sapiens-RNCMPT00160-PBM/HughesRNA(0.811) More Information \| Similar Motifs Found | motif file (matrix) |
| 21 \* | C G T A A C G T A C T G A G T C C G T A A C G T A C T G A C G T A C G T A C T G | 1e-9 | -2.089e+01 | 1.11% | 0.15% | 386.3bp (218.5bp) | FUS3/MA0565.2/Jaspar(0.841) More Information \| Similar Motifs Found | motif file (matrix) |
| 22 \* | A C T G C T A G A G T C C G T A A G C T A G T C A G T C A C T G | 1e-8 | -1.955e+01 | 4.50% | 1.95% | 450.8bp (300.0bp) | PB0077.1\_Spdef\_1/Jaspar(0.725) More Information \| Similar Motifs Found | motif file (matrix) |
| 23 \* | A C G T A G T C C G T A A C G T A C G T A C G T C G T A C G T A C G T A A C T G | 1e-8 | -1.875e+01 | 1.03% | 0.14% | 376.2bp (263.3bp) | eve/dmmpmm(Bergman)/fly(0.788) More Information \| Similar Motifs Found | motif file (matrix) |
| 24 \* | C G T A A C T G A G T C A C T G A T G C A T C G A T G C A C T G | 1e-8 | -1.848e+01 | 6.27% | 3.22% | 306.8bp (396.0bp) | RBM8A(RRM)/Homo\_sapiens-RNCMPT00056-PBM/HughesRNA(0.877) More Information \| Similar Motifs Found | motif file (matrix) |
| 25 \* | A G T C A C G T A C G T A C G T A C T G A C G T A G C T A G C T | 1e-7 | -1.795e+01 | 22.07% | 16.27% | 432.6bp (431.9bp) | SOX10/MA0442.2/Jaspar(0.948) More Information \| Similar Motifs Found | motif file (matrix) |
| 26 \* | A C G T A G T C A C G T A G T C A C T G A G T C A G T C A G T C | 1e-7 | -1.794e+01 | 3.32% | 1.28% | 460.7bp (415.2bp) | hkb/MA0450.1/Jaspar(0.800) More Information \| Similar Motifs Found | motif file (matrix) |
| 27 \* | A C T G C G T A A C G T A C T G A C G T A C G T A G T C A C T G | 1e-6 | -1.498e+01 | 2.29% | 0.80% | 389.6bp (463.4bp) | YBX1(CSD)/Homo\_sapiens-RNCMPT00083-PBM/HughesRNA(0.848) More Information \| Similar Motifs Found | motif file (matrix) |
