## Supplemental Table 1 for "The Establishment of Cell-Type Specific Gene Regulation in the Sea Urchin Embryo": Hpf12_Veg1_homerResults.html

homer\_motifs\_wbackground/512cell\_Veg1/ - Homer de novo Motif Results


### Homer *de novo* Motif Results (homer\_motifs\_wbackground/512cell\_Veg1/)

Known Motif Enrichment Results  
Gene Ontology Enrichment Results  
If Homer is having trouble matching a motif to a known motif, try copy/pasting the matrix file into
STAMP  
More information on motif finding results: HOMER
| Description of Results
| Tips
  
Total target sequences = 236  
Total background sequences = 3362  
\* - possible false positive  

|  |  |  |  |  |  |  |  |  |
| --- | --- | --- | --- | --- | --- | --- | --- | --- |
| Rank | Motif | P-value | log P-pvalue | % of Targets | % of Background | STD(Bg STD) | Best Match/Details | Motif File |
| 1 | A G T C A G C T C T A G A C T G A G T C A C G T A G T C A C G T A T G C A G C T | 1e-12 | -2.801e+01 | 8.90% | 1.16% | 348.3bp (409.6bp) | brk/dmmpmm(Bergman)/fly(0.822) More Information \| Similar Motifs Found | motif file (matrix) |
| 2 \* | C G T A A C T G A G T C A T G C A G C T A T G C C G T A G A C T T C G A A G T C | 1e-11 | -2.736e+01 | 7.20% | 0.69% | 496.3bp (406.8bp) | PUF68(RRM)/Drosophila\_melanogaster-RNCMPT00141-PBM/HughesRNA(0.753) More Information \| Similar Motifs Found | motif file (matrix) |
| 3 \* | A C G T C T A G A C G T A C T G A G T C C G T A A C T G C G T A A T G C A C T G | 1e-10 | -2.318e+01 | 6.36% | 0.67% | 318.8bp (314.9bp) | PB0044.1\_Mtf1\_1/Jaspar(0.726) More Information \| Similar Motifs Found | motif file (matrix) |
| 4 \* | A T C G A T G C C G T A A T C G A C T G A C G T G T C A A C T G A G T C C T G A | 1e-9 | -2.210e+01 | 6.78% | 0.84% | 432.5bp (578.3bp) | PB0155.1\_Osr2\_2/Jaspar(0.831) More Information \| Similar Motifs Found | motif file (matrix) |
| 5 \* | C T A G A C G T A C T G C G T A G T C A G T A C C T A G A C T G A G T C A T C G | 1e-8 | -1.977e+01 | 3.81% | 0.22% | 416.9bp (445.2bp) | AT3G10030(Trihelix)/colamp-AT3G10030-DAP-Seq(GSE60143)/Homer(0.761) More Information \| Similar Motifs Found | motif file (matrix) |
| 6 \* | G T C A T A G C A G C T A C G T T C G A T C G A T C G A T G C A T A C G T A G C | 1e-8 | -1.969e+01 | 2.97% | 0.10% | 355.8bp (306.1bp) | KHDRBS1(KH)/Mus\_musculus-RNCMPT00062-PBM/HughesRNA(0.850) More Information \| Similar Motifs Found | motif file (matrix) |
| 7 \* | C T A G C G A T A G C T C G T A T A G C C G T A C A G T T G C A C A T G C A T G | 1e-8 | -1.952e+01 | 18.64% | 7.08% | 455.6bp (418.3bp) | PF13\_0315(RRM)/Plasmodium\_falciparum-RNCMPT00200-PBM/HughesRNA(0.708) More Information \| Similar Motifs Found | motif file (matrix) |
| 8 \* | A C T G A C G T A G T C A G T C A C G T A T G C A C T G C G T A A C T G A C G T | 1e-8 | -1.863e+01 | 3.81% | 0.25% | 346.9bp (506.6bp) | XBP1(MacIsaac)/Yeast(0.805) More Information \| Similar Motifs Found | motif file (matrix) |
| 9 \* | A C T G C G T A A G T C A C T G C G T A A C T G A C T G A C G T A C G T A C T G | 1e-7 | -1.817e+01 | 2.12% | 0.03% | 533.8bp (0.0bp) | FXR2(KH)/Homo\_sapiens-RNCMPT00020-PBM/HughesRNA(0.669) More Information \| Similar Motifs Found | motif file (matrix) |
| 10 \* | A C G T C G T A C G T A A C G T A C T G A T C G A C T G A C T G A G T C C G T A | 1e-7 | -1.817e+01 | 2.12% | 0.06% | 147.9bp (44.8bp) | Isl1/MA1608.1/Jaspar(0.716) More Information \| Similar Motifs Found | motif file (matrix) |
| 11 \* | C G T A A G T C A G T C A C G T A C G T C G T A A G T C A G T C A C G T A C G T | 1e-7 | -1.774e+01 | 2.97% | 0.14% | 209.3bp (469.2bp) | SD0001.1\_at\_AC\_acceptor/Jaspar(0.799) More Information \| Similar Motifs Found | motif file (matrix) |
| 12 \* | G T C A G T A C C A T G C G A T A C T G A C G T A C T G T C A G C A G T G T C A | 1e-7 | -1.694e+01 | 13.56% | 4.58% | 319.9bp (489.6bp) | BZR1/MA0550.2/Jaspar(0.770) More Information \| Similar Motifs Found | motif file (matrix) |
| 13 \* | A T C G G C T A C G T A G A T C G A C T T A G C A G C T A G C T A G T C C T G A | 1e-7 | -1.691e+01 | 8.90% | 2.12% | 389.7bp (417.5bp) | NR1I3/MA1534.1/Jaspar(0.677) More Information \| Similar Motifs Found | motif file (matrix) |
| 14 \* | A G T C A G T C T C G A C G T A A C T G A C G T A G T C C G T A A C T G G C T A | 1e-7 | -1.683e+01 | 4.66% | 0.53% | 244.8bp (307.9bp) | ASH1/Literature(Harbison)/Yeast(0.765) More Information \| Similar Motifs Found | motif file (matrix) |
| 15 \* | A G T C A C G T A G T C C G T A C G T A C T A G A G T C A C G T A C T G A C G T | 1e-7 | -1.627e+01 | 18.64% | 7.91% | 409.2bp (432.1bp) | bHLH130(bHLH)/col-bHLH130-DAP-Seq(GSE60143)/Homer(0.684) More Information \| Similar Motifs Found | motif file (matrix) |
| 16 \* | A C T G C A G T A C T G T A C G A G T C A C T G A C T G A G C T C A T G C G T A | 1e-6 | -1.582e+01 | 5.51% | 0.84% | 269.1bp (333.3bp) | ABI4(1)(AP2/EREBP)/Zea mays/AthaMap(0.841) More Information \| Similar Motifs Found | motif file (matrix) |
| 17 \* | G C A T A G T C G T C A A G T C G T C A G T C A A T C G C T G A A C G T A T G C | 1e-6 | -1.525e+01 | 10.59% | 3.22% | 445.5bp (507.7bp) | MITF(bHLH)/MastCells-MITF-ChIP-Seq(GSE48085)/Homer(0.753) More Information \| Similar Motifs Found | motif file (matrix) |
| 18 \* | A C T G C G T A A G T C A C T G C G A T A G T C A T G C C G T A A C T G A C G T | 1e-6 | -1.476e+01 | 2.12% | 0.09% | 181.2bp (362.4bp) | TGA10(bZIP)/colamp-TGA10-DAP-Seq(GSE60143)/Homer(0.672) More Information \| Similar Motifs Found | motif file (matrix) |
| 19 \* | A C G T A C G T A C G T A C T G A C T G A G T C A C T G A C T G A G T C A C G T | 1e-5 | -1.321e+01 | 2.54% | 0.17% | 264.7bp (429.5bp) | ERF105(AP2EREBP)/colamp-ERF105-DAP-Seq(GSE60143)/Homer(0.898) More Information \| Similar Motifs Found | motif file (matrix) |
| 20 \* | A C G T A C G T A T C G A G T C G T C A A C T G A C G T C T G A A G T C A C G T | 1e-5 | -1.308e+01 | 4.66% | 0.76% | 243.5bp (343.2bp) | PH0116.1\_Nkx2-9/Jaspar(0.719) More Information \| Similar Motifs Found | motif file (matrix) |
| 21 \* | A C T G A C T G C G T A A C T G A G T C A C G T A G T C A T C G C G T A A C T G | 1e-5 | -1.218e+01 | 2.54% | 0.20% | 264.1bp (194.4bp) | ZNF135/MA1587.1/Jaspar(0.826) More Information \| Similar Motifs Found | motif file (matrix) |
| 22 \* | A G T C C G T A C T G A A C G T C G T A C G T A C G T A G T C A | 1e-4 | -1.135e+01 | 22.46% | 12.39% | 371.0bp (424.9bp) | HOXB13/MA0901.2/Jaspar(0.952) More Information \| Similar Motifs Found | motif file (matrix) |
| 23 \* | A G T C C G T A A G T C G T C A A C G T A G T C | 1e-4 | -1.116e+01 | 63.98% | 50.26% | 413.9bp (420.5bp) | twi/dmmpmm(Papatsenko)/fly(0.776) More Information \| Similar Motifs Found | motif file (matrix) |
| 24 \* | C G T A C G T A C G T A A C T G A G T C A C T G C G T A C G T A | 1e-4 | -1.026e+01 | 8.90% | 3.26% | 372.0bp (469.6bp) | IRF7/MA0772.1/Jaspar(0.798) More Information \| Similar Motifs Found | motif file (matrix) |
| 25 \* | A T C G A T C G C A G T T G C A G C T A G T C A G C T A C T A G A C G T A C T G | 1e-3 | -9.014e+00 | 7.63% | 2.78% | 416.3bp (371.5bp) | AT5G47660/MA1365.2/Jaspar(0.770) More Information \| Similar Motifs Found | motif file (matrix) |
| 26 \* | A G C T C T G A C A T G A C G T C T G A A T C G A C G T C G T A A C T G A G C T | 1e-2 | -5.795e+00 | 5.93% | 2.56% | 213.0bp (585.6bp) | An\_0287(RRM)/Aspergillus\_nidulans-RNCMPT00287-PBM/HughesRNA(0.715) More Information \| Similar Motifs Found | motif file (matrix) |
