## Supplemental Table 1 for "The Establishment of Cell-Type Specific Gene Regulation in the Sea Urchin Embryo": Hpf12_Veg2_homerResults.html

homer\_motifs\_wbackground/512cell\_Veg2/ - Homer de novo Motif Results


### Homer *de novo* Motif Results (homer\_motifs\_wbackground/512cell\_Veg2/)

Known Motif Enrichment Results  
Gene Ontology Enrichment Results  
If Homer is having trouble matching a motif to a known motif, try copy/pasting the matrix file into
STAMP  
More information on motif finding results: HOMER
| Description of Results
| Tips
  
Total target sequences = 683  
Total background sequences = 3523  
\* - possible false positive  

|  |  |  |  |  |  |  |  |  |
| --- | --- | --- | --- | --- | --- | --- | --- | --- |
| Rank | Motif | P-value | log P-pvalue | % of Targets | % of Background | STD(Bg STD) | Best Match/Details | Motif File |
| 1 | C T A G T A C G G A C T C T G A G T A C A T G C G T C A C A T G A G T C C T G A | 1e-69 | -1.606e+02 | 36.60% | 10.81% | 416.5bp (437.0bp) | GCM1/MA0646.1/Jaspar(0.780) More Information \| Similar Motifs Found | motif file (matrix) |
| 2 | A C G T C G T A A C T G A G T C A C G T A C T G A C T G A G T C A C G T A G T C | 1e-30 | -7.093e+01 | 2.78% | 0.05% | 222.3bp (139.0bp) | bZIP18(bZIP)/colamp-bZIP18-DAP-Seq(GSE60143)/Homer(0.709) More Information \| Similar Motifs Found | motif file (matrix) |
| 3 | C G T A A G T C A G T C A G T C C G A T A G T C C G T A A C G T C G T A A C G T | 1e-25 | -5.888e+01 | 3.81% | 0.17% | 395.2bp (330.0bp) | PUF68(RRM)/Drosophila\_melanogaster-RNCMPT00141-PBM/HughesRNA(0.797) More Information \| Similar Motifs Found | motif file (matrix) |
| 4 | A C G T A C G T A G T C A G T C A G T C C G T A A T G C C T A G | 1e-20 | -4.823e+01 | 17.86% | 6.94% | 485.4bp (484.6bp) | Rbpj1(?)/Panc1-Rbpj1-ChIP-Seq(GSE47459)/Homer(0.967) More Information \| Similar Motifs Found | motif file (matrix) |
| 5 | A C T G A C G T C G T A A G T C A G T C A C G T A G T C C A G T A G T C A G T C | 1e-19 | -4.416e+01 | 1.90% | 0.02% | 318.2bp (0.0bp) | SRSF2(RRM)/Homo\_sapiens-RNCMPT00072-PBM/HughesRNA(0.704) More Information \| Similar Motifs Found | motif file (matrix) |
| 6 | A G T C G T A C A G T C A G C T G T C A C T A G A C G T T G C A | 1e-19 | -4.383e+01 | 26.65% | 13.54% | 437.8bp (442.0bp) | TBP3(MYBrelated)/col-TBP3-DAP-Seq(GSE60143)/Homer(0.711) More Information \| Similar Motifs Found | motif file (matrix) |
| 7 | C T G A G T A C G T A C A C T G T G C A A C G T A G T C A T C G A G T C C G T A | 1e-17 | -4.005e+01 | 2.34% | 0.10% | 414.8bp (462.2bp) | wc-1/MA1437.1/Jaspar(0.750) More Information \| Similar Motifs Found | motif file (matrix) |
| 8 | A G C T C G T A A C G T C G T A C G A T A C G T A T G C T G A C | 1e-16 | -3.758e+01 | 37.92% | 23.60% | 430.1bp (433.8bp) | MOT2/MA0379.1/Jaspar(0.806) More Information \| Similar Motifs Found | motif file (matrix) |
| 9 | C G T A A G T C A C G T A C G T C G T A C G T A C G T A C G T A A C T G A G T C | 1e-15 | -3.671e+01 | 2.20% | 0.10% | 318.9bp (85.5bp) | bap/dmmpmm(Noyes\_hd)/fly(0.767) More Information \| Similar Motifs Found | motif file (matrix) |
| 10 | C G A T A C G T A C T G C G T A A C G T A G T C C G T A A T G C A G T C C G T A | 1e-15 | -3.541e+01 | 4.54% | 0.69% | 271.9bp (459.4bp) | PH0016.1\_Cux1\_1/Jaspar(0.781) More Information \| Similar Motifs Found | motif file (matrix) |
| 11 | A T C G T G A C C T G A A T C G G T A C A T C G A T C G G T A C | 1e-14 | -3.304e+01 | 24.16% | 13.12% | 549.5bp (443.1bp) | NHLH1/MA0048.2/Jaspar(0.737) More Information \| Similar Motifs Found | motif file (matrix) |
| 12 | A C T G A C G T A C G T A C G T A C T G C T A G C G T A A C T G C G T A A C G T | 1e-14 | -3.238e+01 | 3.22% | 0.35% | 272.6bp (423.8bp) | YNR063W/MA0432.1/Jaspar(0.671) More Information \| Similar Motifs Found | motif file (matrix) |
| 13 | A C T G A C G T A G T C A G T C A G T C A C G T A C G T A C G T A C T G C G T A | 1e-13 | -3.223e+01 | 2.34% | 0.16% | 176.6bp (258.8bp) | Tcf4(HMG)/Hct116-Tcf4-ChIP-Seq(SRA012054)/Homer(0.775) More Information \| Similar Motifs Found | motif file (matrix) |
| 14 | C G T A A C G T A G T C A C G T A C G T A C G T | 1e-13 | -3.217e+01 | 67.20% | 52.71% | 486.8bp (442.1bp) | ARR2/MA0949.1/Jaspar(0.837) More Information \| Similar Motifs Found | motif file (matrix) |
| 15 | A T G C A C G T C T G A A G C T A T C G T A C G T A C G T A C G T G C A G T C A | 1e-13 | -3.180e+01 | 1.76% | 0.08% | 433.4bp (433.8bp) | ESRP2(RRM)/Homo\_sapiens-RNCMPT00150-PBM/HughesRNA(0.779) More Information \| Similar Motifs Found | motif file (matrix) |
| 16 | A G T C C G T A C G T A A C T G C G T A A G T C A G T C A G T C A C G T A G T C | 1e-13 | -3.023e+01 | 1.90% | 0.11% | 202.7bp (248.0bp) | LRF(Zf)/Erythroblasts-ZBTB7A-ChIP-Seq(GSE74977)/Homer(0.770) More Information \| Similar Motifs Found | motif file (matrix) |
| 17 \* | C G T A A C T G C G T A A G T C A G T C C G T A C G T A A C T G A C G T A C T G | 1e-11 | -2.711e+01 | 1.76% | 0.09% | 404.7bp (314.6bp) | BHLH112/MA0961.1/Jaspar(0.702) More Information \| Similar Motifs Found | motif file (matrix) |
| 18 \* | C G T A A G T C A G T C A C G T A G T C A C G T A C T G A C G T A C G T C G T A | 1e-11 | -2.667e+01 | 1.90% | 0.11% | 421.4bp (756.0bp) | AT3G10030/MA1662.1/Jaspar(0.796) More Information \| Similar Motifs Found | motif file (matrix) |
| 19 \* | C G T A A C T G C G T A A G T C A C G T A C G T A C T G A C G T C G T A C G T A | 1e-11 | -2.664e+01 | 2.05% | 0.14% | 410.5bp (256.3bp) | HMRA2/MA0318.1/Jaspar(0.678) More Information \| Similar Motifs Found | motif file (matrix) |
| 20 \* | A C T G A G T C A T C G A T C G A T G C A T G C C G T A C G T A G C T A G T C A | 1e-11 | -2.664e+01 | 2.05% | 0.16% | 226.7bp (395.1bp) | ERF105(AP2EREBP)/colamp-ERF105-DAP-Seq(GSE60143)/Homer(0.758) More Information \| Similar Motifs Found | motif file (matrix) |
| 21 \* | A G C T A C G T A C T G G C T A C G T A C G T A A G T C C G T A A G T C A G T C | 1e-10 | -2.453e+01 | 3.22% | 0.52% | 440.6bp (330.7bp) | DIG1/DIG1\_YPD/32-STE12(Harbison)/Yeast(0.820) More Information \| Similar Motifs Found | motif file (matrix) |
| 22 \* | G C T A A G C T T C G A A C G T C T A G A C G T C T G A A G T C G T C A A C G T | 1e-10 | -2.372e+01 | 25.33% | 15.65% | 447.1bp (479.4bp) | PUM(PUF)/Drosophila\_melanogaster-RNCMPT00046-PBM/HughesRNA(0.804) More Information \| Similar Motifs Found | motif file (matrix) |
| 23 \* | A G T C A G T C G T A C A G T C A T C G T C G A A G C T A G C T | 1e-9 | -2.229e+01 | 12.88% | 6.30% | 395.4bp (451.3bp) | NHP10/MA0344.1/Jaspar(0.810) More Information \| Similar Motifs Found | motif file (matrix) |
| 24 \* | A C T G A C G T A C G T A C G T C G T A C G T A A C T G A G T C A G T C A G T C | 1e-9 | -2.110e+01 | 1.46% | 0.10% | 429.5bp (240.3bp) | Ct/dmmpmm(Noyes\_hd)/fly(0.743) More Information \| Similar Motifs Found | motif file (matrix) |
| 25 \* | A C T G A C G T C G T A A C T G A C G T A C G T A C G T A C G T C G T A A C G T | 1e-9 | -2.107e+01 | 1.61% | 0.12% | 443.9bp (241.1bp) | abd-B/dmmpmm(Noyes)/fly(0.755) More Information \| Similar Motifs Found | motif file (matrix) |
| 26 \* | A C T G A G T C A C T G A C G T A G C T A C T G A C T G A C T G | 1e-9 | -2.078e+01 | 8.05% | 3.24% | 436.0bp (475.5bp) | YPR022C/MA0436.1/Jaspar(0.713) More Information \| Similar Motifs Found | motif file (matrix) |
| 27 \* | A C G T C G T A A C G T C G T A C G A T C G T A A C G T C G T A | 1e-7 | -1.654e+01 | 30.01% | 21.33% | 438.4bp (432.8bp) | TBP(- other)/several species/AthaMap(0.907) More Information \| Similar Motifs Found | motif file (matrix) |
| 28 \* | A C T G A C T G A C G T A C T G A G T C C G T A A G T C A G T C | 1e-6 | -1.562e+01 | 2.05% | 0.35% | 410.2bp (195.6bp) | AT3G51470(DBP)/col-AT3G51470-DAP-Seq(GSE60143)/Homer(0.884) More Information \| Similar Motifs Found | motif file (matrix) |
| 29 \* | A C G T A C T G A C T G A G T C A G T C A C T G C G T A A C T G | 1e-5 | -1.314e+01 | 6.44% | 2.96% | 306.3bp (406.6bp) | pho/MA1460.1/Jaspar(0.767) More Information \| Similar Motifs Found | motif file (matrix) |
| 30 \* | A C T G A C T G A C T G A C G T A C G T A C T G A G T C A G T C | 1e-5 | -1.295e+01 | 2.78% | 0.77% | 651.9bp (506.0bp) | RFX1/MA0365.1/Jaspar(0.773) More Information \| Similar Motifs Found | motif file (matrix) |
| 31 \* | A C T G A C G T A G T C A C G T A C T G A C G T C T G A A C G T A C T G A C T G | 1e-4 | -1.113e+01 | 27.38% | 20.64% | 478.7bp (441.0bp) | ENOX1(RRM)/Homo\_sapiens-RNCMPT00149-PBM/HughesRNA(0.684) More Information \| Similar Motifs Found | motif file (matrix) |
| 32 \* | A G T C C G T A A C T G C G T A A C T G C G T A A G T C A G T C | 1e-4 | -1.062e+01 | 3.66% | 1.44% | 312.6bp (531.2bp) | SRSF10(RRM)/Homo\_sapiens-RNCMPT00090-PBM/HughesRNA(0.762) More Information \| Similar Motifs Found | motif file (matrix) |
