## Supplemental Table 1 for "The Establishment of Cell-Type Specific Gene Regulation in the Sea Urchin Embryo": Hpf20_ArboralNSM_homerResults.html

homer\_motifs\_wbackground/Hpf20\_ArboralNSM/ - Homer de novo Motif Results


### Homer *de novo* Motif Results (homer\_motifs\_wbackground/Hpf20\_ArboralNSM/)

Known Motif Enrichment Results  
Gene Ontology Enrichment Results  
If Homer is having trouble matching a motif to a known motif, try copy/pasting the matrix file into
STAMP  
More information on motif finding results: HOMER
| Description of Results
| Tips
  
Total target sequences = 2410  
Total background sequences = 6579  
\* - possible false positive  

|  |  |  |  |  |  |  |  |  |
| --- | --- | --- | --- | --- | --- | --- | --- | --- |
| Rank | Motif | P-value | log P-pvalue | % of Targets | % of Background | STD(Bg STD) | Best Match/Details | Motif File |
| 1 | T A C G G A C T C T G A A G T C A G T C G T C A C A T G A G T C C G T A C G A T | 1e-499 | -1.150e+03 | 53.53% | 12.75% | 393.2bp (523.3bp) | GCM1/MA0646.1/Jaspar(0.883) More Information \| Similar Motifs Found | motif file (matrix) |
| 2 | T G A C T G A C A C T G A C T G G C T A G C T A T C A G A G C T | 1e-36 | -8.494e+01 | 46.80% | 34.18% | 468.3bp (459.0bp) | Ets21C/MA0916.1/Jaspar(0.983) More Information \| Similar Motifs Found | motif file (matrix) |
| 3 | C G T A C G A T C G A T C G T A C G T A C G A T G C T A C G A T G A C T C G T A | 1e-26 | -6.028e+01 | 19.25% | 11.71% | 400.1bp (485.9bp) | Arid3b/MA0601.1/Jaspar(0.803) More Information \| Similar Motifs Found | motif file (matrix) |
| 4 | C G T A G A C T A C G T C G T A C G T A A C G T G C A T C G A T G T C A A G C T | 1e-23 | -5.393e+01 | 28.34% | 19.78% | 444.9bp (457.6bp) | tll/dmmpmm(Bergman)/fly(0.833) More Information \| Similar Motifs Found | motif file (matrix) |
| 5 | A C T G C A T G A C T G T A G C A G T C A G T C A G T C A T C G C G A T G T A C | 1e-21 | -5.046e+01 | 2.99% | 0.74% | 551.9bp (475.4bp) | PLAGL2/MA1548.1/Jaspar(0.878) More Information \| Similar Motifs Found | motif file (matrix) |
| 6 | T C G A C G T A C G T A C G T A G C T A G C A T T G C A A C G T A C T G T C A G | 1e-20 | -4.674e+01 | 34.36% | 25.77% | 461.5bp (497.4bp) | Smp\_067420(RRM)/Schistosoma\_mansoni-RNCMPT00232-PBM/HughesRNA(0.768) More Information \| Similar Motifs Found | motif file (matrix) |
| 7 | A T C G C G T A A C G T G T C A C G A T A C T G C T G A A C G T C G T A C T G A | 1e-20 | -4.613e+01 | 3.90% | 1.26% | 375.6bp (369.3bp) | Mecom/MA0029.1/Jaspar(0.845) More Information \| Similar Motifs Found | motif file (matrix) |
| 8 | T G A C A C G T A C G T C G T A T G A C C G T A C A G T C T G A G C A T G T C A | 1e-15 | -3.659e+01 | 10.66% | 6.24% | 437.3bp (461.5bp) | Cf2/dmmpmm(Bergman)/fly(0.803) More Information \| Similar Motifs Found | motif file (matrix) |
| 9 | A C G T A C T G C T G A A C G T C G T A A T C G A C G T C T A G T C A G A C G T | 1e-15 | -3.477e+01 | 4.40% | 1.83% | 440.0bp (609.2bp) | FZF1/MA0298.1/Jaspar(0.779) More Information \| Similar Motifs Found | motif file (matrix) |
| 10 | G C A T C T G A A C T G A C G T A G T C C G T A A C G T C G T A G T A C A G T C | 1e-14 | -3.434e+01 | 1.33% | 0.22% | 612.3bp (190.4bp) | PH0162.1\_Six2/Jaspar(0.651) More Information \| Similar Motifs Found | motif file (matrix) |
| 11 | C G T A C G T A A G T C A C G T A C G T C G T A A C G T A C T G C G T A A C G T | 1e-13 | -3.095e+01 | 0.75% | 0.06% | 709.3bp (333.6bp) | CRC(C2C2YABBY)/col-CRC-DAP-Seq(GSE60143)/Homer(0.719) More Information \| Similar Motifs Found | motif file (matrix) |
| 12 | A C T G A G T C A G T C A C G T A C T G A C G T A G T C A C G T A C G T A G T C | 1e-12 | -2.856e+01 | 1.49% | 0.34% | 570.4bp (309.8bp) | ENOX1(RRM)/Homo\_sapiens-RNCMPT00149-PBM/HughesRNA(0.732) More Information \| Similar Motifs Found | motif file (matrix) |
| 13 \* | A C T G A C G T C G T A A C T G A G T C A G T C A C G T C G T A A G T C A G T C | 1e-10 | -2.441e+01 | 0.50% | 0.04% | 334.2bp (0.0bp) | HRB27C(RRM)/Drosophila\_melanogaster-RNCMPT00093-PBM/HughesRNA(0.704) More Information \| Similar Motifs Found | motif file (matrix) |
| 14 \* | C G T A C G T A A G T C A G T C C G T A C G T A A G T C A G T C C G T A A C T G | 1e-8 | -2.056e+01 | 0.62% | 0.09% | 424.4bp (387.9bp) | P(MYB)/Zea mays/AthaMap(0.811) More Information \| Similar Motifs Found | motif file (matrix) |
| 15 \* | A G T C A C G T A C T G A C T G C G T A A G T C A C G T A C T G | 1e-8 | -1.907e+01 | 2.57% | 1.13% | 458.0bp (349.3bp) | TEAD1/MA0090.3/Jaspar(0.760) More Information \| Similar Motifs Found | motif file (matrix) |
| 16 \* | A G T C A T G C G T C A C G A T C G T A C G A T C T A G A C T G | 1e-8 | -1.905e+01 | 11.91% | 8.48% | 422.9bp (515.9bp) | TCF21(var.2)/MA1568.1/Jaspar(0.914) More Information \| Similar Motifs Found | motif file (matrix) |
| 17 \* | A C G T A G T C A C T G A C G T A C T G C G T A A G T C A C G T | 1e-8 | -1.866e+01 | 4.36% | 2.39% | 480.2bp (455.1bp) | Npas4(bHLH)/Neuron-Npas4-ChIP-Seq(GSE127793)/Homer(0.878) More Information \| Similar Motifs Found | motif file (matrix) |
| 18 \* | A T G C A G C T C G T A A G T C A G T C A C G T A G T C A C G T A G T C A T G C | 1e-5 | -1.318e+01 | 2.86% | 1.57% | 429.3bp (455.4bp) | SRSF2(RRM)/Homo\_sapiens-RNCMPT00072-PBM/HughesRNA(0.746) More Information \| Similar Motifs Found | motif file (matrix) |
| 19 \* | A G T C A C G T A G T C A G T C A G T C A G T C A G T C A G T C | 1e-5 | -1.278e+01 | 9.96% | 7.41% | 444.6bp (492.5bp) | VEZF1/MA1578.1/Jaspar(0.844) More Information \| Similar Motifs Found | motif file (matrix) |
| 20 \* | A C G T A C T G A G T C A C T G C G T A A C G T A G T C A C T G | 1e-5 | -1.275e+01 | 3.49% | 2.05% | 535.9bp (462.5bp) | wc-1/MA1437.1/Jaspar(0.799) More Information \| Similar Motifs Found | motif file (matrix) |
| 21 \* | A C T G A G T C C G T A A C T G A G T C C G T A | 1e-4 | -1.078e+01 | 28.26% | 24.59% | 443.3bp (435.9bp) | Zfp57(Zf)/H1-ZFP57.HA-ChIP-Seq(GSE115387)/Homer(0.785) More Information \| Similar Motifs Found | motif file (matrix) |
| 22 \* | A C T G A C T G A G C T A C T G A C T G C T G A A C G T A C T G | 1e-3 | -8.713e+00 | 10.29% | 8.21% | 456.2bp (404.2bp) | ZNF354C/MA0130.1/Jaspar(0.789) More Information \| Similar Motifs Found | motif file (matrix) |
| 23 \* | C G T A A C T G C G T A A C T G C G T A A C T G C G T A A C T G | 1e-2 | -6.586e+00 | 18.76% | 16.44% | 466.4bp (409.7bp) | Trl/MA0205.2/Jaspar(0.937) More Information \| Similar Motifs Found | motif file (matrix) |
