## Supplemental Table 1 for "The Establishment of Cell-Type Specific Gene Regulation in the Sea Urchin Embryo": Hpf20_EctoApical_homerResults.html

homer\_motifs\_wbackground/Hpf20\_EctoApical/ - Homer de novo Motif Results


### Homer *de novo* Motif Results (homer\_motifs\_wbackground/Hpf20\_EctoApical/)

Known Motif Enrichment Results  
Gene Ontology Enrichment Results  
If Homer is having trouble matching a motif to a known motif, try copy/pasting the matrix file into
STAMP  
More information on motif finding results: HOMER
| Description of Results
| Tips
  
Total target sequences = 1072  
Total background sequences = 8624  
\* - possible false positive  

|  |  |  |  |  |  |  |  |  |
| --- | --- | --- | --- | --- | --- | --- | --- | --- |
| Rank | Motif | P-value | log P-pvalue | % of Targets | % of Background | STD(Bg STD) | Best Match/Details | Motif File |
| 1 | C A T G A T G C A G T C G A C T G C T A G C A T C A G T T C A G C G A T A G T C | 1e-35 | -8.281e+01 | 56.81% | 37.75% | 365.4bp (450.1bp) | SOX15/MA1152.1/Jaspar(0.864) More Information \| Similar Motifs Found | motif file (matrix) |
| 2 | C G A T C T A G C G T A A C G T A C G T A C T G C T G A A G T C C T G A A C G T | 1e-34 | -7.961e+01 | 16.32% | 5.72% | 359.2bp (478.4bp) | Pknox1(Homeobox)/ES-Prep1-ChIP-Seq(GSE63282)/Homer(0.950) More Information \| Similar Motifs Found | motif file (matrix) |
| 3 | T G C A A G T C C G A T G A C T G C A T A G T C G T C A A G T C A G C T C A G T | 1e-27 | -6.262e+01 | 20.80% | 9.64% | 428.1bp (484.2bp) | PRDM1(Zf)/Hela-PRDM1-ChIP-Seq(GSE31477)/Homer(0.946) More Information \| Similar Motifs Found | motif file (matrix) |
| 4 | T A G C T A G C T A C G G C T A A G C T T A G C T C G A C T G A C G A T C G T A | 1e-18 | -4.346e+01 | 9.14% | 3.22% | 402.4bp (548.5bp) | HNF6(Homeobox)/Liver-Hnf6-ChIP-Seq(ERP000394)/Homer(0.852) More Information \| Similar Motifs Found | motif file (matrix) |
| 5 | A T C G A G T C G T C A C G T A A C G T A G C T C G T A A C T G | 1e-14 | -3.437e+01 | 27.15% | 17.37% | 395.8bp (458.5bp) | HOXB2/MA0902.2/Jaspar(0.906) More Information \| Similar Motifs Found | motif file (matrix) |
| 6 \* | C G T A G T C A T C A G C G T A A T C G C A G T A C G T A T G C A C G T T G A C | 1e-11 | -2.666e+01 | 3.92% | 1.09% | 471.4bp (444.4bp) | ZNF274/MA1592.1/Jaspar(0.781) More Information \| Similar Motifs Found | motif file (matrix) |
| 7 \* | C G A T T G A C C G A T C T G A T A G C A G C T A G C T T A C G G T C A A C T G | 1e-11 | -2.656e+01 | 6.81% | 2.73% | 397.3bp (586.4bp) | tin/dmmpmm(Noyes\_hd)/fly(0.686) More Information \| Similar Motifs Found | motif file (matrix) |
| 8 \* | G T C A G T A C C G T A C A G T A C T G T C A G G T A C C G T A C G T A G T A C | 1e-10 | -2.319e+01 | 7.00% | 3.07% | 365.2bp (441.4bp) | RFX3/MA0798.2/Jaspar(0.817) More Information \| Similar Motifs Found | motif file (matrix) |
| 9 \* | C T A G A C T G G T C A A C G T C G T A A G T C A C G T A C G T C G T A C G T A | 1e-9 | -2.251e+01 | 1.12% | 0.08% | 412.5bp (242.1bp) | bap/MA0211.1/Jaspar(0.786) More Information \| Similar Motifs Found | motif file (matrix) |
| 10 \* | A T C G A C T G T A G C C G A T A C T G C G A T T A G C A C T G C T G A A C G T | 1e-9 | -2.203e+01 | 2.99% | 0.78% | 301.2bp (410.1bp) | DDF2(AP2EREBP)/col-DDF2-DAP-Seq(GSE60143)/Homer(0.683) More Information \| Similar Motifs Found | motif file (matrix) |
| 11 \* | A T C G G C T A C T A G G A C T C G T A C T A G A C T G A C G T A T C G A T C G | 1e-9 | -2.143e+01 | 6.06% | 2.59% | 427.9bp (435.2bp) | sna/dmmpmm(SeSiMCMC)/fly(0.784) More Information \| Similar Motifs Found | motif file (matrix) |
| 12 \* | A T C G G T A C C G T A A C G T A G T C C T G A A G T C C G A T A C G T A C T G | 1e-8 | -2.058e+01 | 4.38% | 1.59% | 359.8bp (582.2bp) | vnd/dmmpmm(Papatsenko)/fly(0.726) More Information \| Similar Motifs Found | motif file (matrix) |
| 13 \* | C G T A C A G T A G T C A C G T A G C T A G T C A G T C C G T A A G T C A T C G | 1e-8 | -1.849e+01 | 4.20% | 1.59% | 330.7bp (534.4bp) | GCR1/MA0304.1/Jaspar(0.789) More Information \| Similar Motifs Found | motif file (matrix) |
| 14 \* | C G T A A C T G A C T G C G A T A G T C A G T C A C G T C G A T | 1e-7 | -1.800e+01 | 6.44% | 3.08% | 330.9bp (466.0bp) | ttk/dmmpmm(Pollard)/fly(0.811) More Information \| Similar Motifs Found | motif file (matrix) |
| 15 \* | A C T G C G T A C G T A A C T G A G T C A G T C | 1e-7 | -1.729e+01 | 30.88% | 23.60% | 395.6bp (489.9bp) | che-1/MA0260.1/Jaspar(0.919) More Information \| Similar Motifs Found | motif file (matrix) |
| 16 \* | C T G A A G T C A T G C A T G C G T A C T A G C A T G C C A G T | 1e-6 | -1.576e+01 | 11.85% | 7.39% | 408.1bp (488.9bp) | PB0025.1\_Glis2\_1/Jaspar(0.870) More Information \| Similar Motifs Found | motif file (matrix) |
| 17 \* | A C G T C G T A A C T G C G T A A C T G G T C A G T A C A G T C A C T G C G T A | 1e-6 | -1.409e+01 | 0.93% | 0.12% | 281.2bp (235.2bp) | PRDM14(Zf)/H1-PRDM14-ChIP-Seq(GSE22767)/Homer(0.743) More Information \| Similar Motifs Found | motif file (matrix) |
| 18 \* | A G T C C G T A A C G T A G T C A G C T A G T C A C T G A C G T A G T C A T G C | 1e-5 | -1.251e+01 | 0.75% | 0.09% | 223.3bp (1054.2bp) | FXR2(KH)/Homo\_sapiens-RNCMPT00020-PBM/HughesRNA(0.746) More Information \| Similar Motifs Found | motif file (matrix) |
| 19 \* | A C G T C G T A A G T C A C G T C G T A A C G T C G T A C G T A A G T C C G T A | 1e-5 | -1.251e+01 | 0.75% | 0.09% | 279.7bp (257.7bp) | Caup/dmmpmm(Noyes\_hd)/fly(0.753) More Information \| Similar Motifs Found | motif file (matrix) |
| 20 \* | G A C T A T C G T A G C G T A C A G C T A C G T C T G A T G C A | 1e-5 | -1.242e+01 | 7.74% | 4.60% | 426.9bp (485.6bp) | TBP3(MYBrelated)/col-TBP3-DAP-Seq(GSE60143)/Homer(0.741) More Information \| Similar Motifs Found | motif file (matrix) |
| 21 \* | A G T C C G T A A G T C A C T G A C T G A G T C A C G T A G T C | 1e-4 | -1.148e+01 | 2.24% | 0.81% | 267.9bp (400.3bp) | bHLH157(bHLH)/col-bHLH157-DAP-Seq(GSE60143)/Homer(0.740) More Information \| Similar Motifs Found | motif file (matrix) |
| 22 \* | A G T C A G T C A C G T A C T G A G T C A C G T A C T G A C G T | 1e-4 | -1.141e+01 | 4.38% | 2.21% | 408.6bp (463.9bp) | Zic1::Zic2/MA1628.1/Jaspar(0.935) More Information \| Similar Motifs Found | motif file (matrix) |
| 23 \* | A C T G A G T C A C T G A G T C A C T G A G T C A G T C C G T A | 1e-4 | -1.129e+01 | 3.26% | 1.45% | 279.4bp (359.4bp) | DPL-1(E2F)/cElegans-Adult-ChIP-Seq(modEncode)/Homer(0.920) More Information \| Similar Motifs Found | motif file (matrix) |
| 24 \* | A G T C A C G T A G C T C T A G A G T C A T C G A C T G C T G A | 1e-4 | -1.087e+01 | 9.05% | 5.85% | 380.5bp (483.8bp) | CG7903(RRM)/Drosophila\_melanogaster-RNCMPT00144-PBM/HughesRNA(0.778) More Information \| Similar Motifs Found | motif file (matrix) |
| 25 \* | A C G T A G T C C G T A A T C G C G T A C G T A C G T A A G T C | 1e-4 | -1.031e+01 | 5.32% | 3.00% | 388.4bp (444.1bp) | kni/dmmpmm(Down)/fly(0.698) More Information \| Similar Motifs Found | motif file (matrix) |
| 26 \* | A G T C A C G T A G T C A C G T A C T G A C G T A G T C A G T C A C G T A G T C | 1e-2 | -5.151e+00 | 0.47% | 0.11% | 307.6bp (447.1bp) | CNOT4(RRM)/Homo\_sapiens-RNCMPT00156-PBM/HughesRNA(0.755) More Information \| Similar Motifs Found | motif file (matrix) |
| 27 \* | C G T A A C G T A G T C A G T C A C G T A C T G A C T G A C G T C G T A A C T G | 1e-1 | -3.985e+00 | 0.37% | 0.10% | 196.2bp (425.3bp) | SPDEF(ETS)/VCaP-SPDEF-ChIP-Seq(SRA014231)/Homer(0.750) More Information \| Similar Motifs Found | motif file (matrix) |
