## Supplemental Table 1 for "The Establishment of Cell-Type Specific Gene Regulation in the Sea Urchin Embryo": Hpf20_EctoArboral_homerResults.html

homer\_motifs\_wbackground/Hpf20\_EctoArboral/ - Homer de novo Motif Results


### Homer *de novo* Motif Results (homer\_motifs\_wbackground/Hpf20\_EctoArboral/)

Known Motif Enrichment Results  
Gene Ontology Enrichment Results  
If Homer is having trouble matching a motif to a known motif, try copy/pasting the matrix file into
STAMP  
More information on motif finding results: HOMER
| Description of Results
| Tips
  
Total target sequences = 1095  
Total background sequences = 8465  
\* - possible false positive  

|  |  |  |  |  |  |  |  |  |
| --- | --- | --- | --- | --- | --- | --- | --- | --- |
| Rank | Motif | P-value | log P-pvalue | % of Targets | % of Background | STD(Bg STD) | Best Match/Details | Motif File |
| 1 | C T A G G T C A A C T G C G T A C T G A C G T A A C T G A C G T | 1e-37 | -8.652e+01 | 62.83% | 43.40% | 451.2bp (490.3bp) | PTBP1(RRM)/Homo\_sapiens-RNCMPT00269-PBM/HughesRNA(0.853) More Information \| Similar Motifs Found | motif file (matrix) |
| 2 | A C G T A C T G C G T A C T G A A C G T C G T A A C T G A T C G A C T G C G T A | 1e-29 | -6.805e+01 | 6.12% | 1.03% | 302.8bp (401.7bp) | HNRNPA2B1(RRM)/Homo\_sapiens-RNCMPT00024-PBM/HughesRNA(0.745) More Information \| Similar Motifs Found | motif file (matrix) |
| 3 | A C T G C G T A A C G T A G T C C G T A A G T C A C G T A C T G A C T G C T G A | 1e-28 | -6.657e+01 | 4.75% | 0.60% | 455.3bp (487.0bp) | PB0195.1\_Zbtb3\_2/Jaspar(0.704) More Information \| Similar Motifs Found | motif file (matrix) |
| 4 | A C G T A G T C A C G T C G T A C G T A A T G C C G T A C G T A A G T C A G T C | 1e-24 | -5.738e+01 | 2.56% | 0.15% | 253.5bp (321.7bp) | YBX2(CSD)/Homo\_sapiens-RNCMPT00084-PBM/HughesRNA(0.695) More Information \| Similar Motifs Found | motif file (matrix) |
| 5 | A G T C A G T C A G T C A C T G A C G T A C T G A C T G G T C A A G T C A G T C | 1e-24 | -5.685e+01 | 1.55% | 0.03% | 214.5bp (702.1bp) | TCP16(TCP)/colamp-TCP16-DAP-Seq(GSE60143)/Homer(0.753) More Information \| Similar Motifs Found | motif file (matrix) |
| 6 | T G C A C A G T A T C G A C T G G T A C A C G T A G C T A G T C | 1e-24 | -5.580e+01 | 29.13% | 16.63% | 442.4bp (489.8bp) | che-1/MA0260.1/Jaspar(0.849) More Information \| Similar Motifs Found | motif file (matrix) |
| 7 | A C T G A C T G C G T A A C G T C G T A C G T A C G T A A C G T A G T C A G T C | 1e-23 | -5.393e+01 | 1.64% | 0.04% | 317.8bp (60.1bp) | EDS1/MA0294.1/Jaspar(0.733) More Information \| Similar Motifs Found | motif file (matrix) |
| 8 | A T G C A G C T C G T A A G C T G C A T C T A G C G A T T C G A | 1e-23 | -5.326e+01 | 50.14% | 35.32% | 434.2bp (462.2bp) | SOX2/MA0143.4/Jaspar(0.776) More Information \| Similar Motifs Found | motif file (matrix) |
| 9 | A G T C A C G T A G T C A G T C A G T C A G T C A G T C A C G T A C T G A G T C | 1e-19 | -4.531e+01 | 1.55% | 0.05% | 380.1bp (88.6bp) | CTCFL/MA1102.2/Jaspar(0.771) More Information \| Similar Motifs Found | motif file (matrix) |
| 10 | A C G T G T A C A G C T A C T G C G T A A C T G A T C G A C T G A G C T G C A T | 1e-19 | -4.457e+01 | 4.02% | 0.69% | 489.6bp (536.0bp) | THI2/THI2\_Thi-/[](Harbison)/Yeast(0.698) More Information \| Similar Motifs Found | motif file (matrix) |
| 11 | A T G C A G T C A C G T C G A T A C G T A C G T C T G A C G A T | 1e-18 | -4.329e+01 | 46.67% | 33.52% | 408.3bp (443.5bp) | KHDRBS1(KH)/Mus\_musculus-RNCMPT00062-PBM/HughesRNA(0.894) More Information \| Similar Motifs Found | motif file (matrix) |
| 12 | C G A T A C G T A G T C A C G T A C G T A C T G C G T A A G T C A G T C C G T A | 1e-17 | -4.097e+01 | 4.02% | 0.76% | 327.9bp (495.7bp) | WRKY63/MA1092.1/Jaspar(0.771) More Information \| Similar Motifs Found | motif file (matrix) |
| 13 | A C G T A G T C C T A G C G T A C T A G A C T G A G T C C G T A A C T G A G T C | 1e-17 | -4.096e+01 | 3.38% | 0.53% | 318.1bp (302.1bp) | XBP1(MacIsaac)/Yeast(0.717) More Information \| Similar Motifs Found | motif file (matrix) |
| 14 | C G A T C G T A A C T G A C G T A C T G A C T G A C G T A C G T A G T C A G T C | 1e-17 | -4.064e+01 | 2.28% | 0.21% | 374.3bp (278.2bp) | ARALYDRAFT\_496250/MA1096.1/Jaspar(0.771) More Information \| Similar Motifs Found | motif file (matrix) |
| 15 | A G T C C G T A C T G A C T A G A T C G A C T G C G T A A C G T A C T G C T A G | 1e-17 | -4.011e+01 | 4.29% | 0.89% | 449.2bp (421.7bp) | PB0098.1\_Zfp410\_1/Jaspar(0.738) More Information \| Similar Motifs Found | motif file (matrix) |
| 16 | C G T A C G T A C G T A A C T G A C T G A C G T A C G T A C T G A C G T A G T C | 1e-17 | -3.981e+01 | 1.74% | 0.11% | 360.4bp (283.6bp) | Pp\_0206(RRM)/Physcomitrella\_patens-RNCMPT00206-PBM/HughesRNA(0.791) More Information \| Similar Motifs Found | motif file (matrix) |
| 17 | C T G A A C T G A T G C A C T G G T C A A C G T G T C A T G A C C G T A C G T A | 1e-17 | -3.926e+01 | 4.38% | 0.95% | 337.1bp (476.8bp) | Dref/dmmpmm(Pollard)/fly(0.676) More Information \| Similar Motifs Found | motif file (matrix) |
| 18 | C G T A C T G A C G T A A C T G C G T A A C T G A G T C C G T A A G T C C G T A | 1e-16 | -3.840e+01 | 5.11% | 1.31% | 324.4bp (438.3bp) | NCU08034(RRM)/Neurospora\_crassa-RNCMPT00209-PBM/HughesRNA(0.746) More Information \| Similar Motifs Found | motif file (matrix) |
| 19 | C G T A A C T G A C G T A C T G A G T C A C T G A C G T A C G T A C G T A G T C | 1e-16 | -3.705e+01 | 2.10% | 0.19% | 248.2bp (231.0bp) | CST6(MacIsaac)/Yeast(0.651) More Information \| Similar Motifs Found | motif file (matrix) |
| 20 | A G T C A G T C A G T C A C G T A C G T C G T A C G T A A C G T C G T A A G T C | 1e-15 | -3.576e+01 | 2.19% | 0.23% | 359.9bp (368.7bp) | PHO2(MacIsaac)/Yeast(0.715) More Information \| Similar Motifs Found | motif file (matrix) |
| 21 | C G T A A C T G C G T A A C T G A C T G A C G T A C G T G T C A A C G T A C G T | 1e-15 | -3.462e+01 | 3.20% | 0.57% | 295.9bp (518.5bp) | NR1H4/MA1110.1/Jaspar(0.695) More Information \| Similar Motifs Found | motif file (matrix) |
| 22 | A C G T A G T C A C G T A C G T A G T C C G T A C G T A A T C G A T G C C G T A | 1e-13 | -3.054e+01 | 3.65% | 0.86% | 333.1bp (555.9bp) | AT5G25475(ABI3VP1)/col-AT5G25475-DAP-Seq(GSE60143)/Homer(0.663) More Information \| Similar Motifs Found | motif file (matrix) |
| 23 | C G A T A G T C A C G T A C G T A T C G A G T C A C G T C G T A A C G T C G T A | 1e-12 | -2.863e+01 | 3.56% | 0.87% | 377.4bp (346.1bp) | GAT3(MacIsaac)/Yeast(0.630) More Information \| Similar Motifs Found | motif file (matrix) |
| 24 \* | A G T C A C T G A C T G A C T G C G T A A C T G A C G T A G T C A C G T A C G T | 1e-10 | -2.496e+01 | 0.91% | 0.04% | 352.7bp (105.8bp) | PB0203.1\_Zfp691\_2/Jaspar(0.704) More Information \| Similar Motifs Found | motif file (matrix) |
| 25 \* | A C T G A C T G A G T C C G T A A C T G A C G T A T C G A G T C | 1e-10 | -2.357e+01 | 7.85% | 3.65% | 452.4bp (462.8bp) | PB0091.1\_Zbtb3\_1/Jaspar(0.756) More Information \| Similar Motifs Found | motif file (matrix) |
| 26 \* | A C G T A G T C C G A T C G T A C G T A C G T A C G T A A C G T A C T G C G T A | 1e-9 | -2.222e+01 | 3.38% | 0.99% | 391.2bp (534.7bp) | Lm\_0212(RRM)/Leishmania\_major-RNCMPT00212-PBM/HughesRNA(0.776) More Information \| Similar Motifs Found | motif file (matrix) |
| 27 \* | A G T C A C G T C G T A A G T C A G T C C G T A A C G T A C G T | 1e-9 | -2.198e+01 | 7.03% | 3.21% | 361.0bp (484.9bp) | schlank/MA0193.1/Jaspar(0.747) More Information \| Similar Motifs Found | motif file (matrix) |
| 28 \* | C G T A A G T C A G T C C G T A A C T G C G T A C G T A A C T G | 1e-8 | -1.967e+01 | 5.48% | 2.34% | 337.2bp (464.2bp) | OSR2/MA1646.1/Jaspar(0.785) More Information \| Similar Motifs Found | motif file (matrix) |
| 29 \* | A C G T A C T G A C T G A C T G A C G T A C T G | 1e-6 | -1.511e+01 | 36.99% | 29.88% | 441.4bp (470.9bp) | AFT2/AFT2\_H2O2Lo/10-RCS1[~AFT2](Harbison)/Yeast(0.866) More Information \| Similar Motifs Found | motif file (matrix) |
| 30 \* | A C G T A C G T A C T G A C G T A C T G A C G T C G T A C G T A | 1e-5 | -1.287e+01 | 10.59% | 6.83% | 356.5bp (464.6bp) | CEBPD/MA0836.2/Jaspar(0.836) More Information \| Similar Motifs Found | motif file (matrix) |
| 31 \* | C G T A A C G T C G T A A C G T C G T A A C G T C G T A A C G T | 1e-2 | -5.336e+00 | 7.12% | 5.26% | 438.2bp (455.4bp) | SPT15/MA0386.1/Jaspar(0.897) More Information \| Similar Motifs Found | motif file (matrix) |
