## Supplemental Table 1 for "The Establishment of Cell-Type Specific Gene Regulation in the Sea Urchin Embryo": Hpf20_EctoNearApical_homerResults.html

homer\_motifs\_wbackground/Hpf20\_EctoNearApical/ - Homer de novo Motif Results


### Homer *de novo* Motif Results (homer\_motifs\_wbackground/Hpf20\_EctoNearApical/)

Known Motif Enrichment Results  
Gene Ontology Enrichment Results  
If Homer is having trouble matching a motif to a known motif, try copy/pasting the matrix file into
STAMP  
More information on motif finding results: HOMER
| Description of Results
| Tips
  
Total target sequences = 291  
Total background sequences = 9650  
\* - possible false positive  

|  |  |  |  |  |  |  |  |  |
| --- | --- | --- | --- | --- | --- | --- | --- | --- |
| Rank | Motif | P-value | log P-pvalue | % of Targets | % of Background | STD(Bg STD) | Best Match/Details | Motif File |
| 1 | A T C G A G T C A G T C A C T G A C T G A G T C A G T C C G T A A T G C A C G T | 1e-13 | -3.063e+01 | 4.47% | 0.20% | 499.2bp (424.1bp) | RAP211(AP2EREBP)/colamp-RAP211-DAP-Seq(GSE60143)/Homer(0.757) More Information \| Similar Motifs Found | motif file (matrix) |
| 2 \* | A T C G T G C A A C G T A T G C T G A C C G A T T G A C G T A C A T G C G C T A | 1e-10 | -2.515e+01 | 12.71% | 3.44% | 411.5bp (479.2bp) | GATA15(C2C2gata)/col-GATA15-DAP-Seq(GSE60143)/Homer(0.648) More Information \| Similar Motifs Found | motif file (matrix) |
| 3 \* | A G C T G C T A C T A G C G T A G T A C A C G T C T A G G A T C A G C T A T G C | 1e-9 | -2.291e+01 | 6.19% | 0.85% | 448.8bp (386.5bp) | PB0203.1\_Zfp691\_2/Jaspar(0.640) More Information \| Similar Motifs Found | motif file (matrix) |
| 4 \* | A T C G A C T G C A T G T G A C A T G C T C A G A T C G C A T G T A C G G C T A | 1e-9 | -2.279e+01 | 25.43% | 11.81% | 455.6bp (476.2bp) | HINFP(Zf)/K562-HINFP.eGFP-ChIP-Seq(Encode)/Homer(0.754) More Information \| Similar Motifs Found | motif file (matrix) |
| 5 \* | C G T A A C T G C G A T A T C G C T A G C G A T A C G T C G T A G T C A C T A G | 1e-9 | -2.218e+01 | 9.62% | 2.27% | 493.2bp (386.4bp) | GT-1/MA1020.1/Jaspar(0.736) More Information \| Similar Motifs Found | motif file (matrix) |
| 6 \* | G T A C A G T C C T G A A T G C G C A T A G C T A C G T A G T C | 1e-9 | -2.201e+01 | 51.20% | 33.37% | 404.6bp (475.7bp) | PTBP1(RRM)/Homo\_sapiens-RNCMPT00269-PBM/HughesRNA(0.780) More Information \| Similar Motifs Found | motif file (matrix) |
| 7 \* | A G T C A G T C C G T A C A G T G C T A A G C T G T A C C T G A A T C G A T G C | 1e-8 | -2.047e+01 | 8.59% | 1.98% | 369.1bp (481.6bp) | PB0022.1\_Gata5\_1/Jaspar(0.699) More Information \| Similar Motifs Found | motif file (matrix) |
| 8 \* | A G T C A C G T A G C T A C T G G T A C G T C A G A T C C A G T A C G T G C T A | 1e-8 | -2.018e+01 | 7.56% | 1.55% | 406.2bp (408.1bp) | byn/dmmpmm(Pollard)/fly(0.822) More Information \| Similar Motifs Found | motif file (matrix) |
| 9 \* | A G T C A C G T A G T C A C G T C G T A C G T A G T C A A G T C C G T A A T G C | 1e-8 | -1.842e+01 | 3.44% | 0.27% | 518.6bp (445.2bp) | Foxf1/MA1606.1/Jaspar(0.729) More Information \| Similar Motifs Found | motif file (matrix) |
| 10 \* | C A T G C G T A A C T G C G T A A C G T A C T G A C T G A C T G A T G C A T G C | 1e-7 | -1.821e+01 | 5.50% | 0.89% | 501.8bp (532.3bp) | ARR10/MA0121.1/Jaspar(0.700) More Information \| Similar Motifs Found | motif file (matrix) |
| 11 \* | C G T A A C G T A C T G A C T G C G T A A C G T C G T A C G T A A G T C A G T C | 1e-7 | -1.725e+01 | 1.37% | 0.01% | 309.5bp (0.0bp) | At5g47390(MYBrelated)/col-At5g47390-DAP-Seq(GSE60143)/Homer(0.797) More Information \| Similar Motifs Found | motif file (matrix) |
| 12 \* | A C G T A G T C G T C A G T C A C G A T C G A T A G T C G C A T C T G A A T C G | 1e-5 | -1.358e+01 | 7.22% | 2.10% | 345.4bp (475.5bp) | Hnf6b(Homeobox)/LNCaP-Hnf6b-ChIP-Seq(GSE106305)/Homer(0.698) More Information \| Similar Motifs Found | motif file (matrix) |
| 13 \* | C T A G A C G T C G A T C G T A A G T C T C A G C T G A G A C T G T C A C T G A | 1e-5 | -1.349e+01 | 6.19% | 1.59% | 396.5bp (413.6bp) | HOXA9/MA0594.2/Jaspar(0.748) More Information \| Similar Motifs Found | motif file (matrix) |
| 14 \* | C G T A A C G T A C T G A C T G A C G T A G T C A G T C A G T C A G T C C G T A | 1e-5 | -1.290e+01 | 1.37% | 0.04% | 141.0bp (133.5bp) | TCP17(TCP)/col-TCP17-DAP-Seq(GSE60143)/Homer(0.876) More Information \| Similar Motifs Found | motif file (matrix) |
| 15 \* | G T C A A G C T T G A C A C G T G T C A A G C T C G T A A C G T C G T A G A T C | 1e-5 | -1.164e+01 | 19.93% | 11.15% | 457.0bp (421.6bp) | SeqBias: TA-repeat(0.850) More Information \| Similar Motifs Found | motif file (matrix) |
| 16 \* | C G T A C G A T A G T C C G T A A C T G A C T G A C G T A C T G A C G T A C T G | 1e-3 | -9.179e+00 | 1.72% | 0.16% | 192.0bp (358.5bp) | ZEB2(Zf)/SNU398-ZEB2-ChIP-Seq(GSE103048)/Homer(0.818) More Information \| Similar Motifs Found | motif file (matrix) |
| 17 \* | A C T G C G T A C G T A A C G T C G T A A G T C | 1e-3 | -8.624e+00 | 47.08% | 36.68% | 416.5bp (442.7bp) | KANADI1(Myb)/Seedling-KAN1-ChIP-Seq(GSE48081)/Homer(0.759) More Information \| Similar Motifs Found | motif file (matrix) |
| 18 \* | A T C G C G T A C T A G T C G A T C A G C T G A T A C G A T G C C T G A C G A T | 1e-3 | -8.066e+00 | 12.03% | 6.45% | 357.9bp (495.0bp) | Trl(Zf)/S2-GAGAfactor-ChIP-Seq(GSE40646)/Homer(0.769) More Information \| Similar Motifs Found | motif file (matrix) |
| 19 \* | G T C A C T A G A C G T A C T G A G T C A G T C A C G T C G T A C G T A A G T C | 1e-2 | -6.790e+00 | 2.06% | 0.40% | 247.9bp (355.2bp) | MYB116(MYB)/colamp-MYB116-DAP-Seq(GSE60143)/Homer(0.868) More Information \| Similar Motifs Found | motif file (matrix) |
| 20 \* | A C G T A C T G A C G T A G T C A G T C A C G T A G T C A G T C C G T A A C G T | 1e-2 | -5.369e+00 | 1.03% | 0.12% | 276.3bp (376.9bp) | SRSF1(RRM)/Homo\_sapiens-RNCMPT00106-PBM/HughesRNA(0.776) More Information \| Similar Motifs Found | motif file (matrix) |
