## Supplemental Table 1 for "The Establishment of Cell-Type Specific Gene Regulation in the Sea Urchin Embryo": Hpf20_EctoOral_homerResults.html

homer\_motifs\_wbackground/Hpf20\_EctoOral/ - Homer de novo Motif Results


### Homer *de novo* Motif Results (homer\_motifs\_wbackground/Hpf20\_EctoOral/)

Known Motif Enrichment Results  
Gene Ontology Enrichment Results  
If Homer is having trouble matching a motif to a known motif, try copy/pasting the matrix file into
STAMP  
More information on motif finding results: HOMER
| Description of Results
| Tips
  
Total target sequences = 739  
Total background sequences = 8960  
\* - possible false positive  

|  |  |  |  |  |  |  |  |  |
| --- | --- | --- | --- | --- | --- | --- | --- | --- |
| Rank | Motif | P-value | log P-pvalue | % of Targets | % of Background | STD(Bg STD) | Best Match/Details | Motif File |
| 1 | A C G T A T C G C G T A C G T A C G T A C T G A A C T G G T C A | 1e-18 | -4.222e+01 | 53.18% | 37.09% | 377.6bp (440.2bp) | At2g41835(C2H2)/col-At2g41835-DAP-Seq(GSE60143)/Homer(0.765) More Information \| Similar Motifs Found | motif file (matrix) |
| 2 | T G A C C A T G G A T C A G T C A G C T C G T A G A C T A C G T A C T G A C G T | 1e-17 | -3.984e+01 | 29.63% | 16.82% | 387.9bp (427.1bp) | SOX15/MA1152.1/Jaspar(0.881) More Information \| Similar Motifs Found | motif file (matrix) |
| 3 | C T G A C G T A G A T C C G A T A T C G C T A G A T G C C T A G A C G T A C T G | 1e-13 | -3.169e+01 | 2.03% | 0.12% | 409.5bp (327.0bp) | T11I18.17/MA0936.1/Jaspar(0.759) More Information \| Similar Motifs Found | motif file (matrix) |
| 4 | C G T A C G T A A C T G A C T G A C T G A C G T A C G T A C G T G A T C A C G T | 1e-13 | -3.105e+01 | 4.06% | 0.70% | 353.3bp (387.6bp) | TRP2(MYBrelated)/colamp-TRP2-DAP-Seq(GSE60143)/Homer(0.804) More Information \| Similar Motifs Found | motif file (matrix) |
| 5 | G T C A G A T C G T C A A G T C A G C T A G C T A C G T A G T C | 1e-12 | -2.926e+01 | 39.38% | 27.01% | 398.4bp (483.2bp) | PTBP1(RRM)/Homo\_sapiens-RNCMPT00269-PBM/HughesRNA(0.806) More Information \| Similar Motifs Found | motif file (matrix) |
| 6 \* | A G T C G A C T G C T A C G A T A G T C A T C G C G T A C G A T | 1e-11 | -2.546e+01 | 26.52% | 16.65% | 412.3bp (457.4bp) | pnr/MA0536.1/Jaspar(0.867) More Information \| Similar Motifs Found | motif file (matrix) |
| 7 \* | A C T G A T G C A G C T C A G T T C G A G T C A T G C A A C G T A C T G C A T G | 1e-9 | -2.157e+01 | 11.10% | 5.31% | 376.6bp (540.0bp) | BH2/dmmpmm(Noyes\_hd)/fly(0.686) More Information \| Similar Motifs Found | motif file (matrix) |
| 8 \* | A C G T C G T A A C T G A G T C A C T G A C T G C G T A A C G T A C G T C G T A | 1e-9 | -2.131e+01 | 0.95% | 0.03% | 256.2bp (98.5bp) | PH0126.1\_Obox6/Jaspar(0.808) More Information \| Similar Motifs Found | motif file (matrix) |
| 9 \* | A C G T A C T G A C G T A T G C A G T C A G T C A C G T C G T A A C G T A C G T | 1e-8 | -1.962e+01 | 2.30% | 0.36% | 384.2bp (298.9bp) | HNRNPA2B1(RRM)/Homo\_sapiens-RNCMPT00024-PBM/HughesRNA(0.767) More Information \| Similar Motifs Found | motif file (matrix) |
| 10 \* | C G T A C G T A A C G T A C T G C T G A A C G T A T C G A G T C A C T G C G T A | 1e-8 | -1.955e+01 | 2.57% | 0.47% | 315.2bp (358.9bp) | mab-3/MA0262.1/Jaspar(0.723) More Information \| Similar Motifs Found | motif file (matrix) |
| 11 \* | A G T C A C G T A G T C A G T C A C G T A T C G A C G T C G T A A C G T A C T G | 1e-7 | -1.707e+01 | 1.89% | 0.29% | 345.5bp (327.5bp) | ENOX1(RRM)/Homo\_sapiens-RNCMPT00149-PBM/HughesRNA(0.667) More Information \| Similar Motifs Found | motif file (matrix) |
| 12 \* | A T G C G C T A A C T G C G T A C T G A C G T A C G T A A T C G A G T C A G T C | 1e-6 | -1.610e+01 | 3.92% | 1.25% | 423.8bp (495.3bp) | PTBP1(RRM)/Homo\_sapiens-RNCMPT00268-PBM/HughesRNA(0.750) More Information \| Similar Motifs Found | motif file (matrix) |
| 13 \* | C G T A C G T A A C T G A C T G C G T A A G T C A C G T A C T G A C T G C T G A | 1e-6 | -1.556e+01 | 1.62% | 0.22% | 412.0bp (351.4bp) | ZFX(Zf)/mES-Zfx-ChIP-Seq(GSE11431)/Homer(0.668) More Information \| Similar Motifs Found | motif file (matrix) |
| 14 \* | G T A C A C G T A C G T C T G A G A T C C G A T C T G A A C T G A T G C A G T C | 1e-6 | -1.453e+01 | 1.22% | 0.12% | 342.6bp (330.5bp) | MSI1(RRM)/Homo\_sapiens-RNCMPT00041-PBM/HughesRNA(0.765) More Information \| Similar Motifs Found | motif file (matrix) |
| 15 \* | A C T G A C G T C T A G G T C A A G C T C G T A C G T A C G T A A C T G A C T G | 1e-6 | -1.419e+01 | 2.84% | 0.79% | 393.6bp (411.0bp) | KHDRBS3(KH)/Homo\_sapiens-RNCMPT00034-PBM/HughesRNA(0.736) More Information \| Similar Motifs Found | motif file (matrix) |
| 16 \* | A C T G A C G T A C G T A G T C A C G T C G T A C G T A A C T G A C T G A C G T | 1e-5 | -1.226e+01 | 0.81% | 0.06% | 362.1bp (139.0bp) | PB0194.1\_Zbtb12\_2/Jaspar(0.824) More Information \| Similar Motifs Found | motif file (matrix) |
| 17 \* | A T C G A T G C T C G A A C T G A T G C A T G C G T C A A C T G A T G C A G T C | 1e-4 | -1.025e+01 | 2.30% | 0.72% | 239.2bp (386.7bp) | Pax8(Paired,Homeobox)/Thyroid-Pax8-ChIP-Seq(GSE26938)/Homer(0.727) More Information \| Similar Motifs Found | motif file (matrix) |
| 18 \* | A C T G A C G T A C G T C T A G A G T C A G C T A C T G C G T A A G T C A G T C | 1e-4 | -1.002e+01 | 0.95% | 0.12% | 322.2bp (411.4bp) | MafA(bZIP)/Islet-MafA-ChIP-Seq(GSE30298)/Homer(0.749) More Information \| Similar Motifs Found | motif file (matrix) |
| 19 \* | A G T C A G C T A G T C A C T G A C G T A C T G A G T C C T G A A C T G A C G T | 1e-3 | -8.726e+00 | 0.68% | 0.07% | 305.2bp (377.3bp) | HBI1/MA1025.1/Jaspar(0.687) More Information \| Similar Motifs Found | motif file (matrix) |
| 20 \* | C A T G A T C G C G A T A C T G A T C G G C A T A T C G C T G A A G C T C T A G | 1e-3 | -7.424e+00 | 8.80% | 5.77% | 379.4bp (527.3bp) | Run/dmmpmm(Papatsenko)/fly(0.765) More Information \| Similar Motifs Found | motif file (matrix) |
