## Supplemental Table 1 for "The Establishment of Cell-Type Specific Gene Regulation in the Sea Urchin Embryo": Hpf20_EctoVeg_homerResults.html

homer\_motifs\_wbackground/Hpf20\_EctoVeg/ - Homer de novo Motif Results


### Homer *de novo* Motif Results (homer\_motifs\_wbackground/Hpf20\_EctoVeg/)

Known Motif Enrichment Results  
Gene Ontology Enrichment Results  
If Homer is having trouble matching a motif to a known motif, try copy/pasting the matrix file into
STAMP  
More information on motif finding results: HOMER
| Description of Results
| Tips
  
Total target sequences = 441  
Total background sequences = 9532  
\* - possible false positive  

|  |  |  |  |  |  |  |  |  |
| --- | --- | --- | --- | --- | --- | --- | --- | --- |
| Rank | Motif | P-value | log P-pvalue | % of Targets | % of Background | STD(Bg STD) | Best Match/Details | Motif File |
| 1 \* | C G T A A C T G C G T A C G T A G T C A G C T A T G A C C A G T G T A C C G A T | 1e-11 | -2.718e+01 | 10.20% | 2.99% | 353.9bp (478.1bp) | PABPC3(RRM)/Homo\_sapiens-RNCMPT00153-PBM/HughesRNA(0.702) More Information \| Similar Motifs Found | motif file (matrix) |
| 2 \* | A C T G G A T C C T G A C A G T A G C T G A C T G T C A C G T A C G T A C T A G | 1e-11 | -2.631e+01 | 43.31% | 27.94% | 430.6bp (480.7bp) | HDG7(HB)/col-HDG7-DAP-Seq(GSE60143)/Homer(0.799) More Information \| Similar Motifs Found | motif file (matrix) |
| 3 \* | A T G C A G C T C T G A A G C T C G A T T C A G C G T A T C A G C T G A C T G A | 1e-10 | -2.348e+01 | 28.12% | 15.89% | 350.0bp (463.8bp) | cebp-1/MA1444.1/Jaspar(0.659) More Information \| Similar Motifs Found | motif file (matrix) |
| 4 \* | T C G A T G A C G A T C C A T G G A T C T G C A A T G C A T G C A G T C G T C A | 1e-10 | -2.304e+01 | 11.79% | 4.32% | 364.0bp (466.3bp) | RCS1/RCS1\_H2O2Hi/35-RCS1(Harbison)/Yeast(0.810) More Information \| Similar Motifs Found | motif file (matrix) |
| 5 \* | G T C A A C G T A C G T C G T A A C G T A G T C A C G T A C G T A G T C A C T G | 1e-9 | -2.257e+01 | 1.81% | 0.06% | 490.7bp (174.5bp) | GATA2/MA0036.3/Jaspar(0.733) More Information \| Similar Motifs Found | motif file (matrix) |
| 6 \* | A T G C G A C T G A C T G C A T C G A T C T G A A G T C C G A T C T G A A G T C | 1e-9 | -2.159e+01 | 13.38% | 5.51% | 362.2bp (486.3bp) | br-Z2/dmmpmm(SeSiMCMC)/fly(0.734) More Information \| Similar Motifs Found | motif file (matrix) |
| 7 \* | A C T G C G T A A G C T A T C G A C T G A G T C A C G T G T C A C G T A A G T C | 1e-9 | -2.115e+01 | 1.81% | 0.07% | 222.4bp (485.9bp) | GT-1/MA1020.1/Jaspar(0.678) More Information \| Similar Motifs Found | motif file (matrix) |
| 8 \* | C G T A C G T A A G C T C T G A T C A G A G C T C T G A A T C G A C G T A C T G | 1e-9 | -2.086e+01 | 11.79% | 4.60% | 365.7bp (475.8bp) | br(var.2)/MA0011.1/Jaspar(0.764) More Information \| Similar Motifs Found | motif file (matrix) |
| 9 \* | T G C A T A G C C T A G A T C G G A C T G A C T T A C G G C T A T C A G A C G T | 1e-8 | -2.059e+01 | 19.05% | 9.61% | 361.5bp (464.7bp) | Unknown3/Arabidopsis-Promoters/Homer(0.721) More Information \| Similar Motifs Found | motif file (matrix) |
| 10 \* | A C T G T G A C A C G T A C G T A T C G C G T A A C G T A C T G G T A C A C T G | 1e-8 | -2.021e+01 | 1.13% | 0.02% | 152.8bp (0.0bp) | PH0134.1\_Pbx1/Jaspar(0.696) More Information \| Similar Motifs Found | motif file (matrix) |
| 11 \* | A T C G G T A C C T A G T A G C A C G T A C G T A G T C T C G A A C T G A G C T | 1e-8 | -1.886e+01 | 6.80% | 1.96% | 325.1bp (489.0bp) | PB0091.1\_Zbtb3\_1/Jaspar(0.679) More Information \| Similar Motifs Found | motif file (matrix) |
| 12 \* | G T C A G T A C C G T A G T C A A G C T G T C A A C T G T G A C T C A G T C G A | 1e-8 | -1.859e+01 | 14.97% | 7.07% | 330.3bp (411.3bp) | YOX1/Literature(Harbison)/Yeast(0.754) More Information \| Similar Motifs Found | motif file (matrix) |
| 13 \* | A G T C G A T C C G A T C T A G C T A G C G T A A G T C A G T C A G T C A G T C | 1e-8 | -1.844e+01 | 3.17% | 0.43% | 370.9bp (443.9bp) | NRG1(MacIsaac)/Yeast(0.736) More Information \| Similar Motifs Found | motif file (matrix) |
| 14 \* | A C G T A C T G A C T G C G T A A G C T C G T A C G T A A C T G C G T A A C T G | 1e-7 | -1.833e+01 | 2.49% | 0.24% | 381.0bp (392.5bp) | AT5G61620(MYBrelated)/colamp-AT5G61620-DAP-Seq(GSE60143)/Homer(0.863) More Information \| Similar Motifs Found | motif file (matrix) |
| 15 \* | A C T G A G T C A G T C A T G C C T A G G T A C A G C T A G T C | 1e-7 | -1.831e+01 | 10.66% | 4.24% | 506.3bp (435.5bp) | KLF15/MA1513.1/Jaspar(0.734) More Information \| Similar Motifs Found | motif file (matrix) |
| 16 \* | T A C G T C A G A T C G G T C A G T A C A C G T C G T A C T G A G C A T T G A C | 1e-7 | -1.795e+01 | 14.29% | 6.72% | 412.8bp (449.9bp) | PH0130.1\_Otx2/Jaspar(0.761) More Information \| Similar Motifs Found | motif file (matrix) |
| 17 \* | A G T C C G T A C T A G G C A T A C G T A G T C A G C T C T G A A C G T A C T G | 1e-7 | -1.719e+01 | 5.67% | 1.53% | 358.6bp (391.1bp) | GATA11/MA1014.1/Jaspar(0.723) More Information \| Similar Motifs Found | motif file (matrix) |
| 18 \* | C G T A G A T C C G A T C T A G C T A G T C G A G A T C C T A G C T G A G A C T | 1e-7 | -1.626e+01 | 10.43% | 4.40% | 422.3bp (487.9bp) | PB0134.1\_Hnf4a\_2/Jaspar(0.644) More Information \| Similar Motifs Found | motif file (matrix) |
| 19 \* | C G T A A G T C A C G T A G T C A G T C A G T C A G T C A G T C A C T G C G T A | 1e-5 | -1.231e+01 | 1.13% | 0.06% | 270.2bp (301.1bp) | RDS2/MA0362.1/Jaspar(0.732) More Information \| Similar Motifs Found | motif file (matrix) |
| 20 \* | C G T A A C G T A C T G A C G T A C T G A C T G A C T G A C G T A C T G A C T G | 1e-5 | -1.221e+01 | 1.36% | 0.10% | 547.5bp (558.4bp) | At1g77640(AP2EREBP)/col-At1g77640-DAP-Seq(GSE60143)/Homer(0.777) More Information \| Similar Motifs Found | motif file (matrix) |
| 21 \* | A C G T A C G T A G T C A G C T C G T A A G T C C G T A A G T C C T G A C G A T | 1e-5 | -1.195e+01 | 5.90% | 2.18% | 339.9bp (506.5bp) | RBM28(RRM)/Homo\_sapiens-RNCMPT00049-PBM/HughesRNA(0.705) More Information \| Similar Motifs Found | motif file (matrix) |
| 22 \* | C T A G A T G C C T A G A C G T C G A T C G T A A C T G A C T G A C T G C G A T | 1e-4 | -1.134e+01 | 3.63% | 1.00% | 356.5bp (611.9bp) | HNRNPA1L2(RRM)/Homo\_sapiens-RNCMPT00023-PBM/HughesRNA(0.779) More Information \| Similar Motifs Found | motif file (matrix) |
| 23 \* | G T A C A G T C A C T G A G C T A C T G A G T C C G T A C G T A C G T A A C G T | 1e-4 | -1.053e+01 | 3.17% | 0.83% | 316.8bp (501.7bp) | Pr\_0249(RRM)/Phytophthora\_ramorum-RNCMPT00249-PBM/HughesRNA(0.706) More Information \| Similar Motifs Found | motif file (matrix) |
| 24 \* | A C T G C G T A A G T C A G T C C G T A A C T G | 1e-2 | -5.349e+00 | 33.11% | 27.39% | 407.1bp (472.5bp) | ACE2/ACE2\_YPD/2-SWI5(Harbison)/Yeast(0.808) More Information \| Similar Motifs Found | motif file (matrix) |
| 25 \* | A C G T A C G T C G T A C G T A A C G T A C G T C G T A A G T C A C T G A C T G | 1e-2 | -4.731e+00 | 0.68% | 0.10% | 276.5bp (427.8bp) | Lim1/MA0194.1/Jaspar(0.834) More Information \| Similar Motifs Found | motif file (matrix) |
