## Supplemental Table 1 for "The Establishment of Cell-Type Specific Gene Regulation in the Sea Urchin Embryo": Hpf20_EndoVeg1_homerResults.html

homer\_motifs\_wbackground/Hpf20\_EndoVeg1/ - Homer de novo Motif Results


### Homer *de novo* Motif Results (homer\_motifs\_wbackground/Hpf20\_EndoVeg1/)

Known Motif Enrichment Results  
Gene Ontology Enrichment Results  
If Homer is having trouble matching a motif to a known motif, try copy/pasting the matrix file into
STAMP  
More information on motif finding results: HOMER
| Description of Results
| Tips
  
Total target sequences = 648  
Total background sequences = 9163  
\* - possible false positive  

|  |  |  |  |  |  |  |  |  |
| --- | --- | --- | --- | --- | --- | --- | --- | --- |
| Rank | Motif | P-value | log P-pvalue | % of Targets | % of Background | STD(Bg STD) | Best Match/Details | Motif File |
| 1 | G T A C A T C G A T C G G T C A T A C G G A C T A T G C C G T A G A C T T G C A | 1e-16 | -3.748e+01 | 18.52% | 8.21% | 445.9bp (468.2bp) | RTG3(MacIsaac)/Yeast(0.789) More Information \| Similar Motifs Found | motif file (matrix) |
| 2 | A T C G C G T A A C G T A G T C A C T G A C G T A C T G A C T G A C T G G T C A | 1e-15 | -3.651e+01 | 2.01% | 0.06% | 307.2bp (580.8bp) | Su(H)/dmmpmm(Papatsenko)/fly(0.713) More Information \| Similar Motifs Found | motif file (matrix) |
| 3 | T G A C C G T A A C G T G C A T G T C A C G T A T G C A T G A C G T A C C A T G | 1e-13 | -2.994e+01 | 13.27% | 5.52% | 357.1bp (446.3bp) | zen/dmmpmm(SeSiMCMC)/fly(0.892) More Information \| Similar Motifs Found | motif file (matrix) |
| 4 | A C T G T C G A C T A G G C A T A T C G C G T A T C A G C A G T T A G C T C G A | 1e-12 | -2.945e+01 | 11.27% | 4.30% | 453.0bp (428.4bp) | fos-1/MA1448.1/Jaspar(0.744) More Information \| Similar Motifs Found | motif file (matrix) |
| 5 | C G T A C T G A A G T C A G T C A C G T G T A C A G T C A G C T C G A T A C G T | 1e-12 | -2.865e+01 | 8.33% | 2.66% | 418.7bp (446.5bp) | dof4.5/MA1269.1/Jaspar(0.681) More Information \| Similar Motifs Found | motif file (matrix) |
| 6 | A T G C G A T C A G T C A G C T T A C G A C G T A C G T A C G T C T A G C T G A | 1e-12 | -2.842e+01 | 8.02% | 2.51% | 451.0bp (499.5bp) | REF6(Zf)/Arabidopsis-REF6-ChIP-Seq(GSE106942)/Homer(0.748) More Information \| Similar Motifs Found | motif file (matrix) |
| 7 | A C T G G T A C A G C T A G T C G T C A A C T G C G T A A G T C G T A C T C A G | 1e-12 | -2.777e+01 | 3.70% | 0.57% | 383.9bp (415.7bp) | Smad2(MAD)/ES-SMAD2-ChIP-Seq(GSE29422)/Homer(0.667) More Information \| Similar Motifs Found | motif file (matrix) |
| 8 | C A T G T A G C T C G A T A G C A G T C G T C A C G A T G C A T A T G C G C T A | 1e-12 | -2.771e+01 | 8.49% | 2.81% | 424.1bp (497.6bp) | PB0137.1\_Irf3\_2/Jaspar(0.756) More Information \| Similar Motifs Found | motif file (matrix) |
| 9 \* | A G T C A G T C C G A T C G A T A G C T C A T G C T A G A C G T | 1e-11 | -2.684e+01 | 45.83% | 32.67% | 406.4bp (448.0bp) | Sf3b4(RRM)/Danio\_rerio-RNCMPT00224-PBM/HughesRNA(0.750) More Information \| Similar Motifs Found | motif file (matrix) |
| 10 \* | G A C T A T G C C A G T A T G C A T C G A G T C A C T G A C G T A C G T G C A T | 1e-11 | -2.641e+01 | 17.13% | 8.61% | 405.5bp (466.6bp) | STB1/STB1\_YPD/1-SWI4,1-SWI6(Harbison)/Yeast(0.762) More Information \| Similar Motifs Found | motif file (matrix) |
| 11 \* | C T G A G C A T T C A G T G A C C G T A G A T C A C G T A G C T T C A G A T C G | 1e-11 | -2.582e+01 | 7.72% | 2.51% | 392.4bp (437.4bp) | Bapx1(Homeobox)/VertebralCol-Bapx1-ChIP-Seq(GSE36672)/Homer(0.700) More Information \| Similar Motifs Found | motif file (matrix) |
| 12 \* | C G T A C G T A C G T A C G T A A C T G C G T A A G T C G T A C A C G T A C T G | 1e-10 | -2.434e+01 | 3.24% | 0.49% | 272.5bp (431.3bp) | PABPN1(RRM)/Homo\_sapiens-RNCMPT00157-PBM/HughesRNA(0.697) More Information \| Similar Motifs Found | motif file (matrix) |
| 13 \* | A G T C G A C T A G T C G C A T C A G T G T C A C T A G G T A C T C G A C G T A | 1e-9 | -2.299e+01 | 23.30% | 13.93% | 407.0bp (442.5bp) | PB0056.1\_Rfxdc2\_1/Jaspar(0.727) More Information \| Similar Motifs Found | motif file (matrix) |
| 14 \* | A T G C A C G T A C G T A G T C A C T G A G T C A C T G C G T A C G T A A G T C | 1e-9 | -2.283e+01 | 1.85% | 0.14% | 350.3bp (228.3bp) | SWI4/MA0401.1/Jaspar(0.695) More Information \| Similar Motifs Found | motif file (matrix) |
| 15 \* | G T C A G C T A T G C A A C T G A T G C A G C T G C A T G A C T A T C G A C G T | 1e-9 | -2.277e+01 | 12.35% | 5.71% | 322.9bp (436.6bp) | SOX10/MA0442.2/Jaspar(0.773) More Information \| Similar Motifs Found | motif file (matrix) |
| 16 \* | G A T C C G A T T A C G T C G A G A T C T G A C T C G A C T A G G C A T A G T C | 1e-9 | -2.226e+01 | 6.33% | 2.00% | 445.3bp (458.0bp) | Bcl11a(Zf)/HSPC-BCL11A-ChIP-Seq(GSE104676)/Homer(0.726) More Information \| Similar Motifs Found | motif file (matrix) |
| 17 \* | C G T A A G T C A G C T A C T G A C T G A C G T A C G T A G T C A G T C A C G T | 1e-9 | -2.193e+01 | 1.85% | 0.15% | 424.1bp (412.2bp) | grh/dmmpmm(SeSiMCMC)/fly(0.731) More Information \| Similar Motifs Found | motif file (matrix) |
| 18 \* | A G T C C T G A C G A T G C T A T A G C A T G C T G A C T G C A G C A T T C G A | 1e-8 | -1.984e+01 | 11.88% | 5.77% | 467.0bp (453.7bp) | NCU02182/MA1434.1/Jaspar(0.740) More Information \| Similar Motifs Found | motif file (matrix) |
| 19 \* | A G C T A G T C A C T G A C G T G T A C C G A T C T A G C A T G A G T C G T C A | 1e-7 | -1.807e+01 | 5.09% | 1.62% | 373.1bp (512.7bp) | Hand1::Tcf3/MA0092.1/Jaspar(0.822) More Information \| Similar Motifs Found | motif file (matrix) |
| 20 \* | A C G T A C T G A G T C A G T C A C T G C G T A C G T A A C G T A C G T A G T C | 1e-7 | -1.788e+01 | 1.23% | 0.08% | 325.3bp (491.3bp) | PB0171.1\_Sox18\_2/Jaspar(0.667) More Information \| Similar Motifs Found | motif file (matrix) |
| 21 \* | T C G A A G C T T C G A A T G C A G C T A G C T T A G C C A T G T C G A A G C T | 1e-7 | -1.764e+01 | 1.08% | 0.05% | 520.1bp (418.2bp) | XBP1/Literature(Harbison)/Yeast(0.683) More Information \| Similar Motifs Found | motif file (matrix) |
| 22 \* | C G T A A C T G C G T A C G A T C T A G A C G T A C G T A G T C A G T C A C T G | 1e-6 | -1.593e+01 | 3.40% | 0.88% | 439.0bp (569.6bp) | RGT1/MA0367.1/Jaspar(0.715) More Information \| Similar Motifs Found | motif file (matrix) |
| 23 \* | A T C G G A T C G A T C A C T G G T A C A T G C A C T G T G A C | 1e-5 | -1.310e+01 | 13.73% | 8.27% | 458.8bp (461.8bp) | ERF094/MA1049.1/Jaspar(0.907) More Information \| Similar Motifs Found | motif file (matrix) |
