## Supplemental Table 1 for "The Establishment of Cell-Type Specific Gene Regulation in the Sea Urchin Embryo": Hpf20_EndoVeg2_homerResults.html

homer\_motifs\_wbackground/Hpf20\_EndoVeg2/ - Homer de novo Motif Results


### Homer *de novo* Motif Results (homer\_motifs\_wbackground/Hpf20\_EndoVeg2/)

Known Motif Enrichment Results  
Gene Ontology Enrichment Results  
If Homer is having trouble matching a motif to a known motif, try copy/pasting the matrix file into
STAMP  
More information on motif finding results: HOMER
| Description of Results
| Tips
  
Total target sequences = 1873  
Total background sequences = 7515  
\* - possible false positive  

|  |  |  |  |  |  |  |  |  |
| --- | --- | --- | --- | --- | --- | --- | --- | --- |
| Rank | Motif | P-value | log P-pvalue | % of Targets | % of Background | STD(Bg STD) | Best Match/Details | Motif File |
| 1 | A C G T C T A G A C G T A C G T A C G T C T A G G T A C G C A T G A C T C G A T | 1e-73 | -1.700e+02 | 68.82% | 47.95% | 401.7bp (455.8bp) | PHA-4(Forkhead)/cElegans-Embryos-PHA4-ChIP-Seq(modEncode)/Homer(0.938) More Information \| Similar Motifs Found | motif file (matrix) |
| 2 | A G C T C T A G G A C T A C G T T C A G C T G A A G T C C T G A G A T C G C T A | 1e-22 | -5.178e+01 | 45.22% | 34.17% | 409.7bp (479.5bp) | caup/MA0217.1/Jaspar(0.755) More Information \| Similar Motifs Found | motif file (matrix) |
| 3 | C G T A T A C G A C G T G C A T A G T C A G C T T G C A C A G T A C T G C T A G | 1e-17 | -4.055e+01 | 4.70% | 1.60% | 415.9bp (409.0bp) | PB0194.1\_Zbtb12\_2/Jaspar(0.676) More Information \| Similar Motifs Found | motif file (matrix) |
| 4 | G C T A A C G T C T A G C T G A A T G C A C G T A G T C C G T A A G C T C T G A | 1e-17 | -4.013e+01 | 17.03% | 10.45% | 470.7bp (470.9bp) | JDP2/MA0655.1/Jaspar(0.971) More Information \| Similar Motifs Found | motif file (matrix) |
| 5 | A C T G C A T G T C A G C T A G G T C A C G A T C G A T C T G A C G A T C A T G | 1e-17 | -3.995e+01 | 6.35% | 2.62% | 348.1bp (454.6bp) | PITX2/MA1547.1/Jaspar(0.817) More Information \| Similar Motifs Found | motif file (matrix) |
| 6 | T G A C C G A T A C G T A G C T T A C G C G A T C T A G C G A T G T C A A C G T | 1e-17 | -3.944e+01 | 7.85% | 3.61% | 434.7bp (409.9bp) | SOX10/MA0442.2/Jaspar(0.773) More Information \| Similar Motifs Found | motif file (matrix) |
| 7 | A T C G C T G A C T A G T A C G C G T A A G T C C T G A A G T C A T C G A G C T | 1e-14 | -3.424e+01 | 4.59% | 1.73% | 378.9bp (427.8bp) | HAC1/MA0310.1/Jaspar(0.864) More Information \| Similar Motifs Found | motif file (matrix) |
| 8 | T C A G C T A G G T A C A C T G A C T G A G T C A C G T T G C A | 1e-13 | -3.174e+01 | 25.20% | 18.14% | 391.6bp (448.4bp) | UME6/UME6\_YPD/51-UME6(Harbison)/Yeast(0.842) More Information \| Similar Motifs Found | motif file (matrix) |
| 9 \* | G A T C C A G T T G C A G A T C A G C T A T G C G T A C C G A T A G T C G A T C | 1e-10 | -2.526e+01 | 7.96% | 4.43% | 399.6bp (426.2bp) | SRSF1(RRM)/Homo\_sapiens-RNCMPT00163-PBM/HughesRNA(0.834) More Information \| Similar Motifs Found | motif file (matrix) |
| 10 \* | C G T A C G T A A C T G C G T A A G C T A G T C A C T G A C G T A C T G A C T G | 1e-10 | -2.355e+01 | 0.69% | 0.07% | 674.7bp (224.2bp) | BEAF-32B/dmmpmm(Pollard)/fly(0.665) More Information \| Similar Motifs Found | motif file (matrix) |
| 11 \* | A C G T C G A T A C G T C G T A A G T C A C T G C G T A A G T C | 1e-9 | -2.299e+01 | 14.31% | 9.68% | 427.6bp (454.1bp) | PH0065.1\_Hoxc10/Jaspar(0.956) More Information \| Similar Motifs Found | motif file (matrix) |
| 12 \* | G C A T T A C G C A T G A T C G A C G T G T A C C A G T A C T G | 1e-9 | -2.132e+01 | 9.24% | 5.69% | 410.4bp (442.2bp) | LRF(Zf)/Erythroblasts-ZBTB7A-ChIP-Seq(GSE74977)/Homer(0.753) More Information \| Similar Motifs Found | motif file (matrix) |
| 13 \* | A T G C C G T A A C T G A C T G A C G T A C T G T G C A T C A G | 1e-9 | -2.078e+01 | 14.04% | 9.68% | 462.0bp (478.3bp) | ZEB1/MA0103.3/Jaspar(0.933) More Information \| Similar Motifs Found | motif file (matrix) |
| 14 \* | C T G A A T C G C T A G G T A C A G T C C G A T C T G A C T G A | 1e-8 | -2.069e+01 | 28.83% | 22.84% | 425.5bp (446.5bp) | ZFX(Zf)/mES-Zfx-ChIP-Seq(GSE11431)/Homer(0.820) More Information \| Similar Motifs Found | motif file (matrix) |
| 15 \* | C G T A A C T G A G T C A C G T C G T A A C G T C G T A C G A T A G T C A G T C | 1e-8 | -2.020e+01 | 1.12% | 0.21% | 271.9bp (394.9bp) | MOT2/MA0379.1/Jaspar(0.718) More Information \| Similar Motifs Found | motif file (matrix) |
| 16 \* | T C G A A G C T G T A C T C G A C T G A A T G C C G T A A C G T A G T C G T C A | 1e-8 | -1.967e+01 | 18.26% | 13.44% | 497.3bp (460.0bp) | YBX1(CSD)/Homo\_sapiens-RNCMPT00116-PBM/HughesRNA(0.696) More Information \| Similar Motifs Found | motif file (matrix) |
| 17 \* | A G T C A C T G A C T G A C T G A G T C A G T C A G C T A C G T | 1e-8 | -1.919e+01 | 2.35% | 0.85% | 450.9bp (340.9bp) | SKN7(MacIsaac)/Yeast(0.746) More Information \| Similar Motifs Found | motif file (matrix) |
| 18 \* | A C T G A C T G C G T A A C G T A C T G A C G T A G T C A C G T A C G T C G T A | 1e-8 | -1.847e+01 | 0.59% | 0.05% | 310.6bp (145.0bp) | sd/dmmpmm(Bergman)/fly(0.660) More Information \| Similar Motifs Found | motif file (matrix) |
| 19 \* | A G C T A C G T A G T C A T C G A C T G T G C A G T C A A G T C | 1e-7 | -1.775e+01 | 8.44% | 5.34% | 414.9bp (445.3bp) | STB4/MA0391.1/Jaspar(0.801) More Information \| Similar Motifs Found | motif file (matrix) |
| 20 \* | G T A C A C T G A C G T A G T C A C G T A C T G A T G C A G C T | 1e-7 | -1.765e+01 | 9.77% | 6.43% | 466.1bp (480.9bp) | Smad4(MAD)/ESC-SMAD4-ChIP-Seq(GSE29422)/Homer(0.846) More Information \| Similar Motifs Found | motif file (matrix) |
| 21 \* | A C T G A C G T C G T A A C G T A G T C A G T C A G T C A C G T A C G T A C T G | 1e-6 | -1.611e+01 | 0.48% | 0.04% | 340.3bp (487.0bp) | Ebf2/MA1604.1/Jaspar(0.689) More Information \| Similar Motifs Found | motif file (matrix) |
| 22 \* | A C G T C G T A A G T C A C T G A C G T A C G T A C G T C G T A A C G T A G T C | 1e-6 | -1.611e+01 | 0.48% | 0.04% | 360.8bp (296.6bp) | KHDRBS3(KH)/Homo\_sapiens-RNCMPT00034-PBM/HughesRNA(0.801) More Information \| Similar Motifs Found | motif file (matrix) |
| 23 \* | A C T G A G T C C T G A G T A C A C G T G T A C A C T G A C T G | 1e-6 | -1.483e+01 | 7.69% | 5.00% | 409.5bp (437.4bp) | Adf1/dmmpmm(Pollard)/fly(0.709) More Information \| Similar Motifs Found | motif file (matrix) |
| 24 \* | A C G T A G T C A C G T C G T A A G T C A G T C A C G T A C T G C T G A A C G T | 1e-5 | -1.365e+01 | 0.75% | 0.16% | 397.0bp (342.1bp) | Zelda(Zf)/Embryo-zld-ChIP-Seq(GSE65441)/Homer(0.848) More Information \| Similar Motifs Found | motif file (matrix) |
| 25 \* | A C T G A G T C A G T C A C G T C G T A A C T G | 1e-5 | -1.355e+01 | 20.34% | 16.19% | 409.8bp (422.5bp) | ZNF711(Zf)/SHSY5Y-ZNF711-ChIP-Seq(GSE20673)/Homer(0.814) More Information \| Similar Motifs Found | motif file (matrix) |
| 26 \* | A G T C A C T G A G T C A C T G A T G C A C T G | 1e-4 | -1.083e+01 | 6.73% | 4.60% | 416.1bp (408.5bp) | LARK(RRM,Znf)/Drosophila\_melanogaster-RNCMPT00097-PBM/HughesRNA(0.983) More Information \| Similar Motifs Found | motif file (matrix) |
| 27 \* | C G T A A C T G A C G T C G T A A G T C A C G T C G T A A C T G | 1e-4 | -9.912e+00 | 3.04% | 1.74% | 374.5bp (532.0bp) | SPL11(SBP)/col100-SPL11-DAP-Seq(GSE60143)/Homer(0.749) More Information \| Similar Motifs Found | motif file (matrix) |
| 28 \* | C A G T T C A G T C G A C A G T C A T G C T A G C G A T A T C G | 1e-2 | -6.250e+00 | 35.40% | 32.23% | 485.6bp (447.8bp) | PB0196.1\_Zbtb7b\_2/Jaspar(0.786) More Information \| Similar Motifs Found | motif file (matrix) |
| 29 \* | A C T G A C T G C A G T A C T G A C G T C G T A C G T A A C G T A C T G A C T G | 1e-2 | -4.843e+00 | 0.85% | 0.43% | 342.1bp (543.5bp) | prd/dmmpmm(Pollard)/fly(0.767) More Information \| Similar Motifs Found | motif file (matrix) |
| 30 \* | C G T A A C G T A C T G A C G T C G T A A G T C C G T A A C G T | 1e-1 | -4.043e+00 | 18.31% | 16.47% | 435.5bp (540.0bp) | PUM(PUF)/Drosophila\_melanogaster-RNCMPT00046-PBM/HughesRNA(0.863) More Information \| Similar Motifs Found | motif file (matrix) |
