## Supplemental Table 1 for "The Establishment of Cell-Type Specific Gene Regulation in the Sea Urchin Embryo": Hpf20_GermCells_homerResults.html

homer\_motifs\_wbackground/Hpf20\_GermCells/ - Homer de novo Motif Results


### Homer *de novo* Motif Results (homer\_motifs\_wbackground/Hpf20\_GermCells/)

Known Motif Enrichment Results  
Gene Ontology Enrichment Results  
If Homer is having trouble matching a motif to a known motif, try copy/pasting the matrix file into
STAMP  
More information on motif finding results: HOMER
| Description of Results
| Tips
  
Total target sequences = 393  
Total background sequences = 9474  
\* - possible false positive  

|  |  |  |  |  |  |  |  |  |
| --- | --- | --- | --- | --- | --- | --- | --- | --- |
| Rank | Motif | P-value | log P-pvalue | % of Targets | % of Background | STD(Bg STD) | Best Match/Details | Motif File |
| 1 \* | G T A C G A C T A C T G A T C G C G T A G T A C C T A G C G T A A C T G A G T C | 1e-10 | -2.501e+01 | 3.05% | 0.19% | 268.8bp (330.4bp) | FXR2(KH)/Homo\_sapiens-RNCMPT00020-PBM/HughesRNA(0.750) More Information \| Similar Motifs Found | motif file (matrix) |
| 2 \* | A C T G A C G T G T C A A T G C T C A G T A C G A T G C G C A T A C G T A T C G | 1e-9 | -2.232e+01 | 3.31% | 0.29% | 319.5bp (475.5bp) | POPTR\_0002s00440g/MA0955.1/Jaspar(0.743) More Information \| Similar Motifs Found | motif file (matrix) |
| 3 \* | C T G A A G T C T A C G A G T C A C G T A T G C G T A C G C A T A T C G C G T A | 1e-9 | -2.115e+01 | 4.33% | 0.62% | 604.0bp (638.6bp) | MYB88(MYB)/col-MYB88-DAP-Seq(GSE60143)/Homer(0.825) More Information \| Similar Motifs Found | motif file (matrix) |
| 4 \* | A G T C C T G A A G C T T G C A G A T C G T A C T A G C A C G T A T C G A G T C | 1e-7 | -1.831e+01 | 7.89% | 2.39% | 499.2bp (489.5bp) | TEAD3/MA0808.1/Jaspar(0.665) More Information \| Similar Motifs Found | motif file (matrix) |
| 5 \* | A C T G T C G A C T A G T G A C A T C G A C G T C G T A C T A G A G T C T C A G | 1e-7 | -1.660e+01 | 3.31% | 0.47% | 380.6bp (357.1bp) | RBM28(RRM)/Homo\_sapiens-RNCMPT00049-PBM/HughesRNA(0.687) More Information \| Similar Motifs Found | motif file (matrix) |
| 6 \* | T A C G C T A G G A T C A C T G G A T C G A T C C A G T C A T G T C A G C T G A | 1e-6 | -1.486e+01 | 7.89% | 2.81% | 222.8bp (464.0bp) | AT5G23930(mTERF)/col-AT5G23930-DAP-Seq(GSE60143)/Homer(0.720) More Information \| Similar Motifs Found | motif file (matrix) |
| 7 \* | G A C T T G C A G T A C T A G C C A G T G T C A G C A T T A G C G C T A T A G C | 1e-6 | -1.399e+01 | 5.60% | 1.64% | 392.7bp (659.2bp) | MYB46/MA1040.1/Jaspar(0.790) More Information \| Similar Motifs Found | motif file (matrix) |
| 8 \* | G T C A A C T G G A C T G T A C C T G A A C T G C G T A T A G C A G C T T A C G | 1e-5 | -1.304e+01 | 5.09% | 1.46% | 274.7bp (446.2bp) | YML081W(MacIsaac)/Yeast(0.780) More Information \| Similar Motifs Found | motif file (matrix) |
| 9 \* | A G T C A G C T A G T C G C A T A G T C C G T A T A C G G T A C C T G A A C T G | 1e-5 | -1.262e+01 | 4.07% | 1.00% | 421.1bp (385.7bp) | Zic2/MA1629.1/Jaspar(0.759) More Information \| Similar Motifs Found | motif file (matrix) |
| 10 \* | C G T A A T C G A C T G A G T C A C G T T A G C A C G T A G T C C G T A G T C A | 1e-5 | -1.167e+01 | 3.31% | 0.74% | 307.4bp (550.6bp) | Nkx2-5(var.2)/MA0503.1/Jaspar(0.774) More Information \| Similar Motifs Found | motif file (matrix) |
| 11 \* | C A G T G T C A T A G C G T C A A T G C G T C A C G T A C T A G A C G T C G T A | 1e-4 | -1.129e+01 | 12.47% | 6.53% | 507.9bp (436.7bp) | HNRNPL(RRM)/Homo\_sapiens-RNCMPT00091-PBM/HughesRNA(0.732) More Information \| Similar Motifs Found | motif file (matrix) |
| 12 \* | G A T C C A G T A C T G C T A G G T C A C T G A G T C A G A T C A C G T T C A G | 1e-4 | -1.127e+01 | 27.99% | 19.12% | 429.9bp (520.4bp) | prd/dmmpmm(Down)/fly(0.733) More Information \| Similar Motifs Found | motif file (matrix) |
| 13 \* | T A G C C G T A G C A T T C G A G A C T A C T G A G T C C A T G A G C T A T C G | 1e-4 | -1.062e+01 | 21.88% | 14.18% | 362.6bp (470.0bp) | Bhlha15/MA0607.1/Jaspar(0.815) More Information \| Similar Motifs Found | motif file (matrix) |
| 14 \* | G C T A A T C G A G T C T C A G A G T C A T C G T A C G G A C T | 1e-4 | -1.062e+01 | 11.70% | 6.14% | 502.3bp (508.7bp) | RBM4(RRM,Znf)/Homo\_sapiens-RNCMPT00052-PBM/HughesRNA(0.819) More Information \| Similar Motifs Found | motif file (matrix) |
| 15 \* | A C T G A C T G A T G C A C T G A C G T G C T A G T C A A C T G A C T G A T C G | 1e-4 | -1.014e+01 | 2.80% | 0.62% | 199.4bp (603.6bp) | POPTR\_0002s00440g/MA0955.1/Jaspar(0.683) More Information \| Similar Motifs Found | motif file (matrix) |
| 16 \* | C G A T A T G C A C G T C G T A A C T G T C G A A G C T A G T C A C G T C G T A | 1e-3 | -8.880e+00 | 7.38% | 3.47% | 402.2bp (491.3bp) | GATA6/MA1396.1/Jaspar(0.912) More Information \| Similar Motifs Found | motif file (matrix) |
| 17 \* | C A T G C T G A G T C A T C G A C G A T G C T A G C A T C G A T T A C G C A G T | 1e-3 | -8.532e+00 | 20.87% | 14.18% | 503.0bp (472.5bp) | cad/dmmpmm(Down)/fly(0.794) More Information \| Similar Motifs Found | motif file (matrix) |
| 18 \* | A G C T A C G T G T A C C G T A A G T C A G T C C T G A G T A C G C T A G T A C | 1e-3 | -8.188e+00 | 4.33% | 1.62% | 546.2bp (430.1bp) | Run/dmmpmm(Papatsenko)/fly(0.780) More Information \| Similar Motifs Found | motif file (matrix) |
| 19 \* | C G T A C T G A A G T C A C T G T A C G T A G C T A G C A C T G | 1e-3 | -7.661e+00 | 11.96% | 7.19% | 396.3bp (528.6bp) | RDS1(MacIsaac)/Yeast(0.853) More Information \| Similar Motifs Found | motif file (matrix) |
| 20 \* | A T G C A C T G A G T C A C G T C G T A A C T G C G T A C G T A G T A C A C T G | 1e-2 | -6.323e+00 | 1.78% | 0.44% | 207.7bp (344.6bp) | ZBTB12/MA1649.1/Jaspar(0.716) More Information \| Similar Motifs Found | motif file (matrix) |
| 21 \* | G T C A G T C A T G C A A C T G G T C A A T C G A T G C T C G A A G T C G C T A | 1e-2 | -6.238e+00 | 10.43% | 6.48% | 314.7bp (426.3bp) | NCU08034(RRM)/Neurospora\_crassa-RNCMPT00209-PBM/HughesRNA(0.718) More Information \| Similar Motifs Found | motif file (matrix) |
| 22 \* | C G T A A T G C C G A T A T G C A G T C A C G T T G C A A C T G C A T G T G C A | 1e-1 | -4.403e+00 | 0.51% | 0.04% | 86.5bp (202.2bp) | PDR3/Literature(Harbison)/Yeast(0.694) More Information \| Similar Motifs Found | motif file (matrix) |
| 23 \* | C A G T C G T A C A T G T A C G A T C G T A G C G T A C C G T A | 1e-1 | -4.240e+00 | 15.78% | 11.96% | 499.4bp (494.0bp) | brk/dmmpmm(Bergman)/fly(0.783) More Information \| Similar Motifs Found | motif file (matrix) |
| 24 \* | A T C G C G T A C G T A A C T G T G C A G T C A A C T G C T G A C G T A A C T G | 1e-1 | -3.523e+00 | 8.14% | 5.72% | 225.2bp (492.6bp) | Unknown4/Arabidopsis-Promoters/Homer(0.858) More Information \| Similar Motifs Found | motif file (matrix) |
| 25 \* | A C G T G T A C A G T C A C G T A G T C G T A C A C G T A G T C A T G C A G T C | 1e-1 | -2.908e+00 | 2.29% | 1.23% | 342.0bp (413.0bp) | ZNF263/MA0528.2/Jaspar(0.887) More Information \| Similar Motifs Found | motif file (matrix) |
| 26 \* | A T C G C G T A G A C T A G T C A T G C C T A G G C A T C T G A | 1e-1 | -2.876e+00 | 15.01% | 12.22% | 401.7bp (535.7bp) | GATA20(C2C2gata)/colamp-GATA20-DAP-Seq(GSE60143)/Homer(0.763) More Information \| Similar Motifs Found | motif file (matrix) |
| 27 \* | A G T C A G T C C G T A A C G T A G T C A C G T A G T C A G T C A C G T A G T C | 1e0 | -1.727e+00 | 0.76% | 0.37% | 76.4bp (549.7bp) | SRSF2(RRM)/Homo\_sapiens-RNCMPT00072-PBM/HughesRNA(0.764) More Information \| Similar Motifs Found | motif file (matrix) |
