## Supplemental Table 1 for "The Establishment of Cell-Type Specific Gene Regulation in the Sea Urchin Embryo": Hpf20_OralNSM_homerResults.html

homer\_motifs\_wbackground/Hpf20\_OralNSM/ - Homer de novo Motif Results


### Homer *de novo* Motif Results (homer\_motifs\_wbackground/Hpf20\_OralNSM/)

Known Motif Enrichment Results  
Gene Ontology Enrichment Results  
If Homer is having trouble matching a motif to a known motif, try copy/pasting the matrix file into
STAMP  
More information on motif finding results: HOMER
| Description of Results
| Tips
  
Total target sequences = 1396  
Total background sequences = 7751  
\* - possible false positive  

|  |  |  |  |  |  |  |  |  |
| --- | --- | --- | --- | --- | --- | --- | --- | --- |
| Rank | Motif | P-value | log P-pvalue | % of Targets | % of Background | STD(Bg STD) | Best Match/Details | Motif File |
| 1 | G C A T C T A G T G A C C A G T A C T G A C T G A G C T C G T A A T G C A C G T | 1e-81 | -1.886e+02 | 36.39% | 15.33% | 401.5bp (450.1bp) | GCM1/MA0646.1/Jaspar(0.893) More Information \| Similar Motifs Found | motif file (matrix) |
| 2 | A G C T C T G A A G T C A G C T C G A T G T A C A G T C A C G T A T C G A C T G | 1e-60 | -1.389e+02 | 38.40% | 19.36% | 449.6bp (474.1bp) | Etv2(ETS)/ES-ER71-ChIP-Seq(GSE59402)/Homer(0.953) More Information \| Similar Motifs Found | motif file (matrix) |
| 3 | A G T C A T G C C T G A C A T G T A C G G C T A A G C T C T G A G C A T A C G T | 1e-19 | -4.533e+01 | 31.02% | 20.53% | 399.0bp (449.6bp) | AT3G10113(MYBrelated)/col-AT3G10113-DAP-Seq(GSE60143)/Homer(0.743) More Information \| Similar Motifs Found | motif file (matrix) |
| 4 | A C T G T A G C C T G A T C A G C G T A A G C T T C G A T G C A A T G C C G A T | 1e-14 | -3.305e+01 | 10.32% | 5.12% | 450.2bp (448.2bp) | GATA5/MA0766.2/Jaspar(0.785) More Information \| Similar Motifs Found | motif file (matrix) |
| 5 | A G T C A G T C A G T C A T C G A G C T G T C A A C G T T G C A A C G T T C G A | 1e-14 | -3.272e+01 | 3.30% | 0.82% | 415.0bp (630.2bp) | RBMS1(RRM)/Homo\_sapiens-RNCMPT00152-PBM/HughesRNA(0.756) More Information \| Similar Motifs Found | motif file (matrix) |
| 6 | G A T C A T G C C T A G A T C G G T C A C A G T T G A C C A T G T A C G G T A C | 1e-12 | -2.828e+01 | 8.95% | 4.48% | 494.2bp (418.8bp) | ASH1(MacIsaac)/Yeast(0.759) More Information \| Similar Motifs Found | motif file (matrix) |
| 7 | A C G T A G T C A C T G A T G C A C T G A G C T A C T G A T G C A G T C A C G T | 1e-12 | -2.791e+01 | 0.86% | 0.05% | 250.3bp (505.9bp) | h/dmmpmm(Bergman)/fly(0.821) More Information \| Similar Motifs Found | motif file (matrix) |
| 8 \* | C G T A C G T A A C G T A G C T C G T A C G T A A G T C A C T G C G T A C G T A | 1e-9 | -2.221e+01 | 1.29% | 0.19% | 439.6bp (575.4bp) | SPT2(MacIsaac)/Yeast(0.813) More Information \| Similar Motifs Found | motif file (matrix) |
| 9 \* | G A T C T G A C T C G A A G T C C T A G C G T A T G A C C G T A A T G C A G T C | 1e-9 | -2.132e+01 | 9.46% | 5.39% | 465.8bp (469.9bp) | bZIP911(1)(bZIP)/Antirrhinum majus/AthaMap(0.740) More Information \| Similar Motifs Found | motif file (matrix) |
| 10 \* | A C T G A C G T A C G T C G A T C A T G A C G T A C G T A C G T | 1e-8 | -1.986e+01 | 37.32% | 29.96% | 450.6bp (464.0bp) | HuR(RRM)/Homo\_sapiens-RNCMPT00112-PBM/HughesRNA(0.912) More Information \| Similar Motifs Found | motif file (matrix) |
| 11 \* | A C G T A C G T A C G T A C T G C G T A A G T C C G A T A C T G A C G T A C T G | 1e-8 | -1.933e+01 | 1.93% | 0.50% | 400.0bp (412.7bp) | WRKY40/MA1085.2/Jaspar(0.765) More Information \| Similar Motifs Found | motif file (matrix) |
| 12 \* | A G T C A C G T A C G T A C T G A G T C A C G T A C T G A G T C C G T A A C T G | 1e-8 | -1.911e+01 | 0.57% | 0.03% | 211.3bp (326.8bp) | ZFP57/MA1583.1/Jaspar(0.722) More Information \| Similar Motifs Found | motif file (matrix) |
| 13 \* | C G T A A C T G A C T G A C G T C G A T C G T A A C T G A C T G | 1e-7 | -1.816e+01 | 8.81% | 5.17% | 488.0bp (534.1bp) | HRB27C(RRM)/Drosophila\_melanogaster-RNCMPT00028-PBM/HughesRNA(0.812) More Information \| Similar Motifs Found | motif file (matrix) |
| 14 \* | A C T G A C T G A C T G A T C G A C T G A C T G A T C G A C T G A G C T C T A G | 1e-6 | -1.510e+01 | 14.47% | 10.17% | 484.9bp (459.9bp) | Maz(Zf)/HepG2-Maz-ChIP-Seq(GSE31477)/Homer(0.932) More Information \| Similar Motifs Found | motif file (matrix) |
| 15 \* | A C G T C T G A A G T C A G T C A C T G C G T A A C G T A C G T A C T G C G T A | 1e-6 | -1.457e+01 | 0.64% | 0.07% | 453.3bp (262.3bp) | PB0188.1\_Tcf7l2\_2/Jaspar(0.762) More Information \| Similar Motifs Found | motif file (matrix) |
| 16 \* | T A G C G T A C A G T C A G C T G T C A T A G C G A T C G C A T A G T C A G C T | 1e-6 | -1.446e+01 | 29.51% | 23.79% | 478.2bp (462.8bp) | HRB98DE(RRM)/Drosophila\_melanogaster-RNCMPT00096-PBM/HughesRNA(0.788) More Information \| Similar Motifs Found | motif file (matrix) |
| 17 \* | C G T A A C T G A C G T A C T G C G T A A G T C A C T G A C G T A C G T C G T A | 1e-6 | -1.388e+01 | 0.57% | 0.05% | 511.9bp (750.7bp) | TGA1(bZIP)/Arabidopsis thaliana/AthaMap(0.831) More Information \| Similar Motifs Found | motif file (matrix) |
| 18 \* | C G T A A C T G C G T A A G T C A G T C A C T G C G T A C G T A A C G T C G T A | 1e-5 | -1.332e+01 | 0.50% | 0.04% | 243.9bp (195.5bp) | SOX12/MA1561.1/Jaspar(0.651) More Information \| Similar Motifs Found | motif file (matrix) |
| 19 \* | C G T A A G T C A C T G A C G T A C T G C G T A A G T C C G T A A C T G C G T A | 1e-5 | -1.332e+01 | 0.50% | 0.04% | 185.5bp (181.5bp) | CBF1(MacIsaac)/Yeast(0.778) More Information \| Similar Motifs Found | motif file (matrix) |
| 20 \* | A G T C A G T C C G T A C G T A A G T C C G T A A C T G A C G T | 1e-5 | -1.318e+01 | 3.72% | 1.83% | 466.2bp (462.3bp) | RAV1(1)(AP2/EREBP)/Arabidopsis thaliana/AthaMap(0.758) More Information \| Similar Motifs Found | motif file (matrix) |
| 21 \* | C T G A C G T A C G T A C G T A G T C A A C G T A T C G C G T A | 1e-5 | -1.209e+01 | 35.32% | 29.83% | 435.6bp (457.6bp) | Lm\_0212(RRM)/Leishmania\_major-RNCMPT00212-PBM/HughesRNA(0.868) More Information \| Similar Motifs Found | motif file (matrix) |
| 22 \* | A C T G A G T C A G T C A T C G A C T G A G T C A C T G A C T G | 1e-5 | -1.151e+01 | 4.15% | 2.26% | 372.1bp (425.0bp) | ERF6/MA1006.1/Jaspar(0.790) More Information \| Similar Motifs Found | motif file (matrix) |
| 23 \* | A G T C A G T C A C G T A G T C A G T C C G T A A C G T A G T C | 1e-2 | -5.808e+00 | 3.87% | 2.60% | 386.9bp (456.6bp) | Pp\_0237(RRM)/Physcomitrella\_patens-RNCMPT00237-PBM/HughesRNA(0.832) More Information \| Similar Motifs Found | motif file (matrix) |
