## Supplemental Table 1 for "The Establishment of Cell-Type Specific Gene Regulation in the Sea Urchin Embryo": Hpf20_PMCs_homerResults.html

homer\_motifs\_wbackground/Hpf20\_PMCs/ - Homer de novo Motif Results


### Homer *de novo* Motif Results (homer\_motifs\_wbackground/Hpf20\_PMCs/)

Known Motif Enrichment Results  
Gene Ontology Enrichment Results  
If Homer is having trouble matching a motif to a known motif, try copy/pasting the matrix file into
STAMP  
More information on motif finding results: HOMER
| Description of Results
| Tips
  
Total target sequences = 2530  
Total background sequences = 9828  
\* - possible false positive  

|  |  |  |  |  |  |  |  |  |
| --- | --- | --- | --- | --- | --- | --- | --- | --- |
| Rank | Motif | P-value | log P-pvalue | % of Targets | % of Background | STD(Bg STD) | Best Match/Details | Motif File |
| 1 | T A G C C T G A T A G C T G C A A T C G T C A G G T C A G C T A T C A G G A C T | 1e-146 | -3.383e+02 | 66.44% | 40.93% | 367.9bp (463.2bp) | ERG(ETS)/VCaP-ERG-ChIP-Seq(GSE14097)/Homer(0.974) More Information \| Similar Motifs Found | motif file (matrix) |
| 2 | T G C A A C G T A C T G C G T A T A G C C A G T G T A C C G T A A G C T T G A C | 1e-43 | -9.953e+01 | 27.75% | 16.71% | 378.5bp (488.7bp) | BATF(bZIP)/Th17-BATF-ChIP-Seq(GSE39756)/Homer(0.991) More Information \| Similar Motifs Found | motif file (matrix) |
| 3 | G A C T T G C A T C G A A C G T A G C T C T G A T C A G A G T C C T G A T G A C | 1e-31 | -7.353e+01 | 40.47% | 29.41% | 352.7bp (441.5bp) | VSX1/MA0725.1/Jaspar(0.971) More Information \| Similar Motifs Found | motif file (matrix) |
| 4 | G A C T G T C A C G T A A G T C C T G A G C A T A G T C G A T C C A T G A C T G | 1e-14 | -3.373e+01 | 6.60% | 3.41% | 345.5bp (484.5bp) | SPDEF/MA0686.1/Jaspar(0.816) More Information \| Similar Motifs Found | motif file (matrix) |
| 5 | A C G T A C G T A G T C A G T C A T G C A G T C A G T C A C G T C G A T A C T G | 1e-14 | -3.240e+01 | 1.23% | 0.21% | 257.1bp (340.3bp) | MZF1(var.2)/MA0057.1/Jaspar(0.796) More Information \| Similar Motifs Found | motif file (matrix) |
| 6 \* | A C G T C G T A G T C A A C T G A C G T A G T C A C G T C G T A A C T G A G T C | 1e-10 | -2.530e+01 | 0.43% | 0.03% | 405.6bp (32.8bp) | Smad4/MA1153.1/Jaspar(0.707) More Information \| Similar Motifs Found | motif file (matrix) |
| 7 \* | A C G T G T A C T C A G T A G C A C G T C T A G T G C A G T A C T A G C A C T G | 1e-10 | -2.440e+01 | 1.58% | 0.45% | 396.1bp (474.9bp) | HVH21(HD-KNOTTED)/Hordeum vulgare/AthaMap(0.700) More Information \| Similar Motifs Found | motif file (matrix) |
| 8 \* | C G T A A G T C A C T G C G T A A C T G A C T G A G T C A G T C A G T C A C G T | 1e-9 | -2.223e+01 | 0.40% | 0.02% | 251.0bp (0.0bp) | Zac1(Zf)/Neuro2A-Plagl1-ChIP-Seq(GSE75942)/Homer(0.716) More Information \| Similar Motifs Found | motif file (matrix) |
| 9 \* | C G T A C G T A A C T G C G T A A C G T A C G T C G T A A G T C A C G T A G T C | 1e-8 | -2.061e+01 | 0.47% | 0.04% | 508.1bp (527.3bp) | MATR3(RRM)/Homo\_sapiens-RNCMPT00037-PBM/HughesRNA(0.711) More Information \| Similar Motifs Found | motif file (matrix) |
| 10 \* | G T A C G T C A A G T C A C T G A T C G A G T C A C G T C G T A A C T G A C T G | 1e-8 | -1.865e+01 | 0.55% | 0.07% | 305.9bp (528.4bp) | MYB70/MA1393.2/Jaspar(0.693) More Information \| Similar Motifs Found | motif file (matrix) |
| 11 \* | C G T A A G T C A G T C A C T G A C T G C G T A A C T G C G T A A C T G C T G A | 1e-7 | -1.621e+01 | 0.47% | 0.07% | 353.9bp (515.6bp) | OAF1/MA0348.1/Jaspar(0.715) More Information \| Similar Motifs Found | motif file (matrix) |
| 12 \* | A C G T C G T A A G T C C G T A A G T C A C G T C G T A A C G T A G T C A G T C | 1e-6 | -1.583e+01 | 0.36% | 0.04% | 198.1bp (182.7bp) | AT4G00250(GeBP)/col-AT4G00250-DAP-Seq(GSE60143)/Homer(0.677) More Information \| Similar Motifs Found | motif file (matrix) |
| 13 \* | C T A G A C G T T A G C T G C A G C A T G C T A G C T A G A T C G A C T A G C T | 1e-5 | -1.381e+01 | 23.24% | 19.41% | 394.9bp (469.8bp) | Tb\_0218(RRM)/Trypanosoma\_brucei-RNCMPT00218-PBM/HughesRNA(0.713) More Information \| Similar Motifs Found | motif file (matrix) |
| 14 \* | A G T C C G T A A G T C A C G T A G T C A G T C A G T C A C G T C G T A A G T C | 1e-5 | -1.347e+01 | 0.36% | 0.05% | 228.8bp (253.2bp) | HNRNPA2B1(RRM)/Homo\_sapiens-RNCMPT00024-PBM/HughesRNA(0.767) More Information \| Similar Motifs Found | motif file (matrix) |
| 15 \* | T A C G A G C T A T G C T C A G A T C G A C G T T A G C G C A T T A C G T A G C | 1e-5 | -1.259e+01 | 5.38% | 3.59% | 382.6bp (501.3bp) | AT1G77200(AP2EREBP)/colamp-AT1G77200-DAP-Seq(GSE60143)/Homer(0.753) More Information \| Similar Motifs Found | motif file (matrix) |
| 16 \* | C G T A A G C T C A T G C G T A A G T C A T C G G C A T A G T C G T C A A C G T | 1e-5 | -1.189e+01 | 4.39% | 2.83% | 435.9bp (436.1bp) | ATF7/MA0834.1/Jaspar(0.965) More Information \| Similar Motifs Found | motif file (matrix) |
| 17 \* | T A G C T G C A G T A C T C A G C A G T A C T G G T C A A G T C | 1e-4 | -1.057e+01 | 25.14% | 21.75% | 393.9bp (469.9bp) | CBF1(MacIsaac)/Yeast(0.923) More Information \| Similar Motifs Found | motif file (matrix) |
| 18 \* | A C T G A T G C A C T G C A G T C T G A C G T A A C T G A C T G | 1e-4 | -9.578e+00 | 2.69% | 1.63% | 424.4bp (540.5bp) | PCBP2(KH)/Homo\_sapiens-RNCMPT00044-PBM/HughesRNA(0.707) More Information \| Similar Motifs Found | motif file (matrix) |
| 19 \* | A G C T T G A C C T G A A T G C C G T A A G T C G T A C C G A T | 1e-2 | -6.427e+00 | 38.74% | 35.90% | 381.2bp (470.6bp) | TBX1/MA0805.1/Jaspar(0.991) More Information \| Similar Motifs Found | motif file (matrix) |
| 20 \* | A C T G C G T A A C G T C G T A A C G T A T C G | 1e-1 | -2.766e+00 | 37.00% | 35.52% | 403.5bp (462.9bp) | CCA(Myb)/Arabidopsis-CCA.GFP-ChIP-Seq(GSE70533)/Homer(0.871) More Information \| Similar Motifs Found | motif file (matrix) |
| 21 \* | C G A T A C T G A G T C A G T C C G T A C G T A | 1e0 | -1.454e+00 | 35.85% | 35.15% | 403.5bp (471.6bp) | NFIA/MA0670.1/Jaspar(0.969) More Information \| Similar Motifs Found | motif file (matrix) |
| 22 \* | A C T G A C T G A C G T A C G T A C T G A C G T A G T C A C T G A G T C A G T C | 1e0 | -1.290e+00 | 0.08% | 0.05% | 244.9bp (114.2bp) | ARF29/MA1692.1/Jaspar(0.733) More Information \| Similar Motifs Found | motif file (matrix) |
| 23 \* | A C G T C G T A C G T A A G T C G A C T A T G C | 1e0 | -1.746e-01 | 28.77% | 29.66% | 388.4bp (444.1bp) | MYB65/MA1177.1/Jaspar(0.817) More Information \| Similar Motifs Found | motif file (matrix) |
