## Supplemental Table 1 for "The Establishment of Cell-Type Specific Gene Regulation in the Sea Urchin Embryo": Hpf24_ArboralNSM_homerResults.html

homer\_motifs\_wbackground/Hpf24\_ArboralNSM/ - Homer de novo Motif Results


### Homer *de novo* Motif Results (homer\_motifs\_wbackground/Hpf24\_ArboralNSM/)

Known Motif Enrichment Results  
Gene Ontology Enrichment Results  
If Homer is having trouble matching a motif to a known motif, try copy/pasting the matrix file into
STAMP  
More information on motif finding results: HOMER
| Description of Results
| Tips
  
Total target sequences = 5943  
Total background sequences = 23342  
\* - possible false positive  

|  |  |  |  |  |  |  |  |  |
| --- | --- | --- | --- | --- | --- | --- | --- | --- |
| Rank | Motif | P-value | log P-pvalue | % of Targets | % of Background | STD(Bg STD) | Best Match/Details | Motif File |
| 1 | T C A G T A C G G A C T T C G A T A G C A G T C G T C A A C T G A G T C C G T A | 1e-1138 | -2.622e+03 | 45.99% | 9.64% | 353.2bp (424.6bp) | GCM1/MA0646.1/Jaspar(0.875) More Information \| Similar Motifs Found | motif file (matrix) |
| 2 | C G T A C A G T C A G T C G T A C G T A C G A T C G T A C G A T A C G T G C T A | 1e-43 | -1.009e+02 | 32.63% | 24.58% | 401.5bp (452.7bp) | Arid3b/MA0601.1/Jaspar(0.839) More Information \| Similar Motifs Found | motif file (matrix) |
| 3 | G C T A G C A T C T G A G A C T T A C G G T C A G C T A G C T A G C A T C A G T | 1e-41 | -9.458e+01 | 43.08% | 34.61% | 414.5bp (452.3bp) | br-Z1/dmmpmm(Down)/fly(0.712) More Information \| Similar Motifs Found | motif file (matrix) |
| 4 | G A T C C T A G T A C G A G T C C T A G A T G C A G T C T G A C C A T G C A T G | 1e-40 | -9.288e+01 | 23.44% | 16.66% | 424.4bp (418.0bp) | brk/dmmpmm(Bigfoot)/fly(0.752) More Information \| Similar Motifs Found | motif file (matrix) |
| 5 | G T C A G C T A C G A T G C A T A C G T C G T A A G C T A C T G | 1e-37 | -8.534e+01 | 49.79% | 41.55% | 413.8bp (443.1bp) | cad/dmmpmm(Bergman)/fly(0.859) More Information \| Similar Motifs Found | motif file (matrix) |
| 6 | C A G T A C T G T G C A C G T A C G T A G A C T C G T A A C G T | 1e-34 | -8.015e+01 | 57.97% | 49.94% | 418.0bp (442.9bp) | DIG1/DIG1\_YPD/32-STE12(Harbison)/Yeast(0.754) More Information \| Similar Motifs Found | motif file (matrix) |
| 7 | C T G A C G T A A G C T G A C T C G T A T C G A G A T C C A G T A C T G A T C G | 1e-33 | -7.806e+01 | 6.87% | 3.56% | 439.7bp (394.4bp) | PH0176.1\_Vsx1/Jaspar(0.699) More Information \| Similar Motifs Found | motif file (matrix) |
| 8 | A T G C C A T G C T A G T C G A C G A T G T C A A C G T T A C G T C G A G A T C | 1e-33 | -7.702e+01 | 18.31% | 12.76% | 437.0bp (451.3bp) | Replumless(BLH)/Arabidopsis-RPL.GFP-ChIP-Seq(GSE78727)/Homer(0.762) More Information \| Similar Motifs Found | motif file (matrix) |
| 9 | A G C T T C A G A G C T C T G A A T G C C G T A A G C T T A C G C G A T C G T A | 1e-30 | -6.912e+01 | 29.24% | 22.81% | 435.9bp (481.3bp) | PH0084.1\_Irx3\_2/Jaspar(0.864) More Information \| Similar Motifs Found | motif file (matrix) |
| 10 | A C G T A C G T A C G T C A G T A C G T A C G T A C G T A C G T C G A T C G A T | 1e-19 | -4.447e+01 | 32.73% | 27.37% | 414.6bp (457.9bp) | VRN1(ABI3VP1)/col-VRN1-DAP-Seq(GSE60143)/Homer(0.945) More Information \| Similar Motifs Found | motif file (matrix) |
| 11 | G T C A A G T C G T A C C T A G A C T G T C A G A G C T A C T G A G T C A C T G | 1e-16 | -3.854e+01 | 0.52% | 0.07% | 289.8bp (244.3bp) | ETS2/MA1484.1/Jaspar(0.707) More Information \| Similar Motifs Found | motif file (matrix) |
| 12 | A G T C A T G C C T A G T C G A A T G C T A G C G A T C T A C G | 1e-15 | -3.513e+01 | 9.51% | 6.75% | 413.6bp (383.5bp) | ERF018/MA1048.1/Jaspar(0.797) More Information \| Similar Motifs Found | motif file (matrix) |
| 13 | A G C T A C T G A G T C G T C A C G T A A C T G | 1e-13 | -3.051e+01 | 76.44% | 72.19% | 417.8bp (424.0bp) | MBNL1(Znf)/Homo\_sapiens-RNCMPT00038-PBM/HughesRNA(0.800) More Information \| Similar Motifs Found | motif file (matrix) |
| 14 \* | A T C G A T C G C G T A C G T A A C G T G T C A C G T A A G T C | 1e-11 | -2.537e+01 | 13.55% | 10.75% | 419.9bp (420.8bp) | Tb\_0218(RRM)/Trypanosoma\_brucei-RNCMPT00218-PBM/HughesRNA(0.763) More Information \| Similar Motifs Found | motif file (matrix) |
| 15 \* | A G T C A G T C A G T C A G T C A G T C A G T C A G T C A G T C | 1e-10 | -2.325e+01 | 17.70% | 14.68% | 430.1bp (430.9bp) | Maz(Zf)/HepG2-Maz-ChIP-Seq(GSE31477)/Homer(0.974) More Information \| Similar Motifs Found | motif file (matrix) |
| 16 \* | A G T C A C T G A C T G A C G T A G T C A G T C C G T A A C G T A C G T A C T G | 1e-9 | -2.117e+01 | 0.19% | 0.02% | 174.7bp (332.2bp) | SOX2/MA0143.4/Jaspar(0.691) More Information \| Similar Motifs Found | motif file (matrix) |
| 17 \* | T C A G C G T A C A T G T G C A C A T G C T G A T C A G A G T C A C T G C G T A | 1e-8 | -1.982e+01 | 16.74% | 14.04% | 433.1bp (358.8bp) | SeqBias: GA-repeat(0.873) More Information \| Similar Motifs Found | motif file (matrix) |
| 18 \* | C T G A A T G C G T C A A T G C G C A T A G T C G T C A G T A C T G C A A T G C | 1e-6 | -1.486e+01 | 11.96% | 9.98% | 356.1bp (421.3bp) | SeqBias: CA-repeat(0.823) More Information \| Similar Motifs Found | motif file (matrix) |
| 19 \* | A C T G A C T G A C G T A C T G A C G T A G T C A C T G C G T A A G T C A G T C | 1e-5 | -1.277e+01 | 0.19% | 0.03% | 316.8bp (824.8bp) | PB0117.1\_Eomes\_2/Jaspar(0.755) More Information \| Similar Motifs Found | motif file (matrix) |
| 20 \* | C G T A C G T A C G T A A G T C A G T C C G T A A C T G A C G T C G T A A G T C | 1e-4 | -9.325e+00 | 0.25% | 0.08% | 161.5bp (264.0bp) | AT1G76870(Trihelix)/col-AT1G76870-DAP-Seq(GSE60143)/Homer(0.797) More Information \| Similar Motifs Found | motif file (matrix) |
| 21 \* | A G T C A C G T A G T C A G T C C G T A A C G T A G T C A G T C C G T A A G T C | 1e0 | -1.245e+00 | 0.15% | 0.12% | 371.0bp (349.3bp) | Pp\_0237(RRM)/Physcomitrella\_patens-RNCMPT00237-PBM/HughesRNA(0.768) More Information \| Similar Motifs Found | motif file (matrix) |
| 22 \* | A G T C A T G C A C G T A G T C A G T C A C G T A G T C A G T C A C G T A G T C | 1e0 | -9.961e-01 | 0.94% | 0.90% | 488.5bp (402.5bp) | TF3A(C2H2)/col-TF3A-DAP-Seq(GSE60143)/Homer(0.827) More Information \| Similar Motifs Found | motif file (matrix) |
