## Supplemental Table 1 for "The Establishment of Cell-Type Specific Gene Regulation in the Sea Urchin Embryo": Hpf24_EctoApical_homerResults.html

homer\_motifs\_wbackground/Hpf24\_EctoApical/ - Homer de novo Motif Results


### Homer *de novo* Motif Results (homer\_motifs\_wbackground/Hpf24\_EctoApical/)

Known Motif Enrichment Results  
Gene Ontology Enrichment Results  
If Homer is having trouble matching a motif to a known motif, try copy/pasting the matrix file into
STAMP  
More information on motif finding results: HOMER
| Description of Results
| Tips
  
Total target sequences = 2702  
Total background sequences = 29426  
\* - possible false positive  

|  |  |  |  |  |  |  |  |  |
| --- | --- | --- | --- | --- | --- | --- | --- | --- |
| Rank | Motif | P-value | log P-pvalue | % of Targets | % of Background | STD(Bg STD) | Best Match/Details | Motif File |
| 1 | C G T A A G T C C G T A C G T A A C G T C T G A A C T G G T C A | 1e-58 | -1.339e+02 | 45.67% | 30.83% | 341.8bp (445.0bp) | SOX9/MA0077.1/Jaspar(0.920) More Information \| Similar Motifs Found | motif file (matrix) |
| 2 | A G C T C G T A A C G T A G C T A C T G C G T A A C G T A G C T | 1e-54 | -1.259e+02 | 38.27% | 24.64% | 340.6bp (448.5bp) | ONECUT1/MA0679.2/Jaspar(0.940) More Information \| Similar Motifs Found | motif file (matrix) |
| 3 | A G C T A C T G A G C T A G T C G T C A T G C A C G A T A G T C G C T A G T A C | 1e-31 | -7.325e+01 | 22.32% | 13.87% | 360.3bp (460.7bp) | Pknox1(Homeobox)/ES-Prep1-ChIP-Seq(GSE63282)/Homer(0.921) More Information \| Similar Motifs Found | motif file (matrix) |
| 4 | C G T A A G T C C A G T A C G T C A G T A G T C G C T A G A T C G A C T C G A T | 1e-28 | -6.522e+01 | 32.83% | 23.39% | 368.2bp (461.5bp) | PRDM1(Zf)/Hela-PRDM1-ChIP-Seq(GSE31477)/Homer(0.950) More Information \| Similar Motifs Found | motif file (matrix) |
| 5 | T A G C G A C T C G T A G C T A C A G T A C T G T C G A C T A G T A G C G A T C | 1e-22 | -5.066e+01 | 24.35% | 16.95% | 363.2bp (449.8bp) | En1(Homeobox)/SUM149-EN1-ChIP-Seq(GSE120957)/Homer(0.916) More Information \| Similar Motifs Found | motif file (matrix) |
| 6 | G A T C A C G T C G T A G T C A A C G T A G T C G A T C A G T C | 1e-19 | -4.554e+01 | 44.04% | 35.43% | 332.9bp (441.4bp) | OTX2/MA0712.2/Jaspar(0.961) More Information \| Similar Motifs Found | motif file (matrix) |
| 7 | T A G C T C A G C G T A A C T G A C T G A C T G A C T G A C T G A C G T A G T C | 1e-18 | -4.246e+01 | 11.99% | 7.19% | 364.7bp (470.6bp) | MSN4/MA0342.1/Jaspar(0.843) More Information \| Similar Motifs Found | motif file (matrix) |
| 8 | T A G C C G A T G A C T T C G A G C T A C A T G C T A G C G T A A T C G T A G C | 1e-16 | -3.904e+01 | 23.13% | 16.75% | 324.2bp (468.5bp) | RPH1/Literature(Harbison)/Yeast(0.726) More Information \| Similar Motifs Found | motif file (matrix) |
| 9 | G A C T A G T C C T G A G T C A A C G T A C G T T C A G C G T A C T A G C T G A | 1e-15 | -3.602e+01 | 6.51% | 3.34% | 372.7bp (410.0bp) | tin/dmmpmm(Pollard)/fly(0.754) More Information \| Similar Motifs Found | motif file (matrix) |
| 10 | T G A C G T A C A G T C G A C T A C T G G A T C A C T G A C T G C A T G A C T G | 1e-14 | -3.421e+01 | 1.07% | 0.15% | 450.2bp (350.0bp) | Zic2(Zf)/ESC-Zic2-ChIP-Seq(SRP197560)/Homer(0.807) More Information \| Similar Motifs Found | motif file (matrix) |
| 11 | T C A G C G T A C A T G A G T C A G T C G C A T G A T C C G A T A C G T C G T A | 1e-13 | -3.178e+01 | 50.89% | 43.60% | 372.4bp (448.1bp) | PH0115.1\_Nkx2-6/Jaspar(0.757) More Information \| Similar Motifs Found | motif file (matrix) |
| 12 | A G C T A C T G G T C A G T A C T G C A C T A G C T A G C A G T | 1e-13 | -3.150e+01 | 23.80% | 17.99% | 347.3bp (475.8bp) | unc-62/MA0918.1/Jaspar(0.935) More Information \| Similar Motifs Found | motif file (matrix) |
| 13 | A C T G C G T A C G T A A C G T A C T G A C T G A C G T A G T C A C T G A C T G | 1e-12 | -2.980e+01 | 0.30% | 0.00% | 222.3bp (0.0bp) | ERF018/MA1048.1/Jaspar(0.796) More Information \| Similar Motifs Found | motif file (matrix) |
| 14 \* | A T C G A C T G C G T A A T G C G C T A G T C A C G T A C T G A | 1e-10 | -2.348e+01 | 39.01% | 33.10% | 361.6bp (466.0bp) | IDD2(C2H2)/colamp-IDD2-DAP-Seq(GSE60143)/Homer(0.778) More Information \| Similar Motifs Found | motif file (matrix) |
| 15 \* | A C G T A G T C A G T C C G T A A C T G C G T A A T C G A G T C A C T G A C G T | 1e-9 | -2.216e+01 | 0.33% | 0.02% | 320.4bp (261.1bp) | ZBTB26/MA1579.1/Jaspar(0.707) More Information \| Similar Motifs Found | motif file (matrix) |
| 16 \* | A T G C C G T A G C A T A T C G T C G A A T G C C T G A T C G A A T G C C T A G | 1e-8 | -2.057e+01 | 26.20% | 21.36% | 358.2bp (469.8bp) | FXR1(KH)/Homo\_sapiens-RNCMPT00161-PBM/HughesRNA(0.777) More Information \| Similar Motifs Found | motif file (matrix) |
| 17 \* | A G T C C G T A C G T A A C T G C G T A C G T A A C G T C G T A A G T C C G T A | 1e-8 | -1.910e+01 | 1.18% | 0.35% | 263.0bp (312.2bp) | KANADI1(Myb)/Seedling-KAN1-ChIP-Seq(GSE48081)/Homer(0.705) More Information \| Similar Motifs Found | motif file (matrix) |
| 18 \* | A C T G C G T A C T A G A C G T C G T A C G T A A C T G A T C G A C T G A C G T | 1e-6 | -1.481e+01 | 0.78% | 0.20% | 282.5bp (553.1bp) | PRDM1/MA0508.3/Jaspar(0.671) More Information \| Similar Motifs Found | motif file (matrix) |
| 19 \* | A G T C A T C G A C T G A T G C T G C A C G T A A T G C A G C T G T A C C A T G | 1e-6 | -1.390e+01 | 0.44% | 0.07% | 332.6bp (264.6bp) | bHLH130(bHLH)/col-bHLH130-DAP-Seq(GSE60143)/Homer(0.691) More Information \| Similar Motifs Found | motif file (matrix) |
| 20 \* | C G T A A G T C A C G T A G T C A G T C C G T A C G T A A G T C A C T G C G T A | 1e-5 | -1.319e+01 | 0.26% | 0.02% | 399.0bp (413.5bp) | Pp\_0237(RRM)/Physcomitrella\_patens-RNCMPT00237-PBM/HughesRNA(0.779) More Information \| Similar Motifs Found | motif file (matrix) |
| 21 \* | C G T A A C T G A G T C A T G C C G T A A C G T A G T C A G T C | 1e-4 | -1.047e+01 | 2.18% | 1.23% | 312.8bp (484.5bp) | G3BP2(RRM)/Homo\_sapiens-RNCMPT00021-PBM/HughesRNA(0.690) More Information \| Similar Motifs Found | motif file (matrix) |
| 22 \* | A C T G A C G T A C G T A G C T A G T C C G T A A C G T A G T C | 1e-3 | -9.125e+00 | 6.74% | 5.09% | 374.1bp (496.3bp) | STE12/MA0393.1/Jaspar(0.829) More Information \| Similar Motifs Found | motif file (matrix) |
| 23 \* | A G T C C G T A A G C T C G T A A C G T C G T A A C G T C G T A A C G T G T C A | 1e-2 | -5.613e+00 | 2.70% | 1.94% | 352.3bp (489.8bp) | Cf2/dmmpmm(Bergman)/fly(0.847) More Information \| Similar Motifs Found | motif file (matrix) |
