## Supplemental Table 1 for "The Establishment of Cell-Type Specific Gene Regulation in the Sea Urchin Embryo": Hpf24_EctoArboral_homerResults.html

homer\_motifs\_wbackground/Hpf24\_EctoArboral/ - Homer de novo Motif Results


### Homer *de novo* Motif Results (homer\_motifs\_wbackground/Hpf24\_EctoArboral/)

Known Motif Enrichment Results  
Gene Ontology Enrichment Results  
If Homer is having trouble matching a motif to a known motif, try copy/pasting the matrix file into
STAMP  
More information on motif finding results: HOMER
| Description of Results
| Tips
  
Total target sequences = 3294  
Total background sequences = 31300  
\* - possible false positive  

|  |  |  |  |  |  |  |  |  |
| --- | --- | --- | --- | --- | --- | --- | --- | --- |
| Rank | Motif | P-value | log P-pvalue | % of Targets | % of Background | STD(Bg STD) | Best Match/Details | Motif File |
| 1 | G C T A A G T C C G T A C G T A C G A T C T G A T A C G T C A G | 1e-101 | -2.328e+02 | 42.68% | 25.50% | 376.1bp (407.4bp) | SOX15/MA1152.1/Jaspar(0.889) More Information \| Similar Motifs Found | motif file (matrix) |
| 2 | C G T A C G T A A C G T A G T C A C G T A C T G C G T A A C T G A C T G A C T G | 1e-82 | -1.890e+02 | 2.52% | 0.10% | 347.6bp (348.5bp) | MET31/Literature(Harbison)/Yeast(0.668) More Information \| Similar Motifs Found | motif file (matrix) |
| 3 | A C T G A G T C A C G T A C T G A G T C A G T C A C G T A G T C A C T G C G T A | 1e-75 | -1.745e+02 | 2.13% | 0.07% | 323.8bp (392.0bp) | XBP1(MacIsaac)/Yeast(0.732) More Information \| Similar Motifs Found | motif file (matrix) |
| 4 | A G T C A G T C A C T G A C T G A C T G C G T A A C T G A C G T A G T C A C G T | 1e-67 | -1.550e+02 | 1.22% | 0.01% | 463.0bp (85.4bp) | PB0203.1\_Zfp691\_2/Jaspar(0.710) More Information \| Similar Motifs Found | motif file (matrix) |
| 5 | A G C T G A T C T G A C G T C A A C T G A C G T T C A G C T G A C G A T T A C G | 1e-63 | -1.467e+02 | 3.95% | 0.56% | 413.9bp (364.2bp) | tin/dmmpmm(Bigfoot)/fly(0.757) More Information \| Similar Motifs Found | motif file (matrix) |
| 6 | T G A C A T G C G C T A A G C T A C G T A T G C C T G A C G T A | 1e-63 | -1.466e+02 | 58.57% | 43.86% | 413.9bp (428.6bp) | NKX2-2/MA1645.1/Jaspar(0.799) More Information \| Similar Motifs Found | motif file (matrix) |
| 7 | C G T A A C T G A C G T A T C G A G T C A C T G A C G T A C G T A C G T A G T C | 1e-61 | -1.409e+02 | 1.82% | 0.07% | 312.3bp (214.8bp) | STE12/MA0393.1/Jaspar(0.652) More Information \| Similar Motifs Found | motif file (matrix) |
| 8 | A C G T A G T C A C G T C G T A T C A G A G T C C G T A C G T A A G T C A G T C | 1e-57 | -1.329e+02 | 2.61% | 0.24% | 416.3bp (419.4bp) | RFX1/MA0365.1/Jaspar(0.787) More Information \| Similar Motifs Found | motif file (matrix) |
| 9 | A C G T A C G T A G T C A C G T A C G T A C T G C G T A A G T C A G T C C G T A | 1e-56 | -1.294e+02 | 2.92% | 0.33% | 369.9bp (430.5bp) | WRKY63/MA1092.1/Jaspar(0.762) More Information \| Similar Motifs Found | motif file (matrix) |
| 10 | C G T A A G T C A C G T A C G T A C G T A G T C G C A T G A T C | 1e-52 | -1.202e+02 | 51.40% | 38.25% | 440.0bp (443.8bp) | PTBP1(RRM)/Homo\_sapiens-RNCMPT00269-PBM/HughesRNA(0.917) More Information \| Similar Motifs Found | motif file (matrix) |
| 11 | A C T G A C T G A C T G C G T A A C G T A C T G C G T A A C G T A C T G A G T C | 1e-45 | -1.057e+02 | 1.82% | 0.13% | 289.5bp (374.2bp) | AGL42/MA1201.1/Jaspar(0.808) More Information \| Similar Motifs Found | motif file (matrix) |
| 12 | A C T G A C T G A C T G C G T A A C G T A C T G A C T G G T C A A C G T A C G T | 1e-44 | -1.022e+02 | 2.22% | 0.24% | 277.3bp (413.5bp) | PB0098.1\_Zfp410\_1/Jaspar(0.734) More Information \| Similar Motifs Found | motif file (matrix) |
| 13 | A C T G A C T G A C T G A T G C C T G A A C T G G C T A A C T G A G T C G T C A | 1e-43 | -1.003e+02 | 4.40% | 1.07% | 375.9bp (458.2bp) | Sp5(Zf)/mES-Sp5.Flag-ChIP-Seq(GSE72989)/Homer(0.743) More Information \| Similar Motifs Found | motif file (matrix) |
| 14 | A C T G G T C A A C G T C G A T G T C A C T G A G C A T A G T C G T A C A G C T | 1e-41 | -9.576e+01 | 3.31% | 0.64% | 325.7bp (432.3bp) | PH0138.1\_Pitx2/Jaspar(0.810) More Information \| Similar Motifs Found | motif file (matrix) |
| 15 | C T A G T C A G C T A G T G A C A G T C A G C T A G C T A G C T | 1e-39 | -9.133e+01 | 18.96% | 11.07% | 385.4bp (438.2bp) | DOF5.7/MA0984.1/Jaspar(0.681) More Information \| Similar Motifs Found | motif file (matrix) |
| 16 | A C T G A C G T C G T A A C T G A C G T A C G T C G A T C G T A C G T A A C T G | 1e-38 | -8.951e+01 | 2.13% | 0.26% | 307.5bp (375.1bp) | Rbm42(RRM)/Xenopus\_tropicalis-RNCMPT00282-PBM/HughesRNA(0.749) More Information \| Similar Motifs Found | motif file (matrix) |
| 17 | A G T C A C G T C G T A C G T A C G T A C G T A A C G T A C T G C G T A C G T A | 1e-35 | -8.283e+01 | 2.40% | 0.38% | 441.5bp (365.6bp) | Lm\_0212(RRM)/Leishmania\_major-RNCMPT00212-PBM/HughesRNA(0.766) More Information \| Similar Motifs Found | motif file (matrix) |
| 18 | G T A C T A G C C T G A G T A C T C A G A G T C T G A C G A T C | 1e-34 | -8.043e+01 | 10.81% | 5.32% | 417.2bp (412.4bp) | KLF6/MA1517.1/Jaspar(0.953) More Information \| Similar Motifs Found | motif file (matrix) |
| 19 | C G T A C G T A C G T A A C T G C G T A A C T G A G T C C G T A A G T C C G T A | 1e-32 | -7.576e+01 | 2.34% | 0.40% | 293.9bp (354.0bp) | NCU08034(RRM)/Neurospora\_crassa-RNCMPT00209-PBM/HughesRNA(0.722) More Information \| Similar Motifs Found | motif file (matrix) |
| 20 | C G T A A C T G A C G T A C T G A C T G A C G T A C G T A T G C | 1e-30 | -6.988e+01 | 19.20% | 12.15% | 414.4bp (415.1bp) | OPI1/MA0349.1/Jaspar(0.761) More Information \| Similar Motifs Found | motif file (matrix) |
| 21 | A C G T C G T A C G T A A C G T C G T A C G T A C G T A C G T A A C T G A C T G | 1e-28 | -6.627e+01 | 2.28% | 0.44% | 326.1bp (402.6bp) | cad/dmmpmm(Noyes\_hd)/fly(0.808) More Information \| Similar Motifs Found | motif file (matrix) |
| 22 | C G T A C G T A A C T G A G T C A C G T A C G T A G T C A G C T | 1e-28 | -6.574e+01 | 12.73% | 7.19% | 425.3bp (413.1bp) | SFL1(MacIsaac)/Yeast(0.880) More Information \| Similar Motifs Found | motif file (matrix) |
| 23 | T C G A T C A G A G C T A T G C A T C G T G C A G A T C A C T G | 1e-18 | -4.200e+01 | 7.59% | 4.17% | 359.0bp (416.4bp) | WRKY42(WRKY)/colamp-WRKY42-DAP-Seq(GSE60143)/Homer(0.785) More Information \| Similar Motifs Found | motif file (matrix) |
| 24 | A C T G A G T C A G T C A G T C C G T A A G T C | 1e-17 | -4.038e+01 | 20.72% | 15.07% | 364.5bp (445.1bp) | TCP23/MA1066.1/Jaspar(0.856) More Information \| Similar Motifs Found | motif file (matrix) |
| 25 | A C T G A G T C A G T C A G T C A G T C C G T A A C T G A C T G | 1e-17 | -3.926e+01 | 2.31% | 0.72% | 379.7bp (544.8bp) | TFAP2C(var.2)/MA0814.2/Jaspar(0.812) More Information \| Similar Motifs Found | motif file (matrix) |
| 26 | C G T A A C G T A C G T A C G T C G T A C G T A C G T A A C G T | 1e-13 | -3.036e+01 | 6.93% | 4.11% | 402.2bp (449.9bp) | AT2G20110(CPP)/colamp-AT2G20110-DAP-Seq(GSE60143)/Homer(0.917) More Information \| Similar Motifs Found | motif file (matrix) |
