## Supplemental Table 1 for "The Establishment of Cell-Type Specific Gene Regulation in the Sea Urchin Embryo": Hpf24_EctoNearApical_homerResults.html

homer\_motifs\_wbackground/Hpf24\_EctoNearApical/ - Homer de novo Motif Results


### Homer *de novo* Motif Results (homer\_motifs\_wbackground/Hpf24\_EctoNearApical/)

Known Motif Enrichment Results  
Gene Ontology Enrichment Results  
If Homer is having trouble matching a motif to a known motif, try copy/pasting the matrix file into
STAMP  
More information on motif finding results: HOMER
| Description of Results
| Tips
  
Total target sequences = 1983  
Total background sequences = 30528  
\* - possible false positive  

|  |  |  |  |  |  |  |  |  |
| --- | --- | --- | --- | --- | --- | --- | --- | --- |
| Rank | Motif | P-value | log P-pvalue | % of Targets | % of Background | STD(Bg STD) | Best Match/Details | Motif File |
| 1 | T G C A T A G C G A C T G C T A C A G T A G T C T A C G T G C A A C G T G A C T | 1e-53 | -1.231e+02 | 29.75% | 15.86% | 324.4bp (446.2bp) | HNF6(Homeobox)/Liver-Hnf6-ChIP-Seq(ERP000394)/Homer(0.980) More Information \| Similar Motifs Found | motif file (matrix) |
| 2 | G A T C T G C A A G T C A G C T A G C T A C G T A G T C G C T A G A T C G A C T | 1e-32 | -7.413e+01 | 43.82% | 31.07% | 374.0bp (471.2bp) | PRDM1(Zf)/Hela-PRDM1-ChIP-Seq(GSE31477)/Homer(0.887) More Information \| Similar Motifs Found | motif file (matrix) |
| 3 | G A T C A C G T C G T A G T C A A C G T A G T C G T A C T A G C | 1e-26 | -6.210e+01 | 46.34% | 34.50% | 319.6bp (433.5bp) | OTX2/MA0712.2/Jaspar(0.967) More Information \| Similar Motifs Found | motif file (matrix) |
| 4 | C T G A A T C G C G T A A T G C C G T A C G T A C G A T C T G A A C T G T A C G | 1e-23 | -5.501e+01 | 49.97% | 38.68% | 349.9bp (437.4bp) | Sox21(HMG)/ESC-SOX21-ChIP-Seq(GSE110505)/Homer(0.869) More Information \| Similar Motifs Found | motif file (matrix) |
| 5 | C A T G G T A C G C A T G A C T G C T A G C T A T C A G C G T A T G C A A C T G | 1e-15 | -3.656e+01 | 42.06% | 33.21% | 349.5bp (458.7bp) | PEND/MA0127.1/Jaspar(0.705) More Information \| Similar Motifs Found | motif file (matrix) |
| 6 | C G T A C T G A C G T A A C T G A C T G T G A C C G A T A C G T | 1e-13 | -3.057e+01 | 57.29% | 48.93% | 380.6bp (450.1bp) | Kr/dmmpmm(Papatsenko)/fly(0.848) More Information \| Similar Motifs Found | motif file (matrix) |
| 7 | A C T G A G C T C G T A C G T A A C G T C G T A A C T G A C T G A T C G A C T G | 1e-12 | -2.944e+01 | 1.56% | 0.29% | 376.5bp (387.7bp) | YER130C/MA0423.1/Jaspar(0.801) More Information \| Similar Motifs Found | motif file (matrix) |
| 8 | A G T C A G T C G T C A G C A T G T A C C T G A C A T G A C T G T C G A T A C G | 1e-12 | -2.813e+01 | 3.78% | 1.47% | 336.4bp (551.3bp) | HAP3(CCAATHAP3)/col-HAP3-DAP-Seq(GSE60143)/Homer(0.660) More Information \| Similar Motifs Found | motif file (matrix) |
| 9 \* | G C A T A T C G C G T A A G C T A T C G G A T C G A C T G A T C | 1e-11 | -2.726e+01 | 22.59% | 16.50% | 408.8bp (468.1bp) | YBX1(CSD)/Homo\_sapiens-RNCMPT00116-PBM/HughesRNA(0.782) More Information \| Similar Motifs Found | motif file (matrix) |
| 10 \* | G A C T T G C A C T G A C A G T A T C G T C G A G C A T A T C G T G C A T A C G | 1e-10 | -2.379e+01 | 16.84% | 11.87% | 370.2bp (461.6bp) | AGL42/MA1201.1/Jaspar(0.780) More Information \| Similar Motifs Found | motif file (matrix) |
| 11 \* | C G A T A C G T C T G A T G C A A T C G A T G C C T G A C T G A A G T C A T G C | 1e-10 | -2.345e+01 | 1.21% | 0.23% | 564.0bp (523.4bp) | CEBPB/MA0466.2/Jaspar(0.678) More Information \| Similar Motifs Found | motif file (matrix) |
| 12 \* | T C G A T C A G T A G C A C G T G A T C G A C T G A C T C G A T | 1e-10 | -2.322e+01 | 34.19% | 27.61% | 388.5bp (463.3bp) | DOF5.7/MA0984.1/Jaspar(0.750) More Information \| Similar Motifs Found | motif file (matrix) |
| 13 \* | C G T A C G T A G T A C A C G T A C T G A C T G A G T C T G C A T C A G G T C A | 1e-10 | -2.307e+01 | 2.77% | 1.02% | 294.8bp (463.6bp) | PB0195.1\_Zbtb3\_2/Jaspar(0.748) More Information \| Similar Motifs Found | motif file (matrix) |
| 14 \* | A G T C C T G A A C G T A C T G A C T G A G T C A G T C C G T A A C T G A C G T | 1e-8 | -2.020e+01 | 0.76% | 0.10% | 195.5bp (272.4bp) | SKN7/MA0381.1/Jaspar(0.682) More Information \| Similar Motifs Found | motif file (matrix) |
| 15 \* | A G T C A C T G A C T G A C G T A C T G A G T C A C T G C G T A C G T A A G T C | 1e-7 | -1.819e+01 | 0.35% | 0.02% | 210.0bp (198.6bp) | ABI4(2)(AP2/EREBP)/Zea mays/AthaMap(0.717) More Information \| Similar Motifs Found | motif file (matrix) |
| 16 \* | A C T G C G T A C G T A A C T G A G T C A C T G A C G T A C G T A G T C C G T A | 1e-5 | -1.209e+01 | 0.50% | 0.08% | 263.1bp (254.3bp) | Lm\_0254(RRM)/Leishmania\_major-RNCMPT00254-PBM/HughesRNA(0.734) More Information \| Similar Motifs Found | motif file (matrix) |
| 17 \* | A C G T A G T C C G T A A C T G A C G T C T A G C G T A A C G T A C T G A C T G | 1e-3 | -7.510e+00 | 0.91% | 0.37% | 664.9bp (516.2bp) | Unknown2/Drosophila-Promoters/Homer(0.752) More Information \| Similar Motifs Found | motif file (matrix) |
