## Supplemental Table 1 for "The Establishment of Cell-Type Specific Gene Regulation in the Sea Urchin Embryo": Hpf24_EctoOral_homerResults.html

homer\_motifs\_wbackground/Hpf24\_EctoOral/ - Homer de novo Motif Results


### Homer *de novo* Motif Results (homer\_motifs\_wbackground/Hpf24\_EctoOral/)

Known Motif Enrichment Results  
Gene Ontology Enrichment Results  
If Homer is having trouble matching a motif to a known motif, try copy/pasting the matrix file into
STAMP  
More information on motif finding results: HOMER
| Description of Results
| Tips
  
Total target sequences = 2352  
Total background sequences = 29382  
\* - possible false positive  

|  |  |  |  |  |  |  |  |  |
| --- | --- | --- | --- | --- | --- | --- | --- | --- |
| Rank | Motif | P-value | log P-pvalue | % of Targets | % of Background | STD(Bg STD) | Best Match/Details | Motif File |
| 1 | A T G C A G C T G C T A G A C T A G C T A T C G A G C T A T G C T G A C T A C G | 1e-56 | -1.295e+02 | 45.28% | 29.73% | 338.0bp (447.3bp) | Sox21(HMG)/ESC-SOX21-ChIP-Seq(GSE110505)/Homer(0.944) More Information \| Similar Motifs Found | motif file (matrix) |
| 2 | A G C T A G T C A G T C C G T A A C G T A C G T A G T C C G T A | 1e-37 | -8.621e+01 | 52.98% | 39.81% | 337.5bp (447.5bp) | PB0028.1\_Hbp1\_1/Jaspar(0.821) More Information \| Similar Motifs Found | motif file (matrix) |
| 3 | G A C T C G T A A C G T A G C T A C T G C G T A A C G T A G C T | 1e-32 | -7.561e+01 | 42.47% | 30.70% | 339.9bp (455.0bp) | ONECUT1/MA0679.2/Jaspar(0.945) More Information \| Similar Motifs Found | motif file (matrix) |
| 4 | C T G A T C G A G T C A G T A C G A T C G A T C A G T C G A C T G A C T G C A T | 1e-22 | -5.156e+01 | 35.29% | 26.08% | 378.9bp (442.5bp) | Kr/dmmpmm(SeSiMCMC)/fly(0.814) More Information \| Similar Motifs Found | motif file (matrix) |
| 5 | G A C T G C T A G T C A A T G C G T C A G T A C A G C T A G C T A G C T A G T C | 1e-20 | -4.710e+01 | 26.15% | 18.30% | 412.3bp (463.6bp) | Kr/dmmpmm(Down)/fly(0.774) More Information \| Similar Motifs Found | motif file (matrix) |
| 6 | G A T C G A C T T G C A G T C A C G A T T G A C T G A C A G T C | 1e-18 | -4.288e+01 | 55.06% | 45.86% | 379.4bp (456.5bp) | OTX2/MA0712.2/Jaspar(0.928) More Information \| Similar Motifs Found | motif file (matrix) |
| 7 | C T A G T A C G A T G C C G A T C G A T G A C T A G T C C G T A G T A C C G A T | 1e-18 | -4.216e+01 | 22.07% | 15.16% | 373.0bp (443.7bp) | PRDM1(Zf)/Hela-PRDM1-ChIP-Seq(GSE31477)/Homer(0.837) More Information \| Similar Motifs Found | motif file (matrix) |
| 8 | A G T C A G C T G C A T C G T A C T G A A T C G G T C A C G T A | 1e-17 | -4.121e+01 | 27.64% | 20.11% | 355.2bp (479.3bp) | bap/dmmpmm(Noyes\_hd)/fly(0.734) More Information \| Similar Motifs Found | motif file (matrix) |
| 9 | C T A G T A C G C T G A C T A G G C T A C G T A T C A G G T C A A T C G A G T C | 1e-15 | -3.657e+01 | 2.51% | 0.69% | 367.8bp (503.8bp) | TOD6?/SacCer-Promoters/Homer(0.741) More Information \| Similar Motifs Found | motif file (matrix) |
| 10 | T G A C C G T A A G C T T A G C A C T G T A C G T A G C C G A T A C G T C T G A | 1e-12 | -2.915e+01 | 4.17% | 1.82% | 342.5bp (413.9bp) | bcd/dmmpmm(Bigfoot)/fly(0.704) More Information \| Similar Motifs Found | motif file (matrix) |
| 11 | T G A C C A T G G T A C C A G T G A C T C G T A | 1e-12 | -2.891e+01 | 57.36% | 49.91% | 354.5bp (473.6bp) | bcd-1/dmmpmm(Noyes)/fly(0.821) More Information \| Similar Motifs Found | motif file (matrix) |
| 12 \* | C G T A A T C G A G T C A C T G A C G T A C G T A G T C A T C G | 1e-10 | -2.311e+01 | 4.29% | 2.13% | 288.2bp (450.8bp) | Lm\_0254(RRM)/Leishmania\_major-RNCMPT00254-PBM/HughesRNA(0.816) More Information \| Similar Motifs Found | motif file (matrix) |
| 13 \* | C G T A T A C G C T A G G A C T A G T C A C G T A T C G C T G A A C T G A T G C | 1e-9 | -2.092e+01 | 2.17% | 0.81% | 248.2bp (486.0bp) | POL013.1\_MED-1/Jaspar(0.679) More Information \| Similar Motifs Found | motif file (matrix) |
| 14 \* | G T A C A G T C A C T G C G T A G T C A G A C T A C G T A G T C G T C A A G T C | 1e-8 | -2.011e+01 | 3.10% | 1.43% | 295.6bp (417.3bp) | Tb\_0251(RRM)/Trypanosoma\_brucei-RNCMPT00251-PBM/HughesRNA(0.681) More Information \| Similar Motifs Found | motif file (matrix) |
| 15 \* | A C T G A G C T A C G T G T A C A G T C G A T C C G T A A C G T A G C T T A C G | 1e-8 | -1.909e+01 | 4.17% | 2.21% | 454.7bp (502.6bp) | RBPJ/MA1116.1/Jaspar(0.737) More Information \| Similar Motifs Found | motif file (matrix) |
| 16 \* | C G T A C G A T A G T C A G T C A C G T A G T C | 1e-7 | -1.840e+01 | 53.06% | 47.26% | 381.3bp (490.8bp) | RUNX3/MA0684.2/Jaspar(0.801) More Information \| Similar Motifs Found | motif file (matrix) |
| 17 \* | A G T C A C G T A C G T C G T A A G T C A C G T C G T A A C T G C T A G A G T C | 1e-7 | -1.712e+01 | 0.81% | 0.17% | 377.4bp (505.1bp) | MSI1(RRM)/Homo\_sapiens-RNCMPT00041-PBM/HughesRNA(0.756) More Information \| Similar Motifs Found | motif file (matrix) |
| 18 \* | A C G T A C T G A G C T A C G T A G T C A C T G A G T C C T A G A C T G A T C G | 1e-7 | -1.677e+01 | 0.72% | 0.14% | 491.8bp (493.3bp) | ZBTB14/MA1650.1/Jaspar(0.736) More Information \| Similar Motifs Found | motif file (matrix) |
| 19 \* | G C T A A T C G A C T G A C G T T C G A C G A T A G T C C T G A | 1e-6 | -1.477e+01 | 10.54% | 7.68% | 404.2bp (465.2bp) | Six4/dmmpmm(Noyes\_hd)/fly(0.840) More Information \| Similar Motifs Found | motif file (matrix) |
| 20 \* | C T A G T C A G C G T A A G T C A G T C C A G T A C G T A C G T | 1e-6 | -1.427e+01 | 59.18% | 54.20% | 380.9bp (469.6bp) | PB0137.1\_Irf3\_2/Jaspar(0.786) More Information \| Similar Motifs Found | motif file (matrix) |
| 21 \* | A G T C A C G T A G T C A C G T A G T C C G T A A G T C A C T G A C T G A G T C | 1e-4 | -1.064e+01 | 0.17% | 0.01% | 667.0bp (126.3bp) | Su(H)/dmmpmm(Bergman)/fly(0.707) More Information \| Similar Motifs Found | motif file (matrix) |
| 22 \* | C G T A A C G T G T C A A C G T C G T A A C G T C G T A A C G T C G T A A G T C | 1e-4 | -9.321e+00 | 4.38% | 2.97% | 286.7bp (442.0bp) | SeqBias: TA-repeat(0.873) More Information \| Similar Motifs Found | motif file (matrix) |
| 23 \* | C G T A A G T C A G T C A C G T C G T A A C G T A C G T A G T C | 1e-2 | -6.295e+00 | 3.23% | 2.27% | 249.6bp (352.9bp) | SRS7(SRS)/colamp-SRS7-DAP-Seq(GSE60143)/Homer(0.775) More Information \| Similar Motifs Found | motif file (matrix) |
| 24 \* | A C G T A G T C A C G T C G T A C G T A A C T G A C T G A C G T | 1e-2 | -5.736e+00 | 98.72% | 97.95% | 389.6bp (465.5bp) | Pp\_0206(RRM)/Physcomitrella\_patens-RNCMPT00206-PBM/HughesRNA(0.718) More Information \| Similar Motifs Found | motif file (matrix) |
| 25 \* | A G T C C G T A A C T G A C T G A C G T A C G T A G T C A C T G A C T G A G T C | 1e-1 | -3.181e+00 | 0.09% | 0.02% | 139.8bp (499.7bp) | OPI1(MacIsaac)/Yeast(0.708) More Information \| Similar Motifs Found | motif file (matrix) |
| 26 \* | A G T C A C G T A G T C A G T C A C G T A C T G A C G T C G T A | 1e-1 | -2.685e+00 | 2.89% | 2.39% | 497.5bp (514.5bp) | SNAI2/MA0745.2/Jaspar(0.729) More Information \| Similar Motifs Found | motif file (matrix) |
