## Supplemental Table 1 for "The Establishment of Cell-Type Specific Gene Regulation in the Sea Urchin Embryo": Hpf24_EctoVeg_homerResults.html

homer\_motifs\_wbackground/Hpf24\_EctoVeg/ - Homer de novo Motif Results


### Homer *de novo* Motif Results (homer\_motifs\_wbackground/Hpf24\_EctoVeg/)

Known Motif Enrichment Results  
Gene Ontology Enrichment Results  
If Homer is having trouble matching a motif to a known motif, try copy/pasting the matrix file into
STAMP  
More information on motif finding results: HOMER
| Description of Results
| Tips
  
Total target sequences = 1612  
Total background sequences = 32541  
\* - possible false positive  

|  |  |  |  |  |  |  |  |  |
| --- | --- | --- | --- | --- | --- | --- | --- | --- |
| Rank | Motif | P-value | log P-pvalue | % of Targets | % of Background | STD(Bg STD) | Best Match/Details | Motif File |
| 1 | G A C T A G C T G A T C A G C T C G T A G A C T C A G T T C A G G C A T G C A T | 1e-20 | -4.680e+01 | 41.13% | 30.12% | 345.0bp (425.0bp) | PB0073.1\_Sox7\_1/Jaspar(0.854) More Information \| Similar Motifs Found | motif file (matrix) |
| 2 | C T G A A G C T A T G C A T C G C G T A C A G T G C A T T A G C C G T A A G T C | 1e-15 | -3.543e+01 | 42.93% | 33.26% | 360.3bp (420.8bp) | ceh-28/MA1445.1/Jaspar(0.743) More Information \| Similar Motifs Found | motif file (matrix) |
| 3 | A G T C A G T C A G T C C G T A A G T C A C T G C G T A A C G T A G T C A C G T | 1e-15 | -3.500e+01 | 0.81% | 0.03% | 334.1bp (973.2bp) | YPR022C/MA0436.1/Jaspar(0.700) More Information \| Similar Motifs Found | motif file (matrix) |
| 4 | A C G T C G T A C G T A A C G T A G T C A G T C | 1e-15 | -3.472e+01 | 49.38% | 39.55% | 348.2bp (411.4bp) | bcd/MA0212.1/Jaspar(0.998) More Information \| Similar Motifs Found | motif file (matrix) |
| 5 | C G A T G T A C G C T A C G T A T C G A T C A G G C T A T C A G T G A C G A C T | 1e-14 | -3.255e+01 | 47.15% | 37.71% | 402.1bp (435.9bp) | TCF7/MA0769.2/Jaspar(0.772) More Information \| Similar Motifs Found | motif file (matrix) |
| 6 | A C T G C T G A C G A T A C T G C T G A A T C G G A C T G T A C C G T A C G A T | 1e-13 | -3.195e+01 | 9.99% | 5.25% | 370.5bp (424.4bp) | NFE2/MA0841.1/Jaspar(0.982) More Information \| Similar Motifs Found | motif file (matrix) |
| 7 | C A T G C A G T A C T G G A T C A C T G T C G A T G C A C A T G T A G C C G T A | 1e-13 | -3.032e+01 | 5.02% | 1.96% | 359.4bp (371.1bp) | Tbr1(T-box)/Cortex-Tbr1-ChIP-Seq(GSE71384)/Homer(0.797) More Information \| Similar Motifs Found | motif file (matrix) |
| 8 | A C G T G T C A G T C A C G T A G T C A C T A G A G T C A G T C | 1e-12 | -2.958e+01 | 58.50% | 49.39% | 368.2bp (423.0bp) | MYNN(Zf)/HEK293-MYNN.eGFP-ChIP-Seq(Encode)/Homer(0.844) More Information \| Similar Motifs Found | motif file (matrix) |
| 9 | G T C A A G C T T G A C G T C A C G T A C G T A C T A G T C A G C T G A C A G T | 1e-12 | -2.921e+01 | 32.94% | 24.86% | 397.4bp (441.4bp) | PB0084.1\_Tcf7l2\_1/Jaspar(0.862) More Information \| Similar Motifs Found | motif file (matrix) |
| 10 \* | G T C A G C T A G A T C G T A C A G T C G T A C G A T C A G C T G C A T A C T G | 1e-10 | -2.402e+01 | 27.48% | 20.65% | 324.5bp (410.6bp) | MSN4(MacIsaac)/Yeast(0.790) More Information \| Similar Motifs Found | motif file (matrix) |
| 11 \* | C T A G T C A G T C G A T A G C G T A C C T G A G A C T C G T A T G C A T G C A | 1e-10 | -2.387e+01 | 48.14% | 40.12% | 379.2bp (433.7bp) | caudal(Homeobox)/Drosophila-Embryos-ChIP-Chip(modEncode)/Homer(0.794) More Information \| Similar Motifs Found | motif file (matrix) |
| 12 \* | T C G A A C T G T C A G G C T A A C G T A G T C A G C T G A C T G C A T T C G A | 1e-10 | -2.372e+01 | 2.73% | 0.85% | 369.7bp (411.6bp) | DOF2(C2C2(Zn) Dof)/Zea mays/AthaMap(0.664) More Information \| Similar Motifs Found | motif file (matrix) |
| 13 \* | A T G C A C G T A C G T A C G T C G T A A G T C A C T G C G T A | 1e-10 | -2.305e+01 | 28.78% | 21.98% | 394.1bp (432.7bp) | HOXB9/MA1503.1/Jaspar(0.881) More Information \| Similar Motifs Found | motif file (matrix) |
| 14 \* | G T A C G T C A G T A C C T G A G T C A A G C T C T G A G T A C T A G C T G A C | 1e-9 | -2.076e+01 | 6.82% | 3.67% | 383.9bp (421.5bp) | UNC-75(RRM)/Caenorhabditis\_elegans-RNCMPT00081-PBM/HughesRNA(0.773) More Information \| Similar Motifs Found | motif file (matrix) |
| 15 \* | A T G C A C G T A C G T G T A C A C T G A G T C A C T G C G T A C G T A A G T C | 1e-8 | -1.999e+01 | 0.74% | 0.07% | 280.7bp (215.9bp) | SWI4/MA0401.1/Jaspar(0.697) More Information \| Similar Motifs Found | motif file (matrix) |
| 16 \* | C G T A A C T G C G T A C G T A A C G T A C G T A G T C A C T G A C T G A G T C | 1e-8 | -1.852e+01 | 0.43% | 0.02% | 104.3bp (254.5bp) | RRTF1(AP2EREBP)/colamp-RRTF1-DAP-Seq(GSE60143)/Homer(0.654) More Information \| Similar Motifs Found | motif file (matrix) |
| 17 \* | A C T G C G T A A C T G A G T C A G T C C G T A A G T C C T G A | 1e-7 | -1.637e+01 | 3.72% | 1.74% | 422.6bp (417.0bp) | MET32/MA0334.1/Jaspar(0.853) More Information \| Similar Motifs Found | motif file (matrix) |
| 18 \* | A C G T A T C G C G T A T C G A C G A T G A C T A G T C C G T A C G T A G A C T | 1e-5 | -1.374e+01 | 9.37% | 6.29% | 376.7bp (422.8bp) | PB0068.1\_Sox1\_1/Jaspar(0.809) More Information \| Similar Motifs Found | motif file (matrix) |
| 19 \* | A C T G A C T G A C G T C G T A C G T A A C T G A C T G C G T A A G T C A C G T | 1e-5 | -1.324e+01 | 0.37% | 0.02% | 350.4bp (297.4bp) | PCBP1(KH)/Homo\_sapiens-RNCMPT00186-PBM/HughesRNA(0.744) More Information \| Similar Motifs Found | motif file (matrix) |
| 20 \* | A C T G A G T C A C T G A C G T A C G T A G T C A C T G A C T G A C T G A G T C | 1e-4 | -1.122e+01 | 0.37% | 0.03% | 130.5bp (381.7bp) | Lm\_0254(RRM)/Leishmania\_major-RNCMPT00254-PBM/HughesRNA(0.708) More Information \| Similar Motifs Found | motif file (matrix) |
| 21 \* | A C G T A G C T C G T A A G T C G T C A A G T C A G T C A C G T | 1e-4 | -1.101e+01 | 12.03% | 8.93% | 399.5bp (418.8bp) | TBX4/MA0806.1/Jaspar(0.902) More Information \| Similar Motifs Found | motif file (matrix) |
| 22 \* | A G T C A C T G A G T C A G T C T C A G A C G T A C G T T C G A | 1e-4 | -1.015e+01 | 9.31% | 6.69% | 392.7bp (424.1bp) | PUCHI(AP2EREBP)/colamp-PUCHI-DAP-Seq(GSE60143)/Homer(0.853) More Information \| Similar Motifs Found | motif file (matrix) |
| 23 \* | A C T G A C G T A G T C A C G T A C T G A G T C A C G T C G T A A C G T C T A G | 1e-4 | -9.627e+00 | 1.36% | 0.53% | 305.4bp (282.6bp) | ZNF317/MA1593.1/Jaspar(0.757) More Information \| Similar Motifs Found | motif file (matrix) |
| 24 \* | A C G T A G T C A C T G A C T G A C T G A C G T A T C G A C G T A G T C C T G A | 1e-1 | -4.140e+00 | 0.50% | 0.20% | 290.3bp (441.6bp) | AFT2/MA0270.1/Jaspar(0.764) More Information \| Similar Motifs Found | motif file (matrix) |
| 25 \* | A C T G C G A T C T G A A T C G A G C T C T G A A C T G G A C T C G T A A T C G | 1e0 | -1.297e+00 | 0.81% | 0.66% | 359.4bp (461.3bp) | An\_0287(RRM)/Aspergillus\_nidulans-RNCMPT00287-PBM/HughesRNA(0.709) More Information \| Similar Motifs Found | motif file (matrix) |
