## Supplemental Table 1 for "The Establishment of Cell-Type Specific Gene Regulation in the Sea Urchin Embryo": Hpf24_EndoVeg1_homerResults.html

homer\_motifs\_wbackground/Hpf24\_EndoVeg1/ - Homer de novo Motif Results


### Homer *de novo* Motif Results (homer\_motifs\_wbackground/Hpf24\_EndoVeg1/)

Known Motif Enrichment Results  
Gene Ontology Enrichment Results  
If Homer is having trouble matching a motif to a known motif, try copy/pasting the matrix file into
STAMP  
More information on motif finding results: HOMER
| Description of Results
| Tips
  
Total target sequences = 4048  
Total background sequences = 27775  
\* - possible false positive  

|  |  |  |  |  |  |  |  |  |
| --- | --- | --- | --- | --- | --- | --- | --- | --- |
| Rank | Motif | P-value | log P-pvalue | % of Targets | % of Background | STD(Bg STD) | Best Match/Details | Motif File |
| 1 | G C A T T C G A C G T A C T A G G A T C G T C A G T C A C T G A A G T C G C T A | 1e-138 | -3.186e+02 | 52.79% | 33.57% | 369.8bp (426.8bp) | fkh/dmmpmm(Noyes)/fly(0.957) More Information \| Similar Motifs Found | motif file (matrix) |
| 2 | C A T G C T G A A C G T A C T G G T C A A T C G A C G T G T A C C G T A G C A T | 1e-33 | -7.626e+01 | 15.54% | 9.51% | 391.4bp (432.6bp) | BATF(bZIP)/Th17-BATF-ChIP-Seq(GSE39756)/Homer(0.992) More Information \| Similar Motifs Found | motif file (matrix) |
| 3 | C A G T A C G T A C G T C G T A A T G C G C A T C T G A C T G A A G T C A G T C | 1e-24 | -5.604e+01 | 3.36% | 1.21% | 367.5bp (356.8bp) | ARR1/Literature(Harbison)/Yeast(0.759) More Information \| Similar Motifs Found | motif file (matrix) |
| 4 | A C T G C T G A G T C A A C G T A C G T A G T C C T G A A G T C C T G A C G T A | 1e-22 | -5.144e+01 | 3.85% | 1.58% | 363.9bp (398.0bp) | Tb\_0251(RRM)/Trypanosoma\_brucei-RNCMPT00251-PBM/HughesRNA(0.795) More Information \| Similar Motifs Found | motif file (matrix) |
| 5 | T G A C A C G T C G T A A C G T A C G T A C T G C G T A A G T C | 1e-20 | -4.753e+01 | 29.67% | 23.23% | 400.4bp (413.5bp) | Hnf6b(Homeobox)/LNCaP-Hnf6b-ChIP-Seq(GSE106305)/Homer(0.835) More Information \| Similar Motifs Found | motif file (matrix) |
| 6 | A G T C A G T C C G A T A G C T A G C T C T A G A C G T T C A G | 1e-20 | -4.732e+01 | 33.30% | 26.61% | 398.2bp (419.0bp) | SOX10/MA0442.2/Jaspar(0.882) More Information \| Similar Motifs Found | motif file (matrix) |
| 7 | A G T C C G T A C G T A C G T A A G T C A G T C A C G T A G T C A G T C A C G T | 1e-20 | -4.732e+01 | 2.50% | 0.83% | 351.0bp (353.4bp) | Eip74EF/dmmpmm(SeSiMCMC)/fly(0.662) More Information \| Similar Motifs Found | motif file (matrix) |
| 8 | C T G A A T G C T G A C C G T A C G T A A C T G C T A G A C G T C A G T C G A T | 1e-20 | -4.710e+01 | 5.19% | 2.54% | 394.8bp (408.2bp) | HuR(RRM)/Homo\_sapiens-RNCMPT00136-PBM/HughesRNA(0.579) More Information \| Similar Motifs Found | motif file (matrix) |
| 9 | C T G A C G T A C G T A G A T C A C T G G A T C C T A G G C A T G T A C C T G A | 1e-20 | -4.650e+01 | 3.21% | 1.26% | 381.5bp (343.3bp) | MBP1::SWI6/MA0330.1/Jaspar(0.810) More Information \| Similar Motifs Found | motif file (matrix) |
| 10 | A C G T C T A G G T C A A G T C A C T G A C G T C G T A A G T C A C T G C G T A | 1e-16 | -3.901e+01 | 1.63% | 0.46% | 385.0bp (362.5bp) | SPL7/MA1060.1/Jaspar(0.805) More Information \| Similar Motifs Found | motif file (matrix) |
| 11 | A T C G A G C T C T G A C G T A A C T G A T C G G A C T G C A T C G A T A T C G | 1e-14 | -3.398e+01 | 2.69% | 1.14% | 345.8bp (419.7bp) | Pp\_0206(RRM)/Physcomitrella\_patens-RNCMPT00206-PBM/HughesRNA(0.762) More Information \| Similar Motifs Found | motif file (matrix) |
| 12 | C T A G A T C G C A G T A G T C A G T C T C G A C G A T T A G C C A T G G A C T | 1e-14 | -3.357e+01 | 4.47% | 2.37% | 352.6bp (404.6bp) | TCP16(TCP)/colamp-TCP16-DAP-Seq(GSE60143)/Homer(0.689) More Information \| Similar Motifs Found | motif file (matrix) |
| 13 \* | A G T C A G T C A G T C C G T A A G T C A C T G C G T A A C G T A G T C A C G T | 1e-11 | -2.736e+01 | 0.32% | 0.02% | 610.9bp (220.5bp) | YPR022C/MA0436.1/Jaspar(0.700) More Information \| Similar Motifs Found | motif file (matrix) |
| 14 \* | A G T C A C T G A G C T A G T C A G T C C G T A A C T G A C G T A C T G A C G T | 1e-11 | -2.552e+01 | 0.44% | 0.05% | 365.9bp (255.0bp) | SOX2/MA0143.4/Jaspar(0.660) More Information \| Similar Motifs Found | motif file (matrix) |
| 15 \* | C G T A A G T C G C T A C G T A A C T G A G T C A G T C A G T C | 1e-9 | -2.185e+01 | 11.24% | 8.41% | 369.1bp (412.7bp) | ZNF682/MA1599.1/Jaspar(0.740) More Information \| Similar Motifs Found | motif file (matrix) |
| 16 \* | A C T G C T G A G T C A C G A T C T A G G T C A A T G C A T G C | 1e-9 | -2.156e+01 | 24.70% | 20.70% | 414.4bp (426.9bp) | NR1H4/MA1110.1/Jaspar(0.754) More Information \| Similar Motifs Found | motif file (matrix) |
| 17 \* | A C T G A C G T A G C T G T A C A T G C A G T C G T C A A G C T | 1e-9 | -2.092e+01 | 13.96% | 10.89% | 396.3bp (419.2bp) | RBPJ/MA1116.1/Jaspar(0.804) More Information \| Similar Motifs Found | motif file (matrix) |
| 18 \* | A C T G A C T G C G T A C G T A A C G T A G T C A C G T A G T C A G T C A C T G | 1e-9 | -2.073e+01 | 0.20% | 0.01% | 331.1bp (212.7bp) | PDR8/MA0354.1/Jaspar(0.821) More Information \| Similar Motifs Found | motif file (matrix) |
| 19 \* | C G T A C A T G G T C A A T G C C G A T G T A C G C A T C T A G | 1e-7 | -1.816e+01 | 14.30% | 11.42% | 402.2bp (449.7bp) | ARG80(MacIsaac)/Yeast(0.760) More Information \| Similar Motifs Found | motif file (matrix) |
| 20 \* | C G T A A C G T C G T A T G C A A G T C A C T G A T C G A C T G | 1e-7 | -1.726e+01 | 7.83% | 5.75% | 431.9bp (411.0bp) | STP4/MA0397.1/Jaspar(0.715) More Information \| Similar Motifs Found | motif file (matrix) |
| 21 \* | A C T G A C G T A G T C C G T A A C G T C G T A C G T A C T G A | 1e-7 | -1.710e+01 | 8.92% | 6.70% | 435.2bp (434.7bp) | caudal(Homeobox)/Drosophila-Embryos-ChIP-Chip(modEncode)/Homer(0.866) More Information \| Similar Motifs Found | motif file (matrix) |
| 22 \* | T A G C A C G T T A G C A G T C A C G T C G T A A G T C C G A T C T G A G T A C | 1e-7 | -1.685e+01 | 3.06% | 1.83% | 465.5bp (407.2bp) | MSI(RRM)/Drosophila\_melanogaster-RNCMPT00099-PBM/HughesRNA(0.752) More Information \| Similar Motifs Found | motif file (matrix) |
| 23 \* | A C G T A C G T A C T G A C G T A C G T A G T C | 1e-6 | -1.488e+01 | 66.43% | 62.67% | 394.8bp (417.9bp) | Ot\_0263(RRM)/Ostreococcus\_tauri-RNCMPT00263-PBM/HughesRNA(0.831) More Information \| Similar Motifs Found | motif file (matrix) |
| 24 \* | C T G A C A G T C T G A A T G C A G C T A G T C T G A C C G T A | 1e-6 | -1.483e+01 | 29.79% | 26.31% | 431.8bp (424.5bp) | Pp\_0237(RRM)/Physcomitrella\_patens-RNCMPT00237-PBM/HughesRNA(0.788) More Information \| Similar Motifs Found | motif file (matrix) |
| 25 \* | C G T A A C T G A C G T A G T C A C T G A C T G A C T G A C G T C G T A A C T G | 1e-4 | -1.120e+01 | 0.12% | 0.01% | 298.1bp (209.6bp) | STB2(MacIsaac)/Yeast(0.685) More Information \| Similar Motifs Found | motif file (matrix) |
| 26 \* | C T A G A C G T A C G T C G T A C G T A A G T C | 1e-1 | -3.847e+00 | 51.01% | 49.41% | 414.8bp (412.8bp) | PB0081.1\_Tcf1\_1/Jaspar(0.852) More Information \| Similar Motifs Found | motif file (matrix) |
| 27 \* | A G T C A C T G A C T G A C G T A C G T A C T G A C G T A G T C A G T C A C G T | 1e0 | -2.151e+00 | 0.05% | 0.02% | 327.1bp (46.8bp) | SRSF1(RRM)/Homo\_sapiens-RNCMPT00110-PBM/HughesRNA(0.758) More Information \| Similar Motifs Found | motif file (matrix) |
| 28 \* | A G T C A C G T A G T C A G T C A G T C A C G T | 1e0 | -8.818e-01 | 32.88% | 32.71% | 431.5bp (408.0bp) | HNRNPA2B1(RRM)/Homo\_sapiens-RNCMPT00024-PBM/HughesRNA(0.833) More Information \| Similar Motifs Found | motif file (matrix) |
