## Supplemental Table 1 for "The Establishment of Cell-Type Specific Gene Regulation in the Sea Urchin Embryo": Hpf24_EndoVeg2_homerResults.html

homer\_motifs\_wbackground/Hpf24\_EndoVeg2/ - Homer de novo Motif Results


### Homer *de novo* Motif Results (homer\_motifs\_wbackground/Hpf24\_EndoVeg2/)

Known Motif Enrichment Results  
Gene Ontology Enrichment Results  
If Homer is having trouble matching a motif to a known motif, try copy/pasting the matrix file into
STAMP  
More information on motif finding results: HOMER
| Description of Results
| Tips
  
Total target sequences = 4235  
Total background sequences = 26895  
\* - possible false positive  

|  |  |  |  |  |  |  |  |  |
| --- | --- | --- | --- | --- | --- | --- | --- | --- |
| Rank | Motif | P-value | log P-pvalue | % of Targets | % of Background | STD(Bg STD) | Best Match/Details | Motif File |
| 1 | C G A T T C A G A C G T C A G T C A G T C T A G G A T C G C A T A G T C C G T A | 1e-147 | -3.392e+02 | 58.44% | 38.72% | 369.1bp (430.1bp) | fkh/dmmpmm(Noyes)/fly(0.919) More Information \| Similar Motifs Found | motif file (matrix) |
| 2 | T C G A C A G T T C G A C G A T A C G T C T A G C T G A T A G C C G A T A G T C | 1e-21 | -4.973e+01 | 5.69% | 2.88% | 334.0bp (391.6bp) | Hnf6b(Homeobox)/LNCaP-Hnf6b-ChIP-Seq(GSE106305)/Homer(0.781) More Information \| Similar Motifs Found | motif file (matrix) |
| 3 | A T G C G A C T C G T A T C A G A T G C G A C T G T A C A G C T T A C G G C T A | 1e-17 | -4.019e+01 | 4.20% | 2.06% | 392.1bp (432.3bp) | AGP1(GATA)/Nicotiana tabacum/AthaMap(0.737) More Information \| Similar Motifs Found | motif file (matrix) |
| 4 | G T A C G T A C T G C A G A C T A G T C A G C T G A T C A G T C G T A C G C A T | 1e-15 | -3.647e+01 | 12.99% | 9.15% | 413.5bp (412.7bp) | MYB30(MYB)/colamp-MYB30-DAP-Seq(GSE60143)/Homer(0.676) More Information \| Similar Motifs Found | motif file (matrix) |
| 5 | T C A G G T A C G A C T G C A T G C T A C T A G A T G C G T A C C A G T T G C A | 1e-14 | -3.354e+01 | 14.52% | 10.63% | 385.3bp (401.5bp) | HRB27C(RRM)/Drosophila\_melanogaster-RNCMPT00093-PBM/HughesRNA(0.695) More Information \| Similar Motifs Found | motif file (matrix) |
| 6 | A T C G A C G T A G T C A C G T A C T G A G C T A G T C C G T A C T A G A G T C | 1e-14 | -3.249e+01 | 1.35% | 0.40% | 448.4bp (425.3bp) | achi/MA0207.1/Jaspar(0.818) More Information \| Similar Motifs Found | motif file (matrix) |
| 7 | A G T C G T A C A C T G G A T C A C T G T G C A C G A T A C T G | 1e-13 | -3.148e+01 | 13.32% | 9.70% | 398.3bp (381.1bp) | RBM4(RRM,Znf)/Homo\_sapiens-RNCMPT00052-PBM/HughesRNA(0.730) More Information \| Similar Motifs Found | motif file (matrix) |
| 8 | T G C A A C T G A G T C T C A G C A T G G A C T A G C T C G T A C T A G A G C T | 1e-12 | -2.886e+01 | 7.82% | 5.18% | 372.0bp (401.5bp) | HOW(KH)/Drosophila\_melanogaster-RNCMPT00118-PBM/HughesRNA(0.747) More Information \| Similar Motifs Found | motif file (matrix) |
| 9 | A T C G A C G T A C T G C A G T T C A G C G A T | 1e-12 | -2.819e+01 | 49.78% | 44.32% | 387.2bp (417.8bp) | SM(RRM)/Drosophila\_melanogaster-RNCMPT00069-PBM/HughesRNA(0.896) More Information \| Similar Motifs Found | motif file (matrix) |
| 10 | C T A G C G T A T G C A A G T C C G A T A C G T A C G T C T A G | 1e-12 | -2.775e+01 | 35.70% | 30.63% | 400.1bp (415.8bp) | NR1I3/MA1534.1/Jaspar(0.840) More Information \| Similar Motifs Found | motif file (matrix) |
| 11 \* | C G T A T G C A A G T C C G T A A C G T A C T G A C T G A C T G | 1e-11 | -2.650e+01 | 11.83% | 8.70% | 409.8bp (427.4bp) | REI1/MA0364.1/Jaspar(0.743) More Information \| Similar Motifs Found | motif file (matrix) |
| 12 \* | A C G T T G A C A C G T C A G T C G T A A C G T A G T C A C G T A C G T G T C A | 1e-9 | -2.234e+01 | 6.87% | 4.70% | 351.0bp (503.5bp) | GATA6/MA1104.2/Jaspar(0.786) More Information \| Similar Motifs Found | motif file (matrix) |
| 13 \* | C A G T C T A G T A G C C T A G G A C T G T A C C G A T A T C G | 1e-8 | -2.049e+01 | 30.70% | 26.58% | 402.5bp (429.9bp) | ARG80(MacIsaac)/Yeast(0.921) More Information \| Similar Motifs Found | motif file (matrix) |
| 14 \* | T C A G G T C A A C T G A C T G A G C T A C T G G T C A A C T G | 1e-8 | -2.042e+01 | 22.79% | 19.10% | 410.0bp (413.7bp) | ZKSCAN5/MA1652.1/Jaspar(0.808) More Information \| Similar Motifs Found | motif file (matrix) |
| 15 \* | A C T G A C T G A C T G C G T A A C G T C G T A A G T C A G T C A T G C A G T C | 1e-8 | -2.031e+01 | 0.24% | 0.02% | 130.9bp (376.4bp) | dif/Rel/dmmpmm(Bergman)/fly(0.814) More Information \| Similar Motifs Found | motif file (matrix) |
| 16 \* | G A T C G T A C A G C T G A T C T A G C G A C T A G T C T G A C A G C T G T A C | 1e-8 | -2.030e+01 | 4.82% | 3.11% | 399.2bp (485.4bp) | TF3A(C2H2)/col-TF3A-DAP-Seq(GSE60143)/Homer(0.821) More Information \| Similar Motifs Found | motif file (matrix) |
| 17 \* | A G T C A G T C A G T C A C T G A C T G A C T G C G T A C G T A A C T G A C T G | 1e-8 | -1.998e+01 | 0.21% | 0.01% | 335.3bp (194.3bp) | PUT3/MA0358.1/Jaspar(0.837) More Information \| Similar Motifs Found | motif file (matrix) |
| 18 \* | C G A T C G T A A G T C A G T C A G T C C G T A C G T A A G C T | 1e-8 | -1.981e+01 | 8.74% | 6.42% | 408.2bp (431.7bp) | NCU02182/MA1434.1/Jaspar(0.808) More Information \| Similar Motifs Found | motif file (matrix) |
| 19 \* | C A T G A G T C C G T A T C G A A C T G G A T C C G T A C T A G A C T G A T G C | 1e-7 | -1.790e+01 | 2.69% | 1.54% | 384.4bp (417.1bp) | MBNL1(Znf)/Homo\_sapiens-RNCMPT00038-PBM/HughesRNA(0.739) More Information \| Similar Motifs Found | motif file (matrix) |
| 20 \* | A C G T A C G T A C G T G T C A A C G T A G C T A C G T A C T G | 1e-7 | -1.790e+01 | 30.67% | 26.85% | 396.5bp (419.3bp) | ZC3H14(Znf)/Homo\_sapiens-RNCMPT00086-PBM/HughesRNA(0.842) More Information \| Similar Motifs Found | motif file (matrix) |
| 21 \* | A C G T A G T C G T A C A G T C C G T A A C T G C G T A C T A G | 1e-7 | -1.712e+01 | 7.89% | 5.84% | 426.9bp (445.2bp) | Stat5b/MA1625.1/Jaspar(0.806) More Information \| Similar Motifs Found | motif file (matrix) |
| 22 \* | A G T C A C G T A C T G A C T G A C G T A G T C A C G T A C T G A C G T C G T A | 1e-6 | -1.490e+01 | 0.43% | 0.10% | 436.1bp (405.2bp) | ERF018/MA1048.1/Jaspar(0.683) More Information \| Similar Motifs Found | motif file (matrix) |
| 23 \* | A G T C A C G T C G T A A C T G A C T G A C T G C G T A A C G T A G T C A C G T | 1e-6 | -1.417e+01 | 0.21% | 0.02% | 143.1bp (409.1bp) | HNRNPA2B1(RRM)/Homo\_sapiens-RNCMPT00024-PBM/HughesRNA(0.730) More Information \| Similar Motifs Found | motif file (matrix) |
| 24 \* | A C G T A G T C A C G T A C G T A C T G C G T A A C G T A G T C | 1e-6 | -1.405e+01 | 4.79% | 3.37% | 413.2bp (345.4bp) | SNRNP70K(RRM)/Drosophila\_melanogaster-RNCMPT00143-PBM/HughesRNA(0.945) More Information \| Similar Motifs Found | motif file (matrix) |
| 25 \* | A C T G A C G T C G T A A G T C C G T A A C G T C G T A C T G A | 1e-5 | -1.362e+01 | 12.73% | 10.44% | 404.7bp (442.0bp) | YAP6(MacIsaac)/Yeast(0.826) More Information \| Similar Motifs Found | motif file (matrix) |
| 26 \* | A G T C C G T A A G T C A C T G A G T C A C T G A C G T C T G A A G T C T C A G | 1e-5 | -1.238e+01 | 0.85% | 0.36% | 331.7bp (290.5bp) | Ahr::Arnt/MA0006.1/Jaspar(0.745) More Information \| Similar Motifs Found | motif file (matrix) |
| 27 \* | A C G T A G T C A G T C A C G T A G T C A C T G A C G T A C T G A C T G A C G T | 1e-3 | -6.932e+00 | 0.14% | 0.03% | 492.8bp (247.0bp) | PIF7(bHLH)/col-PIF7-DAP-Seq(GSE60143)/Homer(0.718) More Information \| Similar Motifs Found | motif file (matrix) |
| 28 \* | A C T G A G T C A C T G C G T A A C T G A C G T A C G T A C G T A C T G A C G T | 1e-2 | -5.703e+00 | 0.17% | 0.05% | 175.2bp (355.2bp) | Tv\_0258(RRM)/Trichomonas\_vaginalis-RNCMPT00258-PBM/HughesRNA(0.670) More Information \| Similar Motifs Found | motif file (matrix) |
| 29 \* | T G A C A G C T A C G T C G T A T A G C A C G T | 1e-1 | -3.935e+00 | 27.32% | 25.91% | 390.8bp (408.8bp) | Tb\_0220(RRM)/Trypanosoma\_brucei-RNCMPT00220-PBM/HughesRNA(0.919) More Information \| Similar Motifs Found | motif file (matrix) |
| 30 \* | A G T C A C G T A G T C A C G T A C G T A C G T | 1e0 | -2.974e-01 | 56.60% | 57.08% | 407.5bp (413.5bp) | DOF5.7/MA0984.1/Jaspar(0.837) More Information \| Similar Motifs Found | motif file (matrix) |
