## Supplemental Table 1 for "The Establishment of Cell-Type Specific Gene Regulation in the Sea Urchin Embryo": Hpf24_NSM_homerResults.html

homer\_motifs\_wbackground/Hpf24\_NSM/ - Homer de novo Motif Results


### Homer *de novo* Motif Results (homer\_motifs\_wbackground/Hpf24\_NSM/)

Known Motif Enrichment Results  
Gene Ontology Enrichment Results  
If Homer is having trouble matching a motif to a known motif, try copy/pasting the matrix file into
STAMP  
More information on motif finding results: HOMER
| Description of Results
| Tips
  
Total target sequences = 4862  
Total background sequences = 24261  
\* - possible false positive  

|  |  |  |  |  |  |  |  |  |
| --- | --- | --- | --- | --- | --- | --- | --- | --- |
| Rank | Motif | P-value | log P-pvalue | % of Targets | % of Background | STD(Bg STD) | Best Match/Details | Motif File |
| 1 | T C A G T A C G A G C T T C G A G A T C G T A C G T A C A C T G A G T C C G T A | 1e-819 | -1.887e+03 | 45.45% | 10.74% | 363.6bp (421.1bp) | GCM1/MA0646.1/Jaspar(0.882) More Information \| Similar Motifs Found | motif file (matrix) |
| 2 | C G A T C G T A A C G T T A C G T G C A T A C G C A T G A C T G G A T C G T C A | 1e-58 | -1.356e+02 | 13.37% | 6.79% | 416.9bp (370.9bp) | PUF68(RRM)/Drosophila\_melanogaster-RNCMPT00141-PBM/HughesRNA(0.799) More Information \| Similar Motifs Found | motif file (matrix) |
| 3 | C A G T T A G C G C T A A C G T C G T A C G A T G T A C G C A T | 1e-42 | -9.835e+01 | 47.02% | 37.32% | 427.6bp (413.7bp) | GATA6/MA1104.2/Jaspar(0.820) More Information \| Similar Motifs Found | motif file (matrix) |
| 4 | A C T G C G T A A T G C G T A C A C T G A C T G C G T A G C T A C T A G A C G T | 1e-41 | -9.495e+01 | 21.74% | 14.50% | 428.8bp (399.8bp) | Eip74EF/MA0026.1/Jaspar(0.960) More Information \| Similar Motifs Found | motif file (matrix) |
| 5 | G C T A C G A T C G A T C G T A C G T A C G A T C G T A C G A T G C A T G T C A | 1e-29 | -6.741e+01 | 51.46% | 43.37% | 417.5bp (423.3bp) | SHEP(RRM)/Drosophila\_melanogaster-RNCMPT00068-PBM/HughesRNA(0.862) More Information \| Similar Motifs Found | motif file (matrix) |
| 6 | A G T C C G T A C G A T C G A T A C G T A C G T A C G T A C G T | 1e-26 | -6.026e+01 | 39.14% | 31.87% | 426.2bp (429.1bp) | Smp\_067420(RRM)/Schistosoma\_mansoni-RNCMPT00232-PBM/HughesRNA(0.872) More Information \| Similar Motifs Found | motif file (matrix) |
| 7 | A C G T T C A G A G C T C T G A A T G C C G T A G A C T C T A G G A C T G C T A | 1e-17 | -4.041e+01 | 54.11% | 47.91% | 436.7bp (472.0bp) | PH0084.1\_Irx3\_2/Jaspar(0.881) More Information \| Similar Motifs Found | motif file (matrix) |
| 8 | A C G T G A C T C T G A C G A T G A C T C G A T C G A T C G T A | 1e-16 | -3.910e+01 | 43.50% | 37.54% | 417.2bp (421.5bp) | HuR(RRM)/Homo\_sapiens-RNCMPT00032-PBM/HughesRNA(0.901) More Information \| Similar Motifs Found | motif file (matrix) |
| 9 | A C T G A G T C A G T C A C T G A G T C A G T C A C T G C G T A A G T C C G T A | 1e-15 | -3.538e+01 | 0.23% | 0.01% | 579.9bp (18.0bp) | ERF021/MA1233.2/Jaspar(0.935) More Information \| Similar Motifs Found | motif file (matrix) |
| 10 | A C T G A C T G C G A T C G T A A C T G A C T G A C T G A C G T | 1e-12 | -2.868e+01 | 36.30% | 31.45% | 418.8bp (407.6bp) | HRB98DE(RRM)/Drosophila\_melanogaster-RNCMPT00095-PBM/HughesRNA(0.899) More Information \| Similar Motifs Found | motif file (matrix) |
| 11 \* | A C G T A G T C A C T G G T A C C G T A A C G T C G T A A C G T | 1e-11 | -2.565e+01 | 9.46% | 6.88% | 401.3bp (399.6bp) | KAN1/MA1027.1/Jaspar(0.717) More Information \| Similar Motifs Found | motif file (matrix) |
| 12 \* | A G T C C G T A A G T C A C T G A C T G A G T C A G T C C G T A A G T C C T A G | 1e-10 | -2.344e+01 | 0.49% | 0.09% | 203.8bp (282.9bp) | KLF11/MA1512.1/Jaspar(0.737) More Information \| Similar Motifs Found | motif file (matrix) |
| 13 \* | A C T G C G T A A C T G A C T G A C G T A G T C C G T A A C T G A C T G C G T A | 1e-9 | -2.178e+01 | 0.29% | 0.03% | 426.3bp (542.2bp) | COUP-TFII(NR)/Artia-Nr2f2-ChIP-Seq(GSE46497)/Homer(0.783) More Information \| Similar Motifs Found | motif file (matrix) |
| 14 \* | A C T G A G T C A G T C C G T A A G T C A T G C A C T G A G T C | 1e-9 | -2.163e+01 | 3.70% | 2.27% | 454.2bp (355.7bp) | AT3G57600(AP2EREBP)/col-AT3G57600-DAP-Seq(GSE60143)/Homer(0.873) More Information \| Similar Motifs Found | motif file (matrix) |
| 15 \* | A G T C A G T C A G T C A G T C A G T C A G T C A G T C A G T C | 1e-9 | -2.118e+01 | 18.94% | 15.69% | 423.9bp (422.8bp) | Maz(Zf)/HepG2-Maz-ChIP-Seq(GSE31477)/Homer(0.974) More Information \| Similar Motifs Found | motif file (matrix) |
| 16 \* | A C G T C G T A C G T A C G T A A C T G C T A G A C G T C G T A | 1e-9 | -2.095e+01 | 14.34% | 11.48% | 440.8bp (426.2bp) | Adof1(C2C2dof)/col-Adof1-DAP-Seq(GSE60143)/Homer(0.743) More Information \| Similar Motifs Found | motif file (matrix) |
| 17 \* | C G T A C G T A A C G T A G T C C G T A C G T A A G T C A C G T A C T G A C T G | 1e-8 | -2.051e+01 | 1.03% | 0.38% | 379.2bp (408.3bp) | MYB/MA0100.3/Jaspar(0.751) More Information \| Similar Motifs Found | motif file (matrix) |
| 18 \* | A C G T A G T C A C G T A G T C A C G T A G T C A C G T A G T C A C G T A G T C | 1e-6 | -1.424e+01 | 13.23% | 10.99% | 446.5bp (367.4bp) | SeqBias: GA-repeat(0.994) More Information \| Similar Motifs Found | motif file (matrix) |
| 19 \* | A T C G A T C G C G T A C G T A A G T C A C T G A C G T A C T G A G T C A G T C | 1e-5 | -1.317e+01 | 0.41% | 0.12% | 424.7bp (357.9bp) | BHLH104/MA0960.1/Jaspar(0.847) More Information \| Similar Motifs Found | motif file (matrix) |
| 20 \* | A C T G A T C G C T G A A C T G A C T G G T C A A T G C A C T G C T G A A C T G | 1e-4 | -1.028e+01 | 0.72% | 0.34% | 357.1bp (309.7bp) | ZNF135/MA1587.1/Jaspar(0.783) More Information \| Similar Motifs Found | motif file (matrix) |
| 21 \* | A C T G C G T A A C G T A C T G A C T G C G T A A C T G A C T G | 1e-1 | -4.593e+00 | 2.94% | 2.41% | 421.6bp (422.1bp) | Pp\_0237(RRM)/Physcomitrella\_patens-RNCMPT00237-PBM/HughesRNA(0.832) More Information \| Similar Motifs Found | motif file (matrix) |
