## Supplemental Table 1 for "The Establishment of Cell-Type Specific Gene Regulation in the Sea Urchin Embryo": Hpf24_OralNSM_homerResults.html

homer\_motifs\_wbackground/Hpf24\_OralNSM/ - Homer de novo Motif Results


### Homer *de novo* Motif Results (homer\_motifs\_wbackground/Hpf24\_OralNSM/)

Known Motif Enrichment Results  
Gene Ontology Enrichment Results  
If Homer is having trouble matching a motif to a known motif, try copy/pasting the matrix file into
STAMP  
More information on motif finding results: HOMER
| Description of Results
| Tips
  
Total target sequences = 4967  
Total background sequences = 26998  
\* - possible false positive  

|  |  |  |  |  |  |  |  |  |
| --- | --- | --- | --- | --- | --- | --- | --- | --- |
| Rank | Motif | P-value | log P-pvalue | % of Targets | % of Background | STD(Bg STD) | Best Match/Details | Motif File |
| 1 | C T G A A G T C A C G T A C G T A G T C A G T C A C G T A C T G C T A G G C A T | 1e-563 | -1.299e+03 | 51.52% | 19.21% | 357.4bp (419.5bp) | EHF(ETS)/LoVo-EHF-ChIP-Seq(GSE49402)/Homer(0.980) More Information \| Similar Motifs Found | motif file (matrix) |
| 2 | G C A T T G A C C G A T T G C A C T A G T C G A A G C T T G A C A C G T T G C A | 1e-52 | -1.215e+02 | 8.44% | 3.67% | 397.9bp (444.1bp) | GATA6(C2C2gata)/col200-GATA6-DAP-Seq(GSE60143)/Homer(0.872) More Information \| Similar Motifs Found | motif file (matrix) |
| 3 | A C T G A T C G C T A G T C G A C T G A C G T A C A G T A G T C T G A C T G A C | 1e-33 | -7.713e+01 | 8.25% | 4.33% | 398.2bp (416.8bp) | dl/dmmpmm(Papatsenko)/fly(0.818) More Information \| Similar Motifs Found | motif file (matrix) |
| 4 | C T A G A G C T C T G A A G T C A T G C C T A G G C A T C T G A A G T C A T C G | 1e-31 | -7.150e+01 | 6.40% | 3.12% | 362.8bp (474.2bp) | POPTR\_0002s00440g/MA0955.1/Jaspar(0.777) More Information \| Similar Motifs Found | motif file (matrix) |
| 5 | C A G T G T C A T C A G C G T A A G T C A C G T C G T A T C A G | 1e-27 | -6.387e+01 | 14.40% | 9.50% | 429.4bp (404.0bp) | GATA1/MA0035.4/Jaspar(0.726) More Information \| Similar Motifs Found | motif file (matrix) |
| 6 | A G T C T C A G A G C T C T G A G T A C A C T G A G C T C T A G | 1e-26 | -5.994e+01 | 34.89% | 27.95% | 379.0bp (434.9bp) | ARNT::HIF1A/MA0259.1/Jaspar(0.871) More Information \| Similar Motifs Found | motif file (matrix) |
| 7 | A G T C C G A T C A T G A T G C G C A T C T A G A G T C C G A T C T A G A T G C | 1e-24 | -5.660e+01 | 9.46% | 5.75% | 413.9bp (421.7bp) | SeqBias: GCW-triplet(0.817) More Information \| Similar Motifs Found | motif file (matrix) |
| 8 | A C T G A G T C A C T G A G T C G T C A A C G T A C T G A G T C | 1e-23 | -5.444e+01 | 10.43% | 6.58% | 356.4bp (461.4bp) | NRF(NRF)/Promoter/Homer(0.852) More Information \| Similar Motifs Found | motif file (matrix) |
| 9 | T G A C C A T G A C T G C G T A A G C T C G T A A G C T T A G C | 1e-23 | -5.410e+01 | 30.70% | 24.39% | 411.5bp (429.5bp) | SD0003.1\_at\_AC\_acceptor/Jaspar(0.806) More Information \| Similar Motifs Found | motif file (matrix) |
| 10 | A G T C C G T A A T G C A G C T A T C G G A T C C G T A A T G C A G C T T C A G | 1e-22 | -5.253e+01 | 8.86% | 5.39% | 418.8bp (386.0bp) | PB0091.1\_Zbtb3\_1/Jaspar(0.878) More Information \| Similar Motifs Found | motif file (matrix) |
| 11 | C T G A C G T A C G T A C A G T C G A T C G A T A T G C C T G A | 1e-21 | -4.889e+01 | 50.29% | 43.53% | 412.5bp (422.9bp) | CG17838(RRM)/Drosophila\_melanogaster-RNCMPT00131-PBM/HughesRNA(0.870) More Information \| Similar Motifs Found | motif file (matrix) |
| 12 | A T G C C A G T A G T C C T G A T C A G G T A C A G C T A G T C C G T A A C T G | 1e-20 | -4.747e+01 | 4.07% | 1.95% | 322.7bp (379.4bp) | ZNF416(Zf)/HEK293-ZNF416.GFP-ChIP-Seq(GSE58341)/Homer(0.627) More Information \| Similar Motifs Found | motif file (matrix) |
| 13 | T G A C A G C T G C T A T A C G G T A C A C G T G T C A C A T G A G T C A C G T | 1e-18 | -4.195e+01 | 7.89% | 4.95% | 383.4bp (399.2bp) | DAZAP1(RRM)/Homo\_sapiens-RNCMPT00013-PBM/HughesRNA(0.753) More Information \| Similar Motifs Found | motif file (matrix) |
| 14 | A T G C T A G C T A G C T A C G T G A C A T C G A T G C A T G C | 1e-18 | -4.151e+01 | 13.65% | 9.76% | 372.0bp (395.6bp) | RBM4(RRM,Znf)/Homo\_sapiens-RNCMPT00113-PBM/HughesRNA(0.872) More Information \| Similar Motifs Found | motif file (matrix) |
| 15 | T C A G A T C G A G T C T A G C G C A T C T A G A C T G T G A C | 1e-15 | -3.539e+01 | 12.16% | 8.76% | 368.9bp (402.7bp) | SKN7(MacIsaac)/Yeast(0.784) More Information \| Similar Motifs Found | motif file (matrix) |
| 16 | A G T C C G T A A C G T A C T G C G T A A C G T A G T C G C T A | 1e-14 | -3.376e+01 | 21.88% | 17.52% | 407.8bp (428.0bp) | GATA15(C2C2gata)/col-GATA15-DAP-Seq(GSE60143)/Homer(0.846) More Information \| Similar Motifs Found | motif file (matrix) |
| 17 | C G A T T A G C G C T A T A G C C G T A T G A C G C A T G T A C G C T A G A T C | 1e-12 | -2.917e+01 | 14.66% | 11.27% | 409.4bp (409.8bp) | SeqBias: GA-repeat(0.723) More Information \| Similar Motifs Found | motif file (matrix) |
| 18 \* | C G T A C G T A C G T A C G T A C G T A C G T A C G T A C G T A T C G A C G T A | 1e-11 | -2.622e+01 | 41.47% | 36.75% | 392.7bp (422.2bp) | SeqBias: polyA-repeat(0.957) More Information \| Similar Motifs Found | motif file (matrix) |
| 19 \* | T A G C T G C A C A T G G C T A A C T G G T A C A T G C G C T A T A C G C T G A | 1e-10 | -2.493e+01 | 14.68% | 11.55% | 418.1bp (396.1bp) | GATA6/MA1396.1/Jaspar(0.721) More Information \| Similar Motifs Found | motif file (matrix) |
| 20 \* | A C T G A C T G A G T C C G T A A C T G A G T C C G T A A G T C | 1e-10 | -2.351e+01 | 1.69% | 0.76% | 369.4bp (392.3bp) | Unknown3/Drosophila-Promoters/Homer(0.740) More Information \| Similar Motifs Found | motif file (matrix) |
| 21 \* | G T A C A T G C C G A T C G T A T G C A A G T C A G C T A G T C A G C T G T A C | 1e-5 | -1.226e+01 | 0.95% | 0.46% | 390.8bp (332.8bp) | MYB116(MYB)/colamp-MYB116-DAP-Seq(GSE60143)/Homer(0.686) More Information \| Similar Motifs Found | motif file (matrix) |
| 22 \* | C G T A C G T A A C G T A C G T C G T A C G T A | 1e-4 | -9.479e+00 | 57.88% | 55.20% | 413.3bp (412.8bp) | hbn/MA0226.1/Jaspar(0.958) More Information \| Similar Motifs Found | motif file (matrix) |
| 23 \* | A C T G A G T C A G T C C G T A A C T G A G T C A G T C C G T A A C T G A G T C | 1e-2 | -4.988e+00 | 0.46% | 0.26% | 341.6bp (335.3bp) | Vts1p(SAM)/Saccharomyces\_cerevisiae-RNCMPT00111-PBM/HughesRNA(0.736) More Information \| Similar Motifs Found | motif file (matrix) |
| 24 \* | A G T C A C G T A G T C A G T C C G T A A C G T A G T C A G T C C G T A A G T C | 1e-1 | -2.420e+00 | 0.20% | 0.12% | 338.6bp (364.6bp) | Pp\_0237(RRM)/Physcomitrella\_patens-RNCMPT00237-PBM/HughesRNA(0.768) More Information \| Similar Motifs Found | motif file (matrix) |
