## Supplemental Table 1 for "The Establishment of Cell-Type Specific Gene Regulation in the Sea Urchin Embryo": Hpf24_PMCs_homerResults.html

homer\_motifs\_wbackground/Hpf24\_PMCs/ - Homer de novo Motif Results


### Homer *de novo* Motif Results (homer\_motifs\_wbackground/Hpf24\_PMCs/)

Known Motif Enrichment Results  
Gene Ontology Enrichment Results  
If Homer is having trouble matching a motif to a known motif, try copy/pasting the matrix file into
STAMP  
More information on motif finding results: HOMER
| Description of Results
| Tips
  
Total target sequences = 6316  
Total background sequences = 32039  
\* - possible false positive  

|  |  |  |  |  |  |  |  |  |
| --- | --- | --- | --- | --- | --- | --- | --- | --- |
| Rank | Motif | P-value | log P-pvalue | % of Targets | % of Background | STD(Bg STD) | Best Match/Details | Motif File |
| 1 | T A C G C T G A T G A C T C G A A C T G A C T G C G T A C G T A T C A G A G C T | 1e-299 | -6.906e+02 | 61.19% | 38.11% | 362.7bp (421.8bp) | Etv2(ETS)/ES-ER71-ChIP-Seq(GSE59402)/Homer(0.961) More Information \| Similar Motifs Found | motif file (matrix) |
| 2 | T C G A A G C T C A T G G C T A T A C G C G A T G T A C C T G A A G C T G T A C | 1e-140 | -3.234e+02 | 36.35% | 22.31% | 369.1bp (437.2bp) | Fra1(bZIP)/BT549-Fra1-ChIP-Seq(GSE46166)/Homer(0.983) More Information \| Similar Motifs Found | motif file (matrix) |
| 3 | A G T C G A C T G T C A C T G A A G C T A C G T C T G A C T A G A G T C C T G A | 1e-118 | -2.731e+02 | 35.97% | 23.02% | 347.0bp (402.1bp) | dsc-1/MA0919.1/Jaspar(0.986) More Information \| Similar Motifs Found | motif file (matrix) |
| 4 | T C A G G A C T A G C T T G C A C T A G A G T C T C G A A T G C | 1e-49 | -1.146e+02 | 35.16% | 26.63% | 367.5bp (425.9bp) | prd/dmmpmm(Bigfoot)/fly(0.864) More Information \| Similar Motifs Found | motif file (matrix) |
| 5 | A G T C A C T G A G T C A G T C A C T G C G T A A C G T A C G T A C T G A G T C | 1e-19 | -4.386e+01 | 0.21% | 0.01% | 387.8bp (68.3bp) | PUCHI(AP2EREBP)/colamp-PUCHI-DAP-Seq(GSE60143)/Homer(0.665) More Information \| Similar Motifs Found | motif file (matrix) |
| 6 | A G T C G T A C A G T C A G T C A C G T A G C T T C A G A G T C | 1e-18 | -4.176e+01 | 20.23% | 16.04% | 375.0bp (429.5bp) | MSN4(MacIsaac)/Yeast(0.862) More Information \| Similar Motifs Found | motif file (matrix) |
| 7 | C T G A A C T G A C T G A C G T A C T G A G C T A C T G C G T A | 1e-16 | -3.692e+01 | 19.17% | 15.32% | 375.9bp (434.9bp) | TBX15/MA0803.1/Jaspar(0.999) More Information \| Similar Motifs Found | motif file (matrix) |
| 8 | T C G A A C G T T A C G C T G A C A G T T C A G G C A T T G A C G C T A C G A T | 1e-15 | -3.532e+01 | 4.04% | 2.35% | 376.8bp (463.7bp) | JUN/MA0488.1/Jaspar(0.950) More Information \| Similar Motifs Found | motif file (matrix) |
| 9 \* | T C A G G C A T G T A C C G T A A G T C C T A G A C G T A T C G | 1e-11 | -2.739e+01 | 53.13% | 48.72% | 396.7bp (435.0bp) | Usf2(bHLH)/C2C12-Usf2-ChIP-Seq(GSE36030)/Homer(0.919) More Information \| Similar Motifs Found | motif file (matrix) |
| 10 \* | A C T G A G T C A C T G A C G T C G T A C G T A A C T G A C T G C G A T A G T C | 1e-11 | -2.726e+01 | 0.27% | 0.03% | 185.0bp (277.6bp) | POPTR\_0002s00440g/MA0955.1/Jaspar(0.664) More Information \| Similar Motifs Found | motif file (matrix) |
| 11 \* | A G T C C G T A A C T G A G T C A C G T A C T G A C G T A G T C A C T G C G T A | 1e-10 | -2.383e+01 | 0.17% | 0.01% | 258.0bp (272.4bp) | PL0019.1\_hlh-1/Jaspar(0.842) More Information \| Similar Motifs Found | motif file (matrix) |
| 12 \* | A C G T A C T G C G T A C G T A A G T C A C T G A C G T C T G A A C T G A T C G | 1e-10 | -2.375e+01 | 0.38% | 0.07% | 360.7bp (323.2bp) | Lm\_0254(RRM)/Leishmania\_major-RNCMPT00254-PBM/HughesRNA(0.702) More Information \| Similar Motifs Found | motif file (matrix) |
| 13 \* | A C G T A C G T A G T C A G T C A C T G A G T C A C G T A G C T A C G T A C G T | 1e-10 | -2.318e+01 | 0.66% | 0.20% | 375.9bp (414.0bp) | RDR1/MA0360.1/Jaspar(0.781) More Information \| Similar Motifs Found | motif file (matrix) |
| 14 \* | C G T A A C G T C G T A C G T A A G T C A G T C A C G T A C T G C G T A A G T C | 1e-9 | -2.301e+01 | 0.35% | 0.06% | 322.4bp (376.1bp) | SIX1/MA1118.1/Jaspar(0.765) More Information \| Similar Motifs Found | motif file (matrix) |
| 15 \* | A G T C A C T G A T G C C G T A A C G T A C G T A G T C C G T A A G T C C G T A | 1e-5 | -1.323e+01 | 1.31% | 0.75% | 405.7bp (419.2bp) | Tb\_0251(RRM)/Trypanosoma\_brucei-RNCMPT00251-PBM/HughesRNA(0.812) More Information \| Similar Motifs Found | motif file (matrix) |
| 16 \* | T C G A A G T C C T G A A G C T G T C A A G T C T C G A A C G T A C T G A G C T | 1e-5 | -1.297e+01 | 10.85% | 9.14% | 411.2bp (445.9bp) | PH0082.1\_Irx2/Jaspar(0.759) More Information \| Similar Motifs Found | motif file (matrix) |
| 17 \* | G A C T C T A G G C T A A G T C C G T A C A T G T G C A A G T C | 1e-5 | -1.282e+01 | 46.03% | 43.18% | 389.6bp (443.6bp) | achi/MA0207.1/Jaspar(0.873) More Information \| Similar Motifs Found | motif file (matrix) |
| 18 \* | A G T C A G T C A C G T A C T G A C G T A G T C T G A C C G T A A C T G A C T G | 1e-5 | -1.156e+01 | 0.44% | 0.17% | 392.5bp (362.9bp) | FMR1(KH)/Homo\_sapiens-RNCMPT00016-PBM/HughesRNA(0.757) More Information \| Similar Motifs Found | motif file (matrix) |
