## Supplemental Table 1 for "The Establishment of Cell-Type Specific Gene Regulation in the Sea Urchin Embryo": Hpf40_CPCs_homerResults.html

homer\_motifs\_wbackground/Hpf40\_CPCs/ - Homer de novo Motif Results


### Homer *de novo* Motif Results (homer\_motifs\_wbackground/Hpf40\_CPCs/)

Known Motif Enrichment Results  
Gene Ontology Enrichment Results  
If Homer is having trouble matching a motif to a known motif, try copy/pasting the matrix file into
STAMP  
More information on motif finding results: HOMER
| Description of Results
| Tips
  
Total target sequences = 326  
Total background sequences = 8109  
\* - possible false positive  

|  |  |  |  |  |  |  |  |  |
| --- | --- | --- | --- | --- | --- | --- | --- | --- |
| Rank | Motif | P-value | log P-pvalue | % of Targets | % of Background | STD(Bg STD) | Best Match/Details | Motif File |
| 1 \* | C G T A A T C G T G A C C T G A A G T C G T C A C A G T A G T C C G T A C G T A | 1e-10 | -2.327e+01 | 9.20% | 2.20% | 450.9bp (408.9bp) | Tcf7(HMG)/GM12878-TCF7-ChIP-Seq(Encode)/Homer(0.742) More Information \| Similar Motifs Found | motif file (matrix) |
| 2 \* | C G T A C G A T A G T C A G C T C T A G A C T G A C T G A T C G C G T A G T C A | 1e-8 | -1.981e+01 | 8.59% | 2.25% | 250.5bp (413.9bp) | STOP1(C2H2)/colamp-STOP1-DAP-Seq(GSE60143)/Homer(0.717) More Information \| Similar Motifs Found | motif file (matrix) |
| 3 \* | A G T C G A C T A T G C A G T C T A G C G T C A C G T A A T G C A C G T A G C T | 1e-7 | -1.796e+01 | 5.83% | 1.17% | 379.2bp (459.7bp) | bHLH130(bHLH)/col-bHLH130-DAP-Seq(GSE60143)/Homer(0.634) More Information \| Similar Motifs Found | motif file (matrix) |
| 4 \* | A G T C G T C A A G T C C G A T A T C G G T A C G T C A T A G C A C T G A G T C | 1e-6 | -1.580e+01 | 7.36% | 2.09% | 195.3bp (398.3bp) | PB0091.1\_Zbtb3\_1/Jaspar(0.835) More Information \| Similar Motifs Found | motif file (matrix) |
| 5 \* | G T C A C G A T C A G T T A G C C G T A C G T A T A C G T C G A A G C T A T G C | 1e-6 | -1.545e+01 | 10.74% | 4.04% | 350.7bp (436.3bp) | MATR3(RRM)/Homo\_sapiens-RNCMPT00037-PBM/HughesRNA(0.701) More Information \| Similar Motifs Found | motif file (matrix) |
| 6 \* | A C T G C G T A G T A C C T A G A C G T C A T G C T G A A C T G A G C T A C T G | 1e-6 | -1.525e+01 | 5.52% | 1.25% | 213.6bp (511.7bp) | MSANTD3/MA1523.1/Jaspar(0.721) More Information \| Similar Motifs Found | motif file (matrix) |
| 7 \* | G T C A C G T A C G A T C A G T G A C T C T A G C G T A C T A G G C A T C G A T | 1e-6 | -1.400e+01 | 9.51% | 3.55% | 220.4bp (355.1bp) | lin-54/MA1450.1/Jaspar(0.674) More Information \| Similar Motifs Found | motif file (matrix) |
| 8 \* | T C G A G T A C C A G T A T G C C T G A G A C T T C A G A T G C C G T A G A C T | 1e-6 | -1.385e+01 | 21.78% | 12.24% | 286.5bp (425.4bp) | ASD-1(RRM)/Caenorhabditis\_elegans-RNCMPT00180-PBM/HughesRNA(0.798) More Information \| Similar Motifs Found | motif file (matrix) |
| 9 \* | C G T A A G T C C A G T G T C A A G T C G C T A G T A C G A T C A G C T T G A C | 1e-5 | -1.306e+01 | 6.13% | 1.77% | 330.6bp (411.3bp) | RBM28(RRM)/Homo\_sapiens-RNCMPT00049-PBM/HughesRNA(0.746) More Information \| Similar Motifs Found | motif file (matrix) |
| 10 \* | C G T A C G T A C T A G T C A G G T C A A G T C C G A T A C T G A C G T G T C A | 1e-5 | -1.208e+01 | 3.68% | 0.72% | 264.4bp (426.0bp) | RLR1(MacIsaac)/Yeast(0.694) More Information \| Similar Motifs Found | motif file (matrix) |
| 11 \* | C A T G C A T G G A T C A C G T T G C A T G C A A T G C T C G A A C G T A C G T | 1e-4 | -1.125e+01 | 14.11% | 7.27% | 332.9bp (421.1bp) | Gfi1b(Zf)/HPC7-Gfi1b-ChIP-Seq(GSE22178)/Homer(0.663) More Information \| Similar Motifs Found | motif file (matrix) |
| 12 \* | A G T C A C G T A C T G A C G T A G T C A C G T A C G T C G T A A C T G A C G T | 1e-4 | -9.764e+00 | 1.23% | 0.07% | 280.4bp (256.7bp) | CG2931(RRM)/Drosophila\_melanogaster-RNCMPT00147-PBM/HughesRNA(0.761) More Information \| Similar Motifs Found | motif file (matrix) |
| 13 \* | A G T C A G T C C G T A A C G T A C G T A T G C A G T C A G T C A C G T A C T G | 1e-4 | -9.405e+00 | 4.91% | 1.59% | 171.2bp (386.2bp) | PCBP1(KH)/Mus\_musculus-RNCMPT00239-PBM/HughesRNA(0.743) More Information \| Similar Motifs Found | motif file (matrix) |
| 14 \* | A G C T C T G A A C T G A G C T C G T A A T C G C A G T C T G A T C A G C G A T | 1e-4 | -9.251e+00 | 5.52% | 1.95% | 266.0bp (499.2bp) | MSI(RRM)/Drosophila\_melanogaster-RNCMPT00099-PBM/HughesRNA(0.723) More Information \| Similar Motifs Found | motif file (matrix) |
| 15 \* | C G T A A C G T C G T A A C T G C G T A A C G T G T A C A C G T | 1e-3 | -8.258e+00 | 13.50% | 7.77% | 337.8bp (385.3bp) | AGP1(GATA)/Nicotiana tabacum/AthaMap(0.881) More Information \| Similar Motifs Found | motif file (matrix) |
| 16 \* | C T A G T A C G C T G A C T A G C T A G G T A C C A G T T C A G T A G C A C T G | 1e-3 | -7.335e+00 | 19.63% | 13.15% | 315.1bp (540.5bp) | CRZ1(MacIsaac)/Yeast(0.810) More Information \| Similar Motifs Found | motif file (matrix) |
| 17 \* | A G T C A C T G C T A G A T C G A G T C C G T A G A C T A T G C A G T C C G T A | 1e-2 | -6.738e+00 | 3.99% | 1.47% | 210.8bp (414.6bp) | GCR1(MacIsaac)/Yeast(0.752) More Information \| Similar Motifs Found | motif file (matrix) |
| 18 \* | A C T G A G T C A T C G G T A C C T A G A T G C C A G T T G C A A C G T C G T A | 1e-2 | -6.159e+00 | 7.98% | 4.30% | 272.5bp (403.2bp) | DPL-1(E2F)/cElegans-Adult-ChIP-Seq(modEncode)/Homer(0.733) More Information \| Similar Motifs Found | motif file (matrix) |
| 19 \* | A C G T C G T A C G T A C G T A A G T C A C G T | 1e-1 | -4.376e+00 | 43.87% | 37.67% | 362.4bp (402.2bp) | br(var.3)/MA0012.1/Jaspar(0.826) More Information \| Similar Motifs Found | motif file (matrix) |
| 20 \* | A G T C A G C T C G T A A C T G A G T C A C T G A C T G A T C G | 1e-1 | -3.221e+00 | 1.53% | 0.57% | 101.2bp (352.1bp) | STP3/MA0396.1/Jaspar(0.719) More Information \| Similar Motifs Found | motif file (matrix) |
| 21 \* | A C G T C G T A A C G T A G T C A C G T A C T G A C T G A C T G A G T C A G T C | 1e-1 | -2.558e+00 | 0.31% | 0.04% | 128.1bp (127.9bp) | SKN7(MacIsaac)/Yeast(0.789) More Information \| Similar Motifs Found | motif file (matrix) |
| 22 \* | C G T A C G T A A G T C C G T A A C T G A C G T A C T G A G T C A G T C A G T C | 1e0 | -2.173e+00 | 0.31% | 0.05% | 176.9bp (149.7bp) | PB0099.1\_Zfp691\_1/Jaspar(0.735) More Information \| Similar Motifs Found | motif file (matrix) |
| 23 \* | A C T G A C G T A G T C C G T A A T G C A G T C | 1e0 | -1.016e+00 | 38.65% | 37.56% | 330.4bp (469.6bp) | ASHR1(ND)/col-ASHR1-DAP-Seq(GSE60143)/Homer(0.847) More Information \| Similar Motifs Found | motif file (matrix) |
