## Supplemental Table 1 for "The Establishment of Cell-Type Specific Gene Regulation in the Sea Urchin Embryo": Hpf40_EctoApical_homerResults.html

homer\_motifs\_wbackground/Hpf40\_EctoApical/ - Homer de novo Motif Results


### Homer *de novo* Motif Results (homer\_motifs\_wbackground/Hpf40\_EctoApical/)

Known Motif Enrichment Results  
Gene Ontology Enrichment Results  
If Homer is having trouble matching a motif to a known motif, try copy/pasting the matrix file into
STAMP  
More information on motif finding results: HOMER
| Description of Results
| Tips
  
Total target sequences = 740  
Total background sequences = 7540  
\* - possible false positive  

|  |  |  |  |  |  |  |  |  |
| --- | --- | --- | --- | --- | --- | --- | --- | --- |
| Rank | Motif | P-value | log P-pvalue | % of Targets | % of Background | STD(Bg STD) | Best Match/Details | Motif File |
| 1 | C T G A C G T A A C G T A G T C C T G A C G T A A C G T C G T A G T C A G C A T | 1e-34 | -8.039e+01 | 40.00% | 20.02% | 348.9bp (425.6bp) | ONECUT1/MA0679.2/Jaspar(0.945) More Information \| Similar Motifs Found | motif file (matrix) |
| 2 | C G T A A T G C C T G A C G T A A G C T C T G A C T A G T C A G | 1e-23 | -5.439e+01 | 51.76% | 33.60% | 358.1bp (402.3bp) | SOX15/MA1152.1/Jaspar(0.891) More Information \| Similar Motifs Found | motif file (matrix) |
| 3 | T C A G C T A G C T G A A C G T G C A T C T G A C T A G A G T C | 1e-18 | -4.179e+01 | 27.70% | 15.04% | 372.2bp (465.8bp) | CRX(Homeobox)/Retina-Crx-ChIP-Seq(GSE20012)/Homer(0.968) More Information \| Similar Motifs Found | motif file (matrix) |
| 4 | G T C A C T A G C G A T C A T G C T G A T G A C T G C A C A T G A T C G G A T C | 1e-16 | -3.753e+01 | 28.65% | 16.38% | 422.7bp (454.6bp) | PBX3/MA1114.1/Jaspar(0.873) More Information \| Similar Motifs Found | motif file (matrix) |
| 5 | T A G C T G A C G A T C A C G T A T C G G A C T A C G T T C A G C T A G C T A G | 1e-16 | -3.748e+01 | 28.11% | 15.96% | 421.4bp (448.7bp) | Unknown-ESC-element(?)/mES-Nanog-ChIP-Seq(GSE11724)/Homer(0.727) More Information \| Similar Motifs Found | motif file (matrix) |
| 6 | A C G T A G T C C G T A C G T A C T A G G C T A A C T G T C A G | 1e-15 | -3.673e+01 | 54.59% | 39.57% | 386.7bp (414.1bp) | ceh-22/MA0264.1/Jaspar(0.851) More Information \| Similar Motifs Found | motif file (matrix) |
| 7 \* | A C G T C A T G T C A G A G C T C A G T A C T G T A C G T G A C G T C A C A T G | 1e-11 | -2.730e+01 | 7.70% | 2.63% | 310.5bp (484.0bp) | PB0029.1\_Hic1\_1/Jaspar(0.710) More Information \| Similar Motifs Found | motif file (matrix) |
| 8 \* | C A T G C T A G A C G T A C T G C A T G A C G T A C T G C A T G A C T G A C G T | 1e-11 | -2.672e+01 | 6.76% | 2.14% | 325.7bp (524.8bp) | Run/dmmpmm(Papatsenko)/fly(0.772) More Information \| Similar Motifs Found | motif file (matrix) |
| 9 \* | A C T G T G A C C A G T A G T C C G T A C T G A A C G T A G C T C G T A C T A G | 1e-11 | -2.597e+01 | 6.08% | 1.82% | 402.1bp (380.5bp) | YHP1/MA0426.1/Jaspar(0.788) More Information \| Similar Motifs Found | motif file (matrix) |
| 10 \* | C G A T A C G T A T C G G T C A T C A G C G A T A C G T C A T G C G T A T C G A | 1e-11 | -2.578e+01 | 10.81% | 4.67% | 396.1bp (476.8bp) | NKX2-2/MA1645.1/Jaspar(0.714) More Information \| Similar Motifs Found | motif file (matrix) |
| 11 \* | T A G C C G T A G C T A T G A C C T A G G C T A C A G T T A C G A C T G C T G A | 1e-11 | -2.573e+01 | 13.24% | 6.34% | 384.8bp (431.7bp) | CG2950(KH)/Drosophila\_melanogaster-RNCMPT00007-PBM/HughesRNA(0.684) More Information \| Similar Motifs Found | motif file (matrix) |
| 12 \* | A C T G C G T A A C T G A C T G C G A T A T C G A C G T C T G A C T G A G T A C | 1e-9 | -2.195e+01 | 2.84% | 0.50% | 336.5bp (321.2bp) | TBX15/MA0803.1/Jaspar(0.799) More Information \| Similar Motifs Found | motif file (matrix) |
| 13 \* | A C G T A G T C A C G T C T G A C G T A A C G T A C T G C G T A C T G A A C T G | 1e-9 | -2.102e+01 | 2.97% | 0.58% | 390.6bp (370.0bp) | zen/MA0256.1/Jaspar(0.780) More Information \| Similar Motifs Found | motif file (matrix) |
| 14 \* | A G T C G T C A C G A T A G T C T G A C G A T C T A C G A G T C C G T A C T A G | 1e-8 | -1.985e+01 | 4.59% | 1.38% | 296.0bp (390.4bp) | UGA3/MA0410.1/Jaspar(0.748) More Information \| Similar Motifs Found | motif file (matrix) |
| 15 \* | A G T C C G T A C G T A A C T G A C G T A C G T A C G T A G T C A G T C A C G T | 1e-6 | -1.544e+01 | 0.95% | 0.06% | 218.3bp (353.3bp) | bHLH130(bHLH)/col-bHLH130-DAP-Seq(GSE60143)/Homer(0.706) More Information \| Similar Motifs Found | motif file (matrix) |
| 16 \* | C G T A A C T G A G T C C G T A A C G T A G T C A G T C A C G T | 1e-6 | -1.408e+01 | 3.51% | 1.15% | 328.3bp (461.3bp) | G3BP2(RRM)/Homo\_sapiens-RNCMPT00021-PBM/HughesRNA(0.741) More Information \| Similar Motifs Found | motif file (matrix) |
| 17 \* | C G T A C G T A C G T A A G T C A C G T C G T A A G T C C G T A A C T G A C T G | 1e-5 | -1.277e+01 | 0.95% | 0.09% | 254.8bp (228.7bp) | Rbm42(RRM)/Xenopus\_tropicalis-RNCMPT00282-PBM/HughesRNA(0.739) More Information \| Similar Motifs Found | motif file (matrix) |
| 18 \* | A C T G C G T A A C T G A C G T A C T G A T G C C G T A A C G T | 1e-5 | -1.274e+01 | 11.89% | 7.19% | 457.1bp (450.0bp) | z/dmmpmm(Down)/fly(0.752) More Information \| Similar Motifs Found | motif file (matrix) |
| 19 \* | C G T A A C T G A C T G C G T A A G T C G T C A A C T G C G T A A T C G C G A T | 1e-4 | -9.290e+00 | 3.24% | 1.35% | 415.1bp (572.9bp) | CNOT4(RRM)/Homo\_sapiens-RNCMPT00156-PBM/HughesRNA(0.801) More Information \| Similar Motifs Found | motif file (matrix) |
