## Supplemental Table 1 for "The Establishment of Cell-Type Specific Gene Regulation in the Sea Urchin Embryo": Hpf40_EctoArboral_homerResults.html

homer\_motifs\_wbackground/Hpf40\_EctoArboral/ - Homer de novo Motif Results


### Homer *de novo* Motif Results (homer\_motifs\_wbackground/Hpf40\_EctoArboral/)

Known Motif Enrichment Results  
Gene Ontology Enrichment Results  
If Homer is having trouble matching a motif to a known motif, try copy/pasting the matrix file into
STAMP  
More information on motif finding results: HOMER
| Description of Results
| Tips
  
Total target sequences = 971  
Total background sequences = 7500  
\* - possible false positive  

|  |  |  |  |  |  |  |  |  |
| --- | --- | --- | --- | --- | --- | --- | --- | --- |
| Rank | Motif | P-value | log P-pvalue | % of Targets | % of Background | STD(Bg STD) | Best Match/Details | Motif File |
| 1 | G T A C A T G C A C G T A C G T A C G T A G T C C G T A C G A T | 1e-43 | -9.946e+01 | 63.71% | 41.56% | 442.2bp (408.2bp) | LEF1(HMG)/H1-LEF1-ChIP-Seq(GSE64758)/Homer(0.800) More Information \| Similar Motifs Found | motif file (matrix) |
| 2 | A G T C G C A T G C T A A C G T C G A T T A G C C T A G A T G C | 1e-37 | -8.557e+01 | 35.88% | 18.41% | 421.8bp (415.6bp) | Tb\_0251(RRM)/Trypanosoma\_brucei-RNCMPT00251-PBM/HughesRNA(0.748) More Information \| Similar Motifs Found | motif file (matrix) |
| 3 | A G C T C G T A G A C T C G T A C T G A C T G A C T A G C A T G | 1e-35 | -8.276e+01 | 55.57% | 35.65% | 439.2bp (426.2bp) | POL012.1\_TATA-Box/Jaspar(0.778) More Information \| Similar Motifs Found | motif file (matrix) |
| 4 | C G T A C G T A A C G T A G T C A G C T C A T G C G T A A C T G A C T G A C T G | 1e-35 | -8.202e+01 | 3.81% | 0.18% | 317.0bp (772.3bp) | MET31/Literature(Harbison)/Yeast(0.657) More Information \| Similar Motifs Found | motif file (matrix) |
| 5 | A C G T A C G T A T G C C G T A C G T A A C G T C T A G A C T G | 1e-32 | -7.379e+01 | 34.33% | 18.28% | 441.4bp (388.0bp) | EGL-5(Homeobox)/cElegans-L3-EGL5-ChIP-Seq(modEncode)/Homer(0.780) More Information \| Similar Motifs Found | motif file (matrix) |
| 6 | C T A G T A G C C G T A G C A T A C G T T G A C A T G C T C G A | 1e-29 | -6.892e+01 | 66.91% | 48.63% | 461.0bp (411.5bp) | TEC1/MA0406.1/Jaspar(0.833) More Information \| Similar Motifs Found | motif file (matrix) |
| 7 | G A T C A G C T A T G C G C A T C G T A A T C G A G C T A C T G | 1e-29 | -6.837e+01 | 72.06% | 54.17% | 434.5bp (407.3bp) | tin/dmmpmm(SeSiMCMC)/fly(0.785) More Information \| Similar Motifs Found | motif file (matrix) |
| 8 | A C T G A C T G A C T G A G T C C G T A A C T G A C G T C T A G A G T C C G T A | 1e-26 | -6.140e+01 | 2.58% | 0.10% | 379.4bp (158.3bp) | PB0091.1\_Zbtb3\_1/Jaspar(0.704) More Information \| Similar Motifs Found | motif file (matrix) |
| 9 | A C G T A C T G A G C T C A T G A C T G A C T G A T G C C T A G A C G T A G T C | 1e-25 | -5.871e+01 | 3.09% | 0.20% | 342.0bp (628.9bp) | Foxn1/MA1684.1/Jaspar(0.696) More Information \| Similar Motifs Found | motif file (matrix) |
| 10 | A C T G A C T G C G T A A C T G A C G T A C T G A C T G A C G T A C G T A C T G | 1e-23 | -5.515e+01 | 2.16% | 0.07% | 236.6bp (372.1bp) | MYB94(MYB)/col-MYB94-DAP-Seq(GSE60143)/Homer(0.704) More Information \| Similar Motifs Found | motif file (matrix) |
| 11 | A C T G A C T G G T C A A G C T A C G T C T G A A C T G C G T A A C T G A C G T | 1e-20 | -4.781e+01 | 3.71% | 0.45% | 385.1bp (408.6bp) | oc/dmmpmm(Bergman)/fly(0.803) More Information \| Similar Motifs Found | motif file (matrix) |
| 12 | C T G A C G T A C G T A A C T G C G T A A C T G A G T C C G T A A G T C C G T A | 1e-20 | -4.736e+01 | 4.12% | 0.58% | 450.9bp (416.6bp) | NCU08034(RRM)/Neurospora\_crassa-RNCMPT00209-PBM/HughesRNA(0.710) More Information \| Similar Motifs Found | motif file (matrix) |
| 13 | A C G T A G T C C T A G C G T A A C T G C T A G A G T C C G T A A T C G A G T C | 1e-20 | -4.686e+01 | 3.71% | 0.46% | 339.0bp (328.8bp) | XBP1(MacIsaac)/Yeast(0.716) More Information \| Similar Motifs Found | motif file (matrix) |
| 14 | C G T A C G T A A G C T C G T A C G T A A G T C C G T A C T G A | 1e-20 | -4.647e+01 | 86.80% | 74.60% | 456.3bp (425.2bp) | PB0119.1\_Foxa2\_2/Jaspar(0.912) More Information \| Similar Motifs Found | motif file (matrix) |
| 15 | A C G T A G T C A C G T A C T G A C T G A C G T A G T C C G T A C G T A A C T G | 1e-19 | -4.405e+01 | 2.27% | 0.14% | 355.4bp (583.5bp) | WRKY63/MA1092.1/Jaspar(0.762) More Information \| Similar Motifs Found | motif file (matrix) |
| 16 | C T A G A C T G C G T A A C T G A C G T A C T G A G T C A C G T A C T G C G T A | 1e-17 | -4.006e+01 | 1.13% | 0.02% | 384.7bp (0.0bp) | MafA(bZIP)/Islet-MafA-ChIP-Seq(GSE30298)/Homer(0.708) More Information \| Similar Motifs Found | motif file (matrix) |
| 17 | C G T A A C T G A G C T A C T G C A T G C T A G T G C A A C T G A C G T G C A T | 1e-16 | -3.799e+01 | 4.54% | 0.94% | 340.3bp (420.5bp) | STZ/MA1372.1/Jaspar(0.740) More Information \| Similar Motifs Found | motif file (matrix) |
| 18 | A C T G A C G T C G T A A C G T A G T C C G T A A C T G A G C T | 1e-15 | -3.577e+01 | 68.76% | 56.03% | 466.0bp (439.4bp) | so/MA0246.1/Jaspar(0.873) More Information \| Similar Motifs Found | motif file (matrix) |
| 19 | A G T C A G T C A G T C C G T A A C G T A G T C A G T C A C T G A C G T C G T A | 1e-15 | -3.517e+01 | 1.34% | 0.04% | 352.0bp (316.2bp) | SPL3/MA0577.2/Jaspar(0.696) More Information \| Similar Motifs Found | motif file (matrix) |
| 20 | A C G T A C T G A C G T A G T C A C T G A T G C C G T A C G T A A C G T A C T G | 1e-15 | -3.513e+01 | 69.38% | 56.81% | 484.5bp (411.1bp) | CEBP:AP1(bZIP)/ThioMac-CEBPb-ChIP-Seq(GSE21512)/Homer(0.730) More Information \| Similar Motifs Found | motif file (matrix) |
| 21 | A G T C A C G T A C G T A C T G A G T C A C G T C T G A A C G T C G T A A G T C | 1e-15 | -3.473e+01 | 1.86% | 0.13% | 387.8bp (337.1bp) | MBNL1(Znf)/Homo\_sapiens-RNCMPT00038-PBM/HughesRNA(0.645) More Information \| Similar Motifs Found | motif file (matrix) |
| 22 | A G T C A C G T C T A G A C T G A C T G A C T G A G T C A C T G | 1e-14 | -3.329e+01 | 69.69% | 57.51% | 482.0bp (402.9bp) | PB0143.1\_Klf7\_2/Jaspar(0.744) More Information \| Similar Motifs Found | motif file (matrix) |
| 23 | C G T A A G T C A C T G A C T G A C G T G T A C C G A T A G T C | 1e-14 | -3.268e+01 | 35.98% | 24.81% | 448.1bp (416.8bp) | MYB52/MA1171.1/Jaspar(0.707) More Information \| Similar Motifs Found | motif file (matrix) |
| 24 | C G T A A G T C C G T A C G T A A C G T C G T A A G T C A G T C | 1e-12 | -2.952e+01 | 65.05% | 53.44% | 428.4bp (414.5bp) | PB0183.1\_Sry\_2/Jaspar(0.740) More Information \| Similar Motifs Found | motif file (matrix) |
| 25 | A G T C A C G T A G T C A C T G C G T A A C T G A C G T A C G T A C G T A C G T | 1e-12 | -2.881e+01 | 1.03% | 0.04% | 494.8bp (46.4bp) | XBP1(MacIsaac)/Yeast(0.705) More Information \| Similar Motifs Found | motif file (matrix) |
