## Supplemental Table 1 for "The Establishment of Cell-Type Specific Gene Regulation in the Sea Urchin Embryo": Hpf40_EctoCiliaryBand_homerResults.html

homer\_motifs\_wbackground/Hpf40\_EctoCiliaryBand/ - Homer de novo Motif Results


### Homer *de novo* Motif Results (homer\_motifs\_wbackground/Hpf40\_EctoCiliaryBand/)

Known Motif Enrichment Results  
Gene Ontology Enrichment Results  
If Homer is having trouble matching a motif to a known motif, try copy/pasting the matrix file into
STAMP  
More information on motif finding results: HOMER
| Description of Results
| Tips
  
Total target sequences = 692  
Total background sequences = 7775  
\* - possible false positive  

|  |  |  |  |  |  |  |  |  |
| --- | --- | --- | --- | --- | --- | --- | --- | --- |
| Rank | Motif | P-value | log P-pvalue | % of Targets | % of Background | STD(Bg STD) | Best Match/Details | Motif File |
| 1 | C T G A C G T A A C G T A G T C C T G A C G T A A C G T C T G A | 1e-108 | -2.489e+02 | 64.02% | 24.19% | 333.1bp (414.3bp) | ONECUT1/MA0679.2/Jaspar(0.939) More Information \| Similar Motifs Found | motif file (matrix) |
| 2 | G T C A A T G C G A T C A C T G G T C A G C T A G T A C A C T G C G T A C T G A | 1e-12 | -2.825e+01 | 4.91% | 1.07% | 309.3bp (397.6bp) | Lm\_0254(RRM)/Leishmania\_major-RNCMPT00254-PBM/HughesRNA(0.732) More Information \| Similar Motifs Found | motif file (matrix) |
| 3 \* | C G T A A C G T A G T C A C T G C G T A G T A C | 1e-11 | -2.692e+01 | 55.20% | 42.02% | 352.5bp (456.7bp) | ceh-48/MA0921.1/Jaspar(0.813) More Information \| Similar Motifs Found | motif file (matrix) |
| 4 \* | C T A G C T A G C A T G T G C A A C T G C G T A A G C T C G T A A T G C A G T C | 1e-10 | -2.498e+01 | 3.76% | 0.72% | 278.6bp (444.7bp) | ARF10/MA1685.1/Jaspar(0.718) More Information \| Similar Motifs Found | motif file (matrix) |
| 5 \* | G T C A A G T C C T A G A T C G G T C A A C G T A C T G G T C A C T A G C G A T | 1e-10 | -2.329e+01 | 4.19% | 0.95% | 326.4bp (578.5bp) | RGT1/Literature(Harbison)/Yeast(0.689) More Information \| Similar Motifs Found | motif file (matrix) |
| 6 \* | A C T G A T C G T C G A T C G A A T G C G C T A A T C G T A G C A G C T C T A G | 1e-9 | -2.269e+01 | 5.78% | 1.77% | 365.8bp (443.5bp) | Ptf1a(bHLH)/Panc1-Ptf1a-ChIP-Seq(GSE47459)/Homer(0.857) More Information \| Similar Motifs Found | motif file (matrix) |
| 7 \* | A G C T C G T A A C G T A G T C A C G T A C G T A C T G A C G T A C G T A C T G | 1e-9 | -2.198e+01 | 3.18% | 0.59% | 385.8bp (304.8bp) | Aef1/dmmpmm(Bergman)/fly(0.801) More Information \| Similar Motifs Found | motif file (matrix) |
| 8 \* | C G T A T C A G A G C T G T A C A G T C C T G A C T G A T A G C G A C T T G A C | 1e-9 | -2.074e+01 | 12.28% | 6.08% | 352.9bp (418.9bp) | PB0114.1\_Egr1\_2/Jaspar(0.739) More Information \| Similar Motifs Found | motif file (matrix) |
| 9 \* | C G T A C G A T G T C A C T G A A C T G C G T A G T A C A T G C A G C T A T C G | 1e-8 | -1.910e+01 | 3.76% | 0.94% | 299.5bp (378.1bp) | At5g22890(C2H2)/col-At5g22890-DAP-Seq(GSE60143)/Homer(0.752) More Information \| Similar Motifs Found | motif file (matrix) |
| 10 \* | A G T C A C T G C G T A C G T A C T A G G T C A C G T A A C G T C T A G A C T G | 1e-8 | -1.887e+01 | 2.02% | 0.26% | 407.3bp (319.6bp) | SOX14/MA1562.1/Jaspar(0.767) More Information \| Similar Motifs Found | motif file (matrix) |
| 11 \* | A T C G C A T G A T G C T G A C A T G C T A C G A C T G A C G T T C A G G A C T | 1e-8 | -1.858e+01 | 4.48% | 1.33% | 443.9bp (352.4bp) | AFT2/MA0270.1/Jaspar(0.674) More Information \| Similar Motifs Found | motif file (matrix) |
| 12 \* | A C G T A C T G C G T A C G T A C G T A A C T G A C G T A C T G | 1e-7 | -1.722e+01 | 12.86% | 7.02% | 370.9bp (431.4bp) | AT1G69570(C2C2dof)/col-AT1G69570-DAP-Seq(GSE60143)/Homer(0.800) More Information \| Similar Motifs Found | motif file (matrix) |
| 13 \* | A T C G C G A T G C A T C G T A C T A G C A T G T C G A G C T A A T C G A T C G | 1e-7 | -1.629e+01 | 10.12% | 5.14% | 428.9bp (438.6bp) | MYB116(MYB)/colamp-MYB116-DAP-Seq(GSE60143)/Homer(0.729) More Information \| Similar Motifs Found | motif file (matrix) |
| 14 \* | A C G T A C T G A G T C C G T A A C T G A C T G C G T A A C T G A G T C A C G T | 1e-6 | -1.475e+01 | 0.87% | 0.05% | 276.7bp (282.6bp) | SOK2/SOK2\_BUT14/4-SUT1(Harbison)/Yeast(0.734) More Information \| Similar Motifs Found | motif file (matrix) |
| 15 \* | C G T A C G T A A G T C C G T A A C G T C G T A A C G T C G T A A C T G C G T A | 1e-6 | -1.391e+01 | 1.30% | 0.15% | 426.4bp (541.4bp) | MOT2/MA0379.1/Jaspar(0.706) More Information \| Similar Motifs Found | motif file (matrix) |
| 16 \* | G T C A C A T G A C T G C G T A T A C G A T C G G C T A C A T G A T G C T C A G | 1e-5 | -1.221e+01 | 10.12% | 5.77% | 264.7bp (545.6bp) | SRSF1(RRM)/Homo\_sapiens-RNCMPT00163-PBM/HughesRNA(0.724) More Information \| Similar Motifs Found | motif file (matrix) |
| 17 \* | C G T A A G T C A G T C A G T C A C G T C G T A | 1e-4 | -1.071e+01 | 28.90% | 22.20% | 367.9bp (424.1bp) | MUB(KH)/Drosophila\_melanogaster-RNCMPT00137-PBM/HughesRNA(0.927) More Information \| Similar Motifs Found | motif file (matrix) |
