## Supplemental Table 1 for "The Establishment of Cell-Type Specific Gene Regulation in the Sea Urchin Embryo": Hpf40_EctoOral_homerResults.html

homer\_motifs\_wbackground/Hpf40\_EctoOral/ - Homer de novo Motif Results


### Homer *de novo* Motif Results (homer\_motifs\_wbackground/Hpf40\_EctoOral/)

Known Motif Enrichment Results  
Gene Ontology Enrichment Results  
If Homer is having trouble matching a motif to a known motif, try copy/pasting the matrix file into
STAMP  
More information on motif finding results: HOMER
| Description of Results
| Tips
  
Total target sequences = 333  
Total background sequences = 8179  
\* - possible false positive  

|  |  |  |  |  |  |  |  |  |
| --- | --- | --- | --- | --- | --- | --- | --- | --- |
| Rank | Motif | P-value | log P-pvalue | % of Targets | % of Background | STD(Bg STD) | Best Match/Details | Motif File |
| 1 | C T G A C T G A A C G T A G T C C T A G C T G A C G A T T C A G | 1e-24 | -5.736e+01 | 66.67% | 38.32% | 364.8bp (420.6bp) | CUX1/MA0754.1/Jaspar(0.926) More Information \| Similar Motifs Found | motif file (matrix) |
| 2 \* | C A G T T A C G G C A T C A T G T G C A A C G T T C A G G C T A T C A G C A T G | 1e-10 | -2.352e+01 | 13.21% | 4.31% | 392.2bp (418.5bp) | POU6F1(var.2)/MA1549.1/Jaspar(0.693) More Information \| Similar Motifs Found | motif file (matrix) |
| 3 \* | A C G T G A T C A T C G A T C G A G T C C T G A A G T C A C T G A G T C A C T G | 1e-9 | -2.296e+01 | 3.30% | 0.20% | 152.7bp (348.3bp) | h/dmmpmm(Bergman)/fly(0.818) More Information \| Similar Motifs Found | motif file (matrix) |
| 4 \* | C T G A G T C A C G T A A T G C T C G A T C A G C T G A T G C A G T C A A T C G | 1e-9 | -2.216e+01 | 24.32% | 11.88% | 396.6bp (411.2bp) | Stat2/MA1623.1/Jaspar(0.909) More Information \| Similar Motifs Found | motif file (matrix) |
| 5 \* | A G T C C G T A C G A T A C G T A C T G C G T A A T G C A C T G A C T G T G A C | 1e-9 | -2.117e+01 | 5.11% | 0.73% | 180.4bp (316.2bp) | ERF11/MA1001.2/Jaspar(0.715) More Information \| Similar Motifs Found | motif file (matrix) |
| 6 \* | C G A T C A T G C A T G A G T C G A T C C A T G G C T A C G T A C T A G A G T C | 1e-8 | -2.043e+01 | 9.01% | 2.44% | 237.5bp (387.0bp) | pho/MA1460.1/Jaspar(0.698) More Information \| Similar Motifs Found | motif file (matrix) |
| 7 \* | A T G C T G A C G T A C A C G T A C T G T C G A G T A C G T C A | 1e-8 | -2.040e+01 | 41.44% | 26.28% | 453.7bp (442.2bp) | vis/MA0252.1/Jaspar(0.816) More Information \| Similar Motifs Found | motif file (matrix) |
| 8 \* | A C G T A G T C A G T C C T A G C G T A A G T C A G T C A G T C G T A C A C G T | 1e-8 | -1.849e+01 | 3.00% | 0.23% | 263.7bp (273.6bp) | btd/MA0443.1/Jaspar(0.717) More Information \| Similar Motifs Found | motif file (matrix) |
| 9 \* | C A G T T G A C A C T G A G T C C G T A G A C T C A T G G T C A G T A C G T A C | 1e-6 | -1.471e+01 | 9.61% | 3.53% | 372.0bp (447.1bp) | USF1/MA0093.3/Jaspar(0.743) More Information \| Similar Motifs Found | motif file (matrix) |
| 10 \* | A C G T C G T A C G T A A C G T A G T C A G T C | 1e-2 | -6.315e+00 | 63.66% | 55.65% | 374.8bp (434.2bp) | bcd/MA0212.1/Jaspar(0.977) More Information \| Similar Motifs Found | motif file (matrix) |
| 11 \* | A G T C A G T C A C G T A C G T C G T A A G T C A G T C A G T C A G T C A C G T | 1e-2 | -4.796e+00 | 0.90% | 0.13% | 267.8bp (453.5bp) | PCBP1(KH)/Homo\_sapiens-RNCMPT00186-PBM/HughesRNA(0.744) More Information \| Similar Motifs Found | motif file (matrix) |
