## Supplemental Table 1 for "The Establishment of Cell-Type Specific Gene Regulation in the Sea Urchin Embryo": Hpf40_EctoStomodeum_homerResults.html

homer\_motifs\_wbackground/Hpf40\_EctoStomodeum/ - Homer de novo Motif Results


### Homer *de novo* Motif Results (homer\_motifs\_wbackground/Hpf40\_EctoStomodeum/)

Known Motif Enrichment Results  
Gene Ontology Enrichment Results  
If Homer is having trouble matching a motif to a known motif, try copy/pasting the matrix file into
STAMP  
More information on motif finding results: HOMER
| Description of Results
| Tips
  
Total target sequences = 362  
Total background sequences = 8196  
\* - possible false positive  

|  |  |  |  |  |  |  |  |  |
| --- | --- | --- | --- | --- | --- | --- | --- | --- |
| Rank | Motif | P-value | log P-pvalue | % of Targets | % of Background | STD(Bg STD) | Best Match/Details | Motif File |
| 1 \* | C T G A G T C A T G A C A C T G C T G A G C A T A C G T T A G C C T A G A G C T | 1e-8 | -1.915e+01 | 10.50% | 3.58% | 419.8bp (377.9bp) | ARR2/MA0949.1/Jaspar(0.704) More Information \| Similar Motifs Found | motif file (matrix) |
| 2 \* | A G T C G T A C G T A C A C G T A C T G C G T A G T C A A T G C C T G A A G C T | 1e-8 | -1.895e+01 | 6.63% | 1.58% | 645.6bp (304.0bp) | USV1/MA0413.1/Jaspar(0.739) More Information \| Similar Motifs Found | motif file (matrix) |
| 3 \* | A G C T G C A T C T G A A C G T A C G T A C T G A C T G A G T C G T C A G T C A | 1e-7 | -1.748e+01 | 8.01% | 2.41% | 287.6bp (365.2bp) | NFIA/MA0670.1/Jaspar(0.784) More Information \| Similar Motifs Found | motif file (matrix) |
| 4 \* | A T C G A T G C C T A G C G A T A C G T A C G T C G T A G T C A C T A G C A G T | 1e-7 | -1.730e+01 | 11.60% | 4.52% | 457.9bp (369.9bp) | B-H1/MA0168.1/Jaspar(0.782) More Information \| Similar Motifs Found | motif file (matrix) |
| 5 \* | A G T C C T G A A T G C C G T A T C A G T C A G C G A T A G T C G T C A G T A C | 1e-7 | -1.690e+01 | 4.70% | 0.90% | 257.5bp (430.6bp) | PB0053.1\_Rara\_1/Jaspar(0.788) More Information \| Similar Motifs Found | motif file (matrix) |
| 6 \* | A T C G T G A C C A T G C G A T G T C A G T A C A C G T G T C A T C G A A G C T | 1e-6 | -1.574e+01 | 4.97% | 1.10% | 245.3bp (336.0bp) | CAD1(MacIsaac)/Yeast(0.840) More Information \| Similar Motifs Found | motif file (matrix) |
| 7 \* | T A C G C G A T A T G C C G T A A C G T C T G A A C G T C G A T T G A C A T C G | 1e-6 | -1.566e+01 | 6.91% | 2.03% | 225.2bp (462.0bp) | KAN1/MA1027.1/Jaspar(0.696) More Information \| Similar Motifs Found | motif file (matrix) |
| 8 \* | A G T C T A G C G T C A C G A T A C G T G A T C A C G T A C T G G T C A C T G A | 1e-6 | -1.555e+01 | 7.18% | 2.19% | 412.0bp (395.8bp) | YML081W(MacIsaac)/Yeast(0.832) More Information \| Similar Motifs Found | motif file (matrix) |
| 9 \* | C G T A T A G C A C T G A G C T C G T A G T C A C G T A A C T G C G A T A C T G | 1e-6 | -1.530e+01 | 5.25% | 1.25% | 356.8bp (390.7bp) | AT1G69570(C2C2dof)/col-AT1G69570-DAP-Seq(GSE60143)/Homer(0.710) More Information \| Similar Motifs Found | motif file (matrix) |
| 10 \* | C T G A A G T C A C G T A G T C G T A C G T A C G T C A G T A C G T C A A G T C | 1e-6 | -1.434e+01 | 5.80% | 1.60% | 653.6bp (492.9bp) | WT1(Zf)/Kidney-WT1-ChIP-Seq(GSE90016)/Homer(0.866) More Information \| Similar Motifs Found | motif file (matrix) |
| 11 \* | A T C G T A G C A C G T T A C G A G C T A G C T A G C T G A C T A T C G A T C G | 1e-5 | -1.163e+01 | 1.93% | 0.20% | 354.0bp (522.2bp) | ZNF341/MA1655.1/Jaspar(0.701) More Information \| Similar Motifs Found | motif file (matrix) |
| 12 \* | A G T C A C G T A G T C A G T C A T C G C T A G C G T A C T A G A C G T A C G T | 1e-4 | -1.109e+01 | 3.04% | 0.61% | 362.8bp (416.5bp) | GSM1/MA0308.1/Jaspar(0.857) More Information \| Similar Motifs Found | motif file (matrix) |
| 13 \* | A G T C A C G T A C G T C G T A A C G T A C T G A G T C C G T A A C G T A C T G | 1e-4 | -1.094e+01 | 1.38% | 0.09% | 75.0bp (257.7bp) | unc-86/MA0926.1/Jaspar(0.823) More Information \| Similar Motifs Found | motif file (matrix) |
| 14 \* | C T G A C T A G T C G A A C G T C A T G T C A G C T A G G T A C C G A T C T A G | 1e-4 | -1.068e+01 | 14.36% | 7.90% | 429.9bp (392.8bp) | PCF/Arabidopsis-Promoters/Homer(0.754) More Information \| Similar Motifs Found | motif file (matrix) |
| 15 \* | A C G T C G A T A C T G T A C G A T C G C T G A A G T C A C G T A C G T C G A T | 1e-4 | -9.941e+00 | 9.94% | 4.87% | 376.4bp (424.0bp) | HNF4A/MA0114.4/Jaspar(0.741) More Information \| Similar Motifs Found | motif file (matrix) |
| 16 \* | A C T G A C G T A G T C A C G T A T C G A C T G A C G T A C T G A C G T A C G T | 1e-4 | -9.759e+00 | 1.38% | 0.11% | 306.4bp (303.1bp) | ZNF317/MA1593.1/Jaspar(0.754) More Information \| Similar Motifs Found | motif file (matrix) |
| 17 \* | C G T A A C T G A C T G A C T G A C T G C G T A A C T G C G T A | 1e-4 | -9.493e+00 | 10.77% | 5.58% | 305.0bp (426.3bp) | RGM1/MA0366.1/Jaspar(0.754) More Information \| Similar Motifs Found | motif file (matrix) |
| 18 \* | A C G T C A G T C G A T A C G T A C G T C G T A C T G A C T G A C T G A G T C A | 1e-3 | -9.067e+00 | 11.33% | 6.10% | 328.0bp (430.6bp) | Unknown6/Drosophila-Promoters/Homer(0.792) More Information \| Similar Motifs Found | motif file (matrix) |
| 19 \* | G T C A A G T C C T A G G A C T A C T G C A T G G T A C C G A T | 1e-3 | -7.578e+00 | 23.76% | 16.88% | 327.5bp (428.5bp) | ABI5(bZIP)/col-ABI5-DAP-Seq(GSE60143)/Homer(0.884) More Information \| Similar Motifs Found | motif file (matrix) |
| 20 \* | T A C G G C T A T G C A C T A G C T G A G C T A A G C T T G C A G C T A A C T G | 1e-2 | -6.703e+00 | 20.72% | 14.70% | 421.1bp (417.4bp) | KAN2(G2like)/colamp-KAN2-DAP-Seq(GSE60143)/Homer(0.761) More Information \| Similar Motifs Found | motif file (matrix) |
| 21 \* | A G T C A C G T A C G T A C T G A C T G C G T A C G T A A G T C | 1e-2 | -5.533e+00 | 3.59% | 1.52% | 389.1bp (413.7bp) | Stat5a/MA1624.1/Jaspar(0.778) More Information \| Similar Motifs Found | motif file (matrix) |
| 22 \* | C T A G G T A C G C T A A G T C C G T A A T C G C T G A A G T C C T A G A G T C | 1e-2 | -5.189e+00 | 7.18% | 4.18% | 448.9bp (449.8bp) | PB0130.1\_Gm397\_2/Jaspar(0.785) More Information \| Similar Motifs Found | motif file (matrix) |
| 23 \* | A G T C C G T A C G T A A C G T A C T G A C T G A G T C A G T C C G T A C G T A | 1e-1 | -4.326e+00 | 0.83% | 0.14% | 214.9bp (370.1bp) | pho/dmmpmm(Bergman)/fly(0.775) More Information \| Similar Motifs Found | motif file (matrix) |
| 24 \* | C G T A A C T G A G T C C G T A A C T G A C T G | 1e-1 | -3.620e+00 | 45.86% | 40.71% | 403.2bp (414.6bp) | Zic1::Zic2/MA1628.1/Jaspar(0.890) More Information \| Similar Motifs Found | motif file (matrix) |
| 25 \* | A C T G C G T A C T G A A C G T A G C T A T G C C T A G A G T C A C T G C G T A | 1e-1 | -3.361e+00 | 0.83% | 0.20% | 126.6bp (399.9bp) | PB0136.1\_IRC900814\_2/Jaspar(0.626) More Information \| Similar Motifs Found | motif file (matrix) |
| 26 \* | A G T C C G T A A C G T A C T G A C G T A C T G | 1e-1 | -3.001e+00 | 42.54% | 38.18% | 365.7bp (395.0bp) | MXI1/MA1108.2/Jaspar(0.995) More Information \| Similar Motifs Found | motif file (matrix) |
