## Supplemental Table 1 for "The Establishment of Cell-Type Specific Gene Regulation in the Sea Urchin Embryo": Hpf40_FilopodialCells_homerResults.html

homer\_motifs\_wbackground/Hpf40\_FilopodialCells/ - Homer de novo Motif Results


### Homer *de novo* Motif Results (homer\_motifs\_wbackground/Hpf40\_FilopodialCells/)

Known Motif Enrichment Results  
Gene Ontology Enrichment Results  
If Homer is having trouble matching a motif to a known motif, try copy/pasting the matrix file into
STAMP  
More information on motif finding results: HOMER
| Description of Results
| Tips
  
Total target sequences = 365  
Total background sequences = 8052  
\* - possible false positive  

|  |  |  |  |  |  |  |  |  |
| --- | --- | --- | --- | --- | --- | --- | --- | --- |
| Rank | Motif | P-value | log P-pvalue | % of Targets | % of Background | STD(Bg STD) | Best Match/Details | Motif File |
| 1 \* | A G T C C G T A G T A C A C G T G C A T G T A C G T A C T A G C C G T A C A T G | 1e-9 | -2.265e+01 | 5.75% | 0.97% | 297.4bp (388.1bp) | RBPJ/MA1116.1/Jaspar(0.752) More Information \| Similar Motifs Found | motif file (matrix) |
| 2 \* | T G C A A C G T C G A T C T G A G A T C C A G T T G C A T A C G G C A T T C G A | 1e-9 | -2.237e+01 | 12.05% | 4.06% | 300.5bp (423.7bp) | ARR1/Literature(Harbison)/Yeast(0.645) More Information \| Similar Motifs Found | motif file (matrix) |
| 3 \* | A C G T A C T G G T A C C G T A A C G T A C T G C A G T A G C T T A C G A C T G | 1e-8 | -2.040e+01 | 8.77% | 2.50% | 321.3bp (443.3bp) | FUS3/MA0565.2/Jaspar(0.830) More Information \| Similar Motifs Found | motif file (matrix) |
| 4 \* | A C T G C T A G C T G A G T C A A C G T T A C G C A G T A G T C G T C A C G A T | 1e-8 | -1.845e+01 | 6.58% | 1.61% | 242.4bp (397.4bp) | TEAD2/MA1121.1/Jaspar(0.800) More Information \| Similar Motifs Found | motif file (matrix) |
| 5 \* | G A C T G T A C G C A T C G T A G A C T G A C T A C G T A C T G A C T G C T G A | 1e-7 | -1.811e+01 | 12.33% | 4.85% | 292.9bp (428.3bp) | br(var.2)/MA0011.1/Jaspar(0.693) More Information \| Similar Motifs Found | motif file (matrix) |
| 6 \* | G T C A A C G T C G A T A C G T A G T C G T A C C A G T C A G T C G A T G T A C | 1e-7 | -1.789e+01 | 20.00% | 10.18% | 362.2bp (431.0bp) | EWS:ERG-fusion(ETS)/CADO\_ES1-EWS:ERG-ChIP-Seq(SRA014231)/Homer(0.852) More Information \| Similar Motifs Found | motif file (matrix) |
| 7 \* | A C G T A C T G A C T G C G T A A C G T A C T G A C G T C G T A C G T A A G T C | 1e-6 | -1.593e+01 | 2.47% | 0.22% | 324.0bp (216.6bp) | PF13\_0315(RRM)/Plasmodium\_falciparum-RNCMPT00200-PBM/HughesRNA(0.710) More Information \| Similar Motifs Found | motif file (matrix) |
| 8 \* | A G T C A C T G C G A T T G C A A C T G C T A G G T C A C G A T C G T A A G C T | 1e-6 | -1.578e+01 | 4.38% | 0.85% | 335.7bp (468.5bp) | HRB27C(RRM)/Drosophila\_melanogaster-RNCMPT00093-PBM/HughesRNA(0.718) More Information \| Similar Motifs Found | motif file (matrix) |
| 9 \* | C G T A A G T C A G T C A C G T A G C T A C T G C A G T C G A T A G T C A G T C | 1e-6 | -1.489e+01 | 4.93% | 1.15% | 241.4bp (420.6bp) | PB0200.1\_Zfp187\_2/Jaspar(0.740) More Information \| Similar Motifs Found | motif file (matrix) |
| 10 \* | A G T C A C T G A C G T C G T A A G T C C G T A A G T C A T G C | 1e-5 | -1.311e+01 | 15.89% | 8.38% | 267.5bp (406.2bp) | SPL12/MA1057.1/Jaspar(0.791) More Information \| Similar Motifs Found | motif file (matrix) |
| 11 \* | A C G T C G T A A C T G A C G T C G T A A C G T C G A T C T A G T A C G C G T A | 1e-4 | -9.355e+00 | 3.01% | 0.73% | 447.1bp (488.1bp) | ZNF410/MA0752.1/Jaspar(0.744) More Information \| Similar Motifs Found | motif file (matrix) |
| 12 \* | A C G T A C T G C G A T C T A G T G A C G A T C C G A T G C A T | 1e-3 | -8.647e+00 | 32.88% | 24.46% | 338.2bp (418.7bp) | MEC-8(RRM)/Caenorhabditis\_elegans-RNCMPT00181-PBM/HughesRNA(0.803) More Information \| Similar Motifs Found | motif file (matrix) |
| 13 \* | A C G T C T A G A G T C A C G T A C G T C G A T C T A G C G A T A G T C A G T C | 1e-3 | -8.633e+00 | 3.29% | 0.92% | 310.5bp (332.7bp) | CG5213(RRM)/Drosophila\_melanogaster-RNCMPT00010-PBM/HughesRNA(0.681) More Information \| Similar Motifs Found | motif file (matrix) |
| 14 \* | A C G T A G T C T C A G A G T C C G T A A C G T A C T G A C T G A C T G A G T C | 1e-3 | -7.622e+00 | 6.30% | 2.90% | 197.3bp (422.4bp) | NFY(CCAAT)/Promoter/Homer(0.690) More Information \| Similar Motifs Found | motif file (matrix) |
| 15 \* | A G T C A C G T A G T C A C G T A C T G A C G T A G T C A C T G A G T C A C G T | 1e-2 | -6.424e+00 | 0.82% | 0.07% | 126.7bp (634.1bp) | CNOT4(RRM)/Homo\_sapiens-RNCMPT00156-PBM/HughesRNA(0.755) More Information \| Similar Motifs Found | motif file (matrix) |
| 16 \* | T G A C G A C T C T G A G C A T T A G C A C G T T C G A C G A T T G A C G A C T | 1e-2 | -5.742e+00 | 4.66% | 2.19% | 392.2bp (475.8bp) | LHY(Myb)/Seedling-LHY-ChIP-Seq(GSE52175)/Homer(0.689) More Information \| Similar Motifs Found | motif file (matrix) |
| 17 \* | A C T G A C T G A C T G C G T A A C T G A G T C | 1e-1 | -4.542e+00 | 21.64% | 16.88% | 381.9bp (456.5bp) | POL013.1\_MED-1/Jaspar(0.905) More Information \| Similar Motifs Found | motif file (matrix) |
| 18 \* | A C T G A G C T T A C G A C G T T C G A A C G T A T C G A G C T A C T G G C A T | 1e-1 | -4.456e+00 | 18.63% | 14.22% | 353.6bp (428.9bp) | SeqBias: CA-repeat(0.807) More Information \| Similar Motifs Found | motif file (matrix) |
| 19 \* | A C T G A C T G A C G T A C T G A C G T A C G T A C G T C G T A C G T A A C T G | 1e-1 | -2.447e+00 | 0.27% | 0.03% | 273.8bp (309.4bp) | Ct/dmmpmm(Noyes\_hd)/fly(0.722) More Information \| Similar Motifs Found | motif file (matrix) |
| 20 \* | C G T A C G T A A G T C A C T G A G T C A C T G C G T A A C T G A C T G A C G T | 1e0 | -2.063e+00 | 0.27% | 0.05% | 44.5bp (128.6bp) | MBP1::SWI6/MA0330.1/Jaspar(0.721) More Information \| Similar Motifs Found | motif file (matrix) |
| 21 \* | A G T C A C G T A C G T T G A C C G A T C A G T A G T C C G A T A C G T A T G C | 1e0 | -1.905e+00 | 7.12% | 5.72% | 323.8bp (401.0bp) | Unknown4/Arabidopsis-Promoters/Homer(0.846) More Information \| Similar Motifs Found | motif file (matrix) |
| 22 \* | A C G T A C G T A C G T A G T C C G T A C G T A A C G T A C T G A C T G A C G T | 1e0 | -1.904e+00 | 10.96% | 9.25% | 297.7bp (464.9bp) | At2g41835(C2H2)/col-At2g41835-DAP-Seq(GSE60143)/Homer(0.701) More Information \| Similar Motifs Found | motif file (matrix) |
| 23 \* | A C T G A C T G A C G T A C T G A C G T A C G T C G T A A C T G A C T G A G T C | 1e0 | -1.798e+00 | 0.27% | 0.05% | 193.0bp (91.8bp) | MYB116(MYB)/colamp-MYB116-DAP-Seq(GSE60143)/Homer(0.772) More Information \| Similar Motifs Found | motif file (matrix) |
| 24 \* | A C T G A C G T C G T A A C G T A C T G A C T G A C G T A C T G C G T A C G T A | 1e0 | -1.094e+00 | 0.27% | 0.12% | 200.7bp (390.8bp) | ASHR1(ND)/col-ASHR1-DAP-Seq(GSE60143)/Homer(0.773) More Information \| Similar Motifs Found | motif file (matrix) |
