## Supplemental Table 1 for "The Establishment of Cell-Type Specific Gene Regulation in the Sea Urchin Embryo": Hpf40_Foregut_homerResults.html

homer\_motifs\_wbackground/Hpf40\_Foregut/ - Homer de novo Motif Results


### Homer *de novo* Motif Results (homer\_motifs\_wbackground/Hpf40\_Foregut/)

Known Motif Enrichment Results  
Gene Ontology Enrichment Results  
If Homer is having trouble matching a motif to a known motif, try copy/pasting the matrix file into
STAMP  
More information on motif finding results: HOMER
| Description of Results
| Tips
  
Total target sequences = 579  
Total background sequences = 7578  
\* - possible false positive  

|  |  |  |  |  |  |  |  |  |
| --- | --- | --- | --- | --- | --- | --- | --- | --- |
| Rank | Motif | P-value | log P-pvalue | % of Targets | % of Background | STD(Bg STD) | Best Match/Details | Motif File |
| 1 | C G T A T C A G C G T A C G T A C A G T T C G A T A C G A G T C G T A C T C A G | 1e-16 | -3.710e+01 | 29.36% | 15.66% | 354.5bp (453.7bp) | PABPN1(RRM)/Homo\_sapiens-RNCMPT00157-PBM/HughesRNA(0.797) More Information \| Similar Motifs Found | motif file (matrix) |
| 2 | A C G T A C T G C G T A C T G A G T C A G T C A C T A G T C G A C T A G G T C A | 1e-15 | -3.524e+01 | 21.24% | 9.91% | 411.8bp (450.8bp) | CDF3(C2C2dof)/colamp-CDF3-DAP-Seq(GSE60143)/Homer(0.771) More Information \| Similar Motifs Found | motif file (matrix) |
| 3 | A G C T G A C T A T C G C A G T A G T C G C A T G T C A G T A C C A G T G T A C | 1e-14 | -3.439e+01 | 32.12% | 18.33% | 431.6bp (409.5bp) | FOXN3/MA1489.1/Jaspar(0.755) More Information \| Similar Motifs Found | motif file (matrix) |
| 4 | A T C G G A C T G T C A G T C A C T G A A G T C | 1e-14 | -3.370e+01 | 74.78% | 59.15% | 383.4bp (422.5bp) | Foxo3(Forkhead)/U2OS-Foxo3-ChIP-Seq(E-MTAB-2701)/Homer(0.778) More Information \| Similar Motifs Found | motif file (matrix) |
| 5 | C T G A C T A G C A T G T C A G A C G T A C G T G A C T A C G T | 1e-14 | -3.301e+01 | 43.87% | 28.64% | 417.7bp (426.4bp) | TRP2(MYBrelated)/colamp-TRP2-DAP-Seq(GSE60143)/Homer(0.872) More Information \| Similar Motifs Found | motif file (matrix) |
| 6 | A G T C G T A C A G T C G T C A A C G T A G C T A G T C G C T A | 1e-14 | -3.228e+01 | 50.95% | 35.31% | 368.0bp (426.1bp) | PB0170.1\_Sox17\_2/Jaspar(0.829) More Information \| Similar Motifs Found | motif file (matrix) |
| 7 | C G T A A C G T G T A C G T A C A G T C A G T C A G T C C G T A A G C T A G C T | 1e-13 | -3.199e+01 | 6.04% | 1.19% | 356.2bp (415.0bp) | Znf281/MA1630.1/Jaspar(0.707) More Information \| Similar Motifs Found | motif file (matrix) |
| 8 | C T G A G T C A G C T A T A G C T G C A A G T C G T C A T C A G T A C G A T C G | 1e-13 | -3.087e+01 | 9.33% | 2.82% | 330.6bp (421.3bp) | REF6(Zf)/Arabidopsis-REF6-ChIP-Seq(GSE106942)/Homer(0.670) More Information \| Similar Motifs Found | motif file (matrix) |
| 9 | A C G T C T A G T C G A A T C G C A T G G C A T G C A T G A T C A C G T A C G T | 1e-13 | -3.077e+01 | 7.60% | 1.94% | 315.5bp (454.8bp) | RUNX3/MA0684.2/Jaspar(0.774) More Information \| Similar Motifs Found | motif file (matrix) |
| 10 | A C G T A G C T C A T G T C A G C A G T A G T C T A G C A T G C A G C T A G C T | 1e-12 | -2.964e+01 | 6.74% | 1.61% | 401.2bp (427.6bp) | TCP5/MA1067.1/Jaspar(0.747) More Information \| Similar Motifs Found | motif file (matrix) |
| 11 | A C G T A C G T A G T C G T C A C T A G C T G A C G T A A G T C A C T G A T C G | 1e-12 | -2.840e+01 | 5.35% | 1.07% | 341.9bp (403.3bp) | YML081W(MacIsaac)/Yeast(0.666) More Information \| Similar Motifs Found | motif file (matrix) |
| 12 \* | G T C A G T C A C G T A G T A C G T A C A T G C A C T G A T G C G T A C G C T A | 1e-11 | -2.757e+01 | 14.68% | 6.39% | 340.2bp (421.8bp) | ACE2(MacIsaac)/Yeast(0.804) More Information \| Similar Motifs Found | motif file (matrix) |
| 13 \* | C A G T A C G T T G C A G T A C A C G T G C T A A G T C T G C A T A C G A T G C | 1e-11 | -2.742e+01 | 6.56% | 1.64% | 348.5bp (485.4bp) | br-Z2/dmmpmm(SeSiMCMC)/fly(0.705) More Information \| Similar Motifs Found | motif file (matrix) |
| 14 \* | C A G T C G T A A G C T C G A T A T C G A C G T C T A G C T G A A C T G A C T G | 1e-11 | -2.560e+01 | 11.23% | 4.37% | 443.8bp (372.7bp) | MATA1(MacIsaac)/Yeast(0.744) More Information \| Similar Motifs Found | motif file (matrix) |
| 15 \* | G C T A T C G A A T C G G A T C G A T C G C A T A G T C G A C T A G C T A G T C | 1e-10 | -2.450e+01 | 49.40% | 35.92% | 351.6bp (428.3bp) | Crx/MA0467.1/Jaspar(0.701) More Information \| Similar Motifs Found | motif file (matrix) |
| 16 \* | G C T A C T A G T C G A A T C G A T C G G C A T A C T G G C A T G A C T T G C A | 1e-10 | -2.343e+01 | 8.98% | 3.23% | 451.3bp (511.4bp) | Tbx5(T-box)/HL1-Tbx5.biotin-ChIP-Seq(GSE21529)/Homer(0.872) More Information \| Similar Motifs Found | motif file (matrix) |
| 17 \* | A C G T T C A G A C G T A C G T A G C T A C T G A C G T A C G T A T G C C T G A | 1e-9 | -2.296e+01 | 16.58% | 8.35% | 289.9bp (434.4bp) | Tv\_0258(RRM)/Trichomonas\_vaginalis-RNCMPT00258-PBM/HughesRNA(0.752) More Information \| Similar Motifs Found | motif file (matrix) |
| 18 \* | A C T G C A T G G T A C T A G C T C A G A C T G T G C A A C G T A C T G C G T A | 1e-7 | -1.729e+01 | 4.66% | 1.33% | 306.5bp (411.4bp) | RBM47(RRM)/Gallus\_gallus-RNCMPT00279-PBM/HughesRNA(0.714) More Information \| Similar Motifs Found | motif file (matrix) |
| 19 \* | C T A G C G T A G C A T C G T A C T A G C T G A A G C T A C G T G T A C C G A T | 1e-7 | -1.670e+01 | 5.87% | 2.03% | 350.2bp (436.4bp) | At1g25550(G2like)/colamp-At1g25550-DAP-Seq(GSE60143)/Homer(0.730) More Information \| Similar Motifs Found | motif file (matrix) |
| 20 \* | A C G T A C T G A C G T A G T C A C T G G T C A G T C A A G T C C G T A C G T A | 1e-6 | -1.596e+01 | 1.55% | 0.14% | 309.8bp (434.2bp) | SOX14/MA1562.1/Jaspar(0.707) More Information \| Similar Motifs Found | motif file (matrix) |
| 21 \* | A C G T A C T G A C G T A T C G A C T G A C G T A G T C A G T C | 1e-6 | -1.558e+01 | 9.67% | 4.57% | 358.2bp (461.6bp) | TCP3(TCP)/colamp-TCP3-DAP-Seq(GSE60143)/Homer(0.885) More Information \| Similar Motifs Found | motif file (matrix) |
| 22 \* | A C T G A C G T A C T G C G T A A G T C C G T A C G T A A G T C C G T A A C G T | 1e-5 | -1.167e+01 | 1.04% | 0.09% | 232.5bp (356.0bp) | CREB1/MA0018.4/Jaspar(0.746) More Information \| Similar Motifs Found | motif file (matrix) |
| 23 \* | A G T C A C T G A C T G C G T A A T C G C T G A C A T G C G T A | 1e-4 | -1.129e+01 | 10.02% | 5.53% | 344.8bp (455.6bp) | OAF1/MA0348.1/Jaspar(0.778) More Information \| Similar Motifs Found | motif file (matrix) |
| 24 \* | A G T C C G T A A G T C A C T G A G T C A C T G A C G T C G T A A G T C A C T G | 1e-3 | -8.371e+00 | 0.86% | 0.10% | 205.2bp (138.6bp) | TCFL5/MA0632.2/Jaspar(0.735) More Information \| Similar Motifs Found | motif file (matrix) |
