## Supplemental Table 1 for "The Establishment of Cell-Type Specific Gene Regulation in the Sea Urchin Embryo": Hpf40_Forgut_homerResults.html

homer\_motifs\_wbackground/Hpf40\_Forgut/ - Homer de novo Motif Results


### Homer *de novo* Motif Results (homer\_motifs\_wbackground/Hpf40\_Forgut/)

Known Motif Enrichment Results  
Gene Ontology Enrichment Results  
If Homer is having trouble matching a motif to a known motif, try copy/pasting the matrix file into
STAMP  
More information on motif finding results: HOMER
| Description of Results
| Tips
  
Total target sequences = 490  
Total background sequences = 8030  
\* - possible false positive  

|  |  |  |  |  |  |  |  |  |
| --- | --- | --- | --- | --- | --- | --- | --- | --- |
| Rank | Motif | P-value | log P-pvalue | % of Targets | % of Background | STD(Bg STD) | Best Match/Details | Motif File |
| 1 \* | C G T A C T A G A G T C C G T A C G T A T G C A A G T C C G T A | 1e-9 | -2.273e+01 | 46.94% | 33.07% | 341.1bp (389.4bp) | PHA-4(Forkhead)/cElegans-Embryos-PHA4-ChIP-Seq(modEncode)/Homer(0.907) More Information \| Similar Motifs Found | motif file (matrix) |
| 2 \* | A C T G T A G C G T C A G T A C C G A T T A C G C G T A C G T A A C G T A T G C | 1e-9 | -2.263e+01 | 7.14% | 1.98% | 250.2bp (388.6bp) | Ap1/dmmpmm(Papatsenko)/fly(0.712) More Information \| Similar Motifs Found | motif file (matrix) |
| 3 \* | C G T A A C T G A C G T A C G T A G T C A T C G A C T G A C T G C G T A C G T A | 1e-9 | -2.215e+01 | 1.63% | 0.05% | 287.8bp (325.2bp) | Hsf/MA1458.1/Jaspar(0.727) More Information \| Similar Motifs Found | motif file (matrix) |
| 4 \* | G C T A G T C A G T A C C T G A C A G T A G T C A T C G A C T G C G T A C G T A | 1e-9 | -2.136e+01 | 7.14% | 2.09% | 288.0bp (380.5bp) | PB0115.1\_Ehf\_2/Jaspar(0.778) More Information \| Similar Motifs Found | motif file (matrix) |
| 5 \* | A T G C A C T G A G T C A C G T C T G A C G A T A C T G C T G A A C T G C G T A | 1e-9 | -2.121e+01 | 2.65% | 0.26% | 428.2bp (470.6bp) | PUF68(RRM)/Drosophila\_melanogaster-RNCMPT00141-PBM/HughesRNA(0.727) More Information \| Similar Motifs Found | motif file (matrix) |
| 6 \* | A C T G T C A G A T C G C G T A A C T G A C T G C A G T C G T A C T A G A C T G | 1e-9 | -2.095e+01 | 8.16% | 2.67% | 515.4bp (440.7bp) | vfl/MA1462.1/Jaspar(0.762) More Information \| Similar Motifs Found | motif file (matrix) |
| 7 \* | C G T A C G T A A G T C G C T A A G T C A T C G A G T C C G T A A G T C C G T A | 1e-8 | -1.904e+01 | 3.06% | 0.43% | 247.1bp (467.1bp) | Tp\_0225(RRM)/Thalassiosira\_pseudonana-RNCMPT00225-PBM/HughesRNA(0.775) More Information \| Similar Motifs Found | motif file (matrix) |
| 8 \* | A G T C A C T G G T A C A C T G A C G T A C T G A C G T A G C T C G T A G T C A | 1e-8 | -1.873e+01 | 4.49% | 0.99% | 376.9bp (447.5bp) | Tp\_0225(RRM)/Thalassiosira\_pseudonana-RNCMPT00225-PBM/HughesRNA(0.749) More Information \| Similar Motifs Found | motif file (matrix) |
| 9 \* | C A T G C G T A A C G T A G T C A G T C A G T C A G T C C G T A C G T A A C T G | 1e-7 | -1.783e+01 | 1.63% | 0.09% | 157.4bp (281.9bp) | ESRP2(RRM)/Homo\_sapiens-RNCMPT00150-PBM/HughesRNA(0.790) More Information \| Similar Motifs Found | motif file (matrix) |
| 10 \* | G A T C A C T G G T C A C T A G C G T A G T C A A C G T A C T G C T G A T A G C | 1e-7 | -1.754e+01 | 5.51% | 1.55% | 309.8bp (381.5bp) | TARDBP(RRM)/Homo\_sapiens-RNCMPT00076-PBM/HughesRNA(0.756) More Information \| Similar Motifs Found | motif file (matrix) |
| 11 \* | A G T C C G T A A C T G A C T G A C G T A C G T C G T A A C T G C G T A C G T A | 1e-7 | -1.713e+01 | 1.43% | 0.07% | 200.0bp (137.5bp) | HRB27C(RRM)/Drosophila\_melanogaster-RNCMPT00028-PBM/HughesRNA(0.717) More Information \| Similar Motifs Found | motif file (matrix) |
| 12 \* | A T G C C G T A A T G C G T A C A G T C C G A T A C G T A C T G C A G T C A G T | 1e-7 | -1.636e+01 | 5.92% | 1.86% | 223.4bp (474.0bp) | Kr/dmmpmm(SeSiMCMC)/fly(0.711) More Information \| Similar Motifs Found | motif file (matrix) |
| 13 \* | C G T A C A G T G C T A C G T A A G C T C T G A C T G A C T A G G T A C G A C T | 1e-7 | -1.612e+01 | 22.45% | 13.69% | 314.7bp (387.1bp) | QKR58E-1(KH)/Drosophila\_melanogaster-RNCMPT00142-PBM/HughesRNA(0.771) More Information \| Similar Motifs Found | motif file (matrix) |
| 14 \* | C G T A A C G T A C T G A G T C A C G T C G T A C T G A A G C T A C T G A C G T | 1e-6 | -1.522e+01 | 2.04% | 0.23% | 286.9bp (634.2bp) | Oct6(POU,Homeobox)/NPC-Pou3f1-ChIP-Seq(GSE35496)/Homer(0.802) More Information \| Similar Motifs Found | motif file (matrix) |
| 15 \* | C G A T G T A C G A T C G T A C A C G T T G C A A T C G G A C T G C T A G A T C | 1e-5 | -1.334e+01 | 8.98% | 4.12% | 313.8bp (417.5bp) | HNRNPA2B1(RRM)/Homo\_sapiens-RNCMPT00024-PBM/HughesRNA(0.758) More Information \| Similar Motifs Found | motif file (matrix) |
| 16 \* | A G T C A G T C A C G T A G T C A G T C C G T A A C G T A C T G A C G T A G T C | 1e-4 | -1.012e+01 | 0.82% | 0.04% | 255.1bp (140.5bp) | YY2/MA0748.2/Jaspar(0.675) More Information \| Similar Motifs Found | motif file (matrix) |
| 17 \* | A C T G A C T G A C G T C G T A A C T G A G T C A C T G A C G T A C G T A C G T | 1e-4 | -1.012e+01 | 0.82% | 0.04% | 141.7bp (454.5bp) | PB0154.1\_Osr1\_2/Jaspar(0.723) More Information \| Similar Motifs Found | motif file (matrix) |
| 18 \* | C T A G A C T G G A T C A C G T A C T G G T A C A G C T A C T G | 1e-4 | -9.892e+00 | 12.24% | 7.22% | 367.6bp (391.8bp) | ERF15(AP2EREBP)/colamp-ERF15-DAP-Seq(GSE60143)/Homer(0.810) More Information \| Similar Motifs Found | motif file (matrix) |
| 19 \* | A C T G C T A G A G C T C A T G A T C G C G T A A C T G C T A G | 1e-3 | -7.379e+00 | 8.37% | 4.87% | 372.2bp (476.5bp) | Pp\_0237(RRM)/Physcomitrella\_patens-RNCMPT00237-PBM/HughesRNA(0.811) More Information \| Similar Motifs Found | motif file (matrix) |
| 20 \* | T C G A C T A G A T C G T G C A T A G C A G T C C G T A A G C T G T A C G T A C | 1e-3 | -7.346e+00 | 1.02% | 0.14% | 179.4bp (605.3bp) | GLI3(Zf)/Limb-GLI3-ChIP-Chip(GSE11077)/Homer(0.805) More Information \| Similar Motifs Found | motif file (matrix) |
| 21 \* | C T A G C T A G C A G T T C A G C T A G G A C T C G T A T A C G C G A T C T A G | 1e-1 | -3.793e+00 | 5.51% | 3.64% | 310.3bp (415.8bp) | ZBTB7C/MA0695.1/Jaspar(0.721) More Information \| Similar Motifs Found | motif file (matrix) |
| 22 \* | C G T A A G T C A C T G A C G T A C T G C G A T | 1e-1 | -3.431e+00 | 41.63% | 37.48% | 349.4bp (402.1bp) | HAC1/MA0310.1/Jaspar(0.941) More Information \| Similar Motifs Found | motif file (matrix) |
| 23 \* | C G T A G T A C A G T C C T G A A G T C A T G C C G T A A T C G | 1e-1 | -2.771e+00 | 5.92% | 4.37% | 338.8bp (381.5bp) | BRU-3(RRM)/Drosophila\_melanogaster-RNCMPT00122-PBM/HughesRNA(0.788) More Information \| Similar Motifs Found | motif file (matrix) |
