## Supplemental Table 1 for "The Establishment of Cell-Type Specific Gene Regulation in the Sea Urchin Embryo": Hpf40_GermCells_homerResults.html

homer\_motifs\_wbackground/Hpf40\_GermCells/ - Homer de novo Motif Results


### Homer *de novo* Motif Results (homer\_motifs\_wbackground/Hpf40\_GermCells/)

Known Motif Enrichment Results  
Gene Ontology Enrichment Results  
If Homer is having trouble matching a motif to a known motif, try copy/pasting the matrix file into
STAMP  
More information on motif finding results: HOMER
| Description of Results
| Tips
  
Total target sequences = 364  
Total background sequences = 8265  
\* - possible false positive  

|  |  |  |  |  |  |  |  |  |
| --- | --- | --- | --- | --- | --- | --- | --- | --- |
| Rank | Motif | P-value | log P-pvalue | % of Targets | % of Background | STD(Bg STD) | Best Match/Details | Motif File |
| 1 \* | A G T C A C T G A C G T C T A G C T A G A C G T A C T G C T A G A C G T T C G A | 1e-9 | -2.076e+01 | 4.67% | 0.68% | 321.1bp (514.9bp) | BRU-3(RRM)/Drosophila\_melanogaster-RNCMPT00122-PBM/HughesRNA(0.710) More Information \| Similar Motifs Found | motif file (matrix) |
| 2 \* | A G T C T C A G C G T A C G T A A T G C A C G T A C G T C T G A A C T G C A G T | 1e-8 | -2.041e+01 | 3.85% | 0.45% | 312.8bp (348.1bp) | EXC-7(RRM)/Caenorhabditis\_elegans-RNCMPT00014-PBM/HughesRNA(0.728) More Information \| Similar Motifs Found | motif file (matrix) |
| 3 \* | C G A T C T A G A C T G G T C A A T C G T G A C C G T A C G A T C T A G G A C T | 1e-8 | -2.011e+01 | 6.04% | 1.24% | 283.3bp (404.1bp) | EIF-2ALPHA(S1)/Drosophila\_melanogaster-RNCMPT00273-PBM/HughesRNA(0.793) More Information \| Similar Motifs Found | motif file (matrix) |
| 4 \* | G T C A C G A T A T C G C G A T A G T C C G A T C G T A C T G A T A G C G C T A | 1e-8 | -1.929e+01 | 7.69% | 2.07% | 273.8bp (409.4bp) | Tbox:Smad(T-box,MAD)/ESCd5-Smad2\_3-ChIP-Seq(GSE29422)/Homer(0.707) More Information \| Similar Motifs Found | motif file (matrix) |
| 5 \* | T A C G C G T A C G T A C G T A A C G T C T A G G A T C C T A G A G C T C G T A | 1e-8 | -1.861e+01 | 6.04% | 1.34% | 412.0bp (426.4bp) | CST6(MacIsaac)/Yeast(0.754) More Information \| Similar Motifs Found | motif file (matrix) |
| 6 \* | T A C G A C G T A G T C C A T G A C T G C G T A C T A G T A C G C T G A C G T A | 1e-8 | -1.849e+01 | 4.95% | 0.90% | 329.7bp (527.5bp) | LIN28A(CSD)/Homo\_sapiens-RNCMPT00162-PBM/HughesRNA(0.802) More Information \| Similar Motifs Found | motif file (matrix) |
| 7 \* | C G T A A C T G C G A T C T A G G A T C G T C A C G T A G T A C C G A T A G C T | 1e-7 | -1.786e+01 | 7.14% | 1.93% | 261.7bp (365.4bp) | byn/dmmpmm(SeSiMCMC)/fly(0.827) More Information \| Similar Motifs Found | motif file (matrix) |
| 8 \* | A C G T C G A T A T G C T A C G G T C A A C T G A C T G A G T C A C G T A C G T | 1e-7 | -1.767e+01 | 5.49% | 1.19% | 202.4bp (430.0bp) | CRZ1/MA0285.1/Jaspar(0.683) More Information \| Similar Motifs Found | motif file (matrix) |
| 9 \* | T A G C A G T C C G T A G T A C A C G T C T G A G C A T T A G C C T G A A G C T | 1e-6 | -1.475e+01 | 7.14% | 2.27% | 301.4bp (416.1bp) | FZF1/MA0298.1/Jaspar(0.748) More Information \| Similar Motifs Found | motif file (matrix) |
| 10 \* | A C G T A T C G C G T A A T G C C G T A A G T C G A T C T G C A A C T G C T G A | 1e-6 | -1.434e+01 | 9.07% | 3.44% | 213.2bp (439.0bp) | Six4/dmmpmm(Noyes\_hd)/fly(0.740) More Information \| Similar Motifs Found | motif file (matrix) |
| 11 \* | A G T C A C G T G T C A A T G C A C T G G A C T C G T A A C T G G C T A G T C A | 1e-5 | -1.299e+01 | 3.02% | 0.50% | 237.7bp (318.7bp) | Hrp1p(RRM)/Saccharomyces\_cerevisiae-RNCMPT00031-PBM/HughesRNA(0.645) More Information \| Similar Motifs Found | motif file (matrix) |
| 12 \* | C G T A A C T G C G T A A C G T A G T C A C T G A C T G A C G T A C G T A C T G | 1e-4 | -1.139e+01 | 1.10% | 0.05% | 54.6bp (346.7bp) | GATA14(C2C2gata)/col-GATA14-DAP-Seq(GSE60143)/Homer(0.721) More Information \| Similar Motifs Found | motif file (matrix) |
| 13 \* | A C G T C A T G C T A G A C G T A C T G T C G A A G T C A C T G C T A G A C G T | 1e-4 | -1.134e+01 | 4.12% | 1.07% | 206.9bp (515.6bp) | ASHR1(ND)/col-ASHR1-DAP-Seq(GSE60143)/Homer(0.760) More Information \| Similar Motifs Found | motif file (matrix) |
| 14 \* | C G A T T C G A C A T G C T G A C A G T T A G C A G C T C G T A T C A G C G A T | 1e-3 | -8.988e+00 | 19.23% | 12.39% | 336.4bp (399.4bp) | GATA6(C2C2gata)/col200-GATA6-DAP-Seq(GSE60143)/Homer(0.858) More Information \| Similar Motifs Found | motif file (matrix) |
| 15 \* | G A C T C T G A C T G A C G A T G C T A C G T A G C A T T C G A A C G T A G C T | 1e-3 | -8.779e+00 | 17.03% | 10.66% | 346.9bp (402.8bp) | Arid5a/MA0602.1/Jaspar(0.794) More Information \| Similar Motifs Found | motif file (matrix) |
| 16 \* | A G T C T G C A A G T C A G T C C G T A A G T C A G T C A T G C A C G T C G T A | 1e-3 | -8.748e+00 | 4.67% | 1.66% | 257.7bp (417.7bp) | KLF4/MA0039.4/Jaspar(0.775) More Information \| Similar Motifs Found | motif file (matrix) |
| 17 \* | A G T C G A T C A G T C A G T C G T A C T G A C A T G C A C T G A C T G A C T G | 1e-3 | -7.834e+00 | 9.62% | 5.20% | 327.8bp (430.3bp) | PB0103.1\_Zic3\_1/Jaspar(0.870) More Information \| Similar Motifs Found | motif file (matrix) |
| 18 \* | A T C G T C A G A G C T A C T G A C G T C G T A A C T G A T C G | 1e-2 | -6.460e+00 | 21.98% | 15.95% | 356.8bp (421.9bp) | RBM28(RRM)/Homo\_sapiens-RNCMPT00049-PBM/HughesRNA(0.772) More Information \| Similar Motifs Found | motif file (matrix) |
| 19 \* | A C G T A C G T A G T C C G T A A C G T C G T A C G T A A C T G A G T C A T G C | 1e-2 | -6.166e+00 | 1.37% | 0.24% | 174.1bp (361.8bp) | INO4(MacIsaac)/Yeast(0.684) More Information \| Similar Motifs Found | motif file (matrix) |
| 20 \* | T A G C G T C A C A G T G C T A G A C T G C A T A G C T A G C T A T G C C G T A | 1e-2 | -5.917e+00 | 20.05% | 14.57% | 256.1bp (404.7bp) | PABP(RRM)/Drosophila\_melanogaster-RNCMPT00139-PBM/HughesRNA(0.713) More Information \| Similar Motifs Found | motif file (matrix) |
| 21 \* | A C G T A G T C G T A C G A C T C A G T A T C G C T A G C T G A | 1e-2 | -5.717e+00 | 37.91% | 31.08% | 332.8bp (417.2bp) | Nr5a2(NR)/mES-Nr5a2-ChIP-Seq(GSE19019)/Homer(0.725) More Information \| Similar Motifs Found | motif file (matrix) |
| 22 \* | A C T G G C T A C T A G A C G T T G A C C G A T C G T A T C A G | 1e-2 | -5.662e+00 | 10.71% | 6.79% | 283.5bp (394.4bp) | Smad4/MA1153.1/Jaspar(0.780) More Information \| Similar Motifs Found | motif file (matrix) |
| 23 \* | A G T C A C G T C G T A A C T G A G T C A C G T A C G T C G T A A G T C A C G T | 1e-2 | -4.831e+00 | 0.55% | 0.04% | 220.7bp (290.3bp) | CAD1/CAD1\_YPD/2-CAD1,2-YAP1(Harbison)/Yeast(0.681) More Information \| Similar Motifs Found | motif file (matrix) |
| 24 \* | G T C A G T A C C G T A C A T G G C A T G T A C A G C T A C T G A G C T A G C T | 1e-1 | -4.433e+00 | 2.75% | 1.18% | 255.6bp (387.5bp) | PGR(NR)/EndoStromal-PGR-ChIP-Seq(GSE69539)/Homer(0.791) More Information \| Similar Motifs Found | motif file (matrix) |
| 25 \* | A G T C C G T A C G T A A C T G C G T A A C T G C G T A A G T C A C G T A C G T | 1e-1 | -4.228e+00 | 1.10% | 0.26% | 326.3bp (347.0bp) | NAC078/MA1677.1/Jaspar(0.705) More Information \| Similar Motifs Found | motif file (matrix) |
| 26 \* | C A T G G T A C C G T A T A G C G T C A A G C T C G A T T A G C | 1e-1 | -3.879e+00 | 13.74% | 10.25% | 375.0bp (408.0bp) | AT3G24120/MA1388.1/Jaspar(0.791) More Information \| Similar Motifs Found | motif file (matrix) |
| 27 \* | C G T A A C T G C G T A A G T C C G T A A G T C C G T A A G T C A C G T A C T G | 1e-1 | -2.460e+00 | 0.55% | 0.14% | 167.1bp (270.5bp) | Nv\_0278(RRM)/Nematostella\_vectensis-RNCMPT00278-PBM/HughesRNA(0.756) More Information \| Similar Motifs Found | motif file (matrix) |
