## Supplemental Table 1 for "The Establishment of Cell-Type Specific Gene Regulation in the Sea Urchin Embryo": Hpf40_GlobularCells_homerResults.html

homer\_motifs\_wbackground/Hpf40\_GlobularCells/ - Homer de novo Motif Results


### Homer *de novo* Motif Results (homer\_motifs\_wbackground/Hpf40\_GlobularCells/)

Known Motif Enrichment Results  
Gene Ontology Enrichment Results  
If Homer is having trouble matching a motif to a known motif, try copy/pasting the matrix file into
STAMP  
More information on motif finding results: HOMER
| Description of Results
| Tips
  
Total target sequences = 346  
Total background sequences = 8085  
\* - possible false positive  

|  |  |  |  |  |  |  |  |  |
| --- | --- | --- | --- | --- | --- | --- | --- | --- |
| Rank | Motif | P-value | log P-pvalue | % of Targets | % of Background | STD(Bg STD) | Best Match/Details | Motif File |
| 1 \* | G T A C C A T G A C G T G T C A A C G T A G T C G T A C A C G T A C T G A G T C | 1e-9 | -2.078e+01 | 5.20% | 0.82% | 308.1bp (350.0bp) | ttk/dmmpmm(SeSiMCMC)/fly(0.733) More Information \| Similar Motifs Found | motif file (matrix) |
| 2 \* | G T A C A C G T A C T G A C T G G A T C A G T C A C G T A C T G C T A G G T C A | 1e-8 | -1.913e+01 | 6.65% | 1.50% | 392.3bp (372.0bp) | SKN7(MacIsaac)/Yeast(0.706) More Information \| Similar Motifs Found | motif file (matrix) |
| 3 \* | C G T A G T C A G T C A G T C A A C G T A C G T C G A T A T G C A C T G C G A T | 1e-8 | -1.880e+01 | 9.54% | 2.98% | 249.5bp (502.6bp) | CG17838(RRM)/Drosophila\_melanogaster-RNCMPT00131-PBM/HughesRNA(0.806) More Information \| Similar Motifs Found | motif file (matrix) |
| 4 \* | A C T G G T A C C G T A T C A G A T C G A C T G A T G C A C T G T A G C A C T G | 1e-8 | -1.855e+01 | 4.62% | 0.73% | 231.1bp (404.7bp) | CTCFL/MA1102.2/Jaspar(0.758) More Information \| Similar Motifs Found | motif file (matrix) |
| 5 \* | C G T A G T C A A C T G C G A T A T G C C A G T G T C A A G C T G T C A A T G C | 1e-7 | -1.759e+01 | 8.67% | 2.67% | 293.4bp (394.1bp) | ZSCAN29/MA1602.1/Jaspar(0.720) More Information \| Similar Motifs Found | motif file (matrix) |
| 6 \* | A G T C A C G T A C T G A C G T C G T A A G T C A T G C T C A G A T C G A G T C | 1e-7 | -1.715e+01 | 2.89% | 0.27% | 326.1bp (346.2bp) | LEU3/LEU3\_SM/47-LEU3(Harbison)/Yeast(0.748) More Information \| Similar Motifs Found | motif file (matrix) |
| 7 \* | T A G C C T A G A G C T T C G A T C G A G C T A G A T C A G C T G T C A A T G C | 1e-6 | -1.521e+01 | 1.45% | 0.05% | 323.5bp (294.9bp) | ARF16(ARF)/col-ARF16-DAP-Seq(GSE60143)/Homer(0.725) More Information \| Similar Motifs Found | motif file (matrix) |
| 8 \* | A G T C G C T A G T A C A C G T G T A C A G C T T A G C C G T A A G T C G T C A | 1e-5 | -1.303e+01 | 8.67% | 3.32% | 293.4bp (434.9bp) | Su(H)/dmmpmm(Bergman)/fly(0.709) More Information \| Similar Motifs Found | motif file (matrix) |
| 9 \* | G T A C A C T G G C T A C G T A C G A T A C G T A G T C A C T G C T G A C G A T | 1e-5 | -1.256e+01 | 2.89% | 0.44% | 141.0bp (471.8bp) | AT5G22990(C2H2)/col-AT5G22990-DAP-Seq(GSE60143)/Homer(0.790) More Information \| Similar Motifs Found | motif file (matrix) |
| 10 \* | C G T A C T G A C T G A G T C A G C A T G C A T C G T A C G T A C T G A C T G A | 1e-5 | -1.240e+01 | 15.90% | 8.41% | 326.4bp (425.6bp) | tll/dmmpmm(Bergman)/fly(0.825) More Information \| Similar Motifs Found | motif file (matrix) |
| 11 \* | C A T G T G C A C T A G G T A C A G C T A C G T G T A C A G T C G C A T A T C G | 1e-4 | -1.132e+01 | 13.29% | 6.81% | 351.6bp (456.7bp) | ETV4/MA0764.2/Jaspar(0.817) More Information \| Similar Motifs Found | motif file (matrix) |
| 12 \* | C T A G C G A T A G T C A C G T C G T A A C T G C T G A A G C T A G T C A C G T | 1e-3 | -8.525e+00 | 9.54% | 4.86% | 389.1bp (418.5bp) | GATA6/MA1396.1/Jaspar(0.913) More Information \| Similar Motifs Found | motif file (matrix) |
| 13 \* | A C T G A C G T A C G T A C G T A G T C A G T C A C T G C T G A A C G T C G T A | 1e-3 | -8.070e+00 | 0.87% | 0.05% | 467.1bp (192.3bp) | ERT1/MA0420.1/Jaspar(0.759) More Information \| Similar Motifs Found | motif file (matrix) |
| 14 \* | A C G T C G T A A C T G A C G T C G T A A C G T A C T G A C G T G T A C A C T G | 1e-3 | -7.581e+00 | 1.45% | 0.18% | 188.6bp (445.7bp) | DDF1(AP2EREBP)/col-DDF1-DAP-Seq(GSE60143)/Homer(0.700) More Information \| Similar Motifs Found | motif file (matrix) |
| 15 \* | C G T A C G A T A C T G C G T A A C T G C T A G C T A G C G A T A T G C A C T G | 1e-3 | -7.427e+00 | 2.31% | 0.53% | 345.2bp (444.9bp) | NRG1(MacIsaac)/Yeast(0.687) More Information \| Similar Motifs Found | motif file (matrix) |
| 16 \* | A C G T A G T C A C G T A G T C A C G T A G T C A C T G C G T A A G T C A G T C | 1e-3 | -7.238e+00 | 0.87% | 0.06% | 265.7bp (215.0bp) | SRSF10(RRM)/Homo\_sapiens-RNCMPT00090-PBM/HughesRNA(0.680) More Information \| Similar Motifs Found | motif file (matrix) |
| 17 \* | A C G T G T A C C A G T C A T G A T G C A G C T A C G T G T A C G A C T A G C T | 1e-2 | -6.124e+00 | 17.63% | 12.23% | 350.6bp (423.5bp) | Unknown4/Arabidopsis-Promoters/Homer(0.761) More Information \| Similar Motifs Found | motif file (matrix) |
| 18 \* | C T A G A G C T A G T C A T G C A G C T C T G A A C T G C T A G A T C G A C T G | 1e-2 | -5.959e+00 | 2.89% | 0.99% | 264.1bp (435.4bp) | MSN4/MA0342.1/Jaspar(0.697) More Information \| Similar Motifs Found | motif file (matrix) |
| 19 \* | A G T C A C T G A C G T A C T G A G T C A C G T A G T C A C G T C G T A A C G T | 1e-2 | -4.897e+00 | 0.58% | 0.04% | 91.8bp (140.9bp) | kni/MA0451.1/Jaspar(0.711) More Information \| Similar Motifs Found | motif file (matrix) |
| 20 \* | C A G T T G A C G T A C A T C G A G T C T G C A A G C T T A C G | 1e-2 | -4.820e+00 | 31.21% | 25.35% | 306.0bp (458.2bp) | CG7903(RRM)/Drosophila\_melanogaster-RNCMPT00144-PBM/HughesRNA(0.805) More Information \| Similar Motifs Found | motif file (matrix) |
| 21 \* | A G T C A C G T C G T A A C G T A C G T A G T C C G T A A C G T C G T A C G T A | 1e-1 | -3.725e+00 | 2.31% | 1.00% | 332.6bp (436.8bp) | Brn2(POU,Homeobox)/NPC-Brn2-ChIP-Seq(GSE35496)/Homer(0.729) More Information \| Similar Motifs Found | motif file (matrix) |
| 22 \* | A C T G G T A C C G T A G T C A G A C T A C G T A C T G A C G T A G T C A C T G | 1e-1 | -3.100e+00 | 1.73% | 0.75% | 144.0bp (425.0bp) | ARF8/MA0944.1/Jaspar(0.735) More Information \| Similar Motifs Found | motif file (matrix) |
| 23 \* | A C T G A C T G A C G T C G A T C G T A C T A G A C T G A C T G C G T A A C T G | 1e0 | -2.134e+00 | 0.87% | 0.35% | 257.7bp (356.0bp) | HNRNPA1(RRM)/Homo\_sapiens-RNCMPT00022-PBM/HughesRNA(0.738) More Information \| Similar Motifs Found | motif file (matrix) |
| 24 \* | A C G T A C G T A C T G A G T C A C G T A G T C A G T C A C T G | 1e0 | -1.849e+00 | 3.18% | 2.24% | 278.9bp (488.9bp) | POL013.1\_MED-1/Jaspar(0.854) More Information \| Similar Motifs Found | motif file (matrix) |
