## Supplemental Table 1 for "The Establishment of Cell-Type Specific Gene Regulation in the Sea Urchin Embryo": Hpf40_Hindgut_homerResults.html

homer\_motifs\_wbackground/Hpf40\_Hindgut/ - Homer de novo Motif Results


### Homer *de novo* Motif Results (homer\_motifs\_wbackground/Hpf40\_Hindgut/)

Known Motif Enrichment Results  
Gene Ontology Enrichment Results  
If Homer is having trouble matching a motif to a known motif, try copy/pasting the matrix file into
STAMP  
More information on motif finding results: HOMER
| Description of Results
| Tips
  
Total target sequences = 1279  
Total background sequences = 6853  
\* - possible false positive  

|  |  |  |  |  |  |  |  |  |
| --- | --- | --- | --- | --- | --- | --- | --- | --- |
| Rank | Motif | P-value | log P-pvalue | % of Targets | % of Background | STD(Bg STD) | Best Match/Details | Motif File |
| 1 | C A G T T C A G G C A T C A G T A C G T C T A G G T A C G C A T A G T C C G T A | 1e-65 | -1.516e+02 | 57.70% | 34.11% | 341.3bp (424.1bp) | fkh/dmmpmm(Noyes)/fly(0.962) More Information \| Similar Motifs Found | motif file (matrix) |
| 2 | T C G A G C T A A C T G C G T A A C G T C G T A G C T A T C A G | 1e-30 | -7.077e+01 | 50.12% | 34.27% | 366.0bp (446.1bp) | GATA3/MA0037.3/Jaspar(0.971) More Information \| Similar Motifs Found | motif file (matrix) |
| 3 | T C G A C G T A C G A T A T G C C G T A A C G T G A C T C T G A G T C A A G T C | 1e-26 | -6.084e+01 | 30.34% | 17.95% | 357.3bp (429.3bp) | HNF1b(Homeobox)/PDAC-HNF1B-ChIP-Seq(GSE64557)/Homer(0.881) More Information \| Similar Motifs Found | motif file (matrix) |
| 4 | G C A T A C T G T C A G C G T A A T G C A C G T G C A T G A C T C T A G C T A G | 1e-17 | -4.079e+01 | 5.63% | 1.67% | 279.3bp (342.3bp) | HNF4A/MA0114.4/Jaspar(0.986) More Information \| Similar Motifs Found | motif file (matrix) |
| 5 | G A T C C T A G T G C A A T G C T A G C G C A T C G A T A C G T T C A G T C A G | 1e-16 | -3.701e+01 | 7.74% | 3.03% | 372.5bp (435.0bp) | HNF4A(var.2)/MA1494.1/Jaspar(0.825) More Information \| Similar Motifs Found | motif file (matrix) |
| 6 | G T A C G C T A A T G C A G C T A T C G C A T G C A G T T C G A C A G T T G A C | 1e-15 | -3.645e+01 | 5.79% | 1.90% | 354.7bp (424.4bp) | AT1G76870(Trihelix)/col-AT1G76870-DAP-Seq(GSE60143)/Homer(0.691) More Information \| Similar Motifs Found | motif file (matrix) |
| 7 | A C T G C G A T A G T C A G C T C T G A C G T A C G T A A C T G | 1e-12 | -2.906e+01 | 13.37% | 7.49% | 348.4bp (419.5bp) | HNF4A(var.2)/MA1494.1/Jaspar(0.731) More Information \| Similar Motifs Found | motif file (matrix) |
| 8 \* | C G T A G C A T A C T G C G T A A T C G A C G T A G T C C G T A | 1e-11 | -2.606e+01 | 15.40% | 9.38% | 376.2bp (443.4bp) | FOS/MA0476.1/Jaspar(0.959) More Information \| Similar Motifs Found | motif file (matrix) |
| 9 \* | A C G T C T G A A C T G A C T G A C G T A G T C A T G C G C T A G T A C A C T G | 1e-9 | -2.217e+01 | 1.02% | 0.09% | 280.5bp (268.1bp) | TCP16(TCP)/colamp-TCP16-DAP-Seq(GSE60143)/Homer(0.721) More Information \| Similar Motifs Found | motif file (matrix) |
| 10 \* | G T C A A T C G A T G C A C G T A T C G A C G T A C G T A C T G C G T A A G C T | 1e-9 | -2.182e+01 | 1.95% | 0.41% | 448.4bp (570.0bp) | SCRT1/MA0743.2/Jaspar(0.713) More Information \| Similar Motifs Found | motif file (matrix) |
| 11 \* | A G C T A C T G C G T A A C T G G T C A C G T A A C G T C G T A C G T A A G T C | 1e-9 | -2.172e+01 | 0.94% | 0.08% | 361.2bp (292.1bp) | KAN2(G2like)/colamp-KAN2-DAP-Seq(GSE60143)/Homer(0.744) More Information \| Similar Motifs Found | motif file (matrix) |
| 12 \* | A C T G A T C G A C T G A C T G A C T G A G T C A C G T A C T G A C G T A C G T | 1e-8 | -2.045e+01 | 0.55% | 0.02% | 197.2bp (126.8bp) | ZNF341/MA1655.1/Jaspar(0.799) More Information \| Similar Motifs Found | motif file (matrix) |
| 13 \* | G T C A C G T A A G T C C T A G C G T A A C T G C G T A A G T C A G T C A G T C | 1e-8 | -1.909e+01 | 1.17% | 0.17% | 436.4bp (407.3bp) | LRF(Zf)/Erythroblasts-ZBTB7A-ChIP-Seq(GSE74977)/Homer(0.748) More Information \| Similar Motifs Found | motif file (matrix) |
| 14 \* | A C G T G T A C A C G T A G T C A C T G A G T C C T A G A C T G | 1e-7 | -1.714e+01 | 5.16% | 2.48% | 381.6bp (422.5bp) | ZBTB33/MA0527.1/Jaspar(0.795) More Information \| Similar Motifs Found | motif file (matrix) |
| 15 \* | A C G T A C T G A C T G A C T G A C T G A G T C C G T A A C G T C G T A A C G T | 1e-7 | -1.613e+01 | 0.70% | 0.06% | 146.0bp (431.4bp) | PB0143.1\_Klf7\_2/Jaspar(0.718) More Information \| Similar Motifs Found | motif file (matrix) |
| 16 \* | A G T C A G T C A G T C A G T C A C G T C G T A A C G T A G T C C G T A A C T G | 1e-6 | -1.576e+01 | 0.55% | 0.03% | 362.5bp (162.2bp) | YER130C/MA0423.1/Jaspar(0.805) More Information \| Similar Motifs Found | motif file (matrix) |
| 17 \* | A C G T C G T A A C G T C G T A A G T C C G T A A G T C A C T G | 1e-6 | -1.416e+01 | 4.77% | 2.41% | 308.1bp (388.6bp) | MATALPHA2/MA0328.2/Jaspar(0.837) More Information \| Similar Motifs Found | motif file (matrix) |
| 18 \* | A C G T C G T A A C G T C G T A A C T G A C G T A C T G A C T G A C T G A C T G | 1e-5 | -1.309e+01 | 0.55% | 0.06% | 470.3bp (244.1bp) | MIG1(MacIsaac)/Yeast(0.775) More Information \| Similar Motifs Found | motif file (matrix) |
| 19 \* | C A T G A C G T C T A G C T A G C A G T C T A G C T A G G A C T C A T G A C T G | 1e-5 | -1.213e+01 | 5.16% | 2.86% | 507.0bp (376.2bp) | Znf281/MA1630.1/Jaspar(0.714) More Information \| Similar Motifs Found | motif file (matrix) |
| 20 \* | A C G T A C T G A C G T A C T G A G T C A C G T C G T A A C T G | 1e-4 | -1.037e+01 | 2.74% | 1.28% | 306.7bp (399.8bp) | NCU08034(RRM)/Neurospora\_crassa-RNCMPT00209-PBM/HughesRNA(0.875) More Information \| Similar Motifs Found | motif file (matrix) |
| 21 \* | C G T A A C G T C G T A C G T A A C T G A C G T C G T A A C G T A C G T A G T C | 1e-3 | -7.795e+00 | 0.47% | 0.08% | 289.9bp (205.1bp) | br-Z4/dmmpmm(SeSiMCMC)/fly(0.754) More Information \| Similar Motifs Found | motif file (matrix) |
| 22 \* | C G T A A G T C A C T G C G T A C G T A C G T A A C G T C G T A A C G T A C G T | 1e-2 | -5.215e+00 | 0.63% | 0.22% | 361.1bp (511.9bp) | cad/dmmpmm(Down)/fly(0.735) More Information \| Similar Motifs Found | motif file (matrix) |
