## Supplemental Table 1 for "The Establishment of Cell-Type Specific Gene Regulation in the Sea Urchin Embryo": Hpf40_Midgut_homerResults.html

homer\_motifs\_wbackground/Hpf40\_Midgut/ - Homer de novo Motif Results


### Homer *de novo* Motif Results (homer\_motifs\_wbackground/Hpf40\_Midgut/)

Known Motif Enrichment Results  
Gene Ontology Enrichment Results  
If Homer is having trouble matching a motif to a known motif, try copy/pasting the matrix file into
STAMP  
More information on motif finding results: HOMER
| Description of Results
| Tips
  
Total target sequences = 834  
Total background sequences = 7421  
\* - possible false positive  

|  |  |  |  |  |  |  |  |  |
| --- | --- | --- | --- | --- | --- | --- | --- | --- |
| Rank | Motif | P-value | log P-pvalue | % of Targets | % of Background | STD(Bg STD) | Best Match/Details | Motif File |
| 1 | G C A T T C G A C G A T C T A G G A T C G C T A G T C A G C T A G A T C C T G A | 1e-22 | -5.268e+01 | 56.83% | 39.69% | 343.1bp (432.8bp) | fkh/dmmpmm(Noyes)/fly(0.910) More Information \| Similar Motifs Found | motif file (matrix) |
| 2 | C T G A C T G A A G C T T A C G C G T A C A G T A C G T C T G A G C T A A G T C | 1e-21 | -4.852e+01 | 8.75% | 2.29% | 278.5bp (323.6bp) | HNF1b(Homeobox)/PDAC-HNF1B-ChIP-Seq(GSE64557)/Homer(0.915) More Information \| Similar Motifs Found | motif file (matrix) |
| 3 | G T C A G C T A T G A C A G T C C G A T T C A G G T C A A C G T C G T A G C T A | 1e-14 | -3.294e+01 | 32.13% | 20.62% | 361.6bp (457.4bp) | pqm-1/MA1703.1/Jaspar(0.874) More Information \| Similar Motifs Found | motif file (matrix) |
| 4 | G C A T A G C T A C T G A C T G C G T A A G T C C G A T A C G T A G C T A C T G | 1e-12 | -2.959e+01 | 5.40% | 1.46% | 280.1bp (406.9bp) | HNF4A/MA0114.4/Jaspar(0.969) More Information \| Similar Motifs Found | motif file (matrix) |
| 5 \* | A T C G A C T G G C A T A G C T C T A G A G T C A C G T A G C T | 1e-11 | -2.570e+01 | 46.88% | 35.42% | 384.9bp (421.4bp) | YAP5(MacIsaac)/Yeast(0.722) More Information \| Similar Motifs Found | motif file (matrix) |
| 6 \* | C G T A A G T C A C G T C T A G A C T G G A C T C A G T G T C A A G C T C T G A | 1e-10 | -2.306e+01 | 6.12% | 2.18% | 316.5bp (400.5bp) | br-Z4/dmmpmm(Bigfoot)/fly(0.703) More Information \| Similar Motifs Found | motif file (matrix) |
| 7 \* | G C A T G T C A C A G T G T A C C G A T A G T C G A C T A C T G T A C G C G A T | 1e-9 | -2.267e+01 | 10.55% | 5.07% | 297.2bp (419.0bp) | SRSF10(RRM)/Homo\_sapiens-RNCMPT00090-PBM/HughesRNA(0.705) More Information \| Similar Motifs Found | motif file (matrix) |
| 8 \* | A C G T A G T C G T C A C G T A C G T A A T G C A C G T A T G C C T G A A C G T | 1e-9 | -2.247e+01 | 4.44% | 1.29% | 400.7bp (597.1bp) | pan/dmmpmm(Papatsenko)/fly(0.692) More Information \| Similar Motifs Found | motif file (matrix) |
| 9 \* | C A G T G T A C A C G T T G A C A T C G C G A T A C T G C G T A A T C G A C G T | 1e-9 | -2.220e+01 | 4.80% | 1.48% | 337.6bp (443.0bp) | MSANTD3/MA1523.1/Jaspar(0.713) More Information \| Similar Motifs Found | motif file (matrix) |
| 10 \* | A C G T A C G T A C T G A C G T A G T C A C T G A G T C A C G T A G T C A C T G | 1e-8 | -1.981e+01 | 0.72% | 0.03% | 329.7bp (46.2bp) | MAC1(MacIsaac)/Yeast(0.770) More Information \| Similar Motifs Found | motif file (matrix) |
| 11 \* | C G T A C G T A A G T C A C T G A C T G A C G T A G T C A C G T A C G T C G T A | 1e-8 | -1.970e+01 | 1.20% | 0.09% | 270.8bp (157.0bp) | MYB3R1(MYB)/col-MYB3R1-DAP-Seq(GSE60143)/Homer(0.746) More Information \| Similar Motifs Found | motif file (matrix) |
| 12 \* | C G T A A G T C A C T G A G T C C G T A C G T A C G T A A C T G A G T C C G T A | 1e-7 | -1.645e+01 | 0.84% | 0.05% | 409.6bp (325.9bp) | FHL1/MA0295.1/Jaspar(0.731) More Information \| Similar Motifs Found | motif file (matrix) |
| 13 \* | C G T A A G T C C G T A A C G T A G T C A G T C C G T A A C T G A C T G A C G T | 1e-6 | -1.574e+01 | 0.72% | 0.03% | 165.2bp (109.5bp) | SPDEF/MA0686.1/Jaspar(0.802) More Information \| Similar Motifs Found | motif file (matrix) |
| 14 \* | A T G C A C T G C T G A C T A G C G A T A C T G C G T A A C T G A T C G G A T C | 1e-6 | -1.558e+01 | 2.52% | 0.64% | 293.1bp (553.8bp) | Rbm24(RRM)/Tetraodon\_nigroviridis-RNCMPT00285-PBM/HughesRNA(0.713) More Information \| Similar Motifs Found | motif file (matrix) |
| 15 \* | C G T A A C T G C G T A A C G T C T G A A C T G A T G C C G A T C G T A C G T A | 1e-5 | -1.217e+01 | 1.92% | 0.49% | 262.3bp (388.8bp) | HNRNPAB(RRM)/Tetraodon\_nigroviridis-RNCMPT00245-PBM/HughesRNA(0.743) More Information \| Similar Motifs Found | motif file (matrix) |
| 16 \* | A G C T C A T G A G T C C T A G T A C G A G T C T A G C T C A G A G T C G A T C | 1e-5 | -1.177e+01 | 6.12% | 3.14% | 336.8bp (416.4bp) | ERF2(AP2EREBP)/colamp-ERF2-DAP-Seq(GSE60143)/Homer(0.773) More Information \| Similar Motifs Found | motif file (matrix) |
| 17 \* | A G T C A C T G C G T A A G T C A C G T C G T A A C T G A G T C | 1e-4 | -1.068e+01 | 1.32% | 0.27% | 423.0bp (445.7bp) | ARF4/MA1697.1/Jaspar(0.700) More Information \| Similar Motifs Found | motif file (matrix) |
| 18 \* | A C T G A C G T A C G T A C G T A C G T C G T A C G T A A C G T A G T C A C T G | 1e-4 | -1.051e+01 | 0.60% | 0.05% | 764.0bp (228.2bp) | CG34031/dmmpmm(Noyes\_hd)/fly(0.753) More Information \| Similar Motifs Found | motif file (matrix) |
| 19 \* | C G T A G T C A A C T G A C T G A C G T A G T C A C G T C G T A A C T G A C G T | 1e-2 | -5.983e+00 | 0.84% | 0.22% | 287.5bp (457.2bp) | NR6A1/MA1541.1/Jaspar(0.690) More Information \| Similar Motifs Found | motif file (matrix) |
| 20 \* | G T A C G C A T T A G C A C G T A G T C C G A T T C A G A G C T A G T C G A C T | 1e-1 | -3.559e+00 | 4.92% | 3.59% | 400.9bp (409.9bp) | GAGA-repeat/Arabidopsis-Promoters/Homer(0.804) More Information \| Similar Motifs Found | motif file (matrix) |
