## Supplemental Table 1 for "The Establishment of Cell-Type Specific Gene Regulation in the Sea Urchin Embryo": Hpf40_NeuronSERO_homerResults.html

homer\_motifs\_wbackground/Hpf40\_NeuronSERO/ - Homer de novo Motif Results


### Homer *de novo* Motif Results (homer\_motifs\_wbackground/Hpf40\_NeuronSERO/)

Known Motif Enrichment Results  
Gene Ontology Enrichment Results  
If Homer is having trouble matching a motif to a known motif, try copy/pasting the matrix file into
STAMP  
More information on motif finding results: HOMER
| Description of Results
| Tips
  
Total target sequences = 439  
Total background sequences = 7873  
\* - possible false positive  

|  |  |  |  |  |  |  |  |  |
| --- | --- | --- | --- | --- | --- | --- | --- | --- |
| Rank | Motif | P-value | log P-pvalue | % of Targets | % of Background | STD(Bg STD) | Best Match/Details | Motif File |
| 1 \* | G A T C A C T G A G T C C T G A A C T G C G T A A T G C C T A G G C T A A C G T | 1e-9 | -2.239e+01 | 7.74% | 2.13% | 387.3bp (405.0bp) | Adf1/dmmpmm(Pollard)/fly(0.767) More Information \| Similar Motifs Found | motif file (matrix) |
| 2 \* | C G T A A C T G G A C T A C G T C G T A C A T G A C T G G T A C A C T G A C G T | 1e-9 | -2.083e+01 | 3.64% | 0.50% | 356.5bp (402.5bp) | MYB116(MYB)/colamp-MYB116-DAP-Seq(GSE60143)/Homer(0.777) More Information \| Similar Motifs Found | motif file (matrix) |
| 3 \* | A G C T A C G T G A C T A C T G G T C A T C G A G A T C A G T C T C G A A C G T | 1e-8 | -1.930e+01 | 11.39% | 4.56% | 394.3bp (500.1bp) | FOS/MA0476.1/Jaspar(0.671) More Information \| Similar Motifs Found | motif file (matrix) |
| 4 \* | A C G T C G T A C T G A A C T G A C T G A G T C A G T C C G T A A C G T C G T A | 1e-7 | -1.661e+01 | 1.82% | 0.12% | 276.0bp (362.8bp) | SKN7/MA0381.1/Jaspar(0.687) More Information \| Similar Motifs Found | motif file (matrix) |
| 5 \* | A C G T A G C T A G T C C G T A C T G A A G T C C T G A A G T C C G T A A C T G | 1e-7 | -1.639e+01 | 4.10% | 0.86% | 388.7bp (353.5bp) | BRUNOL6(RRM)/Homo\_sapiens-RNCMPT00187-PBM/HughesRNA(0.714) More Information \| Similar Motifs Found | motif file (matrix) |
| 6 \* | G T C A A C G T G T C A C G T A C G T A A G T C C T A G A G T C C T A G A G C T | 1e-6 | -1.576e+01 | 4.10% | 0.90% | 252.7bp (479.1bp) | CAMTA1(CAMTA)/col-CAMTA1-DAP-Seq(GSE60143)/Homer(0.815) More Information \| Similar Motifs Found | motif file (matrix) |
| 7 \* | A G T C A G C T A C T G A G T C A C T G C G T A C T G A G T A C C G T A A C T G | 1e-6 | -1.521e+01 | 2.05% | 0.20% | 592.5bp (354.1bp) | CHA4(MacIsaac)/Yeast(0.673) More Information \| Similar Motifs Found | motif file (matrix) |
| 8 \* | A G C T A T G C A C T G A T C G G T A C A T C G A T C G C G T A A T C G A G C T | 1e-5 | -1.354e+01 | 3.19% | 0.64% | 287.6bp (378.6bp) | CHA4/MA0283.1/Jaspar(0.787) More Information \| Similar Motifs Found | motif file (matrix) |
| 9 \* | A C G T A G T C A G T C C G T A A G T C A C G T A C G T A C T G A C T G A G T C | 1e-4 | -1.142e+01 | 1.14% | 0.07% | 225.0bp (258.0bp) | BHLH112/MA0961.1/Jaspar(0.821) More Information \| Similar Motifs Found | motif file (matrix) |
| 10 \* | A G C T G C A T T G A C C G A T T C G A G T A C A T C G C G A T C A G T A C T G | 1e-4 | -1.044e+01 | 14.58% | 8.65% | 251.8bp (432.5bp) | ZNF136/MA1588.1/Jaspar(0.652) More Information \| Similar Motifs Found | motif file (matrix) |
| 11 \* | C T G A C T G A C T G A A T G C C G T A G T A C A C G T A C T G G T A C A G T C | 1e-4 | -9.530e+00 | 2.96% | 0.81% | 381.1bp (349.1bp) | PB0195.1\_Zbtb3\_2/Jaspar(0.693) More Information \| Similar Motifs Found | motif file (matrix) |
| 12 \* | C G T A A C T G C G T A A C G T C G T A A C T G A C G T C G T A A T G C G C A T | 1e-3 | -8.515e+00 | 0.91% | 0.07% | 137.2bp (376.6bp) | HNRNPAB(RRM)/Tetraodon\_nigroviridis-RNCMPT00245-PBM/HughesRNA(0.793) More Information \| Similar Motifs Found | motif file (matrix) |
| 13 \* | G T A C G C T A T A G C A G C T A G T C G T C A A G T C G C A T A G T C T G C A | 1e-3 | -8.342e+00 | 10.71% | 6.23% | 310.3bp (468.9bp) | Egr1(Zf)/K562-Egr1-ChIP-Seq(GSE32465)/Homer(0.790) More Information \| Similar Motifs Found | motif file (matrix) |
| 14 \* | A T C G C T A G C G T A C T A G C T G A C T G A A C G T G T C A G T C A C A T G | 1e-3 | -8.061e+00 | 12.98% | 8.11% | 344.5bp (422.2bp) | Unknown4/Arabidopsis-Promoters/Homer(0.656) More Information \| Similar Motifs Found | motif file (matrix) |
| 15 \* | C T G A C G A T A G T C T G A C G A C T T C G A T G C A G C A T G C A T C G T A | 1e-3 | -7.975e+00 | 19.82% | 13.84% | 323.2bp (418.8bp) | VSX2/MA0726.1/Jaspar(0.821) More Information \| Similar Motifs Found | motif file (matrix) |
| 16 \* | C G T A C G A T C G T A C G T A C G T A A C G T A C G T A G T C | 1e-3 | -7.944e+00 | 7.06% | 3.61% | 267.1bp (427.5bp) | abd-A/dmmpmm(Down)/fly(0.812) More Information \| Similar Motifs Found | motif file (matrix) |
| 17 \* | C T A G A G T C G A C T A C G T A C T G A G T C T G C A A C G T A C T G A G T C | 1e-3 | -7.419e+00 | 11.39% | 7.04% | 286.2bp (466.0bp) | HNRPLL(RRM)/Homo\_sapiens-RNCMPT00178-PBM/HughesRNA(0.766) More Information \| Similar Motifs Found | motif file (matrix) |
| 18 \* | A C T G A C T G C G T A C G T A A G T C A G T C A C G T A G T C A C G T A G T C | 1e-2 | -6.461e+00 | 0.68% | 0.06% | 131.2bp (520.0bp) | NRG1/NRG1\_H2O2Hi/[](Harbison)/Yeast(0.654) More Information \| Similar Motifs Found | motif file (matrix) |
| 19 \* | A C G T A C T G A C T G C G T A A C T G C G T A A C T G A C G T A C G T C G T A | 1e-2 | -6.461e+00 | 0.68% | 0.06% | 675.6bp (331.4bp) | ZNF652/MA1657.1/Jaspar(0.684) More Information \| Similar Motifs Found | motif file (matrix) |
| 20 \* | C G T A A G T C A G T C A G T C A C T G C G T A A C T G A C G T A C G T A C T G | 1e-2 | -5.832e+00 | 0.68% | 0.06% | 562.6bp (181.2bp) | RDS2/MA0362.1/Jaspar(0.666) More Information \| Similar Motifs Found | motif file (matrix) |
| 21 \* | A G T C A T C G A C G T A G T C C T A G A T G C A C T G G T A C | 1e-2 | -5.575e+00 | 2.73% | 1.11% | 302.6bp (405.1bp) | RSF1(RRM)/Drosophila\_melanogaster-RNCMPT00061-PBM/HughesRNA(0.763) More Information \| Similar Motifs Found | motif file (matrix) |
| 22 \* | A C T G A G T C C G T A A C G T C T G A A C G T A C T G C G T A A G T C A G T C | 1e-2 | -4.788e+00 | 0.91% | 0.19% | 190.6bp (355.3bp) | NEUROG1/MA0623.2/Jaspar(0.702) More Information \| Similar Motifs Found | motif file (matrix) |
| 23 \* | A C G T A G T C A C T G A C G T A C T G A T G C A C G T A G T C A C G T C G T A | 1e-1 | -4.379e+00 | 0.46% | 0.04% | 97.0bp (435.0bp) | kni/MA0451.1/Jaspar(0.718) More Information \| Similar Motifs Found | motif file (matrix) |
| 24 \* | A C G T C T A G A G C T A C T G G A T C T C A G A G T C C T A G | 1e-1 | -3.552e+00 | 15.72% | 12.54% | 249.3bp (409.3bp) | ZBTB14/MA1650.1/Jaspar(0.746) More Information \| Similar Motifs Found | motif file (matrix) |
| 25 \* | G T C A C G A T C G A T C G T A A C G T C G A T T G C A A G T C A G C T C G T A | 1e-1 | -3.509e+00 | 9.57% | 7.09% | 298.2bp (422.2bp) | Antp/dmmpmm(Down)/fly(0.785) More Information \| Similar Motifs Found | motif file (matrix) |
| 26 \* | A C G T A C T G A C G T A C G T A G T C A C G T A C T G C G T A A G T C A C G T | 1e-1 | -3.430e+00 | 0.46% | 0.07% | 43.4bp (250.4bp) | Ot\_0263(RRM)/Ostreococcus\_tauri-RNCMPT00263-PBM/HughesRNA(0.743) More Information \| Similar Motifs Found | motif file (matrix) |
| 27 \* | A G T C C G T A A C G T A G T C C G T A A C G T A G T C C G T A A C G T A G T C | 1e0 | -1.166e+00 | 3.19% | 2.73% | 359.6bp (431.9bp) | ZML2(C2C2gata)/col-ZML2-DAP-Seq(GSE60143)/Homer(0.956) More Information \| Similar Motifs Found | motif file (matrix) |
