## Supplemental Table 1 for "The Establishment of Cell-Type Specific Gene Regulation in the Sea Urchin Embryo": Hpf40_PigmentCells_homerResults.html

homer\_motifs\_wbackground/Hpf40\_PigmentCells/ - Homer de novo Motif Results


### Homer *de novo* Motif Results (homer\_motifs\_wbackground/Hpf40\_PigmentCells/)

Known Motif Enrichment Results  
Gene Ontology Enrichment Results  
If Homer is having trouble matching a motif to a known motif, try copy/pasting the matrix file into
STAMP  
More information on motif finding results: HOMER
| Description of Results
| Tips
  
Total target sequences = 568  
Total background sequences = 7955  
\* - possible false positive  

|  |  |  |  |  |  |  |  |  |
| --- | --- | --- | --- | --- | --- | --- | --- | --- |
| Rank | Motif | P-value | log P-pvalue | % of Targets | % of Background | STD(Bg STD) | Best Match/Details | Motif File |
| 1 | C A T G A C G T T C A G G T A C C G A T A C T G A T C G A G C T G T C A A G T C | 1e-74 | -1.719e+02 | 31.16% | 6.05% | 372.4bp (438.1bp) | ACE2/ACE2\_YPD/2-SWI5(Harbison)/Yeast(0.879) More Information \| Similar Motifs Found | motif file (matrix) |
| 2 \* | T A C G A T C G T G C A T A G C G C T A C A T G A G C T C G T A C G A T T A C G | 1e-11 | -2.550e+01 | 2.64% | 0.23% | 372.7bp (395.2bp) | ZBTB32/MA1580.1/Jaspar(0.725) More Information \| Similar Motifs Found | motif file (matrix) |
| 3 \* | C G T A C G A T G A C T G A C T T A G C C A T G C T A G A T C G A T C G T A G C | 1e-9 | -2.234e+01 | 13.91% | 6.49% | 388.9bp (398.8bp) | RDS2/MA0362.1/Jaspar(0.712) More Information \| Similar Motifs Found | motif file (matrix) |
| 4 \* | T C A G A G T C T A G C T C G A A T C G A G T C G T A C G A T C T C A G C A G T | 1e-9 | -2.167e+01 | 6.34% | 1.87% | 264.2bp (411.4bp) | Vts1p(SAM)/Saccharomyces\_cerevisiae-RNCMPT00082-PBM/HughesRNA(0.744) More Information \| Similar Motifs Found | motif file (matrix) |
| 5 \* | G T A C A G C T C G T A G C A T C T A G A G C T A G T C C T G A T G A C A T G C | 1e-8 | -1.965e+01 | 6.34% | 2.02% | 477.3bp (379.1bp) | RPN4/RPN4\_H2O2Lo/[](Harbison)/Yeast(0.643) More Information \| Similar Motifs Found | motif file (matrix) |
| 6 \* | T G A C A T G C G T C A A T G C C T G A C G T A G T C A C G T A C G T A A C G T | 1e-8 | -1.869e+01 | 6.87% | 2.40% | 336.0bp (494.4bp) | HNRNPC(RRM)/Homo\_sapiens-RNCMPT00025-PBM/HughesRNA(0.775) More Information \| Similar Motifs Found | motif file (matrix) |
| 7 \* | G A C T C G T A A C T G C A G T C A G T G A C T G C A T A G T C G T A C C G T A | 1e-8 | -1.858e+01 | 10.74% | 4.85% | 338.6bp (424.6bp) | NFATC2/MA0152.1/Jaspar(0.688) More Information \| Similar Motifs Found | motif file (matrix) |
| 8 \* | A G T C A C G T C G T A G T C A A C G T C G T A A C G T C G T A C G T A A C T G | 1e-7 | -1.841e+01 | 1.06% | 0.04% | 144.0bp (317.5bp) | SHEP(RRM)/Drosophila\_melanogaster-RNCMPT00174-PBM/HughesRNA(0.794) More Information \| Similar Motifs Found | motif file (matrix) |
| 9 \* | G T A C T G C A G T C A G T C A A C G T A G C T A C G T T C A G T C G A A C G T | 1e-6 | -1.548e+01 | 7.04% | 2.82% | 324.9bp (521.6bp) | pan/dmmpmm(Bigfoot)/fly(0.786) More Information \| Similar Motifs Found | motif file (matrix) |
| 10 \* | T C G A A C G T C T G A G C A T T A G C G C T A C G A T C A T G C G T A C G A T | 1e-6 | -1.517e+01 | 28.70% | 19.83% | 379.6bp (435.0bp) | GAT3(MacIsaac)/Yeast(0.773) More Information \| Similar Motifs Found | motif file (matrix) |
| 11 \* | A C T G C G T A A C G T A C T G A C T G A G T C A G T C A G T C A C G T A G T C | 1e-6 | -1.466e+01 | 0.88% | 0.03% | 150.2bp (120.0bp) | pho/MA1460.1/Jaspar(0.691) More Information \| Similar Motifs Found | motif file (matrix) |
| 12 \* | A C T G C G T A A C G T C G T A A G T C A G T C A G T C C G T A A C G T A C G T | 1e-6 | -1.437e+01 | 1.06% | 0.05% | 223.5bp (188.3bp) | PB0059.1\_Six6\_1/Jaspar(0.770) More Information \| Similar Motifs Found | motif file (matrix) |
| 13 \* | A C G T C G T A C G T A C G T A A C T G C G T A A C G T A C G T A C T G A G T C | 1e-5 | -1.309e+01 | 1.06% | 0.06% | 148.4bp (343.6bp) | HAHB4(HD-ZIP)/Helianthus anuus/AthaMap(0.739) More Information \| Similar Motifs Found | motif file (matrix) |
| 14 \* | A G T C A C G T A G T C C G T A A C T G A G T C A G T C A C G T A T G C A C T G | 1e-5 | -1.206e+01 | 1.06% | 0.08% | 159.7bp (550.4bp) | CRZ1(MacIsaac)/Yeast(0.767) More Information \| Similar Motifs Found | motif file (matrix) |
| 15 \* | A C G T A C G T A C T G A C G T A C G T A G T C A C G T C G T A A C T G C G T A | 1e-4 | -1.045e+01 | 1.06% | 0.11% | 167.7bp (229.4bp) | ZBTB12(Zf)/HEK293-ZBTB12.GFP-ChIP-Seq(GSE58341)/Homer(0.829) More Information \| Similar Motifs Found | motif file (matrix) |
| 16 \* | A C T G C G T A C G T A A C T G A C G T A G T C A G T C A C G T C G T A A C G T | 1e-4 | -9.524e+00 | 0.70% | 0.04% | 314.3bp (177.6bp) | ZmHOX2a(1)(HD-HOX)/Zea mays/AthaMap(0.675) More Information \| Similar Motifs Found | motif file (matrix) |
| 17 \* | G C A T A G C T G C A T A C T G A G C T G T A C C A T G C T G A T G C A T G C A | 1e-3 | -7.312e+00 | 4.23% | 2.02% | 337.6bp (391.0bp) | PB0032.1\_IRC900814\_1/Jaspar(0.718) More Information \| Similar Motifs Found | motif file (matrix) |
| 18 \* | A G T C C G T A A G T C A C T G A C T G A G T C C G T A A G T C C G T A G T C A | 1e-2 | -6.580e+00 | 0.53% | 0.05% | 121.0bp (284.0bp) | NAC92/MA1044.1/Jaspar(0.689) More Information \| Similar Motifs Found | motif file (matrix) |
| 19 \* | G A C T G A T C C G A T G A T C A G C T G A T C G C A T G T A C A G C T C A T G | 1e0 | -1.595e+00 | 16.20% | 14.87% | 431.2bp (371.5bp) | SeqBias: GA-repeat(0.855) More Information \| Similar Motifs Found | motif file (matrix) |
