## Supplemental Table 1 for "The Establishment of Cell-Type Specific Gene Regulation in the Sea Urchin Embryo": Hpf40_PMCs_homerResults.html

homer\_motifs\_wbackground/Hpf40\_PMCs/ - Homer de novo Motif Results


### Homer *de novo* Motif Results (homer\_motifs\_wbackground/Hpf40\_PMCs/)

Known Motif Enrichment Results  
Gene Ontology Enrichment Results  
If Homer is having trouble matching a motif to a known motif, try copy/pasting the matrix file into
STAMP  
More information on motif finding results: HOMER
| Description of Results
| Tips
  
Total target sequences = 338  
Total background sequences = 8111  
\* - possible false positive  

|  |  |  |  |  |  |  |  |  |
| --- | --- | --- | --- | --- | --- | --- | --- | --- |
| Rank | Motif | P-value | log P-pvalue | % of Targets | % of Background | STD(Bg STD) | Best Match/Details | Motif File |
| 1 \* | A C T G A C G T C G T A A C G T A G C T A C G T A G T C A C T G A G T C A C G T | 1e-9 | -2.117e+01 | 2.37% | 0.09% | 198.2bp (407.5bp) | SWI4(MacIsaac)/Yeast(0.616) More Information \| Similar Motifs Found | motif file (matrix) |
| 2 \* | C T G A G T A C A C T G A C T G G T A C A C G T A G C T A C T G A C T G C G T A | 1e-8 | -2.046e+01 | 5.93% | 1.08% | 247.5bp (468.7bp) | ZML1(C2C2gata)/colamp-ZML1-DAP-Seq(GSE60143)/Homer(0.703) More Information \| Similar Motifs Found | motif file (matrix) |
| 3 \* | A C T G A C T G C T A G A T C G T G C A A G C T A C T G A C T G G T A C T A G C | 1e-8 | -2.031e+01 | 3.56% | 0.32% | 254.2bp (434.8bp) | ZNF467(Zf)/HEK293-ZNF467.GFP-ChIP-Seq(GSE58341)/Homer(0.745) More Information \| Similar Motifs Found | motif file (matrix) |
| 4 \* | C G T A A C T G A C T G C G T A C G T A A C G T A C T G A C G T A C T G A C T G | 1e-7 | -1.649e+01 | 1.78% | 0.07% | 275.2bp (265.9bp) | TEAD2/MA1121.1/Jaspar(0.857) More Information \| Similar Motifs Found | motif file (matrix) |
| 5 \* | G T A C C T G A T C A G A G T C T C G A A C G T G T A C C G A T G A T C A G T C | 1e-7 | -1.633e+01 | 5.64% | 1.26% | 402.0bp (470.7bp) | PRDM9(Zf)/Testis-DMC1-ChIP-Seq(GSE35498)/Homer(0.674) More Information \| Similar Motifs Found | motif file (matrix) |
| 6 \* | G T C A A C T G C G T A C G T A A G C T A C T G A C G T A G T C C G T A A G C T | 1e-6 | -1.605e+01 | 4.15% | 0.67% | 428.8bp (609.2bp) | TEC1(MacIsaac)/Yeast(0.765) More Information \| Similar Motifs Found | motif file (matrix) |
| 7 \* | C T G A C T A G C T A G G A T C A G C T G T C A G A C T C T G A C G A T T A C G | 1e-6 | -1.605e+01 | 8.61% | 2.77% | 296.4bp (406.1bp) | MOT2/MA0379.1/Jaspar(0.711) More Information \| Similar Motifs Found | motif file (matrix) |
| 8 \* | T G C A G A T C T A G C C T A G T G A C G T A C G C A T G T A C C G A T C T G A | 1e-5 | -1.368e+01 | 12.17% | 5.38% | 276.1bp (391.2bp) | MSN2/MSN2\_H2O2Hi/1-MSN2(Harbison)/Yeast(0.710) More Information \| Similar Motifs Found | motif file (matrix) |
| 9 \* | A G T C G C T A C G T A T A C G C G T A A C G T C T G A A G C T C T G A A G T C | 1e-5 | -1.300e+01 | 12.46% | 5.73% | 319.1bp (424.5bp) | Cf2/dmmpmm(Bergman)/fly(0.726) More Information \| Similar Motifs Found | motif file (matrix) |
| 10 \* | A G T C C G T A C G T A C G T A C G T A A G T C C G T A C T A G A T G C C T G A | 1e-5 | -1.297e+01 | 5.93% | 1.73% | 225.0bp (474.0bp) | Pp\_0229(RRM)/Physcomitrella\_patens-RNCMPT00229-PBM/HughesRNA(0.769) More Information \| Similar Motifs Found | motif file (matrix) |
| 11 \* | T G A C G C A T T G C A T A C G G A C T A T G C T A G C A C T G G C A T T G C A | 1e-5 | -1.276e+01 | 7.12% | 2.40% | 277.1bp (373.1bp) | SPL5/MA1059.2/Jaspar(0.697) More Information \| Similar Motifs Found | motif file (matrix) |
| 12 \* | G T C A G T A C C T G A C T A G T C G A A G T C A T C G A T G C T G C A G A T C | 1e-5 | -1.173e+01 | 13.95% | 7.09% | 282.6bp (424.4bp) | PB0151.1\_Myf6\_2/Jaspar(0.819) More Information \| Similar Motifs Found | motif file (matrix) |
| 13 \* | A G T C C G T A A G T C C G T A A C G T C G A T A C T G A C G T | 1e-3 | -9.187e+00 | 18.40% | 11.39% | 319.4bp (411.9bp) | dsx/dmmpmm(Bergman)/fly(0.832) More Information \| Similar Motifs Found | motif file (matrix) |
| 14 \* | G T A C C T A G T G C A T A C G C A G T C T A G G C T A T C G A G T C A G A T C | 1e-3 | -8.558e+00 | 7.42% | 3.33% | 487.2bp (399.4bp) | STE12/MA0393.1/Jaspar(0.729) More Information \| Similar Motifs Found | motif file (matrix) |
| 15 \* | C G T A A C G T C G T A C G T A A G T C A C T G A C G T C G T A A C G T C G T A | 1e-3 | -8.273e+00 | 0.89% | 0.04% | 334.3bp (172.5bp) | NAC043/MA1045.1/Jaspar(0.736) More Information \| Similar Motifs Found | motif file (matrix) |
| 16 \* | A T G C C A T G G C T A A G C T G A C T A G T C T C G A C A G T G C T A G T C A | 1e-3 | -7.908e+00 | 17.80% | 11.44% | 241.4bp (442.5bp) | unc-86/MA0926.1/Jaspar(0.706) More Information \| Similar Motifs Found | motif file (matrix) |
| 17 \* | G T A C T C G A A G T C A C T G A C G T A C T G C G T A A G T C | 1e-3 | -7.865e+00 | 8.90% | 4.52% | 264.4bp (413.7bp) | CBF1(MacIsaac)/Yeast(0.990) More Information \| Similar Motifs Found | motif file (matrix) |
| 18 \* | C A T G A C T G A C T G A G T C C T A G T A C G G A T C G T C A | 1e-3 | -7.026e+00 | 19.88% | 13.61% | 318.2bp (442.7bp) | NtERF2(AP2/EREBP)/Nicotiana tabacum/AthaMap(0.792) More Information \| Similar Motifs Found | motif file (matrix) |
| 19 \* | C G T A C G T A A C T G G T C A A C G T A G T C C G T A A C T G | 1e-2 | -6.149e+00 | 5.04% | 2.28% | 411.2bp (420.3bp) | ZmHOX2a(2)(HD-HOX)/Zea mays/AthaMap(0.776) More Information \| Similar Motifs Found | motif file (matrix) |
| 20 \* | A C T G A G T C G T C A A G T C A C T G C G T A A C T G A C G T C G T A A C G T | 1e-2 | -5.476e+00 | 0.89% | 0.10% | 199.3bp (427.0bp) | TUT1(RRM,Znf)/Homo\_sapiens-RNCMPT00075-PBM/HughesRNA(0.685) More Information \| Similar Motifs Found | motif file (matrix) |
| 21 \* | C G T A A G T C A G T C A G T C A C G T C G T A A G T C A G T C A C G T A G T C | 1e0 | -1.551e+00 | 0.30% | 0.07% | 58.4bp (401.9bp) | HRB98DE(RRM)/Drosophila\_melanogaster-RNCMPT00095-PBM/HughesRNA(0.825) More Information \| Similar Motifs Found | motif file (matrix) |
