## Supplemental Table 1 for "The Establishment of Cell-Type Specific Gene Regulation in the Sea Urchin Embryo": Hpf72_EctoApical_homerResults.html

homer\_motifs\_wbackground/Hpf72\_EctoApical/ - Homer de novo Motif Results


### Homer *de novo* Motif Results (homer\_motifs\_wbackground/Hpf72\_EctoApical/)

Known Motif Enrichment Results  
Gene Ontology Enrichment Results  
If Homer is having trouble matching a motif to a known motif, try copy/pasting the matrix file into
STAMP  
More information on motif finding results: HOMER
| Description of Results
| Tips
  
Total target sequences = 957  
Total background sequences = 15372  
\* - possible false positive  

|  |  |  |  |  |  |  |  |  |
| --- | --- | --- | --- | --- | --- | --- | --- | --- |
| Rank | Motif | P-value | log P-pvalue | % of Targets | % of Background | STD(Bg STD) | Best Match/Details | Motif File |
| 1 | A C G T C G T A A C G T A G C T A C T G C G T A A C G T A G C T | 1e-118 | -2.730e+02 | 54.65% | 20.49% | 327.9bp (443.8bp) | ONECUT1/MA0679.2/Jaspar(0.937) More Information \| Similar Motifs Found | motif file (matrix) |
| 2 | A C G T G A T C C A T G C T G A A C G T A G C T A T C G G A C T A G C T G T A C | 1e-16 | -3.765e+01 | 16.93% | 8.49% | 318.8bp (438.5bp) | SOX9/MA0077.1/Jaspar(0.834) More Information \| Similar Motifs Found | motif file (matrix) |
| 3 | A T G C C T G A G C T A C A G T T C G A T C A G T A G C G T A C A G T C G A T C | 1e-15 | -3.589e+01 | 14.42% | 6.87% | 340.2bp (412.3bp) | HAP1(MacIsaac)/Yeast(0.640) More Information \| Similar Motifs Found | motif file (matrix) |
| 4 | A T C G A C G T C A T G C G T A G T A C C G T A A T C G A G T C A G T C A G T C | 1e-12 | -2.891e+01 | 4.18% | 1.03% | 265.0bp (509.4bp) | Meis1(Homeobox)/MastCells-Meis1-ChIP-Seq(GSE48085)/Homer(0.785) More Information \| Similar Motifs Found | motif file (matrix) |
| 5 \* | A T G C T G C A C G T A C T A G C A T G C A T G A C T G A C T G G C A T C A T G | 1e-11 | -2.639e+01 | 27.59% | 18.47% | 356.1bp (457.8bp) | MSN4(MacIsaac)/Yeast(0.823) More Information \| Similar Motifs Found | motif file (matrix) |
| 6 \* | A G T C A G T C G A C T A C T G A C G T A G T C C G T A G C T A C A G T A T G C | 1e-11 | -2.631e+01 | 4.81% | 1.44% | 332.3bp (458.3bp) | Pbx3(Homeobox)/GM12878-PBX3-ChIP-Seq(GSE32465)/Homer(0.898) More Information \| Similar Motifs Found | motif file (matrix) |
| 7 \* | C T G A A G C T G C A T A T C G C G T A G A C T G A T C A G T C T G C A C A G T | 1e-11 | -2.624e+01 | 7.84% | 3.22% | 306.8bp (372.2bp) | CUX1(Homeobox)/K562-CUX1-ChIP-Seq(GSE92882)/Homer(0.764) More Information \| Similar Motifs Found | motif file (matrix) |
| 8 \* | A T C G A G C T C T A G A C T G G C A T C A T G G T C A C G T A T A G C A C T G | 1e-10 | -2.440e+01 | 12.43% | 6.55% | 391.1bp (476.5bp) | AGL55/MA1202.1/Jaspar(0.731) More Information \| Similar Motifs Found | motif file (matrix) |
| 9 \* | C A G T C T A G C G A T A G T C C T A G C T G A C T G A G T A C C G T A C A G T | 1e-9 | -2.118e+01 | 6.48% | 2.71% | 333.6bp (454.6bp) | MGA1/MA0336.1/Jaspar(0.657) More Information \| Similar Motifs Found | motif file (matrix) |
| 10 \* | C A T G C T A G A T G C A C G T G T A C G A T C A G C T G T A C | 1e-9 | -2.114e+01 | 27.48% | 19.34% | 404.3bp (469.8bp) | SRSF1(RRM)/Homo\_sapiens-RNCMPT00163-PBM/HughesRNA(0.781) More Information \| Similar Motifs Found | motif file (matrix) |
| 11 \* | A G T C A G T C A C G T A C G T C T A G A G T C A G T C A G T C | 1e-8 | -2.022e+01 | 15.57% | 9.47% | 373.5bp (449.5bp) | PCBP1(KH)/Mus\_musculus-RNCMPT00239-PBM/HughesRNA(0.757) More Information \| Similar Motifs Found | motif file (matrix) |
| 12 \* | C G T A C A G T A T G C A C G T A T G C A G T C G A T C G T C A A G C T G T A C | 1e-8 | -1.987e+01 | 4.60% | 1.65% | 346.3bp (463.8bp) | pho/MA1460.1/Jaspar(0.673) More Information \| Similar Motifs Found | motif file (matrix) |
| 13 \* | A G C T A C G T T A G C A C T G G T A C C G T A A T G C G A C T A C T G C G T A | 1e-8 | -1.929e+01 | 4.18% | 1.44% | 217.4bp (417.6bp) | STB4/STB4\_YPD/[](Harbison)/Yeast(0.686) More Information \| Similar Motifs Found | motif file (matrix) |
| 14 \* | G A C T G C T A C G T A A T G C C T G A C T G A A C T G G C A T T C A G G T A C | 1e-7 | -1.663e+01 | 18.60% | 12.55% | 375.7bp (438.8bp) | sna/MA0086.2/Jaspar(0.739) More Information \| Similar Motifs Found | motif file (matrix) |
| 15 \* | C G T A A C G T A T G C A C G T A G T C C G T A C G T A A C T G | 1e-6 | -1.506e+01 | 14.11% | 9.09% | 452.8bp (464.8bp) | vnd/dmmpmm(Noyes\_hd)/fly(0.792) More Information \| Similar Motifs Found | motif file (matrix) |
| 16 \* | A G T C C G T A A C G T A C G T A C T G A C G T | 1e-5 | -1.152e+01 | 63.22% | 56.37% | 357.8bp (432.5bp) | SOX13/MA1120.1/Jaspar(0.970) More Information \| Similar Motifs Found | motif file (matrix) |
| 17 \* | A G T C A C G T A C G T A C T G A G T C A C T G C G T A A G T C C G T A A C T G | 1e-4 | -1.004e+01 | 0.42% | 0.02% | 134.6bp (255.4bp) | CEBP:AP1(bZIP)/ThioMac-CEBPb-ChIP-Seq(GSE21512)/Homer(0.721) More Information \| Similar Motifs Found | motif file (matrix) |
| 18 \* | C G T A A C G T A C G T A G T C A C G T A G T C A C G T C G T A | 1e-3 | -8.828e+00 | 5.22% | 2.99% | 381.7bp (434.7bp) | RNP4F(RRM)/Drosophila\_melanogaster-RNCMPT00060-PBM/HughesRNA(0.788) More Information \| Similar Motifs Found | motif file (matrix) |
| 19 \* | A G T C A G T C A C T G C G T A A C T G A C G T A C G T A C T G A C T G A G T C | 1e-2 | -5.527e+00 | 0.31% | 0.03% | 552.0bp (787.9bp) | NFIX/MA0671.1/Jaspar(0.679) More Information \| Similar Motifs Found | motif file (matrix) |
| 20 \* | A C G T C G T A A G T C A G T C A G T C A T C G A T C G A C G T A G T C A G T C | 1e-2 | -4.866e+00 | 1.67% | 0.84% | 108.9bp (494.6bp) | REB1(MacIsaac)/Yeast(0.753) More Information \| Similar Motifs Found | motif file (matrix) |
