## Supplemental Table 1 for "The Establishment of Cell-Type Specific Gene Regulation in the Sea Urchin Embryo": Hpf72_EctoArboral_homerResults.html

homer\_motifs\_wbackground/Hpf72\_EctoArboral/ - Homer de novo Motif Results


### Homer *de novo* Motif Results (homer\_motifs\_wbackground/Hpf72\_EctoArboral/)

Known Motif Enrichment Results  
Gene Ontology Enrichment Results  
If Homer is having trouble matching a motif to a known motif, try copy/pasting the matrix file into
STAMP  
More information on motif finding results: HOMER
| Description of Results
| Tips
  
Total target sequences = 2552  
Total background sequences = 13714  
\* - possible false positive  

|  |  |  |  |  |  |  |  |  |
| --- | --- | --- | --- | --- | --- | --- | --- | --- |
| Rank | Motif | P-value | log P-pvalue | % of Targets | % of Background | STD(Bg STD) | Best Match/Details | Motif File |
| 1 | A G T C A C G T A T C G C G T A C G T A A C G T C T G A A C T G | 1e-85 | -1.975e+02 | 51.72% | 32.80% | 408.4bp (400.8bp) | vvl/dmmpmm(Bigfoot)/fly(0.761) More Information \| Similar Motifs Found | motif file (matrix) |
| 2 | C A T G A G C T A G T C G A T C C T G A A T G C A G C T A T C G T G C A T C G A | 1e-55 | -1.277e+02 | 63.32% | 47.79% | 415.8bp (425.4bp) | NKX2-5/MA0063.2/Jaspar(0.802) More Information \| Similar Motifs Found | motif file (matrix) |
| 3 | A G T C C G T A G T A C A C G T A C T G A C T G C G T A A C T G A G T C C T A G | 1e-46 | -1.076e+02 | 3.17% | 0.37% | 389.7bp (299.7bp) | tin/dmmpmm(SeSiMCMC)/fly(0.732) More Information \| Similar Motifs Found | motif file (matrix) |
| 4 | G A T C A T C G A G T C G C T A A G C T T A C G C T G A T C G A | 1e-46 | -1.063e+02 | 66.89% | 52.88% | 435.7bp (407.9bp) | ASD-1(RRM)/Caenorhabditis\_elegans-RNCMPT00180-PBM/HughesRNA(0.796) More Information \| Similar Motifs Found | motif file (matrix) |
| 5 | T C G A T C G A C T G A T C A G A T C G C G T A G C A T A G C T | 1e-45 | -1.042e+02 | 62.77% | 48.78% | 436.1bp (423.7bp) | Kr/dmmpmm(Papatsenko)/fly(0.856) More Information \| Similar Motifs Found | motif file (matrix) |
| 6 | A C T G A C T G C G T A T G A C A C G T A C G T C G A T A C G T C G T A A C T G | 1e-42 | -9.779e+01 | 6.97% | 2.06% | 388.4bp (383.7bp) | MYNN(Zf)/HEK293-MYNN.eGFP-ChIP-Seq(Encode)/Homer(0.772) More Information \| Similar Motifs Found | motif file (matrix) |
| 7 | C G T A C G T A A C G T A G T C A C G T A C T G C G T A A C T G A C T G A C T G | 1e-40 | -9.381e+01 | 1.65% | 0.08% | 335.4bp (287.7bp) | MET31/Literature(Harbison)/Yeast(0.668) More Information \| Similar Motifs Found | motif file (matrix) |
| 8 | A G T C A C G T A C G T A C G T A G T C G T C A T C G A C T A G | 1e-40 | -9.343e+01 | 34.87% | 23.12% | 415.4bp (434.8bp) | vnd/dmmpmm(Noyes\_hd)/fly(0.748) More Information \| Similar Motifs Found | motif file (matrix) |
| 9 | C G T A A C G T A G T C A C G T A G T C A C T G C G T A A C T G A C T G A G T C | 1e-36 | -8.307e+01 | 1.41% | 0.06% | 340.9bp (212.4bp) | XBP1(MacIsaac)/Yeast(0.833) More Information \| Similar Motifs Found | motif file (matrix) |
| 10 | A T C G C G T A A T C G A C G T A C T G A G T C | 1e-33 | -7.825e+01 | 52.00% | 39.98% | 429.0bp (403.0bp) | z/dmmpmm(Down)/fly(0.834) More Information \| Similar Motifs Found | motif file (matrix) |
| 11 | A G T C A C T G A C T G A C G T A C G T A G T C | 1e-31 | -7.221e+01 | 55.33% | 43.73% | 480.0bp (430.9bp) | OPI1/MA0349.1/Jaspar(0.967) More Information \| Similar Motifs Found | motif file (matrix) |
| 12 | A G T C A G C T A C G T A C T G A G T C A C G T A C T G A G C T A G T C A C G T | 1e-30 | -6.947e+01 | 1.72% | 0.16% | 354.1bp (271.5bp) | ARF34/MA1693.1/Jaspar(0.807) More Information \| Similar Motifs Found | motif file (matrix) |
| 13 | C G T A A G T C A G C T A C T G C G T A A C G T A T G C A G T C | 1e-30 | -6.912e+01 | 47.65% | 36.52% | 448.3bp (425.1bp) | Initiator/Drosophila-Promoters/Homer(0.771) More Information \| Similar Motifs Found | motif file (matrix) |
| 14 | C T G A G C T A C T G A A G T C A C G T A G T C G A C T G T A C | 1e-29 | -6.804e+01 | 64.03% | 52.87% | 447.1bp (415.7bp) | CG11360(KH)/Drosophila\_melanogaster-RNCMPT00129-PBM/HughesRNA(0.810) More Information \| Similar Motifs Found | motif file (matrix) |
| 15 | A G T C C G T A C G T A C G T A C G T A A C T G | 1e-29 | -6.781e+01 | 62.85% | 51.68% | 424.8bp (421.0bp) | Sf3b4(RRM)/Danio\_rerio-RNCMPT00224-PBM/HughesRNA(0.998) More Information \| Similar Motifs Found | motif file (matrix) |
| 16 | C G T A C T G A C G T A A C T G A C T G G T A C A C G T A G T C A C G T A G T C | 1e-29 | -6.760e+01 | 61.29% | 50.09% | 444.0bp (430.2bp) | dof42(C2C2dof)/col-dof42-DAP-Seq(GSE60143)/Homer(0.656) More Information \| Similar Motifs Found | motif file (matrix) |
| 17 | C T G A A C T G A C G T A G T C A G T C A G T C A C G T C G T A | 1e-27 | -6.421e+01 | 53.61% | 42.71% | 431.7bp (430.4bp) | HNRNPA2B1(RRM)/Homo\_sapiens-RNCMPT00024-PBM/HughesRNA(0.863) More Information \| Similar Motifs Found | motif file (matrix) |
| 18 | A G C T A C G T A C T G C G A T C T G A A G C T C G T A T G C A | 1e-26 | -6.183e+01 | 52.94% | 42.27% | 450.6bp (423.7bp) | Cf2-II/dmmpmm(Pollard)/fly(0.780) More Information \| Similar Motifs Found | motif file (matrix) |
| 19 | A G T C C T G A A G T C C G T A C G T A G C A T A C T G C T A G | 1e-25 | -5.839e+01 | 32.88% | 23.69% | 457.3bp (428.9bp) | PB0168.1\_Sox14\_2/Jaspar(0.884) More Information \| Similar Motifs Found | motif file (matrix) |
| 20 | A G T C A C T G A C G T C G T A C A T G A G T C C G T A G T C A C G A T A T G C | 1e-24 | -5.664e+01 | 2.98% | 0.69% | 336.6bp (376.5bp) | PB0055.1\_Rfx4\_1/Jaspar(0.703) More Information \| Similar Motifs Found | motif file (matrix) |
| 21 | A C G T A T G C C G T A A T G C C G A T A G T C A C T G C G A T C G T A C G A T | 1e-24 | -5.637e+01 | 2.82% | 0.62% | 377.0bp (421.7bp) | NAC055/MA0937.1/Jaspar(0.726) More Information \| Similar Motifs Found | motif file (matrix) |
| 22 | A C T G A C T G C G T A A C T G A C G T A C T G A C T G A C G T A C G T A C T G | 1e-24 | -5.616e+01 | 1.29% | 0.11% | 302.0bp (452.6bp) | MYB94(MYB)/col-MYB94-DAP-Seq(GSE60143)/Homer(0.704) More Information \| Similar Motifs Found | motif file (matrix) |
| 23 | A G T C A C G T C T A G A C T G A C G T A G T C C G T A C G T A A C T G C G T A | 1e-23 | -5.520e+01 | 1.49% | 0.15% | 401.0bp (527.0bp) | WRKY63/MA1092.1/Jaspar(0.761) More Information \| Similar Motifs Found | motif file (matrix) |
| 24 | A C T G A G C T A C G T C G T A A G T C A T C G A C T G A C G T | 1e-21 | -4.918e+01 | 11.48% | 6.35% | 457.3bp (382.8bp) | OSR1/MA1542.1/Jaspar(0.763) More Information \| Similar Motifs Found | motif file (matrix) |
| 25 | A C G T C G T A A C T G A C G T A C G T C G A T C G T A C G T A A C T G A G T C | 1e-21 | -4.879e+01 | 1.53% | 0.20% | 355.0bp (279.6bp) | br(var.3)/MA0012.1/Jaspar(0.744) More Information \| Similar Motifs Found | motif file (matrix) |
| 26 | A C T G C G T A A G T C A C T G A C G T A G T C A G T C A C G T | 1e-20 | -4.781e+01 | 5.92% | 2.49% | 408.5bp (405.7bp) | TGA10(bZIP)/colamp-TGA10-DAP-Seq(GSE60143)/Homer(0.816) More Information \| Similar Motifs Found | motif file (matrix) |
| 27 | A G T C A C G T A G T C A C G T C G T A C G T A A C G T A G T C A G T C C G T A | 1e-19 | -4.509e+01 | 60.97% | 51.94% | 422.3bp (424.2bp) | dve/MA0915.1/Jaspar(0.807) More Information \| Similar Motifs Found | motif file (matrix) |
| 28 | C G T A A C T G A G T C A C T G C G T A A C G T C G T A C G T A C G T A C G T A | 1e-17 | -4.056e+01 | 1.25% | 0.16% | 382.7bp (297.9bp) | PH0064.1\_Hoxb9/Jaspar(0.807) More Information \| Similar Motifs Found | motif file (matrix) |
| 29 | A C T G A C T G A C G T C G T A A C T G C G T A C G T A A C T G C G T A C G T A | 1e-15 | -3.490e+01 | 1.25% | 0.20% | 336.4bp (464.6bp) | PABPN1(RRM)/Homo\_sapiens-RNCMPT00157-PBM/HughesRNA(0.746) More Information \| Similar Motifs Found | motif file (matrix) |
| 30 | A C G T A C T G A G C T A C T G A G T C A C G T A G T C A C G T A C G T A C G T | 1e-13 | -3.114e+01 | 1.84% | 0.48% | 319.7bp (348.8bp) | NCU08034(RRM)/Neurospora\_crassa-RNCMPT00209-PBM/HughesRNA(0.718) More Information \| Similar Motifs Found | motif file (matrix) |
| 31 \* | A T C G C G T A A C T G A C T G A C G T A C G T C G T A A C G T | 1e-9 | -2.280e+01 | 6.66% | 3.98% | 467.9bp (460.2bp) | HRB27C(RRM)/Drosophila\_melanogaster-RNCMPT00028-PBM/HughesRNA(0.776) More Information \| Similar Motifs Found | motif file (matrix) |
