## Supplemental Table 1 for "The Establishment of Cell-Type Specific Gene Regulation in the Sea Urchin Embryo": Hpf72_EctoCiliaryBand_homerResults.html

homer\_motifs\_wbackground/Hpf72\_EctoCiliaryBand/ - Homer de novo Motif Results


### Homer *de novo* Motif Results (homer\_motifs\_wbackground/Hpf72\_EctoCiliaryBand/)

Known Motif Enrichment Results  
Gene Ontology Enrichment Results  
If Homer is having trouble matching a motif to a known motif, try copy/pasting the matrix file into
STAMP  
More information on motif finding results: HOMER
| Description of Results
| Tips
  
Total target sequences = 794  
Total background sequences = 15313  
\* - possible false positive  

|  |  |  |  |  |  |  |  |  |
| --- | --- | --- | --- | --- | --- | --- | --- | --- |
| Rank | Motif | P-value | log P-pvalue | % of Targets | % of Background | STD(Bg STD) | Best Match/Details | Motif File |
| 1 | C G T A C T G A C A G T T G A C A T G C A C T G G T A C G A C T G A C T G C T A | 1e-41 | -9.465e+01 | 37.28% | 17.12% | 328.7bp (453.2bp) | Pax7(Paired,Homeobox),long/Myoblast-Pax7-ChIP-Seq(GSE25064)/Homer(0.765) More Information \| Similar Motifs Found | motif file (matrix) |
| 2 | T A G C C T A G T A C G T G C A G C A T C G A T G C T A C A G T G A T C G T A C | 1e-25 | -5.858e+01 | 61.96% | 43.31% | 401.9bp (466.1bp) | DPRX/MA1480.1/Jaspar(0.853) More Information \| Similar Motifs Found | motif file (matrix) |
| 3 | C G T A C G T A A C T G A T G C A G T C G C A T C A G T G C T A | 1e-22 | -5.070e+01 | 42.32% | 26.27% | 382.5bp (473.3bp) | Nr2e3/MA0164.1/Jaspar(0.733) More Information \| Similar Motifs Found | motif file (matrix) |
| 4 | C T A G A C T G G C T A A C G T C A G T C G T A G C T A A C T G | 1e-20 | -4.646e+01 | 40.43% | 25.28% | 395.8bp (479.4bp) | Oc/dmmpmm(Noyes\_hd)/fly(0.905) More Information \| Similar Motifs Found | motif file (matrix) |
| 5 | T C G A A T C G G C T A A C T G G T A C A C G T A G C T C G T A | 1e-18 | -4.203e+01 | 35.52% | 21.75% | 330.1bp (449.5bp) | bcd-1/dmmpmm(Noyes)/fly(0.748) More Information \| Similar Motifs Found | motif file (matrix) |
| 6 | C T G A A C T G C T G A A C G T A C G T A C G T A C T G T A G C | 1e-15 | -3.669e+01 | 49.24% | 34.98% | 390.2bp (445.6bp) | ARR10/MA0121.1/Jaspar(0.805) More Information \| Similar Motifs Found | motif file (matrix) |
| 7 | A G T C A G C T C G T A A C G T A C G T A C T G G C A T A C T G | 1e-15 | -3.598e+01 | 53.65% | 39.32% | 413.6bp (425.2bp) | PB0168.1\_Sox14\_2/Jaspar(0.856) More Information \| Similar Motifs Found | motif file (matrix) |
| 8 | C T G A A C G T A G T C A G T C C A G T G C A T C G A T G A T C G T C A C G T A | 1e-13 | -3.069e+01 | 36.40% | 24.46% | 362.3bp (452.1bp) | RME1/MA0370.1/Jaspar(0.754) More Information \| Similar Motifs Found | motif file (matrix) |
| 9 \* | A C T G A T G C G T A C A G C T T G C A C G A T A C G T G T A C T C A G A C T G | 1e-11 | -2.664e+01 | 2.52% | 0.32% | 284.0bp (365.4bp) | TaMYB80(MYB)/Triticum aestivum/AthaMap(0.638) More Information \| Similar Motifs Found | motif file (matrix) |
| 10 \* | A G T C A C G T A C T G G A T C A G T C A G T C A C G T C G T A G T C A C A G T | 1e-10 | -2.407e+01 | 4.79% | 1.34% | 318.6bp (461.5bp) | TBP3(MYBrelated)/col-TBP3-DAP-Seq(GSE60143)/Homer(0.827) More Information \| Similar Motifs Found | motif file (matrix) |
| 11 \* | A C G T A G T C G T A C C G T A A C T G A C T G G A T C G C T A A C T G C G A T | 1e-9 | -2.255e+01 | 4.41% | 1.22% | 376.8bp (399.8bp) | Tag/dmmpmm(Papatsenko)/fly(0.728) More Information \| Similar Motifs Found | motif file (matrix) |
| 12 \* | C G T A C G T A G T C A A C T G A C G T C G T A C T A G C T A G | 1e-9 | -2.137e+01 | 19.40% | 11.81% | 359.1bp (472.4bp) | CG5213(RRM)/Drosophila\_melanogaster-RNCMPT00010-PBM/HughesRNA(0.872) More Information \| Similar Motifs Found | motif file (matrix) |
| 13 \* | A C T G A C G T C G T A A G C T C G T A A C T G C G T A C G T A C G A T T G C A | 1e-8 | -2.039e+01 | 3.90% | 1.07% | 341.7bp (466.1bp) | Tb\_0219(RRM)/Trypanosoma\_brucei-RNCMPT00219-PBM/HughesRNA(0.692) More Information \| Similar Motifs Found | motif file (matrix) |
| 14 \* | A C G T A C G T A G T C A C T G A G T C A T G C A C T G A C T G A C T G A G T C | 1e-8 | -1.964e+01 | 0.63% | 0.00% | 482.8bp (0.0bp) | SUT1(MacIsaac)/Yeast(0.730) More Information \| Similar Motifs Found | motif file (matrix) |
| 15 \* | A G T C A G T C T C G A A T G C A C G T A C T G C T G A A C G T A T G C A G T C | 1e-8 | -1.959e+01 | 5.04% | 1.71% | 400.6bp (502.6bp) | RHOXF1/MA0719.1/Jaspar(0.702) More Information \| Similar Motifs Found | motif file (matrix) |
| 16 \* | G T C A T G C A T C G A G T A C A C T G A C T G A C T G G T C A | 1e-8 | -1.895e+01 | 23.68% | 15.82% | 355.3bp (474.8bp) | TYE7/MA0409.1/Jaspar(0.704) More Information \| Similar Motifs Found | motif file (matrix) |
| 17 \* | A C G T A G T C C G T A A C T G C G T A A C G T A C G T A G T C C G T A A G T C | 1e-6 | -1.596e+01 | 0.88% | 0.05% | 267.6bp (202.9bp) | Tb\_0251(RRM)/Trypanosoma\_brucei-RNCMPT00251-PBM/HughesRNA(0.733) More Information \| Similar Motifs Found | motif file (matrix) |
| 18 \* | T A C G T C A G C G A T A T C G A C G T C G T A T G C A G T A C T G C A T C A G | 1e-6 | -1.515e+01 | 8.06% | 4.06% | 386.9bp (474.0bp) | ovo/dmmpmm(Bergman)/fly(0.710) More Information \| Similar Motifs Found | motif file (matrix) |
| 19 \* | C G A T A C T G A C T G A C T G C T G A A C G T A C T G G T C A | 1e-6 | -1.439e+01 | 12.72% | 7.68% | 367.6bp (467.4bp) | PB0098.1\_Zfp410\_1/Jaspar(0.795) More Information \| Similar Motifs Found | motif file (matrix) |
| 20 \* | T C G A T G C A A T C G T C G A T C G A G C A T A G C T G T A C T A G C T C A G | 1e-4 | -9.897e+00 | 1.51% | 0.37% | 216.8bp (614.2bp) | dl-B/dmmpmm(Bergman)/fly(0.743) More Information \| Similar Motifs Found | motif file (matrix) |
| 21 \* | A G T C A C T G A C G T A C T G A G T C A C G T A G T C A C G T C G T A A C G T | 1e-2 | -5.528e+00 | 0.38% | 0.04% | 152.8bp (146.4bp) | kni/MA0451.1/Jaspar(0.711) More Information \| Similar Motifs Found | motif file (matrix) |
| 22 \* | A G T C A C T G G T C A A C T G A C T G A G T C A G T C C G A T | 1e-1 | -3.482e+00 | 2.77% | 1.79% | 393.5bp (384.5bp) | ZNF711(Zf)/SHSY5Y-ZNF711-ChIP-Seq(GSE20673)/Homer(0.874) More Information \| Similar Motifs Found | motif file (matrix) |
| 23 \* | C T G A A G T C A G T C C T G A A G T C A G T C G T C A A G T C A G T C C G T A | 1e0 | -1.599e+00 | 2.27% | 1.82% | 454.0bp (375.1bp) | ZBTB7C/MA0695.1/Jaspar(0.739) More Information \| Similar Motifs Found | motif file (matrix) |
