## Supplemental Table 1 for "The Establishment of Cell-Type Specific Gene Regulation in the Sea Urchin Embryo": Hpf72_EctoOral_homerResults.html

homer\_motifs\_wbackground/Hpf72\_EctoOral/ - Homer de novo Motif Results


### Homer *de novo* Motif Results (homer\_motifs\_wbackground/Hpf72\_EctoOral/)

Known Motif Enrichment Results  
Gene Ontology Enrichment Results  
If Homer is having trouble matching a motif to a known motif, try copy/pasting the matrix file into
STAMP  
More information on motif finding results: HOMER
| Description of Results
| Tips
  
Total target sequences = 1971  
Total background sequences = 14303  
\* - possible false positive  

|  |  |  |  |  |  |  |  |  |
| --- | --- | --- | --- | --- | --- | --- | --- | --- |
| Rank | Motif | P-value | log P-pvalue | % of Targets | % of Background | STD(Bg STD) | Best Match/Details | Motif File |
| 1 | A C G T C G T A A C G T A G C T A C T G C G T A A C G T A G C T | 1e-404 | -9.324e+02 | 60.02% | 16.81% | 362.3bp (450.1bp) | ONECUT1/MA0679.2/Jaspar(0.936) More Information \| Similar Motifs Found | motif file (matrix) |
| 2 | C A G T A G T C C G A T C G T A A C G T A C G T A C T G A C G T | 1e-45 | -1.049e+02 | 44.14% | 28.99% | 363.8bp (418.0bp) | SOX15/MA1152.1/Jaspar(0.882) More Information \| Similar Motifs Found | motif file (matrix) |
| 3 | C G T A A C G T A G T C A C T G C G T A G T C A | 1e-27 | -6.410e+01 | 55.96% | 43.56% | 412.6bp (470.8bp) | ceh-48/MA0921.1/Jaspar(0.813) More Information \| Similar Motifs Found | motif file (matrix) |
| 4 | A G T C T C A G A C T G A T C G T G A C G A T C T C A G A C T G C T G A T A G C | 1e-21 | -4.879e+01 | 14.51% | 8.05% | 345.6bp (481.2bp) | SKN7/SKN7\_H2O2Lo/[](Harbison)/Yeast(0.754) More Information \| Similar Motifs Found | motif file (matrix) |
| 5 | A G C T T A G C A G T C A C T G A C G T A C T G G T C A A C T G G T C A G T C A | 1e-18 | -4.246e+01 | 2.99% | 0.72% | 520.7bp (369.9bp) | Su(H)/dmmpmm(Papatsenko)/fly(0.748) More Information \| Similar Motifs Found | motif file (matrix) |
| 6 | C G T A A G T C A C G T A C G T A C G T A G T C | 1e-18 | -4.159e+01 | 49.77% | 39.97% | 420.5bp (454.9bp) | PTBP1(RRM)/Homo\_sapiens-RNCMPT00269-PBM/HughesRNA(0.881) More Information \| Similar Motifs Found | motif file (matrix) |
| 7 | G C T A C T A G A C T G C T A G A C T G C T A G A C G T A T C G | 1e-17 | -3.948e+01 | 41.91% | 32.69% | 416.0bp (461.2bp) | MSN2/MA0341.1/Jaspar(0.804) More Information \| Similar Motifs Found | motif file (matrix) |
| 8 | C G A T G A C T G T A C A C G T C A G T A G C T A G C T T A G C C G T A G C T A | 1e-16 | -3.801e+01 | 20.35% | 13.50% | 402.1bp (444.0bp) | At2g41835(C2H2)/col-At2g41835-DAP-Seq(GSE60143)/Homer(0.781) More Information \| Similar Motifs Found | motif file (matrix) |
| 9 | A C G T C G T A T C A G T C G A G A C T A T G C A C G T C G T A A C T G C G T A | 1e-16 | -3.747e+01 | 7.15% | 3.30% | 417.2bp (463.2bp) | GATA6/MA1396.1/Jaspar(0.905) More Information \| Similar Motifs Found | motif file (matrix) |
| 10 | A G T C G A C T C A G T T C G A A C T G G A T C G T A C G A C T A C G T C G A T | 1e-15 | -3.601e+01 | 14.10% | 8.55% | 400.4bp (451.0bp) | At5g08520/MA1399.1/Jaspar(0.660) More Information \| Similar Motifs Found | motif file (matrix) |
| 11 | T A C G G T A C C A T G T A G C T A C G A G T C T A G C A C T G T G A C C A T G | 1e-15 | -3.486e+01 | 13.24% | 7.95% | 292.6bp (431.8bp) | STP1(MacIsaac)/Yeast(0.787) More Information \| Similar Motifs Found | motif file (matrix) |
| 12 | A G C T G T A C A G C T A G T C A T G C A C G T C T G A A G C T C G A T A T G C | 1e-14 | -3.445e+01 | 4.11% | 1.48% | 390.6bp (430.4bp) | PB0128.1\_Gcm1\_2/Jaspar(0.782) More Information \| Similar Motifs Found | motif file (matrix) |
| 13 | C T G A A T G C C T A G G A T C C T A G C A T G A C T G A T C G A C T G G A T C | 1e-13 | -3.213e+01 | 3.20% | 1.02% | 459.5bp (402.6bp) | PB0010.1\_Egr1\_1/Jaspar(0.787) More Information \| Similar Motifs Found | motif file (matrix) |
| 14 | A T G C G T A C C G T A G A C T A C T G C T A G T G C A C G A T C G T A A C T G | 1e-12 | -2.811e+01 | 3.04% | 1.04% | 413.2bp (425.5bp) | HNRNPAB(RRM)/Tetraodon\_nigroviridis-RNCMPT00245-PBM/HughesRNA(0.694) More Information \| Similar Motifs Found | motif file (matrix) |
| 15 | A G T C A T G C A G T C A C G T G T C A A C G T A C T G A C G T A C G T A G T C | 1e-12 | -2.795e+01 | 1.12% | 0.15% | 462.8bp (283.1bp) | Sox4(HMG)/proB-Sox4-ChIP-Seq(GSE50066)/Homer(0.736) More Information \| Similar Motifs Found | motif file (matrix) |
| 16 | C G T A C G T A A G T C G T A C A G T C C G T A T G C A A G T C A C T G A C T G | 1e-12 | -2.781e+01 | 1.47% | 0.28% | 355.8bp (611.8bp) | MYB55/MA1041.1/Jaspar(0.770) More Information \| Similar Motifs Found | motif file (matrix) |
| 17 \* | A G T C C A T G C G T A A G T C A C T G A C T G A C G T A C G T A G T C A T G C | 1e-11 | -2.660e+01 | 0.41% | 0.01% | 393.7bp (0.0bp) | OPI1/MA0349.1/Jaspar(0.741) More Information \| Similar Motifs Found | motif file (matrix) |
| 18 \* | A G T C A G T C C G T A A G T C A C T G A C G T A G T C A T G C C G T A A C T G | 1e-10 | -2.306e+01 | 0.61% | 0.04% | 662.6bp (152.5bp) | ABF4/MA1659.1/Jaspar(0.761) More Information \| Similar Motifs Found | motif file (matrix) |
| 19 \* | T G A C G T A C G T C A G T A C A G T C C G T A A G T C G T A C C G T A G T C A | 1e-9 | -2.269e+01 | 6.44% | 3.52% | 405.2bp (418.9bp) | schlank/MA0193.1/Jaspar(0.775) More Information \| Similar Motifs Found | motif file (matrix) |
| 20 \* | A G T C A C G T A C G T A C T G A C T G C G T A C G T A A G T C A C G T A C G T | 1e-9 | -2.202e+01 | 0.71% | 0.08% | 328.6bp (284.8bp) | TCF7L1/MA1421.1/Jaspar(0.704) More Information \| Similar Motifs Found | motif file (matrix) |
| 21 \* | A G T C A C T G A C G T A C T G A C G T A C G T A G T C C G T A A C G T A C G T | 1e-9 | -2.104e+01 | 0.81% | 0.11% | 434.5bp (273.7bp) | lin-14/MA0261.1/Jaspar(0.717) More Information \| Similar Motifs Found | motif file (matrix) |
| 22 \* | A G T C A G T C C G T A C G T A A C T G A G T C A C G T A G T C A T G C A G T C | 1e-8 | -1.948e+01 | 0.51% | 0.04% | 150.4bp (325.6bp) | KLF17/MA1514.1/Jaspar(0.717) More Information \| Similar Motifs Found | motif file (matrix) |
| 23 \* | A G T C A C T G A G T C A G T C A C T G A G C T C G T A C G T A | 1e-7 | -1.680e+01 | 2.44% | 1.01% | 327.2bp (416.2bp) | ERF115(AP2EREBP)/colamp-ERF115-DAP-Seq(GSE60143)/Homer(0.841) More Information \| Similar Motifs Found | motif file (matrix) |
| 24 \* | A C T G C G T A A C T G A G T C C G T A C G T A A G T C A C T G C G T A C G T A | 1e-6 | -1.508e+01 | 0.36% | 0.03% | 447.6bp (1078.7bp) | CG2950(KH)/Drosophila\_melanogaster-RNCMPT00007-PBM/HughesRNA(0.772) More Information \| Similar Motifs Found | motif file (matrix) |
