## Supplemental Table 1 for "The Establishment of Cell-Type Specific Gene Regulation in the Sea Urchin Embryo": Hpf72_FilopodialCells_homerResults.html

homer\_motifs\_wbackground/Hpf72\_FilopodialCells/ - Homer de novo Motif Results


### Homer *de novo* Motif Results (homer\_motifs\_wbackground/Hpf72\_FilopodialCells/)

Known Motif Enrichment Results  
Gene Ontology Enrichment Results  
If Homer is having trouble matching a motif to a known motif, try copy/pasting the matrix file into
STAMP  
More information on motif finding results: HOMER
| Description of Results
| Tips
  
Total target sequences = 855  
Total background sequences = 16129  
\* - possible false positive  

|  |  |  |  |  |  |  |  |  |
| --- | --- | --- | --- | --- | --- | --- | --- | --- |
| Rank | Motif | P-value | log P-pvalue | % of Targets | % of Background | STD(Bg STD) | Best Match/Details | Motif File |
| 1 | C T A G T G A C T G C A A C T G A C T G G T C A C G T A T C A G A G C T A C T G | 1e-42 | -9.750e+01 | 42.57% | 21.58% | 312.8bp (427.4bp) | Etv2(ETS)/ES-ER71-ChIP-Seq(GSE59402)/Homer(0.987) More Information \| Similar Motifs Found | motif file (matrix) |
| 2 \* | G T C A G T A C T A G C A C G T A T C G C A T G A C G T C A G T A C T G C A T G | 1e-9 | -2.272e+01 | 2.81% | 0.54% | 274.3bp (427.6bp) | grh/dmmpmm(Bigfoot)/fly(0.731) More Information \| Similar Motifs Found | motif file (matrix) |
| 3 \* | C G A T A T C G C T A G A C G T C T A G C G T A C A T G C G T A G C T A C A G T | 1e-7 | -1.666e+01 | 7.02% | 3.29% | 285.9bp (365.7bp) | AGL55/MA1202.1/Jaspar(0.796) More Information \| Similar Motifs Found | motif file (matrix) |
| 4 \* | A C T G A G T C C T A G A G T C C G A T A T C G A G T C C T G A G A T C C G A T | 1e-6 | -1.522e+01 | 3.74% | 1.32% | 299.9bp (365.0bp) | ZC3H10(Znf)/Homo\_sapiens-RNCMPT00085-PBM/HughesRNA(0.794) More Information \| Similar Motifs Found | motif file (matrix) |
| 5 \* | A C G T A G T C A C T G C G T A A C T G A T C G A C T G C G T A A C T G A G C T | 1e-6 | -1.389e+01 | 1.17% | 0.15% | 245.2bp (343.6bp) | HNRNPA2B1(RRM)/Homo\_sapiens-RNCMPT00024-PBM/HughesRNA(0.633) More Information \| Similar Motifs Found | motif file (matrix) |
| 6 \* | A C T G T C A G A G T C A C T G C G T A C G T A A C G T A C G T A G T C A C T G | 1e-5 | -1.225e+01 | 0.47% | 0.02% | 205.8bp (105.9bp) | AT5G22990(C2H2)/col-AT5G22990-DAP-Seq(GSE60143)/Homer(0.733) More Information \| Similar Motifs Found | motif file (matrix) |
| 7 \* | C G T A G C T A C G A T T A C G C G T A A C T G C T A G G T A C C G T A C A T G | 1e-4 | -1.094e+01 | 6.67% | 3.68% | 251.3bp (429.0bp) | FOS/MA0476.1/Jaspar(0.731) More Information \| Similar Motifs Found | motif file (matrix) |
| 8 \* | A G T C A C T G G T C A A T G C C G A T A G T C A G T C A C G T A C T G A C T G | 1e-4 | -1.043e+01 | 0.82% | 0.10% | 199.9bp (499.7bp) | CTCFL/MA1102.2/Jaspar(0.754) More Information \| Similar Motifs Found | motif file (matrix) |
| 9 \* | C A G T C A T G C G A T A T G C T C G A A G T C C G A T A C T G T C A G A T G C | 1e-4 | -9.672e+00 | 10.64% | 7.02% | 379.4bp (413.4bp) | NR1H4/MA1110.1/Jaspar(0.623) More Information \| Similar Motifs Found | motif file (matrix) |
| 10 \* | A G T C G T A C A T C G A T C G A C T G A C G T G T A C C G T A | 1e-4 | -9.605e+00 | 15.79% | 11.39% | 287.8bp (425.7bp) | PB0153.1\_Nr2f2\_2/Jaspar(0.907) More Information \| Similar Motifs Found | motif file (matrix) |
| 11 \* | A C G T A C T G A T C G T A G C T C G A C T G A T C A G G A C T G A C T G T A C | 1e-4 | -9.425e+00 | 10.18% | 6.69% | 297.2bp (430.8bp) | SD0002.1\_at\_AC\_acceptor/Jaspar(0.727) More Information \| Similar Motifs Found | motif file (matrix) |
| 12 \* | G T C A A C T G G A C T C G A T A T G C G C T A A C T G A C G T A G C T A T G C | 1e-4 | -9.221e+00 | 13.68% | 9.68% | 285.9bp (414.0bp) | Initiator/Drosophila-Promoters/Homer(0.734) More Information \| Similar Motifs Found | motif file (matrix) |
| 13 \* | A T C G C A G T C A G T T C G A A G C T T G C A A T C G A C T G G T C A G C T A | 1e-3 | -8.982e+00 | 9.01% | 5.82% | 408.3bp (428.2bp) | STAT1/MA0137.3/Jaspar(0.730) More Information \| Similar Motifs Found | motif file (matrix) |
| 14 \* | T C A G G A C T T G A C T C G A A T G C A T C G G C A T A T C G T C G A C A G T | 1e-3 | -8.501e+00 | 18.71% | 14.27% | 418.4bp (411.0bp) | SREBF2(var.2)/MA0828.1/Jaspar(0.912) More Information \| Similar Motifs Found | motif file (matrix) |
| 15 \* | A C G T C G T A T C A G A C G T A C G T A G C T A G T C C T A G | 1e-2 | -6.649e+00 | 20.12% | 16.17% | 313.2bp (428.8bp) | IRF6/MA1509.1/Jaspar(0.848) More Information \| Similar Motifs Found | motif file (matrix) |
| 16 \* | T A C G A G T C C G A T C G T A C G A T C G T A G A T C C T A G | 1e-2 | -5.345e+00 | 14.85% | 11.85% | 367.1bp (407.0bp) | Cf2-II/dmmpmm(Pollard)/fly(0.709) More Information \| Similar Motifs Found | motif file (matrix) |
| 17 \* | A G T C A C G T A C T G A C G T C G T A A C G T A C G T C G T A A C T G A C T G | 1e-1 | -4.368e+00 | 0.35% | 0.06% | 391.2bp (374.0bp) | PUF68(RRM)/Drosophila\_melanogaster-RNCMPT00141-PBM/HughesRNA(0.699) More Information \| Similar Motifs Found | motif file (matrix) |
| 18 \* | A C T G A C T G A C G T A G T C A G T C A C G T A C G T C G T A C G A T A C T G | 1e0 | -1.676e+00 | 0.35% | 0.18% | 228.8bp (372.0bp) | ttk/dmmpmm(Bigfoot)/fly(0.724) More Information \| Similar Motifs Found | motif file (matrix) |
| 19 \* | G A T C A C T G A C G T C G T A A T G C C T A G C G T A C G T A | 1e0 | -1.644e+00 | 15.20% | 14.12% | 258.6bp (409.1bp) | SPL5(SBP)/colamp-SPL5-DAP-Seq(GSE60143)/Homer(0.865) More Information \| Similar Motifs Found | motif file (matrix) |
| 20 \* | C G T A T A C G C T G A C G A T C G T A A C T G C G T A C T A G C G T A A C T G | 1e0 | -8.277e-01 | 15.09% | 14.85% | 363.5bp (411.0bp) | Trl/dmmpmm(Down)/fly(0.801) More Information \| Similar Motifs Found | motif file (matrix) |
| 21 \* | A C T G A C T G A C T G A C T G A C T G A C T G A C T G A C T G | 1e0 | -2.008e-01 | 13.33% | 14.37% | 284.9bp (440.0bp) | Maz(Zf)/HepG2-Maz-ChIP-Seq(GSE31477)/Homer(0.974) More Information \| Similar Motifs Found | motif file (matrix) |
