## Supplemental Table 1 for "The Establishment of Cell-Type Specific Gene Regulation in the Sea Urchin Embryo": Hpf72_Forgut_homerResults.html

homer\_motifs\_wbackground/Hpf72\_Forgut/ - Homer de novo Motif Results


### Homer *de novo* Motif Results (homer\_motifs\_wbackground/Hpf72\_Forgut/)

Known Motif Enrichment Results  
Gene Ontology Enrichment Results  
If Homer is having trouble matching a motif to a known motif, try copy/pasting the matrix file into
STAMP  
More information on motif finding results: HOMER
| Description of Results
| Tips
  
Total target sequences = 1586  
Total background sequences = 14596  
\* - possible false positive  

|  |  |  |  |  |  |  |  |  |
| --- | --- | --- | --- | --- | --- | --- | --- | --- |
| Rank | Motif | P-value | log P-pvalue | % of Targets | % of Background | STD(Bg STD) | Best Match/Details | Motif File |
| 1 | C T G A G C A T T C A G A C G T A C G T C A G T C T A G G A T C G C A T A G T C | 1e-83 | -1.923e+02 | 65.83% | 41.58% | 386.0bp (434.7bp) | PHA-4(Forkhead)/cElegans-Embryos-PHA4-ChIP-Seq(modEncode)/Homer(0.947) More Information \| Similar Motifs Found | motif file (matrix) |
| 2 | C A G T A T G C A G C T C G T A A C G T C A G T T A C G C G A T | 1e-15 | -3.517e+01 | 60.59% | 50.53% | 386.2bp (418.1bp) | SOX15/MA1152.1/Jaspar(0.854) More Information \| Similar Motifs Found | motif file (matrix) |
| 3 | C G T A T G C A A C T G A G C T C A T G A G T C A C G T T G A C G C A T A T G C | 1e-13 | -3.063e+01 | 3.66% | 1.15% | 360.8bp (427.0bp) | Trl/dmmpmm(Pollard)/fly(0.731) More Information \| Similar Motifs Found | motif file (matrix) |
| 4 \* | G T C A A C T G C G T A C G T A C G T A A C T G C G A T G T C A A C G T C G T A | 1e-11 | -2.698e+01 | 5.42% | 2.33% | 389.7bp (372.0bp) | PTBP1(RRM)/Homo\_sapiens-RNCMPT00269-PBM/HughesRNA(0.776) More Information \| Similar Motifs Found | motif file (matrix) |
| 5 \* | C T A G T G A C G C A T G T A C C T A G G C T A A C T G A C G T T A C G C G A T | 1e-11 | -2.621e+01 | 2.96% | 0.90% | 561.6bp (435.4bp) | Rbm24(RRM)/Tetraodon\_nigroviridis-RNCMPT00285-PBM/HughesRNA(0.725) More Information \| Similar Motifs Found | motif file (matrix) |
| 6 \* | A G T C C G T A G A T C G T A C A C G T A T C G T G C A T C G A A T G C A G C T | 1e-10 | -2.404e+01 | 3.72% | 1.40% | 357.7bp (410.8bp) | NGA4(ABI3VP1)/col-NGA4-DAP-Seq(GSE60143)/Homer(0.812) More Information \| Similar Motifs Found | motif file (matrix) |
| 7 \* | A C G T A C G T C T G A A G T C C T A G A C G T C G T A C G T A | 1e-10 | -2.393e+01 | 30.33% | 23.20% | 383.3bp (420.8bp) | CIN5/CIN5\_H2O2Lo/[](Harbison)/Yeast(0.961) More Information \| Similar Motifs Found | motif file (matrix) |
| 8 \* | C G T A G C A T C G T A T A G C T A C G A G T C A C T G C T G A A G T C C T A G | 1e-10 | -2.369e+01 | 5.30% | 2.41% | 363.9bp (489.2bp) | PB0179.1\_Sp100\_2/Jaspar(0.728) More Information \| Similar Motifs Found | motif file (matrix) |
| 9 \* | T A C G A T C G T A G C A C G T T A C G C T A G C G A T A G C T T A G C A C G T | 1e-10 | -2.354e+01 | 2.46% | 0.71% | 418.8bp (495.3bp) | SWI5/MA0402.1/Jaspar(0.657) More Information \| Similar Motifs Found | motif file (matrix) |
| 10 \* | T G C A C T G A A G C T C A T G C A G T C T G A G T A C A T C G G T A C T G C A | 1e-9 | -2.220e+01 | 11.73% | 7.31% | 444.6bp (440.7bp) | SPL13/MA1321.1/Jaspar(0.748) More Information \| Similar Motifs Found | motif file (matrix) |
| 11 \* | C G A T A C T G C G T A A C T G A C T G C T A G C G T A A C G T A G T C C G A T | 1e-8 | -2.061e+01 | 1.58% | 0.36% | 391.8bp (358.5bp) | HAP3(CCAATHAP3)/col-HAP3-DAP-Seq(GSE60143)/Homer(0.680) More Information \| Similar Motifs Found | motif file (matrix) |
| 12 \* | A T G C A C T G A C G T A C G T A T C G C G T A C G T A C G T A C T G A A C T G | 1e-8 | -2.037e+01 | 3.34% | 1.30% | 307.0bp (387.7bp) | At2g41835(C2H2)/col-At2g41835-DAP-Seq(GSE60143)/Homer(0.809) More Information \| Similar Motifs Found | motif file (matrix) |
| 13 \* | C G T A A G T C A C G T A G T C A G T C A G T C A G T C A C T G A C T G A T G C | 1e-8 | -2.000e+01 | 0.38% | 0.01% | 268.5bp (223.8bp) | NHP10/MA0344.1/Jaspar(0.867) More Information \| Similar Motifs Found | motif file (matrix) |
| 14 \* | A C G T A C G T A C G T A C G T A G T C A C T G C G T A C G T A A C G T A C T G | 1e-8 | -1.988e+01 | 0.50% | 0.02% | 268.6bp (179.4bp) | YER051W(MacIsaac)/Yeast(0.774) More Information \| Similar Motifs Found | motif file (matrix) |
| 15 \* | A G C T T A C G A T C G C A T G A T C G A C G T A T G C T A G C T A G C A C G T | 1e-7 | -1.726e+01 | 0.57% | 0.05% | 267.7bp (237.9bp) | PCF5(TCP)/Oryza sativa/AthaMap(0.775) More Information \| Similar Motifs Found | motif file (matrix) |
| 16 \* | A C T G C G T A A C T G A C G T A C T G A G T C A G T C C G T A A C G T A C G T | 1e-6 | -1.435e+01 | 0.76% | 0.12% | 387.9bp (514.5bp) | pho/dmmpmm(Bergman)/fly(0.775) More Information \| Similar Motifs Found | motif file (matrix) |
| 17 \* | C G T A C G T A A G T C A C G T A G T C C G T A C G T A A C T G A G T C A G T C | 1e-5 | -1.359e+01 | 0.38% | 0.02% | 112.8bp (181.6bp) | EXC-7(RRM)/Caenorhabditis\_elegans-RNCMPT00014-PBM/HughesRNA(0.717) More Information \| Similar Motifs Found | motif file (matrix) |
| 18 \* | G C T A A G T C A G T C G T C A A G T C A T C G A T G C A C T G A C G T C G T A | 1e-5 | -1.253e+01 | 0.82% | 0.17% | 334.6bp (296.1bp) | PL0008.1\_hlh-29/Jaspar(0.794) More Information \| Similar Motifs Found | motif file (matrix) |
