## Supplemental Table 1 for "The Establishment of Cell-Type Specific Gene Regulation in the Sea Urchin Embryo": Hpf72_GlobularCells_homerResults.html

homer\_motifs\_wbackground/Hpf72\_GlobularCells/ - Homer de novo Motif Results


### Homer *de novo* Motif Results (homer\_motifs\_wbackground/Hpf72\_GlobularCells/)

Known Motif Enrichment Results  
Gene Ontology Enrichment Results  
If Homer is having trouble matching a motif to a known motif, try copy/pasting the matrix file into
STAMP  
More information on motif finding results: HOMER
| Description of Results
| Tips
  
Total target sequences = 255  
Total background sequences = 16653  
\* - possible false positive  

|  |  |  |  |  |  |  |  |  |
| --- | --- | --- | --- | --- | --- | --- | --- | --- |
| Rank | Motif | P-value | log P-pvalue | % of Targets | % of Background | STD(Bg STD) | Best Match/Details | Motif File |
| 1 \* | A C G T G T C A C A G T G A C T C G T A C G T A C A T G G C T A G A C T T C A G | 1e-9 | -2.209e+01 | 13.33% | 3.78% | 400.0bp (430.1bp) | Arid3b/MA0601.1/Jaspar(0.766) More Information \| Similar Motifs Found | motif file (matrix) |
| 2 \* | G A C T G T C A T A G C C G A T C T A G A C G T A C T G C G T A C G T A C T A G | 1e-8 | -2.044e+01 | 8.63% | 1.75% | 394.1bp (420.5bp) | Tb\_0251(RRM)/Trypanosoma\_brucei-RNCMPT00251-PBM/HughesRNA(0.719) More Information \| Similar Motifs Found | motif file (matrix) |
| 3 \* | C G A T G C A T A T G C A T G C C T G A T A C G C A T G T G C A T G C A C G A T | 1e-8 | -1.937e+01 | 10.98% | 2.97% | 246.6bp (449.0bp) | STAT1/MA0137.3/Jaspar(0.788) More Information \| Similar Motifs Found | motif file (matrix) |
| 4 \* | G T A C C T G A A C T G G C T A T G A C C G T A C G A T T A C G T C G A G T A C | 1e-8 | -1.931e+01 | 10.59% | 2.78% | 521.3bp (402.5bp) | ARF27/MA1691.1/Jaspar(0.663) More Information \| Similar Motifs Found | motif file (matrix) |
| 5 \* | A T G C G T C A G T A C A G C T C A G T A C G T G A C T T G C A A G T C C G T A | 1e-7 | -1.753e+01 | 26.67% | 13.58% | 367.0bp (410.9bp) | CDF3/MA0974.2/Jaspar(0.742) More Information \| Similar Motifs Found | motif file (matrix) |
| 6 \* | G C A T T G A C A C G T T C G A A G T C T A G C C G A T A C T G A G T C C T G A | 1e-6 | -1.491e+01 | 8.63% | 2.40% | 519.0bp (407.8bp) | vfl/MA1462.1/Jaspar(0.862) More Information \| Similar Motifs Found | motif file (matrix) |
| 7 \* | G T A C G A C T G T C A C G T A A C T G G T C A A G T C A G T C A T G C A G C T | 1e-6 | -1.445e+01 | 6.27% | 1.35% | 299.1bp (458.9bp) | LRF(Zf)/Erythroblasts-ZBTB7A-ChIP-Seq(GSE74977)/Homer(0.775) More Information \| Similar Motifs Found | motif file (matrix) |
| 8 \* | A C T G A G T C A C T G A G T C A C G T C G T A A C G T A G T C A C T G A C T G | 1e-5 | -1.228e+01 | 2.35% | 0.16% | 263.2bp (394.6bp) | STP4/MA0397.1/Jaspar(0.751) More Information \| Similar Motifs Found | motif file (matrix) |
| 9 \* | G C A T A T C G A G C T A C G T T A C G A T G C A C G T A T G C G T C A T A C G | 1e-4 | -9.884e+00 | 4.71% | 1.15% | 333.8bp (400.3bp) | ACE2/ACE2\_YPD/2-SWI5(Harbison)/Yeast(0.630) More Information \| Similar Motifs Found | motif file (matrix) |
| 10 \* | A G T C T C G A A G T C A G T C C G T A A C T G A G T C A C G T | 1e-3 | -9.119e+00 | 19.61% | 11.46% | 401.9bp (423.9bp) | Tcf12(bHLH)/GM12878-Tcf12-ChIP-Seq(GSE32465)/Homer(0.795) More Information \| Similar Motifs Found | motif file (matrix) |
| 11 \* | G T A C A C T G C G T A C A T G T A C G T A G C A G T C A G C T G T A C A T C G | 1e-3 | -8.581e+00 | 1.18% | 0.04% | 86.6bp (214.0bp) | ZNF711(Zf)/SHSY5Y-ZNF711-ChIP-Seq(GSE20673)/Homer(0.708) More Information \| Similar Motifs Found | motif file (matrix) |
| 12 \* | A C G T C G T A C G T A A C G T A G T C A C G T C G T A A G T C | 1e-3 | -8.164e+00 | 8.24% | 3.49% | 283.1bp (472.0bp) | Tb\_0252(RRM)/Trypanosoma\_brucei-RNCMPT00252-PBM/HughesRNA(0.814) More Information \| Similar Motifs Found | motif file (matrix) |
| 13 \* | G C A T A G C T A T G C G T A C G A T C A T C G C A T G C A T G | 1e-3 | -7.675e+00 | 30.59% | 21.54% | 291.9bp (435.8bp) | PUT3/MA0358.1/Jaspar(0.897) More Information \| Similar Motifs Found | motif file (matrix) |
| 14 \* | A C G T A C G T C G T A A G C T A C T G A T C G A G T C C G T A | 1e-3 | -7.605e+00 | 6.27% | 2.39% | 293.2bp (440.5bp) | CDX1/MA0878.2/Jaspar(0.781) More Information \| Similar Motifs Found | motif file (matrix) |
| 15 \* | A G T C A G T C C G T A A C T G C G T A A C G T A C G T A G T C A C G T A C T G | 1e-2 | -6.704e+00 | 8.24% | 3.92% | 230.1bp (426.0bp) | At1g25550(G2like)/colamp-At1g25550-DAP-Seq(GSE60143)/Homer(0.782) More Information \| Similar Motifs Found | motif file (matrix) |
| 16 \* | A G T C C G T A A C G T A G T C A G T C A C G T | 1e-1 | -3.922e+00 | 37.25% | 31.02% | 416.1bp (425.2bp) | G3BP2(RRM)/Homo\_sapiens-RNCMPT00021-PBM/HughesRNA(0.870) More Information \| Similar Motifs Found | motif file (matrix) |
| 17 \* | A G T C A G T C A G T C A G T C A C G T A T G C A G T C A G T C A C G T A G T C | 1e-1 | -3.227e+00 | 3.53% | 1.78% | 566.4bp (478.6bp) | ZNF148/MA1653.1/Jaspar(0.876) More Information \| Similar Motifs Found | motif file (matrix) |
| 18 \* | A C T G A C G T A G T C C G T A C G T A A C T G A C T G C G T A A C T G A G T C | 1e0 | -1.864e+00 | 0.39% | 0.07% | 406.8bp (442.9bp) | RBM5(RRM)/Homo\_sapiens-RNCMPT00154-PBM/HughesRNA(0.730) More Information \| Similar Motifs Found | motif file (matrix) |
| 19 \* | A C T G A C G T A T C G A C T G A C G T T C A G | 1e0 | -1.799e+00 | 59.61% | 56.39% | 399.2bp (418.5bp) | Run/dmmpmm(Papatsenko)/fly(0.843) More Information \| Similar Motifs Found | motif file (matrix) |
| 20 \* | C G T A A C T G A G T C A C T G A G T C A C T G | 1e0 | -9.925e-01 | 10.98% | 10.21% | 381.7bp (391.5bp) | FUS(RRM)/Homo\_sapiens-RNCMPT00018-PBM/HughesRNA(0.883) More Information \| Similar Motifs Found | motif file (matrix) |
