## Supplemental Table 1 for "The Establishment of Cell-Type Specific Gene Regulation in the Sea Urchin Embryo": Hpf72_Hindgut_homerResults.html

homer\_motifs\_wbackground/Hpf72\_Hindgut/ - Homer de novo Motif Results


### Homer *de novo* Motif Results (homer\_motifs\_wbackground/Hpf72\_Hindgut/)

Known Motif Enrichment Results  
Gene Ontology Enrichment Results  
If Homer is having trouble matching a motif to a known motif, try copy/pasting the matrix file into
STAMP  
More information on motif finding results: HOMER
| Description of Results
| Tips
  
Total target sequences = 1456  
Total background sequences = 14792  
\* - possible false positive  

|  |  |  |  |  |  |  |  |  |
| --- | --- | --- | --- | --- | --- | --- | --- | --- |
| Rank | Motif | P-value | log P-pvalue | % of Targets | % of Background | STD(Bg STD) | Best Match/Details | Motif File |
| 1 | G C A T T C G A C G T A A C T G A G T C G T C A G T C A C G T A A G T C C G T A | 1e-39 | -8.993e+01 | 64.49% | 47.32% | 348.0bp (420.5bp) | fkh/dmmpmm(Noyes)/fly(0.962) More Information \| Similar Motifs Found | motif file (matrix) |
| 2 | A C G T A G T C C T G A C T G A C T G A A C T G A C G T G A T C A G T C C T G A | 1e-34 | -7.942e+01 | 17.51% | 7.62% | 397.5bp (422.0bp) | HNF4G/MA0484.2/Jaspar(0.931) More Information \| Similar Motifs Found | motif file (matrix) |
| 3 | A C T G C G A T A C G T C G T A G T C A G C A T A T G C C G T A A C G T A G C T | 1e-26 | -6.196e+01 | 13.05% | 5.50% | 326.7bp (433.9bp) | HNF1b(Homeobox)/PDAC-HNF1B-ChIP-Seq(GSE64557)/Homer(0.918) More Information \| Similar Motifs Found | motif file (matrix) |
| 4 | A C T G A C G T A C G T A C G T C G T A A C G T A T C G C T G A A T G C A G C T | 1e-22 | -5.253e+01 | 37.02% | 25.20% | 344.8bp (432.6bp) | caudal(Homeobox)/Drosophila-Embryos-ChIP-Chip(modEncode)/Homer(0.927) More Information \| Similar Motifs Found | motif file (matrix) |
| 5 | G T C A A G T C G T A C A G C T C A G T A G C T T C A G C T A G G C T A T A G C | 1e-16 | -3.831e+01 | 9.41% | 4.27% | 432.1bp (398.2bp) | HNF4A(var.2)/MA1494.1/Jaspar(0.755) More Information \| Similar Motifs Found | motif file (matrix) |
| 6 | T C G A A C G T A C T G G T C A T A G C C A G T G T A C C G T A G A C T T G A C | 1e-14 | -3.340e+01 | 27.68% | 19.21% | 368.3bp (431.8bp) | JUND/MA0491.2/Jaspar(0.972) More Information \| Similar Motifs Found | motif file (matrix) |
| 7 | A T G C T G A C A G C T G C T A T C A G G A T C G C T A T C G A T G A C T G A C | 1e-13 | -3.124e+01 | 18.68% | 11.83% | 379.4bp (414.5bp) | PB0055.1\_Rfx4\_1/Jaspar(0.884) More Information \| Similar Motifs Found | motif file (matrix) |
| 8 | G A T C A T C G G A C T G A C T T G C A G A C T A T G C C G A T G A T C A C T G | 1e-13 | -3.058e+01 | 20.33% | 13.27% | 361.4bp (408.1bp) | GATA3(Zf)/iTreg-Gata3-ChIP-Seq(GSE20898)/Homer(0.836) More Information \| Similar Motifs Found | motif file (matrix) |
| 9 \* | T C G A T A G C A G C T T A G C T A G C C T A G G C A T T C A G G A C T A C T G | 1e-10 | -2.364e+01 | 7.69% | 3.97% | 356.1bp (393.3bp) | YLR278C/MA0430.1/Jaspar(0.714) More Information \| Similar Motifs Found | motif file (matrix) |
| 10 \* | G T C A A T C G G A T C T C G A G C T A G C A T A T C G A T G C A G C T T A G C | 1e-9 | -2.130e+01 | 8.24% | 4.54% | 361.8bp (420.3bp) | hkb/dmmpmm(Pollard)/fly(0.648) More Information \| Similar Motifs Found | motif file (matrix) |
| 11 \* | C G A T A C T G G T C A A G T C A C T G A G C T A G T C C G T A G A C T A T G C | 1e-9 | -2.097e+01 | 9.62% | 5.61% | 398.7bp (427.6bp) | FOSB::JUN/MA1127.1/Jaspar(0.961) More Information \| Similar Motifs Found | motif file (matrix) |
| 12 \* | C A T G C G A T C A G T T C G A A T C G A C T G C G T A A T G C C A T G T G A C | 1e-7 | -1.779e+01 | 1.85% | 0.51% | 324.0bp (438.7bp) | ZmHOX2a(1)(HD-HOX)/Zea mays/AthaMap(0.759) More Information \| Similar Motifs Found | motif file (matrix) |
| 13 \* | G A C T C A T G A T G C A G T C A T G C T A C G A T G C G A T C C T G A C G A T | 1e-7 | -1.772e+01 | 2.68% | 0.96% | 401.1bp (405.9bp) | pho/MA1460.1/Jaspar(0.742) More Information \| Similar Motifs Found | motif file (matrix) |
| 14 \* | C T G A A G T C C G A T A G T C G T A C A C G T G A C T A T C G T C A G A G C T | 1e-7 | -1.716e+01 | 3.78% | 1.66% | 378.9bp (364.1bp) | RBM5(RRM)/Homo\_sapiens-RNCMPT00154-PBM/HughesRNA(0.732) More Information \| Similar Motifs Found | motif file (matrix) |
| 15 \* | A G T C G A T C A T C G A C T G A G T C C T G A C T G A C G A T T G A C A C G T | 1e-6 | -1.431e+01 | 3.02% | 1.31% | 404.5bp (432.9bp) | ERF6/MA1006.1/Jaspar(0.670) More Information \| Similar Motifs Found | motif file (matrix) |
| 16 \* | G A C T T C G A C T A G C G T A G T A C A G T C C T A G C T G A | 1e-5 | -1.195e+01 | 16.83% | 12.83% | 359.9bp (417.6bp) | ZAP1(WRKY(Zn))/Arabidopsis thaliana/AthaMap(0.794) More Information \| Similar Motifs Found | motif file (matrix) |
| 17 \* | C G T A A C G T A C G T C G T A C G T A A C G T A G T C A C G T A C T G C G T A | 1e-5 | -1.194e+01 | 1.03% | 0.25% | 255.3bp (455.0bp) | bcd-1/dmmpmm(Noyes)/fly(0.728) More Information \| Similar Motifs Found | motif file (matrix) |
| 18 \* | A C T G C G T A C G T A A G T C A C T G A C T G C G T A C G T A A G T C A C T G | 1e-4 | -9.780e+00 | 0.34% | 0.03% | 234.6bp (80.0bp) | ERT1/MA0420.1/Jaspar(0.758) More Information \| Similar Motifs Found | motif file (matrix) |
