## Supplemental Table 1 for "The Establishment of Cell-Type Specific Gene Regulation in the Sea Urchin Embryo": Hpf72_LeftPouch_homerResults.html

homer\_motifs\_wbackground/Hpf72\_LeftPouch/ - Homer de novo Motif Results


### Homer *de novo* Motif Results (homer\_motifs\_wbackground/Hpf72\_LeftPouch/)

Known Motif Enrichment Results  
Gene Ontology Enrichment Results  
If Homer is having trouble matching a motif to a known motif, try copy/pasting the matrix file into
STAMP  
More information on motif finding results: HOMER
| Description of Results
| Tips
  
Total target sequences = 373  
Total background sequences = 16561  
\* - possible false positive  

|  |  |  |  |  |  |  |  |  |
| --- | --- | --- | --- | --- | --- | --- | --- | --- |
| Rank | Motif | P-value | log P-pvalue | % of Targets | % of Background | STD(Bg STD) | Best Match/Details | Motif File |
| 1 \* | A C T G C T G A A C T G A C T G A C G T A C T G C G T A A C G T A C T G A C T G | 1e-6 | -1.506e+01 | 3.75% | 0.66% | 228.9bp (592.7bp) | HAP3(CCAATHAP3)/col-HAP3-DAP-Seq(GSE60143)/Homer(0.740) More Information \| Similar Motifs Found | motif file (matrix) |
| 2 \* | C G A T T C G A C G A T A C T G A C T G G T C A C G T A C A G T A C G T A T G C | 1e-6 | -1.422e+01 | 7.24% | 2.43% | 305.5bp (419.8bp) | TEAD(TEA)/Fibroblast-PU.1-ChIP-Seq(Unpublished)/Homer(0.743) More Information \| Similar Motifs Found | motif file (matrix) |
| 3 \* | A T C G C T G A G C T A T G C A C T A G A G C T G T C A A G T C A C T G T A G C | 1e-6 | -1.420e+01 | 16.35% | 8.50% | 322.4bp (439.9bp) | CG5213(RRM)/Drosophila\_melanogaster-RNCMPT00010-PBM/HughesRNA(0.778) More Information \| Similar Motifs Found | motif file (matrix) |
| 4 \* | G T A C G C A T A C G T A T G C A C G T G T C A G C A T G C A T G T C A T G C A | 1e-5 | -1.361e+01 | 20.64% | 11.95% | 336.7bp (449.5bp) | br-Z2/dmmpmm(Pollard)/fly(0.784) More Information \| Similar Motifs Found | motif file (matrix) |
| 5 \* | C G T A G T C A G C T A C T A G A C G T T C A G C G T A T C G A G T C A G A C T | 1e-5 | -1.311e+01 | 32.44% | 21.98% | 342.1bp (441.0bp) | tll/dmmpmm(Papatsenko)/fly(0.792) More Information \| Similar Motifs Found | motif file (matrix) |
| 6 \* | C G A T A G C T T G A C C T G A C G A T C A G T T A C G A G T C A C T G C G A T | 1e-4 | -1.077e+01 | 6.97% | 2.77% | 307.2bp (427.7bp) | Hoxa10(Homeobox)/ChickenMSG-Hoxa10.Flag-ChIP-Seq(GSE86088)/Homer(0.796) More Information \| Similar Motifs Found | motif file (matrix) |
| 7 \* | T G A C A C T G T G C A T G C A A T C G A T C G T G C A A T C G T G C A A T C G | 1e-4 | -1.017e+01 | 7.51% | 3.22% | 221.9bp (512.1bp) | RBM5(RRM)/Homo\_sapiens-RNCMPT00154-PBM/HughesRNA(0.907) More Information \| Similar Motifs Found | motif file (matrix) |
| 8 \* | A C G T G A T C G C T A C T A G T A G C C G A T A T C G T C G A T A C G T A G C | 1e-4 | -9.919e+00 | 14.48% | 8.30% | 357.2bp (430.4bp) | PL0001.1\_hlh-11/Jaspar(0.839) More Information \| Similar Motifs Found | motif file (matrix) |
| 9 \* | A G C T C G T A A C T G C G T A A G C T A G T C C A G T G T C A A T C G G C A T | 1e-4 | -9.620e+00 | 7.77% | 3.50% | 328.8bp (427.2bp) | GATA6(C2C2gata)/col200-GATA6-DAP-Seq(GSE60143)/Homer(0.919) More Information \| Similar Motifs Found | motif file (matrix) |
| 10 \* | C A G T T G C A A G T C A G T C C G A T G T A C C A G T G T C A A T G C A C G T | 1e-4 | -9.222e+00 | 22.52% | 15.11% | 338.8bp (464.2bp) | RUNX3/MA0684.2/Jaspar(0.703) More Information \| Similar Motifs Found | motif file (matrix) |
| 11 \* | C T G A A T C G A C G T A G C T C A T G G A T C A C T G C G T A A G C T C G A T | 1e-3 | -8.183e+00 | 4.29% | 1.54% | 366.4bp (526.7bp) | PB0037.1\_Isgf3g\_1/Jaspar(0.757) More Information \| Similar Motifs Found | motif file (matrix) |
| 12 \* | T A C G C G A T G A T C T G A C C T A G A C T G A C T G A T C G C A G T A C T G | 1e-3 | -7.525e+00 | 12.60% | 7.64% | 320.2bp (454.8bp) | AFT2/MA0270.1/Jaspar(0.699) More Information \| Similar Motifs Found | motif file (matrix) |
| 13 \* | A C T G A C T G A G T C A C G T A C G T A C G T A G T C A C G T A G T C A C T G | 1e-3 | -7.074e+00 | 0.80% | 0.05% | 123.7bp (328.1bp) | GCR1/Literature(Harbison)/Yeast(0.767) More Information \| Similar Motifs Found | motif file (matrix) |
| 14 \* | T G C A A G T C G T A C G T C A G A C T A G T C A T G C A C G T C G T A A T G C | 1e-2 | -6.653e+00 | 4.02% | 1.62% | 311.9bp (481.8bp) | G3BP2(RRM)/Homo\_sapiens-RNCMPT00021-PBM/HughesRNA(0.718) More Information \| Similar Motifs Found | motif file (matrix) |
| 15 \* | A C G T A C T G C G T A A G C T A C G T C G T A A C T G A C G T | 1e-2 | -6.373e+00 | 6.43% | 3.31% | 343.9bp (444.7bp) | PFF0320c(RRM)/Plasmodium\_falciparum-RNCMPT00235-PBM/HughesRNA(0.838) More Information \| Similar Motifs Found | motif file (matrix) |
| 16 \* | C G A T A C G T C G T A A G T C C T A G A C G T A C T G A C T G | 1e-2 | -6.158e+00 | 7.24% | 3.95% | 305.6bp (415.6bp) | ABF4/MA1659.1/Jaspar(0.789) More Information \| Similar Motifs Found | motif file (matrix) |
| 17 \* | A G T C A C T G A G T C A C T G C G T A A C T G C G T A C G T A A C T G C G T A | 1e-2 | -6.130e+00 | 0.54% | 0.02% | 60.4bp (410.3bp) | ZBTB33/MA0527.1/Jaspar(0.760) More Information \| Similar Motifs Found | motif file (matrix) |
| 18 \* | A T G C C A T G A C G T A T G C T A G C G C T A A G T C T G A C A T G C C T G A | 1e-2 | -5.933e+00 | 1.07% | 0.16% | 220.4bp (867.0bp) | Gli2(Zf)/GM2-Gli2-ChIP-Chip(GSE112702)/Homer(0.724) More Information \| Similar Motifs Found | motif file (matrix) |
| 19 \* | A C T G A C G T C G T A C T G A C T G A A G T C A C T G A G T C A C T G G T C A | 1e-1 | -4.042e+00 | 0.80% | 0.15% | 217.6bp (801.8bp) | STB1/STB1\_YPD/1-SWI4,1-SWI6(Harbison)/Yeast(0.756) More Information \| Similar Motifs Found | motif file (matrix) |
| 20 \* | C A T G C G A T A T C G A T C G A C G T C A G T A T C G G C A T C G A T C T A G | 1e-1 | -3.846e+00 | 12.60% | 9.32% | 367.3bp (501.9bp) | Aef1/dmmpmm(Bergman)/fly(0.764) More Information \| Similar Motifs Found | motif file (matrix) |
| 21 \* | C G T A C G T A C G T A C G T A C G T A C G T A | 1e0 | -4.844e-01 | 58.98% | 59.59% | 324.2bp (456.6bp) | TIAR-1(RRM)/Caenorhabditis\_elegans-RNCMPT00256-PBM/HughesRNA(0.988) More Information \| Similar Motifs Found | motif file (matrix) |
| 22 \* | G T C A A C T G C G T A C G T A A C G T A C T G | 1e0 | -4.694e-01 | 43.43% | 44.13% | 355.6bp (444.9bp) | TEC1(MacIsaac)/Yeast(0.956) More Information \| Similar Motifs Found | motif file (matrix) |
