## Supplemental Table 1 for "The Establishment of Cell-Type Specific Gene Regulation in the Sea Urchin Embryo": Hpf72_Midgut_homerResults.html

homer\_motifs\_wbackground/Hpf72\_Midgut/ - Homer de novo Motif Results


### Homer *de novo* Motif Results (homer\_motifs\_wbackground/Hpf72\_Midgut/)

Known Motif Enrichment Results  
Gene Ontology Enrichment Results  
If Homer is having trouble matching a motif to a known motif, try copy/pasting the matrix file into
STAMP  
More information on motif finding results: HOMER
| Description of Results
| Tips
  
Total target sequences = 1574  
Total background sequences = 14509  
\* - possible false positive  

|  |  |  |  |  |  |  |  |  |
| --- | --- | --- | --- | --- | --- | --- | --- | --- |
| Rank | Motif | P-value | log P-pvalue | % of Targets | % of Background | STD(Bg STD) | Best Match/Details | Motif File |
| 1 | G A C T A G C T C T A G C T G A G T C A A T G C A G C T G A C T A G C T C T A G | 1e-63 | -1.469e+02 | 36.59% | 18.44% | 386.6bp (452.6bp) | HNF4A/MA0114.4/Jaspar(0.920) More Information \| Similar Motifs Found | motif file (matrix) |
| 2 | G C A T T C A G C G A T C A T G G A T C G T C A G T C A C T G A A G T C G C T A | 1e-58 | -1.358e+02 | 68.42% | 48.11% | 363.5bp (430.4bp) | fkh/dmmpmm(Noyes)/fly(0.922) More Information \| Similar Motifs Found | motif file (matrix) |
| 3 | G A T C C T G A A T G C G C A T A C T G T C G A A C G T T C G A C T G A T A C G | 1e-52 | -1.199e+02 | 43.77% | 25.95% | 369.4bp (424.8bp) | Gata4(Zf)/Heart-Gata4-ChIP-Seq(GSE35151)/Homer(0.975) More Information \| Similar Motifs Found | motif file (matrix) |
| 4 | C T A G C G A T A C G T G T C A T G C A G C A T T A G C C G T A A C G T A G C T | 1e-24 | -5.554e+01 | 7.43% | 2.48% | 328.0bp (466.0bp) | HNF1b(Homeobox)/PDAC-HNF1B-ChIP-Seq(GSE64557)/Homer(0.916) More Information \| Similar Motifs Found | motif file (matrix) |
| 5 | A C T G A T C G A C T G T C A G C G T A G C T A C G A T G A T C A G T C A G T C | 1e-19 | -4.428e+01 | 5.65% | 1.83% | 358.2bp (492.2bp) | dif/Rel/dmmpmm(Bergman)/fly(0.974) More Information \| Similar Motifs Found | motif file (matrix) |
| 6 | C G A T T A C G A C G T A G T C C G T A C G T A A G C T G T C A G C A T G C A T | 1e-16 | -3.863e+01 | 14.04% | 7.75% | 367.4bp (423.8bp) | slp1/dmmpmm(Papatsenko)/fly(0.838) More Information \| Similar Motifs Found | motif file (matrix) |
| 7 | G T A C T G A C T G C A C G T A C A T G C T A G A C G T G A C T | 1e-16 | -3.797e+01 | 43.01% | 32.88% | 426.0bp (426.1bp) | HAP2/MA0313.1/Jaspar(0.689) More Information \| Similar Motifs Found | motif file (matrix) |
| 8 | C T G A A C G T A C G T C T G A T G A C C T A G A C G T A G T C C G T A A C G T | 1e-12 | -2.767e+01 | 9.21% | 4.92% | 409.7bp (419.2bp) | FOSL2::JUN(var.2)/MA1131.1/Jaspar(0.921) More Information \| Similar Motifs Found | motif file (matrix) |
| 9 \* | A C G T A T C G G C T A G T A C T A C G G T C A T G A C A C T G T A G C T C G A | 1e-11 | -2.550e+01 | 3.94% | 1.46% | 347.7bp (455.5bp) | cnc::maf-S/MA0530.1/Jaspar(0.747) More Information \| Similar Motifs Found | motif file (matrix) |
| 10 \* | A G C T C T G A A C T G A C T G G C A T A G T C A G T C G T C A G A T C C T A G | 1e-10 | -2.465e+01 | 6.48% | 3.16% | 427.4bp (382.4bp) | TCP16(TCP)/colamp-TCP16-DAP-Seq(GSE60143)/Homer(0.851) More Information \| Similar Motifs Found | motif file (matrix) |
| 11 \* | C G T A A T G C G T C A A C G T G T C A A C T G G T A C A G C T G T C A C T A G | 1e-9 | -2.261e+01 | 9.15% | 5.24% | 355.9bp (430.6bp) | SOX10/MA0442.2/Jaspar(0.719) More Information \| Similar Motifs Found | motif file (matrix) |
| 12 \* | A C T G G C A T A G T C A G C T G C T A G T C A G T C A A T C G | 1e-9 | -2.101e+01 | 41.74% | 34.37% | 413.0bp (431.1bp) | Smad3(MAD)/NPC-Smad3-ChIP-Seq(GSE36673)/Homer(0.659) More Information \| Similar Motifs Found | motif file (matrix) |
| 13 \* | A G T C C G T A T A C G G T C A A T C G A T G C A T C G G T C A | 1e-8 | -2.062e+01 | 28.02% | 21.59% | 389.4bp (412.6bp) | Klf4(Zf)/mES-Klf4-ChIP-Seq(GSE11431)/Homer(0.625) More Information \| Similar Motifs Found | motif file (matrix) |
| 14 \* | T G A C A C G T G T C A A T C G A G T C G A C T C G T A C T G A | 1e-8 | -1.992e+01 | 20.46% | 14.93% | 354.0bp (411.6bp) | PH0040.1\_Hmbox1/Jaspar(0.800) More Information \| Similar Motifs Found | motif file (matrix) |
| 15 \* | C G T A A C G T A G T C A C G T A G T C A C T G G T C A A C T G C G T A A C G T | 1e-5 | -1.349e+01 | 0.51% | 0.05% | 316.1bp (454.4bp) | ZBED1/MA0749.1/Jaspar(0.742) More Information \| Similar Motifs Found | motif file (matrix) |
| 16 \* | A G T C A C G T A C G T A C T G A G T C A C T G A C G T C G T A C G T A A G T C | 1e-5 | -1.197e+01 | 0.38% | 0.03% | 122.4bp (267.3bp) | CEBPB/MA0466.2/Jaspar(0.788) More Information \| Similar Motifs Found | motif file (matrix) |
| 17 \* | G C A T A C G T C G T A A C T G A C G T A C T G A T C G A T C G A G C T A C T G | 1e-5 | -1.174e+01 | 1.08% | 0.30% | 407.9bp (460.4bp) | CRZ1(MacIsaac)/Yeast(0.706) More Information \| Similar Motifs Found | motif file (matrix) |
| 18 \* | A C T G A C T G G T C A A C G T C G T A A C G T A C G T A G C T | 1e-4 | -1.070e+01 | 23.44% | 19.26% | 425.6bp (460.5bp) | AT3G10113(MYBrelated)/col-AT3G10113-DAP-Seq(GSE60143)/Homer(0.939) More Information \| Similar Motifs Found | motif file (matrix) |
| 19 \* | A C G T C G T A A C G T C G T A A G C T C G T A A C G T G T C A A G C T C G T A | 1e-3 | -8.454e+00 | 5.02% | 3.30% | 276.5bp (429.5bp) | SeqBias: TA-repeat(0.984) More Information \| Similar Motifs Found | motif file (matrix) |
| 20 \* | A C T G C G T A C G T A A G T C A C T G A C T G C G T A C G T A A G T C A T C G | 1e-3 | -8.174e+00 | 0.38% | 0.06% | 258.5bp (666.8bp) | ERT1/MA0420.1/Jaspar(0.756) More Information \| Similar Motifs Found | motif file (matrix) |
| 21 \* | A C G T A C T G A G T C A C G T A C G T A C G T A C T G A C G T A C G T A C G T | 1e-1 | -4.465e+00 | 1.21% | 0.67% | 348.5bp (363.1bp) | SOX10/MA0442.2/Jaspar(0.840) More Information \| Similar Motifs Found | motif file (matrix) |
