## Supplemental Table 1 for "The Establishment of Cell-Type Specific Gene Regulation in the Sea Urchin Embryo": Hpf72_MuscleCells_homerResults.html

homer\_motifs\_wbackground/Hpf72\_MuscleCells/ - Homer de novo Motif Results


### Homer *de novo* Motif Results (homer\_motifs\_wbackground/Hpf72\_MuscleCells/)

Known Motif Enrichment Results  
Gene Ontology Enrichment Results  
If Homer is having trouble matching a motif to a known motif, try copy/pasting the matrix file into
STAMP  
More information on motif finding results: HOMER
| Description of Results
| Tips
  
Total target sequences = 442  
Total background sequences = 15889  
\* - possible false positive  

|  |  |  |  |  |  |  |  |  |
| --- | --- | --- | --- | --- | --- | --- | --- | --- |
| Rank | Motif | P-value | log P-pvalue | % of Targets | % of Background | STD(Bg STD) | Best Match/Details | Motif File |
| 1 \* | A C T G A G T C A C T G A C G T A T C G C G T A C A T G T A G C C T A G T A C G | 1e-8 | -2.069e+01 | 4.30% | 0.72% | 260.2bp (574.3bp) | EGR1/MA0162.4/Jaspar(0.665) More Information \| Similar Motifs Found | motif file (matrix) |
| 2 \* | A C T G G T A C T A C G T C G A C G A T A G C T T A C G A G T C C T G A G T A C | 1e-8 | -2.030e+01 | 6.56% | 1.72% | 248.5bp (469.2bp) | Ddit3::Cebpa/MA0019.1/Jaspar(0.799) More Information \| Similar Motifs Found | motif file (matrix) |
| 3 \* | G T A C T A G C A G C T A T C G A G T C C T A G A T G C G C A T C G T A C G T A | 1e-7 | -1.746e+01 | 3.85% | 0.70% | 273.6bp (519.3bp) | STP3/MA0396.1/Jaspar(0.774) More Information \| Similar Motifs Found | motif file (matrix) |
| 4 \* | A G T C T G C A G C T A C G T A C G T A A G T C G T A C T A C G C G T A T G A C | 1e-7 | -1.642e+01 | 10.41% | 4.37% | 198.8bp (419.3bp) | CEJ1(AP2EREBP)/col-CEJ1-DAP-Seq(GSE60143)/Homer(0.754) More Information \| Similar Motifs Found | motif file (matrix) |
| 5 \* | A C T G C G T A G T A C T A C G A C T G A C G T G T C A A C T G T A C G C G T A | 1e-6 | -1.585e+01 | 2.49% | 0.30% | 205.7bp (420.7bp) | OSR1/MA1542.1/Jaspar(0.662) More Information \| Similar Motifs Found | motif file (matrix) |
| 6 \* | A T G C C A T G C G T A A G C T G T A C A C T G A T C G A G T C A C T G C G T A | 1e-6 | -1.535e+01 | 2.94% | 0.46% | 230.1bp (433.2bp) | wc-1/MA1437.1/Jaspar(0.738) More Information \| Similar Motifs Found | motif file (matrix) |
| 7 \* | A G T C C T A G C G A T A T C G A G C T A G T C G A T C G T A C T C G A C A T G | 1e-6 | -1.499e+01 | 5.66% | 1.72% | 300.6bp (459.0bp) | PL0004.1\_hlh-27/Jaspar(0.762) More Information \| Similar Motifs Found | motif file (matrix) |
| 8 \* | G C A T T A G C C T G A C G A T G T C A T G A C C A T G T G A C T G C A A T G C | 1e-6 | -1.414e+01 | 4.98% | 1.44% | 309.6bp (474.4bp) | PB0143.1\_Klf7\_2/Jaspar(0.676) More Information \| Similar Motifs Found | motif file (matrix) |
| 9 \* | C G T A C T G A G A C T A C G T A C G T C A G T G A C T T G C A C G T A G C A T | 1e-5 | -1.310e+01 | 22.85% | 14.53% | 323.4bp (406.2bp) | cad/dmmpmm(Down)/fly(0.802) More Information \| Similar Motifs Found | motif file (matrix) |
| 10 \* | T C A G T G C A T G A C A T C G T C G A C A G T A C T G T G C A A C T G C A T G | 1e-5 | -1.175e+01 | 14.71% | 8.39% | 286.9bp (459.9bp) | ZML2(C2C2gata)/col-ZML2-DAP-Seq(GSE60143)/Homer(0.773) More Information \| Similar Motifs Found | motif file (matrix) |
| 11 \* | A C T G C G T A A C T G A C G T G T A C C G T A C T G A A C G T A G T C C G T A | 1e-5 | -1.160e+01 | 3.39% | 0.86% | 230.6bp (413.1bp) | DUX(Homeobox)/C2C12-Dux-ChIP-Seq(GSE87279)/Homer(0.803) More Information \| Similar Motifs Found | motif file (matrix) |
| 12 \* | A G T C C G T A C G A T A C T G A C G T C T A G C A T G G T C A A G T C C T A G | 1e-4 | -1.117e+01 | 5.88% | 2.27% | 285.9bp (410.4bp) | MXI1/MA1108.2/Jaspar(0.776) More Information \| Similar Motifs Found | motif file (matrix) |
| 13 \* | T C A G T C A G A T C G C T A G A C G T G T C A T A C G A C G T G C A T C T A G | 1e-4 | -1.062e+01 | 4.98% | 1.80% | 331.5bp (417.0bp) | Rbm42(RRM)/Xenopus\_tropicalis-RNCMPT00282-PBM/HughesRNA(0.770) More Information \| Similar Motifs Found | motif file (matrix) |
| 14 \* | A G T C A G C T C T A G C G T A T G C A T A G C A G C T T A C G T C A G G C T A | 1e-4 | -1.051e+01 | 6.33% | 2.65% | 266.7bp (418.9bp) | RORA/MA0071.1/Jaspar(0.675) More Information \| Similar Motifs Found | motif file (matrix) |
| 15 \* | C T A G C G T A A C T G A T G C G T C A A C T G C G T A C G T A A C G T A C T G | 1e-4 | -9.986e+00 | 3.85% | 1.23% | 204.0bp (427.0bp) | TEC1(MacIsaac)/Yeast(0.748) More Information \| Similar Motifs Found | motif file (matrix) |
| 16 \* | C G A T C G A T T A G C T A C G A C G T A G C T C A T G C A G T T G C A G A T C | 1e-4 | -9.298e+00 | 11.09% | 6.27% | 252.7bp (473.3bp) | RBP1(RRM)/Drosophila\_melanogaster-RNCMPT00058-PBM/HughesRNA(0.773) More Information \| Similar Motifs Found | motif file (matrix) |
| 17 \* | A G C T T A G C G A C T C G A T A C T G A C T G G C A T T G A C C A G T T A G C | 1e-3 | -8.975e+00 | 15.84% | 10.13% | 327.0bp (458.6bp) | eor-1/MA0543.1/Jaspar(0.745) More Information \| Similar Motifs Found | motif file (matrix) |
| 18 \* | A T C G T C A G A G C T A G C T A G C T T A C G A G C T G A T C C T A G T G A C | 1e-3 | -8.058e+00 | 14.03% | 8.97% | 331.2bp (434.3bp) | KHDRBS3(KH)/Homo\_sapiens-RNCMPT00034-PBM/HughesRNA(0.695) More Information \| Similar Motifs Found | motif file (matrix) |
| 19 \* | G T A C G T A C C G A T G C T A A G T C G T A C C G A T A G T C | 1e-2 | -6.637e+00 | 16.97% | 11.99% | 375.3bp (463.8bp) | HRB98DE(RRM)/Drosophila\_melanogaster-RNCMPT00094-PBM/HughesRNA(0.794) More Information \| Similar Motifs Found | motif file (matrix) |
| 20 \* | C T A G T G A C A G T C A G T C C T G A G T C A T C A G A T G C | 1e-2 | -5.286e+00 | 12.22% | 8.52% | 283.8bp (461.0bp) | MBNL1(Znf)/Homo\_sapiens-RNCMPT00038-PBM/HughesRNA(0.775) More Information \| Similar Motifs Found | motif file (matrix) |
| 21 \* | T C G A G A C T G T C A G C A T C G A T G C T A C G T A A C G T C T G A C T G A | 1e-2 | -4.769e+00 | 15.38% | 11.52% | 303.5bp (433.5bp) | Arid3b/MA0601.1/Jaspar(0.855) More Information \| Similar Motifs Found | motif file (matrix) |
| 22 \* | C T A G C G A T C T A G C G A T C T A G C G T A A C T G C G T A A C T G C T G A | 1e-1 | -3.391e+00 | 5.43% | 3.62% | 298.6bp (412.2bp) | Rbm24(RRM)/Tetraodon\_nigroviridis-RNCMPT00285-PBM/HughesRNA(0.760) More Information \| Similar Motifs Found | motif file (matrix) |
| 23 \* | T G A C T C A G T A G C T C G A T G A C C T A G A T G C G C T A | 1e-1 | -3.101e+00 | 10.63% | 8.25% | 297.7bp (446.4bp) | RBM8A(RRM)/Homo\_sapiens-RNCMPT00056-PBM/HughesRNA(0.771) More Information \| Similar Motifs Found | motif file (matrix) |
| 24 \* | A G T C A C T G A C T G C G T A A C T G A C T G | 1e-1 | -2.659e+00 | 17.19% | 14.58% | 339.2bp (548.7bp) | LIN28A(CSD)/Homo\_sapiens-RNCMPT00162-PBM/HughesRNA(0.956) More Information \| Similar Motifs Found | motif file (matrix) |
| 25 \* | G A C T G A T C A C T G G A C T A T G C A C G T C A G T A G T C G A C T G A C T | 1e0 | -2.282e+00 | 17.42% | 15.13% | 318.2bp (474.6bp) | Unknown4/Arabidopsis-Promoters/Homer(0.802) More Information \| Similar Motifs Found | motif file (matrix) |
