## Supplemental Table 1 for "The Establishment of Cell-Type Specific Gene Regulation in the Sea Urchin Embryo": Hpf72_Neurons_homerResults.html

homer\_motifs\_wbackground/Hpf72\_Neurons/ - Homer de novo Motif Results


### Homer *de novo* Motif Results (homer\_motifs\_wbackground/Hpf72\_Neurons/)

Known Motif Enrichment Results  
Gene Ontology Enrichment Results  
If Homer is having trouble matching a motif to a known motif, try copy/pasting the matrix file into
STAMP  
More information on motif finding results: HOMER
| Description of Results
| Tips
  
Total target sequences = 742  
Total background sequences = 14649  
\* - possible false positive  

|  |  |  |  |  |  |  |  |  |
| --- | --- | --- | --- | --- | --- | --- | --- | --- |
| Rank | Motif | P-value | log P-pvalue | % of Targets | % of Background | STD(Bg STD) | Best Match/Details | Motif File |
| 1 | A C T G G C A T A C T G C G A T A C T G C G A T C T A G C G A T C T A G G C A T | 1e-28 | -6.462e+01 | 24.39% | 10.17% | 278.2bp (501.8bp) | SeqBias: CA-repeat(0.898) More Information \| Similar Motifs Found | motif file (matrix) |
| 2 | G C T A A G C T C A T G G T A C G C A T T C G A T C G A G A C T A C T G G T C A | 1e-22 | -5.270e+01 | 38.14% | 21.93% | 334.7bp (476.4bp) | Oct6(POU,Homeobox)/NPC-Pou3f1-ChIP-Seq(GSE35496)/Homer(0.911) More Information \| Similar Motifs Found | motif file (matrix) |
| 3 | C G A T T C A G G T C A C G T A A C G T G T A C C G T A C T G A A C G T G C T A | 1e-18 | -4.258e+01 | 29.11% | 16.04% | 384.0bp (470.4bp) | ONECUT1/MA0679.2/Jaspar(0.795) More Information \| Similar Motifs Found | motif file (matrix) |
| 4 | C G T A C T A G G A T C C T A G G C T A A T C G G A C T A T C G | 1e-16 | -3.710e+01 | 65.23% | 50.14% | 357.8bp (556.1bp) | FHY3/MA0557.1/Jaspar(0.783) More Information \| Similar Motifs Found | motif file (matrix) |
| 5 | T C G A G C A T C T A G G T A C C G T A A G C T T C A G G T A C G T A C A C T G | 1e-15 | -3.541e+01 | 61.86% | 47.07% | 328.2bp (496.4bp) | ABI3/MA0564.1/Jaspar(0.849) More Information \| Similar Motifs Found | motif file (matrix) |
| 6 | C T A G A C T G C G T A C A G T C T A G T C G A C G A T C T A G | 1e-14 | -3.406e+01 | 50.94% | 36.67% | 347.6bp (527.7bp) | AGL42/MA1201.1/Jaspar(0.870) More Information \| Similar Motifs Found | motif file (matrix) |
| 7 | C G T A T A G C C G T A A T C G C T G A A C G T C G T A T A C G T G C A C T A G | 1e-13 | -3.057e+01 | 21.56% | 11.85% | 368.3bp (580.4bp) | Trl/MA0205.2/Jaspar(0.754) More Information \| Similar Motifs Found | motif file (matrix) |
| 8 | G C A T C T G A G C T A A G C T A C T G T G C A T C A G G C T A A G C T A C T G | 1e-12 | -2.981e+01 | 28.44% | 17.47% | 343.4bp (455.0bp) | POU6F1(var.2)/MA1549.1/Jaspar(0.802) More Information \| Similar Motifs Found | motif file (matrix) |
| 9 \* | G A C T A T G C A C G T A C T G A G T C A T G C A G C T A C T G A T G C A T C G | 1e-10 | -2.506e+01 | 4.45% | 1.06% | 372.9bp (518.4bp) | SOK2(MacIsaac)/Yeast(0.786) More Information \| Similar Motifs Found | motif file (matrix) |
| 10 \* | G T C A C G A T G C A T T C G A G T C A G C T A A T G C C G T A C T A G A T G C | 1e-10 | -2.451e+01 | 6.60% | 2.20% | 335.1bp (371.6bp) | Arid3a/MA0151.1/Jaspar(0.744) More Information \| Similar Motifs Found | motif file (matrix) |
| 11 \* | C T A G C A T G A G T C A C G T T C G A A C T G A G T C C A T G | 1e-10 | -2.437e+01 | 48.25% | 36.39% | 359.4bp (535.3bp) | STP3/MA0396.1/Jaspar(0.834) More Information \| Similar Motifs Found | motif file (matrix) |
| 12 \* | C A T G A G T C G A C T A G C T A T C G A T G C A C G T G A T C T A C G G T A C | 1e-10 | -2.413e+01 | 7.41% | 2.68% | 356.0bp (418.2bp) | MBNL1(Znf)/Homo\_sapiens-RNCMPT00038-PBM/HughesRNA(0.764) More Information \| Similar Motifs Found | motif file (matrix) |
| 13 \* | T A C G A G T C C A T G G T A C C G T A G C A T C G A T T C G A A G T C C T A G | 1e-10 | -2.338e+01 | 21.29% | 12.76% | 330.2bp (524.2bp) | SKO1/MA0382.1/Jaspar(0.749) More Information \| Similar Motifs Found | motif file (matrix) |
| 14 \* | A G C T A G T C A T C G A G T C A C T G C T A G A G T C A C T G A G T C A G T C | 1e-9 | -2.297e+01 | 3.23% | 0.62% | 279.3bp (362.8bp) | STP1(MacIsaac)/Yeast(0.831) More Information \| Similar Motifs Found | motif file (matrix) |
| 15 \* | G C A T A T G C G C A T G T A C A C G T G T A C G A C T G C A T G A T C A G T C | 1e-9 | -2.235e+01 | 26.01% | 16.84% | 411.6bp (546.6bp) | Trl/MA0205.2/Jaspar(0.853) More Information \| Similar Motifs Found | motif file (matrix) |
| 16 \* | C G A T A T G C G A T C A T G C G C A T A T G C A G T C G C T A A G C T T G A C | 1e-8 | -2.064e+01 | 28.98% | 19.74% | 361.9bp (576.5bp) | B52(RRM)/Drosophila\_melanogaster-RNCMPT00134-PBM/HughesRNA(0.732) More Information \| Similar Motifs Found | motif file (matrix) |
| 17 \* | T A C G A C G T C T G A G C A T A C G T A G T C A G T C T G C A A C G T G C T A | 1e-8 | -1.960e+01 | 16.04% | 9.24% | 363.4bp (462.6bp) | HSF1/MA0319.1/Jaspar(0.749) More Information \| Similar Motifs Found | motif file (matrix) |
| 18 \* | T A C G T A G C T G A C G C T A A G C T T C G A G A T C A T C G | 1e-7 | -1.828e+01 | 51.62% | 41.37% | 370.4bp (510.3bp) | NEUROD2/MA0668.1/Jaspar(0.792) More Information \| Similar Motifs Found | motif file (matrix) |
| 19 \* | A G T C T G A C T G C A G C A T C G A T C G T A C T G A C T A G | 1e-6 | -1.606e+01 | 21.97% | 14.77% | 358.8bp (468.2bp) | Tup/dmmpmm(Noyes\_hd)/fly(0.866) More Information \| Similar Motifs Found | motif file (matrix) |
| 20 \* | C T G A A G T C A C G T C G A T A C G T A T G C C T G A A C T G A G T C C T A G | 1e-6 | -1.568e+01 | 2.02% | 0.36% | 220.0bp (327.0bp) | FZF1/MA0298.1/Jaspar(0.736) More Information \| Similar Motifs Found | motif file (matrix) |
| 21 \* | A G T C A C G T A C T G A C G T C G T A A G T C A C T G C T G A | 1e-6 | -1.474e+01 | 5.80% | 2.47% | 290.5bp (525.2bp) | SPL5(SBP)/colamp-SPL5-DAP-Seq(GSE60143)/Homer(0.827) More Information \| Similar Motifs Found | motif file (matrix) |
| 22 \* | C G T A A C T G A C G T G T A C A G T C A G T C A G T C C G T A C G T A A C G T | 1e-6 | -1.434e+01 | 1.48% | 0.21% | 314.8bp (308.7bp) | MZF1/MA0056.2/Jaspar(0.701) More Information \| Similar Motifs Found | motif file (matrix) |
