## Supplemental Table 1 for "The Establishment of Cell-Type Specific Gene Regulation in the Sea Urchin Embryo": Hpf72_PancreasL_homerResults.html

homer\_motifs\_wbackground/Hpf72\_PancreasL/ - Homer de novo Motif Results


### Homer *de novo* Motif Results (homer\_motifs\_wbackground/Hpf72\_PancreasL/)

Known Motif Enrichment Results  
Gene Ontology Enrichment Results  
If Homer is having trouble matching a motif to a known motif, try copy/pasting the matrix file into
STAMP  
More information on motif finding results: HOMER
| Description of Results
| Tips
  
Total target sequences = 343  
Total background sequences = 16404  
\* - possible false positive  

|  |  |  |  |  |  |  |  |  |
| --- | --- | --- | --- | --- | --- | --- | --- | --- |
| Rank | Motif | P-value | log P-pvalue | % of Targets | % of Background | STD(Bg STD) | Best Match/Details | Motif File |
| 1 \* | G A T C A C G T A G T C C G A T A C T G A C T G C G T A C A G T A T G C C G T A | 1e-9 | -2.277e+01 | 7.58% | 1.61% | 371.6bp (453.1bp) | ZBTB26/MA1579.1/Jaspar(0.752) More Information \| Similar Motifs Found | motif file (matrix) |
| 2 \* | A G C T G C A T C G A T G T A C A C T G C T A G A G T C A G T C C G A T A C G T | 1e-9 | -2.161e+01 | 7.29% | 1.58% | 356.7bp (422.6bp) | pho/MA1460.1/Jaspar(0.681) More Information \| Similar Motifs Found | motif file (matrix) |
| 3 \* | G A C T G A C T T C G A C T G A A T G C A G T C G C A T G C A T G T A C G T A C | 1e-9 | -2.129e+01 | 8.45% | 2.13% | 300.3bp (413.4bp) | tll/dmmpmm(Bigfoot)/fly(0.820) More Information \| Similar Motifs Found | motif file (matrix) |
| 4 \* | C G A T A C T G T A G C C G T A C G A T A G T C C G T A C A G T A T C G A C G T | 1e-8 | -1.952e+01 | 9.04% | 2.61% | 341.5bp (404.5bp) | Atf4(bZIP)/MEF-Atf4-ChIP-Seq(GSE35681)/Homer(0.745) More Information \| Similar Motifs Found | motif file (matrix) |
| 5 \* | G A T C C G T A A C G T A C T G A G T C A G C T T A G C A C G T A T C G A C T G | 1e-7 | -1.798e+01 | 4.96% | 0.88% | 333.7bp (427.1bp) | EIF-2ALPHA(S1)/Drosophila\_melanogaster-RNCMPT00273-PBM/HughesRNA(0.728) More Information \| Similar Motifs Found | motif file (matrix) |
| 6 \* | A C G T A C G T A C T G C G T A A G T C A C T G G T A C C A T G C T G A A G T C | 1e-7 | -1.779e+01 | 4.66% | 0.78% | 205.3bp (426.2bp) | exd/MA0222.1/Jaspar(0.697) More Information \| Similar Motifs Found | motif file (matrix) |
| 7 \* | A T G C T A C G A T C G A G T C T A G C C A T G T G A C A T G C T C A G A T G C | 1e-6 | -1.598e+01 | 9.62% | 3.40% | 254.3bp (529.0bp) | CRF4/MA0976.2/Jaspar(0.791) More Information \| Similar Motifs Found | motif file (matrix) |
| 8 \* | T G C A T G C A C A G T A G T C A G C T G T C A A T C G T C G A T A G C T G C A | 1e-6 | -1.534e+01 | 8.16% | 2.65% | 326.4bp (409.7bp) | Smad4/MA1153.1/Jaspar(0.760) More Information \| Similar Motifs Found | motif file (matrix) |
| 9 \* | A C G T C G T A A T C G C T G A A G C T A G T C A C G T C G T A A T C G C T G A | 1e-6 | -1.445e+01 | 19.83% | 10.74% | 369.0bp (430.5bp) | GATA6(C2C2gata)/col200-GATA6-DAP-Seq(GSE60143)/Homer(0.854) More Information \| Similar Motifs Found | motif file (matrix) |
| 10 \* | A T C G A G T C G C A T C T G A G A T C A C G T G T C A G T A C A G C T C G T A | 1e-5 | -1.223e+01 | 11.95% | 5.60% | 492.9bp (415.0bp) | An\_0287(RRM)/Aspergillus\_nidulans-RNCMPT00287-PBM/HughesRNA(0.728) More Information \| Similar Motifs Found | motif file (matrix) |
| 11 \* | A C G T G C T A A C T G A G C T A C T G T G C A C G T A C A T G C G A T A G C T | 1e-4 | -1.131e+01 | 26.24% | 17.04% | 330.0bp (430.9bp) | At5g04390(C2H2)/col200-At5g04390-DAP-Seq(GSE60143)/Homer(0.714) More Information \| Similar Motifs Found | motif file (matrix) |
| 12 \* | T A G C A G T C A G T C G T A C A G T C A T G C A T C G C T A G C T A G A T C G | 1e-4 | -1.114e+01 | 3.21% | 0.63% | 328.5bp (480.4bp) | PB0102.1\_Zic2\_1/Jaspar(0.910) More Information \| Similar Motifs Found | motif file (matrix) |
| 13 \* | C G T A A T G C T A G C C T A G A T C G C T G A A G C T A C G T | 1e-4 | -1.002e+01 | 22.16% | 14.18% | 314.4bp (421.3bp) | Dmbx1/MA0883.1/Jaspar(0.760) More Information \| Similar Motifs Found | motif file (matrix) |
| 14 \* | A C G T A G T C A T G C C T A G T G C A A G T C C G T A G T C A | 1e-4 | -9.700e+00 | 13.99% | 7.78% | 346.2bp (461.7bp) | ARF2(ARF)/col-ARF2-DAP-Seq(GSE60143)/Homer(0.842) More Information \| Similar Motifs Found | motif file (matrix) |
| 15 \* | A C T G A C G T A C G T A G T C A G C T A T C G A C T G A G T C A C G T A C G T | 1e-3 | -9.162e+00 | 4.66% | 1.54% | 250.2bp (449.7bp) | Ot\_0263(RRM)/Ostreococcus\_tauri-RNCMPT00263-PBM/HughesRNA(0.688) More Information \| Similar Motifs Found | motif file (matrix) |
| 16 \* | A G T C C G T A C G T A A G T C A C G T C G T A A C T G A G T C A G T C A G T C | 1e-2 | -6.277e+00 | 0.58% | 0.02% | 215.8bp (188.7bp) | bHLH130/MA1358.1/Jaspar(0.678) More Information \| Similar Motifs Found | motif file (matrix) |
| 17 \* | C G T A A C G T A C T G A C G T A C G T A G T C | 1e-2 | -5.100e+00 | 43.15% | 36.43% | 386.9bp (443.9bp) | lin-14/MA0261.1/Jaspar(0.851) More Information \| Similar Motifs Found | motif file (matrix) |
| 18 \* | A T C G A C G T A C T G C A G T T A C G A C T G A C T G A C T G A C T G A C T G | 1e-1 | -4.377e+00 | 6.71% | 4.02% | 294.0bp (456.3bp) | ZNF740/MA0753.2/Jaspar(0.864) More Information \| Similar Motifs Found | motif file (matrix) |
| 19 \* | A C G T C T A G A C G T C G T A A C T G A C T G A C G T G T C A C G T A A G T C | 1e-1 | -4.344e+00 | 1.75% | 0.56% | 164.3bp (363.0bp) | MYB41(MYB)/col-MYB41-DAP-Seq(GSE60143)/Homer(0.750) More Information \| Similar Motifs Found | motif file (matrix) |
| 20 \* | A G T C A G C T C G T A A C T G A C T G A C T G C G T A C T A G A C T G A C T G | 1e-1 | -4.097e+00 | 1.46% | 0.43% | 199.0bp (423.4bp) | HNRNPH2(RRM)/Homo\_sapiens-RNCMPT00160-PBM/HughesRNA(0.848) More Information \| Similar Motifs Found | motif file (matrix) |
| 21 \* | A C T G A G T C A G T C C G T A A C G T A G T C | 1e-1 | -3.982e+00 | 30.03% | 24.94% | 359.8bp (457.6bp) | YY2/MA0748.2/Jaspar(0.809) More Information \| Similar Motifs Found | motif file (matrix) |
| 22 \* | C G T A A G T C A G T C G C T A A G T C G T C A T C G A A T G C T A G C C G T A | 1e-1 | -3.302e+00 | 6.12% | 4.00% | 268.4bp (425.7bp) | RUNX2(Runt)/PCa-RUNX2-ChIP-Seq(GSE33889)/Homer(0.712) More Information \| Similar Motifs Found | motif file (matrix) |
| 23 \* | A C G T A G T C A G T C A C T G A C T G A C T G A G T C C G T A C G T A A G T C | 1e-1 | -3.196e+00 | 0.29% | 0.01% | 204.2bp (191.7bp) | THAP1/MA0597.1/Jaspar(0.736) More Information \| Similar Motifs Found | motif file (matrix) |
| 24 \* | C G T A A G T C A G T C C G T A A C G T C G T A A G T C C G T A A C T G C G T A | 1e-1 | -2.334e+00 | 0.58% | 0.15% | 184.7bp (347.1bp) | ENOX1(RRM)/Homo\_sapiens-RNCMPT00149-PBM/HughesRNA(0.672) More Information \| Similar Motifs Found | motif file (matrix) |
| 25 \* | A G T C A C G T C G A T A G T C A C G T A C G T A G T C A G C T | 1e0 | -1.679e+00 | 25.95% | 23.76% | 340.6bp (449.2bp) | TRA2(RRM)/Drosophila\_melanogaster-RNCMPT00078-PBM/HughesRNA(0.899) More Information \| Similar Motifs Found | motif file (matrix) |
| 26 \* | A C G T A C G T A C G T A G T C C G T A C G T A A C G T A C T G A C T G A C G T | 1e0 | -9.319e-01 | 0.58% | 0.40% | 171.8bp (362.4bp) | At2g41835(C2H2)/col-At2g41835-DAP-Seq(GSE60143)/Homer(0.701) More Information \| Similar Motifs Found | motif file (matrix) |
| 27 \* | A C G T A G T C C G T A A G T C A G T C C G T A A C T G A C G T A G T C C G T A | 1e0 | -8.138e-01 | 0.29% | 0.17% | 72.4bp (275.9bp) | AGL55/MA1202.1/Jaspar(0.773) More Information \| Similar Motifs Found | motif file (matrix) |
