## Supplemental Table 1 for "The Establishment of Cell-Type Specific Gene Regulation in the Sea Urchin Embryo": Hpf72_PigmentCells_homerResults.html

homer\_motifs\_wbackground/Hpf72\_PigmentCells/ - Homer de novo Motif Results


### Homer *de novo* Motif Results (homer\_motifs\_wbackground/Hpf72\_PigmentCells/)

Known Motif Enrichment Results  
Gene Ontology Enrichment Results  
If Homer is having trouble matching a motif to a known motif, try copy/pasting the matrix file into
STAMP  
More information on motif finding results: HOMER
| Description of Results
| Tips
  
Total target sequences = 2376  
Total background sequences = 13815  
\* - possible false positive  

|  |  |  |  |  |  |  |  |  |
| --- | --- | --- | --- | --- | --- | --- | --- | --- |
| Rank | Motif | P-value | log P-pvalue | % of Targets | % of Background | STD(Bg STD) | Best Match/Details | Motif File |
| 1 | G A C T T C G A G T A C T A G C G T C A C T A G A G T C G T C A G C A T T G A C | 1e-578 | -1.332e+03 | 46.72% | 7.53% | 343.2bp (494.3bp) | ACE2/ACE2\_YPD/2-SWI5(Harbison)/Yeast(0.881) More Information \| Similar Motifs Found | motif file (matrix) |
| 2 | T C G A A G C T C T G A G C A T A T G C G T C A G C T A G C T A C A G T G A C T | 1e-37 | -8.522e+01 | 57.15% | 44.05% | 407.5bp (507.6bp) | PB0105.1\_Arid3a\_2/Jaspar(0.773) More Information \| Similar Motifs Found | motif file (matrix) |
| 3 | C T A G T G A C A T G C G A T C C A T G A G T C T G C A G C A T C T G A T G C A | 1e-25 | -5.965e+01 | 2.65% | 0.48% | 344.1bp (309.1bp) | gcm2/MA0917.1/Jaspar(0.706) More Information \| Similar Motifs Found | motif file (matrix) |
| 4 | C A G T T C A G C G T A G C T A C T G A C G A T C G A T T C A G G C T A C G T A | 1e-16 | -3.880e+01 | 18.56% | 12.47% | 408.8bp (526.2bp) | IRF1(IRF)/PBMC-IRF1-ChIP-Seq(GSE43036)/Homer(0.743) More Information \| Similar Motifs Found | motif file (matrix) |
| 5 | G A T C T C G A A C G T G T C A G A C T A C T G C A G T G A C T C G A T G T A C | 1e-16 | -3.768e+01 | 19.49% | 13.34% | 416.8bp (476.9bp) | BHLHE23/MA0817.1/Jaspar(0.870) More Information \| Similar Motifs Found | motif file (matrix) |
| 6 | A C T G A G T C G C T A C G T A A C G T C G T A C G T A G T C A | 1e-14 | -3.254e+01 | 24.45% | 18.12% | 434.9bp (459.6bp) | CDX2/MA0465.2/Jaspar(0.973) More Information \| Similar Motifs Found | motif file (matrix) |
| 7 | G C T A G T A C C G T A A C G T C G A T G C T A G T C A C G A T | 1e-13 | -3.027e+01 | 23.06% | 17.10% | 418.0bp (512.5bp) | zen/dmmpmm(SeSiMCMC)/fly(0.886) More Information \| Similar Motifs Found | motif file (matrix) |
| 8 | C T G A C G T A C G T A C G T A G C T A C G T A C G A T A G T C G T C A A C G T | 1e-12 | -2.987e+01 | 8.84% | 5.17% | 415.6bp (522.3bp) | hb/dmmpmm(Noyes)/fly(0.807) More Information \| Similar Motifs Found | motif file (matrix) |
| 9 \* | C T A G A C G T A G T C A C T G A C G T C G T A C G T A T C G A | 1e-11 | -2.758e+01 | 15.66% | 10.90% | 408.3bp (418.2bp) | HOXC9/MA0485.2/Jaspar(0.943) More Information \| Similar Motifs Found | motif file (matrix) |
| 10 \* | G T A C G T A C T A C G T A C G T G A C C T G A G C T A C A T G T A G C G C T A | 1e-10 | -2.528e+01 | 3.28% | 1.39% | 330.3bp (402.0bp) | MBNL1(Znf)/Homo\_sapiens-RNCMPT00038-PBM/HughesRNA(0.756) More Information \| Similar Motifs Found | motif file (matrix) |
| 11 \* | A C G T A T C G C G A T G T C A A G T C A C G T A C G T A C T G A C G T A C T G | 1e-10 | -2.403e+01 | 3.03% | 1.27% | 447.2bp (402.7bp) | SPL9(SBP)/colamp-SPL9-DAP-Seq(GSE60143)/Homer(0.751) More Information \| Similar Motifs Found | motif file (matrix) |
| 12 \* | A C G T A C G T A C T G A C G T A G C T A C G T A C G T C G T A | 1e-10 | -2.309e+01 | 16.84% | 12.32% | 368.6bp (539.6bp) | Pp\_0229(RRM)/Physcomitrella\_patens-RNCMPT00229-PBM/HughesRNA(0.778) More Information \| Similar Motifs Found | motif file (matrix) |
| 13 \* | A G T C A C T G C G T A G T C A G T C A A G T C A G T C A C T G A G T C A G T C | 1e-9 | -2.226e+01 | 0.46% | 0.03% | 266.5bp (189.6bp) | PB0034.1\_Irf4\_1/Jaspar(0.738) More Information \| Similar Motifs Found | motif file (matrix) |
| 14 \* | A C G T A C G T A G T C A C T G C G T A A C G T C G T A A G T C A C G T A C T G | 1e-8 | -1.895e+01 | 1.05% | 0.26% | 309.5bp (466.5bp) | TUT1(RRM,Znf)/Homo\_sapiens-RNCMPT00075-PBM/HughesRNA(0.809) More Information \| Similar Motifs Found | motif file (matrix) |
| 15 \* | A C T G A C T G A C T G A C T G A C T G A C T G A C T G A C T G A C T G A C T G | 1e-8 | -1.847e+01 | 16.71% | 12.70% | 527.2bp (493.1bp) | SeqBias: polyC-repeat(1.000) More Information \| Similar Motifs Found | motif file (matrix) |
| 16 \* | C G T A A G T C A C G T A C G T A C T G A G T C C G T A C G T A A C G T A C G T | 1e-7 | -1.822e+01 | 0.88% | 0.19% | 405.3bp (432.6bp) | bHLH80/MA1357.1/Jaspar(0.714) More Information \| Similar Motifs Found | motif file (matrix) |
| 17 \* | A C T G A C G T A C G T A C T G A C T G A G C T C G A T A C T G A C T G A C G T | 1e-6 | -1.383e+01 | 2.02% | 0.93% | 487.6bp (338.5bp) | AT1G24250(Orphan)/col-AT1G24250-DAP-Seq(GSE60143)/Homer(0.799) More Information \| Similar Motifs Found | motif file (matrix) |
| 18 \* | G T C A A G T C A C T G C G T A A C T G C G T A C G T A A G T C A C T G A C G T | 1e-5 | -1.263e+01 | 0.55% | 0.12% | 192.1bp (461.8bp) | AT5G22990(C2H2)/col-AT5G22990-DAP-Seq(GSE60143)/Homer(0.687) More Information \| Similar Motifs Found | motif file (matrix) |
| 19 \* | A C G T A G T C A C T G A G T C A C G T A G T C A C G T A G T C A C G T A G T C | 1e-5 | -1.184e+01 | 17.55% | 14.33% | 450.1bp (365.9bp) | SeqBias: GA-repeat(0.857) More Information \| Similar Motifs Found | motif file (matrix) |
| 20 \* | C T G A C G T A A C T G A C T G A G T C C G T A A C G T C G T A | 1e-5 | -1.166e+01 | 5.98% | 4.10% | 390.8bp (462.6bp) | unc-86/MA0926.1/Jaspar(0.759) More Information \| Similar Motifs Found | motif file (matrix) |
| 21 \* | C T A G A G T C C T A G G C A T A C T G A C G T A T C G C A G T A C T G C G A T | 1e-4 | -1.107e+01 | 2.90% | 1.68% | 446.6bp (364.4bp) | Rbm38(RRM)/Danio\_rerio-RNCMPT00283-PBM/HughesRNA(0.797) More Information \| Similar Motifs Found | motif file (matrix) |
| 22 \* | C G A T C G T A C A T G C G A T G T C A T A C G A C G T G C T A T C A G G A C T | 1e-4 | -1.067e+01 | 8.54% | 6.39% | 384.7bp (487.9bp) | MSI(RRM)/Drosophila\_melanogaster-RNCMPT00040-PBM/HughesRNA(0.734) More Information \| Similar Motifs Found | motif file (matrix) |
| 23 \* | A C G T A C G T A G T C C G T A A C G T C G T A T C G A C G A T | 1e-4 | -9.639e+00 | 9.64% | 7.47% | 392.7bp (489.7bp) | Pdp1/MA1702.1/Jaspar(0.781) More Information \| Similar Motifs Found | motif file (matrix) |
| 24 \* | A C T G C G T A C G T A A G T C A C T G A G T C A C G T C G T A A C G T C G T A | 1e-2 | -6.251e+00 | 0.21% | 0.04% | 218.4bp (343.4bp) | Lm\_0254(RRM)/Leishmania\_major-RNCMPT00254-PBM/HughesRNA(0.719) More Information \| Similar Motifs Found | motif file (matrix) |
