## Supplemental Table 1 for "The Establishment of Cell-Type Specific Gene Regulation in the Sea Urchin Embryo": Hpf72_PMCs_homerResults.html

homer\_motifs\_wbackground/Hpf72\_PMCs/ - Homer de novo Motif Results


### Homer *de novo* Motif Results (homer\_motifs\_wbackground/Hpf72\_PMCs/)

Known Motif Enrichment Results  
Gene Ontology Enrichment Results  
If Homer is having trouble matching a motif to a known motif, try copy/pasting the matrix file into
STAMP  
More information on motif finding results: HOMER
| Description of Results
| Tips
  
Total target sequences = 905  
Total background sequences = 15789  
\* - possible false positive  

|  |  |  |  |  |  |  |  |  |
| --- | --- | --- | --- | --- | --- | --- | --- | --- |
| Rank | Motif | P-value | log P-pvalue | % of Targets | % of Background | STD(Bg STD) | Best Match/Details | Motif File |
| 1 | T A G C T C G A T A G C T G C A A C T G T A C G G C T A C G T A T C A G A G C T | 1e-42 | -9.856e+01 | 40.88% | 20.63% | 376.1bp (466.1bp) | Etv2(ETS)/ES-ER71-ChIP-Seq(GSE59402)/Homer(0.953) More Information \| Similar Motifs Found | motif file (matrix) |
| 2 | T C G A A C G T A T C G G C T A A T C G A C G T T A G C G T C A A G T C C T A G | 1e-21 | -4.978e+01 | 22.10% | 10.88% | 369.2bp (425.1bp) | Smad2::Smad3/MA1622.1/Jaspar(0.902) More Information \| Similar Motifs Found | motif file (matrix) |
| 3 \* | C T G A A T G C C G A T T G C A A T C G T A G C T A C G C A T G T G A C T C G A | 1e-10 | -2.319e+01 | 3.65% | 0.93% | 205.1bp (347.8bp) | Zfp57(Zf)/H1-ZFP57.HA-ChIP-Seq(GSE115387)/Homer(0.711) More Information \| Similar Motifs Found | motif file (matrix) |
| 4 \* | A G C T C G T A C G T A A C G T A C G T C T G A C T G A G T A C | 1e-9 | -2.270e+01 | 35.36% | 25.82% | 349.9bp (418.1bp) | en/dmmpmm(Noyes\_hd)/fly(0.970) More Information \| Similar Motifs Found | motif file (matrix) |
| 5 \* | G C T A T A G C G A C T A G C T C A G T G A T C G T C A T C A G A T G C T C A G | 1e-8 | -2.007e+01 | 4.75% | 1.67% | 222.1bp (407.7bp) | tll/dmmpmm(Pollard)/fly(0.645) More Information \| Similar Motifs Found | motif file (matrix) |
| 6 \* | T A G C T C A G C T G A T A G C G C T A C A T G A C G T A G C T A T C G G T A C | 1e-8 | -1.911e+01 | 22.54% | 15.27% | 349.4bp (451.7bp) | MYOD1/MA0499.2/Jaspar(0.826) More Information \| Similar Motifs Found | motif file (matrix) |
| 7 \* | G C A T T C A G G T A C A C T G A C G T C G T A C T G A A G C T | 1e-8 | -1.907e+01 | 19.56% | 12.75% | 382.4bp (424.5bp) | YAP3/MA0416.1/Jaspar(0.845) More Information \| Similar Motifs Found | motif file (matrix) |
| 8 \* | C G T A A C T G A C G T G A C T A G C T A T C G A T C G T G A C C G T A G T C A | 1e-8 | -1.883e+01 | 7.73% | 3.65% | 301.1bp (440.6bp) | NFIA/MA0670.1/Jaspar(0.763) More Information \| Similar Motifs Found | motif file (matrix) |
| 9 \* | A C G T A T C G C T G A A G C T A T C G A C G T G T A C C T G A A G C T C T A G | 1e-6 | -1.601e+01 | 4.86% | 1.99% | 267.2bp (469.7bp) | CREB1/MA0018.4/Jaspar(0.913) More Information \| Similar Motifs Found | motif file (matrix) |
| 10 \* | G A C T A C T G T G C A A T G C G C T A A T C G T A G C C G A T T A C G A C T G | 1e-5 | -1.331e+01 | 16.24% | 11.06% | 359.3bp (474.4bp) | bZIP52/MA1343.1/Jaspar(0.878) More Information \| Similar Motifs Found | motif file (matrix) |
| 11 \* | C G T A A C G T C G T A A C G T A G T C A G T C A G T C A G T C A G T C C G T A | 1e-4 | -1.035e+01 | 0.77% | 0.10% | 427.9bp (222.7bp) | Znf281/MA1630.1/Jaspar(0.725) More Information \| Similar Motifs Found | motif file (matrix) |
| 12 \* | T A G C T C G A A T C G C G T A A T C G C A T G T A C G C T A G C T A G C T G A | 1e-4 | -9.581e+00 | 8.84% | 5.65% | 317.8bp (470.2bp) | ARF10/MA1685.1/Jaspar(0.668) More Information \| Similar Motifs Found | motif file (matrix) |
| 13 \* | C G T A A C T G A T C G A C T G A G T C C G T A A T C G A C G T | 1e-3 | -8.818e+00 | 3.87% | 1.95% | 417.0bp (440.4bp) | TBP3(MYBrelated)/col-TBP3-DAP-Seq(GSE60143)/Homer(0.773) More Information \| Similar Motifs Found | motif file (matrix) |
| 14 \* | A G T C A G T C A G T C A C G T G C T A G T A C G T A C A G T C | 1e-3 | -8.697e+00 | 16.57% | 12.43% | 390.9bp (497.2bp) | HRB98DE(RRM)/Drosophila\_melanogaster-RNCMPT00096-PBM/HughesRNA(0.843) More Information \| Similar Motifs Found | motif file (matrix) |
| 15 \* | C G A T A C T G C G A T C G T A A G T C A G T C A C G T A C T G C G T A A C T G | 1e-3 | -8.556e+00 | 0.99% | 0.22% | 342.3bp (468.3bp) | vfl/MA1462.1/Jaspar(0.734) More Information \| Similar Motifs Found | motif file (matrix) |
| 16 \* | C G T A C G T A A C G T A C T G C G T A A C G T A C G T A C G T | 1e-3 | -7.116e+00 | 11.49% | 8.40% | 320.6bp (427.8bp) | ATHB-5/MA0110.3/Jaspar(0.857) More Information \| Similar Motifs Found | motif file (matrix) |
| 17 \* | T C A G G A C T T A G C C G T A A G T C T C G A A C G T A C T G | 1e-2 | -6.582e+00 | 27.51% | 23.19% | 332.0bp (431.5bp) | CBF1(MacIsaac)/Yeast(0.899) More Information \| Similar Motifs Found | motif file (matrix) |
| 18 \* | A C G T A C T G A C G T A G T C A G T C C G T A C G T A A C G T | 1e-2 | -6.232e+00 | 4.97% | 3.13% | 369.8bp (410.2bp) | HAP2/Literature(Harbison)/Yeast(0.791) More Information \| Similar Motifs Found | motif file (matrix) |
| 19 \* | G T A C A C G T C A G T G T C A G T C A A T G C C A T G G T A C | 1e-2 | -5.655e+00 | 6.08% | 4.14% | 398.7bp (401.9bp) | AT3G10030(Trihelix)/colamp-AT3G10030-DAP-Seq(GSE60143)/Homer(0.782) More Information \| Similar Motifs Found | motif file (matrix) |
| 20 \* | A C T G C G A T G T A C T A C G A T G C A T G C A T C G C T G A | 1e-1 | -4.555e+00 | 3.43% | 2.18% | 352.8bp (449.4bp) | RDS2/MA0362.1/Jaspar(0.805) More Information \| Similar Motifs Found | motif file (matrix) |
| 21 \* | A G T C A G T C A C T G A C T G C G T A A G T C C G T A C G T A A G T C A G T C | 1e-1 | -3.791e+00 | 0.22% | 0.03% | 87.6bp (206.9bp) | RGT1/Literature(Harbison)/Yeast(0.707) More Information \| Similar Motifs Found | motif file (matrix) |
