## Supplemental Table 1 for "The Establishment of Cell-Type Specific Gene Regulation in the Sea Urchin Embryo": Hpf72_Sphincters_homerResults.html

homer\_motifs\_wbackground/Hpf72\_Sphincters/ - Homer de novo Motif Results


### Homer *de novo* Motif Results (homer\_motifs\_wbackground/Hpf72\_Sphincters/)

Known Motif Enrichment Results  
Gene Ontology Enrichment Results  
If Homer is having trouble matching a motif to a known motif, try copy/pasting the matrix file into
STAMP  
More information on motif finding results: HOMER
| Description of Results
| Tips
  
Total target sequences = 405  
Total background sequences = 16782  
\* - possible false positive  

|  |  |  |  |  |  |  |  |  |
| --- | --- | --- | --- | --- | --- | --- | --- | --- |
| Rank | Motif | P-value | log P-pvalue | % of Targets | % of Background | STD(Bg STD) | Best Match/Details | Motif File |
| 1 \* | A C G T A T C G A T C G C G T A A G T C C G T A T G A C A C T G C G T A C G T A | 1e-8 | -2.056e+01 | 5.68% | 1.19% | 534.0bp (389.7bp) | FMR1(KH)/Homo\_sapiens-RNCMPT00016-PBM/HughesRNA(0.710) More Information \| Similar Motifs Found | motif file (matrix) |
| 2 \* | C G T A G T A C G C T A A G C T A T G C G C T A T C G A G A T C C G T A C T G A | 1e-8 | -1.906e+01 | 11.11% | 4.22% | 354.2bp (422.2bp) | RBP1-LIKE(RRM)/Drosophila\_melanogaster-RNCMPT00127-PBM/HughesRNA(0.730) More Information \| Similar Motifs Found | motif file (matrix) |
| 3 \* | C G T A A T C G C G T A C T A G T A C G A T C G A T G C C G T A A C T G A G T C | 1e-7 | -1.808e+01 | 4.20% | 0.73% | 229.1bp (359.2bp) | Erra(NR)/HepG2-Erra-ChIP-Seq(GSE31477)/Homer(0.700) More Information \| Similar Motifs Found | motif file (matrix) |
| 4 \* | A T G C A T C G G T C A A C T G A G C T A C G T C G T A A G T C A C T G A C T G | 1e-7 | -1.798e+01 | 2.96% | 0.33% | 238.3bp (316.5bp) | IRF6/MA1509.1/Jaspar(0.763) More Information \| Similar Motifs Found | motif file (matrix) |
| 5 \* | G A T C C A T G C T A G C G A T C G A T A C G T G T A C G A T C C G A T C G A T | 1e-7 | -1.706e+01 | 13.83% | 6.31% | 304.1bp (408.0bp) | ETS:E-box(ETS,bHLH)/HPC7-Scl-ChIP-Seq(GSE22178)/Homer(0.753) More Information \| Similar Motifs Found | motif file (matrix) |
| 6 \* | A G C T A C G T C G A T T A C G G A C T T A G C C A G T G C A T T G A C A T G C | 1e-6 | -1.598e+01 | 10.86% | 4.53% | 427.6bp (441.5bp) | GCR1/MA0304.1/Jaspar(0.725) More Information \| Similar Motifs Found | motif file (matrix) |
| 7 \* | C T G A C G T A T C G A G C A T A C G T A C G T A C G T A C G T A T C G C G T A | 1e-6 | -1.558e+01 | 10.12% | 4.12% | 507.0bp (417.6bp) | Ng\_0261(RRM)/Naegleria\_gruberi-RNCMPT00261-PBM/HughesRNA(0.784) More Information \| Similar Motifs Found | motif file (matrix) |
| 8 \* | A G C T T C A G A G T C C G T A C G T A G A C T A G C T G T C A G C A T C A T G | 1e-6 | -1.473e+01 | 18.52% | 10.26% | 338.9bp (409.1bp) | YHP1/Literature(Harbison)/Yeast(0.824) More Information \| Similar Motifs Found | motif file (matrix) |
| 9 \* | A G T C C G T A C G T A A C G T A T G C A T G C A C T G A C G T G T C A C G T A | 1e-5 | -1.289e+01 | 3.46% | 0.73% | 223.8bp (524.0bp) | Eip74EF/dmmpmm(Bergman)/fly(0.679) More Information \| Similar Motifs Found | motif file (matrix) |
| 10 \* | A G T C C G T A T C A G A C T G C G A T A C T G A C G T C T A G C G T A C T A G | 1e-5 | -1.206e+01 | 4.44% | 1.26% | 237.2bp (427.7bp) | TBX1/MA0805.1/Jaspar(0.874) More Information \| Similar Motifs Found | motif file (matrix) |
| 11 \* | C A G T C T G A T C A G A T C G T A G C G T A C G C A T G T A C G T C A A G C T | 1e-5 | -1.194e+01 | 6.17% | 2.23% | 361.8bp (422.9bp) | ZFX(Zf)/mES-Zfx-ChIP-Seq(GSE11431)/Homer(0.707) More Information \| Similar Motifs Found | motif file (matrix) |
| 12 \* | C G T A A C T G A G C T A G T C A C T G A G T C A C T G A G T C A C G T A C T G | 1e-5 | -1.192e+01 | 2.72% | 0.49% | 231.6bp (389.1bp) | ZC3H10(Znf)/Homo\_sapiens-RNCMPT00085-PBM/HughesRNA(0.701) More Information \| Similar Motifs Found | motif file (matrix) |
| 13 \* | A C G T A C G T A C G T C G T A A G T C A G T C A G T C A C G T | 1e-4 | -1.063e+01 | 7.41% | 3.20% | 261.8bp (420.9bp) | REB1/MA0363.1/Jaspar(0.822) More Information \| Similar Motifs Found | motif file (matrix) |
| 14 \* | A G C T G T A C C A T G A G C T T G C A G A T C C A T G A C G T G T C A T A G C | 1e-4 | -1.045e+01 | 14.81% | 8.61% | 368.6bp (400.5bp) | Gmeb1/MA0615.1/Jaspar(0.755) More Information \| Similar Motifs Found | motif file (matrix) |
| 15 \* | T A C G T G C A G A T C C T G A G C T A G T C A A G C T A G C T A T C G T A C G | 1e-4 | -9.943e+00 | 9.63% | 4.86% | 327.4bp (423.6bp) | HAP2/Literature(Harbison)/Yeast(0.734) More Information \| Similar Motifs Found | motif file (matrix) |
| 16 \* | A C T G A G T C C G T A A C G T A C G T A G T C C G T A A G T C A G T C A C T G | 1e-3 | -8.090e+00 | 0.99% | 0.08% | 73.2bp (319.7bp) | Tb\_0251(RRM)/Trypanosoma\_brucei-RNCMPT00251-PBM/HughesRNA(0.704) More Information \| Similar Motifs Found | motif file (matrix) |
| 17 \* | A C G T C G T A A G T C C G T A A T C G A G C T A C G T A G T C A G T C A C G T | 1e-2 | -6.609e+00 | 2.72% | 0.90% | 157.9bp (423.8bp) | Eip74EF/dmmpmm(SeSiMCMC)/fly(0.735) More Information \| Similar Motifs Found | motif file (matrix) |
| 18 \* | A G T C C T G A G T A C A T G C C T G A G A T C A G T C C T G A G A T C G T A C | 1e-1 | -3.869e+00 | 19.01% | 15.17% | 351.7bp (439.8bp) | Run/dmmpmm(Papatsenko)/fly(0.790) More Information \| Similar Motifs Found | motif file (matrix) |
| 19 \* | A C T G A G T C A C T G A C T G C G T A A C T G A G T C C G T A | 1e0 | -2.160e+00 | 3.95% | 2.83% | 513.4bp (434.5bp) | POL013.1\_MED-1/Jaspar(0.839) More Information \| Similar Motifs Found | motif file (matrix) |
| 20 \* | A G T C A C G T A C T G A G T C A C G T C G T A | 1e0 | -9.830e-01 | 26.91% | 26.11% | 390.0bp (407.7bp) | UME6/UME6\_YPD/51-UME6(Harbison)/Yeast(0.722) More Information \| Similar Motifs Found | motif file (matrix) |
| 21 \* | A G T C A G T C A G T C A C G T A G T C C G T A | 1e0 | -3.966e-01 | 26.67% | 27.55% | 370.4bp (429.6bp) | z/dmmpmm(Bigfoot)/fly(0.795) More Information \| Similar Motifs Found | motif file (matrix) |
